# Comprehensive comparative genomics analysis for the emerging human pathogen *Streptococcus dysgalactiae* subsp. *equisimilis* (SDSE): A case study and Pan-subspecies genomic analysis

**DOI:** 10.1101/2021.09.13.460118

**Authors:** Amr T. M. Saeb, Hamsa T. Tayeb

**Author notes:** Corresponding Authors: Amr T. M. Saeb, Ph.D. College of Medicine, King Saud University, KSA. Zifo RnD Solutions, New Market, United Kingdom (Current address). Hamsa T. Tayeb, Ph.D. Genetics Department, King Faisal Specialist Hospital and Research Center, Riyadh, Saudi Arabia.

## Abstract

**Background:** *Streptococcus dysgalactiae* subsp. *equisimilis* (SDSE) is the causal agent of various diseases that include wound infection, erysipelas, cellulitis, life-threatening necrotizing fasciitis, and streptococcal toxic shock syndrome. It is capable of infecting both humans and animals. In this investigation, we present a comprehensive genomic analysis for the SDSE strain SCDR1 that belongs to Lancefield group G, emm type (stG6) and (MLST) sequence type (ST44) that is the first time to be documented in Saudi Arabia and the middle east. Besides, we present the most comprehensive comparative genomics analysis for the emerging human pathogen SDSE.

**Methodology:** We utilized next-generation sequencing techniques (NGS), bioinformatics, phylogenetic analysis, and comparative pathogenomics to characterize SCDR1 and all publicly available SDES genomes.

**Results:** We found that SCDR1 consisted of a circular genome of 2179136 bp. Comparative analyses among bacterial genomes indicated that SCDR1 was most closely related to AC-2713 and GGS_124. Genome annotation of SCDR-1 strain showed that it contains many genes with homology to known virulence factors, including genes involved in cellular invasion, Antiphagocytosis, immune evasion, invasion of skin and soft tissue, host mortality and tissue damage, toxins, pore-forming proteins, cytotoxins, beta-hemolysis agents. Two CRISPR arrays were identified in SCDR1 that are consist of 35 CRISPR repeats and 33 CRISPR spacers. Two CAS systems were observed in the SCDR-1 genome, namely, CAS-TypeIIA and CAS-TypeIC. SDSE core Resistome is consisting of 22 genes, including folA, ***gyrA***, ***gyrB,*** and ***FabK***. SDSE core Virulome consisting of 38 genes including, ***fba***, ***fbp54***, ***gidA,*** and ***lsp***.

**Conclusion:** Our study confirmed that the SDSE strains possess different characteristics in producing virulence factors for pathogenicity to humans and based on its genome sequence and close relationship with GAS. Our study shed light on the proposed pathogenic mechanisms of SDSE and may form the basis of molecular epidemiological research on these highly virulent bacteria.

## Introduction

*Streptococcus dysgalactiae* subsp. *equisimilis* (SDSE) is an emerging human pathogen that causes severe invasive streptococcal diseases (Matsue et al., 2020). It belongs to β-hemolytic streptococci and possesses Lancefield group C or G antigens and rarely, A antigen (Takahashi et al., 2011). SDSE was suggested as a new taxon among human streptococcal isolates in 1996. It is a common colonizer of the pharynx, skin, gastrointestinal tract, and female genital tract (Yung et al., 2019). However, SDSE is the causal agent of various diseases that include wound infection, erysipelas, cellulitis, streptococcal toxic shock syndrome (STSS), and life-threatening necrotizing fasciitis (Rantala, 2014). Besides, SDSE is associated with pharyngitis, acute post-streptococcal glomerulonephritis, and acute rheumatic fever (Reid et al., 1985; Haidan et al., 2000). These clinical characteristics of diseases produced by SDSE infection are closely similar to *Streptococcus pyogenes* (GAS) infections (Brandt and Spellerberg, 2009). Several studies suggested a unique link between group G streptococcal bacteremia and invasive infection with underlying conditions, such as alcoholism, diabetes mellitus, or cardiovascular (Armstrong et al., 1970). In addition, numerous studies reported the invasive pathogenicity of SDSE, especially with older adults with underlying diseases such as diabetes mellitus. Therefore, it was found that the mean age of adult patients with invasive SDSE infection is older than those infected with GAS (Cohen-Poradosu et al., 2004; Wajima et al., 2016). Furthermore, animal studies showed that an SDSE strain causes severe pathogenicity to diabetic mice compared with GAS strains (Ogura et al., 2018). SDSE causes skin and soft-tissue infections, including pyoderma, cellulitis, wound infections, abscesses, erysipelas, and necrotizing fasciitis (Broyles et al., 2009). Severe infections often affect injection drug users, elderly patients with underlying immunosuppressive comorbidities or conditions predisposing them to skin breakdown, and burn patients in whom the infections may lead to graft failure (Rider and McGregor, 1994). For invasive SDSE infections, sites of colonization or primary focal infections are most likely ports of entry (Auckenthaler et al., 1983). Invasive infections comprise arthritis, osteomyelitis, pleuropulmonary infections, peritonitis, intra-abdominal and epidural abscesses, meningitis, endocarditis, puerperal septicemia, neonatal infections, necrotizing fascitis, myositis, and streptococcal toxic-like syndrome (Barnham and Weightman, 2004). SDSE has Lancefield group C or G antigens, exhibits strong β-hemolysis, and exerts streptokinase activity upon human plasminogen and proteolytic activity upon human fibrin. Similarly, to group A streptococci, SDSE possesses virulence factors including M protein, streptolysin O, streptolysin S, streptokinase, hyaluronidase, C5a peptidase, and others (Takahashi et al., 2011). In contrast, SDSE lacks several key virulence factors present in GAS, such as superantigens other than *speG*, the cysteine protease *speB*, the hyaluronic synthesis operon *hasA* and *hasB* and an inhibitor of complement activation *sic* (Wajima et al., 2016; Watanabe et al., 2016). It was found that 59% of clinical isolates of SDSE clinical isolates were able to form biofilm in vitro; however, all of the tested isolates were able to form biofilm in vivo (Genteluci et al., 2015). It was found that in vivo passage in mice enhanced biofilm-forming ability in some SDSE isolates. Epidemiological surveillance has been crucial to detect changes in the geographical and temporal variation of GAS associated disease pattern; for this purpose, the M protein gene (*emm)* gene typing is the most widely used genotyping method than 200 *emm* types recognized (Gherardi et al., 2018). The *emm1* GAS, the strain with enhanced virulence, is the dominant type in Europe, North America, and Japan (Wajima et al., 2008; Gherardi et al., 2018). The enhanced virulence and invasion fitness of *emm1* GAS strain is justified by phage mobilization-based diversification and the capability to recognize and adapt to the diverse host environments (Aziz and Kotb, 2008). Consequently, it was reported that *emm1* type is associated with the distribution of phage-associated superantigen genes (*speA*, *speC* and *ssa*) and the number of necrotizing fasciitis cases (Plainvert et al., 2012). On the contrary, prevalent *emm* types of SDSE differ amongst countries. For instance, in Japan, the most prevalent is stG6792 followed by stG485 and stG245 (Wajima et al., 2016; Ikebe et al., 2018). Whereas in Taiwan, the stG10, stG245, stG840, stG6.1, and stG652 strains predominated (Ma et al., 2017). In Jerusalem and Austria stG485 and stG6.1 were predominant, respectively (Cohen-Poradosu et al., 2004; Leitner et al., 2015). In addition, the Multi-Locus Sequence Typing (MLST) showed that 178 SDSE isolates, representing 37 emm types, segregate into 80 distinct sequence types (STs) and that the same ST can harbor different emm types (McMillan et al., 2010). In this investigation we present a comprehensive genomic analysis for the SDSE strain SCDR1 that belong to Lancefield group G, emm type (stG6) and Multilocus sequence typing (MLST) sequence type (ST44) that is first time to be documented in Saudi Arabia and middle east. In addition, we present the most comprehensive comparative genomics analysis for the emerging human pathogen SDSE.

## Methodology

### Bacterial isolate

Streptococcus dysgalactiae subsp. equisimilis strain SCDR-SD1 was isolated from a diabetic ulcer patient in the diabetic foot unit at the University Diabetes Center, King Saud University. SDSE-SCDR1 was collected from a 60 years old’s neuroischaemic diabetic foot ulcer beneath the left fifth metatarsal head with a wound surface area of one cm^2^, and the depth of the wound was 0.5 cm. The wound was debrided before swabbing, and an appropriate wound swab was collected from the patient and was sent for additional microbiological culture and investigations. SDSE was cultured in 5% sheep blood agar or brain–heart infusion medium at 37 °C under 5% CO2 as described Miyoshi-Akiyama et al. (Miyoshi-Akiyama et al., 2003). The species identification was conducted by partially sequencing the 16S rRNA gene described previously (Wang et al., 2016). In vitro susceptibility of isolate SDSE-SCDR1 to antibiotics was determined by disk diffusion following the recommendations of CLSI and as described before (Matuschek et al., 2014; Wang et al., 2016).

### DNA isolation, purification, sequencing, bioinformatics, and phylogenetic analysis

DNA isolation, purification, genome sequencing, bioinformatics, and phylogenetic analysis were performed as described before (Saeb et al., 2017b; Saeb, 2018). The closest reference and representative genomes were identified using Mash/MinHash (Ondov et al., 2016). PATRIC global protein families (PGFams) (Davis et al., 2016) were selected from these genomes to determine this genome’s phylogenetic placement. The protein sequences from these families were aligned with MUSCLE (Edgar, 2004), and the nucleotides for each of those sequences were mapped to the protein alignment. The common set of amino acid and nucleotide alignments were concatenated into a data matrix, and RaxML (Stamatakis, 2014) was used to analyze this matrix, with fast bootstrapping (Stamatakis et al., 2008) was used to generate the support values in the tree.

Furthermore, we performed whole-genome phylogeny based proteomic comparison among *Streptococcus dysgalactiae* subsp. *equisimilis* SCDR-SD1 isolate and other related Proteus mirabilis strains using Proteome Comparison service, a protein sequence-based available at (https://www.patricbrc.org/app/SeqComparison) as described before (Saeb et al., 2017b, 2017a).

### Gene annotation and Pathogenomics analysis

*Streptococcus dysgalactiae* subsp. *equisimilis* SCDR- SD1 genome contigs were annotated using the Prokaryotic Genomes Automatic Annotation Pipeline (PGAAP) available at NCBI (http://www.ncbi.nlm.nih.gov/). Besides, contigs were further annotated using the bacterial bioinformatics database and analysis resource (PATRIC) gene annotation service (https://www.patricbrc.org/app/Annotation). **PubMLST,** the public databases for molecular typing and microbial genome diversity, was used to confirm the MLST analysis available at (https://pubmlst.org/). The PathogenFinder, pathogenicity prediction program available at (https://cge.cbs.dtu.dk/services/PathogenFinder/) was used to examine the nature of *Streptococcus dysgalactiae* subsp. *equisimilis* SCDR-SD1 as a human pathogen. Virulence gene sequences and functions, matching to different main bacterial virulence factors of SDSE were claimed from GenBank and confirmed using virulence factors of the pathogenic bacteria database available at (http://www.mgc.ac.cn/VFs/), the Victors virulence factors search service (http://www.phidias.us/victors/), besides, the PATRIC_VF tool accessible at https://www.patricbrc.org/ as described before (Saeb et al., 2017b). **RASTA-Bacteria,** the Rapid Automated Scan for Toxins and Antitoxins in Bacteria (Sevin and Barloy-Hubler, 2007), and **TASer, a** web tool that locates Toxins and Antitoxins in a genome using a high-performance Computing Cluster (Akarsu et al., 2019), were used for the Toxins and Antitoxins confirmatory analysis. **DrugBank,** available at (https://go.drugbank.com/), was used in the drug target analysis.

### Resistome analysis

*Streptococcus dysgalactiae* subsp. *equisimilis* SCDR-SD1 genome contigs were investigated manually for the presence of antibiotic resistance loci using the PGAAP and PATRIC gene annotation services. Antibiotic resistance loci were further investigated using specialized search tools and services, namely: Gene Search, Antibiotic Resistance, and Genome Feature Finder (https://www.patricbrc.org), CARD (The Comprehensive Antibiotic Resistance Database) accessible at (https://card.mcmaster.ca/), ARDB (Antibiotic Resistance Genes Database) accessible at (https://ardb.cbcb.umd.edu/), and ResFinder 2.1 available at (https://cge.cbs.dtu.dk//services/ResFinder/) as described before (Saeb et al., 2017b; Saeb, 2018).

### Phage, Insertion sequences, and CRISPR-CAS analysis

**PHASTER** (PHAge Search Tool Enhanced Release) available at (https://phaster.ca/), **PHAST** (PHAge Search Tool) available at (http://phast.wishartlab.com/index.html), **MARVEL**, a Tool for

Prediction of Bacteriophage Sequences in Metagenomic Bins (Amgarten et al., 2018) along with PARTIC’s Genome Feature functions were used for prophage search and annotation. Additionally, **ISMapper** was used in identifying transposase insertion sites in the SDSE-SCDR1 genome (Hawkey et al., 2015). **CRISPRCasFinder,** the program enables the easy detection of CRISPRs and ***cas*** genes, available at (https://crisprcas.i2bc.paris-saclay.fr/CrisprCasFinder/Index) along with PARTIC’s Genome Feature functions were used for CRISPRs-cas systems search and annotation.

### Comparative genomics and bioinformatics analysis

All publicly available representative SDSE genome sequences were collected from GenBank and validated using PARTIC deposited genomes. **Supplementary Table 1** represents the taxonomic/genomic information of the 55 publicly available SDSE strains/isolates used in this study. Comparative investigations were performed for all genomes previously described (Saeb et al., 2017a, 2017b): this includes phylogenomic, virulome, resistome, drug targets, prophage, CRISPER-CAS analysis. Additional web recourses used in the analysis are the **Galaxy** (an open-source, web-based platform for data-intensive biomedical research) available at (https://usegalaxy.org/).

**DDBJ Center** provides sharing and analysis services for data from life science researches and advances science, available at (https://www.ddbj.nig.ac.jp/index-e.html).

## Results

### Case description

Our isolate was collected from 60 years old Saudi male visiting the foot unit at the university diabetes center at King Saud University. He was diagnosed with Type two diabetes mellites for more than 20 years with Neuropathy, Retinopathy, Obesity complications. His HbA1C was 7.7, and his BMI is 28.58. He came with a neuroischaemic diabetic foot ulcer beneath the left fifth metatarsal head with a wound surface area of one cm^2^, and the depth of the wound was 0.5 cm. The case of the DFU was under a callus, and the ankle-brachial pressure index (ABPI) was 0.61. The patient was on Insulin mixtard 30U AM/ 30U BT.

### Taxonomic and Genome description

Our *Streptococcus dysgalactiae* subsp. *equisimilis* strain SDSE- SCDR-1 complete genome was deposited in the NCBI-GenBank with GenBank accession number CP033391 under BioProject accession number PRJNA498183, BioSample accession number SAMN10285107 and assembly accession number GCA_009497975.1. The NCBI Taxon ID is 1334 Bacteria; *Terrabacteria* group; *Firmicutes*; *Bacilli*; *Lactobacillales*; *Streptococcaceae*; *Streptococcus*; *Streptococcus dysgalactiae* group; *Streptococcus dysgalactiae subsp. equisimilis*. The Multilocus sequence type (MLST) is *Streptococcus_dysgalactiae_equisimilis.*44. In silico analysis showed that SDSE strain SD- SCDR1 belongs to the emm *stG6.0* group. The genome size is 2179136, the GC Content is 39.87, the number of RefSeq CDS is 1843, and PATRIC CDS is 2065. The number of contigs is one, the Contig L50 is one, and Contig N50 is 2,179,136. The number of tRNAs is 57, rRNA is 15, misc-RNA is 14, regulatory sequences is 6, repeat regions is 90, CRISPR repeats is 35, and CRISPR spacers is 33. This genome was annotated using genetic code 11. The annotation included 375 hypothetical proteins and 1,671 proteins with functional assignments. The proteins with functional assignments included 561 proteins with Enzyme Commission (EC) numbers (Schomburg et al., 2004), 467 with Gene Ontology (GO) assignments (Ashburner et al., 2000), and 383 proteins that were mapped to KEGG pathways (Kanehisa et al., 2016). PATRIC annotation includes two types of protein families (Davis et al., 2016), and this genome has 1,910 proteins that belong to the genus-specific protein families (PLFams) and 1,914 proteins that belong to the cross-genus protein families (PGFams). **Figure 1** designates the graphical demonstration of the genome circulation and annotations. The Genome-genome comparison analysis showed that SDSE strain SD- SCDR1 is closely related to SDSE strain SK1250, strain WCHSDSE-1, and strain UT-5345, with distances values of 0.00399629, 0.00467416, and 0.00687462, respectively (**Table 1).** Whole Genome Alignment of strain SD-SCDR1 against reference genomes of strain NCTC7136, strain WCHSDSE-1 and Streptococcus pyogenes strain MGAS15252 is presented in **Figure 2.**

**Figure 1:**
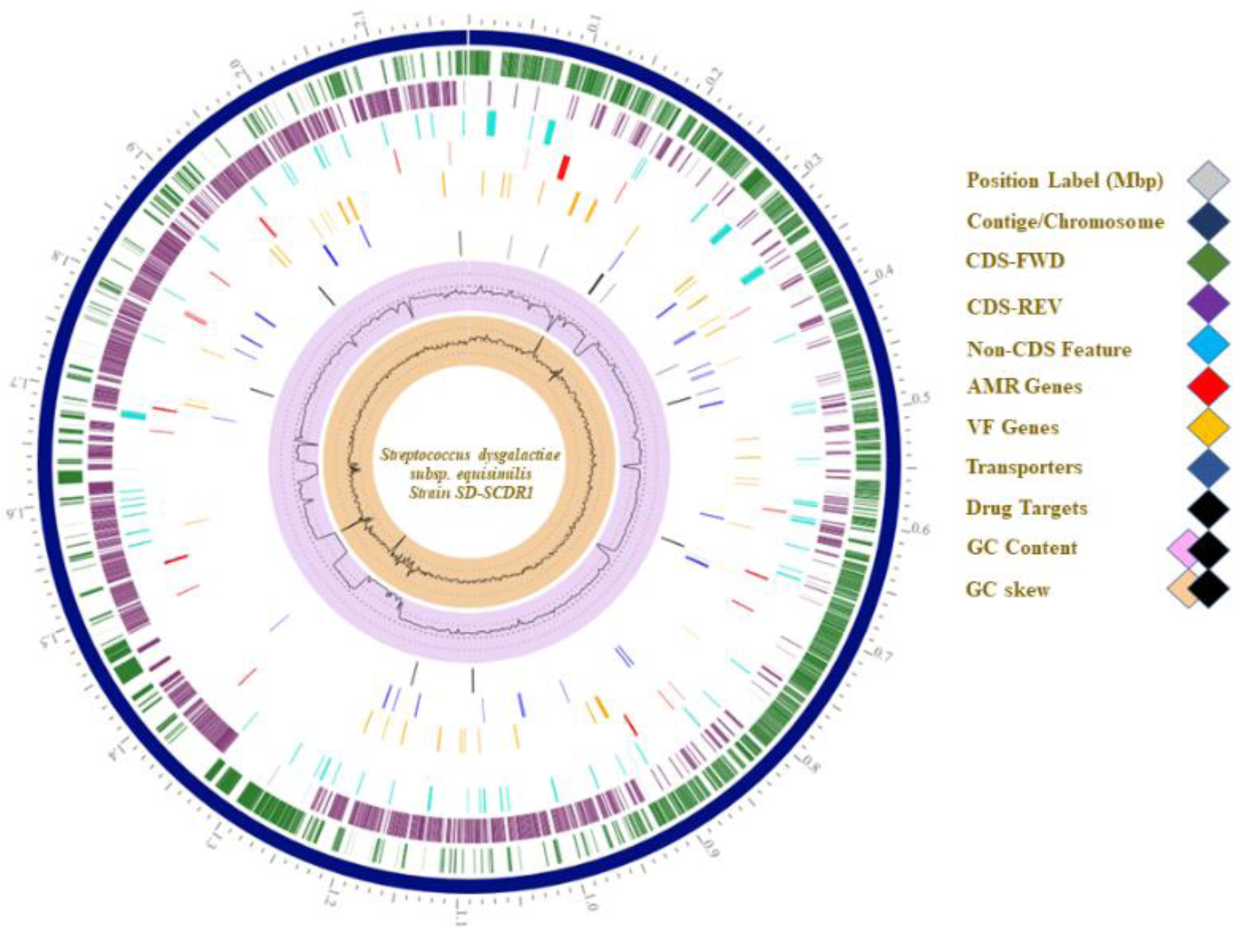
Genome representation shows from outer to inner rings, the contigs, CDS on the forward strand, CDS on the reverse strand, RNA genes, CDS with homology to known antimicrobial resistance genes, CDS with homology to known virulence factors, GC content, and GC skew.

**Figure 2:**
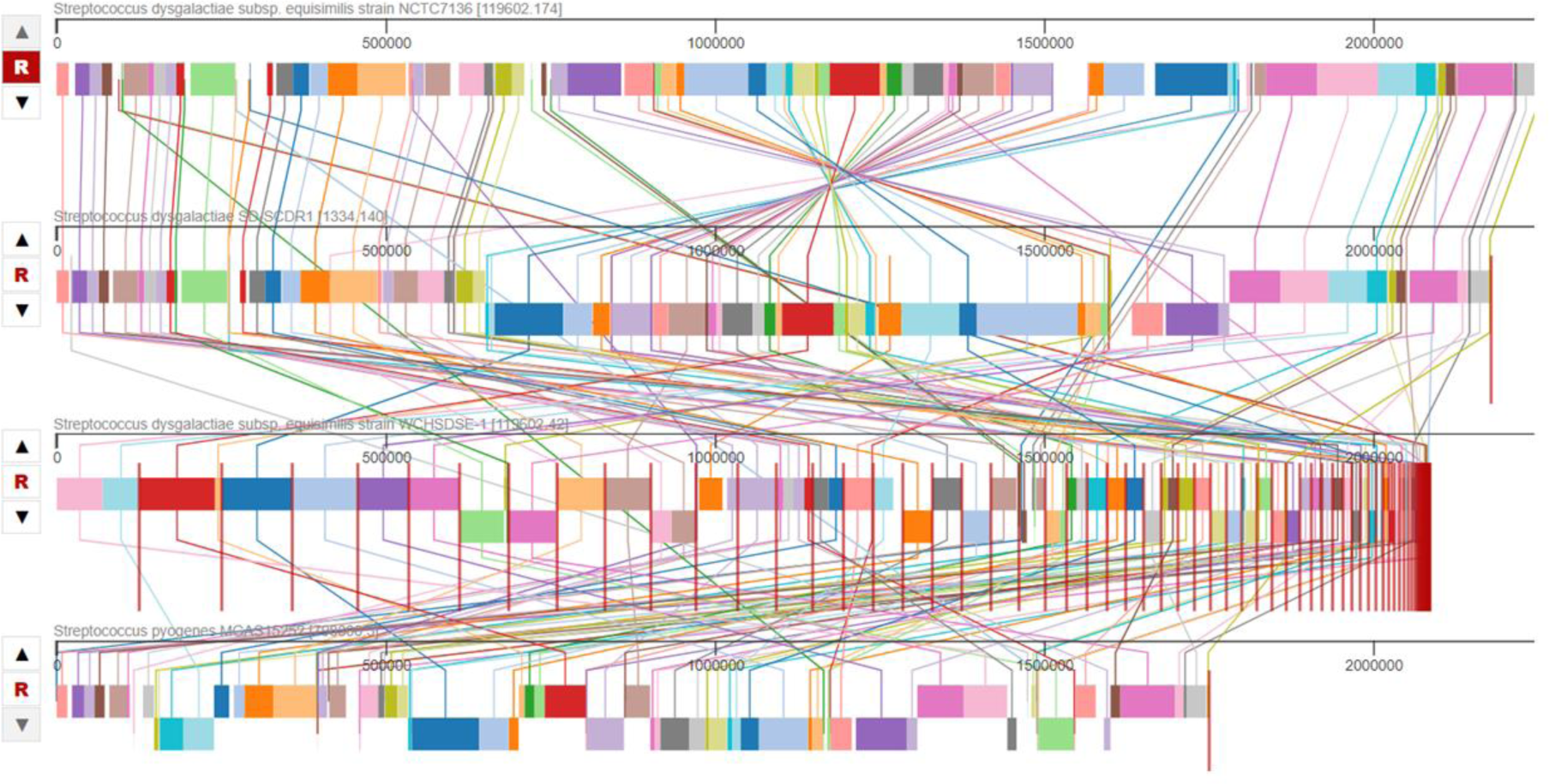
Mauve Multiple Whole Genome Alignment of SDSE strain SD-SCDR1 against reference genomes SDSE strain NCTC7136, SDSE strain WCHSDSE-1 and Streptococcus pyogenes strain MGAS15252.

### Biological Subsystems Analysis

A subsystem is a set of proteins that together implement a specific biological process or structural complex (Overbeek et al., 2005), and PATRIC annotation includes an analysis of the subsystems unique to each genome. An overview of the subsystems for this genome is provided in **Figure 3**. The highest number of genes (342) in our genome is dedicated to metabolism, followed by protein processing (204), then energy production (117). It is noteworthy to mention that 85 genes are dedicated to stress response, defense, and virulence.

**Figure 3:**
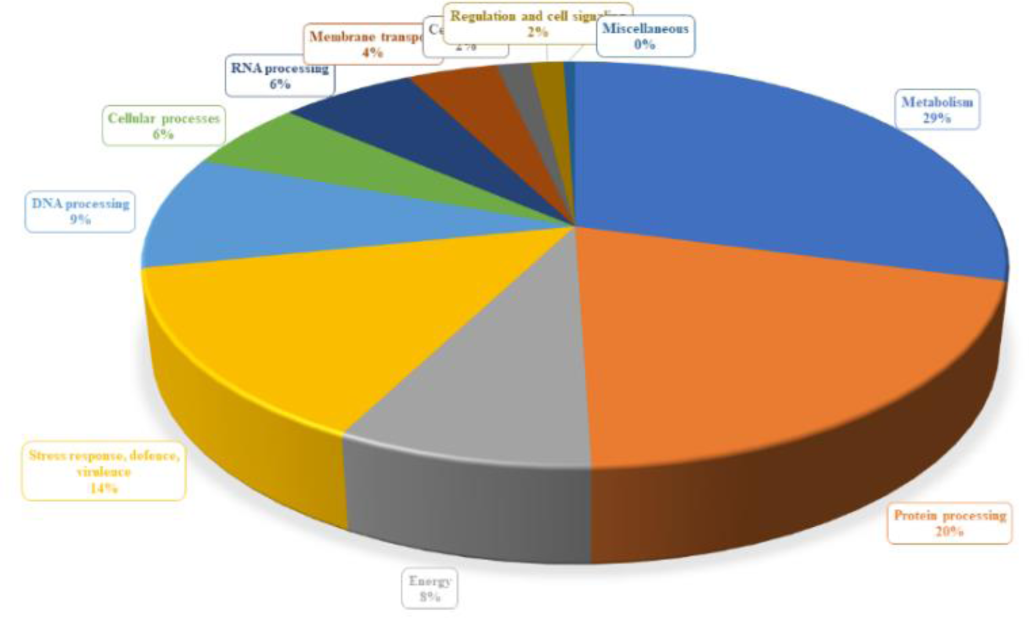
An overview of the biological subsystems and genes for Streptococcus dysgalactiae subsp. equisimilis strain SD-SCDR1 genome

### Phylogenetic analysis

Whole-genome phylogenetic analysis showed that our strain SD-SCDR1 belongs to a clade that contains *Streptococcus dysgalactiae* subsp. *equisimilis* strains, namely the reference strain AC-2713 and GGS 124 (**Figure 4a)**. Besides, within *Streptococcus dysgalactiae* subsp. *equisimilis* taxa, our strain SD-SCDR1 is closely related to both WCHSDSE-1 and strain SK1250 **(Figure 4b)**.

**Figure 4:**
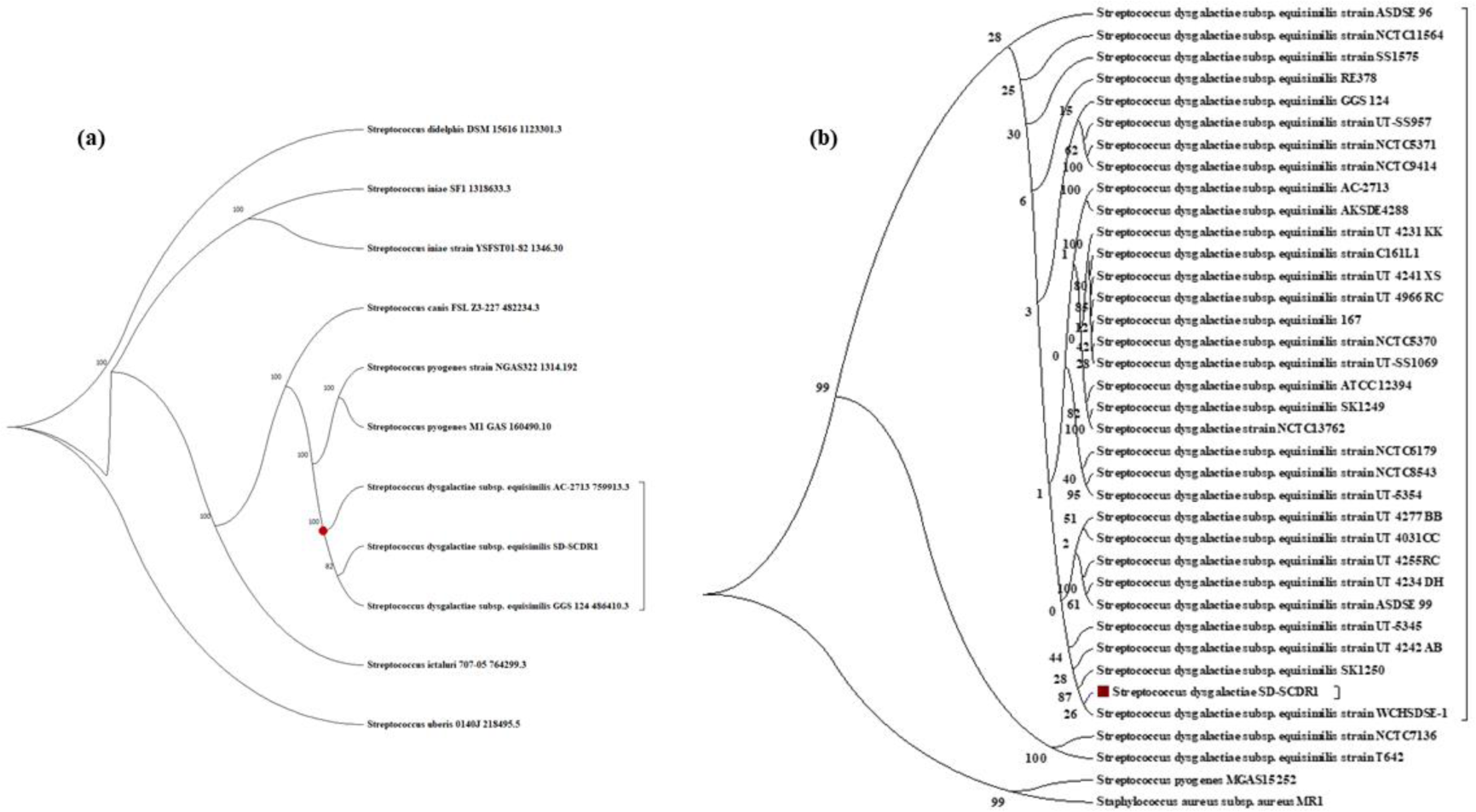
(a) Phylogenetic tree of SDSE based on whole-genome sequences (reference and outgroup genomes) and (b) selected 33 SDSE genome sequences, Streptococcus dysgalactiae subsp. equisimilis strain SD-SCDR1, Streptococcus pyogenes MGAS15252, and Staphylococcus aureus MR1 genomes

### Specialty Genes investigation

Genome annotation of Our SD-SCDR-1 strain showed that it contains many genes with homology to known virulence factors, antibiotic resistance genes, and drug targets.

### Virulence factors genes

**Table 2** represents the observed virulence factors genetic determinants in the SDSE-SCDR-1 genome. Our strains contain genetic determinates involved in adherence to the host cell, cellular invasion, Antiphagocytosis, immune evasion, and skin and soft tissue invasion. In addition to genes responsible for host mortality and tissue damage, toxins, pore-forming proteins, cytotoxins, beta-hemolysis agents, stress resistance thermotolerance, oxidative stress resistance, growth in blood and lung, and many other metabolic, genetic determinants that are crucial for the pathogenesis and virulence.

### Antimicrobial Resistance Genes

A summary of the antimicrobial resistance genes (AMR) genes annotated in our genome and the corresponding AMR mechanism is provided in **Table 3**. It contains resistant genes the confer resistance against 4-quinolones (Fluoroquinolones), Aminocoumarin, Aminoglycoside (Streptomycin), Cycloserine, Daptomycin, Elfamycin (Pulvomycin), Fosfomycin, Fusidic acid, Isoniazid, Mupirocin (Pseudomonic acid), Rifampicin, Vancomycin, Daptomycin, Streptomycin Sulfonamide antibiotics, Tigecycline, Triclosan, and Trimethoprim. It is noteworthy to mention that our strain contains a gene that encodes for the multidrug resistance protein ErmB that confer resistance against Lincosamides, Macrolides, Streptogramins, and Tetracyclines.

### Drug target genes

**Table 4** represents the potential novel drug target determinants in the SDSE-SCDR-1 genome. SDSE-SCDR-1 genome contains 12 different potential novel drug targets (genes) namely, ***metG***, ***glpO***, ***fusA***, ***rplC***, ***parE*** (***gyrB***), ***guaB***, ***rmlC***, ***murI***, ***proC***, ***relA***, ***def***, and ***rmlB***. Against these targets, six experimental drugs were suggested, such as Inosinic Acid, Merimepodib, and Guanosine-5’-Diphosphate. Two were in the investigational stage, namely, Gatifloxacin and REP8839. Moreover, five approved drugs target four-drug targets: Procaine benzylpenicillin, Retapamulin, Spiramycin, Sulfaphenazole, and Curcumin.

### CRISPR and Cas genes/systems information

**Tables 5, 6, and 7** describe CRISPR and Cas genes/systems identified in the SDSE-SCDR-1 genome. Two CRISPR arrays were identified that are consist of 35 CRISPR repeats and 33 CRISPR spacers. Although four potential CRISPR repeats were identified, namely SCDR1-CRISPR1-4, only two gained the highest confidence (evidence) level of 4, which are SCDR1-CRISPR 2 and 3. The length of SCDR1-CRISPR2 is 1286 bases, the repeat length is 36, the spacer length is 19 bases, and the conservation of repeats is 100%. Whereas the length of SCDR1- CRISPR 3 is 962 bases, the repeat length is 32, the spacer length is 14 bases, and the conservation of repeats is 90.625 %. Two CAS systems were observed in the SDSE-SCDR-1 genome, namely, CAS-TypeIIA and CAS-TypeIC. CAS-TypeIIA contains Csn2-0-IIA as a mandatory gene and Cas2-0-I-II-III, Cas1-0-II, and Cas9-1-II (endonuclease Cas9) as accessory genes. While, CAS-TypeIC contains Cas1-0-IC, Cas7-0-IC, Cas8c-0-IC (Csd2/Csh2 family), Cas5-0-IC as mandatory genes, and Cas2-0-I-II-III-V, Cas4-0-I-II (RecB family exonuclease Cas4) and Cas3-0-I as accessory genes.

### Prophage and transposons insertion sequences

**Figures 5 and 8** present the 36.4Kb incomplete prophage containing 19 CDS that encodes for phage proteins. The GC% of the prophage is 36.25 and starts with the left attachment site (***attL***) at CDS position 2119960 and ends with the right attachment site (***attR***) at CDS position 2156433. Our prophage contains protein-coding sequences like CDSs from Lactococcus phage, Javan S145 prophage, Listeria phage, Bacillus virus G, Bacillus phage TsarBomba, and Streptococcus phage 315.1. Observed CDSs encodes for phage antirepressor protein, repressor protein, helix-turn-helix transcriptional regulators, site-specific integrase, transposase, DNA helicase, and phosphodiesterase.

**Figure 5:**
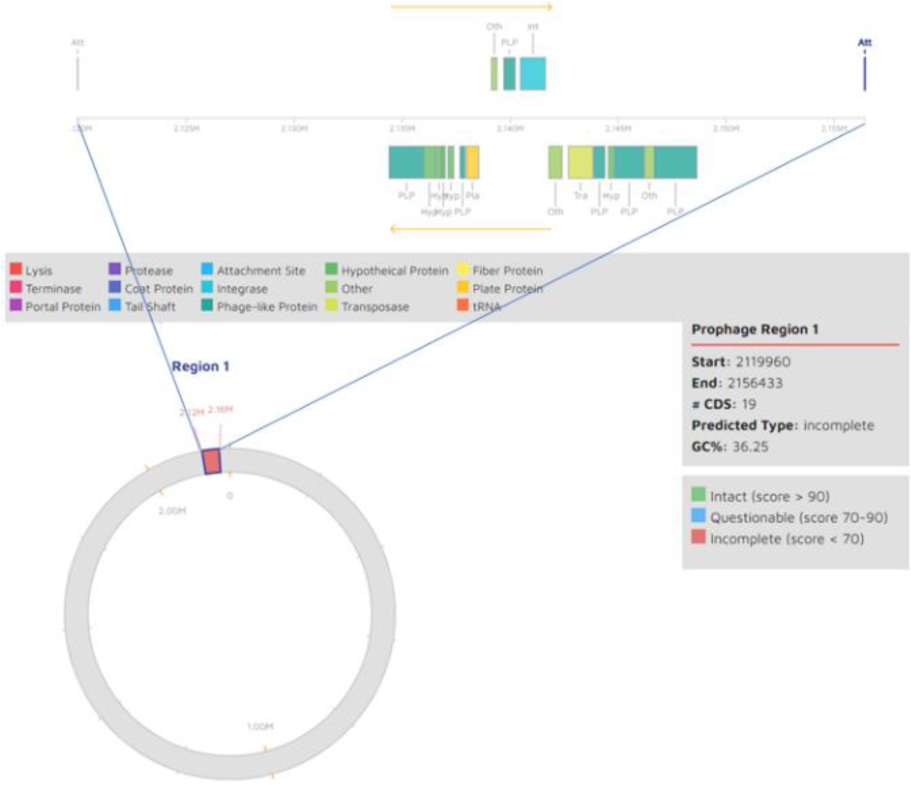
Incomplete prophage region identified in Streptococcus dysgalactiae subsp. equisimilis strain SD-SCDR1 genome

Furthermore, 40 Insertion sequences were observed in the SDSE-SCDR1 genome **(Table 9)**. Amongst these, 20 belong to the insertion sequence (IS) family IS3, eight belongs to IS family IS30, five belongs to the insertion sequence IS family IS1182, four belongs to IS family IS982, two belong to IS family ISAs1, and one belongs to IS family IS481. 36 IS have a *Streptococcus* species origin, two have a *Lactococcus* species origin, and two have an *Actinobacillus* species origin.

### Streptococcus dysgalactiae subsp. equisimilis pan-taxonomic Comparative genomics section

**Supplementary table 1** represents the taxonomic/genomic information of the 55 publicly available SDSE strains/isolates. These include genome name, NCBI taxon ID, genome status, strain MLST, other available typing serovar information, BioProject accession number, BioSample accession number, genome assembly accession number, GenBank accessions number, number of contigs per genome, genome length, GC% content, isolation Source, and host-name. These included both SDSE GGS_124 and AC-2713 strains as reference/representative genomes. We found that both SDSE strain NCTC11565 and strain NCTC5969 genomes are of low quality and should be revisited or removed from the public databases by the submitting authors. For the NCTC11565 genome, the genome quality was inferior; the length was too long, the CDS count was very high (6041 RefSeq CDS), and with a low **PLfam** CDS ratio. We believe that this genome is mixed with a *Pseudomonas aeruginosa* DNA similar to the PAO1 strain. The NCTC5969 genome’s length was too long; the CDS count was high (2850 RefSeq CDS). We believe that this genome is mixed with *Staphylococcus aureus* DNA. Thus, both strains were excluded from our comparative genomics analysis. Nonetheless, the majority of isolates were of human origin; two isolates were collected from horses (genital tract), one isolate was collected from a monkey (throat), and one is from a dog (pleural exudate). Sites of isolation included blood cultures, sterile sites of invasive Diseases, oral, throat (Pharyngitis) or upper respiratory samples, and foot ulcers and Joint fluid samples.

### SDSE strains protein families based phylogenetic analysis

**Figure 6** represents protein families based Maximum likelihood phylogenetic analysis for 60 genomes comprising the publicly available SDSE strains/isolates and related outgroup taxa showed that our isolate (SD-SCDR1) is located in a clade that contains 10 SDSE strains, namely, SK1250, WCHSDSE-1, KNZ01, UT 4242 AB, NCTC11556, NCTC6407, NCTC10321, NCTC6179 and NCTC6181 strains with a bootstrap value of 79. The reference/representative strain SDSE-AC 2713 is located in a clade containing 11 SDSE strains, namely, UT-SS957, NCTC9414, NCTC5371, AKSDE4288, NCTC9603, UT 4255RC, NCTC11557, UT 4234 DH, ASDSE 99, and NCTC11555 strains with a bootstrap value of 91. To build this tree, the number of the protein alignments was 227, the number of aligned amino acids was 72333, the number of CDS alignments was 227, and aligned nucleotides was 210171.

**Figure 6:**
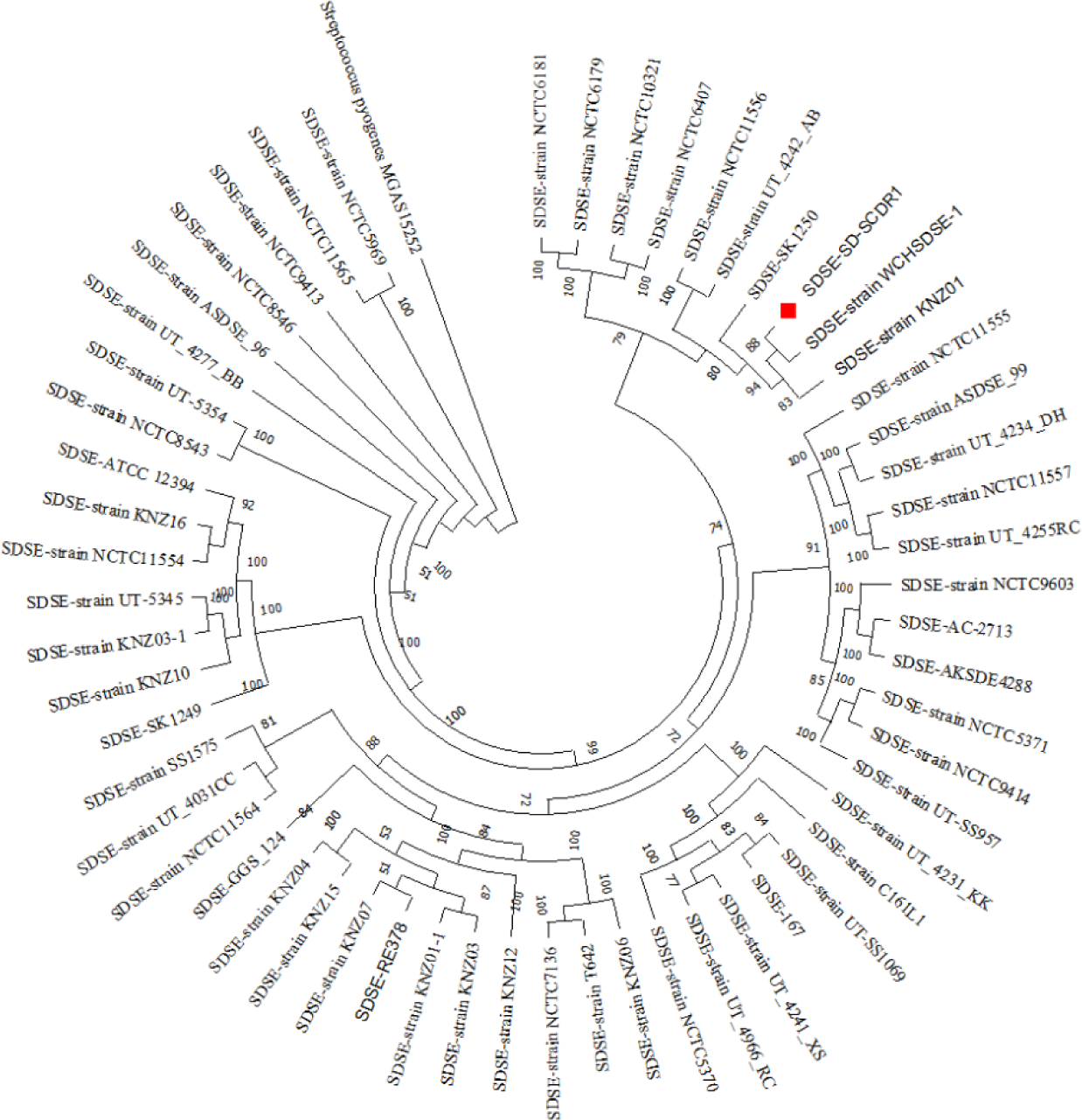
Protein families based on Maximum likelihood phylogenetic analysis for the publicly available SDSE strains/isolates genomes.

**Figure 7 (a and b)** represents protein families phylogeny based proteomic comparison among SDSE-SD- SCDR1 clade containing strains against the reference strain SDSE-AC2713 in both all gene/proteins **(a)** and gene/proteins essential for the bacterial survival **(b)**. These are essential 189 genes recognized for strain SDSE-AC 2713 predicted using LB media’s metabolic flux balance analysis (**Supplementary Table 2**) **(Figure 7b)**. Generally, a high degree of conservation (≥ 90%) was observed in the four tested strains in percent of gene/protein identity and sequence coverage compared to the reference strain SDSE-AC2713. For SDSE-SD-SCDR1 strain only three gene/proteins showed less conservation namely, Pyrroline-5- carboxylate reductase (EC 1.5.1.2) (***proC***) with a sequence coverage of 87%, Isochorismatase (EC 3.3.2.1) with a sequence coverage of 69% and Aspartyl-tRNA (Asn) amidotransferase subunit C (EC 6.3.5.6) @ Glutamyl-tRNA(Gln) amidotransferase subunit C (EC 6.3.5.7) (***gatC***) with a sequence coverage of 87%. Whereas, SDSE SK1250 showed reduced conservation in only two gene/proteins, namely, Undecaprenyl- diphosphatase (EC 3.6.1.27) (***uppP***) with a sequence coverage of 87% and Heptaprenyl diphosphate synthase component I (EC 2.5.1.30) with a sequence coverage of 89%. While in strain KNZ01, four gene/proteins showed less conservation, namely, Threonine synthase (EC 4.2.3.1) (***thrC***) with a percent of gene/protein identity of 70%, Pyrroline-5-carboxylate reductase (EC 1.5.1.2) (***proC***) with a sequence coverage of 87%, Isochorismatase (EC 3.3.2.1) with a sequence coverage of 86% and Heptaprenyl diphosphate synthase component I (EC 2.5.1.30) with a sequence coverage of 89%. For SDSE WCHSDSE- 1, only three gene/proteins showed less conservation, namely, Pyrroline-5-carboxylate reductase (EC 1.5.1.2) (***proC***) with a sequence coverage of 87%, a hypothetical protein (Isochorismatase (EC 3.3.2.1) with a sequence coverage of 86%, and Heptaprenyl diphosphate synthase component I (EC 2.5.1.30) with a sequence coverage of 89%. However, the major four regions showed less conservation (≤ 60% identity and/or coverage) in the non-essential regions of the genome/proteome (**Figure 7b),** namely, 1-4 regions. Region one (starts 265508-ends 287829) contains mobile elements, hypothetical proteins **(*int* and *manR*)**, Integrase gene **(*int*),** Lantibiotic salivaricin A gene and its associated genes/proteins (***salA, salB, salX, salY, salK, salR***), Antitoxin HigA (***pheT***), Toxin HigB, Transposase, PTS system, galactitol-specific IIA component, galactitol-specific IIC component and galactitol-specific IIB component (EC 2.7.1.200), and Triosephosphate isomerase (EC 5.3.1.1) (***tpiA***). Competence-stimulating peptide ABC transporter ATP- binding protein ComA was found in SDSE SD-SCDR1, WCHSDSE-1, and SK1250. Region two (starts 1240732– ends 1345833) contains Transposase TnpA **(*tnpA*)**, mobile element proteins (***tra*)**, site-specific recombinase **(*ccrB*)**, DNA topoisomerase I (EC 5.99.1.2), TrsK-like protein, DNA polymerase IV (EC 2.7.7.7), and replicative DNA helicase (***dnaB***) (EC 3.6.4.12). Besides, several phage proteins were detected including, phage integrase, phage-associated cell wall hydrolase, Holin, phage tail length tape-measure protein T, prophage pi2 protein 37, phage major capsid protein, phage head, head-tail preconnector protease C, phage portal protein, phage terminase, large subunit, phage terminase, small subunit, and phage endonuclease. The third region (starts 1414176– ends 1471124) contains several hypothetical proteins and several phage proteins, such as phage integrase, site-specific recombinase (s), phage-associated cell wall hydrolase, Holin, phage capsid and scaffold, phage endopeptidase, phage tail length tape-measure protein T, prophage pi2 protein 37, phage major capsid protein, prophage Clp protease-like protein, phage portal protein, phage terminase, large subunit, phage terminase, small subunit, phage protein, contains HNH endonuclease motif, phage endonuclease, DNA helicase (EC 3.6.4.12), phage-associated, DNA primase, phage associated, and phage-encoded DNA polymerase I (EC 2.7.7.7). It contains putative UV-damage repair protein UvrX, DNA polymerase IV (EC 2.7.7.7), SWF/SNF family helicase, and S-adenosylmethionine synthetase (EC 2.5.1.6). The fourth region (starts 1597335– ends 1633454) contains numerous hypothetical proteins (***iga***, ***yddE***, ***sspB, int***), mobile element proteins (***tra*),** transcriptional regulator (s), Xre family, Serine endopeptidase ScpC (EC 3.4.21.-), Antirestriction protein, phage repressor protein, conjugal transfer protein, and phage integrase protein.

**Figure 7:**
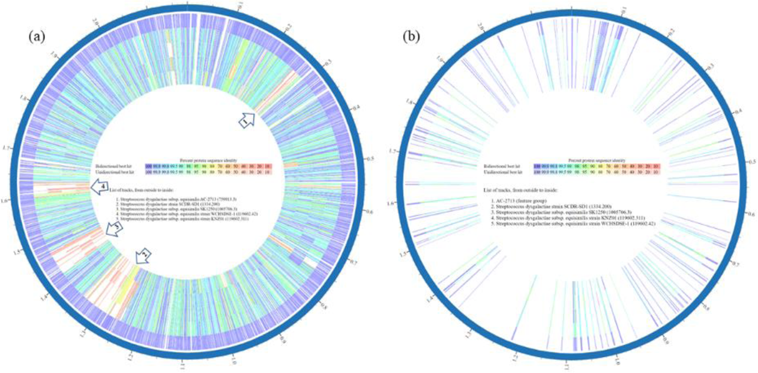
Protein families phylogeny based proteomic comparison among SDSE-SD-SCDR1 clade containing strains (a) All gene/proteins (b) Essential gene/proteins

### *Streptococcus dysgalactiae subsp. equisimilis* pan-taxonomic Antimicrobial Resistance Genes (Resistome)

**Supplementary table** 3 presents antimicrobial resistance genes (AMR) genes annotated in SDSE genomes and their corresponding AMR mechanism. All tested SDSE genomes contains ***folA***, ***Dfr***, ***Ddl***, ***LiaS***, ***LiaR***, ***pgsA***, ***Alr***, ***kasA***, ***gidB***, ***EF-G***, ***EF-Tu***, ***folP***, ***gyrA***, ***gyrB***, Iso-tRNA (***ileS***), ***LiaF***, ***S12p***, ***S10p***, ***rpoB***, ***rpoC***, ***MurA***, and ***FabK***. Besides, all tested SDSE genomes contain ***lmrP*** except for two strains, namely, T642 and C161L1. All the strains hold ***aadK*** except for five strains, namely, NCTC6179, NCTC6181, NCTC9413, NCTC6407, NCTC10321. ***rplF*** gene was missing from 22 SDSE strains, namely, NCTC8543, NCTC11564, SD-SCDR1, NCTC6181, NCTC8546, NCTC9413, NCTC11554, NCTC 11556, NCTC11565, NCTC6407, NCTC10321, NCTC5969, NCTC9603, NCTC11555, NCTC11557, KNZ10 KNZ04, KNZ07, KNZ06, KNZ12, KNZ15, and KNZ16. Some SDSE strains showed some unique AMR phenotypes and genotypes. It was found that only SDSE strains AKSDE4288, strain ASDSE99, strain NCTC11557, and strain UT_4255RC have ***ermA***. Moreover, only strains SDSE KNZ04 and WCHSDSE- 1 have ***ermB***. Also, only two *Streptococcus dysgalactiae* subsp. *equisimilis* strain Sde558 has ***erm(T)***. SDSE strain AC-2713 has ABC-F type ribosomal protection protein **Msr(D)** that confer resistance to the antibiotic telithromycin and MFS efflux pump, **Mef(A)**, which is responsible for resistance against Macrolides, Lincosamides, Streptogramins, Ketolides, Oxazolidinones (MLSKO). Moreover, SDSE strains AKSDE4288, and SK1250 have the tetracycline resistance gene, ***tetO***, that encodes for the ribosomal protection type (O) Tetracycline resistance. Whereas, SDSE strains NCTC11557, UT_4242_AB, UT_4234_DH, KNZ04, KNZ06, ASDSE_99, and ASDSE_96 have Tetracycline resistance gene ***tetM***, that encodes for the ribosomal protection type (M) Tetracycline resistance. Furthermore, SDSE strain KNZ06 has a ***parC*** gene that confers resistance against fluoroquinolone antibiotics. The SDSE strain T642 a point mutation in ***ParE*,** a subunit of topoisomerase IV that decatenates and relaxes DNA to allow access to genes for transcription or translation that prevent Anticoumarin antibiotics from inhibiting DNA synthesis, and thus conferring resistance. Besides, it has the PhoB, phosphate regulon transcriptional regulatory protein (***SphR***), a direct regulator of the macAB efflux pump antibiotic out of a cell to confer resistance. Similarly, SDSE strain C161L1 has **PhoB** resistance against peptide antibiotics. Moreover, strain C161L1 has the ***gyrB*** (parE) gene that confers resistance against aminocoumarin (novobiocin).

### *Streptococcus dysgalactiae subsp. equisimilis* pan-taxonomic virulence factors Genes (Virulome)

**Supplementary table 4** presents virulence factors (**VF**) genes annotated in SDSE genomes and related products. The core Virulome of SDSE consists of 38 genetic determinants. All tested SDSE genomes contains the following virulence factors ***idnO***, ***purB***, ***asnA***, ***clpP***, ***bcaT (ilvE)***, ***ccpA***, ***SpyM3-0013***, ***pyrG***, ***cydA***, ***dltA***, ***BCHE***, ***hasC (galU)***, ***vicK***, ***gidA***, ***covS***, ***lepA***, ***luxS***, ***SPy_1633***, ***ykqC (rnjA)***, ***SP_0095***, ***pflD***, ***lgt***, ***deoB***, ***metQ (atmB)***, ***lsp (lspA)***, ***leuS***, ***purH***, ***guaA***, ***glnA***, ***fba***, ***pstA***, ***pstB***, ***perR***, ***oppA***, ***cpsY***, ***fbp54 (fbp, flpA)****, **perR**.* Moreover, all examined strains contain ***rpoE*** gene, encoding for DNA-directed RNA polymerase delta subunit (EC 2.7.7.6), except for strain 167. Furthermore, all tested strains contain the ***gldA*** gene, encoding for Glycerol dehydrogenase (EC 1.1.1.6), except for two strains: T642 and C161L1. Similarly, all examined SDSE strains contain ***slo*** gene, encoding for Thiol-activated cytolysin/streptolysin O, except for two, namely, NCTC6407 and NCTC10321. Additionally, all examined strains contain the ***lmb*** (***psaA***) gene, which encodes for Laminin-binding surface protein, except for two strains: strain NCTC6407 and strain NCTC10321. All SDSE strains contain both ***scpA*** and ***scpB***, encoding for C5a peptidase (EC 3.4.21.-), except for four strains, namely, NCTC6179, NCTC6181, NCTC6407, and NCTC10321. Besides, all SDSE strains contain the ***ska (skc)*** gene, encoding for Streptokinase, except for four strains, namely, NCTC6407, NCTC10321, NCTC6181, and NCTC6179. Additionally, all SDSE strains contain the ***nga*** gene, encoding for Nicotine adenine dinucleotide glycohydrolase (NADGH) (EC 3.2.2.5), except for four strains NCTC6407, NCTC10321, NCTC6181, and NCTC6179. Moreover, 87% of SDSE strains contain ***sagA***, which encodes for Streptolysin S precursor. Additionally, only 68% of SDSE strains contained ***gbs0630*** and ***gbs0631*** genes. Similarly, only 53% of SDSE strains contain ***soS1A*** that encodes for Ethanolamine utilization protein. Besides, only 51% of SDSE strains contain ***csoS1A*** that encodes for Ethanolamine utilization polyhedral-body-like protein (EutM). Only 47% of SDSE strains contain speg (***spegg4***) gene encodes for Exotoxin, phage associated streptococcal pyrogenic exotoxin G (SpeG). Only 32% of SDSE strains contain ***gbs0632*** that encodes for the adherence sortase-assembled pili hypothetical protein. Furthermore, only 43% of SDSE strains contain the ***ccrB*** gene that encodes for site- specific recombinase. Likewise, only 28% of SDSE strains contain ***SP_0939*** genetic determinant that encodes for DUF1706 domain-containing protein. Interestingly, some SDSE strains showed some VF unique resistance phenotypes and genotypes. Only eight SDSE strains have the ***sda*** gene that encodes for Streptodornase D, the exoenzyme and spreading factor, namely, 167, KNZ16, NCTC11554, UT_4031CC, UT_4242_AB, UT_4277_BB, UT_4966_RC, and WCHSDSE-1. Six strains contain ***SP_1018*** gene which encodes for Thymidine kinase (EC 2.7.1.21) namely, NCTC6181, NCTC10321, NCTC6407, UT_4031CC, ASDSE_96 and NCTC6179. Moreover, six SDSE strains contain ***mf3*** that encodes for the Streptococcal extracellular nuclease 3 (Mitogenic factor 3), namely, NCTC8543, NCTC9414, UT_4231_KK, UT_4241_XS, UT-SS1069, and SK1249. Four SDSE strains have ***emm1/emm*** encodes for Antiphagocytic M protein that plays an important role in Antiphagocytosis, Adherence, and Cellular invasion. These strains are ASDSE_96, KNZ06, KNZ16, and NCTC5371. Similarly, Four SDSE strains have ***salT/ salX*** gene that encodes for Lanthionine biosynthesis protein/ Lantibiotic transport ATP-binding protein, namely, KNZ03, KNZ04, KNZ16, and AC-2713. Four SDSE strains have gene ***EF1623*** encodes for EutM/PduA/PduJ-like protein two clustered with choline trimethylamine-lyase NCTC11554, KNZ03, KNZ04, and KNZ16. Only three isolates have ***SP_1143*** that encodes for Toxin HigB, namely, strains NCTC11564, NCTC8543, and UT-5354. Similarly, three SDSE strains have the ***purl*** gene that encodes for Phosphoribosylformylglycinamidine synthase, glutamine amidotransferase subunit (EC 6.3.5.3), namely, UT-SS1069, C161L1, and strain T642. Additionally, only two strains have the ***mf/spd*** gene that encodes for Streptodornase B, the exoenzyme and spreading factor, namely, NCTC6179and NCTC6181. Additionally, only two strains have the ***rumA1*** gene encoding RNA methyltransferase, TrmA family, namely, AKSDE4288 and AC-2713. Only SDSE strain RE378 has the ***pyrR*** gene that encodes for Pyrimidine operon regulatory protein. Only SDSE strain SK1249 has the ***pepF*** gene that encodes for Oligoendopeptidase F-like protein. Only SDSE strain KNZ16 has the ***lacA*** gene that encodes for Galactose- 6-phosphate isomerase, LacA subunit (EC 5.3.1.26). Only SDSE strain NCTC6179 has the ***spot*** gene thatencodes for Guanosine-3’,5’-bis(diphosphate) 3’-pyrophosphohydrolase (EC 3.1.7.2) / GTP pyrophosphokinase (EC 2.7.6.5), (p)ppGpp synthetase II. Only SEDE strains C161L1 and T642 have the following genes: ***pstC*** gene that encodes for phosphate ABC transporter, permease protein, ***csrR*** gene that encodes for Response regulator CsrR, ***ciaR*** gene that encodes for two-component system response regulator CiaR, the ***txrR*** gene that encodes for two-component response regulator TrxR, and ***trmFO*** gene that encodes for Methylenetetrahydrofolate--tRNA-(uracil-5-)-methyltransferase TrmFO (EC 2.1.1.74) virulence factor. ***Streptococcus dysgalactiae subsp. equisimilis* pan-taxonomic drug targets: Supplementary table 5** presents drug target genes/products annotated in SDSE genomes and proposed drug. All tested SDSE genomes contain the following drug targets, ***def***, ***guaB***, ***gyrB***, ***murI***, ***relA***, ***rmlB***, ***rmlC***, and ***rplC***. Additionally, the ***parE*** gene is observed in all SDSE genomes/strains except for three strains: NCTC7136, NCTC9603, and KNZ06. Additionally, the ***lacG*** gene, which encodes 6-phospho-beta- galactosidase (EC 3.2.1.85) and targeted by beta-D-galactose 6-phosphate, is found in 43% of SDSE genomes/strains. Furthermore, the ***spg*** gene, which codes for Protein G-related-alpha-2 macroglobulin- binding protein (GRAB) and targeted by N-Formylmethionine, is 30% of SDSE genomes/strains. Furthermore, SDSE strain KNZ06 has the ***parC***, which encodes for DNA topoisomerase IV subunit A (EC 5.99.1.3) and is targeted by both Gatifloxacin and Besifloxacin. Finally, SDSE strain SK1249 has Catabolite control protein A and is targeted by the experimental drug 2-Phenylamino-Ethanesulfonic Acid.

### Streptococcus dysgalactiae subsp. equisimilis pan-taxonomic Complete Phage genomes

**Supplementary table 6** and **supplementary figure 1** presents detailed information about the complete phage genomes annotated in SDSE genomes. To date, 16 complete phages are associated with SDSE strains. The species representative SDSE strain AC-2713 has two phages, namely, *Streptococcus* phage Javan122 and Javan123. In comparison, SDSE UT_4231_KK has two phages: *Streptococcus* phage Javan161 and *Streptococcus* satellite phage Javan162. Similarly, SDSE UT_4966_RC has two phages, namely, *Streptococcus* phage Javan164 and Streptococcus satellite phage Javan165. The SDSE strain UT- SS1069 also has two complete prophages: *Streptococcus* phage Javan170 and *Streptococcus* satellite phage Javan171. Only SDSE strain UT-5345 has three complete phages: Streptococcus phage Javan166, Streptococcus satellite phages Javan167 Javan168. SDSE strains SDSE NS3396, SDSE strain ASDSE_99, SDSE UT_4242_AB, SDSE UT-5354, SDSE WCHSDSE-1 each has one complete phage genome. The SDSE associated phage genome lengths ranged between 8144 and 45241 bp for phage Javan167 and phage Javan166, respectively. Besides, the phages GC% content ranged between 33.3 and 41.6. Furthermore, the number of protein-coding sequences (CDS) ranged between 12 and 85 CDSs for Streptococcus satellite phage Javan167 and Streptococcus phage Javan166, respectively.

### Streptococcus dysgalactiae subsp. equisimilis pan-taxonomic CRISPER-CAS systems

**Supplementary table 7** presents detailed information about the CRISPER-CAS systems observed in SDSE genomes. We found that 89% of SDSE strains/genomes have CRISPER-CAS systems associated with genes/products. Two CAS systems were observed in SDSE genomes, namely, CAS-TypeIIA and CAS-TypeIC. The core SDSE-CAS systems in SDSE contains CRISPR-associated endonuclease Cas9, CRISPR-associated helicase Cas3, CRISPR-associated protein Cas1 (II), CRISPR-associated protein Cas1 (IC), CRISPR-associated protein Cas2 (I-II-III), CRISPR-associated protein Cas2 (I-II-III-V), CRISPR-associated protein Cas5, CRISPR-associated protein, Csd1 family, CRISPR-associated protein, Csd2/Csh2 family, CRISPR-associated protein, Csn2 family and CRISPR-associated RecB family exonuclease Cas4. <u>CAS-TypeIIA contains</u> Csn2-0-IIA as a mandatory gene and Cas2 (I-II-III), Cas1-0-II, and Cas9 (1-II)(endonuclease Cas9) as accessory genes. While CAS-TypeIC contains Cas1(-0-IC), Csd2/Csh2 family (Cas7-0-IC, Cas8c-0-IC), Cas5(-0-IC) as mandatory genes and Cas2 (I-II-III-V), RecB family exonuclease Cas4 (Cas4-0-I-II), and Cas3(-0-I) as accessory genes. We found that 36 (68%) SDSE strains comprises the full core SDSE-CAS systems, namely, SS1575, NCTC7136, NCTC8543, SD-SCDR1, NCTC11554, NCTC11556, NCTC11555, NCTC11557, KNZ03, KNZ01, KNZ10, KNZ04, KNZ06, KNZ03, KNZ12, KNZ15, WCHSDSE-1, UT_4277_BB, UT_4234_DH, ASDSE_99, UT_4242_AB, ASDSE_96, UT_4255RC, AKSDE4288, AC-2713, RE378, SK1250, UT-5345, UT-5354, KNZ07, KNZ16, UT_4031CC, T642, NCTC6179, ATCC 12394, and NCTC6181. Besides, seven SDSE strains (13%) contain only CAS-TypeIIA, namely, NCTC5371, NCTC9414, NCTC6407, NCTC10321, NCTC9603, UT-SS957, GGS_124. We were not able to detect any CRISPER-CAS systems associated genes/products in six SDSE strains/genome (11%) namely, 167, C161L1, NCTC5370, UT_4231_KK, UT_4241_XS, UT_4966_RC, and UT-SS1069. Four strains showed different patterns of missing and/or gaining CAS genes/proteins, such as NCTC11564 that is missing both CRISPR-associated helicase Cas3 and CRISPR-associated protein, Csd1 family.

## Discussion

In the present work, we present a comprehensive genomic analysis for the SDSE strain SCDR1 that belongs to Lancefield group G, emm type (*stG*6), and (MLST) sequence type (ST44) isolated from neuroischaemic diabetic foot ulcer. To our knowledge, this is the first time for this emerging pathogen to be documented in Saudi Arabia and the middle east. This isolate was characterized by its extreme beta-hemolytic ability (data not shown). Matsue et al. (Matsue et al., 2020) stated that predominant *emm* types of SDSE varied per country. For example, in central Taiwan, the *stG6.1*, *stG10*, *stG245*, *stG840*, and *stG652* strains predominated (Ma et al., 2017), whereas *stG485* and *stG6.1* were predominant in Jerusalem and Austria, respectively (Cohen-Poradosu et al., 2004; Leitner et al., 2015). Moreover, Shimomura et al. reported that *stG10.0* was the most common emm type in Portugal and Japan, *stG643.0* was the most common in western Norway, and *stG6.0* was the most recurrent in the USA (Shimomura et al., 2011). Rantala et al. conducted a retrospective population-based study of 140 episodes of *Streptococcus dysgalactiae* subsp. *equisimilis* bacteremia occurring in Finland during 1995–2004. Skin and soft tissue infections were more common clinical signs amongst cases caused by frequent emm types. They reported 18 *emm* types (including four *stG6: stG6.0, stG6.1, stG6.3*, and *stG6.4*). Besides, *stG480* (27 isolates), *stG6* (23 isolates), and *stG485* (22 isolates) were the three highest frequent *emm* types and represented 51% of all isolates. In all cases, *stG6* was associated with Cellulitis (Rantala et al., 2010). Moreover, Gherardi et al. investigated the genetic relationships, the virulence profile, and cell invasiveness, as well as the susceptibility to macrolide and tetracycline antibiotics of SDSE strains isolated throughout Italy from invasive and non-invasive diseases and carriers (Gherardi et al., 2014). They found that invasiveness properties seemed to be specifically correlated with the emm subtype. For example, emm subtypes *stG6*, *stC36,* and *stG6792* differed between the group G fibronectin-binding protein gene gfbA-positive WI/NI (weak invasive/ non-invasive) and HI (high invasive) strains. Similarly, *stG6* and *stG480 emm* subtypes encountered within gfbA-negative WI/NI and HI strains were different, except for *st480.0*. Moreover, almost all of the most frequent emm types (*stG6*, *stC6979*, *stG480*, *stG62647*, *stG6792*, *stC839*, *stG485*, *stG643,* and *stC36*, in decreasing order) have also been encountered elsewhere, although with different frequencies, as reported by population genetic studies on SDSE in Australia, Europe and North America (McMillan et al., 2010), reflecting the diffusion of a few successful emm types fit to disseminate in humans. The nominal sample size of the collection does not allow any statistical association between emm type and specific clinical manifestation to be inferred; thus, they cannot confirm some strong associations reported elsewhere (i.e., *stG485*, *stG480,* and *stG6* and invasive infections) (Jensen and Kilian, 2012). For example, while our findings agree with the reported association of *stC839* with non-invasive infections (Pinho et al., 2006) (Pinho et al., 2006), all nine *stG6* strains here examined were isolated from either carriers or non-invasive infections (Gherardi et al., 2014). Lu et al. (Lu et al., 2016) characterized 56 SDSE isolates collected from two tertiary hospitals in Beijing, China. They found that emm types *stG245.0*, *stG652.0*, and *stG485.0*, were prevalent, in disagreement with those reported in Norway (*stG485*, *stG6,* and *stG643* predominated) (Kittang et al., 2011) (Kittang et al., 2011), Japan (*st6792* or subtype *stG6792.3*, *stG485*, and *stG2078* dominated) (Sunaoshi et al., 2010; Takahashi et al., 2010; Takahashi et al., 2011) (Sunaoshi et al., 2010; Takahashi et al., 2010, 2011), and the United States (*stG6*, *stG245*, *stG2078*, and *stG643* dominated) (Broyles et al., 2009) (Broyles et al., 2009), respectively. Wajima et al. (Wajima et al., 2016) gathered β-hemolytic streptococci (1,611 isolates) from invasive streptococcal infections patients in Japan during April 2010– March 2013. SDSE was most frequent (n = 693); 99% of patients with SDSE infections were elderly (mean age 75 years, SD ±15 years). They previously reported that stG485 and stG6792 were more prevalent in isolates from iSDSE infections in Japan, whereas *stG10* and *stG6* were more prevalent in noninvasive strains (Takahashi et al., 2010). The most predominant was *stG6792*, which accounted for 27.1% of isolates, followed by *stG485* (13.3%), *stG245* (10.7%), *stG652* (6.8%), *stG10* (6.2%), and *stG6* (5.5%). To explain relationships among *emm* type and clonal multifaceted CC, the main CC was identified for each *emm* type. Excluding for *stG652* strains, CCs in almost all strains, *stG653 and stG6792,* and all of those in *stG2078,* and *stG4974*, were recognized as CC17. Similarly, strains *in stG6*,*stG245, stG5420, and stG166b*,were allocated to CC25, *stC74a* to CC29, *stG10* to CC15, and *stC6979* to CC129. Numerous *emm* types, *stG485*, *stC36, stC652*, *stG480*, and *stG4222,* belonged to >2 diverse CCs. No significant association was found between *emm* type and the fatality rate for infected patients. Moreover, Wang et al. (Wang et al., 2016) In a previous study, The MLST ST44 of SDSE was associated with five *emm* types (*stG6, stC36*, *stG245*, *stG480*, and *stGL265*) (McMillan et al., 2011). MLST is a nucleotide-based method for characterizing genetic relationships amongst isolates of the same bacterial species. Unlike emm-typing, MLST utilizes multiple housekeeping genes that are considered selectively neutral and located in different parts of the genome. MLST, therefore, provides a better tool for determining evolutionary relationships within the SDSE population than typing using the ***emm*** gene, which is under strong diversifying selective pressure. Recent MLST studies of SDSE isolates from Australia, Portugal, and the USA reported a high degree of genetic diversity in these populations and revealed lateral gene transfer (LGT) of housekeeping alleles was occurring (Ahmad et al., 2009; McMillan et al., 2010). McMillan et al. (McMillan et al., 2011) used MLST to characterize relationships in India’s SDSE population, a country where streptococcal disease is endemic. The study revealed that Indian SDSE isolates have sequence types (STs) predominantly different from those reported from other world regions. MLST resolved the 181 isolates into 52 STs. The most common ST, ST84, was found in 43 isolates, emm-type stg4831. The five next most abundant STs (ST44, ST15, ST89, ST81, and ST107) each individually constitute 5 to 10% of the total collection, and together with ST84, account for 53% of isolates. McMillan et al. [25] investigated the molecular epidemiology, and evolutionary relationships of SDSE isolates collected from Australia, Europe, and North America using MLST analysis in a previous study. They segregated 178 SDSE isolates, representing 37 emm types, into 80 distinct sequence types (STs) that form 17 clonal complexes (CCs). They showed that ST44 is associated with *emm* type *stG6* found mainly in Australia and Europe and belongs to the carbohydrate group G, which all in agreement with our isolate. Our genome-genome analysis showed that SDSE strain SD-SCDR1 is closely related to SDSE strain SK1250, strain WCHSDSE-1 and strain UT-5345, with distances values of 0.00399629, 0.00467416, and 0.00687462, respectively. While whole-genome phylogenetic analysis showed that our strain SD-SCDR1 belongs to a clade that contains *Streptococcus dysgalactiae* subsp. *equisimilis* strains, namely the reference strain AC-2713 and GGS 124. In addition, within *Streptococcus dysgalactiae* subsp. *equisimilis* taxa, our strain SDSE-SCDR1 is closely related to both WCHSDSE-1 and strain SK1250. The SDSE strain WCHSDSE-1 caused an outbreak of tonsillopharyngitis among healthcare workers in China that belongs to the Lancefield group G, *emm* type *stG211.1,* and sequence type 44 (Wang et al., 2016). Whereas strain SK1250 belonged to Lancefield group A and emm type *stG840* and was isolated from the human throat in Sweden in 2001. The relative closeness between SDSE-SCDR1 and both WCHSDSE-1 and SK1250 was also confirmed by in silico genome-to-genome comparisons. Both phylogenetic and genome-to-genome analyses confirmed that the SDSE-SCDR1 strain clustered together more closely than the *Streptococcus dysgalactiae subsp. equisimilis* strain AC-2713 and GGS 124. SDSE AC-2713 isolate possesses the group A cell wall carbohydrate antigen. This isolate was recovered from a blood culture of a 78-year-old man who was suffering from an acute onset of headache, 40°C fever, and chills. His medical history included liver cirrhosis and signs of portal hypertension. He presented local erythema and tenderness of his right foot, and the diagnosis of erysipelas was rendered. The site of bacterial entry was assumed to be a minor skin lesion acquired four days before admission. Antimicrobial therapy with a typical dose of intravenous amoxicillin was administered (Brandt et al., 1999). The patient’s condition improved, and he was discharged six days later on an oral regimen to complete a 10-day course of amoxicillin. Whereas *Streptococcus dysgalactiae subsp. equisimilis GGS_124* was isolated from a patient with Streptococcal toxic shock syndrome (STSS) by Shimomura et al. (Shimomura et al., 2011). This isolate belongs to Lancefield group G, ST67, and is generally used for comparative analysis by Japan’s International Medical Center. It is associated with endocarditis, meningitis, and septicemia in both humans and swine (Streptococcus dysgalactiae subsp. equisimilis GGS_124::Genome Overview). Our SDSE- SCDR-1 isolate’s genome annotation showed that it contains many genes with homology to known virulence factors, antibiotic resistance genes, and drug targets. In general, SDSE shares most of the virulence factor genes of GAS, including streptolysin O, streptokinase, CT-like regions, NADase, and DRS (for “distantly related to SIC”)(Shimomura et al., 2011). Our isolate contains genetic determinates involved in adherence to the host cell, such as ***dltA***, ***lmb***, ***gbs0628***, ***gbs0632***, ***gbs0630***, ***gbs0631***, and ***fbp54***. Similarly, WCHSDSE-1 contains ***lmb***, ***pavA***, ***gfbA,*** and ***gapC*** for adherence (Wang et al., 2016), whereas GGS_124 possesses genes that encode putative adhesion proteins (Shimomura et al., 2011), including proteins similar to putative fibronectin-binding proteins, pullulanase, phosphopyruvate hydratase, laminin-binding protein, internalin protein, and collagen-binding protein, all of which bind to the extracellular matrix. Besides, GGS_124 possesses genes encoding the multifunctional M protein (***stg480.0***) that governs the antiphagocytic and adhesin activities, though the adhesion function of the GGS_124 M protein might be due to a collagen-binding motif (Nitsche et al., 2006; Barroso et al., 2009). Moreover, our isolate contains genetic determinates involved in cellular and tissue invasion, Antiphagocytosis, and immune evasion, such as *emm1*, *emm*, and ***hasC***. Whereas WCHSDSE-1 does not contain Antiphagocytosis or immune evasion genetic determinants (Wang et al., 2016). SDSE-SCDR1 contains ***ska***, encoding for Streptokinase, and ***SpyM3_0013***, encoding for cationic amino acid transporter-APC Superfamily, gene determinants for the invasion of skin and soft tissue, While WCHSDSE-1 contains ***skc*** and ***skg***, encoding for Streptokinase, aiding in spreading of the pathogen. Moreover, GGS_124 also carries a gene for streptokinase (SDEG_0233), similar to streptokinase A of GAS, with 88% amino acid identity. Furthermore, SDSE- SCDR1 contains the ***slo*** gene that encodes for Thiol-activated cytolysin streptolysin O that function as a toxin and pore-forming agent. Additionally, SDSE-SCDR1 contains the ***sagA*** gene that encodes for Streptolysin S precursor (SagA), the oxygen-stable, nonimmunogenic cytotoxin, which causes a zone of beta-hemolysis observed on the surface of blood agar. Similarly, WCHSDSE-1 contains both determinants of streptolysins O and S. Moreover, GGS_124 also has genes encoding streptolysin S (***sagA***) and its biosynthesis proteins (sag BCDEFGHI). Also, GGS_124 harbours a gene for streptolysin O (***slo***) (SDEG_2027), which is necessary for GAS virulence and is essential for the organism to evade from the endosome into the cytosol following the invasion of host cells (Nakagawa et al., 2004). Furthermore, SDSE- SCDR1 contains the ***speg*** gene that encodes for the exotoxin G (phage associated *Streptococcal pyrogenic* exotoxin G (SpeG)), which functions as the membrane-acting toxin (superantigen). Similarly, WCHSDSE- 1 contains exotoxin G encoded by the ***speG***, whereas GGS_124 carries the superantigen gene, exotoxin G variant 4 (*spegg4*) is homologous to GAS streptococcal exotoxin G (SpeG), with 79% amino acid identity. Indeed, superantigen G and streptolysin S genes, carried by SDSE-SCDR1, are considered the most important virulence factors triggering invasive diseases (Abdelsalam et al., 2010). It is worth mentioning that SDSE-SCDR1 possesses ***fbp54*** gene encoding fibronectin/fibrinogen-binding protein. WCHSDSE-1 contains a similar gene, ***gfbA,*** the fibronectin-binding protein-encoding gene. Whereas and GGS_124 possesses genes that encode putative adhesion proteins, including proteins similar to putative fibronectin- binding proteins (SDEG_0161, 1263, and 1984). The genetic redundancy and the exact roles of ***fbp54***, ***gfbA***, ***SDEG_0161***, and ***fnB***, which causes invasive diseases, still not clear and require additional investigation. In general, we detected the presence of 53 potential virulence factors that serve in host mortality and tissue damage, toxins, pore-forming proteins, cytotoxins, beta-hemolysis agents, stress resistance thermotolerance, oxidative stress resistance, growth in blood and lung, and many other metabolic, genetic determinants that is crucial for the pathogenesis and virulence. Most of these virulence factors are present in the invasive strain GGS_124, though these factors need to be experimentally characterized. The variety of virulence factors of SDSE-SCDR1 is similar to SDSE strains causing invasive diseases. SDSE- SCDR1 contains 53 putative virulent factors that contain all the 48 putative virulent factors observed in the invasive strain GGS_124 (data not shown). Nevertheless, invasive and non-invasive strains cannot be distinguished by the range of virulence factors. As prophages may carry virulence factors, we identified and predicted the presence of incomplete prophage region SCDR1 genome. Our prophage contains protein- coding sequences similar to CDSs from Lactococcus phage, Javan S145 prophage, Listeria phage, Bacillus virus G, Bacillus phage TsarBomba, and Streptococcus phage 315.1. As we mentioned before, the ***speg*** gene that encodes for the exotoxin G (phage associated *Streptococcal pyrogenic* exotoxin G (SpeG)) was also detected. However, no other virulence factors identified are present on the prophage. In addition to genetic factors that confer resistance to Fluoroquinolones, Aminocoumarin, Aminoglycoside, Streptomycin, Cycloserine, Daptomycin, Pulvomycin, Fosfomycin, Fusidic acid, Isoniazid, Mupirocin, Rifampicin, Vancomycin, Daptomycin, Streptomycin Sulfonamide antibiotics, Tigecycline, Triclosan, and Trimethoprim, SDSE-SCDR1 contains *lmrP* gene that encodes for the multidrug resistance protein ErmB that confer resistance against Lincosamides, Macrolides, Streptogramins and Tetracyclines (Table 3). Wang et al. stated that isolate WCHSDSE-1 harbors only one known antimicrobial resistance gene, erm(B), mediating resistance to macrolides, clindamycin and streptogramins B13 (Wang et al., 2016). However, we detected the presence of an antibiotic-resistant gene variant or mutant that leads to sulfonamide resistance in WCHSDSE-1 (data not shown). On the other hand, our analysis showed that strain GGS_124 contains several putative gene determinants that confer resistance to Fluoroquinolones, Aminoglycoside (Streptomycin), Cycloserine. Besides, it also contains the ***lmrP*** gene that encodes for the multidrug resistance protein ErmB and the ***folP*** gene variant or mutant that confer sulfonamide resistance (data not shown). Prokaryotes have a system that confers resistance against infection by exogenous DNA, such as viruses. In phage infection, bacteria integrate short fragments derived from phage DNA into their genomes in regions called clusters of the regularly interspaced short palindromic repeat (CRISPR). Consequently, bacteria utilize CRISPR-RNA transcripts and CRISPR-associated proteins (Cas) complexes to identify and interfere with viral propagation (Brouns et al., 2008; Marraffini and Sontheimer, 2008). This system has been detected in GAS (McShan et al., 2008), *S. equi subsp. zooepidemicus* SESZ (Holden et al., 2009), *S. mutans* (van der Ploeg, 2009), and *S. thermophilus* (Horvath and Barrangou, 2010) and SDSE (Shimomura et al., 2011; Wang et al., 2016). Two conserved CRISPR arrays were identified in the SDSE-SCDR-1 genome, namely, SCDR1-CRISPR2 and SCDR1-CRISPR 3. The length of SCDR1-CRISPR2 is 1286 bases; the repeat length is 36, the spacer length is 19 bases. While the length of SCDR1-CRISPR 3 is 962 bases, the repeat length is 32; the spacer length is 14 bases. Additionally, Two CAS systems were observed in the SDSE-SCDR-1 genome, namely, CAS-TypeIIA and CAS-TypeIC. Though it was not mentioned (Wang et al., 2016), WCHSDSE-1 contains one CRISPER array, 19 CRISPR repeats, and 18 CRISPR spacers. Similarly, CAS-TypeIIA and CAS-TypeIC systems were observed in the WCHSDSE-1 genome (data not shown). On the other hand, strain GGS_124 has a CRISPR/Cas system, named CRISPR1/Cas, which consists of CRISPR-associated protein, Csn2 family CRISPR-associated protein Cas2, CRISPR- associated protein Cas1 and, CRISPR-associated endonuclease Cas9 (data not shown). It was suggested that SDSE strains typically have a higher total number of spacers than GAS, suggesting that prophage infection of SDSE was relatively limited, resulting in a lower number of virulence factors positioned in the prophage regions of SDSE genomes. Similar behavior was detected in SESZ when compared with *S. equi subsp. equi* (SESE). For instance, the SESE 4047 genome contains no CRISPR but contains prophage associated genes encoding virulence factors. On the other hand, SESZ MGCS10565 and H70 contain which 26 and 18 spacers, respectively, but do not contain any prophages. Therefore, the CRISPR system in *streptococci* sharing prophages might play a significant role in the spread of virulence factors among species. Alternatively, these virulence factors might not benefit to SDSE during carriage or disease, thus, integration of these explicit phages is not under Darwinian selection (Shimomura et al., 2011).

To present a comprehensive contemporary SDSE pan-taxonomic/subspecies comparative genomics section, we included taxonomic/genomic information of the 55 publicly available SDSE strains/isolates. Our analysis included both SDSE GGS_124 and AC-2713 strains as reference/representative genomes, in addition to our isolate SDSE-SCDR1. As we mentioned earlier, strains NCTC11565 and strain NCTC5969 then excluded from the comparative investigation because of low quality and suggested genetic material contamination with *Pseudomonas aeruginosa* genetic material in case of strain NCTC11565 and *Staphylococcus aureus* strain NCTC5969. Besides, we believe that *Streptococcus dysgalactiae Str.* iSDSE230 and *Streptococcus dysgalactiae Str.* iSDSE140 available in the public database should be grouped with SDSE strains/isolates. Thus, we included 53 strains/genomes were included in our comparative investigation. The majority of isolates were of human origin; however, two isolates were collected from the genital tract of horses, one isolate was collected from a monkey’s throat, and one is from a pleural exudate of a dog. Sites of isolation included blood cultures, sterile sites of invasive diseases, oral, throat (Pharyngitis) or upper respiratory samples, foot ulcers, and Joint fluid samples.

The protein-families-based Maximum likelihood phylogenetic analysis showed that SDSE isolates are distributed amongst six significant clades. The first clade, which contains our isolate SD-SCDR1, contains nine additional SDSE strains: SK1250, WCHSDSE-1, KNZ01, UT 4242 AB, NCTC11556, NCTC6407, NCTC10321, NCTC6179, and NCTC6181 strains with a bootstrap value of 79. Whereas strain SDSE-AC 2713 is located in the second clade that contains 11 SDSE strains, namely, UT-SS957, NCTC9414, NCTC5371, AKSDE4288, NCTC9603, UT_4255RC, NCTC11557, UT_4234_DH, ASDSE 99, and NCTC11555 strains with a bootstrap value of 91. The third clade contains seven SDSE strains: UT_4231_KK, C161L1, UT-SS1069, 167, UT_4241_XS, UT_4966_RC, and strain NCTC5370 with a bootstrap value of 100. The fourth clade contains 14 SDSE strains, namely, the invasive strain GGS_124, SS1575, UT_4031CC, NCTC11564, KNZ04, KNZ15, KNZ07, RE378, KNZ01, KNZ03, KNZ12, NCTC7136, T642, and KNZ06 with a bootstrap value of 88. The fifth clade contains seven SDSE strains: ATCC 12394, KNZ16, NCTC11554, UT-5345, KNZ03-1, KNZ10 strain SK1249. The sixth clade contains two SDSE strains, namely, UT-5354 and NCTC8543. Strains UT_4277_BB, ASDSE_96, NCTC8546, and NCTC9413 each showed an independent phylogenetic lineage. Simultaneously, strains NCTC11565 and NCTC5969 confirmed the suggested low-quality genome by being phylogenetically outcast taxa. There is no doubt that advancements are required in the field of antimicrobial/antibiotic resistance (AMR) and antibiotic susceptibility testing. Both accessory genomic elements and CRISP-Cas-mediated immunity modulate resistance elements’ acquisition (van Belkum and Dunne, 2013; van Belkum et al., 2015). Henceforth, further phenotype-genotype association investigations for critical bacterial pathogens are required. Studies have suggested that genomic antibiograms are as good as phenotypic ones. The initial study involved a mixture of bacterial species resulted in 99.7% accordance between genotypes and phenotypes (Zankari et al., 2012). Additionally, investigations involved *Proteus mirabilis*, *Staphylococcus aureus*, *Escherichia coli*, *Klebsiella pneumoniae,* and *Campylobacter spp.* have expanded these findings (Stoesser et al., 2013; Gordon et al., 2014; Zhao et al., 2016; Saeb et al., 2017b). Full accordance was detected for clonal bacterial taxa such as *Mycobacterium tuberculosis* (Walker et al., 2015). Genomic approaches embrace the potential for developing imminent antibiotic susceptibility testing schemes for routine usage in clinical microbiology laboratories, even though principal knowledge breaches still need to be overcome (Jaillard et al., 2017).

### *Streptococcus dysgalactiae subsp. equisimilis* pan-taxonomic Antimicrobial Resistance Genes (Resistome)

We observed the presence of a common core Resistome amongst SDSE genomes/isolates and strains. It is consisting of ***folA***, ***Dfr***, ***Ddl***, ***LiaS***, ***LiaR***, ***pgsA***, ***Alr***, ***kasA***, ***gidB***, ***EF-G***, ***EF-Tu***, ***folP***, ***gyrA***, ***gyrB***, Iso-tRNA (***ileS***), ***LiaF***, ***S12p***, ***S10p***, ***rpoB***, ***rpoC***, ***MurA***, and ***FabK*** AMR genetic determinants (**Supplementary Table** 3). These AMR genetic determinants potentially confer resistance against Trimethoprim (***folA***, ***Dfr)***, Cycloserine (***Ddl, Alr)***, Daptomycin (***LiaS***, ***LiaR***, ***pgsA, LiaF),*** Isoniazid (***kasA),*** Aminoglycoside (Streptomycin) (***gidB***), Fusidic acid ***(EF-G***), Elfamycin (Pulvomycin) (***EF-Tu***), Sulfonamide antibiotics (***folP***), Fluoroquinolones ***(gyrA, gyrB),*** Mupirocin (Pseudomonic acid) ***(ileS),*** Streptomycin ***(S12p)***, Tigecycline (***S10p)***, Rifampicin, Daptomycin ***(rpoB),*** Rifampicin, Vancomycin, Daptomycin ***(rpoC***), Fosfomycin ***(MurA)*** and, (***FabK***) Triclosan and **Supplementary table 3** and **Table 3**. Besides, all tested SDSE genomes contain ***lmrP*** except for two strains, namely, T642 and C161L1. All the strains hold ***aadK*** except for five strains, namely, NCTC6179, NCTC6181, NCTC9413, NCTC6407, NCTC10321. ***rplF*** gene was missing from 22 SDSE strains, namely, NCTC8543, NCTC11564, SD- SCDR1, NCTC6181, NCTC8546, NCTC9413, NCTC11554, NCTC 11556, NCTC11565, NCTC6407, NCTC10321, NCTC5969, NCTC9603, NCTC11555, NCTC11557, KNZ10 KNZ04, KNZ07, KNZ06, KNZ12, KNZ15, and KNZ16. Some SDSE strains showed some unique AMR phenotypes and genotypes. It was found that only SDSE strains AKSDE4288, strain ASDSE99, strain NCTC11557, and strain UT_4255RC have ***ermA***. Moreover, only strains SDSE KNZ04 and WCHSDSE-1 have ***ermB***. Besides, only two *Streptococcus dysgalactiae subsp. equisimilis* strain Sde558 has ***erm(T)***. SDSE strain AC-2713 has ABC-F type ribosomal protection protein **Msr(D)** that confer resistance to the antibiotic telithromycin and MFS efflux pump, **Mef(A)**, which is responsible for resistance against Macrolides, Lincosamides, Streptogramins, Ketolides, Oxazolidinones (MLSKO). Furthermore, SDSE strains AKSDE4288, and SK1250 have the tetracycline resistance gene, ***tetO***, that encodes for the ribosomal protection type (O) Tetracycline resistance. Whereas, SDSE strains NCTC11557, UT_4242_AB, UT_4234_DH, KNZ04, KNZ06, ASDSE_99, and ASDSE_96 have Tetracycline resistance gene ***tetM***, that encodes for the ribosomal protection type (M) Tetracycline resistance. In addition, SDSE strain KNZ06 has ***parC*** gene that confers resistance against fluoroquinolone antibiotics. The SDSE strain T642 a point mutation in ***ParE*,** a subunit of topoisomerase IV that decatenates and relaxes DNA to allow access to genes for transcription or translation that prevent Anticoumarin antibiotics from inhibiting DNA synthesis, and thus conferring resistance. Besides, it has the PhoB, phosphate regulon transcriptional regulatory protein (***SphR***), a direct regulator of the macAB efflux pump antibiotic out of a cell to confer resistance. Similarly, SDSE strain C161L1 has **PhoB** resistance against peptide antibiotics. Moreover, strain C161L1 has the ***gyrB*** (parE) gene that confers resistance against aminocoumarin (novobiocin). Indeed, elevated rates of macrolide resistance encoded by ***erm*** (erythromycin ribosome methylase) and ***mef*** (macrolide efflux) genes have been described in the last three decades amongst the main streptococcal pathogens, including SDSE (Brandt and Spellerberg, 2009; Broyles et al., 2009; Gherardi et al., 2014). It is worth mentioning that our isolate SDSE- SCDR1 is resistant to erythromycin, clindamycin, tetracycline, and tigecycline; however, it showed sensitivity against cefotaxime and vancomycin. Similarly, SDSE- WCHSDSE-1 was susceptible to penicillin, cefotaxime, levofloxacin, and vancomycin but was resistant to erythromycin, azithromycin, clindamycin, and tetracycline (Wang et al., 2016). As we mentioned above, the genomes of SDSE consists of a core virulence structure (virulome) consisting of 38 genetic determinants. These genetic determinants are involved in the infectivity process, ***idnO, asnA, bcaT, pyrG*, *metQ***, ***pflD,* SP_0095, *ykqC, fba, pstA, deoB, pstB,*** and ***leuS***, Streptococcal pathogenesis and full virulence, ***oppA, vicK, gidA, lgt*,** and ***perR***, invasive skin and soft tissue infections and systemic dissemination of GAS from the skin, **BCHE**, adhesion, tissue invasion, and bacterial virulence ***fbp54, dlt, hasC, lepA, guaA, cydA, SpyM3-0013, luxS***, and colonization ability, ***purB, ccpA, glnA*** (Holmes et al., 2001; Hava and Camilli, 2002; Wang et al., 2005; Zhu et al., 2009). Besides, we observed the presence of ***covS*** associated with hypervirulence (Hollands et al., 2010) amongst all the genomes of SDSE. Furthermore, the stress resistance genes ***clpP, cpsY*** and ***SPy_1633*** were also part of the SDSE core virulome (Shelver et al., 2003; Ibrahim et al., 2005; Kang et al., 2010). Moreover, ***lsp***, the evasive immune mechanism, phagosomal escape, acquisition of zinc and virulence genetics determinant, and ***purH***, involved in the synthesis of purines, required for pathogenesis and virulence, were also observed (Flashner et al., 2004; Moldovan and Fraunholz, 2019) **(Supplementary Table 4 and Table 2).** All tested SDSE genomes contain the eight potential drug targets that can provide a solution for severe untreatable infections (**Supplementary table 5 and table 4)**. Amongst these two are approved drugs and six with experimental drugs. The ***gyrB*** gene encoded for DNA topoisomerase IV subunit B (EC 5.99.1.3) can be targeted by Gatifloxacin, an approved drug, an antibiotic, and a member of the fourth-generation fluoroquinolone family. It functions through inhibiting the bacterial enzymes DNA gyrase and topoisomerase IV. It is indicated for the treatment of bronchitis, sinusitis, community-acquired pneumonia, and skin infections (abscesses, wounds) caused by *S. pneumoniae*, *H. influenzae*, *S. aureus*, *M. pneumoniae*, *C. pneumoniae*, *L. pneumophila*, and *S. pyogenes* (Park-Wyllie et al., 2006). Also, ***rplC*** encode for LSU ribosomal protein L3p (L3e) can be targeted by Retapamulin and Spiramycin, both approved drugs. Retapamulin is an antibiotic for skin infections like impetigo. The FDA approved it in April 2007. Retapamulin acts by selectively inhibiting protein synthesis initiation in bacteria at the bacterial 50S ribosome level against *S. aureus* (methicillin-susceptible isolates only) or *S. pyogenes* (Jones et al., 2006). Spiramycin is a chiefly bacteriostatic macrolide antimicrobial agent with activity against Gram- positive cocci and rods, Gram-negative cocci, and Legionellae *mycoplasmas*, *chlamydiae*, some types of spirochetes, *Toxoplasma gondii,* and *Cryptosporidium*. Spiramycin is a 16-membered macrolide that inhibits translocation by attaching to bacterial 50S ribosomal subunits with an evident 1:1 stoichiometry. This antibiotic is a powerful inhibitor of the binding to the ribosome of both donor and acceptor substrates (Brisson-Noël et al., 1988). Amongst the drug targets with experimental drugs is the ***def*** gene that encodes for peptide deformylase (EC 3.5.1.88) that is targeted by 2-[(Formyl-Hydroxy-Amino)-Methyl]-Heptanoic Acid [1-(2-Hydroxymethyl-Pyrrolidine-1-Carbonyl)-2-Methyl-Propyl]-Amide. This experimental drug removes the formyl group from the N-terminal Met of newly synthesized proteins (Meinnel et al., 1997). Moreover, the ***guaB*** gene encodes for Inosine-5’-monophosphate dehydrogenase/CBS domain is targeted by Merimepodib, an experimental drug, the active inhibitor of inosine monophosphate dehydrogenase (IMPDH). IMPDH inhibition leads to a decline in intracellular guanosine triphosphate (GT) required for DNA and RNA synthesis. It is also affected by Inosinic acid, an experimental drug that catalyzes the transformation of inosine 5’-phosphate (IMP) to xanthosine 5’-phosphate (XMP), essential for the de-novo synthesis of guanine nucleotides. As a result, it plays a vital role in regulating cell growth (Ashbaugh and Wessels, 1995; Markland et al., 2000). **Supplementary Table 6** and **supplementary figure 1** presents detailed information about the complete phage genomes annotated in SDSE genomes. To date, 16 complete phages are associated with SDSE strains. Streptococcal phages are considered critical in horizontal gene transfer, especially in the transport of virulence factors in Streptococci (Ikebe et al., 2002; Holden et al., 2009). We observed the presence of **Streptodornase D** encoding gene ***sda*** in both strains SDSE UT_4242_AB and SDSE UT_4966_RC. This gene is homologous to phage-encoded extracellular streptodornase D (***sda1***) of GAS (PHA01790), responsible of the degradation of neutrophil extracellular traps (NETs), which are composed of granule proteins and chromatin released by neutrophils and can catch and kill surrounding bacteria (Buchanan et al., 2006). Besides, we observed the presence of the pleiotropic regulator of exopolysaccharide synthesis, competence, and biofilm formation Ftr, XRE family homologous with ftr protein of GAS strain SPY5003 in Streptococcus phage Javan161 and Streptococcus phage Javan170. Also, we observed the presence of the Streptococcal extracellular nuclease 3 (Mitogenic factor 3, MF3) encoded by ***SPy1436,*** of GAS serotype M1, in both Streptococcus phage Javan161 and Streptococcus phage Javan170. Mitogenic factor 3 plays a crucial role in the evasion of the innate immune response by degrading neutrophil extracellular traps. Besides, MF3 acts as an extracellular DNase in most clinically isolated GAS strains. Moreover, all GAS strains have extracellular nuclease activity and produce significantly more activity compared with other groups of streptococci, signifying that the activity might contribute to virulence (Wen et al., 2011). Mitogenic factor 3 is also found in Prophage GGS_124.2 of the invasive strain GGS_124 (Shimomura et al., 2011). Also, we observed the presence of pathogenicity island SaPIn2 in Streptococcus phage phi3396. The pathogenicity island SaPIn2 appears to be important to the pathogen, i.e. *S. aureus*, as it is maintained in the genome by two restriction-modification genes in the center of the island. Remnants of gene duplication and transfer may also be found in the transposase gene at the 5 end of SaPIn2. This finding further supports the importance and prevalence of these proteins as agents of host/pathogen interactions (Arcus et al., 2002).

CRISPR relies on small RNAs for sequence-specific detection and cleaving of foreign nucleic acids, including bacteriophages and plasmids. Watanabe et al. reported that All SDSE strains sequenced previously harbor at least one probable CRISPR with more than 20 spacers. In contrast, their newly sequenced 167 strain does not harbor any probable CRISPR or CAS. They suggested that presumably, there is no interference by the CRISPR system in 167, and SDSE 167 may be prone to infection by phages. Moreover, analysis of a larger number of group C SDSE isolates is necessary to determine whether the absence of CRISPR interference is common to these isolates (Watanabe et al., 2013). Thus, in the 55 genomes of SDSE, we discovered that most SDSE strains/genomes (89%) have CRISPER-CAS systems associated with genes/products **(Supplementary Table 7)**.

Moreover, GGS_124, the STSS causing strain, contains a CRISPR/Cas system consisting of an array of genes, can1, cas1, cas2, csn2, and CRISPR (19 direct repeats of 36 bp each and 18 spacer sequences 30 or 32 bp in length). Besides, it was found that the mean number of CRISPR spacers was higher in seven SDSE (18.0 ± 3.3 spacers) than in GAS strains (4.0 ± 1.0 spacers; range, 0 to 9), signifying that prophage infection of SDSE is to some extent constrained, resulting in a lesser number of virulence factors positioned in the prophage regions of SDSE. Then Again, SDSE may contact phages more frequently, with integrated phages having a fitness cost for SDSE (Shimomura et al., 2011). Therefore, the CRISPR system in streptococci sharing prophages may play a significant role in spreading virulence factors among species. On the other hand, these virulence factors might not be beneficial for SDSE throughout the carriage or disease, such that the integration of these individual phages is not under positive selection. We noticed two CAS systems among the investigated SDSE genomes, namely, CAS-TypeIIA and CAS-TypeIC. The signature gene for type II CRISPR-Cas systems is ***cas9***, which encodes a multi-domain protein that combines all the functions of effector complexes and the target DNA cleavage and is essential for the maturation of the CRISPR RNA (crRNA) (Jinek et al., 2012). Type II-A systems encompass an additional gene, known as csn2, which is considered a signature gene for this subtype. The Csn2 protein is not required for interference but has an unclear role in spacer integration (Barrangou et al., 2007). All type I systems contain the signature gene ***cas3,*** which encodes a large protein with a helicase possessing a single-stranded DNA (ssDNA)-stimulated ATPase activity coupled to the unwinding of DNA-DNA and RNA-DNA duplexes [40]. The signature gene for this subtype C is Cas8c and usually lacks the ***cas6*** gene. The Cas5 is catalytically active and replaces the Cas6 function (Makarova and Koonin, 2015). In conclusion, in the present work, we present a comprehensive genomic analysis for the SDSE strain SCDR1 that belong to Lancefield group G, emm type (stG6) and (MLST) sequence type (ST44) isolated from neuroischaemic diabetic foot ulcer, the first time for this emerging pathogen to be documented in Saudi Arabia and middle east. Strain SCDR1 has a superior spectrum of virulence factors with SDSE strains causing invasive diseases, suggesting that strains causing invasive and non-invasive diseases may not be distinguished by the presence or absence of certain virulence factors.

Moreover, we presented the most comprehensive comparative genomics analysis for the emerging human pathogen SDSE. Our study confirmed that the SDSE strains possess different characteristics in producing virulence factors for pathogenicity to humans and their genome sequence and close relationship with GAS. Our study shed light on the proposed pathogenic mechanisms of SDSE and may form the basis of molecular epidemiological research on these highly virulent bacteria.

## Figures titles

**Figure 1**: Genome representation shows from outer to inner rings, the contigs, CDS on the forward strand, CDS on the reverse strand, RNA genes, CDS with homology to known antimicrobial resistance genes, CDS with homology to known virulence factors, GC content, and GC skew.

**Figure 2**: Mauve Multiple Whole Genome Alignment of SDSE strain SD-SCDR1 against reference genomes SDSE strain NCTC7136, SDSE strain WCHSDSE-1 and Streptococcus pyogenes strain MGAS15252.

**Figure 3**: An overview of the biological subsystems and genes for Streptococcus dysgalactiae subsp. equisimilis strain SD-SCDR1 genome.

**Figure 4**: (a) Phylogenetic tree of SDSE based on whole-genome sequences (reference and outgroup genomes) and (b) selected 33 SDSE genome sequences, Streptococcus dysgalactiae subsp. equisimilis strain SD-SCDR1, Streptococcus pyogenes MGAS15252, and Staphylococcus aureus MR1 genomes.

**Figure 5**: incomplete prophage region identified in Streptococcus dysgalactiae subsp. equisimilis strain SD-SCDR1 genome

**Figure 6**: protein families based on Maximum likelihood phylogenetic analysis for the publicly available SDSE strains/isolates genomes.

**Figure 7**: protein families phylogeny based proteomic comparison among SDSE-SD-SCDR1 clade containing strains (a) All gene/proteins (b) Essential gene/proteins

## Tables titles

**Table1**: The genome-genome comparison between SDSE strain SCDR-SD1 and other SDSE strains.

**Table 2**: Virulence factors genetic determinants observed in SD-SCDR-1 genome.

**Table 3**: antimicrobial resistance genes (AMR) genes annotated in SD-SCDR1 genome and corresponding AMR mechanism

**Table 4**: SDSE SD-SCDR1 genome potential novel drug targets.

**Table 5**: SDSE SD-SCDR1 genome CRISPR repeats, spaces, and arrays.

**Table 6**: CRISPR repeats information identified in SD-SCDR-1 genome.

**Table 7**: CAS systems information identified in SD-SCDR-1 genome.

**Table 8**: incomplete prophage region/CDS identified in Streptococcus dysgalactiae subsp. equisimilis strain SD-SCDR1 genome

**Table 9**: 40 Insertion sequences in SDSE-SCDR1 genome

## Supplemental Figures titles

**Supplementary Figure 1**: complete phage identified in *Streptococcus dysgalactiae* subsp. *equisimilis* strain/genomes.

**Supplementary Figure 2**: *Streptococcus dysgalactiae* subsp. *equisimilis* pan-taxonomic CRISPER- CAS-TypeIC systems.

**Supplementary Figure 3**: *Streptococcus dysgalactiae* subsp. *equisimilis* pan-taxonomic CRISPER-CAS- TypeIIA systems.

## Supplementary Tables titles

**Supplementary Table 1**: represents the taxonomic/genomic information of the 55 publicly available SDSE strains/isolates.

**Supplementary Table 2**: Reference strain SDSE-AC 2713 essential 189 genes predicted using LB media’s metabolic flux-balance analysis.

**Supplementary Table 3**: Antimicrobial resistance genes (AMR) genes annotated in SDSE genomes and corresponding AMR mechanism

**Supplementary Table 4**: Major virulence factors (VF) genes annotated in SDSE genomes and corresponding products.

**Supplementary Table 5**: Drug target genes/products annotated in SDSE genomes and proposed drug.

**Supplementary Table 6**: Detailed information for the complete phage genomes annotated in SDSE genomes.

**Supplementary Table 7**: Detailed information on CRISPER-CAS systems detected in SDSE genomes.

## * List of abbreviations

SDSE: Streptococcus dysgalactiae subsp. equisimilis
GAS: Streptococcus pyogenes
MLST: Multilocus sequence typing
PATRIC: Bacterial bioinformatics database and analysis resource
ARDB: Antibiotic Resistance Genes Database
CARD: The Comprehensive Antibiotic Resistance Database
DFU: Diabetic foot ulcer
ABPI: The ankle-brachial pressure index
CRISPR: Clustered regularly interspaced short palindromic repeats
AMR: The antimicrobial resistance genes
MLSKO: Macrolides, Lincosamides, Streptogramins, Ketolides, and Oxazolidinones
VF: Virulence factors
GRAB: G-related-alpha-2 macroglobulin-binding protein
WI/NI: Weak invasive/ Non-invasive
HI: High invasive
STSS: Streptococcal toxic shock syndrome erm: erythromycin ribosome
methylase mef: macrolide efflux

## Declarations

### **\*** Ethics approval and consent to participate

The institutional review board approved this study at King Saud University, College of Medicine Riyadh, Kingdom of Saudi Arabia (IRB-E-16-1839). The subjects provided written informed consent for participating in this study.

### **\*** Consent to publish

All authors have consented to the publication of this manuscript.

### * Availability of data and materials

The genome sequence was submitted to NCBI with an accession number CP033391 under BioProject accession number PRJNA498183, BioSample accession number SAMN10285107 and assembly accession number GCA_009497975.1.

### **\*** Authors’ contributions

**ATMS:** Involved in study conception and design, data analysis, and interpretation and involved in drafting the manuscript or revising it critically for important intellectual content. Preparing the final approval of the version to be published.

**HT:** Involved in study conception and design and drafted the manuscript or revised it critically for important intellectual content. She was involved in preparing the final approval of the version to be published.

## Acknowledgment

The authors want to thank the University Diabetes Center members at King Saud University for their help throughout the study. Besides, we want to acknowledge that NGS experiments and analysis were supported by the Saudi Human Genome Program (SHGP) at KACST and KFSHRC.

---

**Supplementary figure 1:**
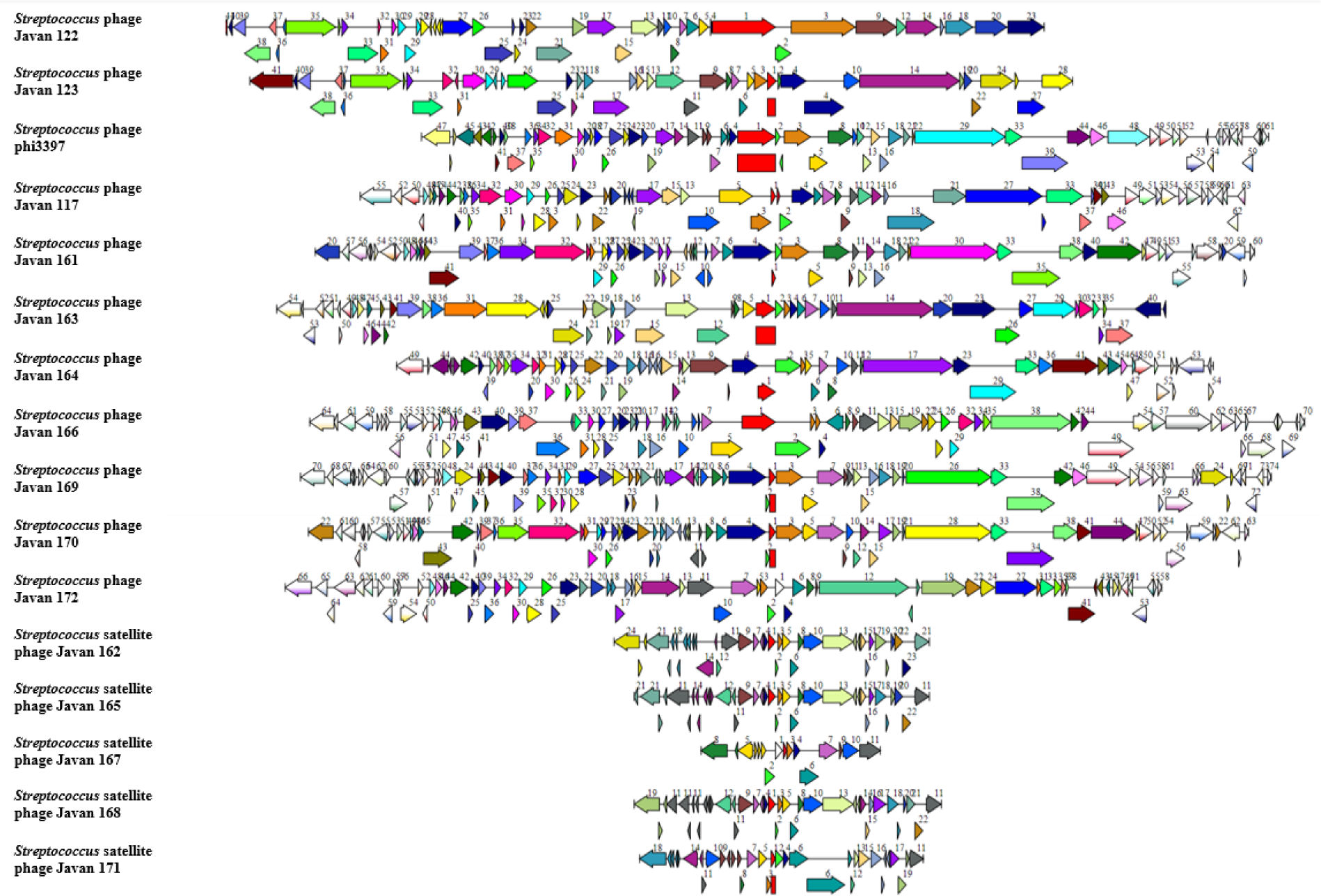
Complete phage identified in Streptococcus dysgalactiae subsp. equisimilis strain/genomes.

**Supplementary figure 2:**
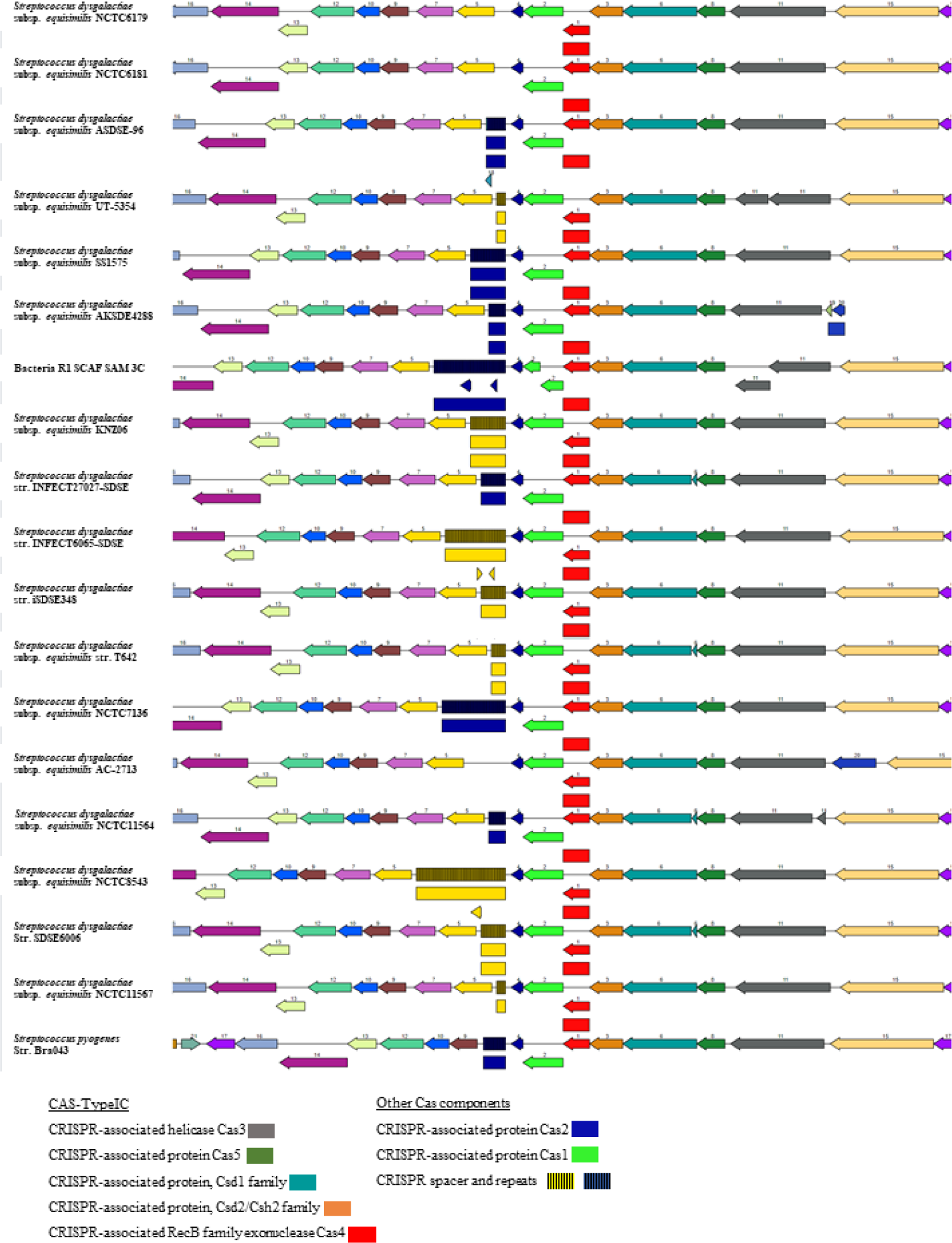
Streptococcus dysgalactiae subsp. equisimilis pan-taxonomic CRISPER- CAS-TypeIC systems.

**Supplementary figure 3:**
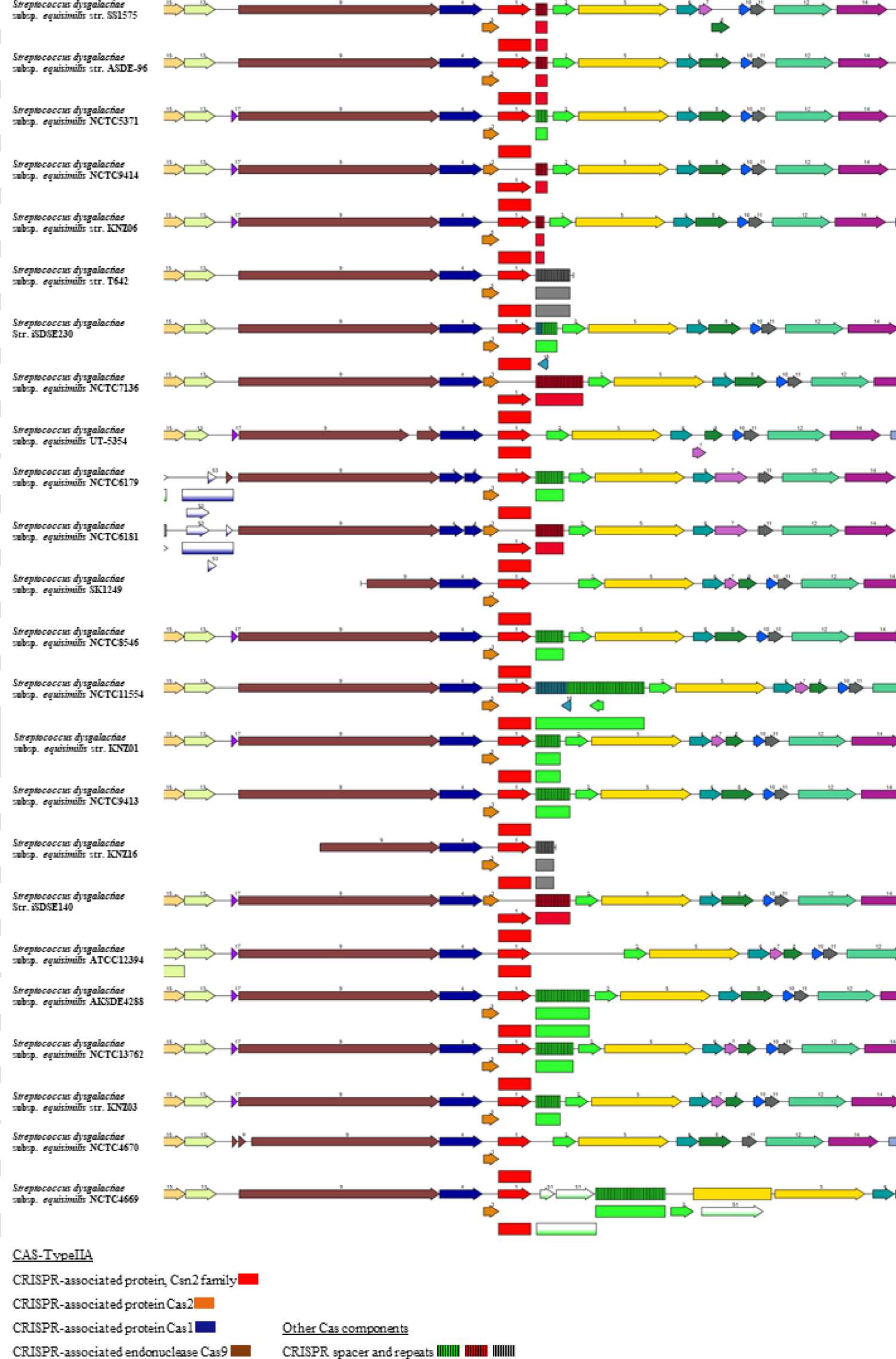
Streptococcus dysgalactiae subsp. equisimilis pan-taxonomic CRISPER-CAS-TypeIIA systems.

