## Supplementary figures and images for "Comprehensive comparative genomics analysis for the emerging human pathogen *Streptococcus dysgalactiae* subsp. *equisimilis* (SDSE): A case study and Pan-subspecies genomic analysis"

### Supplemental Fig 1

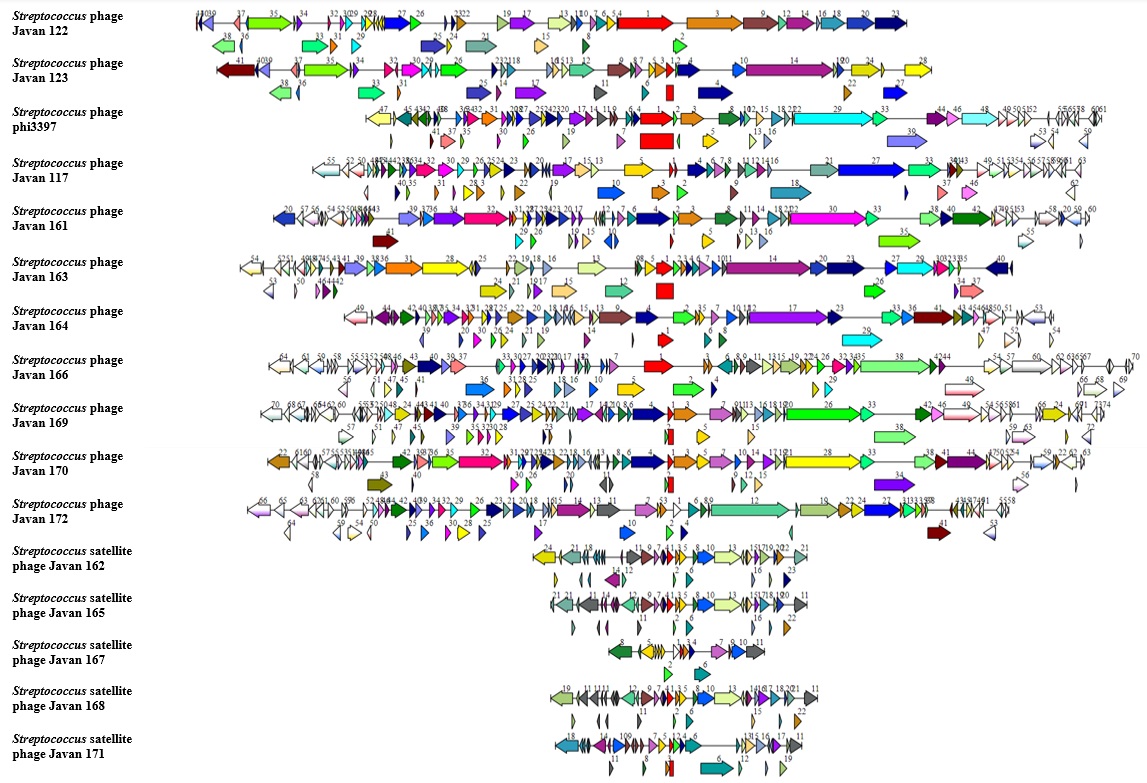

### Supplemental Fig 2

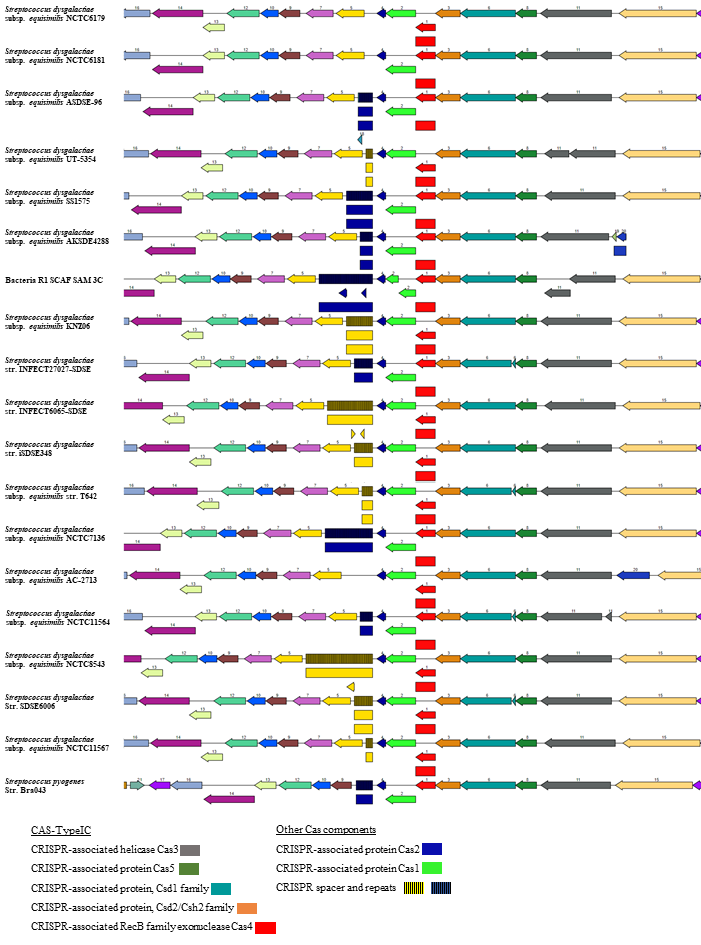

### Supplemental Fig 3

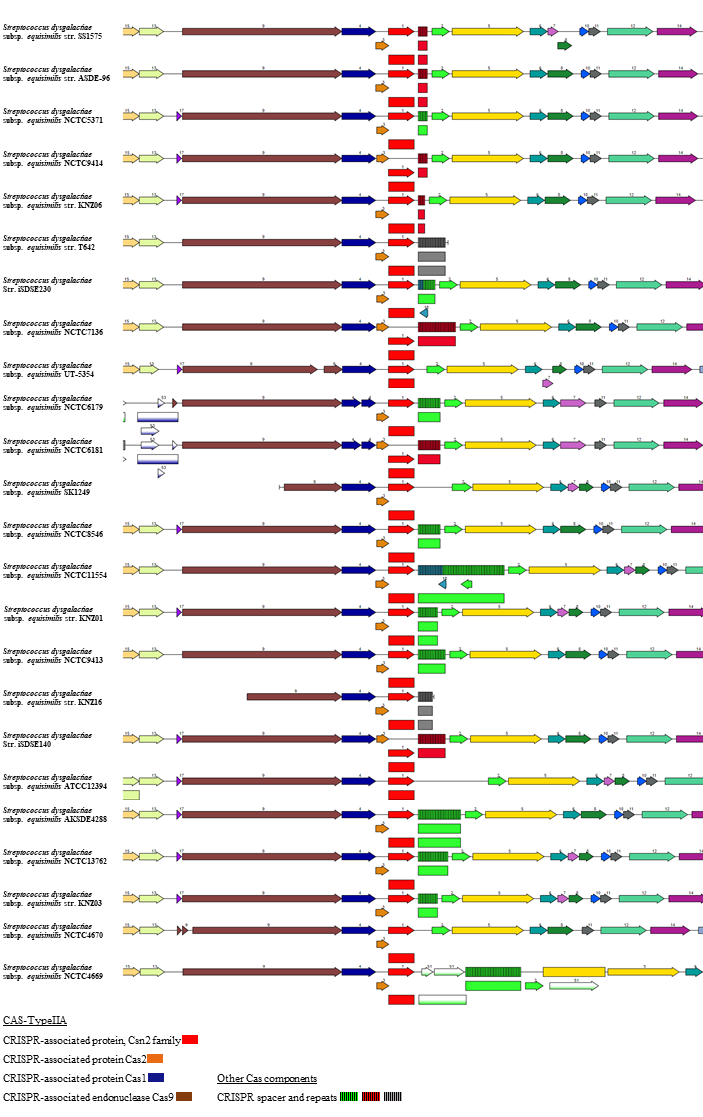
