## Supplemental File 3 for "Comprehensive comparative genomics analysis for the emerging human pathogen *Streptococcus dysgalactiae* subsp. *equisimilis* (SDSE): A case study and Pan-subspecies genomic analysis"

Supplementary Table 3: Antimicrobial resistance genes (AMR) genes annotated in SDSE genomes and corresponding AMR mechanism

| Genome Name | Gene | | Product | Classification and Function | | PubMed ID |
| --- | --- | --- | --- | --- | --- | --- |
| Streptococcus dysgalactiae subsp. equisimilis SD SCDR1 | folA, Dfr | Dihydrofolate reductase (EC 1.5.1.3) | | | antibiotic target in susceptible species | 20169085;25288078 |
| Streptococcus dysgalactiae subsp. equisimilis 167 | dyr | Dihydrofolate reductase (EC 1.5.1.3) | | | antibiotic target in susceptible species | 20169085;25288078 |
| Streptococcus dysgalactiae subsp. equisimilis AC-2713 | dyr | Dihydrofolate reductase (EC 1.5.1.3) | | | antibiotic target in susceptible species | 20169085;25288078 |
| Streptococcus dysgalactiae subsp. equisimilis AKSDE4288 | folA, Dfr | Dihydrofolate reductase (EC 1.5.1.3) | | | antibiotic target in susceptible species | 20169085;25288078 |
| Streptococcus dysgalactiae subsp. equisimilis ATCC 12394 | folA, Dfr | Dihydrofolate reductase (EC 1.5.1.3) | | | antibiotic target in susceptible species | 20169085;25288078 |
| Streptococcus dysgalactiae subsp. equisimilis GGS_124 | dyr | Dihydrofolate reductase (EC 1.5.1.3) | | | antibiotic target in susceptible species | 20169085;25288078 |
| Streptococcus dysgalactiae subsp. equisimilis RE378 | dyr | Dihydrofolate reductase (EC 1.5.1.3) | | | antibiotic target in susceptible species | 20169085;25288078 |
| Streptococcus dysgalactiae subsp. equisimilis SK1249 | folA | Dihydrofolate reductase (EC 1.5.1.3) | | | antibiotic target in susceptible species | 20169085;25288078 |
| Streptococcus dysgalactiae subsp. equisimilis SK1250 | folA | Dihydrofolate reductase (EC 1.5.1.3) | | | antibiotic target in susceptible species | 20169085;25288078 |
| Streptococcus dysgalactiae subsp. equisimilis strain ASDSE_96 | folA, Dfr | Dihydrofolate reductase (EC 1.5.1.3) | | | antibiotic target in susceptible species | 20169085;25288078 |
| Streptococcus dysgalactiae subsp. equisimilis strain ASDSE_99 | folA, Dfr | Dihydrofolate reductase (EC 1.5.1.3) | | | antibiotic target in susceptible species | 20169085;25288078 |
| Streptococcus dysgalactiae subsp. equisimilis strain C161L1 | folA, Dfr | Dihydrofolate reductase (EC 1.5.1.3) | | | antibiotic target in susceptible species | 20169085;25288078 |
| Streptococcus dysgalactiae subsp. equisimilis strain KNZ01 | folA, Dfr | Dihydrofolate reductase (EC 1.5.1.3) | | | antibiotic target in susceptible species | 20169085;25288078 |
| Streptococcus dysgalactiae subsp. equisimilis strain KNZ01 | folA, Dfr | Dihydrofolate reductase (EC 1.5.1.3) | | | antibiotic target in susceptible species | 20169085;25288078 |
| Streptococcus dysgalactiae subsp. equisimilis strain KNZ03 | folA, Dfr | Dihydrofolate reductase (EC 1.5.1.3) | | | antibiotic target in susceptible species | 20169085;25288078 |
| Streptococcus dysgalactiae subsp. equisimilis strain KNZ03 | folA, Dfr | Dihydrofolate reductase (EC 1.5.1.3) | | | antibiotic target in susceptible species | 20169085;25288078 |
| Streptococcus dysgalactiae subsp. equisimilis strain KNZ04 | folA, Dfr | Dihydrofolate reductase (EC 1.5.1.3) | | | antibiotic target in susceptible species | 20169085;25288078 |
| Streptococcus dysgalactiae subsp. equisimilis strain KNZ06 | folA, Dfr | Dihydrofolate reductase (EC 1.5.1.3) | | | antibiotic target in susceptible species | 20169085;25288078 |
| Streptococcus dysgalactiae subsp. equisimilis strain KNZ07 | folA, Dfr | Dihydrofolate reductase (EC 1.5.1.3) | | | antibiotic target in susceptible species | 20169085;25288078 |
| Streptococcus dysgalactiae subsp. equisimilis strain KNZ10 | folA, Dfr | Dihydrofolate reductase (EC 1.5.1.3) | | | antibiotic target in susceptible species | 20169085;25288078 |
| Streptococcus dysgalactiae subsp. equisimilis strain KNZ12 | folA, Dfr | Dihydrofolate reductase (EC 1.5.1.3) | | | antibiotic target in susceptible species | 20169085;25288078 |
| Streptococcus dysgalactiae subsp. equisimilis strain KNZ15 | folA, Dfr | Dihydrofolate reductase (EC 1.5.1.3) | | | antibiotic target in susceptible species | 20169085;25288078 |
| Streptococcus dysgalactiae subsp. equisimilis strain KNZ16 | folA, Dfr | Dihydrofolate reductase (EC 1.5.1.3) | | | antibiotic target in susceptible species | 20169085;25288078 |
| Streptococcus dysgalactiae subsp. equisimilis strain NCTC10321 | folA, Dfr | Dihydrofolate reductase (EC 1.5.1.3) | | | antibiotic target in susceptible species | 20169085;25288078 |
| Streptococcus dysgalactiae subsp. equisimilis strain NCTC11554 | folA, Dfr | Dihydrofolate reductase (EC 1.5.1.3) | | | antibiotic target in susceptible species | 20169085;25288078 |
| Streptococcus dysgalactiae subsp. equisimilis strain NCTC11555 | folA, Dfr | Dihydrofolate reductase (EC 1.5.1.3) | | | antibiotic target in susceptible species | 20169085;25288078 |
| Streptococcus dysgalactiae subsp. equisimilis strain NCTC11556 | folA, Dfr | Dihydrofolate reductase (EC 1.5.1.3) | | | antibiotic target in susceptible species | 20169085;25288078 |
| Streptococcus dysgalactiae subsp. equisimilis strain NCTC11557 | folA, Dfr | Dihydrofolate reductase (EC 1.5.1.3) | | | antibiotic target in susceptible species | 20169085;25288078 |
| Streptococcus dysgalactiae subsp. equisimilis strain NCTC11564 | folA, Dfr | Dihydrofolate reductase (EC 1.5.1.3) | | | antibiotic target in susceptible species | 20169085;25288078 |
| Streptococcus dysgalactiae subsp. equisimilis strain NCTC11565 | folA, Dfr | Dihydrofolate reductase (EC 1.5.1.3) | | | antibiotic target in susceptible species | 20169085;25288078 |
| Streptococcus dysgalactiae subsp. equisimilis strain NCTC5370 | folA, Dfr | Dihydrofolate reductase (EC 1.5.1.3) | | | antibiotic target in susceptible species | 20169085;25288078 |
| Streptococcus dysgalactiae subsp. equisimilis strain NCTC5370 | dyr | Dihydrofolate reductase (EC 1.5.1.3) | | | antibiotic target in susceptible species | 20169085;25288078 |
| Streptococcus dysgalactiae subsp. equisimilis strain NCTC5371 | dyr | Dihydrofolate reductase (EC 1.5.1.3) | | | antibiotic target in susceptible species | 20169085;25288078 |
| Streptococcus dysgalactiae subsp. equisimilis strain NCTC5969 | dfrC | Dihydrofolate reductase (EC 1.5.1.3) | | | antibiotic target replacement protein,trimethoprim resistance gene | 20169085;25288078 |
| Streptococcus dysgalactiae subsp. equisimilis strain NCTC5969 | folA, Dfr | Dihydrofolate reductase (EC 1.5.1.3) | | | antibiotic target in susceptible species | 20169085;25288078 |
| Streptococcus dysgalactiae subsp. equisimilis strain NCTC5969 | folA | Dihydrofolate reductase (EC 1.5.1.3) | | | antibiotic target in susceptible species | 20169085;25288078 |
| Streptococcus dysgalactiae subsp. equisimilis strain NCTC6179 | dyr | Dihydrofolate reductase (EC 1.5.1.3) | | | antibiotic target in susceptible species | 20169085;25288078 |
| Streptococcus dysgalactiae subsp. equisimilis strain NCTC6181 | folA, Dfr | Dihydrofolate reductase (EC 1.5.1.3) | | | antibiotic target in susceptible species | 20169085;25288078 |
| Streptococcus dysgalactiae subsp. equisimilis strain NCTC6407 | folA, Dfr | Dihydrofolate reductase (EC 1.5.1.3) | | | antibiotic target in susceptible species | 20169085;25288078 |
| Streptococcus dysgalactiae subsp. equisimilis strain NCTC7136 | dyr | Dihydrofolate reductase (EC 1.5.1.3) | | | antibiotic target in susceptible species | 20169085;25288078 |
| Streptococcus dysgalactiae subsp. equisimilis strain NCTC8543 | folA, Dfr | Dihydrofolate reductase (EC 1.5.1.3) | | | antibiotic target in susceptible species | 20169085;25288078 |
| Streptococcus dysgalactiae subsp. equisimilis strain NCTC8546 | folA, Dfr | Dihydrofolate reductase (EC 1.5.1.3) | | | antibiotic target in susceptible species | 20169085;25288078 |
| Streptococcus dysgalactiae subsp. equisimilis strain NCTC9413 | folA, Dfr | Dihydrofolate reductase (EC 1.5.1.3) | | | antibiotic target in susceptible species | 20169085;25288078 |
| Streptococcus dysgalactiae subsp. equisimilis strain NCTC9414 | dyr | Dihydrofolate reductase (EC 1.5.1.3) | | | antibiotic target in susceptible species | 20169085;25288078 |
| Streptococcus dysgalactiae subsp. equisimilis strain NCTC9603 | folA, Dfr | Dihydrofolate reductase (EC 1.5.1.3) | | | antibiotic target in susceptible species | 20169085;25288078 |
| Streptococcus dysgalactiae subsp. equisimilis strain SS1575 | folA, Dfr | Dihydrofolate reductase (EC 1.5.1.3) | | | antibiotic target in susceptible species | 20169085;25288078 |
| Streptococcus dysgalactiae subsp. equisimilis strain T642 | folA, Dfr | Dihydrofolate reductase (EC 1.5.1.3) | | | antibiotic target in susceptible species | 20169085;25288078 |
| Streptococcus dysgalactiae subsp. equisimilis strain UT_4031CC | folA, Dfr | Dihydrofolate reductase (EC 1.5.1.3) | | | antibiotic target in susceptible species | 20169085;25288078 |
| Streptococcus dysgalactiae subsp. equisimilis strain UT_4231_KK | folA, Dfr | Dihydrofolate reductase (EC 1.5.1.3) | | | antibiotic target in susceptible species | 20169085;25288078 |
| Streptococcus dysgalactiae subsp. equisimilis strain UT_4234_DH | folA, Dfr | Dihydrofolate reductase (EC 1.5.1.3) | | | antibiotic target in susceptible species | 20169085;25288078 |
| Streptococcus dysgalactiae subsp. equisimilis strain UT_4241_XS | folA, Dfr | Dihydrofolate reductase (EC 1.5.1.3) | | | antibiotic target in susceptible species | 20169085;25288078 |
| Streptococcus dysgalactiae subsp. equisimilis strain UT_4242_AB | folA, Dfr | Dihydrofolate reductase (EC 1.5.1.3) | | | antibiotic target in susceptible species | 20169085;25288078 |
| Streptococcus dysgalactiae subsp. equisimilis strain UT_4255RC | folA, Dfr | Dihydrofolate reductase (EC 1.5.1.3) | | | antibiotic target in susceptible species | 20169085;25288078 |
| Streptococcus dysgalactiae subsp. equisimilis strain UT_4277_BB | folA, Dfr | Dihydrofolate reductase (EC 1.5.1.3) | | | antibiotic target in susceptible species | 20169085;25288078 |
| Streptococcus dysgalactiae subsp. equisimilis strain UT_4966_RC | folA, Dfr | Dihydrofolate reductase (EC 1.5.1.3) | | | antibiotic target in susceptible species | 20169085;25288078 |
| Streptococcus dysgalactiae subsp. equisimilis strain UT-5345 | folA, Dfr | Dihydrofolate reductase (EC 1.5.1.3) | | | antibiotic target in susceptible species | 20169085;25288078 |
| Streptococcus dysgalactiae subsp. equisimilis strain UT-5354 | folA, Dfr | Dihydrofolate reductase (EC 1.5.1.3) | | | antibiotic target in susceptible species | 20169085;25288078 |
| Streptococcus dysgalactiae subsp. equisimilis strain UT-SS1069 | folA, Dfr | Dihydrofolate reductase (EC 1.5.1.3) | | | antibiotic target in susceptible species | 20169085;25288078 |
| Streptococcus dysgalactiae subsp. equisimilis strain UT-SS957 | folA, Dfr | Dihydrofolate reductase (EC 1.5.1.3) | | | antibiotic target in susceptible species | 20169085;25288078 |
| Streptococcus dysgalactiae subsp. equisimilis strain WCHSDSE-1 | folA, Dfr | Dihydrofolate reductase (EC 1.5.1.3) | | | antibiotic target in susceptible species | 20169085;25288078 |
| Streptococcus dysgalactiae subsp. equisimilis SD SCDR1 | Ddl | D-alanine--D-alanine ligase (EC 6.3.2.4) | | | antibiotic target in susceptible species | 24303782;24033232 |
| Streptococcus dysgalactiae subsp. equisimilis 167 | ddl | D-alanine--D-alanine ligase (EC 6.3.2.4) | | | antibiotic target in susceptible species | 24303782;24033232 |
| Streptococcus dysgalactiae subsp. equisimilis AC-2713 | ddl | D-alanine--D-alanine ligase (EC 6.3.2.4) | | | antibiotic target in susceptible species | 24303782;24033232 |
| Streptococcus dysgalactiae subsp. equisimilis AKSDE4288 | ddl | D-alanine--D-alanine ligase (EC 6.3.2.4) | | | antibiotic target in susceptible species | 24303782;24033232 |
| Streptococcus dysgalactiae subsp. equisimilis ATCC 12394 | ddl | D-alanine--D-alanine ligase (EC 6.3.2.4) | | | antibiotic target in susceptible species | 24303782;24033232 |
| Streptococcus dysgalactiae subsp. equisimilis GGS_124 | ddl | D-alanine--D-alanine ligase (EC 6.3.2.4) | | | antibiotic target in susceptible species | 24303782;24033232 |
| Streptococcus dysgalactiae subsp. equisimilis RE378 | ddl | D-alanine--D-alanine ligase (EC 6.3.2.4) | | | antibiotic target in susceptible species | 24303782;24033232 |
| Streptococcus dysgalactiae subsp. equisimilis SK1249 | ddl | D-alanine--D-alanine ligase (EC 6.3.2.4) | | | antibiotic target in susceptible species | 24303782;24033232 |
| Streptococcus dysgalactiae subsp. equisimilis SK1250 | ddl | D-alanine--D-alanine ligase (EC 6.3.2.4) | | | antibiotic target in susceptible species | 24303782;24033232 |
| Streptococcus dysgalactiae subsp. equisimilis strain ASDSE_96 | ddl | D-alanine--D-alanine ligase (EC 6.3.2.4) | | | antibiotic target in susceptible species | 24303782;24033232 |
| Streptococcus dysgalactiae subsp. equisimilis strain ASDSE_99 | ddl | D-alanine--D-alanine ligase (EC 6.3.2.4) | | | antibiotic target in susceptible species | 24303782;24033232 |
| Streptococcus dysgalactiae subsp. equisimilis strain C161L1 | ddl | D-alanine--D-alanine ligase (EC 6.3.2.4) | | | antibiotic target in susceptible species | 24303782;24033232 |
| Streptococcus dysgalactiae subsp. equisimilis strain KNZ01 | Ddl | D-alanine--D-alanine ligase (EC 6.3.2.4) | | | antibiotic target in susceptible species | 24303782;24033232 |
| Streptococcus dysgalactiae subsp. equisimilis strain KNZ01 | Ddl | D-alanine--D-alanine ligase (EC 6.3.2.4) | | | antibiotic target in susceptible species | 24303782;24033232 |
| Streptococcus dysgalactiae subsp. equisimilis strain KNZ03 | Ddl | D-alanine--D-alanine ligase (EC 6.3.2.4) | | | antibiotic target in susceptible species | 24303782;24033232 |
| Streptococcus dysgalactiae subsp. equisimilis strain KNZ03 | Ddl | D-alanine--D-alanine ligase (EC 6.3.2.4) | | | antibiotic target in susceptible species | 24303782;24033232 |
| Streptococcus dysgalactiae subsp. equisimilis strain KNZ04 | Ddl | D-alanine--D-alanine ligase (EC 6.3.2.4) | | | antibiotic target in susceptible species | 24303782;24033232 |
| Streptococcus dysgalactiae subsp. equisimilis strain KNZ06 | Ddl | D-alanine--D-alanine ligase (EC 6.3.2.4) | | | antibiotic target in susceptible species | 24303782;24033232 |
| Streptococcus dysgalactiae subsp. equisimilis strain KNZ07 | Ddl | D-alanine--D-alanine ligase (EC 6.3.2.4) | | | antibiotic target in susceptible species | 24303782;24033232 |
| Streptococcus dysgalactiae subsp. equisimilis strain KNZ10 | Ddl | D-alanine--D-alanine ligase (EC 6.3.2.4) | | | antibiotic target in susceptible species | 24303782;24033232 |
| Streptococcus dysgalactiae subsp. equisimilis strain KNZ12 | Ddl | D-alanine--D-alanine ligase (EC 6.3.2.4) | | | antibiotic target in susceptible species | 24303782;24033232 |
| Streptococcus dysgalactiae subsp. equisimilis strain KNZ15 | Ddl | D-alanine--D-alanine ligase (EC 6.3.2.4) | | | antibiotic target in susceptible species | 24303782;24033232 |
| Streptococcus dysgalactiae subsp. equisimilis strain KNZ16 | Ddl | D-alanine--D-alanine ligase (EC 6.3.2.4) | | | antibiotic target in susceptible species | 24303782;24033232 |
| Streptococcus dysgalactiae subsp. equisimilis strain NCTC10321 | Ddl | D-alanine--D-alanine ligase (EC 6.3.2.4) | | | antibiotic target in susceptible species | 24303782;24033232 |
| Streptococcus dysgalactiae subsp. equisimilis strain NCTC10321 | Ddl | D-alanine--D-alanine ligase (EC 6.3.2.4) | | | antibiotic target in susceptible species | 24303782;24033232 |
| Streptococcus dysgalactiae subsp. equisimilis strain NCTC11554 | Ddl | D-alanine--D-alanine ligase (EC 6.3.2.4) | | | antibiotic target in susceptible species | 24303782;24033232 |
| Streptococcus dysgalactiae subsp. equisimilis strain NCTC11555 | Ddl | D-alanine--D-alanine ligase (EC 6.3.2.4) | | | antibiotic target in susceptible species | 24303782;24033232 |
| Streptococcus dysgalactiae subsp. equisimilis strain NCTC11556 | Ddl | D-alanine--D-alanine ligase (EC 6.3.2.4) | | | antibiotic target in susceptible species | 24303782;24033232 |
| Streptococcus dysgalactiae subsp. equisimilis strain NCTC11557 | Ddl | D-alanine--D-alanine ligase (EC 6.3.2.4) | | | antibiotic target in susceptible species | 24303782;24033232 |
| Streptococcus dysgalactiae subsp. equisimilis strain NCTC11564 | Ddl | D-alanine--D-alanine ligase (EC 6.3.2.4) | | | antibiotic target in susceptible species | 24303782;24033232 |
| Streptococcus dysgalactiae subsp. equisimilis strain NCTC11565 | Ddl | D-alanine--D-alanine ligase (EC 6.3.2.4) | | | antibiotic target in susceptible species | 24303782;24033232 |
| Streptococcus dysgalactiae subsp. equisimilis strain NCTC11565 | Ddl | D-alanine--D-alanine ligase (EC 6.3.2.4) | | | antibiotic target in susceptible species | 24303782;24033232 |
| Streptococcus dysgalactiae subsp. equisimilis strain NCTC5370 | ddl | D-alanine--D-alanine ligase (EC 6.3.2.4) | | | antibiotic target in susceptible species | 24303782;24033232 |
| Streptococcus dysgalactiae subsp. equisimilis strain NCTC5371 | ddl | D-alanine--D-alanine ligase (EC 6.3.2.4) | | | antibiotic target in susceptible species | 24303782;24033232 |
| Streptococcus dysgalactiae subsp. equisimilis strain NCTC5969 | Ddl | D-alanine--D-alanine ligase (EC 6.3.2.4) | | | antibiotic target in susceptible species | 24303782;24033232 |
| Streptococcus dysgalactiae subsp. equisimilis strain NCTC6179 | ddl | D-alanine--D-alanine ligase (EC 6.3.2.4) | | | antibiotic target in susceptible species | 24303782;24033232 |
| Streptococcus dysgalactiae subsp. equisimilis strain NCTC6181 | Ddl | D-alanine--D-alanine ligase (EC 6.3.2.4) | | | antibiotic target in susceptible species | 24303782;24033232 |
| Streptococcus dysgalactiae subsp. equisimilis strain NCTC6407 | Ddl | D-alanine--D-alanine ligase (EC 6.3.2.4) | | | antibiotic target in susceptible species | 24303782;24033232 |
| Streptococcus dysgalactiae subsp. equisimilis strain NCTC6407 | Ddl | D-alanine--D-alanine ligase (EC 6.3.2.4) | | | antibiotic target in susceptible species | 24303782;24033232 |
| Streptococcus dysgalactiae subsp. equisimilis strain NCTC7136 | ddl | D-alanine--D-alanine ligase (EC 6.3.2.4) | | | antibiotic target in susceptible species | 24303782;24033232 |
| Streptococcus dysgalactiae subsp. equisimilis strain NCTC8543 | Ddl | D-alanine--D-alanine ligase (EC 6.3.2.4) | | | antibiotic target in susceptible species | 24303782;24033232 |
| Streptococcus dysgalactiae subsp. equisimilis strain NCTC8546 | Ddl | D-alanine--D-alanine ligase (EC 6.3.2.4) | | | antibiotic target in susceptible species | 24303782;24033232 |
| Streptococcus dysgalactiae subsp. equisimilis strain NCTC9413 | Ddl | D-alanine--D-alanine ligase (EC 6.3.2.4) | | | antibiotic target in susceptible species | 24303782;24033232 |
| Streptococcus dysgalactiae subsp. equisimilis strain NCTC9414 | ddl | D-alanine--D-alanine ligase (EC 6.3.2.4) | | | antibiotic target in susceptible species | 24303782;24033232 |
| Streptococcus dysgalactiae subsp. equisimilis strain NCTC9603 | Ddl | D-alanine--D-alanine ligase (EC 6.3.2.4) | | | antibiotic target in susceptible species | 24303782;24033232 |
| Streptococcus dysgalactiae subsp. equisimilis strain SS1575 | ddl | D-alanine--D-alanine ligase (EC 6.3.2.4) | | | antibiotic target in susceptible species | 24303782;24033232 |
| Streptococcus dysgalactiae subsp. equisimilis strain T642 | ddl | D-alanine--D-alanine ligase (EC 6.3.2.4) | | | antibiotic target in susceptible species | 24303782;24033232 |
| Streptococcus dysgalactiae subsp. equisimilis strain UT_4031CC | ddl | D-alanine--D-alanine ligase (EC 6.3.2.4) | | | antibiotic target in susceptible species | 24303782;24033232 |
| Streptococcus dysgalactiae subsp. equisimilis strain UT_4231_KK | ddl | D-alanine--D-alanine ligase (EC 6.3.2.4) | | | antibiotic target in susceptible species | 24303782;24033232 |
| Streptococcus dysgalactiae subsp. equisimilis strain UT_4234_DH | ddl | D-alanine--D-alanine ligase (EC 6.3.2.4) | | | antibiotic target in susceptible species | 24303782;24033232 |
| Streptococcus dysgalactiae subsp. equisimilis strain UT_4241_XS | ddl | D-alanine--D-alanine ligase (EC 6.3.2.4) | | | antibiotic target in susceptible species | 24303782;24033232 |
| Streptococcus dysgalactiae subsp. equisimilis strain UT_4242_AB | ddl | D-alanine--D-alanine ligase (EC 6.3.2.4) | | | antibiotic target in susceptible species | 24303782;24033232 |
| Streptococcus dysgalactiae subsp. equisimilis strain UT_4255RC | ddl | D-alanine--D-alanine ligase (EC 6.3.2.4) | | | antibiotic target in susceptible species | 24303782;24033232 |
| Streptococcus dysgalactiae subsp. equisimilis strain UT_4277_BB | ddl | D-alanine--D-alanine ligase (EC 6.3.2.4) | | | antibiotic target in susceptible species | 24303782;24033232 |
| Streptococcus dysgalactiae subsp. equisimilis strain UT_4966_RC | ddl | D-alanine--D-alanine ligase (EC 6.3.2.4) | | | antibiotic target in susceptible species | 24303782;24033232 |
| Streptococcus dysgalactiae subsp. equisimilis strain UT_4966_RC | ddl | D-alanine--D-alanine ligase (EC 6.3.2.4) | | | antibiotic target in susceptible species | 24303782;24033232 |
| Streptococcus dysgalactiae subsp. equisimilis strain UT-5345 | ddl | D-alanine--D-alanine ligase (EC 6.3.2.4) | | | antibiotic target in susceptible species | 24303782;24033232 |
| Streptococcus dysgalactiae subsp. equisimilis strain UT-5354 | ddl | D-alanine--D-alanine ligase (EC 6.3.2.4) | | | antibiotic target in susceptible species | 24303782;24033232 |
| Streptococcus dysgalactiae subsp. equisimilis strain UT-SS1069 | ddl | D-alanine--D-alanine ligase (EC 6.3.2.4) | | | antibiotic target in susceptible species | 24303782;24033232 |
| Streptococcus dysgalactiae subsp. equisimilis strain UT-SS957 | ddl | D-alanine--D-alanine ligase (EC 6.3.2.4) | | | antibiotic target in susceptible species | 24303782;24033232 |
| Streptococcus dysgalactiae subsp. equisimilis strain WCHSDSE-1 | ddl | D-alanine--D-alanine ligase (EC 6.3.2.4) | | | antibiotic target in susceptible species | 24303782;24033232 |
| Streptococcus dysgalactiae subsp. equisimilis SD SCDR1 | LiaS | Cell envelope stress response system LiaFSR, sensor histidine kinase LiaS(VraS) | | | regulator modulating expression of antibiotic resistance genes | 21899450;26020679 |
| Streptococcus dysgalactiae subsp. equisimilis 167 | yvqE | Cell envelope stress response system LiaFSR, sensor histidine kinase LiaS(VraS) | | | regulator modulating expression of antibiotic resistance genes | 21899450;26020679 |
| Streptococcus dysgalactiae subsp. equisimilis AC-2713 | yvqE | Cell envelope stress response system LiaFSR, sensor histidine kinase LiaS(VraS) | | | regulator modulating expression of antibiotic resistance genes | 21899450;26020679 |
| Streptococcus dysgalactiae subsp. equisimilis AKSDE4288 | LiaS | Cell envelope stress response system LiaFSR, sensor histidine kinase LiaS(VraS) | | | regulator modulating expression of antibiotic resistance genes | 21899450;26020679 |
| Streptococcus dysgalactiae subsp. equisimilis ATCC 12394 | LiaS | Cell envelope stress response system LiaFSR, sensor histidine kinase LiaS(VraS) | | | regulator modulating expression of antibiotic resistance genes | 21899450;26020679 |
| Streptococcus dysgalactiae subsp. equisimilis GGS_124 | yvqE | Cell envelope stress response system LiaFSR, sensor histidine kinase LiaS(VraS) | | | regulator modulating expression of antibiotic resistance genes | 21899450;26020679 |
| Streptococcus dysgalactiae subsp. equisimilis RE378 | yvqE | Cell envelope stress response system LiaFSR, sensor histidine kinase LiaS(VraS) | | | regulator modulating expression of antibiotic resistance genes | 21899450;26020679 |
| Streptococcus dysgalactiae subsp. equisimilis SK1249 | LiaS | Cell envelope stress response system LiaFSR, sensor histidine kinase LiaS(VraS) | | | regulator modulating expression of antibiotic resistance genes | 21899450;26020679 |
| Streptococcus dysgalactiae subsp. equisimilis SK1250 | LiaS | Cell envelope stress response system LiaFSR, sensor histidine kinase LiaS(VraS) | | | regulator modulating expression of antibiotic resistance genes | 21899450;26020679 |
| Streptococcus dysgalactiae subsp. equisimilis strain ASDSE_96 | LiaS | Cell envelope stress response system LiaFSR, sensor histidine kinase LiaS(VraS) | | | regulator modulating expression of antibiotic resistance genes | 21899450;26020679 |
| Streptococcus dysgalactiae subsp. equisimilis strain ASDSE_99 | LiaS | Cell envelope stress response system LiaFSR, sensor histidine kinase LiaS(VraS) | | | regulator modulating expression of antibiotic resistance genes | 21899450;26020679 |
| Streptococcus dysgalactiae subsp. equisimilis strain C161L1 | LiaS | Cell envelope stress response system LiaFSR, sensor histidine kinase LiaS(VraS) | | | regulator modulating expression of antibiotic resistance genes | 21899450;26020679 |
| Streptococcus dysgalactiae subsp. equisimilis strain KNZ01 | LiaS | Cell envelope stress response system LiaFSR, sensor histidine kinase LiaS(VraS) | | | regulator modulating expression of antibiotic resistance genes | 21899450;26020679 |
| Streptococcus dysgalactiae subsp. equisimilis strain KNZ03 | LiaS | Cell envelope stress response system LiaFSR, sensor histidine kinase LiaS(VraS) | | | regulator modulating expression of antibiotic resistance genes | 21899450;26020679 |
| Streptococcus dysgalactiae subsp. equisimilis strain KNZ04 | LiaS | Cell envelope stress response system LiaFSR, sensor histidine kinase LiaS(VraS) | | | regulator modulating expression of antibiotic resistance genes | 21899450;26020679 |
| Streptococcus dysgalactiae subsp. equisimilis strain KNZ06 | LiaS | Cell envelope stress response system LiaFSR, sensor histidine kinase LiaS(VraS) | | | regulator modulating expression of antibiotic resistance genes | 21899450;26020679 |
| Streptococcus dysgalactiae subsp. equisimilis strain KNZ07 | LiaS | Cell envelope stress response system LiaFSR, sensor histidine kinase LiaS(VraS) | | | regulator modulating expression of antibiotic resistance genes | 21899450;26020679 |
| Streptococcus dysgalactiae subsp. equisimilis strain KNZ10 | LiaS | Cell envelope stress response system LiaFSR, sensor histidine kinase LiaS(VraS) | | | regulator modulating expression of antibiotic resistance genes | 21899450;26020679 |
| Streptococcus dysgalactiae subsp. equisimilis strain KNZ12 | LiaS | Cell envelope stress response system LiaFSR, sensor histidine kinase LiaS(VraS) | | | regulator modulating expression of antibiotic resistance genes | 21899450;26020679 |
| Streptococcus dysgalactiae subsp. equisimilis strain KNZ15 | LiaS | Cell envelope stress response system LiaFSR, sensor histidine kinase LiaS(VraS) | | | regulator modulating expression of antibiotic resistance genes | 21899450;26020679 |
| Streptococcus dysgalactiae subsp. equisimilis strain KNZ16 | LiaS | Cell envelope stress response system LiaFSR, sensor histidine kinase LiaS(VraS) | | | regulator modulating expression of antibiotic resistance genes | 21899450;26020679 |
| Streptococcus dysgalactiae subsp. equisimilis strain NCTC10321 | LiaS | Cell envelope stress response system LiaFSR, sensor histidine kinase LiaS(VraS) | | | regulator modulating expression of antibiotic resistance genes | 21899450;26020679 |
| Streptococcus dysgalactiae subsp. equisimilis strain NCTC11554 | LiaS | Cell envelope stress response system LiaFSR, sensor histidine kinase LiaS(VraS) | | | regulator modulating expression of antibiotic resistance genes | 21899450;26020679 |
| Streptococcus dysgalactiae subsp. equisimilis strain NCTC11555 | LiaS | Cell envelope stress response system LiaFSR, sensor histidine kinase LiaS(VraS) | | | regulator modulating expression of antibiotic resistance genes | 21899450;26020679 |
| Streptococcus dysgalactiae subsp. equisimilis strain NCTC11556 | LiaS | Cell envelope stress response system LiaFSR, sensor histidine kinase LiaS(VraS) | | | regulator modulating expression of antibiotic resistance genes | 21899450;26020679 |
| Streptococcus dysgalactiae subsp. equisimilis strain NCTC11557 | LiaS | Cell envelope stress response system LiaFSR, sensor histidine kinase LiaS(VraS) | | | regulator modulating expression of antibiotic resistance genes | 21899450;26020679 |
| Streptococcus dysgalactiae subsp. equisimilis strain NCTC11564 | LiaS | Cell envelope stress response system LiaFSR, sensor histidine kinase LiaS(VraS) | | | regulator modulating expression of antibiotic resistance genes | 21899450;26020679 |
| Streptococcus dysgalactiae subsp. equisimilis strain NCTC5370 | yvqE | Cell envelope stress response system LiaFSR, sensor histidine kinase LiaS(VraS) | | | regulator modulating expression of antibiotic resistance genes | 21899450;26020679 |
| Streptococcus dysgalactiae subsp. equisimilis strain NCTC5371 | yvqE | Cell envelope stress response system LiaFSR, sensor histidine kinase LiaS(VraS) | | | regulator modulating expression of antibiotic resistance genes | 21899450;26020679 |
| Streptococcus dysgalactiae subsp. equisimilis strain NCTC5969 | LiaS | Cell envelope stress response system LiaFSR, sensor histidine kinase LiaS(VraS) | | | regulator modulating expression of antibiotic resistance genes | 21899450;26020679 |
| Streptococcus dysgalactiae subsp. equisimilis strain NCTC6179 | yvqE | Cell envelope stress response system LiaFSR, sensor histidine kinase LiaS(VraS) | | | regulator modulating expression of antibiotic resistance genes | 21899450;26020679 |
| Streptococcus dysgalactiae subsp. equisimilis strain NCTC6181 | LiaS | Cell envelope stress response system LiaFSR, sensor histidine kinase LiaS(VraS) | | | regulator modulating expression of antibiotic resistance genes | 21899450;26020679 |
| Streptococcus dysgalactiae subsp. equisimilis strain NCTC6407 | LiaS | Cell envelope stress response system LiaFSR, sensor histidine kinase LiaS(VraS) | | | regulator modulating expression of antibiotic resistance genes | 21899450;26020679 |
| Streptococcus dysgalactiae subsp. equisimilis strain NCTC7136 | yvqE | Cell envelope stress response system LiaFSR, sensor histidine kinase LiaS(VraS) | | | regulator modulating expression of antibiotic resistance genes | 21899450;26020679 |
| Streptococcus dysgalactiae subsp. equisimilis strain NCTC8543 | LiaS | Cell envelope stress response system LiaFSR, sensor histidine kinase LiaS(VraS) | | | regulator modulating expression of antibiotic resistance genes | 21899450;26020679 |
| Streptococcus dysgalactiae subsp. equisimilis strain NCTC8546 | LiaS | Cell envelope stress response system LiaFSR, sensor histidine kinase LiaS(VraS) | | | regulator modulating expression of antibiotic resistance genes | 21899450;26020679 |
| Streptococcus dysgalactiae subsp. equisimilis strain NCTC9413 | LiaS | Cell envelope stress response system LiaFSR, sensor histidine kinase LiaS(VraS) | | | regulator modulating expression of antibiotic resistance genes | 21899450;26020679 |
| Streptococcus dysgalactiae subsp. equisimilis strain NCTC9414 | yvqE | Cell envelope stress response system LiaFSR, sensor histidine kinase LiaS(VraS) | | | regulator modulating expression of antibiotic resistance genes | 21899450;26020679 |
| Streptococcus dysgalactiae subsp. equisimilis strain NCTC9603 | LiaS | Cell envelope stress response system LiaFSR, sensor histidine kinase LiaS(VraS) | | | regulator modulating expression of antibiotic resistance genes | 21899450;26020679 |
| Streptococcus dysgalactiae subsp. equisimilis strain SS1575 | LiaS | Cell envelope stress response system LiaFSR, sensor histidine kinase LiaS(VraS) | | | regulator modulating expression of antibiotic resistance genes | 21899450;26020679 |
| Streptococcus dysgalactiae subsp. equisimilis strain T642 | LiaS | Cell envelope stress response system LiaFSR, sensor histidine kinase LiaS(VraS) | | | regulator modulating expression of antibiotic resistance genes | 21899450;26020679 |
| Streptococcus dysgalactiae subsp. equisimilis strain UT_4031CC | LiaS | Cell envelope stress response system LiaFSR, sensor histidine kinase LiaS(VraS) | | | regulator modulating expression of antibiotic resistance genes | 21899450;26020679 |
| Streptococcus dysgalactiae subsp. equisimilis strain UT_4231_KK | LiaS | Cell envelope stress response system LiaFSR, sensor histidine kinase LiaS(VraS) | | | regulator modulating expression of antibiotic resistance genes | 21899450;26020679 |
| Streptococcus dysgalactiae subsp. equisimilis strain UT_4234_DH | LiaS | Cell envelope stress response system LiaFSR, sensor histidine kinase LiaS(VraS) | | | regulator modulating expression of antibiotic resistance genes | 21899450;26020679 |
| Streptococcus dysgalactiae subsp. equisimilis strain UT_4241_XS | LiaS | Cell envelope stress response system LiaFSR, sensor histidine kinase LiaS(VraS) | | | regulator modulating expression of antibiotic resistance genes | 21899450;26020679 |
| Streptococcus dysgalactiae subsp. equisimilis strain UT_4242_AB | LiaS | Cell envelope stress response system LiaFSR, sensor histidine kinase LiaS(VraS) | | | regulator modulating expression of antibiotic resistance genes | 21899450;26020679 |
| Streptococcus dysgalactiae subsp. equisimilis strain UT_4255RC | LiaS | Cell envelope stress response system LiaFSR, sensor histidine kinase LiaS(VraS) | | | regulator modulating expression of antibiotic resistance genes | 21899450;26020679 |
| Streptococcus dysgalactiae subsp. equisimilis strain UT_4277_BB | LiaS | Cell envelope stress response system LiaFSR, sensor histidine kinase LiaS(VraS) | | | regulator modulating expression of antibiotic resistance genes | 21899450;26020679 |
| Streptococcus dysgalactiae subsp. equisimilis strain UT_4966_RC | LiaS | Cell envelope stress response system LiaFSR, sensor histidine kinase LiaS(VraS) | | | regulator modulating expression of antibiotic resistance genes | 21899450;26020679 |
| Streptococcus dysgalactiae subsp. equisimilis strain UT-5345 | LiaS | Cell envelope stress response system LiaFSR, sensor histidine kinase LiaS(VraS) | | | regulator modulating expression of antibiotic resistance genes | 21899450;26020679 |
| Streptococcus dysgalactiae subsp. equisimilis strain UT-5354 | LiaS | Cell envelope stress response system LiaFSR, sensor histidine kinase LiaS(VraS) | | | regulator modulating expression of antibiotic resistance genes | 21899450;26020679 |
| Streptococcus dysgalactiae subsp. equisimilis strain UT-SS1069 | LiaS | Cell envelope stress response system LiaFSR, sensor histidine kinase LiaS(VraS) | | | regulator modulating expression of antibiotic resistance genes | 21899450;26020679 |
| Streptococcus dysgalactiae subsp. equisimilis strain UT-SS957 | LiaS | Cell envelope stress response system LiaFSR, sensor histidine kinase LiaS(VraS) | | | regulator modulating expression of antibiotic resistance genes | 21899450;26020679 |
| Streptococcus dysgalactiae subsp. equisimilis strain WCHSDSE-1 | LiaS | Cell envelope stress response system LiaFSR, sensor histidine kinase LiaS(VraS) | | | regulator modulating expression of antibiotic resistance genes | 21899450;26020679 |
| Streptococcus dysgalactiae subsp. equisimilis SD SCDR1 | PgsA | CDP-diacylglycerol--glycerol-3-phosphate 3-phosphatidyltransferase (EC 2.7.8.5) | | | protein altering cell wall charge conferring antibiotic resistance | 22238576 |
| Streptococcus dysgalactiae subsp. equisimilis 167 | pgsA | CDP-diacylglycerol--glycerol-3-phosphate 3-phosphatidyltransferase (EC 2.7.8.5) | | | protein altering cell wall charge conferring antibiotic resistance | 22238576 |
| Streptococcus dysgalactiae subsp. equisimilis AC-2713 | pgsA | CDP-diacylglycerol--glycerol-3-phosphate 3-phosphatidyltransferase (EC 2.7.8.5) | | | protein altering cell wall charge conferring antibiotic resistance | 22238576 |
| Streptococcus dysgalactiae subsp. equisimilis AKSDE4288 | pgsA | CDP-diacylglycerol--glycerol-3-phosphate 3-phosphatidyltransferase (EC 2.7.8.5) | | | protein altering cell wall charge conferring antibiotic resistance | 22238576 |
| Streptococcus dysgalactiae subsp. equisimilis ATCC 12394 | pgsA | CDP-diacylglycerol--glycerol-3-phosphate 3-phosphatidyltransferase (EC 2.7.8.5) | | | protein altering cell wall charge conferring antibiotic resistance | 22238576 |
| Streptococcus dysgalactiae subsp. equisimilis GGS_124 | pgsA | CDP-diacylglycerol--glycerol-3-phosphate 3-phosphatidyltransferase (EC 2.7.8.5) | | | protein altering cell wall charge conferring antibiotic resistance | 22238576 |
| Streptococcus dysgalactiae subsp. equisimilis RE378 | pgsA | CDP-diacylglycerol--glycerol-3-phosphate 3-phosphatidyltransferase (EC 2.7.8.5) | | | protein altering cell wall charge conferring antibiotic resistance | 22238576 |
| Streptococcus dysgalactiae subsp. equisimilis SK1249 | pgsA | CDP-diacylglycerol--glycerol-3-phosphate 3-phosphatidyltransferase (EC 2.7.8.5) | | | protein altering cell wall charge conferring antibiotic resistance | 22238576 |
| Streptococcus dysgalactiae subsp. equisimilis SK1250 | pgsA | CDP-diacylglycerol--glycerol-3-phosphate 3-phosphatidyltransferase (EC 2.7.8.5) | | | protein altering cell wall charge conferring antibiotic resistance | 22238576 |
| Streptococcus dysgalactiae subsp. equisimilis strain ASDSE_96 | pgsA | CDP-diacylglycerol--glycerol-3-phosphate 3-phosphatidyltransferase (EC 2.7.8.5) | | | protein altering cell wall charge conferring antibiotic resistance | 22238576 |
| Streptococcus dysgalactiae subsp. equisimilis strain ASDSE_99 | pgsA | CDP-diacylglycerol--glycerol-3-phosphate 3-phosphatidyltransferase (EC 2.7.8.5) | | | protein altering cell wall charge conferring antibiotic resistance | 22238576 |
| Streptococcus dysgalactiae subsp. equisimilis strain C161L1 | pgsA | CDP-diacylglycerol--glycerol-3-phosphate 3-phosphatidyltransferase (EC 2.7.8.5) | | | protein altering cell wall charge conferring antibiotic resistance | 22238576 |
| Streptococcus dysgalactiae subsp. equisimilis strain KNZ01 | PgsA | CDP-diacylglycerol--glycerol-3-phosphate 3-phosphatidyltransferase (EC 2.7.8.5) | | | protein altering cell wall charge conferring antibiotic resistance | 22238576 |
| Streptococcus dysgalactiae subsp. equisimilis strain KNZ01 | PgsA | CDP-diacylglycerol--glycerol-3-phosphate 3-phosphatidyltransferase (EC 2.7.8.5) | | | protein altering cell wall charge conferring antibiotic resistance | 22238576 |
| Streptococcus dysgalactiae subsp. equisimilis strain KNZ03 | PgsA | CDP-diacylglycerol--glycerol-3-phosphate 3-phosphatidyltransferase (EC 2.7.8.5) | | | protein altering cell wall charge conferring antibiotic resistance | 22238576 |
| Streptococcus dysgalactiae subsp. equisimilis strain KNZ03 | PgsA | CDP-diacylglycerol--glycerol-3-phosphate 3-phosphatidyltransferase (EC 2.7.8.5) | | | protein altering cell wall charge conferring antibiotic resistance | 22238576 |
| Streptococcus dysgalactiae subsp. equisimilis strain KNZ04 | PgsA | CDP-diacylglycerol--glycerol-3-phosphate 3-phosphatidyltransferase (EC 2.7.8.5) | | | protein altering cell wall charge conferring antibiotic resistance | 22238576 |
| Streptococcus dysgalactiae subsp. equisimilis strain KNZ06 | PgsA | CDP-diacylglycerol--glycerol-3-phosphate 3-phosphatidyltransferase (EC 2.7.8.5) | | | protein altering cell wall charge conferring antibiotic resistance | 22238576 |
| Streptococcus dysgalactiae subsp. equisimilis strain KNZ07 | PgsA | CDP-diacylglycerol--glycerol-3-phosphate 3-phosphatidyltransferase (EC 2.7.8.5) | | | protein altering cell wall charge conferring antibiotic resistance | 22238576 |
| Streptococcus dysgalactiae subsp. equisimilis strain KNZ10 | PgsA | CDP-diacylglycerol--glycerol-3-phosphate 3-phosphatidyltransferase (EC 2.7.8.5) | | | protein altering cell wall charge conferring antibiotic resistance | 22238576 |
| Streptococcus dysgalactiae subsp. equisimilis strain KNZ12 | PgsA | CDP-diacylglycerol--glycerol-3-phosphate 3-phosphatidyltransferase (EC 2.7.8.5) | | | protein altering cell wall charge conferring antibiotic resistance | 22238576 |
| Streptococcus dysgalactiae subsp. equisimilis strain KNZ15 | PgsA | CDP-diacylglycerol--glycerol-3-phosphate 3-phosphatidyltransferase (EC 2.7.8.5) | | | protein altering cell wall charge conferring antibiotic resistance | 22238576 |
| Streptococcus dysgalactiae subsp. equisimilis strain KNZ16 | PgsA | CDP-diacylglycerol--glycerol-3-phosphate 3-phosphatidyltransferase (EC 2.7.8.5) | | | protein altering cell wall charge conferring antibiotic resistance | 22238576 |
| Streptococcus dysgalactiae subsp. equisimilis strain NCTC10321 | PgsA | CDP-diacylglycerol--glycerol-3-phosphate 3-phosphatidyltransferase (EC 2.7.8.5) | | | protein altering cell wall charge conferring antibiotic resistance | 22238576 |
| Streptococcus dysgalactiae subsp. equisimilis strain NCTC11554 | PgsA | CDP-diacylglycerol--glycerol-3-phosphate 3-phosphatidyltransferase (EC 2.7.8.5) | | | protein altering cell wall charge conferring antibiotic resistance | 22238576 |
| Streptococcus dysgalactiae subsp. equisimilis strain NCTC11555 | PgsA | CDP-diacylglycerol--glycerol-3-phosphate 3-phosphatidyltransferase (EC 2.7.8.5) | | | protein altering cell wall charge conferring antibiotic resistance | 22238576 |
| Streptococcus dysgalactiae subsp. equisimilis strain NCTC11556 | PgsA | CDP-diacylglycerol--glycerol-3-phosphate 3-phosphatidyltransferase (EC 2.7.8.5) | | | protein altering cell wall charge conferring antibiotic resistance | 22238576 |
| Streptococcus dysgalactiae subsp. equisimilis strain NCTC11557 | PgsA | CDP-diacylglycerol--glycerol-3-phosphate 3-phosphatidyltransferase (EC 2.7.8.5) | | | protein altering cell wall charge conferring antibiotic resistance | 22238576 |
| Streptococcus dysgalactiae subsp. equisimilis strain NCTC11564 | PgsA | CDP-diacylglycerol--glycerol-3-phosphate 3-phosphatidyltransferase (EC 2.7.8.5) | | | protein altering cell wall charge conferring antibiotic resistance | 22238576 |
| Streptococcus dysgalactiae subsp. equisimilis strain NCTC11565 | PgsA | CDP-diacylglycerol--glycerol-3-phosphate 3-phosphatidyltransferase (EC 2.7.8.5) | | | protein altering cell wall charge conferring antibiotic resistance | 22238576 |
| Streptococcus dysgalactiae subsp. equisimilis strain NCTC5370 | pgsA | CDP-diacylglycerol--glycerol-3-phosphate 3-phosphatidyltransferase (EC 2.7.8.5) | | | protein altering cell wall charge conferring antibiotic resistance | 22238576 |
| Streptococcus dysgalactiae subsp. equisimilis strain NCTC5371 | pgsA | CDP-diacylglycerol--glycerol-3-phosphate 3-phosphatidyltransferase (EC 2.7.8.5) | | | protein altering cell wall charge conferring antibiotic resistance | 22238576 |
| Streptococcus dysgalactiae subsp. equisimilis strain NCTC5969 | PgsA | CDP-diacylglycerol--glycerol-3-phosphate 3-phosphatidyltransferase (EC 2.7.8.5) | | | protein altering cell wall charge conferring antibiotic resistance | 22238576 |
| Streptococcus dysgalactiae subsp. equisimilis strain NCTC5969 | pgsA | CDP-diacylglycerol--glycerol-3-phosphate 3-phosphatidyltransferase (EC 2.7.8.5) | | | antibiotic resistant gene variant or mutant,lipopeptide antibiotic resistance gene | 22238576 |
| Streptococcus dysgalactiae subsp. equisimilis strain NCTC6179 | pgsA | CDP-diacylglycerol--glycerol-3-phosphate 3-phosphatidyltransferase (EC 2.7.8.5) | | | protein altering cell wall charge conferring antibiotic resistance | 22238576 |
| Streptococcus dysgalactiae subsp. equisimilis strain NCTC6181 | PgsA | CDP-diacylglycerol--glycerol-3-phosphate 3-phosphatidyltransferase (EC 2.7.8.5) | | | protein altering cell wall charge conferring antibiotic resistance | 22238576 |
| Streptococcus dysgalactiae subsp. equisimilis strain NCTC6407 | PgsA | CDP-diacylglycerol--glycerol-3-phosphate 3-phosphatidyltransferase (EC 2.7.8.5) | | | protein altering cell wall charge conferring antibiotic resistance | 22238576 |
| Streptococcus dysgalactiae subsp. equisimilis strain NCTC7136 | pgsA | CDP-diacylglycerol--glycerol-3-phosphate 3-phosphatidyltransferase (EC 2.7.8.5) | | | protein altering cell wall charge conferring antibiotic resistance | 22238576 |
| Streptococcus dysgalactiae subsp. equisimilis strain NCTC8543 | PgsA | CDP-diacylglycerol--glycerol-3-phosphate 3-phosphatidyltransferase (EC 2.7.8.5) | | | protein altering cell wall charge conferring antibiotic resistance | 22238576 |
| Streptococcus dysgalactiae subsp. equisimilis strain NCTC8546 | PgsA | CDP-diacylglycerol--glycerol-3-phosphate 3-phosphatidyltransferase (EC 2.7.8.5) | | | protein altering cell wall charge conferring antibiotic resistance | 22238576 |
| Streptococcus dysgalactiae subsp. equisimilis strain NCTC9413 | PgsA | CDP-diacylglycerol--glycerol-3-phosphate 3-phosphatidyltransferase (EC 2.7.8.5) | | | protein altering cell wall charge conferring antibiotic resistance | 22238576 |
| Streptococcus dysgalactiae subsp. equisimilis strain NCTC9414 | pgsA | CDP-diacylglycerol--glycerol-3-phosphate 3-phosphatidyltransferase (EC 2.7.8.5) | | | protein altering cell wall charge conferring antibiotic resistance | 22238576 |
| Streptococcus dysgalactiae subsp. equisimilis strain NCTC9603 | PgsA | CDP-diacylglycerol--glycerol-3-phosphate 3-phosphatidyltransferase (EC 2.7.8.5) | | | protein altering cell wall charge conferring antibiotic resistance | 22238576 |
| Streptococcus dysgalactiae subsp. equisimilis strain SS1575 | pgsA | CDP-diacylglycerol--glycerol-3-phosphate 3-phosphatidyltransferase (EC 2.7.8.5) | | | protein altering cell wall charge conferring antibiotic resistance | 22238576 |
| Streptococcus dysgalactiae subsp. equisimilis strain T642 | pgsA | CDP-diacylglycerol--glycerol-3-phosphate 3-phosphatidyltransferase (EC 2.7.8.5) | | | protein altering cell wall charge conferring antibiotic resistance | 22238576 |
| Streptococcus dysgalactiae subsp. equisimilis strain UT_4031CC | pgsA | CDP-diacylglycerol--glycerol-3-phosphate 3-phosphatidyltransferase (EC 2.7.8.5) | | | protein altering cell wall charge conferring antibiotic resistance | 22238576 |
| Streptococcus dysgalactiae subsp. equisimilis strain UT_4231_KK | pgsA | CDP-diacylglycerol--glycerol-3-phosphate 3-phosphatidyltransferase (EC 2.7.8.5) | | | protein altering cell wall charge conferring antibiotic resistance | 22238576 |
| Streptococcus dysgalactiae subsp. equisimilis strain UT_4234_DH | pgsA | CDP-diacylglycerol--glycerol-3-phosphate 3-phosphatidyltransferase (EC 2.7.8.5) | | | protein altering cell wall charge conferring antibiotic resistance | 22238576 |
| Streptococcus dysgalactiae subsp. equisimilis strain UT_4241_XS | pgsA | CDP-diacylglycerol--glycerol-3-phosphate 3-phosphatidyltransferase (EC 2.7.8.5) | | | protein altering cell wall charge conferring antibiotic resistance | 22238576 |
| Streptococcus dysgalactiae subsp. equisimilis strain UT_4242_AB | pgsA | CDP-diacylglycerol--glycerol-3-phosphate 3-phosphatidyltransferase (EC 2.7.8.5) | | | protein altering cell wall charge conferring antibiotic resistance | 22238576 |
| Streptococcus dysgalactiae subsp. equisimilis strain UT_4255RC | pgsA | CDP-diacylglycerol--glycerol-3-phosphate 3-phosphatidyltransferase (EC 2.7.8.5) | | | protein altering cell wall charge conferring antibiotic resistance | 22238576 |
| Streptococcus dysgalactiae subsp. equisimilis strain UT_4277_BB | pgsA | CDP-diacylglycerol--glycerol-3-phosphate 3-phosphatidyltransferase (EC 2.7.8.5) | | | protein altering cell wall charge conferring antibiotic resistance | 22238576 |
| Streptococcus dysgalactiae subsp. equisimilis strain UT_4966_RC | pgsA | CDP-diacylglycerol--glycerol-3-phosphate 3-phosphatidyltransferase (EC 2.7.8.5) | | | protein altering cell wall charge conferring antibiotic resistance | 22238576 |
| Streptococcus dysgalactiae subsp. equisimilis strain UT-5345 | pgsA | CDP-diacylglycerol--glycerol-3-phosphate 3-phosphatidyltransferase (EC 2.7.8.5) | | | protein altering cell wall charge conferring antibiotic resistance | 22238576 |
| Streptococcus dysgalactiae subsp. equisimilis strain UT-5354 | pgsA | CDP-diacylglycerol--glycerol-3-phosphate 3-phosphatidyltransferase (EC 2.7.8.5) | | | protein altering cell wall charge conferring antibiotic resistance | 22238576 |
| Streptococcus dysgalactiae subsp. equisimilis strain UT-SS1069 | pgsA | CDP-diacylglycerol--glycerol-3-phosphate 3-phosphatidyltransferase (EC 2.7.8.5) | | | protein altering cell wall charge conferring antibiotic resistance | 22238576 |
| Streptococcus dysgalactiae subsp. equisimilis strain UT-SS957 | pgsA | CDP-diacylglycerol--glycerol-3-phosphate 3-phosphatidyltransferase (EC 2.7.8.5) | | | protein altering cell wall charge conferring antibiotic resistance | 22238576 |
| Streptococcus dysgalactiae subsp. equisimilis strain WCHSDSE-1 | pgsA | CDP-diacylglycerol--glycerol-3-phosphate 3-phosphatidyltransferase (EC 2.7.8.5) | | | protein altering cell wall charge conferring antibiotic resistance | 22238576 |
| Streptococcus dysgalactiae subsp. equisimilis SD SCDR1 | aadK | Aminoglycoside 6-nucleotidyltransferase, putative | | | antibiotic inactivation enzyme | 17407061 |
| Streptococcus dysgalactiae subsp. equisimilis 167 | aadK | Aminoglycoside 6-nucleotidyltransferase, putative | | | antibiotic inactivation enzyme | 17407061 |
| Streptococcus dysgalactiae subsp. equisimilis AC-2713 | aadK | Aminoglycoside 6-nucleotidyltransferase, putative | | | antibiotic inactivation enzyme | 17407061 |
| Streptococcus dysgalactiae subsp. equisimilis AKSDE4288 | aadK | Aminoglycoside 6-nucleotidyltransferase, putative | | | antibiotic inactivation enzyme | 17407061 |
| Streptococcus dysgalactiae subsp. equisimilis ATCC 12394 | aadK | Aminoglycoside 6-nucleotidyltransferase, putative | | | antibiotic inactivation enzyme | 17407061 |
| Streptococcus dysgalactiae subsp. equisimilis GGS_124 | aadK | Aminoglycoside 6-nucleotidyltransferase, putative | | | antibiotic inactivation enzyme | 17407061 |
| Streptococcus dysgalactiae subsp. equisimilis RE378 | aadK | Aminoglycoside 6-nucleotidyltransferase, putative | | | antibiotic inactivation enzyme | 17407061 |
| Streptococcus dysgalactiae subsp. equisimilis SK1249 | aadK | Aminoglycoside 6-nucleotidyltransferase, putative | | | antibiotic inactivation enzyme | 17407061 |
| Streptococcus dysgalactiae subsp. equisimilis SK1250 | aadK | Aminoglycoside 6-nucleotidyltransferase, putative | | | antibiotic inactivation enzyme | 17407061 |
| Streptococcus dysgalactiae subsp. equisimilis strain ASDSE_96 | aadK | Aminoglycoside 6-nucleotidyltransferase, putative | | | antibiotic inactivation enzyme | 17407061 |
| Streptococcus dysgalactiae subsp. equisimilis strain ASDSE_99 | aadK | Aminoglycoside 6-nucleotidyltransferase, putative | | | antibiotic inactivation enzyme | 17407061 |
| Streptococcus dysgalactiae subsp. equisimilis strain C161L1 | aadK | Aminoglycoside 6-nucleotidyltransferase, putative | | | antibiotic inactivation enzyme | 17407061 |
| Streptococcus dysgalactiae subsp. equisimilis strain KNZ01 | aadK | Aminoglycoside 6-nucleotidyltransferase, putative | | | antibiotic inactivation enzyme | 17407061 |
| Streptococcus dysgalactiae subsp. equisimilis strain KNZ03 | aadK | Aminoglycoside 6-nucleotidyltransferase, putative | | | antibiotic inactivation enzyme | 17407061 |
| Streptococcus dysgalactiae subsp. equisimilis strain KNZ04 | aadK | Aminoglycoside 6-nucleotidyltransferase, putative | | | antibiotic inactivation enzyme | 17407061 |
| Streptococcus dysgalactiae subsp. equisimilis strain KNZ06 | aadK | Aminoglycoside 6-nucleotidyltransferase, putative | | | antibiotic inactivation enzyme | 17407061 |
| Streptococcus dysgalactiae subsp. equisimilis strain KNZ07 | aadK | Aminoglycoside 6-nucleotidyltransferase, putative | | | antibiotic inactivation enzyme | 17407061 |
| Streptococcus dysgalactiae subsp. equisimilis strain KNZ10 | aadK | Aminoglycoside 6-nucleotidyltransferase, putative | | | antibiotic inactivation enzyme | 17407061 |
| Streptococcus dysgalactiae subsp. equisimilis strain KNZ12 | aadK | Aminoglycoside 6-nucleotidyltransferase, putative | | | antibiotic inactivation enzyme | 17407061 |
| Streptococcus dysgalactiae subsp. equisimilis strain KNZ15 | aadK | Aminoglycoside 6-nucleotidyltransferase, putative | | | antibiotic inactivation enzyme | 17407061 |
| Streptococcus dysgalactiae subsp. equisimilis strain KNZ16 | aadK | Aminoglycoside 6-nucleotidyltransferase, putative | | | antibiotic inactivation enzyme | 17407061 |
| Streptococcus dysgalactiae subsp. equisimilis strain NCTC11554 | aadK | Aminoglycoside 6-nucleotidyltransferase, putative | | | antibiotic inactivation enzyme | 17407061 |
| Streptococcus dysgalactiae subsp. equisimilis strain NCTC11555 | aadK | Aminoglycoside 6-nucleotidyltransferase, putative | | | antibiotic inactivation enzyme | 17407061 |
| Streptococcus dysgalactiae subsp. equisimilis strain NCTC11556 | aadK | Aminoglycoside 6-nucleotidyltransferase, putative | | | antibiotic inactivation enzyme | 17407061 |
| Streptococcus dysgalactiae subsp. equisimilis strain NCTC11557 | aadK | Aminoglycoside 6-nucleotidyltransferase, putative | | | antibiotic inactivation enzyme | 17407061 |
| Streptococcus dysgalactiae subsp. equisimilis strain NCTC11564 | aadK | Aminoglycoside 6-nucleotidyltransferase, putative | | | antibiotic inactivation enzyme | 17407061 |
| Streptococcus dysgalactiae subsp. equisimilis strain NCTC5370 | aadK | Aminoglycoside 6-nucleotidyltransferase, putative | | | antibiotic inactivation enzyme | 17407061 |
| Streptococcus dysgalactiae subsp. equisimilis strain NCTC5371 | aadK | Aminoglycoside 6-nucleotidyltransferase, putative | | | antibiotic inactivation enzyme | 17407061 |
| Streptococcus dysgalactiae subsp. equisimilis strain NCTC7136 | aadK | Aminoglycoside 6-nucleotidyltransferase, putative | | | antibiotic inactivation enzyme | 17407061 |
| Streptococcus dysgalactiae subsp. equisimilis strain NCTC8543 | aadK | Aminoglycoside 6-nucleotidyltransferase, putative | | | antibiotic inactivation enzyme | 17407061 |
| Streptococcus dysgalactiae subsp. equisimilis strain NCTC8546 | aadK | Aminoglycoside 6-nucleotidyltransferase, putative | | | antibiotic inactivation enzyme | 17407061 |
| Streptococcus dysgalactiae subsp. equisimilis strain NCTC9414 | aadK | Aminoglycoside 6-nucleotidyltransferase, putative | | | antibiotic inactivation enzyme | 17407061 |
| Streptococcus dysgalactiae subsp. equisimilis strain NCTC9603 | aadK | Aminoglycoside 6-nucleotidyltransferase, putative | | | antibiotic inactivation enzyme | 17407061 |
| Streptococcus dysgalactiae subsp. equisimilis strain SS1575 | aadK | Aminoglycoside 6-nucleotidyltransferase, putative | | | antibiotic inactivation enzyme | 17407061 |
| Streptococcus dysgalactiae subsp. equisimilis strain T642 | aadK | Aminoglycoside 6-nucleotidyltransferase, putative | | | antibiotic inactivation enzyme | 17407061 |
| Streptococcus dysgalactiae subsp. equisimilis strain UT_4031CC | aadK | Aminoglycoside 6-nucleotidyltransferase, putative | | | antibiotic inactivation enzyme | 17407061 |
| Streptococcus dysgalactiae subsp. equisimilis strain UT_4231_KK | aadK | Aminoglycoside 6-nucleotidyltransferase, putative | | | antibiotic inactivation enzyme | 17407061 |
| Streptococcus dysgalactiae subsp. equisimilis strain UT_4234_DH | aadK | Aminoglycoside 6-nucleotidyltransferase, putative | | | antibiotic inactivation enzyme | 17407061 |
| Streptococcus dysgalactiae subsp. equisimilis strain UT_4241_XS | aadK | Aminoglycoside 6-nucleotidyltransferase, putative | | | antibiotic inactivation enzyme | 17407061 |
| Streptococcus dysgalactiae subsp. equisimilis strain UT_4242_AB | aadK | Aminoglycoside 6-nucleotidyltransferase, putative | | | antibiotic inactivation enzyme | 17407061 |
| Streptococcus dysgalactiae subsp. equisimilis strain UT_4255RC | aadK | Aminoglycoside 6-nucleotidyltransferase, putative | | | antibiotic inactivation enzyme | 17407061 |
| Streptococcus dysgalactiae subsp. equisimilis strain UT_4277_BB | aadK | Aminoglycoside 6-nucleotidyltransferase, putative | | | antibiotic inactivation enzyme | 17407061 |
| Streptococcus dysgalactiae subsp. equisimilis strain UT_4966_RC | aadK | Aminoglycoside 6-nucleotidyltransferase, putative | | | antibiotic inactivation enzyme | 17407061 |
| Streptococcus dysgalactiae subsp. equisimilis strain UT-5345 | aadK | Aminoglycoside 6-nucleotidyltransferase, putative | | | antibiotic inactivation enzyme | 17407061 |
| Streptococcus dysgalactiae subsp. equisimilis strain UT-5354 | aadK | Aminoglycoside 6-nucleotidyltransferase, putative | | | antibiotic inactivation enzyme | 17407061 |
| Streptococcus dysgalactiae subsp. equisimilis strain UT-SS1069 | aadK | Aminoglycoside 6-nucleotidyltransferase, putative | | | antibiotic inactivation enzyme | 17407061 |
| Streptococcus dysgalactiae subsp. equisimilis strain UT-SS957 | aadK | Aminoglycoside 6-nucleotidyltransferase, putative | | | antibiotic inactivation enzyme | 17407061 |
| Streptococcus dysgalactiae subsp. equisimilis strain WCHSDSE-1 | aadK | Aminoglycoside 6-nucleotidyltransferase, putative | | | antibiotic inactivation enzyme | 17407061 |
| Streptococcus dysgalactiae subsp. equisimilis strain NCTC11565 | aadK | Aminoglycoside 6-phosphotransferase, putative | | | antibiotic inactivation enzyme | 17407061 |
| Streptococcus dysgalactiae subsp. equisimilis SD SCDR1 | Alr | Alanine racemase (EC 5.1.1.1) | | | antibiotic target in susceptible species | 19748470;24303782 |
| Streptococcus dysgalactiae subsp. equisimilis 167 | alr | Alanine racemase (EC 5.1.1.1) | | | antibiotic target in susceptible species | 19748470;24303782 |
| Streptococcus dysgalactiae subsp. equisimilis AC-2713 | alr | Alanine racemase (EC 5.1.1.1) | | | antibiotic target in susceptible species | 19748470;24303782 |
| Streptococcus dysgalactiae subsp. equisimilis AKSDE4288 | alr | Alanine racemase (EC 5.1.1.1) | | | antibiotic target in susceptible species | 19748470;24303782 |
| Streptococcus dysgalactiae subsp. equisimilis ATCC 12394 | alr | Alanine racemase (EC 5.1.1.1) | | | antibiotic target in susceptible species | 19748470;24303782 |
| Streptococcus dysgalactiae subsp. equisimilis GGS_124 | acpS | Alanine racemase (EC 5.1.1.1) | | | antibiotic target in susceptible species | 19748470;24303782 |
| Streptococcus dysgalactiae subsp. equisimilis RE378 | alr | Alanine racemase (EC 5.1.1.1) | | | antibiotic target in susceptible species | 19748470;24303782 |
| Streptococcus dysgalactiae subsp. equisimilis SK1249 | alr | Alanine racemase (EC 5.1.1.1) | | | antibiotic target in susceptible species | 19748470;24303782 |
| Streptococcus dysgalactiae subsp. equisimilis SK1250 | alr | Alanine racemase (EC 5.1.1.1) | | | antibiotic target in susceptible species | 19748470;24303782 |
| Streptococcus dysgalactiae subsp. equisimilis strain ASDSE_96 | alr | Alanine racemase (EC 5.1.1.1) | | | antibiotic target in susceptible species | 19748470;24303782 |
| Streptococcus dysgalactiae subsp. equisimilis strain ASDSE_99 | alr | Alanine racemase (EC 5.1.1.1) | | | antibiotic target in susceptible species | 19748470;24303782 |
| Streptococcus dysgalactiae subsp. equisimilis strain C161L1 | alr | Alanine racemase (EC 5.1.1.1) | | | antibiotic target in susceptible species | 19748470;24303782 |
| Streptococcus dysgalactiae subsp. equisimilis strain KNZ01 | Alr | Alanine racemase (EC 5.1.1.1) | | | antibiotic target in susceptible species | 19748470;24303782 |
| Streptococcus dysgalactiae subsp. equisimilis strain KNZ03 | Alr | Alanine racemase (EC 5.1.1.1) | | | antibiotic target in susceptible species | 19748470;24303782 |
| Streptococcus dysgalactiae subsp. equisimilis strain KNZ04 | Alr | Alanine racemase (EC 5.1.1.1) | | | antibiotic target in susceptible species | 19748470;24303782 |
| Streptococcus dysgalactiae subsp. equisimilis strain KNZ06 | Alr | Alanine racemase (EC 5.1.1.1) | | | antibiotic target in susceptible species | 19748470;24303782 |
| Streptococcus dysgalactiae subsp. equisimilis strain KNZ07 | Alr | Alanine racemase (EC 5.1.1.1) | | | antibiotic target in susceptible species | 19748470;24303782 |
| Streptococcus dysgalactiae subsp. equisimilis strain KNZ10 | Alr | Alanine racemase (EC 5.1.1.1) | | | antibiotic target in susceptible species | 19748470;24303782 |
| Streptococcus dysgalactiae subsp. equisimilis strain KNZ12 | Alr | Alanine racemase (EC 5.1.1.1) | | | antibiotic target in susceptible species | 19748470;24303782 |
| Streptococcus dysgalactiae subsp. equisimilis strain KNZ15 | Alr | Alanine racemase (EC 5.1.1.1) | | | antibiotic target in susceptible species | 19748470;24303782 |
| Streptococcus dysgalactiae subsp. equisimilis strain KNZ16 | Alr | Alanine racemase (EC 5.1.1.1) | | | antibiotic target in susceptible species | 19748470;24303782 |
| Streptococcus dysgalactiae subsp. equisimilis strain NCTC10321 | Alr | Alanine racemase (EC 5.1.1.1) | | | antibiotic target in susceptible species | 19748470;24303782 |
| Streptococcus dysgalactiae subsp. equisimilis strain NCTC11554 | Alr | Alanine racemase (EC 5.1.1.1) | | | antibiotic target in susceptible species | 19748470;24303782 |
| Streptococcus dysgalactiae subsp. equisimilis strain NCTC11555 | Alr | Alanine racemase (EC 5.1.1.1) | | | antibiotic target in susceptible species | 19748470;24303782 |
| Streptococcus dysgalactiae subsp. equisimilis strain NCTC11556 | Alr | Alanine racemase (EC 5.1.1.1) | | | antibiotic target in susceptible species | 19748470;24303782 |
| Streptococcus dysgalactiae subsp. equisimilis strain NCTC11557 | Alr | Alanine racemase (EC 5.1.1.1) | | | antibiotic target in susceptible species | 19748470;24303782 |
| Streptococcus dysgalactiae subsp. equisimilis strain NCTC11564 | Alr | Alanine racemase (EC 5.1.1.1) | | | antibiotic target in susceptible species | 19748470;24303782 |
| Streptococcus dysgalactiae subsp. equisimilis strain NCTC11565 | Alr | Alanine racemase (EC 5.1.1.1) | | | antibiotic target in susceptible species | 19748470;24303782 |
| Streptococcus dysgalactiae subsp. equisimilis strain NCTC5370 | alr | Alanine racemase (EC 5.1.1.1) | | | antibiotic target in susceptible species | 19748470;24303782 |
| Streptococcus dysgalactiae subsp. equisimilis strain NCTC5371 | alr | Alanine racemase (EC 5.1.1.1) | | | antibiotic target in susceptible species | 19748470;24303782 |
| Streptococcus dysgalactiae subsp. equisimilis strain NCTC5969 | Alr | Alanine racemase (EC 5.1.1.1) | | | antibiotic target in susceptible species | 19748470;24303782 |
| Streptococcus dysgalactiae subsp. equisimilis strain NCTC6179 | alr | Alanine racemase (EC 5.1.1.1) | | | antibiotic target in susceptible species | 19748470;24303782 |
| Streptococcus dysgalactiae subsp. equisimilis strain NCTC6181 | Alr | Alanine racemase (EC 5.1.1.1) | | | antibiotic target in susceptible species | 19748470;24303782 |
| Streptococcus dysgalactiae subsp. equisimilis strain NCTC6407 | Alr | Alanine racemase (EC 5.1.1.1) | | | antibiotic target in susceptible species | 19748470;24303782 |
| Streptococcus dysgalactiae subsp. equisimilis strain NCTC7136 | alr | Alanine racemase (EC 5.1.1.1) | | | antibiotic target in susceptible species | 19748470;24303782 |
| Streptococcus dysgalactiae subsp. equisimilis strain NCTC8543 | Alr | Alanine racemase (EC 5.1.1.1) | | | antibiotic target in susceptible species | 19748470;24303782 |
| Streptococcus dysgalactiae subsp. equisimilis strain NCTC8546 | Alr | Alanine racemase (EC 5.1.1.1) | | | antibiotic target in susceptible species | 19748470;24303782 |
| Streptococcus dysgalactiae subsp. equisimilis strain NCTC9413 | Alr | Alanine racemase (EC 5.1.1.1) | | | antibiotic target in susceptible species | 19748470;24303782 |
| Streptococcus dysgalactiae subsp. equisimilis strain NCTC9414 | alr | Alanine racemase (EC 5.1.1.1) | | | antibiotic target in susceptible species | 19748470;24303782 |
| Streptococcus dysgalactiae subsp. equisimilis strain NCTC9603 | Alr | Alanine racemase (EC 5.1.1.1) | | | antibiotic target in susceptible species | 19748470;24303782 |
| Streptococcus dysgalactiae subsp. equisimilis strain SS1575 | alr | Alanine racemase (EC 5.1.1.1) | | | antibiotic target in susceptible species | 19748470;24303782 |
| Streptococcus dysgalactiae subsp. equisimilis strain T642 | alr | Alanine racemase (EC 5.1.1.1) | | | antibiotic target in susceptible species | 19748470;24303782 |
| Streptococcus dysgalactiae subsp. equisimilis strain UT_4031CC | alr | Alanine racemase (EC 5.1.1.1) | | | antibiotic target in susceptible species | 19748470;24303782 |
| Streptococcus dysgalactiae subsp. equisimilis strain UT_4231_KK | alr | Alanine racemase (EC 5.1.1.1) | | | antibiotic target in susceptible species | 19748470;24303782 |
| Streptococcus dysgalactiae subsp. equisimilis strain UT_4234_DH | alr | Alanine racemase (EC 5.1.1.1) | | | antibiotic target in susceptible species | 19748470;24303782 |
| Streptococcus dysgalactiae subsp. equisimilis strain UT_4241_XS | alr | Alanine racemase (EC 5.1.1.1) | | | antibiotic target in susceptible species | 19748470;24303782 |
| Streptococcus dysgalactiae subsp. equisimilis strain UT_4242_AB | alr | Alanine racemase (EC 5.1.1.1) | | | antibiotic target in susceptible species | 19748470;24303782 |
| Streptococcus dysgalactiae subsp. equisimilis strain UT_4255RC | alr | Alanine racemase (EC 5.1.1.1) | | | antibiotic target in susceptible species | 19748470;24303782 |
| Streptococcus dysgalactiae subsp. equisimilis strain UT_4277_BB | alr | Alanine racemase (EC 5.1.1.1) | | | antibiotic target in susceptible species | 19748470;24303782 |
| Streptococcus dysgalactiae subsp. equisimilis strain UT_4966_RC | alr | Alanine racemase (EC 5.1.1.1) | | | antibiotic target in susceptible species | 19748470;24303782 |
| Streptococcus dysgalactiae subsp. equisimilis strain UT-5345 | alr | Alanine racemase (EC 5.1.1.1) | | | antibiotic target in susceptible species | 19748470;24303782 |
| Streptococcus dysgalactiae subsp. equisimilis strain UT-5354 | alr | Alanine racemase (EC 5.1.1.1) | | | antibiotic target in susceptible species | 19748470;24303782 |
| Streptococcus dysgalactiae subsp. equisimilis strain UT-SS1069 | alr | Alanine racemase (EC 5.1.1.1) | | | antibiotic target in susceptible species | 19748470;24303782 |
| Streptococcus dysgalactiae subsp. equisimilis strain UT-SS957 | alr | Alanine racemase (EC 5.1.1.1) | | | antibiotic target in susceptible species | 19748470;24303782 |
| Streptococcus dysgalactiae subsp. equisimilis strain WCHSDSE-1 | alr | Alanine racemase (EC 5.1.1.1) | | | antibiotic target in susceptible species | 19748470;24303782 |
| Streptococcus dysgalactiae subsp. equisimilis SD SCDR1 | kasA | 3-oxoacyl-[acyl-carrier-protein] synthase, KASII (EC 2.3.1.179) | | | antibiotic target in susceptible species | 10428945 |
| Streptococcus dysgalactiae subsp. equisimilis 167 | fabF | 3-oxoacyl-[acyl-carrier-protein] synthase, KASII (EC 2.3.1.179) | | | antibiotic target in susceptible species | 10428945 |
| Streptococcus dysgalactiae subsp. equisimilis AC-2713 | fabF | 3-oxoacyl-[acyl-carrier-protein] synthase, KASII (EC 2.3.1.179) | | | antibiotic target in susceptible species | 10428945 |
| Streptococcus dysgalactiae subsp. equisimilis AKSDE4288 |  | 3-oxoacyl-[acyl-carrier-protein] synthase, KASII (EC 2.3.1.179) | | | antibiotic target in susceptible species | 10428945 |
| Streptococcus dysgalactiae subsp. equisimilis ATCC 12394 |  | 3-oxoacyl-[acyl-carrier-protein] synthase, KASII (EC 2.3.1.179) | | | antibiotic target in susceptible species | 10428945 |
| Streptococcus dysgalactiae subsp. equisimilis GGS_124 | fabF | 3-oxoacyl-[acyl-carrier-protein] synthase, KASII (EC 2.3.1.179) | | | antibiotic target in susceptible species | 10428945 |
| Streptococcus dysgalactiae subsp. equisimilis RE378 | fabF | 3-oxoacyl-[acyl-carrier-protein] synthase, KASII (EC 2.3.1.179) | | | antibiotic target in susceptible species | 10428945 |
| Streptococcus dysgalactiae subsp. equisimilis SK1249 | fabF | 3-oxoacyl-[acyl-carrier-protein] synthase, KASII (EC 2.3.1.179) | | | antibiotic target in susceptible species | 10428945 |
| Streptococcus dysgalactiae subsp. equisimilis SK1250 | fabF | 3-oxoacyl-[acyl-carrier-protein] synthase, KASII (EC 2.3.1.179) | | | antibiotic target in susceptible species | 10428945 |
| Streptococcus dysgalactiae subsp. equisimilis strain ASDSE_96 |  | 3-oxoacyl-[acyl-carrier-protein] synthase, KASII (EC 2.3.1.179) | | | antibiotic target in susceptible species | 10428945 |
| Streptococcus dysgalactiae subsp. equisimilis strain ASDSE_99 |  | 3-oxoacyl-[acyl-carrier-protein] synthase, KASII (EC 2.3.1.179) | | | antibiotic target in susceptible species | 10428945 |
| Streptococcus dysgalactiae subsp. equisimilis strain C161L1 |  | 3-oxoacyl-[acyl-carrier-protein] synthase, KASII (EC 2.3.1.179) | | | antibiotic target in susceptible species | 10428945 |
| Streptococcus dysgalactiae subsp. equisimilis strain KNZ01 | kasA | 3-oxoacyl-[acyl-carrier-protein] synthase, KASII (EC 2.3.1.179) | | | antibiotic target in susceptible species | 10428945 |
| Streptococcus dysgalactiae subsp. equisimilis strain KNZ03 | kasA | 3-oxoacyl-[acyl-carrier-protein] synthase, KASII (EC 2.3.1.179) | | | antibiotic target in susceptible species | 10428945 |
| Streptococcus dysgalactiae subsp. equisimilis strain KNZ04 | kasA | 3-oxoacyl-[acyl-carrier-protein] synthase, KASII (EC 2.3.1.179) | | | antibiotic target in susceptible species | 10428945 |
| Streptococcus dysgalactiae subsp. equisimilis strain KNZ06 | kasA | 3-oxoacyl-[acyl-carrier-protein] synthase, KASII (EC 2.3.1.179) | | | antibiotic target in susceptible species | 10428945 |
| Streptococcus dysgalactiae subsp. equisimilis strain KNZ07 | kasA | 3-oxoacyl-[acyl-carrier-protein] synthase, KASII (EC 2.3.1.179) | | | antibiotic target in susceptible species | 10428945 |
| Streptococcus dysgalactiae subsp. equisimilis strain KNZ10 | kasA | 3-oxoacyl-[acyl-carrier-protein] synthase, KASII (EC 2.3.1.179) | | | antibiotic target in susceptible species | 10428945 |
| Streptococcus dysgalactiae subsp. equisimilis strain KNZ12 | kasA | 3-oxoacyl-[acyl-carrier-protein] synthase, KASII (EC 2.3.1.179) | | | antibiotic target in susceptible species | 10428945 |
| Streptococcus dysgalactiae subsp. equisimilis strain KNZ15 | kasA | 3-oxoacyl-[acyl-carrier-protein] synthase, KASII (EC 2.3.1.179) | | | antibiotic target in susceptible species | 10428945 |
| Streptococcus dysgalactiae subsp. equisimilis strain KNZ16 | kasA | 3-oxoacyl-[acyl-carrier-protein] synthase, KASII (EC 2.3.1.179) | | | antibiotic target in susceptible species | 10428945 |
| Streptococcus dysgalactiae subsp. equisimilis strain NCTC10321 | kasA | 3-oxoacyl-[acyl-carrier-protein] synthase, KASII (EC 2.3.1.179) | | | antibiotic target in susceptible species | 10428945 |
| Streptococcus dysgalactiae subsp. equisimilis strain NCTC11554 | kasA | 3-oxoacyl-[acyl-carrier-protein] synthase, KASII (EC 2.3.1.179) | | | antibiotic target in susceptible species | 10428945 |
| Streptococcus dysgalactiae subsp. equisimilis strain NCTC11555 | kasA | 3-oxoacyl-[acyl-carrier-protein] synthase, KASII (EC 2.3.1.179) | | | antibiotic target in susceptible species | 10428945 |
| Streptococcus dysgalactiae subsp. equisimilis strain NCTC11556 | kasA | 3-oxoacyl-[acyl-carrier-protein] synthase, KASII (EC 2.3.1.179) | | | antibiotic target in susceptible species | 10428945 |
| Streptococcus dysgalactiae subsp. equisimilis strain NCTC11557 | kasA | 3-oxoacyl-[acyl-carrier-protein] synthase, KASII (EC 2.3.1.179) | | | antibiotic target in susceptible species | 10428945 |
| Streptococcus dysgalactiae subsp. equisimilis strain NCTC11564 | kasA | 3-oxoacyl-[acyl-carrier-protein] synthase, KASII (EC 2.3.1.179) | | | antibiotic target in susceptible species | 10428945 |
| Streptococcus dysgalactiae subsp. equisimilis strain NCTC11565 | kasA | 3-oxoacyl-[acyl-carrier-protein] synthase, KASII (EC 2.3.1.179) | | | antibiotic target in susceptible species | 10428945 |
| Streptococcus dysgalactiae subsp. equisimilis strain NCTC5370 | fabF | 3-oxoacyl-[acyl-carrier-protein] synthase, KASII (EC 2.3.1.179) | | | antibiotic target in susceptible species | 10428945 |
| Streptococcus dysgalactiae subsp. equisimilis strain NCTC5371 | fabF | 3-oxoacyl-[acyl-carrier-protein] synthase, KASII (EC 2.3.1.179) | | | antibiotic target in susceptible species | 10428945 |
| Streptococcus dysgalactiae subsp. equisimilis strain NCTC5969 | kasA | 3-oxoacyl-[acyl-carrier-protein] synthase, KASII (EC 2.3.1.179) | | | antibiotic target in susceptible species | 10428945 |
| Streptococcus dysgalactiae subsp. equisimilis strain NCTC6179 | fabF | 3-oxoacyl-[acyl-carrier-protein] synthase, KASII (EC 2.3.1.179) | | | antibiotic target in susceptible species | 10428945 |
| Streptococcus dysgalactiae subsp. equisimilis strain NCTC6181 | kasA | 3-oxoacyl-[acyl-carrier-protein] synthase, KASII (EC 2.3.1.179) | | | antibiotic target in susceptible species | 10428945 |
| Streptococcus dysgalactiae subsp. equisimilis strain NCTC6407 | kasA | 3-oxoacyl-[acyl-carrier-protein] synthase, KASII (EC 2.3.1.179) | | | antibiotic target in susceptible species | 10428945 |
| Streptococcus dysgalactiae subsp. equisimilis strain NCTC7136 | fabF | 3-oxoacyl-[acyl-carrier-protein] synthase, KASII (EC 2.3.1.179) | | | antibiotic target in susceptible species | 10428945 |
| Streptococcus dysgalactiae subsp. equisimilis strain NCTC8543 | kasA | 3-oxoacyl-[acyl-carrier-protein] synthase, KASII (EC 2.3.1.179) | | | antibiotic target in susceptible species | 10428945 |
| Streptococcus dysgalactiae subsp. equisimilis strain NCTC8546 | kasA | 3-oxoacyl-[acyl-carrier-protein] synthase, KASII (EC 2.3.1.179) | | | antibiotic target in susceptible species | 10428945 |
| Streptococcus dysgalactiae subsp. equisimilis strain NCTC9413 | kasA | 3-oxoacyl-[acyl-carrier-protein] synthase, KASII (EC 2.3.1.179) | | | antibiotic target in susceptible species | 10428945 |
| Streptococcus dysgalactiae subsp. equisimilis strain NCTC9414 | fabF | 3-oxoacyl-[acyl-carrier-protein] synthase, KASII (EC 2.3.1.179) | | | antibiotic target in susceptible species | 10428945 |
| Streptococcus dysgalactiae subsp. equisimilis strain NCTC9603 | kasA | 3-oxoacyl-[acyl-carrier-protein] synthase, KASII (EC 2.3.1.179) | | | antibiotic target in susceptible species | 10428945 |
| Streptococcus dysgalactiae subsp. equisimilis strain SS1575 | fabF | 3-oxoacyl-[acyl-carrier-protein] synthase, KASII (EC 2.3.1.179) | | | antibiotic target in susceptible species | 10428945 |
| Streptococcus dysgalactiae subsp. equisimilis strain T642 | kasA | 3-oxoacyl-[acyl-carrier-protein] synthase, KASII (EC 2.3.1.179) | | | antibiotic target in susceptible species | 10428945 |
| Streptococcus dysgalactiae subsp. equisimilis strain UT_4031CC | kasA | 3-oxoacyl-[acyl-carrier-protein] synthase, KASII (EC 2.3.1.179) | | | antibiotic target in susceptible species | 10428945 |
| Streptococcus dysgalactiae subsp. equisimilis strain UT_4231_KK | kasA | 3-oxoacyl-[acyl-carrier-protein] synthase, KASII (EC 2.3.1.179) | | | antibiotic target in susceptible species | 10428945 |
| Streptococcus dysgalactiae subsp. equisimilis strain UT_4234_DH | kasA | 3-oxoacyl-[acyl-carrier-protein] synthase, KASII (EC 2.3.1.179) | | | antibiotic target in susceptible species | 10428945 |
| Streptococcus dysgalactiae subsp. equisimilis strain UT_4241_XS | kasA | 3-oxoacyl-[acyl-carrier-protein] synthase, KASII (EC 2.3.1.179) | | | antibiotic target in susceptible species | 10428945 |
| Streptococcus dysgalactiae subsp. equisimilis strain UT_4242_AB | kasA | 3-oxoacyl-[acyl-carrier-protein] synthase, KASII (EC 2.3.1.179) | | | antibiotic target in susceptible species | 10428945 |
| Streptococcus dysgalactiae subsp. equisimilis strain UT_4255RC | kasA | 3-oxoacyl-[acyl-carrier-protein] synthase, KASII (EC 2.3.1.179) | | | antibiotic target in susceptible species | 10428945 |
| Streptococcus dysgalactiae subsp. equisimilis strain UT_4277_BB | kasA | 3-oxoacyl-[acyl-carrier-protein] synthase, KASII (EC 2.3.1.179) | | | antibiotic target in susceptible species | 10428945 |
| Streptococcus dysgalactiae subsp. equisimilis strain UT_4966_RC | kasA | 3-oxoacyl-[acyl-carrier-protein] synthase, KASII (EC 2.3.1.179) | | | antibiotic target in susceptible species | 10428945 |
| Streptococcus dysgalactiae subsp. equisimilis strain UT-5345 | kasA | 3-oxoacyl-[acyl-carrier-protein] synthase, KASII (EC 2.3.1.179) | | | antibiotic target in susceptible species | 10428945 |
| Streptococcus dysgalactiae subsp. equisimilis strain UT-5354 | kasA | 3-oxoacyl-[acyl-carrier-protein] synthase, KASII (EC 2.3.1.179) | | | antibiotic target in susceptible species | 10428945 |
| Streptococcus dysgalactiae subsp. equisimilis strain UT-SS1069 | kasA | 3-oxoacyl-[acyl-carrier-protein] synthase, KASII (EC 2.3.1.179) | | | antibiotic target in susceptible species | 10428945 |
| Streptococcus dysgalactiae subsp. equisimilis strain UT-SS957 | kasA | 3-oxoacyl-[acyl-carrier-protein] synthase, KASII (EC 2.3.1.179) | | | antibiotic target in susceptible species | 10428945 |
| Streptococcus dysgalactiae subsp. equisimilis strain WCHSDSE-1 | kasA | 3-oxoacyl-[acyl-carrier-protein] synthase, KASII (EC 2.3.1.179) | | | antibiotic target in susceptible species | 10428945 |
| Streptococcus dysgalactiae subsp. equisimilis SD SCDR1 | gidB | 16S rRNA (guanine(527)-N(7))-methyltransferase (EC 2.1.1.170) | | | gene conferring resistance via absence | 17238915 |
| Streptococcus dysgalactiae subsp. equisimilis 167 | gidB | 16S rRNA (guanine(527)-N(7))-methyltransferase (EC 2.1.1.170) | | | gene conferring resistance via absence | 17238915 |
| Streptococcus dysgalactiae subsp. equisimilis AC-2713 | gidB | 16S rRNA (guanine(527)-N(7))-methyltransferase (EC 2.1.1.170) | | | gene conferring resistance via absence | 17238915 |
| Streptococcus dysgalactiae subsp. equisimilis AKSDE4288 |  | 16S rRNA (guanine(527)-N(7))-methyltransferase (EC 2.1.1.170) | | | gene conferring resistance via absence | 17238915 |
| Streptococcus dysgalactiae subsp. equisimilis ATCC 12394 | gidB | 16S rRNA (guanine(527)-N(7))-methyltransferase (EC 2.1.1.170) | | | gene conferring resistance via absence | 17238915 |
| Streptococcus dysgalactiae subsp. equisimilis GGS_124 | gidB | 16S rRNA (guanine(527)-N(7))-methyltransferase (EC 2.1.1.170) | | | gene conferring resistance via absence | 17238915 |
| Streptococcus dysgalactiae subsp. equisimilis RE378 | gidB | 16S rRNA (guanine(527)-N(7))-methyltransferase (EC 2.1.1.170) | | | gene conferring resistance via absence | 17238915 |
| Streptococcus dysgalactiae subsp. equisimilis SK1249 | rmsG | 16S rRNA (guanine(527)-N(7))-methyltransferase (EC 2.1.1.170) | | | gene conferring resistance via absence | 17238915 |
| Streptococcus dysgalactiae subsp. equisimilis SK1250 | gidB | 16S rRNA (guanine(527)-N(7))-methyltransferase (EC 2.1.1.170) | | | gene conferring resistance via absence | 17238915 |
| Streptococcus dysgalactiae subsp. equisimilis strain ASDSE_96 |  | 16S rRNA (guanine(527)-N(7))-methyltransferase (EC 2.1.1.170) | | | gene conferring resistance via absence | 17238915 |
| Streptococcus dysgalactiae subsp. equisimilis strain ASDSE_99 |  | 16S rRNA (guanine(527)-N(7))-methyltransferase (EC 2.1.1.170) | | | gene conferring resistance via absence | 17238915 |
| Streptococcus dysgalactiae subsp. equisimilis strain C161L1 |  | 16S rRNA (guanine(527)-N(7))-methyltransferase (EC 2.1.1.170) | | | gene conferring resistance via absence | 17238915 |
| Streptococcus dysgalactiae subsp. equisimilis strain KNZ01 | gidB | 16S rRNA (guanine(527)-N(7))-methyltransferase (EC 2.1.1.170) | | | gene conferring resistance via absence | 17238915 |
| Streptococcus dysgalactiae subsp. equisimilis strain KNZ03 | gidB | 16S rRNA (guanine(527)-N(7))-methyltransferase (EC 2.1.1.170) | | | gene conferring resistance via absence | 17238915 |
| Streptococcus dysgalactiae subsp. equisimilis strain KNZ04 | gidB | 16S rRNA (guanine(527)-N(7))-methyltransferase (EC 2.1.1.170) | | | gene conferring resistance via absence | 17238915 |
| Streptococcus dysgalactiae subsp. equisimilis strain KNZ06 | gidB | 16S rRNA (guanine(527)-N(7))-methyltransferase (EC 2.1.1.170) | | | gene conferring resistance via absence | 17238915 |
| Streptococcus dysgalactiae subsp. equisimilis strain KNZ07 | gidB | 16S rRNA (guanine(527)-N(7))-methyltransferase (EC 2.1.1.170) | | | gene conferring resistance via absence | 17238915 |
| Streptococcus dysgalactiae subsp. equisimilis strain KNZ10 | gidB | 16S rRNA (guanine(527)-N(7))-methyltransferase (EC 2.1.1.170) | | | gene conferring resistance via absence | 17238915 |
| Streptococcus dysgalactiae subsp. equisimilis strain KNZ12 | gidB | 16S rRNA (guanine(527)-N(7))-methyltransferase (EC 2.1.1.170) | | | gene conferring resistance via absence | 17238915 |
| Streptococcus dysgalactiae subsp. equisimilis strain KNZ15 | gidB | 16S rRNA (guanine(527)-N(7))-methyltransferase (EC 2.1.1.170) | | | gene conferring resistance via absence | 17238915 |
| Streptococcus dysgalactiae subsp. equisimilis strain KNZ16 | gidB | 16S rRNA (guanine(527)-N(7))-methyltransferase (EC 2.1.1.170) | | | gene conferring resistance via absence | 17238915 |
| Streptococcus dysgalactiae subsp. equisimilis strain NCTC10321 | gidB | 16S rRNA (guanine(527)-N(7))-methyltransferase (EC 2.1.1.170) | | | gene conferring resistance via absence | 17238915 |
| Streptococcus dysgalactiae subsp. equisimilis strain NCTC11554 | gidB | 16S rRNA (guanine(527)-N(7))-methyltransferase (EC 2.1.1.170) | | | gene conferring resistance via absence | 17238915 |
| Streptococcus dysgalactiae subsp. equisimilis strain NCTC11555 | gidB | 16S rRNA (guanine(527)-N(7))-methyltransferase (EC 2.1.1.170) | | | gene conferring resistance via absence | 17238915 |
| Streptococcus dysgalactiae subsp. equisimilis strain NCTC11556 | gidB | 16S rRNA (guanine(527)-N(7))-methyltransferase (EC 2.1.1.170) | | | gene conferring resistance via absence | 17238915 |
| Streptococcus dysgalactiae subsp. equisimilis strain NCTC11557 | gidB | 16S rRNA (guanine(527)-N(7))-methyltransferase (EC 2.1.1.170) | | | gene conferring resistance via absence | 17238915 |
| Streptococcus dysgalactiae subsp. equisimilis strain NCTC11564 | gidB | 16S rRNA (guanine(527)-N(7))-methyltransferase (EC 2.1.1.170) | | | gene conferring resistance via absence | 17238915 |
| Streptococcus dysgalactiae subsp. equisimilis strain NCTC11565 | gidB | 16S rRNA (guanine(527)-N(7))-methyltransferase (EC 2.1.1.170) | | | gene conferring resistance via absence | 17238915 |
| Streptococcus dysgalactiae subsp. equisimilis strain NCTC5370 | gidB | 16S rRNA (guanine(527)-N(7))-methyltransferase (EC 2.1.1.170) | | | gene conferring resistance via absence | 17238915 |
| Streptococcus dysgalactiae subsp. equisimilis strain NCTC5371 | gidB | 16S rRNA (guanine(527)-N(7))-methyltransferase (EC 2.1.1.170) | | | gene conferring resistance via absence | 17238915 |
| Streptococcus dysgalactiae subsp. equisimilis strain NCTC5969 | gidB | 16S rRNA (guanine(527)-N(7))-methyltransferase (EC 2.1.1.170) | | | gene conferring resistance via absence | 17238915 |
| Streptococcus dysgalactiae subsp. equisimilis strain NCTC6179 | gidB | 16S rRNA (guanine(527)-N(7))-methyltransferase (EC 2.1.1.170) | | | gene conferring resistance via absence | 17238915 |
| Streptococcus dysgalactiae subsp. equisimilis strain NCTC6181 | gidB | 16S rRNA (guanine(527)-N(7))-methyltransferase (EC 2.1.1.170) | | | gene conferring resistance via absence | 17238915 |
| Streptococcus dysgalactiae subsp. equisimilis strain NCTC6407 | gidB | 16S rRNA (guanine(527)-N(7))-methyltransferase (EC 2.1.1.170) | | | gene conferring resistance via absence | 17238915 |
| Streptococcus dysgalactiae subsp. equisimilis strain NCTC7136 | gidB | 16S rRNA (guanine(527)-N(7))-methyltransferase (EC 2.1.1.170) | | | gene conferring resistance via absence | 17238915 |
| Streptococcus dysgalactiae subsp. equisimilis strain NCTC8543 | gidB | 16S rRNA (guanine(527)-N(7))-methyltransferase (EC 2.1.1.170) | | | gene conferring resistance via absence | 17238915 |
| Streptococcus dysgalactiae subsp. equisimilis strain NCTC8546 | gidB | 16S rRNA (guanine(527)-N(7))-methyltransferase (EC 2.1.1.170) | | | gene conferring resistance via absence | 17238915 |
| Streptococcus dysgalactiae subsp. equisimilis strain NCTC9413 | gidB | 16S rRNA (guanine(527)-N(7))-methyltransferase (EC 2.1.1.170) | | | gene conferring resistance via absence | 17238915 |
| Streptococcus dysgalactiae subsp. equisimilis strain NCTC9414 | gidB | 16S rRNA (guanine(527)-N(7))-methyltransferase (EC 2.1.1.170) | | | gene conferring resistance via absence | 17238915 |
| Streptococcus dysgalactiae subsp. equisimilis strain NCTC9603 | gidB | 16S rRNA (guanine(527)-N(7))-methyltransferase (EC 2.1.1.170) | | | gene conferring resistance via absence | 17238915 |
| Streptococcus dysgalactiae subsp. equisimilis strain SS1575 | gidB | 16S rRNA (guanine(527)-N(7))-methyltransferase (EC 2.1.1.170) | | | gene conferring resistance via absence | 17238915 |
| Streptococcus dysgalactiae subsp. equisimilis strain T642 | gidB | 16S rRNA (guanine(527)-N(7))-methyltransferase (EC 2.1.1.170) | | | gene conferring resistance via absence | 17238915 |
| Streptococcus dysgalactiae subsp. equisimilis strain UT_4031CC | gidB | 16S rRNA (guanine(527)-N(7))-methyltransferase (EC 2.1.1.170) | | | gene conferring resistance via absence | 17238915 |
| Streptococcus dysgalactiae subsp. equisimilis strain UT_4231_KK | gidB | 16S rRNA (guanine(527)-N(7))-methyltransferase (EC 2.1.1.170) | | | gene conferring resistance via absence | 17238915 |
| Streptococcus dysgalactiae subsp. equisimilis strain UT_4234_DH | gidB | 16S rRNA (guanine(527)-N(7))-methyltransferase (EC 2.1.1.170) | | | gene conferring resistance via absence | 17238915 |
| Streptococcus dysgalactiae subsp. equisimilis strain UT_4241_XS | gidB | 16S rRNA (guanine(527)-N(7))-methyltransferase (EC 2.1.1.170) | | | gene conferring resistance via absence | 17238915 |
| Streptococcus dysgalactiae subsp. equisimilis strain UT_4242_AB | gidB | 16S rRNA (guanine(527)-N(7))-methyltransferase (EC 2.1.1.170) | | | gene conferring resistance via absence | 17238915 |
| Streptococcus dysgalactiae subsp. equisimilis strain UT_4255RC | gidB | 16S rRNA (guanine(527)-N(7))-methyltransferase (EC 2.1.1.170) | | | gene conferring resistance via absence | 17238915 |
| Streptococcus dysgalactiae subsp. equisimilis strain UT_4277_BB | gidB | 16S rRNA (guanine(527)-N(7))-methyltransferase (EC 2.1.1.170) | | | gene conferring resistance via absence | 17238915 |
| Streptococcus dysgalactiae subsp. equisimilis strain UT_4966_RC | gidB | 16S rRNA (guanine(527)-N(7))-methyltransferase (EC 2.1.1.170) | | | gene conferring resistance via absence | 17238915 |
| Streptococcus dysgalactiae subsp. equisimilis strain UT-5345 | gidB | 16S rRNA (guanine(527)-N(7))-methyltransferase (EC 2.1.1.170) | | | gene conferring resistance via absence | 17238915 |
| Streptococcus dysgalactiae subsp. equisimilis strain UT-5354 | gidB | 16S rRNA (guanine(527)-N(7))-methyltransferase (EC 2.1.1.170) | | | gene conferring resistance via absence | 17238915 |
| Streptococcus dysgalactiae subsp. equisimilis strain UT-SS1069 | gidB | 16S rRNA (guanine(527)-N(7))-methyltransferase (EC 2.1.1.170) | | | gene conferring resistance via absence | 17238915 |
| Streptococcus dysgalactiae subsp. equisimilis strain UT-SS957 | gidB | 16S rRNA (guanine(527)-N(7))-methyltransferase (EC 2.1.1.170) | | | gene conferring resistance via absence | 17238915 |
| Streptococcus dysgalactiae subsp. equisimilis strain WCHSDSE-1 | gidB | 16S rRNA (guanine(527)-N(7))-methyltransferase (EC 2.1.1.170) | | | gene conferring resistance via absence | 17238915 |
| Streptococcus dysgalactiae subsp. equisimilis SD SCDR1 | EF-G | Translation elongation factor G | | | antibiotic target in susceptible species | 17980694 |
| Streptococcus dysgalactiae subsp. equisimilis 167 | fus | Translation elongation factor G | | | antibiotic target in susceptible species | 17980694;19289529;19289529;17325218;19289529 |
| Streptococcus dysgalactiae subsp. equisimilis AC-2713 | fusA | Translation elongation factor G | | | antibiotic target in susceptible species | 17980694;19289529;19289529;17325218;19289529 |
| Streptococcus dysgalactiae subsp. equisimilis AKSDE4288 | EF-G | Translation elongation factor G | | | antibiotic target in susceptible species | 17980694;19289529;19289529;17325218;19289529 |
| Streptococcus dysgalactiae subsp. equisimilis ATCC 12394 | EF-G | Translation elongation factor G | | | antibiotic target in susceptible species | 17980694;19289529;19289529;17325218;19289529 |
| Streptococcus dysgalactiae subsp. equisimilis GGS_124 | fus | Translation elongation factor G | | | antibiotic target in susceptible species | 17980694;19289529;19289529;17325218;19289529 |
| Streptococcus dysgalactiae subsp. equisimilis RE378 | fus | Translation elongation factor G | | | antibiotic target in susceptible species | 17980694;19289529;19289529;17325218;19289529 |
| Streptococcus dysgalactiae subsp. equisimilis SK1249 | fusA | Translation elongation factor G | | | antibiotic target in susceptible species | 17980694;19289529;19289529;17325218;19289529 |
| Streptococcus dysgalactiae subsp. equisimilis SK1250 | fusA | Translation elongation factor G | | | antibiotic target in susceptible species | 17980694;19289529;19289529;17325218;19289529 |
| Streptococcus dysgalactiae subsp. equisimilis strain ASDSE_96 | EF-G | Translation elongation factor G | | | antibiotic target in susceptible species | 17980694;19289529;19289529;17325218;19289529 |
| Streptococcus dysgalactiae subsp. equisimilis strain ASDSE_99 | EF-G | Translation elongation factor G | | | antibiotic target in susceptible species | 17980694;19289529;19289529;17325218;19289529 |
| Streptococcus dysgalactiae subsp. equisimilis strain C161L1 | EF-G | Translation elongation factor G | | | antibiotic target in susceptible species | 17980694;19289529;19289529;17325218;19289529 |
| Streptococcus dysgalactiae subsp. equisimilis strain KNZ01 | EF-G | Translation elongation factor G | | | antibiotic target in susceptible species | 17980694 |
| Streptococcus dysgalactiae subsp. equisimilis strain KNZ03 | EF-G | Translation elongation factor G | | | antibiotic target in susceptible species | 17980694 |
| Streptococcus dysgalactiae subsp. equisimilis strain KNZ04 | EF-G | Translation elongation factor G | | | antibiotic target in susceptible species | 17980694 |
| Streptococcus dysgalactiae subsp. equisimilis strain KNZ06 | EF-G | Translation elongation factor G | | | antibiotic target in susceptible species | 17980694 |
| Streptococcus dysgalactiae subsp. equisimilis strain KNZ07 | EF-G | Translation elongation factor G | | | antibiotic target in susceptible species | 17980694 |
| Streptococcus dysgalactiae subsp. equisimilis strain KNZ10 | EF-G | Translation elongation factor G | | | antibiotic target in susceptible species | 17980694 |
| Streptococcus dysgalactiae subsp. equisimilis strain KNZ12 | EF-G | Translation elongation factor G | | | antibiotic target in susceptible species | 17980694 |
| Streptococcus dysgalactiae subsp. equisimilis strain KNZ15 | EF-G | Translation elongation factor G | | | antibiotic target in susceptible species | 17980694 |
| Streptococcus dysgalactiae subsp. equisimilis strain KNZ16 | EF-G | Translation elongation factor G | | | antibiotic target in susceptible species | 17980694 |
| Streptococcus dysgalactiae subsp. equisimilis strain NCTC10321 | EF-G | Translation elongation factor G | | | antibiotic target in susceptible species | 17980694 |
| Streptococcus dysgalactiae subsp. equisimilis strain NCTC11554 | EF-G | Translation elongation factor G | | | antibiotic target in susceptible species | 17980694 |
| Streptococcus dysgalactiae subsp. equisimilis strain NCTC11555 | EF-G | Translation elongation factor G | | | antibiotic target in susceptible species | 17980694 |
| Streptococcus dysgalactiae subsp. equisimilis strain NCTC11556 | EF-G | Translation elongation factor G | | | antibiotic target in susceptible species | 17980694 |
| Streptococcus dysgalactiae subsp. equisimilis strain NCTC11557 | EF-G | Translation elongation factor G | | | antibiotic target in susceptible species | 17980694 |
| Streptococcus dysgalactiae subsp. equisimilis strain NCTC11564 | EF-G | Translation elongation factor G | | | antibiotic target in susceptible species | 17980694 |
| Streptococcus dysgalactiae subsp. equisimilis strain NCTC11565 | EF-G | Translation elongation factor G | | | antibiotic target in susceptible species | 17980694 |
| Streptococcus dysgalactiae subsp. equisimilis strain NCTC5370 | fusA | Translation elongation factor G | | | antibiotic target in susceptible species | 17980694;19289529;19289529;17325218;19289529 |
| Streptococcus dysgalactiae subsp. equisimilis strain NCTC5371 | fusA | Translation elongation factor G | | | antibiotic target in susceptible species | 17980694;19289529;19289529;17325218;19289529 |
| Streptococcus dysgalactiae subsp. equisimilis strain NCTC5969 | EF-G | Translation elongation factor G | | | antibiotic target in susceptible species | 17980694 |
| Streptococcus dysgalactiae subsp. equisimilis strain NCTC6179 | fusA | Translation elongation factor G | | | antibiotic target in susceptible species | 17980694;19289529;19289529;17325218;19289529 |
| Streptococcus dysgalactiae subsp. equisimilis strain NCTC6181 | EF-G | Translation elongation factor G | | | antibiotic target in susceptible species | 17980694 |
| Streptococcus dysgalactiae subsp. equisimilis strain NCTC6407 | EF-G | Translation elongation factor G | | | antibiotic target in susceptible species | 17980694 |
| Streptococcus dysgalactiae subsp. equisimilis strain NCTC7136 | fusA | Translation elongation factor G | | | antibiotic target in susceptible species | 17980694;19289529;19289529;17325218;19289529 |
| Streptococcus dysgalactiae subsp. equisimilis strain NCTC8543 | EF-G | Translation elongation factor G | | | antibiotic target in susceptible species | 17980694 |
| Streptococcus dysgalactiae subsp. equisimilis strain NCTC8546 | EF-G | Translation elongation factor G | | | antibiotic target in susceptible species | 17980694 |
| Streptococcus dysgalactiae subsp. equisimilis strain NCTC9413 | EF-G | Translation elongation factor G | | | antibiotic target in susceptible species | 17980694 |
| Streptococcus dysgalactiae subsp. equisimilis strain NCTC9414 | fusA | Translation elongation factor G | | | antibiotic target in susceptible species | 17980694;19289529;19289529;17325218;19289529 |
| Streptococcus dysgalactiae subsp. equisimilis strain NCTC9603 | EF-G | Translation elongation factor G | | | antibiotic target in susceptible species | 17980694 |
| Streptococcus dysgalactiae subsp. equisimilis strain SS1575 | fusA | Translation elongation factor G | | | antibiotic target in susceptible species | 17980694;19289529;19289529;17325218;19289529 |
| Streptococcus dysgalactiae subsp. equisimilis strain T642 |  | Translation elongation factor G | | | antibiotic target in susceptible species | 17980694;19289529;19289529;17325218;19289529 |
| Streptococcus dysgalactiae subsp. equisimilis strain UT_4031CC |  | Translation elongation factor G | | | antibiotic target in susceptible species | 17980694;19289529;19289529;17325218;19289529 |
| Streptococcus dysgalactiae subsp. equisimilis strain UT_4231_KK |  | Translation elongation factor G | | | antibiotic target in susceptible species | 17980694;19289529;19289529;17325218;19289529 |
| Streptococcus dysgalactiae subsp. equisimilis strain UT_4234_DH | fusA | Translation elongation factor G | | | antibiotic target in susceptible species | 17980694;19289529;19289529;17325218;19289529 |
| Streptococcus dysgalactiae subsp. equisimilis strain UT_4241_XS | fusA | Translation elongation factor G | | | antibiotic target in susceptible species | 17980694;19289529;19289529;17325218;19289529 |
| Streptococcus dysgalactiae subsp. equisimilis strain UT_4242_AB | fusA | Translation elongation factor G | | | antibiotic target in susceptible species | 17980694;19289529;19289529;17325218;19289529 |
| Streptococcus dysgalactiae subsp. equisimilis strain UT_4255RC | fusA | Translation elongation factor G | | | antibiotic target in susceptible species | 17980694;19289529;19289529;17325218;19289529 |
| Streptococcus dysgalactiae subsp. equisimilis strain UT_4277_BB | fusA | Translation elongation factor G | | | antibiotic target in susceptible species | 17980694;19289529;19289529;17325218;19289529 |
| Streptococcus dysgalactiae subsp. equisimilis strain UT_4966_RC | fusA | Translation elongation factor G | | | antibiotic target in susceptible species | 17980694;19289529;19289529;17325218;19289529 |
| Streptococcus dysgalactiae subsp. equisimilis strain UT-5345 | fusA | Translation elongation factor G | | | antibiotic target in susceptible species | 17980694;19289529;19289529;17325218;19289529 |
| Streptococcus dysgalactiae subsp. equisimilis strain UT-5354 | fusA | Translation elongation factor G | | | antibiotic target in susceptible species | 17980694;19289529;19289529;17325218;19289529 |
| Streptococcus dysgalactiae subsp. equisimilis strain UT-SS1069 | fusA | Translation elongation factor G | | | antibiotic target in susceptible species | 17980694;19289529;19289529;17325218;19289529 |
| Streptococcus dysgalactiae subsp. equisimilis strain UT-SS957 | fusA | Translation elongation factor G | | | antibiotic target in susceptible species | 17980694;19289529;19289529;17325218;19289529 |
| Streptococcus dysgalactiae subsp. equisimilis strain WCHSDSE-1 | fusA | Translation elongation factor G | | | antibiotic target in susceptible species | 17980694;19289529;19289529;17325218;19289529 |
| Streptococcus dysgalactiae subsp. equisimilis SD SCDR1 | EF-Tu | Translation elongation factor Tu | | | antibiotic target in susceptible species | 364475;9678602 |
| Streptococcus dysgalactiae subsp. equisimilis 167 | EF-Tu | Translation elongation factor Tu | | | antibiotic target in susceptible species | 364475;9678602 |
| Streptococcus dysgalactiae subsp. equisimilis AC-2713 | tuf | Translation elongation factor Tu | | | antibiotic target in susceptible species | 364475;9678602 |
| Streptococcus dysgalactiae subsp. equisimilis AKSDE4288 | EF-Tu | Translation elongation factor Tu | | | antibiotic target in susceptible species | 364475;9678602 |
| Streptococcus dysgalactiae subsp. equisimilis ATCC 12394 | EF-Tu | Translation elongation factor Tu | | | antibiotic target in susceptible species | 364475;9678602 |
| Streptococcus dysgalactiae subsp. equisimilis GGS_124 | EF-Tu | Translation elongation factor Tu | | | antibiotic target in susceptible species | 364475;9678602 |
| Streptococcus dysgalactiae subsp. equisimilis RE378 | tuf | Translation elongation factor Tu | | | antibiotic target in susceptible species | 364475;9678602 |
| Streptococcus dysgalactiae subsp. equisimilis SK1249 | tuf | Translation elongation factor Tu | | | antibiotic target in susceptible species | 364475;9678602 |
| Streptococcus dysgalactiae subsp. equisimilis SK1250 | tuf | Translation elongation factor Tu | | | antibiotic target in susceptible species | 364475;9678602 |
| Streptococcus dysgalactiae subsp. equisimilis strain ASDSE_96 | EF-Tu | Translation elongation factor Tu | | | antibiotic target in susceptible species | 364475;9678602 |
| Streptococcus dysgalactiae subsp. equisimilis strain ASDSE_99 | EF-Tu | Translation elongation factor Tu | | | antibiotic target in susceptible species | 364475;9678602 |
| Streptococcus dysgalactiae subsp. equisimilis strain C161L1 | EF-Tu | Translation elongation factor Tu | | | antibiotic target in susceptible species | 364475;9678602 |
| Streptococcus dysgalactiae subsp. equisimilis strain KNZ01 | EF-Tu | Translation elongation factor Tu | | | antibiotic target in susceptible species | 364475;9678602 |
| Streptococcus dysgalactiae subsp. equisimilis strain KNZ03 | EF-Tu | Translation elongation factor Tu | | | antibiotic target in susceptible species | 364475;9678602 |
| Streptococcus dysgalactiae subsp. equisimilis strain KNZ04 | EF-Tu | Translation elongation factor Tu | | | antibiotic target in susceptible species | 364475;9678602 |
| Streptococcus dysgalactiae subsp. equisimilis strain KNZ06 | EF-Tu | Translation elongation factor Tu | | | antibiotic target in susceptible species | 364475;9678602 |
| Streptococcus dysgalactiae subsp. equisimilis strain KNZ07 | EF-Tu | Translation elongation factor Tu | | | antibiotic target in susceptible species | 364475;9678602 |
| Streptococcus dysgalactiae subsp. equisimilis strain KNZ10 | EF-Tu | Translation elongation factor Tu | | | antibiotic target in susceptible species | 364475;9678602 |
| Streptococcus dysgalactiae subsp. equisimilis strain KNZ12 | EF-Tu | Translation elongation factor Tu | | | antibiotic target in susceptible species | 364475;9678602 |
| Streptococcus dysgalactiae subsp. equisimilis strain KNZ15 | EF-Tu | Translation elongation factor Tu | | | antibiotic target in susceptible species | 364475;9678602 |
| Streptococcus dysgalactiae subsp. equisimilis strain KNZ16 | EF-Tu | Translation elongation factor Tu | | | antibiotic target in susceptible species | 364475;9678602 |
| Streptococcus dysgalactiae subsp. equisimilis strain NCTC10321 | EF-Tu | Translation elongation factor Tu | | | antibiotic target in susceptible species | 364475;9678602 |
| Streptococcus dysgalactiae subsp. equisimilis strain NCTC11554 | EF-Tu | Translation elongation factor Tu | | | antibiotic target in susceptible species | 364475;9678602 |
| Streptococcus dysgalactiae subsp. equisimilis strain NCTC11555 | EF-Tu | Translation elongation factor Tu | | | antibiotic target in susceptible species | 364475;9678602 |
| Streptococcus dysgalactiae subsp. equisimilis strain NCTC11556 | EF-Tu | Translation elongation factor Tu | | | antibiotic target in susceptible species | 364475;9678602 |
| Streptococcus dysgalactiae subsp. equisimilis strain NCTC11557 | EF-Tu | Translation elongation factor Tu | | | antibiotic target in susceptible species | 364475;9678602 |
| Streptococcus dysgalactiae subsp. equisimilis strain NCTC11564 | EF-Tu | Translation elongation factor Tu | | | antibiotic target in susceptible species | 364475;9678602 |
| Streptococcus dysgalactiae subsp. equisimilis strain NCTC11565 | EF-Tu | Translation elongation factor Tu | | | antibiotic target in susceptible species | 364475;9678602 |
| Streptococcus dysgalactiae subsp. equisimilis strain NCTC5370 | tuf | Translation elongation factor Tu | | | antibiotic target in susceptible species | 364475;9678602 |
| Streptococcus dysgalactiae subsp. equisimilis strain NCTC5371 | tuf | Translation elongation factor Tu | | | antibiotic target in susceptible species | 364475;9678602 |
| Streptococcus dysgalactiae subsp. equisimilis strain NCTC5969 | EF-Tu | Translation elongation factor Tu | | | antibiotic target in susceptible species | 364475;9678602 |
| Streptococcus dysgalactiae subsp. equisimilis strain NCTC6179 | tuf | Translation elongation factor Tu | | | antibiotic target in susceptible species | 364475;9678602 |
| Streptococcus dysgalactiae subsp. equisimilis strain NCTC6181 | EF-Tu | Translation elongation factor Tu | | | antibiotic target in susceptible species | 364475;9678602 |
| Streptococcus dysgalactiae subsp. equisimilis strain NCTC6407 | EF-Tu | Translation elongation factor Tu | | | antibiotic target in susceptible species | 364475;9678602 |
| Streptococcus dysgalactiae subsp. equisimilis strain NCTC7136 | tuf | Translation elongation factor Tu | | | antibiotic target in susceptible species | 364475;9678602 |
| Streptococcus dysgalactiae subsp. equisimilis strain NCTC8543 | EF-Tu | Translation elongation factor Tu | | | antibiotic target in susceptible species | 364475;9678602 |
| Streptococcus dysgalactiae subsp. equisimilis strain NCTC8546 | EF-Tu | Translation elongation factor Tu | | | antibiotic target in susceptible species | 364475;9678602 |
| Streptococcus dysgalactiae subsp. equisimilis strain NCTC9413 | EF-Tu | Translation elongation factor Tu | | | antibiotic target in susceptible species | 364475;9678602 |
| Streptococcus dysgalactiae subsp. equisimilis strain NCTC9414 | tuf | Translation elongation factor Tu | | | antibiotic target in susceptible species | 364475;9678602 |
| Streptococcus dysgalactiae subsp. equisimilis strain NCTC9603 | EF-Tu | Translation elongation factor Tu | | | antibiotic target in susceptible species | 364475;9678602 |
| Streptococcus dysgalactiae subsp. equisimilis strain SS1575 | tuf | Translation elongation factor Tu | | | antibiotic target in susceptible species | 364475;9678602 |
| Streptococcus dysgalactiae subsp. equisimilis strain T642 | EF-Tu | Translation elongation factor Tu | | | antibiotic target in susceptible species | 364475;9678602 |
| Streptococcus dysgalactiae subsp. equisimilis strain UT_4031CC | EF-Tu | Translation elongation factor Tu | | | antibiotic target in susceptible species | 364475;9678602 |
| Streptococcus dysgalactiae subsp. equisimilis strain UT_4231_KK | EF-Tu | Translation elongation factor Tu | | | antibiotic target in susceptible species | 364475;9678602 |
| Streptococcus dysgalactiae subsp. equisimilis strain UT_4234_DH | EF-Tu | Translation elongation factor Tu | | | antibiotic target in susceptible species | 364475;9678602 |
| Streptococcus dysgalactiae subsp. equisimilis strain UT_4241_XS | EF-Tu | Translation elongation factor Tu | | | antibiotic target in susceptible species | 364475;9678602 |
| Streptococcus dysgalactiae subsp. equisimilis strain UT_4242_AB | EF-Tu | Translation elongation factor Tu | | | antibiotic target in susceptible species | 364475;9678602 |
| Streptococcus dysgalactiae subsp. equisimilis strain UT_4255RC | EF-Tu | Translation elongation factor Tu | | | antibiotic target in susceptible species | 364475;9678602 |
| Streptococcus dysgalactiae subsp. equisimilis strain UT_4277_BB | EF-Tu | Translation elongation factor Tu | | | antibiotic target in susceptible species | 364475;9678602 |
| Streptococcus dysgalactiae subsp. equisimilis strain UT_4966_RC | EF-Tu | Translation elongation factor Tu | | | antibiotic target in susceptible species | 364475;9678602 |
| Streptococcus dysgalactiae subsp. equisimilis strain UT-5345 | tuf | Translation elongation factor Tu | | | antibiotic target in susceptible species | 364475;9678602 |
| Streptococcus dysgalactiae subsp. equisimilis strain UT-5354 | tuf | Translation elongation factor Tu | | | antibiotic target in susceptible species | 364475;9678602 |
| Streptococcus dysgalactiae subsp. equisimilis strain UT-SS1069 | tuf | Translation elongation factor Tu | | | antibiotic target in susceptible species | 364475;9678602 |
| Streptococcus dysgalactiae subsp. equisimilis strain UT-SS957 | tuf | Translation elongation factor Tu | | | antibiotic target in susceptible species | 364475;9678602 |
| Streptococcus dysgalactiae subsp. equisimilis strain WCHSDSE-1 | EF-Tu | Translation elongation factor Tu | | | antibiotic target in susceptible species | 364475;9678602 |
| Streptococcus dysgalactiae subsp. equisimilis SD SCDR1 | FabK | Enoyl-[acyl-carrier-protein] reductase [FMN, NADH] (EC 1.3.1.9), FabK => refractory to triclosan | | | antibiotic target replacement protein | 10910344 |
| Streptococcus dysgalactiae subsp. equisimilis 167 | fabK | Enoyl-[acyl-carrier-protein] reductase [FMN, NADH] (EC 1.3.1.9), FabK => refractory to triclosan | | | antibiotic target replacement protein | 10910344 |
| Streptococcus dysgalactiae subsp. equisimilis AC-2713 | fabK | Enoyl-[acyl-carrier-protein] reductase [FMN, NADH] (EC 1.3.1.9), FabK => refractory to triclosan | | | antibiotic target replacement protein | 10910344 |
| Streptococcus dysgalactiae subsp. equisimilis AKSDE4288 | FabK | Enoyl-[acyl-carrier-protein] reductase [FMN, NADH] (EC 1.3.1.9), FabK => refractory to triclosan | | | antibiotic target replacement protein | 10910344 |
| Streptococcus dysgalactiae subsp. equisimilis ATCC 12394 | FabK | Enoyl-[acyl-carrier-protein] reductase [FMN, NADH] (EC 1.3.1.9), FabK => refractory to triclosan | | | antibiotic target replacement protein | 10910344 |
| Streptococcus dysgalactiae subsp. equisimilis GGS_124 | FabK | Enoyl-[acyl-carrier-protein] reductase [FMN, NADH] (EC 1.3.1.9), FabK => refractory to triclosan | | | antibiotic target replacement protein | 10910344 |
| Streptococcus dysgalactiae subsp. equisimilis RE378 | fabK | Enoyl-[acyl-carrier-protein] reductase [FMN, NADH] (EC 1.3.1.9), FabK => refractory to triclosan | | | antibiotic target replacement protein | 10910344 |
| Streptococcus dysgalactiae subsp. equisimilis SK1249 | FabK | Enoyl-[acyl-carrier-protein] reductase [FMN, NADH] (EC 1.3.1.9), FabK => refractory to triclosan | | | antibiotic target replacement protein | 10910344 |
| Streptococcus dysgalactiae subsp. equisimilis SK1250 | FabK | Enoyl-[acyl-carrier-protein] reductase [FMN, NADH] (EC 1.3.1.9), FabK => refractory to triclosan | | | antibiotic target replacement protein | 10910344 |
| Streptococcus dysgalactiae subsp. equisimilis strain ASDSE_96 | FabK | Enoyl-[acyl-carrier-protein] reductase [FMN, NADH] (EC 1.3.1.9), FabK => refractory to triclosan | | | antibiotic target replacement protein | 10910344 |
| Streptococcus dysgalactiae subsp. equisimilis strain C161L1 | FabK | Enoyl-[acyl-carrier-protein] reductase [FMN, NADH] (EC 1.3.1.9), FabK => refractory to triclosan | | | antibiotic target replacement protein | 10910344 |
| Streptococcus dysgalactiae subsp. equisimilis strain KNZ01 | FabK | Enoyl-[acyl-carrier-protein] reductase [FMN, NADH] (EC 1.3.1.9), FabK => refractory to triclosan | | | antibiotic target replacement protein | 10910344 |
| Streptococcus dysgalactiae subsp. equisimilis strain KNZ03 | FabK | Enoyl-[acyl-carrier-protein] reductase [FMN, NADH] (EC 1.3.1.9), FabK => refractory to triclosan | | | antibiotic target replacement protein | 10910344 |
| Streptococcus dysgalactiae subsp. equisimilis strain KNZ04 | FabK | Enoyl-[acyl-carrier-protein] reductase [FMN, NADH] (EC 1.3.1.9), FabK => refractory to triclosan | | | antibiotic target replacement protein | 10910344 |
| Streptococcus dysgalactiae subsp. equisimilis strain KNZ06 | FabK | Enoyl-[acyl-carrier-protein] reductase [FMN, NADH] (EC 1.3.1.9), FabK => refractory to triclosan | | | antibiotic target replacement protein | 10910344 |
| Streptococcus dysgalactiae subsp. equisimilis strain KNZ07 | FabK | Enoyl-[acyl-carrier-protein] reductase [FMN, NADH] (EC 1.3.1.9), FabK => refractory to triclosan | | | antibiotic target replacement protein | 10910344 |
| Streptococcus dysgalactiae subsp. equisimilis strain KNZ10 | FabK | Enoyl-[acyl-carrier-protein] reductase [FMN, NADH] (EC 1.3.1.9), FabK => refractory to triclosan | | | antibiotic target replacement protein | 10910344 |
| Streptococcus dysgalactiae subsp. equisimilis strain KNZ12 | FabK | Enoyl-[acyl-carrier-protein] reductase [FMN, NADH] (EC 1.3.1.9), FabK => refractory to triclosan | | | antibiotic target replacement protein | 10910344 |
| Streptococcus dysgalactiae subsp. equisimilis strain KNZ15 | FabK | Enoyl-[acyl-carrier-protein] reductase [FMN, NADH] (EC 1.3.1.9), FabK => refractory to triclosan | | | antibiotic target replacement protein | 10910344 |
| Streptococcus dysgalactiae subsp. equisimilis strain KNZ16 | FabK | Enoyl-[acyl-carrier-protein] reductase [FMN, NADH] (EC 1.3.1.9), FabK => refractory to triclosan | | | antibiotic target replacement protein | 10910344 |
| Streptococcus dysgalactiae subsp. equisimilis strain NCTC10321 | FabK | Enoyl-[acyl-carrier-protein] reductase [FMN, NADH] (EC 1.3.1.9), FabK => refractory to triclosan | | | antibiotic target replacement protein | 10910344 |
| Streptococcus dysgalactiae subsp. equisimilis strain NCTC11554 | FabK | Enoyl-[acyl-carrier-protein] reductase [FMN, NADH] (EC 1.3.1.9), FabK => refractory to triclosan | | | antibiotic target replacement protein | 10910344 |
| Streptococcus dysgalactiae subsp. equisimilis strain NCTC11555 | FabK | Enoyl-[acyl-carrier-protein] reductase [FMN, NADH] (EC 1.3.1.9), FabK => refractory to triclosan | | | antibiotic target replacement protein | 10910344 |
| Streptococcus dysgalactiae subsp. equisimilis strain NCTC11556 | FabK | Enoyl-[acyl-carrier-protein] reductase [FMN, NADH] (EC 1.3.1.9), FabK => refractory to triclosan | | | antibiotic target replacement protein | 10910344 |
| Streptococcus dysgalactiae subsp. equisimilis strain NCTC11557 | FabK | Enoyl-[acyl-carrier-protein] reductase [FMN, NADH] (EC 1.3.1.9), FabK => refractory to triclosan | | | antibiotic target replacement protein | 10910344 |
| Streptococcus dysgalactiae subsp. equisimilis strain NCTC11564 | FabK | Enoyl-[acyl-carrier-protein] reductase [FMN, NADH] (EC 1.3.1.9), FabK => refractory to triclosan | | | antibiotic target replacement protein | 10910344 |
| Streptococcus dysgalactiae subsp. equisimilis strain NCTC5370 | FabK | Enoyl-[acyl-carrier-protein] reductase [FMN, NADH] (EC 1.3.1.9), FabK => refractory to triclosan | | | antibiotic target replacement protein | 10910344 |
| Streptococcus dysgalactiae subsp. equisimilis strain NCTC5371 | FabK | Enoyl-[acyl-carrier-protein] reductase [FMN, NADH] (EC 1.3.1.9), FabK => refractory to triclosan | | | antibiotic target replacement protein | 10910344 |
| Streptococcus dysgalactiae subsp. equisimilis strain NCTC6179 | FabK | Enoyl-[acyl-carrier-protein] reductase [FMN, NADH] (EC 1.3.1.9), FabK => refractory to triclosan | | | antibiotic target replacement protein | 10910344 |
| Streptococcus dysgalactiae subsp. equisimilis strain NCTC6181 | FabK | Enoyl-[acyl-carrier-protein] reductase [FMN, NADH] (EC 1.3.1.9), FabK => refractory to triclosan | | | antibiotic target replacement protein | 10910344 |
| Streptococcus dysgalactiae subsp. equisimilis strain NCTC6407 | FabK | Enoyl-[acyl-carrier-protein] reductase [FMN, NADH] (EC 1.3.1.9), FabK => refractory to triclosan | | | antibiotic target replacement protein | 10910344 |
| Streptococcus dysgalactiae subsp. equisimilis strain NCTC7136 | FabK | Enoyl-[acyl-carrier-protein] reductase [FMN, NADH] (EC 1.3.1.9), FabK => refractory to triclosan | | | antibiotic target replacement protein | 10910344 |
| Streptococcus dysgalactiae subsp. equisimilis strain NCTC8543 | FabK | Enoyl-[acyl-carrier-protein] reductase [FMN, NADH] (EC 1.3.1.9), FabK => refractory to triclosan | | | antibiotic target replacement protein | 10910344 |
| Streptococcus dysgalactiae subsp. equisimilis strain NCTC8546 | FabK | Enoyl-[acyl-carrier-protein] reductase [FMN, NADH] (EC 1.3.1.9), FabK => refractory to triclosan | | | antibiotic target replacement protein | 10910344 |
| Streptococcus dysgalactiae subsp. equisimilis strain NCTC9413 | FabK | Enoyl-[acyl-carrier-protein] reductase [FMN, NADH] (EC 1.3.1.9), FabK => refractory to triclosan | | | antibiotic target replacement protein | 10910344 |
| Streptococcus dysgalactiae subsp. equisimilis strain NCTC9414 | FabK | Enoyl-[acyl-carrier-protein] reductase [FMN, NADH] (EC 1.3.1.9), FabK => refractory to triclosan | | | antibiotic target replacement protein | 10910344 |
| Streptococcus dysgalactiae subsp. equisimilis strain NCTC9603 | FabK | Enoyl-[acyl-carrier-protein] reductase [FMN, NADH] (EC 1.3.1.9), FabK => refractory to triclosan | | | antibiotic target replacement protein | 10910344 |
| Streptococcus dysgalactiae subsp. equisimilis strain SS1575 | fabK | Enoyl-[acyl-carrier-protein] reductase [FMN, NADH] (EC 1.3.1.9), FabK => refractory to triclosan | | | antibiotic target replacement protein | 10910344 |
| Streptococcus dysgalactiae subsp. equisimilis strain T642 | FabK | Enoyl-[acyl-carrier-protein] reductase [FMN, NADH] (EC 1.3.1.9), FabK => refractory to triclosan | | | antibiotic target replacement protein | 10910344 |
| Streptococcus dysgalactiae subsp. equisimilis strain UT_4031CC | FabK | Enoyl-[acyl-carrier-protein] reductase [FMN, NADH] (EC 1.3.1.9), FabK => refractory to triclosan | | | antibiotic target replacement protein | 10910344 |
| Streptococcus dysgalactiae subsp. equisimilis strain UT_4231_KK | FabK | Enoyl-[acyl-carrier-protein] reductase [FMN, NADH] (EC 1.3.1.9), FabK => refractory to triclosan | | | antibiotic target replacement protein | 10910344 |
| Streptococcus dysgalactiae subsp. equisimilis strain UT_4234_DH | FabK | Enoyl-[acyl-carrier-protein] reductase [FMN, NADH] (EC 1.3.1.9), FabK => refractory to triclosan | | | antibiotic target replacement protein | 10910344 |
| Streptococcus dysgalactiae subsp. equisimilis strain UT_4241_XS | FabK | Enoyl-[acyl-carrier-protein] reductase [FMN, NADH] (EC 1.3.1.9), FabK => refractory to triclosan | | | antibiotic target replacement protein | 10910344 |
| Streptococcus dysgalactiae subsp. equisimilis strain UT_4242_AB | FabK | Enoyl-[acyl-carrier-protein] reductase [FMN, NADH] (EC 1.3.1.9), FabK => refractory to triclosan | | | antibiotic target replacement protein | 10910344 |
| Streptococcus dysgalactiae subsp. equisimilis strain UT_4255RC | FabK | Enoyl-[acyl-carrier-protein] reductase [FMN, NADH] (EC 1.3.1.9), FabK => refractory to triclosan | | | antibiotic target replacement protein | 10910344 |
| Streptococcus dysgalactiae subsp. equisimilis strain UT_4277_BB | FabK | Enoyl-[acyl-carrier-protein] reductase [FMN, NADH] (EC 1.3.1.9), FabK => refractory to triclosan | | | antibiotic target replacement protein | 10910344 |
| Streptococcus dysgalactiae subsp. equisimilis strain UT_4966_RC | FabK | Enoyl-[acyl-carrier-protein] reductase [FMN, NADH] (EC 1.3.1.9), FabK => refractory to triclosan | | | antibiotic target replacement protein | 10910344 |
| Streptococcus dysgalactiae subsp. equisimilis strain UT-5345 | FabK | Enoyl-[acyl-carrier-protein] reductase [FMN, NADH] (EC 1.3.1.9), FabK => refractory to triclosan | | | antibiotic target replacement protein | 10910344 |
| Streptococcus dysgalactiae subsp. equisimilis strain UT-5354 | FabK | Enoyl-[acyl-carrier-protein] reductase [FMN, NADH] (EC 1.3.1.9), FabK => refractory to triclosan | | | antibiotic target replacement protein | 10910344 |
| Streptococcus dysgalactiae subsp. equisimilis strain UT-SS1069 | FabK | Enoyl-[acyl-carrier-protein] reductase [FMN, NADH] (EC 1.3.1.9), FabK => refractory to triclosan | | | antibiotic target replacement protein | 10910344 |
| Streptococcus dysgalactiae subsp. equisimilis strain UT-SS957 | FabK | Enoyl-[acyl-carrier-protein] reductase [FMN, NADH] (EC 1.3.1.9), FabK => refractory to triclosan | | | antibiotic target replacement protein | 10910344 |
| Streptococcus dysgalactiae subsp. equisimilis strain WCHSDSE-1 | FabK | Enoyl-[acyl-carrier-protein] reductase [FMN, NADH] (EC 1.3.1.9), FabK => refractory to triclosan | | | antibiotic target replacement protein | 10910344 |
| Streptococcus dysgalactiae subsp. equisimilis SD SCDR1 | folP | Dihydropteroate synthase (EC 2.5.1.15) | | | antibiotic resistant gene variant or mutant,sulfonamide resistance gene | 15673783 |
| Streptococcus dysgalactiae subsp. equisimilis 167 | folP | Dihydropteroate synthase (EC 2.5.1.15) | | | antibiotic resistant gene variant or mutant,sulfonamide resistance gene | 15673783 |
| Streptococcus dysgalactiae subsp. equisimilis AC-2713 | folP | Dihydropteroate synthase (EC 2.5.1.15) | | | antibiotic resistant gene variant or mutant,sulfonamide resistance gene | 15673783 |
| Streptococcus dysgalactiae subsp. equisimilis AKSDE4288 | folP | Dihydropteroate synthase (EC 2.5.1.15) | | | antibiotic resistant gene variant or mutant,sulfonamide resistance gene | 15673783 |
| Streptococcus dysgalactiae subsp. equisimilis ATCC 12394 | folP | Dihydropteroate synthase (EC 2.5.1.15) | | | antibiotic resistant gene variant or mutant,sulfonamide resistance gene | 15673783 |
| Streptococcus dysgalactiae subsp. equisimilis GGS_124 | folP | Dihydropteroate synthase (EC 2.5.1.15) | | | antibiotic resistant gene variant or mutant,sulfonamide resistance gene | 15673783 |
| Streptococcus dysgalactiae subsp. equisimilis RE378 | folP | Dihydropteroate synthase (EC 2.5.1.15) | | | antibiotic resistant gene variant or mutant,sulfonamide resistance gene | 15673783 |
| Streptococcus dysgalactiae subsp. equisimilis SK1249 | folP | Dihydropteroate synthase (EC 2.5.1.15) | | | antibiotic resistant gene variant or mutant,sulfonamide resistance gene | 15673783 |
| Streptococcus dysgalactiae subsp. equisimilis SK1250 | folP | Dihydropteroate synthase (EC 2.5.1.15) | | | antibiotic resistant gene variant or mutant,sulfonamide resistance gene | 15673783 |
| Streptococcus dysgalactiae subsp. equisimilis strain ASDSE_96 | folP | Dihydropteroate synthase (EC 2.5.1.15) | | | antibiotic resistant gene variant or mutant,sulfonamide resistance gene | 15673783 |
| Streptococcus dysgalactiae subsp. equisimilis strain ASDSE_99 | folP | Dihydropteroate synthase (EC 2.5.1.15) | | | antibiotic resistant gene variant or mutant,sulfonamide resistance gene | 15673783 |
| Streptococcus dysgalactiae subsp. equisimilis strain C161L1 | folP | Dihydropteroate synthase (EC 2.5.1.15) | | | antibiotic resistant gene variant or mutant,sulfonamide resistance gene | 15673783 |
| Streptococcus dysgalactiae subsp. equisimilis strain KNZ01 | folP | Dihydropteroate synthase (EC 2.5.1.15) | | | antibiotic resistant gene variant or mutant,sulfonamide resistance gene | 15673783 |
| Streptococcus dysgalactiae subsp. equisimilis strain KNZ03 | folP | Dihydropteroate synthase (EC 2.5.1.15) | | | antibiotic resistant gene variant or mutant,sulfonamide resistance gene | 15673783 |
| Streptococcus dysgalactiae subsp. equisimilis strain KNZ04 | folP | Dihydropteroate synthase (EC 2.5.1.15) | | | antibiotic resistant gene variant or mutant,sulfonamide resistance gene | 15673783 |
| Streptococcus dysgalactiae subsp. equisimilis strain KNZ06 | folP | Dihydropteroate synthase (EC 2.5.1.15) | | | antibiotic resistant gene variant or mutant,sulfonamide resistance gene | 15673783 |
| Streptococcus dysgalactiae subsp. equisimilis strain KNZ07 | folP | Dihydropteroate synthase (EC 2.5.1.15) | | | antibiotic resistant gene variant or mutant,sulfonamide resistance gene | 15673783 |
| Streptococcus dysgalactiae subsp. equisimilis strain KNZ10 | folP | Dihydropteroate synthase (EC 2.5.1.15) | | | antibiotic resistant gene variant or mutant,sulfonamide resistance gene | 15673783 |
| Streptococcus dysgalactiae subsp. equisimilis strain KNZ12 | folP | Dihydropteroate synthase (EC 2.5.1.15) | | | antibiotic resistant gene variant or mutant,sulfonamide resistance gene | 15673783 |
| Streptococcus dysgalactiae subsp. equisimilis strain KNZ15 | folP | Dihydropteroate synthase (EC 2.5.1.15) | | | antibiotic resistant gene variant or mutant,sulfonamide resistance gene | 15673783 |
| Streptococcus dysgalactiae subsp. equisimilis strain KNZ16 | folP | Dihydropteroate synthase (EC 2.5.1.15) | | | antibiotic resistant gene variant or mutant,sulfonamide resistance gene | 15673783 |
| Streptococcus dysgalactiae subsp. equisimilis strain NCTC10321 | folP | Dihydropteroate synthase (EC 2.5.1.15) | | | antibiotic resistant gene variant or mutant,sulfonamide resistance gene | 15673783 |
| Streptococcus dysgalactiae subsp. equisimilis strain NCTC11554 | folP | Dihydropteroate synthase (EC 2.5.1.15) | | | antibiotic resistant gene variant or mutant,sulfonamide resistance gene | 15673783 |
| Streptococcus dysgalactiae subsp. equisimilis strain NCTC11555 | folP | Dihydropteroate synthase (EC 2.5.1.15) | | | antibiotic resistant gene variant or mutant,sulfonamide resistance gene | 15673783 |
| Streptococcus dysgalactiae subsp. equisimilis strain NCTC11556 | folP | Dihydropteroate synthase (EC 2.5.1.15) | | | antibiotic resistant gene variant or mutant,sulfonamide resistance gene | 15673783 |
| Streptococcus dysgalactiae subsp. equisimilis strain NCTC11557 | folP | Dihydropteroate synthase (EC 2.5.1.15) | | | antibiotic resistant gene variant or mutant,sulfonamide resistance gene | 15673783 |
| Streptococcus dysgalactiae subsp. equisimilis strain NCTC11564 | folP | Dihydropteroate synthase (EC 2.5.1.15) | | | antibiotic resistant gene variant or mutant,sulfonamide resistance gene | 15673783 |
| Streptococcus dysgalactiae subsp. equisimilis strain NCTC11565 | folP | Dihydropteroate synthase (EC 2.5.1.15) | | | antibiotic resistant gene variant or mutant,sulfonamide resistance gene | 15673783 |
| Streptococcus dysgalactiae subsp. equisimilis strain NCTC5370 | folP | Dihydropteroate synthase (EC 2.5.1.15) | | | antibiotic resistant gene variant or mutant,sulfonamide resistance gene | 15673783 |
| Streptococcus dysgalactiae subsp. equisimilis strain NCTC5371 | folP | Dihydropteroate synthase (EC 2.5.1.15) | | | antibiotic resistant gene variant or mutant,sulfonamide resistance gene | 15673783 |
| Streptococcus dysgalactiae subsp. equisimilis strain NCTC5969 | folP | Dihydropteroate synthase (EC 2.5.1.15) | | | antibiotic resistant gene variant or mutant,sulfonamide resistance gene | 15673783 |
| Streptococcus dysgalactiae subsp. equisimilis strain NCTC6179 | folP | Dihydropteroate synthase (EC 2.5.1.15) | | | antibiotic resistant gene variant or mutant,sulfonamide resistance gene | 15673783 |
| Streptococcus dysgalactiae subsp. equisimilis strain NCTC6181 | folP | Dihydropteroate synthase (EC 2.5.1.15) | | | antibiotic resistant gene variant or mutant,sulfonamide resistance gene | 15673783 |
| Streptococcus dysgalactiae subsp. equisimilis strain NCTC6407 | folP | Dihydropteroate synthase (EC 2.5.1.15) | | | antibiotic resistant gene variant or mutant,sulfonamide resistance gene | 15673783 |
| Streptococcus dysgalactiae subsp. equisimilis strain NCTC7136 | folP | Dihydropteroate synthase (EC 2.5.1.15) | | | antibiotic resistant gene variant or mutant,sulfonamide resistance gene | 15673783 |
| Streptococcus dysgalactiae subsp. equisimilis strain NCTC8543 | folP | Dihydropteroate synthase (EC 2.5.1.15) | | | antibiotic resistant gene variant or mutant,sulfonamide resistance gene | 15673783 |
| Streptococcus dysgalactiae subsp. equisimilis strain NCTC8546 | folP | Dihydropteroate synthase (EC 2.5.1.15) | | | antibiotic resistant gene variant or mutant,sulfonamide resistance gene | 15673783 |
| Streptococcus dysgalactiae subsp. equisimilis strain NCTC9413 | folP | Dihydropteroate synthase (EC 2.5.1.15) | | | antibiotic resistant gene variant or mutant,sulfonamide resistance gene | 15673783 |
| Streptococcus dysgalactiae subsp. equisimilis strain NCTC9414 | folP | Dihydropteroate synthase (EC 2.5.1.15) | | | antibiotic resistant gene variant or mutant,sulfonamide resistance gene | 15673783 |
| Streptococcus dysgalactiae subsp. equisimilis strain NCTC9603 | folP | Dihydropteroate synthase (EC 2.5.1.15) | | | antibiotic resistant gene variant or mutant,sulfonamide resistance gene | 15673783 |
| Streptococcus dysgalactiae subsp. equisimilis strain SS1575 | folP | Dihydropteroate synthase (EC 2.5.1.15) | | | antibiotic resistant gene variant or mutant,sulfonamide resistance gene | 15673783 |
| Streptococcus dysgalactiae subsp. equisimilis strain T642 | folP | Dihydropteroate synthase (EC 2.5.1.15) | | | antibiotic resistant gene variant or mutant,sulfonamide resistance gene | 15673783 |
| Streptococcus dysgalactiae subsp. equisimilis strain UT_4031CC | folP | Dihydropteroate synthase (EC 2.5.1.15) | | | antibiotic resistant gene variant or mutant,sulfonamide resistance gene | 15673783 |
| Streptococcus dysgalactiae subsp. equisimilis strain UT_4231_KK | folP | Dihydropteroate synthase (EC 2.5.1.15) | | | antibiotic resistant gene variant or mutant,sulfonamide resistance gene | 15673783 |
| Streptococcus dysgalactiae subsp. equisimilis strain UT_4234_DH | folP | Dihydropteroate synthase (EC 2.5.1.15) | | | antibiotic resistant gene variant or mutant,sulfonamide resistance gene | 15673783 |
| Streptococcus dysgalactiae subsp. equisimilis strain UT_4241_XS | folP | Dihydropteroate synthase (EC 2.5.1.15) | | | antibiotic resistant gene variant or mutant,sulfonamide resistance gene | 15673783 |
| Streptococcus dysgalactiae subsp. equisimilis strain UT_4242_AB | folP | Dihydropteroate synthase (EC 2.5.1.15) | | | antibiotic resistant gene variant or mutant,sulfonamide resistance gene | 15673783 |
| Streptococcus dysgalactiae subsp. equisimilis strain UT_4255RC | folP | Dihydropteroate synthase (EC 2.5.1.15) | | | antibiotic resistant gene variant or mutant,sulfonamide resistance gene | 15673783 |
| Streptococcus dysgalactiae subsp. equisimilis strain UT_4277_BB | folP | Dihydropteroate synthase (EC 2.5.1.15) | | | antibiotic resistant gene variant or mutant,sulfonamide resistance gene | 15673783 |
| Streptococcus dysgalactiae subsp. equisimilis strain UT_4966_RC | folP | Dihydropteroate synthase (EC 2.5.1.15) | | | antibiotic resistant gene variant or mutant,sulfonamide resistance gene | 15673783 |
| Streptococcus dysgalactiae subsp. equisimilis strain UT-5345 | folP | Dihydropteroate synthase (EC 2.5.1.15) | | | antibiotic resistant gene variant or mutant,sulfonamide resistance gene | 15673783 |
| Streptococcus dysgalactiae subsp. equisimilis strain UT-5354 | folP | Dihydropteroate synthase (EC 2.5.1.15) | | | antibiotic resistant gene variant or mutant,sulfonamide resistance gene | 15673783 |
| Streptococcus dysgalactiae subsp. equisimilis strain UT-SS1069 | folP | Dihydropteroate synthase (EC 2.5.1.15) | | | antibiotic resistant gene variant or mutant,sulfonamide resistance gene | 15673783 |
| Streptococcus dysgalactiae subsp. equisimilis strain UT-SS957 | folP | Dihydropteroate synthase (EC 2.5.1.15) | | | antibiotic resistant gene variant or mutant;sulfonamide resistance gene | 15673783 |
| Streptococcus dysgalactiae subsp. equisimilis strain WCHSDSE-1 | folP | Dihydropteroate synthase (EC 2.5.1.15) | | | antibiotic resistant gene variant or mutant;sulfonamide resistance gene | 15673783 |
| Streptococcus dysgalactiae subsp. equisimilis SD SCDR1 | gyrA | DNA gyrase subunit A (EC 5.99.1.3) | | | antibiotic target in susceptible species | 9293187 |
| Streptococcus dysgalactiae subsp. equisimilis 167 | gyrA | DNA gyrase subunit A (EC 5.99.1.3) | | | antibiotic target in susceptible species | 9293187 |
| Streptococcus dysgalactiae subsp. equisimilis AC-2713 | gyrA | DNA gyrase subunit A (EC 5.99.1.3) | | | antibiotic target in susceptible species | 9293187 |
| Streptococcus dysgalactiae subsp. equisimilis AKSDE4288 | gyrA | DNA gyrase subunit A (EC 5.99.1.3) | | | antibiotic target in susceptible species | 9293187 |
| Streptococcus dysgalactiae subsp. equisimilis ATCC 12394 | gyrA | DNA gyrase subunit A (EC 5.99.1.3) | | | antibiotic target in susceptible species | 9293187 |
| Streptococcus dysgalactiae subsp. equisimilis GGS_124 | gyrA | DNA gyrase subunit A (EC 5.99.1.3) | | | antibiotic target in susceptible species | 9293187 |
| Streptococcus dysgalactiae subsp. equisimilis RE378 | gyrA | DNA gyrase subunit A (EC 5.99.1.3) | | | antibiotic target in susceptible species | 9293187 |
| Streptococcus dysgalactiae subsp. equisimilis SK1249 | gyrA | DNA gyrase subunit A (EC 5.99.1.3) | | | antibiotic target in susceptible species | 9293187 |
| Streptococcus dysgalactiae subsp. equisimilis SK1250 | gyrA | DNA gyrase subunit A (EC 5.99.1.3) | | | antibiotic target in susceptible species | 9293187 |
| Streptococcus dysgalactiae subsp. equisimilis strain ASDSE_96 | gyrA | DNA gyrase subunit A (EC 5.99.1.3) | | | antibiotic target in susceptible species | 9293187 |
| Streptococcus dysgalactiae subsp. equisimilis strain ASDSE_99 | gyrA | DNA gyrase subunit A (EC 5.99.1.3) | | | antibiotic target in susceptible species | 9293187 |
| Streptococcus dysgalactiae subsp. equisimilis strain C161L1 | gyrA | DNA gyrase subunit A (EC 5.99.1.3) | | | antibiotic target in susceptible species | 9293187 |
| Streptococcus dysgalactiae subsp. equisimilis strain C161L1 | gyrA | DNA gyrase subunit A (EC 5.99.1.3) | | | antibiotic target in susceptible species | 9293187 |
| Streptococcus dysgalactiae subsp. equisimilis strain C161L1 | gyrA | DNA gyrase subunit A (EC 5.99.1.3) | | | antibiotic target in susceptible species | 9293187 |
| Streptococcus dysgalactiae subsp. equisimilis strain KNZ01 | gyrA | DNA gyrase subunit A (EC 5.99.1.3) | | | antibiotic target in susceptible species | 9293187 |
| Streptococcus dysgalactiae subsp. equisimilis strain KNZ01 | gyrA | DNA gyrase subunit A (EC 5.99.1.3) | | | antibiotic target in susceptible species | 9293187 |
| Streptococcus dysgalactiae subsp. equisimilis strain KNZ03 | gyrA | DNA gyrase subunit A (EC 5.99.1.3) | | | antibiotic target in susceptible species | 9293187 |
| Streptococcus dysgalactiae subsp. equisimilis strain KNZ03 | gyrA | DNA gyrase subunit A (EC 5.99.1.3) | | | antibiotic target in susceptible species | 9293187 |
| Streptococcus dysgalactiae subsp. equisimilis strain KNZ04 | gyrA | DNA gyrase subunit A (EC 5.99.1.3) | | | antibiotic target in susceptible species | 9293187 |
| Streptococcus dysgalactiae subsp. equisimilis strain KNZ06 | gyrA | DNA gyrase subunit A (EC 5.99.1.3) | | | antibiotic target in susceptible species | 9293187 |
| Streptococcus dysgalactiae subsp. equisimilis strain KNZ07 | gyrA | DNA gyrase subunit A (EC 5.99.1.3) | | | antibiotic target in susceptible species | 9293187 |
| Streptococcus dysgalactiae subsp. equisimilis strain KNZ10 | gyrA | DNA gyrase subunit A (EC 5.99.1.3) | | | antibiotic target in susceptible species | 9293187 |
| Streptococcus dysgalactiae subsp. equisimilis strain KNZ12 | gyrA | DNA gyrase subunit A (EC 5.99.1.3) | | | antibiotic target in susceptible species | 9293187 |
| Streptococcus dysgalactiae subsp. equisimilis strain KNZ15 | gyrA | DNA gyrase subunit A (EC 5.99.1.3) | | | antibiotic target in susceptible species | 9293187 |
| Streptococcus dysgalactiae subsp. equisimilis strain KNZ16 | gyrA | DNA gyrase subunit A (EC 5.99.1.3) | | | antibiotic target in susceptible species | 9293187 |
| Streptococcus dysgalactiae subsp. equisimilis strain NCTC10321 | gyrA | DNA gyrase subunit A (EC 5.99.1.3) | | | antibiotic target in susceptible species | 9293187 |
| Streptococcus dysgalactiae subsp. equisimilis strain NCTC11554 | gyrA | DNA gyrase subunit A (EC 5.99.1.3) | | | antibiotic target in susceptible species | 9293187 |
| Streptococcus dysgalactiae subsp. equisimilis strain NCTC11555 | gyrA | DNA gyrase subunit A (EC 5.99.1.3) | | | antibiotic target in susceptible species | 9293187 |
| Streptococcus dysgalactiae subsp. equisimilis strain NCTC11556 | gyrA | DNA gyrase subunit A (EC 5.99.1.3) | | | antibiotic target in susceptible species | 9293187 |
| Streptococcus dysgalactiae subsp. equisimilis strain NCTC11557 | gyrA | DNA gyrase subunit A (EC 5.99.1.3) | | | antibiotic target in susceptible species | 9293187 |
| Streptococcus dysgalactiae subsp. equisimilis strain NCTC11564 | gyrA | DNA gyrase subunit A (EC 5.99.1.3) | | | antibiotic target in susceptible species | 9293187 |
| Streptococcus dysgalactiae subsp. equisimilis strain NCTC11565 | gyrA | DNA gyrase subunit A (EC 5.99.1.3) | | | antibiotic target in susceptible species | 9293187 |
| Streptococcus dysgalactiae subsp. equisimilis strain NCTC11565 | gyrA | DNA gyrase subunit A (EC 5.99.1.3) | | | antibiotic resistant gene variant or mutant,fluoroquinolone resistance gene |  |
| Streptococcus dysgalactiae subsp. equisimilis strain NCTC5370 | gyrA | DNA gyrase subunit A (EC 5.99.1.3) | | | antibiotic target in susceptible species | 9293187 |
| Streptococcus dysgalactiae subsp. equisimilis strain NCTC5371 | gyrA | DNA gyrase subunit A (EC 5.99.1.3) | | | antibiotic target in susceptible species | 9293187 |
| Streptococcus dysgalactiae subsp. equisimilis strain NCTC5969 | gyrA | DNA gyrase subunit A (EC 5.99.1.3) | | | antibiotic resistant gene variant or mutant,fluoroquinolone resistance gene |  |
| Streptococcus dysgalactiae subsp. equisimilis strain NCTC5969 | gyrA | DNA gyrase subunit A (EC 5.99.1.3) | | | antibiotic target in susceptible species | 9293187 |
| Streptococcus dysgalactiae subsp. equisimilis strain NCTC6179 | gyrA | DNA gyrase subunit A (EC 5.99.1.3) | | | antibiotic target in susceptible species | 9293187 |
| Streptococcus dysgalactiae subsp. equisimilis strain NCTC6181 | gyrA | DNA gyrase subunit A (EC 5.99.1.3) | | | antibiotic target in susceptible species | 9293187 |
| Streptococcus dysgalactiae subsp. equisimilis strain NCTC6407 | gyrA | DNA gyrase subunit A (EC 5.99.1.3) | | | antibiotic target in susceptible species | 9293187 |
| Streptococcus dysgalactiae subsp. equisimilis strain NCTC7136 | gyrA | DNA gyrase subunit A (EC 5.99.1.3) | | | antibiotic target in susceptible species | 9293187 |
| Streptococcus dysgalactiae subsp. equisimilis strain NCTC8543 | gyrA | DNA gyrase subunit A (EC 5.99.1.3) | | | antibiotic target in susceptible species | 9293187 |
| Streptococcus dysgalactiae subsp. equisimilis strain NCTC8546 | gyrA | DNA gyrase subunit A (EC 5.99.1.3) | | | antibiotic target in susceptible species | 9293187 |
| Streptococcus dysgalactiae subsp. equisimilis strain NCTC9413 | gyrA | DNA gyrase subunit A (EC 5.99.1.3) | | | antibiotic target in susceptible species | 9293187 |
| Streptococcus dysgalactiae subsp. equisimilis strain NCTC9414 | gyrA | DNA gyrase subunit A (EC 5.99.1.3) | | | antibiotic target in susceptible species | 9293187 |
| Streptococcus dysgalactiae subsp. equisimilis strain NCTC9603 | gyrA | DNA gyrase subunit A (EC 5.99.1.3) | | | antibiotic target in susceptible species | 9293187 |
| Streptococcus dysgalactiae subsp. equisimilis strain SS1575 | gyrA | DNA gyrase subunit A (EC 5.99.1.3) | | | antibiotic target in susceptible species | 9293187 |
| Streptococcus dysgalactiae subsp. equisimilis strain T642 | gyrA | DNA gyrase subunit A (EC 5.99.1.3) | | | ARO:1000001,ARO:3000618 |  |
| Streptococcus dysgalactiae subsp. equisimilis strain T642 | gyrA | DNA gyrase subunit A (EC 5.99.1.3) | | | antibiotic target in susceptible species | 9293187 |
| Streptococcus dysgalactiae subsp. equisimilis strain UT_4031CC | gyrA | DNA gyrase subunit A (EC 5.99.1.3) | | | antibiotic target in susceptible species | 9293187 |
| Streptococcus dysgalactiae subsp. equisimilis strain UT_4231_KK | gyrA | DNA gyrase subunit A (EC 5.99.1.3) | | | antibiotic target in susceptible species | 9293187 |
| Streptococcus dysgalactiae subsp. equisimilis strain UT_4234_DH | gyrA | DNA gyrase subunit A (EC 5.99.1.3) | | | antibiotic target in susceptible species | 9293187 |
| Streptococcus dysgalactiae subsp. equisimilis strain UT_4241_XS | gyrA | DNA gyrase subunit A (EC 5.99.1.3) | | | antibiotic target in susceptible species | 9293187 |
| Streptococcus dysgalactiae subsp. equisimilis strain UT_4242_AB | gyrA | DNA gyrase subunit A (EC 5.99.1.3) | | | antibiotic target in susceptible species | 9293187 |
| Streptococcus dysgalactiae subsp. equisimilis strain UT_4255RC | gyrA | DNA gyrase subunit A (EC 5.99.1.3) | | | antibiotic target in susceptible species | 9293187 |
| Streptococcus dysgalactiae subsp. equisimilis strain UT_4277_BB | gyrA | DNA gyrase subunit A (EC 5.99.1.3) | | | antibiotic target in susceptible species | 9293187 |
| Streptococcus dysgalactiae subsp. equisimilis strain UT_4966_RC | gyrA | DNA gyrase subunit A (EC 5.99.1.3) | | | antibiotic target in susceptible species | 9293187 |
| Streptococcus dysgalactiae subsp. equisimilis strain UT-5345 | gyrA | DNA gyrase subunit A (EC 5.99.1.3) | | | antibiotic target in susceptible species | 9293187 |
| Streptococcus dysgalactiae subsp. equisimilis strain UT-5354 | gyrA | DNA gyrase subunit A (EC 5.99.1.3) | | | antibiotic target in susceptible species | 9293187 |
| Streptococcus dysgalactiae subsp. equisimilis strain UT-SS1069 | gyrA | DNA gyrase subunit A (EC 5.99.1.3) | | | antibiotic target in susceptible species | 9293187 |
| Streptococcus dysgalactiae subsp. equisimilis strain UT-SS957 | gyrA | DNA gyrase subunit A (EC 5.99.1.3) | | | antibiotic target in susceptible species | 9293187 |
| Streptococcus dysgalactiae subsp. equisimilis strain WCHSDSE-1 | gyrA | DNA gyrase subunit A (EC 5.99.1.3) | | | antibiotic target in susceptible species | 9293187 |
| Streptococcus dysgalactiae subsp. equisimilis SD SCDR1 | gyrB | DNA gyrase subunit B (EC 5.99.1.3) | | | antibiotic target in susceptible species | 21693461;22279180;9293187 |
| Streptococcus dysgalactiae subsp. equisimilis 167 | gyrB | DNA gyrase subunit B (EC 5.99.1.3) | | | antibiotic target in susceptible species | 21693461;22279180;9293187 |
| Streptococcus dysgalactiae subsp. equisimilis AC-2713 | gyrB | DNA gyrase subunit B (EC 5.99.1.3) | | | antibiotic target in susceptible species | 21693461;22279180;9293187 |
| Streptococcus dysgalactiae subsp. equisimilis AKSDE4288 | gyrB | DNA gyrase subunit B (EC 5.99.1.3) | | | antibiotic target in susceptible species | 21693461;22279180;9293187 |
| Streptococcus dysgalactiae subsp. equisimilis ATCC 12394 | gyrB | DNA gyrase subunit B (EC 5.99.1.3) | | | antibiotic target in susceptible species | 21693461;22279180;9293187 |
| Streptococcus dysgalactiae subsp. equisimilis ATCC 12394 | gyrB | DNA gyrase subunit B (EC 5.99.1.3) | | | antibiotic target in susceptible species | 21693461;22279180;9293187 |
| Streptococcus dysgalactiae subsp. equisimilis GGS_124 | gyrB | DNA gyrase subunit B (EC 5.99.1.3) | | | antibiotic target in susceptible species | 21693461;22279180;9293187 |
| Streptococcus dysgalactiae subsp. equisimilis RE378 | gyrB | DNA gyrase subunit B (EC 5.99.1.3) | | | antibiotic target in susceptible species | 21693461;22279180;9293187 |
| Streptococcus dysgalactiae subsp. equisimilis SK1249 | gyrB | DNA gyrase subunit B (EC 5.99.1.3) | | | antibiotic target in susceptible species | 21693461;22279180;9293187 |
| Streptococcus dysgalactiae subsp. equisimilis SK1250 | gyrB | DNA gyrase subunit B (EC 5.99.1.3) | | | antibiotic target in susceptible species | 21693461;22279180;9293187 |
| Streptococcus dysgalactiae subsp. equisimilis strain ASDSE_96 | gyrB | DNA gyrase subunit B (EC 5.99.1.3) | | | antibiotic target in susceptible species | 21693461;22279180;9293187 |
| Streptococcus dysgalactiae subsp. equisimilis strain ASDSE_99 | gyrB | DNA gyrase subunit B (EC 5.99.1.3) | | | antibiotic target in susceptible species | 21693461;22279180;9293187 |
| Streptococcus dysgalactiae subsp. equisimilis strain C161L1 | gyrB | DNA gyrase subunit B (EC 5.99.1.3) | | | antibiotic target in susceptible species | 21693461;22279180;9293187 |
| Streptococcus dysgalactiae subsp. equisimilis strain KNZ01 | gyrB | DNA gyrase subunit B (EC 5.99.1.3) | | | antibiotic target in susceptible species | 21693461;22279180;9293187 |
| Streptococcus dysgalactiae subsp. equisimilis strain KNZ01 | gyrB | DNA gyrase subunit B (EC 5.99.1.3) | | | antibiotic target in susceptible species | 21693461;22279180;9293187 |
| Streptococcus dysgalactiae subsp. equisimilis strain KNZ03 | gyrB | DNA gyrase subunit B (EC 5.99.1.3) | | | antibiotic target in susceptible species | 21693461;22279180;9293187 |
| Streptococcus dysgalactiae subsp. equisimilis strain KNZ03 | gyrB | DNA gyrase subunit B (EC 5.99.1.3) | | | antibiotic target in susceptible species | 21693461;22279180;9293187 |
| Streptococcus dysgalactiae subsp. equisimilis strain KNZ04 | gyrB | DNA gyrase subunit B (EC 5.99.1.3) | | | antibiotic target in susceptible species | 21693461;22279180;9293187 |
| Streptococcus dysgalactiae subsp. equisimilis strain KNZ06 | gyrB | DNA gyrase subunit B (EC 5.99.1.3) | | | antibiotic target in susceptible species | 21693461;22279180;9293187 |
| Streptococcus dysgalactiae subsp. equisimilis strain KNZ07 | gyrB | DNA gyrase subunit B (EC 5.99.1.3) | | | antibiotic target in susceptible species | 21693461;22279180;9293187 |
| Streptococcus dysgalactiae subsp. equisimilis strain KNZ10 | gyrB | DNA gyrase subunit B (EC 5.99.1.3) | | | antibiotic target in susceptible species | 21693461;22279180;9293187 |
| Streptococcus dysgalactiae subsp. equisimilis strain KNZ12 | gyrB | DNA gyrase subunit B (EC 5.99.1.3) | | | antibiotic target in susceptible species | 21693461;22279180;9293187 |
| Streptococcus dysgalactiae subsp. equisimilis strain KNZ15 | gyrB | DNA gyrase subunit B (EC 5.99.1.3) | | | antibiotic target in susceptible species | 21693461;22279180;9293187 |
| Streptococcus dysgalactiae subsp. equisimilis strain KNZ16 | gyrB | DNA gyrase subunit B (EC 5.99.1.3) | | | antibiotic target in susceptible species | 21693461;22279180;9293187 |
| Streptococcus dysgalactiae subsp. equisimilis strain NCTC10321 | gyrB | DNA gyrase subunit B (EC 5.99.1.3) | | | antibiotic target in susceptible species | 21693461;22279180;9293187 |
| Streptococcus dysgalactiae subsp. equisimilis strain NCTC11554 | gyrB | DNA gyrase subunit B (EC 5.99.1.3) | | | antibiotic target in susceptible species | 21693461;22279180;9293187 |
| Streptococcus dysgalactiae subsp. equisimilis strain NCTC11555 | gyrB | DNA gyrase subunit B (EC 5.99.1.3) | | | antibiotic target in susceptible species | 21693461;22279180;9293187 |
| Streptococcus dysgalactiae subsp. equisimilis strain NCTC11556 | gyrB | DNA gyrase subunit B (EC 5.99.1.3) | | | antibiotic target in susceptible species | 21693461;22279180;9293187 |
| Streptococcus dysgalactiae subsp. equisimilis strain NCTC11557 | gyrB | DNA gyrase subunit B (EC 5.99.1.3) | | | antibiotic target in susceptible species | 21693461;22279180;9293187 |
| Streptococcus dysgalactiae subsp. equisimilis strain NCTC11564 | gyrB | DNA gyrase subunit B (EC 5.99.1.3) | | | antibiotic target in susceptible species | 21693461;22279180;9293187 |
| Streptococcus dysgalactiae subsp. equisimilis strain NCTC11565 | gyrB | DNA gyrase subunit B (EC 5.99.1.3) | | | antibiotic target in susceptible species | 21693461;22279180;9293187 |
| Streptococcus dysgalactiae subsp. equisimilis strain NCTC5370 | gyrB | DNA gyrase subunit B (EC 5.99.1.3) | | | antibiotic target in susceptible species | 21693461;22279180;9293187 |
| Streptococcus dysgalactiae subsp. equisimilis strain NCTC5371 | gyrB | DNA gyrase subunit B (EC 5.99.1.3) | | | antibiotic target in susceptible species | 21693461;22279180;9293187 |
| Streptococcus dysgalactiae subsp. equisimilis strain NCTC5969 | gyrB | DNA gyrase subunit B (EC 5.99.1.3) | | | antibiotic target in susceptible species | 21693461;22279180;9293187 |
| Streptococcus dysgalactiae subsp. equisimilis strain NCTC5969 | gyrB | DNA gyrase subunit B (EC 5.99.1.3) | | | aminocoumarin resistance gene,antibiotic resistant gene variant or mutant | 21693461;22279180;9293187 |
| Streptococcus dysgalactiae subsp. equisimilis strain NCTC6179 | gyrB | DNA gyrase subunit B (EC 5.99.1.3) | | | antibiotic target in susceptible species | 21693461;22279180;9293187 |
| Streptococcus dysgalactiae subsp. equisimilis strain NCTC6181 | gyrB | DNA gyrase subunit B (EC 5.99.1.3) | | | antibiotic target in susceptible species | 21693461;22279180;9293187 |
| Streptococcus dysgalactiae subsp. equisimilis strain NCTC6407 | gyrB | DNA gyrase subunit B (EC 5.99.1.3) | | | antibiotic target in susceptible species | 21693461;22279180;9293187 |
| Streptococcus dysgalactiae subsp. equisimilis strain NCTC7136 | gyrB | DNA gyrase subunit B (EC 5.99.1.3) | | | antibiotic target in susceptible species | 21693461;22279180;9293187 |
| Streptococcus dysgalactiae subsp. equisimilis strain NCTC8543 | gyrB | DNA gyrase subunit B (EC 5.99.1.3) | | | antibiotic target in susceptible species | 21693461;22279180;9293187 |
| Streptococcus dysgalactiae subsp. equisimilis strain NCTC8546 | gyrB | DNA gyrase subunit B (EC 5.99.1.3) | | | antibiotic target in susceptible species | 21693461;22279180;9293187 |
| Streptococcus dysgalactiae subsp. equisimilis strain NCTC9413 | gyrB | DNA gyrase subunit B (EC 5.99.1.3) | | | antibiotic target in susceptible species | 21693461;22279180;9293187 |
| Streptococcus dysgalactiae subsp. equisimilis strain NCTC9414 | gyrB | DNA gyrase subunit B (EC 5.99.1.3) | | | antibiotic target in susceptible species | 21693461;22279180;9293187 |
| Streptococcus dysgalactiae subsp. equisimilis strain NCTC9603 | gyrB | DNA gyrase subunit B (EC 5.99.1.3) | | | antibiotic target in susceptible species | 21693461;22279180;9293187 |
| Streptococcus dysgalactiae subsp. equisimilis strain SS1575 | gyrB | DNA gyrase subunit B (EC 5.99.1.3) | | | antibiotic target in susceptible species | 21693461;22279180;9293187 |
| Streptococcus dysgalactiae subsp. equisimilis strain T642 | gyrB | DNA gyrase subunit B (EC 5.99.1.3) | | | antibiotic target in susceptible species | 21693461;22279180;9293187 |
| Streptococcus dysgalactiae subsp. equisimilis strain UT_4031CC | gyrB | DNA gyrase subunit B (EC 5.99.1.3) | | | antibiotic target in susceptible species | 21693461;22279180;9293187 |
| Streptococcus dysgalactiae subsp. equisimilis strain UT_4231_KK | gyrB | DNA gyrase subunit B (EC 5.99.1.3) | | | antibiotic target in susceptible species | 21693461;22279180;9293187 |
| Streptococcus dysgalactiae subsp. equisimilis strain UT_4234_DH | gyrB | DNA gyrase subunit B (EC 5.99.1.3) | | | antibiotic target in susceptible species | 21693461;22279180;9293187 |
| Streptococcus dysgalactiae subsp. equisimilis strain UT_4241_XS | gyrB | DNA gyrase subunit B (EC 5.99.1.3) | | | antibiotic target in susceptible species | 21693461;22279180;9293187 |
| Streptococcus dysgalactiae subsp. equisimilis strain UT_4242_AB | gyrB | DNA gyrase subunit B (EC 5.99.1.3) | | | antibiotic target in susceptible species | 21693461;22279180;9293187 |
| Streptococcus dysgalactiae subsp. equisimilis strain UT_4255RC | gyrB | DNA gyrase subunit B (EC 5.99.1.3) | | | antibiotic target in susceptible species | 21693461;22279180;9293187 |
| Streptococcus dysgalactiae subsp. equisimilis strain UT_4277_BB | gyrB | DNA gyrase subunit B (EC 5.99.1.3) | | | antibiotic target in susceptible species | 21693461;22279180;9293187 |
| Streptococcus dysgalactiae subsp. equisimilis strain UT_4966_RC | gyrB | DNA gyrase subunit B (EC 5.99.1.3) | | | antibiotic target in susceptible species | 21693461;22279180;9293187 |
| Streptococcus dysgalactiae subsp. equisimilis strain UT-5345 | gyrB | DNA gyrase subunit B (EC 5.99.1.3) | | | antibiotic target in susceptible species | 21693461;22279180;9293187 |
| Streptococcus dysgalactiae subsp. equisimilis strain UT-5354 | gyrB | DNA gyrase subunit B (EC 5.99.1.3) | | | antibiotic target in susceptible species | 21693461;22279180;9293187 |
| Streptococcus dysgalactiae subsp. equisimilis strain UT-SS1069 | gyrB | DNA gyrase subunit B (EC 5.99.1.3) | | | antibiotic target in susceptible species | 21693461;22279180;9293187 |
| Streptococcus dysgalactiae subsp. equisimilis strain UT-SS1069 | gyrB | DNA gyrase subunit B (EC 5.99.1.3) | | | antibiotic target in susceptible species | 21693461;22279180;9293187 |
| Streptococcus dysgalactiae subsp. equisimilis strain UT-SS957 | gyrB | DNA gyrase subunit B (EC 5.99.1.3) | | | antibiotic target in susceptible species | 21693461;22279180;9293187 |
| Streptococcus dysgalactiae subsp. equisimilis strain WCHSDSE-1 | gyrB | DNA gyrase subunit B (EC 5.99.1.3) | | | antibiotic target in susceptible species | 21693461;22279180;9293187 |
| Streptococcus dysgalactiae subsp. equisimilis SD SCDR1 | Iso-tRNA | Isoleucyl-tRNA synthetase (EC 6.1.1.5) | | | antibiotic target in susceptible species | 7929087 |
| Streptococcus dysgalactiae subsp. equisimilis 167 | Iso-tRNA | Isoleucyl-tRNA synthetase (EC 6.1.1.5) | | | antibiotic target in susceptible species | 7929087 |
| Streptococcus dysgalactiae subsp. equisimilis AC-2713 | ileS | Isoleucyl-tRNA synthetase (EC 6.1.1.5) | | | antibiotic target in susceptible species | 7929087 |
| Streptococcus dysgalactiae subsp. equisimilis AKSDE4288 | ileS | Isoleucyl-tRNA synthetase (EC 6.1.1.5) | | | antibiotic target in susceptible species | 7929087 |
| Streptococcus dysgalactiae subsp. equisimilis ATCC 12394 | ileS | Isoleucyl-tRNA synthetase (EC 6.1.1.5) | | | antibiotic target in susceptible species | 7929087 |
| Streptococcus dysgalactiae subsp. equisimilis GGS_124 | ileS | Isoleucyl-tRNA synthetase (EC 6.1.1.5) | | | antibiotic target in susceptible species | 7929087 |
| Streptococcus dysgalactiae subsp. equisimilis RE378 | ileS | Isoleucyl-tRNA synthetase (EC 6.1.1.5) | | | antibiotic target in susceptible species | 7929087 |
| Streptococcus dysgalactiae subsp. equisimilis SK1249 | ileS | Isoleucyl-tRNA synthetase (EC 6.1.1.5) | | | antibiotic target in susceptible species | 7929087 |
| Streptococcus dysgalactiae subsp. equisimilis SK1250 | ileS | Isoleucyl-tRNA synthetase (EC 6.1.1.5) | | | antibiotic target in susceptible species | 7929087 |
| Streptococcus dysgalactiae subsp. equisimilis strain ASDSE_96 | ileS | Isoleucyl-tRNA synthetase (EC 6.1.1.5) | | | antibiotic target in susceptible species | 7929087 |
| Streptococcus dysgalactiae subsp. equisimilis strain ASDSE_99 | ileS | Isoleucyl-tRNA synthetase (EC 6.1.1.5) | | | antibiotic target in susceptible species | 7929087 |
| Streptococcus dysgalactiae subsp. equisimilis strain C161L1 | ileS | Isoleucyl-tRNA synthetase (EC 6.1.1.5) | | | antibiotic target in susceptible species | 7929087 |
| Streptococcus dysgalactiae subsp. equisimilis strain KNZ01 | Iso-tRNA | Isoleucyl-tRNA synthetase (EC 6.1.1.5) | | | antibiotic target in susceptible species | 7929087 |
| Streptococcus dysgalactiae subsp. equisimilis strain KNZ03 | Iso-tRNA | Isoleucyl-tRNA synthetase (EC 6.1.1.5) | | | antibiotic target in susceptible species | 7929087 |
| Streptococcus dysgalactiae subsp. equisimilis strain KNZ04 | Iso-tRNA | Isoleucyl-tRNA synthetase (EC 6.1.1.5) | | | antibiotic target in susceptible species | 7929087 |
| Streptococcus dysgalactiae subsp. equisimilis strain KNZ06 | Iso-tRNA | Isoleucyl-tRNA synthetase (EC 6.1.1.5) | | | antibiotic target in susceptible species | 7929087 |
| Streptococcus dysgalactiae subsp. equisimilis strain KNZ07 | Iso-tRNA | Isoleucyl-tRNA synthetase (EC 6.1.1.5) | | | antibiotic target in susceptible species | 7929087 |
| Streptococcus dysgalactiae subsp. equisimilis strain KNZ10 | Iso-tRNA | Isoleucyl-tRNA synthetase (EC 6.1.1.5) | | | antibiotic target in susceptible species | 7929087 |
| Streptococcus dysgalactiae subsp. equisimilis strain KNZ12 | Iso-tRNA | Isoleucyl-tRNA synthetase (EC 6.1.1.5) | | | antibiotic target in susceptible species | 7929087 |
| Streptococcus dysgalactiae subsp. equisimilis strain KNZ15 | Iso-tRNA | Isoleucyl-tRNA synthetase (EC 6.1.1.5) | | | antibiotic target in susceptible species | 7929087 |
| Streptococcus dysgalactiae subsp. equisimilis strain KNZ16 | Iso-tRNA | Isoleucyl-tRNA synthetase (EC 6.1.1.5) | | | antibiotic target in susceptible species | 7929087 |
| Streptococcus dysgalactiae subsp. equisimilis strain NCTC10321 | Iso-tRNA | Isoleucyl-tRNA synthetase (EC 6.1.1.5) | | | antibiotic target in susceptible species | 7929087 |
| Streptococcus dysgalactiae subsp. equisimilis strain NCTC11554 | Iso-tRNA | Isoleucyl-tRNA synthetase (EC 6.1.1.5) | | | antibiotic target in susceptible species | 7929087 |
| Streptococcus dysgalactiae subsp. equisimilis strain NCTC11555 | Iso-tRNA | Isoleucyl-tRNA synthetase (EC 6.1.1.5) | | | antibiotic target in susceptible species | 7929087 |
| Streptococcus dysgalactiae subsp. equisimilis strain NCTC11556 | Iso-tRNA | Isoleucyl-tRNA synthetase (EC 6.1.1.5) | | | antibiotic target in susceptible species | 7929087 |
| Streptococcus dysgalactiae subsp. equisimilis strain NCTC11557 | Iso-tRNA | Isoleucyl-tRNA synthetase (EC 6.1.1.5) | | | antibiotic target in susceptible species | 7929087 |
| Streptococcus dysgalactiae subsp. equisimilis strain NCTC11564 | Iso-tRNA | Isoleucyl-tRNA synthetase (EC 6.1.1.5) | | | antibiotic target in susceptible species | 7929087 |
| Streptococcus dysgalactiae subsp. equisimilis strain NCTC11565 | Iso-tRNA | Isoleucyl-tRNA synthetase (EC 6.1.1.5) | | | antibiotic target in susceptible species | 7929087 |
| Streptococcus dysgalactiae subsp. equisimilis strain NCTC5370 | ileS | Isoleucyl-tRNA synthetase (EC 6.1.1.5) | | | antibiotic target in susceptible species | 7929087 |
| Streptococcus dysgalactiae subsp. equisimilis strain NCTC5371 | ileS | Isoleucyl-tRNA synthetase (EC 6.1.1.5) | | | antibiotic target in susceptible species | 7929087 |
| Streptococcus dysgalactiae subsp. equisimilis strain NCTC5969 | Iso-tRNA | Isoleucyl-tRNA synthetase (EC 6.1.1.5) | | | antibiotic target in susceptible species | 7929087 |
| Streptococcus dysgalactiae subsp. equisimilis strain NCTC6179 | ileS | Isoleucyl-tRNA synthetase (EC 6.1.1.5) | | | antibiotic target in susceptible species | 7929087 |
| Streptococcus dysgalactiae subsp. equisimilis strain NCTC6181 | Iso-tRNA | Isoleucyl-tRNA synthetase (EC 6.1.1.5) | | | antibiotic target in susceptible species | 7929087 |
| Streptococcus dysgalactiae subsp. equisimilis strain NCTC6407 | Iso-tRNA | Isoleucyl-tRNA synthetase (EC 6.1.1.5) | | | antibiotic target in susceptible species | 7929087 |
| Streptococcus dysgalactiae subsp. equisimilis strain NCTC7136 | ileS | Isoleucyl-tRNA synthetase (EC 6.1.1.5) | | | antibiotic target in susceptible species | 7929087 |
| Streptococcus dysgalactiae subsp. equisimilis strain NCTC8543 | Iso-tRNA | Isoleucyl-tRNA synthetase (EC 6.1.1.5) | | | antibiotic target in susceptible species | 7929087 |
| Streptococcus dysgalactiae subsp. equisimilis strain NCTC8546 | Iso-tRNA | Isoleucyl-tRNA synthetase (EC 6.1.1.5) | | | antibiotic target in susceptible species | 7929087 |
| Streptococcus dysgalactiae subsp. equisimilis strain NCTC9413 | Iso-tRNA | Isoleucyl-tRNA synthetase (EC 6.1.1.5) | | | antibiotic target in susceptible species | 7929087 |
| Streptococcus dysgalactiae subsp. equisimilis strain NCTC9414 | ileS | Isoleucyl-tRNA synthetase (EC 6.1.1.5) | | | antibiotic target in susceptible species | 7929087 |
| Streptococcus dysgalactiae subsp. equisimilis strain NCTC9603 | Iso-tRNA | Isoleucyl-tRNA synthetase (EC 6.1.1.5) | | | antibiotic target in susceptible species | 7929087 |
| Streptococcus dysgalactiae subsp. equisimilis strain SS1575 | ileS | Isoleucyl-tRNA synthetase (EC 6.1.1.5) | | | antibiotic target in susceptible species | 7929087 |
| Streptococcus dysgalactiae subsp. equisimilis strain T642 | ileS | Isoleucyl-tRNA synthetase (EC 6.1.1.5) | | | antibiotic target in susceptible species | 7929087 |
| Streptococcus dysgalactiae subsp. equisimilis strain UT_4031CC | ileS | Isoleucyl-tRNA synthetase (EC 6.1.1.5) | | | antibiotic target in susceptible species | 7929087 |
| Streptococcus dysgalactiae subsp. equisimilis strain UT_4231_KK | ileS | Isoleucyl-tRNA synthetase (EC 6.1.1.5) | | | antibiotic target in susceptible species | 7929087 |
| Streptococcus dysgalactiae subsp. equisimilis strain UT_4234_DH | ileS | Isoleucyl-tRNA synthetase (EC 6.1.1.5) | | | antibiotic target in susceptible species | 7929087 |
| Streptococcus dysgalactiae subsp. equisimilis strain UT_4241_XS | ileS | Isoleucyl-tRNA synthetase (EC 6.1.1.5) | | | antibiotic target in susceptible species | 7929087 |
| Streptococcus dysgalactiae subsp. equisimilis strain UT_4242_AB | ileS | Isoleucyl-tRNA synthetase (EC 6.1.1.5) | | | antibiotic target in susceptible species | 7929087 |
| Streptococcus dysgalactiae subsp. equisimilis strain UT_4255RC | ileS | Isoleucyl-tRNA synthetase (EC 6.1.1.5) | | | antibiotic target in susceptible species | 7929087 |
| Streptococcus dysgalactiae subsp. equisimilis strain UT_4277_BB | ileS | Isoleucyl-tRNA synthetase (EC 6.1.1.5) | | | antibiotic target in susceptible species | 7929087 |
| Streptococcus dysgalactiae subsp. equisimilis strain UT_4966_RC | ileS | Isoleucyl-tRNA synthetase (EC 6.1.1.5) | | | antibiotic target in susceptible species | 7929087 |
| Streptococcus dysgalactiae subsp. equisimilis strain UT-5345 | ileS | Isoleucyl-tRNA synthetase (EC 6.1.1.5) | | | antibiotic target in susceptible species | 7929087 |
| Streptococcus dysgalactiae subsp. equisimilis strain UT-5354 | ileS | Isoleucyl-tRNA synthetase (EC 6.1.1.5) | | | antibiotic target in susceptible species | 7929087 |
| Streptococcus dysgalactiae subsp. equisimilis strain UT-SS1069 | ileS | Isoleucyl-tRNA synthetase (EC 6.1.1.5) | | | antibiotic target in susceptible species | 7929087 |
| Streptococcus dysgalactiae subsp. equisimilis strain UT-SS957 | ileS | Isoleucyl-tRNA synthetase (EC 6.1.1.5) | | | antibiotic target in susceptible species | 7929087 |
| Streptococcus dysgalactiae subsp. equisimilis strain WCHSDSE-1 | ileS | Isoleucyl-tRNA synthetase (EC 6.1.1.5) | | | antibiotic target in susceptible species | 7929087 |
| Streptococcus dysgalactiae subsp. equisimilis SD SCDR1 | LiaF | Membrane protein LiaF(VraT), specific inhibitor of LiaRS(VraRS) signaling pathway | | | regulator modulating expression of antibiotic resistance genes | 21899450;26020679 |
| Streptococcus dysgalactiae subsp. equisimilis 167 | yvqF | Membrane protein LiaF(VraT), specific inhibitor of LiaRS(VraRS) signaling pathway | | | regulator modulating expression of antibiotic resistance genes | 21899450;26020679 |
| Streptococcus dysgalactiae subsp. equisimilis AC-2713 | yvqF | Membrane protein LiaF(VraT), specific inhibitor of LiaRS(VraRS) signaling pathway | | | regulator modulating expression of antibiotic resistance genes | 21899450;26020679 |
| Streptococcus dysgalactiae subsp. equisimilis AKSDE4288 | LiaF | Membrane protein LiaF(VraT), specific inhibitor of LiaRS(VraRS) signaling pathway | | | regulator modulating expression of antibiotic resistance genes | 21899450;26020679 |
| Streptococcus dysgalactiae subsp. equisimilis ATCC 12394 | LiaF | Membrane protein LiaF(VraT), specific inhibitor of LiaRS(VraRS) signaling pathway | | | regulator modulating expression of antibiotic resistance genes | 21899450;26020679 |
| Streptococcus dysgalactiae subsp. equisimilis GGS_124 | yvqF | Membrane protein LiaF(VraT), specific inhibitor of LiaRS(VraRS) signaling pathway | | | regulator modulating expression of antibiotic resistance genes | 21899450;26020679 |
| Streptococcus dysgalactiae subsp. equisimilis RE378 | yvqF | Membrane protein LiaF(VraT), specific inhibitor of LiaRS(VraRS) signaling pathway | | | regulator modulating expression of antibiotic resistance genes | 21899450;26020679 |
| Streptococcus dysgalactiae subsp. equisimilis SK1249 | LiaF | Membrane protein LiaF(VraT), specific inhibitor of LiaRS(VraRS) signaling pathway | | | regulator modulating expression of antibiotic resistance genes | 21899450;26020679 |
| Streptococcus dysgalactiae subsp. equisimilis SK1250 | LiaF | Membrane protein LiaF(VraT), specific inhibitor of LiaRS(VraRS) signaling pathway | | | regulator modulating expression of antibiotic resistance genes | 21899450;26020679 |
| Streptococcus dysgalactiae subsp. equisimilis strain ASDSE_96 | LiaF | Membrane protein LiaF(VraT), specific inhibitor of LiaRS(VraRS) signaling pathway | | | regulator modulating expression of antibiotic resistance genes | 21899450;26020679 |
| Streptococcus dysgalactiae subsp. equisimilis strain ASDSE_99 | LiaF | Membrane protein LiaF(VraT), specific inhibitor of LiaRS(VraRS) signaling pathway | | | regulator modulating expression of antibiotic resistance genes | 21899450;26020679 |
| Streptococcus dysgalactiae subsp. equisimilis strain C161L1 | LiaF | Membrane protein LiaF(VraT), specific inhibitor of LiaRS(VraRS) signaling pathway | | | regulator modulating expression of antibiotic resistance genes | 21899450;26020679 |
| Streptococcus dysgalactiae subsp. equisimilis strain KNZ01 | LiaF | Membrane protein LiaF(VraT), specific inhibitor of LiaRS(VraRS) signaling pathway | | | regulator modulating expression of antibiotic resistance genes | 21899450;26020679 |
| Streptococcus dysgalactiae subsp. equisimilis strain KNZ03 | LiaF | Membrane protein LiaF(VraT), specific inhibitor of LiaRS(VraRS) signaling pathway | | | regulator modulating expression of antibiotic resistance genes | 21899450;26020679 |
| Streptococcus dysgalactiae subsp. equisimilis strain KNZ04 | LiaF | Membrane protein LiaF(VraT), specific inhibitor of LiaRS(VraRS) signaling pathway | | | regulator modulating expression of antibiotic resistance genes | 21899450;26020679 |
| Streptococcus dysgalactiae subsp. equisimilis strain KNZ06 | LiaF | Membrane protein LiaF(VraT), specific inhibitor of LiaRS(VraRS) signaling pathway | | | regulator modulating expression of antibiotic resistance genes | 21899450;26020679 |
| Streptococcus dysgalactiae subsp. equisimilis strain KNZ07 | LiaF | Membrane protein LiaF(VraT), specific inhibitor of LiaRS(VraRS) signaling pathway | | | regulator modulating expression of antibiotic resistance genes | 21899450;26020679 |
| Streptococcus dysgalactiae subsp. equisimilis strain KNZ10 | LiaF | Membrane protein LiaF(VraT), specific inhibitor of LiaRS(VraRS) signaling pathway | | | regulator modulating expression of antibiotic resistance genes | 21899450;26020679 |
| Streptococcus dysgalactiae subsp. equisimilis strain KNZ12 | LiaF | Membrane protein LiaF(VraT), specific inhibitor of LiaRS(VraRS) signaling pathway | | | regulator modulating expression of antibiotic resistance genes | 21899450;26020679 |
| Streptococcus dysgalactiae subsp. equisimilis strain KNZ15 | LiaF | Membrane protein LiaF(VraT), specific inhibitor of LiaRS(VraRS) signaling pathway | | | regulator modulating expression of antibiotic resistance genes | 21899450;26020679 |
| Streptococcus dysgalactiae subsp. equisimilis strain KNZ16 | LiaF | Membrane protein LiaF(VraT), specific inhibitor of LiaRS(VraRS) signaling pathway | | | regulator modulating expression of antibiotic resistance genes | 21899450;26020679 |
| Streptococcus dysgalactiae subsp. equisimilis strain NCTC10321 | LiaF | Membrane protein LiaF(VraT), specific inhibitor of LiaRS(VraRS) signaling pathway | | | regulator modulating expression of antibiotic resistance genes | 21899450;26020679 |
| Streptococcus dysgalactiae subsp. equisimilis strain NCTC11554 | LiaF | Membrane protein LiaF(VraT), specific inhibitor of LiaRS(VraRS) signaling pathway | | | regulator modulating expression of antibiotic resistance genes | 21899450;26020679 |
| Streptococcus dysgalactiae subsp. equisimilis strain NCTC11555 | LiaF | Membrane protein LiaF(VraT), specific inhibitor of LiaRS(VraRS) signaling pathway | | | regulator modulating expression of antibiotic resistance genes | 21899450;26020679 |
| Streptococcus dysgalactiae subsp. equisimilis strain NCTC11556 | LiaF | Membrane protein LiaF(VraT), specific inhibitor of LiaRS(VraRS) signaling pathway | | | regulator modulating expression of antibiotic resistance genes | 21899450;26020679 |
| Streptococcus dysgalactiae subsp. equisimilis strain NCTC11557 | LiaF | Membrane protein LiaF(VraT), specific inhibitor of LiaRS(VraRS) signaling pathway | | | regulator modulating expression of antibiotic resistance genes | 21899450;26020679 |
| Streptococcus dysgalactiae subsp. equisimilis strain NCTC11564 | LiaF | Membrane protein LiaF(VraT), specific inhibitor of LiaRS(VraRS) signaling pathway | | | regulator modulating expression of antibiotic resistance genes | 21899450;26020679 |
| Streptococcus dysgalactiae subsp. equisimilis strain NCTC5370 | yvqF | Membrane protein LiaF(VraT), specific inhibitor of LiaRS(VraRS) signaling pathway | | | regulator modulating expression of antibiotic resistance genes | 21899450;26020679 |
| Streptococcus dysgalactiae subsp. equisimilis strain NCTC5371 | yvqF | Membrane protein LiaF(VraT), specific inhibitor of LiaRS(VraRS) signaling pathway | | | regulator modulating expression of antibiotic resistance genes | 21899450;26020679 |
| Streptococcus dysgalactiae subsp. equisimilis strain NCTC5969 | LiaF | Membrane protein LiaF(VraT), specific inhibitor of LiaRS(VraRS) signaling pathway | | | regulator modulating expression of antibiotic resistance genes | 21899450;26020679 |
| Streptococcus dysgalactiae subsp. equisimilis strain NCTC6179 | yvqF | Membrane protein LiaF(VraT), specific inhibitor of LiaRS(VraRS) signaling pathway | | | regulator modulating expression of antibiotic resistance genes | 21899450;26020679 |
| Streptococcus dysgalactiae subsp. equisimilis strain NCTC6181 | LiaF | Membrane protein LiaF(VraT), specific inhibitor of LiaRS(VraRS) signaling pathway | | | regulator modulating expression of antibiotic resistance genes | 21899450;26020679 |
| Streptococcus dysgalactiae subsp. equisimilis strain NCTC6407 | LiaF | Membrane protein LiaF(VraT), specific inhibitor of LiaRS(VraRS) signaling pathway | | | regulator modulating expression of antibiotic resistance genes | 21899450;26020679 |
| Streptococcus dysgalactiae subsp. equisimilis strain NCTC7136 | yvqF | Membrane protein LiaF(VraT), specific inhibitor of LiaRS(VraRS) signaling pathway | | | regulator modulating expression of antibiotic resistance genes | 21899450;26020679 |
| Streptococcus dysgalactiae subsp. equisimilis strain NCTC8543 | LiaF | Membrane protein LiaF(VraT), specific inhibitor of LiaRS(VraRS) signaling pathway | | | regulator modulating expression of antibiotic resistance genes | 21899450;26020679 |
| Streptococcus dysgalactiae subsp. equisimilis strain NCTC8546 | LiaF | Membrane protein LiaF(VraT), specific inhibitor of LiaRS(VraRS) signaling pathway | | | regulator modulating expression of antibiotic resistance genes | 21899450;26020679 |
| Streptococcus dysgalactiae subsp. equisimilis strain NCTC9413 | LiaF | Membrane protein LiaF(VraT), specific inhibitor of LiaRS(VraRS) signaling pathway | | | regulator modulating expression of antibiotic resistance genes | 21899450;26020679 |
| Streptococcus dysgalactiae subsp. equisimilis strain NCTC9414 | yvqF | Membrane protein LiaF(VraT), specific inhibitor of LiaRS(VraRS) signaling pathway | | | regulator modulating expression of antibiotic resistance genes | 21899450;26020679 |
| Streptococcus dysgalactiae subsp. equisimilis strain NCTC9603 | LiaF | Membrane protein LiaF(VraT), specific inhibitor of LiaRS(VraRS) signaling pathway | | | regulator modulating expression of antibiotic resistance genes | 21899450;26020679 |
| Streptococcus dysgalactiae subsp. equisimilis strain SS1575 | LiaF | Membrane protein LiaF(VraT), specific inhibitor of LiaRS(VraRS) signaling pathway | | | regulator modulating expression of antibiotic resistance genes | 21899450;26020679 |
| Streptococcus dysgalactiae subsp. equisimilis strain T642 | LiaF | Membrane protein LiaF(VraT), specific inhibitor of LiaRS(VraRS) signaling pathway | | | regulator modulating expression of antibiotic resistance genes | 21899450;26020679 |
| Streptococcus dysgalactiae subsp. equisimilis strain UT_4031CC | LiaF | Membrane protein LiaF(VraT), specific inhibitor of LiaRS(VraRS) signaling pathway | | | regulator modulating expression of antibiotic resistance genes | 21899450;26020679 |
| Streptococcus dysgalactiae subsp. equisimilis strain UT_4231_KK | LiaF | Membrane protein LiaF(VraT), specific inhibitor of LiaRS(VraRS) signaling pathway | | | regulator modulating expression of antibiotic resistance genes | 21899450;26020679 |
| Streptococcus dysgalactiae subsp. equisimilis strain UT_4234_DH | LiaF | Membrane protein LiaF(VraT), specific inhibitor of LiaRS(VraRS) signaling pathway | | | regulator modulating expression of antibiotic resistance genes | 21899450;26020679 |
| Streptococcus dysgalactiae subsp. equisimilis strain UT_4241_XS | LiaF | Membrane protein LiaF(VraT), specific inhibitor of LiaRS(VraRS) signaling pathway | | | regulator modulating expression of antibiotic resistance genes | 21899450;26020679 |
| Streptococcus dysgalactiae subsp. equisimilis strain UT_4242_AB | LiaF | Membrane protein LiaF(VraT), specific inhibitor of LiaRS(VraRS) signaling pathway | | | regulator modulating expression of antibiotic resistance genes | 21899450;26020679 |
| Streptococcus dysgalactiae subsp. equisimilis strain UT_4255RC | LiaF | Membrane protein LiaF(VraT), specific inhibitor of LiaRS(VraRS) signaling pathway | | | regulator modulating expression of antibiotic resistance genes | 21899450;26020679 |
| Streptococcus dysgalactiae subsp. equisimilis strain UT_4277_BB | LiaF | Membrane protein LiaF(VraT), specific inhibitor of LiaRS(VraRS) signaling pathway | | | regulator modulating expression of antibiotic resistance genes | 21899450;26020679 |
| Streptococcus dysgalactiae subsp. equisimilis strain UT_4966_RC | LiaF | Membrane protein LiaF(VraT), specific inhibitor of LiaRS(VraRS) signaling pathway | | | regulator modulating expression of antibiotic resistance genes | 21899450;26020679 |
| Streptococcus dysgalactiae subsp. equisimilis strain UT-5345 | LiaF | Membrane protein LiaF(VraT), specific inhibitor of LiaRS(VraRS) signaling pathway | | | regulator modulating expression of antibiotic resistance genes | 21899450;26020679 |
| Streptococcus dysgalactiae subsp. equisimilis strain UT-5354 | LiaF | Membrane protein LiaF(VraT), specific inhibitor of LiaRS(VraRS) signaling pathway | | | regulator modulating expression of antibiotic resistance genes | 21899450;26020679 |
| Streptococcus dysgalactiae subsp. equisimilis strain UT-SS1069 | LiaF | Membrane protein LiaF(VraT), specific inhibitor of LiaRS(VraRS) signaling pathway | | | regulator modulating expression of antibiotic resistance genes | 21899450;26020679 |
| Streptococcus dysgalactiae subsp. equisimilis strain UT-SS957 | LiaF | Membrane protein LiaF(VraT), specific inhibitor of LiaRS(VraRS) signaling pathway | | | regulator modulating expression of antibiotic resistance genes | 21899450;26020679 |
| Streptococcus dysgalactiae subsp. equisimilis strain WCHSDSE-1 | LiaF | Membrane protein LiaF(VraT), specific inhibitor of LiaRS(VraRS) signaling pathway | | | regulator modulating expression of antibiotic resistance genes | 21899450;26020679 |
| Streptococcus dysgalactiae subsp. equisimilis SD SCDR1 | lmrP | Multidrug resistance protein ErmB | | | efflux pump conferring antibiotic resistance | 11577166 |
| Streptococcus dysgalactiae subsp. equisimilis 167 | lmrP | Multidrug resistance protein ErmB | | | efflux pump conferring antibiotic resistance | 11577166 |
| Streptococcus dysgalactiae subsp. equisimilis AC-2713 | lmrP | Multidrug resistance protein ErmB | | | efflux pump conferring antibiotic resistance | 11577166 |
| Streptococcus dysgalactiae subsp. equisimilis AKSDE4288 | lmrP | Multidrug resistance protein ErmB | | | efflux pump conferring antibiotic resistance | 11577166 |
| Streptococcus dysgalactiae subsp. equisimilis ATCC 12394 | lmrP | Multidrug resistance protein ErmB | | | efflux pump conferring antibiotic resistance | 11577166 |
| Streptococcus dysgalactiae subsp. equisimilis GGS_124 | lmrP | Multidrug resistance protein ErmB | | | efflux pump conferring antibiotic resistance | 11577166 |
| Streptococcus dysgalactiae subsp. equisimilis RE378 | lmrP | Multidrug resistance protein ErmB | | | efflux pump conferring antibiotic resistance | 11577166 |
| Streptococcus dysgalactiae subsp. equisimilis SK1249 | lmrP | Multidrug resistance protein ErmB | | | efflux pump conferring antibiotic resistance | 11577166 |
| Streptococcus dysgalactiae subsp. equisimilis SK1250 | lmrP | Multidrug resistance protein ErmB | | | efflux pump conferring antibiotic resistance | 11577166 |
| Streptococcus dysgalactiae subsp. equisimilis strain ASDSE_96 | lmrP | Multidrug resistance protein ErmB | | | efflux pump conferring antibiotic resistance | 11577166 |
| Streptococcus dysgalactiae subsp. equisimilis strain ASDSE_99 | lmrP | Multidrug resistance protein ErmB | | | efflux pump conferring antibiotic resistance | 11577166 |
| Streptococcus dysgalactiae subsp. equisimilis strain KNZ01 | lmrP | Multidrug resistance protein ErmB | | | efflux pump conferring antibiotic resistance | 11577166 |
| Streptococcus dysgalactiae subsp. equisimilis strain KNZ03 | lmrP | Multidrug resistance protein ErmB | | | efflux pump conferring antibiotic resistance | 11577166 |
| Streptococcus dysgalactiae subsp. equisimilis strain KNZ04 | lmrP | Multidrug resistance protein ErmB | | | efflux pump conferring antibiotic resistance | 11577166 |
| Streptococcus dysgalactiae subsp. equisimilis strain KNZ06 | lmrP | Multidrug resistance protein ErmB | | | efflux pump conferring antibiotic resistance | 11577166 |
| Streptococcus dysgalactiae subsp. equisimilis strain KNZ07 | lmrP | Multidrug resistance protein ErmB | | | efflux pump conferring antibiotic resistance | 11577166 |
| Streptococcus dysgalactiae subsp. equisimilis strain KNZ10 | lmrP | Multidrug resistance protein ErmB | | | efflux pump conferring antibiotic resistance | 11577166 |
| Streptococcus dysgalactiae subsp. equisimilis strain KNZ12 | lmrP | Multidrug resistance protein ErmB | | | efflux pump conferring antibiotic resistance | 11577166 |
| Streptococcus dysgalactiae subsp. equisimilis strain KNZ15 | lmrP | Multidrug resistance protein ErmB | | | efflux pump conferring antibiotic resistance | 11577166 |
| Streptococcus dysgalactiae subsp. equisimilis strain KNZ16 | lmrP | Multidrug resistance protein ErmB | | | efflux pump conferring antibiotic resistance | 11577166 |
| Streptococcus dysgalactiae subsp. equisimilis strain NCTC10321 | lmrP | Multidrug resistance protein ErmB | | | efflux pump conferring antibiotic resistance | 11577166 |
| Streptococcus dysgalactiae subsp. equisimilis strain NCTC11554 | lmrP | Multidrug resistance protein ErmB | | | efflux pump conferring antibiotic resistance | 11577166 |
| Streptococcus dysgalactiae subsp. equisimilis strain NCTC11555 | lmrP | Multidrug resistance protein ErmB | | | efflux pump conferring antibiotic resistance | 11577166 |
| Streptococcus dysgalactiae subsp. equisimilis strain NCTC11556 | lmrP | Multidrug resistance protein ErmB | | | efflux pump conferring antibiotic resistance | 11577166 |
| Streptococcus dysgalactiae subsp. equisimilis strain NCTC11557 | lmrP | Multidrug resistance protein ErmB | | | efflux pump conferring antibiotic resistance | 11577166 |
| Streptococcus dysgalactiae subsp. equisimilis strain NCTC11564 | lmrP | Multidrug resistance protein ErmB | | | efflux pump conferring antibiotic resistance | 11577166 |
| Streptococcus dysgalactiae subsp. equisimilis strain NCTC5370 | lmrP | Multidrug resistance protein ErmB | | | efflux pump conferring antibiotic resistance | 11577166 |
| Streptococcus dysgalactiae subsp. equisimilis strain NCTC5371 | lmrP | Multidrug resistance protein ErmB | | | efflux pump conferring antibiotic resistance | 11577166 |
| Streptococcus dysgalactiae subsp. equisimilis strain NCTC6179 | lmrP | Multidrug resistance protein ErmB | | | efflux pump conferring antibiotic resistance | 11577166 |
| Streptococcus dysgalactiae subsp. equisimilis strain NCTC6181 | lmrP | Multidrug resistance protein ErmB | | | efflux pump conferring antibiotic resistance | 11577166 |
| Streptococcus dysgalactiae subsp. equisimilis strain NCTC6407 | lmrP | Multidrug resistance protein ErmB | | | efflux pump conferring antibiotic resistance | 11577166 |
| Streptococcus dysgalactiae subsp. equisimilis strain NCTC7136 | lmrP | Multidrug resistance protein ErmB | | | efflux pump conferring antibiotic resistance | 11577166 |
| Streptococcus dysgalactiae subsp. equisimilis strain NCTC8543 | lmrP | Multidrug resistance protein ErmB | | | efflux pump conferring antibiotic resistance | 11577166 |
| Streptococcus dysgalactiae subsp. equisimilis strain NCTC8546 | lmrP | Multidrug resistance protein ErmB | | | efflux pump conferring antibiotic resistance | 11577166 |
| Streptococcus dysgalactiae subsp. equisimilis strain NCTC9413 | lmrP | Multidrug resistance protein ErmB | | | efflux pump conferring antibiotic resistance | 11577166 |
| Streptococcus dysgalactiae subsp. equisimilis strain NCTC9414 | lmrP | Multidrug resistance protein ErmB | | | efflux pump conferring antibiotic resistance | 11577166 |
| Streptococcus dysgalactiae subsp. equisimilis strain NCTC9603 | lmrP | Multidrug resistance protein ErmB | | | efflux pump conferring antibiotic resistance | 11577166 |
| Streptococcus dysgalactiae subsp. equisimilis strain SS1575 | lmrP | Multidrug resistance protein ErmB | | | efflux pump conferring antibiotic resistance | 11577166 |
| Streptococcus dysgalactiae subsp. equisimilis strain UT_4031CC | lmrP | Multidrug resistance protein ErmB | | | efflux pump conferring antibiotic resistance | 11577166 |
| Streptococcus dysgalactiae subsp. equisimilis strain UT_4231_KK | lmrP | Multidrug resistance protein ErmB | | | efflux pump conferring antibiotic resistance | 11577166 |
| Streptococcus dysgalactiae subsp. equisimilis strain UT_4234_DH | lmrP | Multidrug resistance protein ErmB | | | efflux pump conferring antibiotic resistance | 11577166 |
| Streptococcus dysgalactiae subsp. equisimilis strain UT_4241_XS | lmrP | Multidrug resistance protein ErmB | | | efflux pump conferring antibiotic resistance | 11577166 |
| Streptococcus dysgalactiae subsp. equisimilis strain UT_4242_AB | lmrP | Multidrug resistance protein ErmB | | | efflux pump conferring antibiotic resistance | 11577166 |
| Streptococcus dysgalactiae subsp. equisimilis strain UT_4255RC | lmrP | Multidrug resistance protein ErmB | | | efflux pump conferring antibiotic resistance | 11577166 |
| Streptococcus dysgalactiae subsp. equisimilis strain UT_4277_BB | lmrP | Multidrug resistance protein ErmB | | | efflux pump conferring antibiotic resistance | 11577166 |
| Streptococcus dysgalactiae subsp. equisimilis strain UT_4966_RC | lmrP | Multidrug resistance protein ErmB | | | efflux pump conferring antibiotic resistance | 11577166 |
| Streptococcus dysgalactiae subsp. equisimilis strain UT-5345 | lmrP | Multidrug resistance protein ErmB | | | efflux pump conferring antibiotic resistance | 11577166 |
| Streptococcus dysgalactiae subsp. equisimilis strain UT-5354 | lmrP | Multidrug resistance protein ErmB | | | efflux pump conferring antibiotic resistance | 11577166 |
| Streptococcus dysgalactiae subsp. equisimilis strain UT-SS1069 | lmrP | Multidrug resistance protein ErmB | | | efflux pump conferring antibiotic resistance | 11577166 |
| Streptococcus dysgalactiae subsp. equisimilis strain UT-SS957 | lmrP | Multidrug resistance protein ErmB | | | efflux pump conferring antibiotic resistance | 11577166 |
| Streptococcus dysgalactiae subsp. equisimilis strain WCHSDSE-1 | lmrP | Multidrug resistance protein ErmB | | | efflux pump conferring antibiotic resistance | 11577166 |
| Streptococcus dysgalactiae subsp. equisimilis SD SCDR1 | rpoB | DNA-directed RNA polymerase beta subunit (EC 2.7.7.6) | | | antibiotic target in susceptible species | 3050121;15047531;16723576 |
| Streptococcus dysgalactiae subsp. equisimilis 167 | rpoB | DNA-directed RNA polymerase beta subunit (EC 2.7.7.6) | | | antibiotic target in susceptible species | 3050121;15047531;16723576 |
| Streptococcus dysgalactiae subsp. equisimilis AC-2713 | rpoB | DNA-directed RNA polymerase beta subunit (EC 2.7.7.6) | | | antibiotic target in susceptible species | 3050121;15047531;16723576 |
| Streptococcus dysgalactiae subsp. equisimilis AKSDE4288 | rpoB | DNA-directed RNA polymerase beta subunit (EC 2.7.7.6) | | | antibiotic target in susceptible species | 3050121;15047531;16723576 |
| Streptococcus dysgalactiae subsp. equisimilis ATCC 12394 | rpoB | DNA-directed RNA polymerase beta subunit (EC 2.7.7.6) | | | antibiotic target in susceptible species | 3050121;15047531;16723576 |
| Streptococcus dysgalactiae subsp. equisimilis GGS_124 | rpoB | DNA-directed RNA polymerase beta subunit (EC 2.7.7.6) | | | antibiotic target in susceptible species | 3050121;15047531;16723576 |
| Streptococcus dysgalactiae subsp. equisimilis RE378 | rpoB | DNA-directed RNA polymerase beta subunit (EC 2.7.7.6) | | | antibiotic target in susceptible species | 3050121;15047531;16723576 |
| Streptococcus dysgalactiae subsp. equisimilis SK1249 | rpoB | DNA-directed RNA polymerase beta subunit (EC 2.7.7.6) | | | antibiotic target in susceptible species | 3050121;15047531;16723576 |
| Streptococcus dysgalactiae subsp. equisimilis SK1250 | rpoB | DNA-directed RNA polymerase beta subunit (EC 2.7.7.6) | | | antibiotic target in susceptible species | 3050121;15047531;16723576 |
| Streptococcus dysgalactiae subsp. equisimilis strain ASDSE_96 | rpoB | DNA-directed RNA polymerase beta subunit (EC 2.7.7.6) | | | antibiotic target in susceptible species | 3050121;15047531;16723576 |
| Streptococcus dysgalactiae subsp. equisimilis strain ASDSE_99 | rpoB | DNA-directed RNA polymerase beta subunit (EC 2.7.7.6) | | | antibiotic target in susceptible species | 3050121;15047531;16723576 |
| Streptococcus dysgalactiae subsp. equisimilis strain C161L1 | rpoB | DNA-directed RNA polymerase beta subunit (EC 2.7.7.6) | | | antibiotic target in susceptible species | 3050121;15047531;16723576 |
| Streptococcus dysgalactiae subsp. equisimilis strain KNZ01 | rpoB | DNA-directed RNA polymerase beta subunit (EC 2.7.7.6) | | | antibiotic target in susceptible species | 3050121;15047531;16723576 |
| Streptococcus dysgalactiae subsp. equisimilis strain KNZ03 | rpoB | DNA-directed RNA polymerase beta subunit (EC 2.7.7.6) | | | antibiotic target in susceptible species | 3050121;15047531;16723576 |
| Streptococcus dysgalactiae subsp. equisimilis strain KNZ04 | rpoB | DNA-directed RNA polymerase beta subunit (EC 2.7.7.6) | | | antibiotic target in susceptible species | 3050121;15047531;16723576 |
| Streptococcus dysgalactiae subsp. equisimilis strain KNZ06 | rpoB | DNA-directed RNA polymerase beta subunit (EC 2.7.7.6) | | | antibiotic target in susceptible species | 3050121;15047531;16723576 |
| Streptococcus dysgalactiae subsp. equisimilis strain KNZ07 | rpoB | DNA-directed RNA polymerase beta subunit (EC 2.7.7.6) | | | antibiotic target in susceptible species | 3050121;15047531;16723576 |
| Streptococcus dysgalactiae subsp. equisimilis strain KNZ10 | rpoB | DNA-directed RNA polymerase beta subunit (EC 2.7.7.6) | | | antibiotic target in susceptible species | 3050121;15047531;16723576 |
| Streptococcus dysgalactiae subsp. equisimilis strain KNZ12 | rpoB | DNA-directed RNA polymerase beta subunit (EC 2.7.7.6) | | | antibiotic target in susceptible species | 3050121;15047531;16723576 |
| Streptococcus dysgalactiae subsp. equisimilis strain KNZ15 | rpoB | DNA-directed RNA polymerase beta subunit (EC 2.7.7.6) | | | antibiotic target in susceptible species | 3050121;15047531;16723576 |
| Streptococcus dysgalactiae subsp. equisimilis strain KNZ16 | rpoB | DNA-directed RNA polymerase beta subunit (EC 2.7.7.6) | | | antibiotic target in susceptible species | 3050121;15047531;16723576 |
| Streptococcus dysgalactiae subsp. equisimilis strain NCTC10321 | rpoB | DNA-directed RNA polymerase beta subunit (EC 2.7.7.6) | | | antibiotic target in susceptible species | 3050121;15047531;16723576 |
| Streptococcus dysgalactiae subsp. equisimilis strain NCTC11554 | rpoB | DNA-directed RNA polymerase beta subunit (EC 2.7.7.6) | | | antibiotic target in susceptible species | 3050121;15047531;16723576 |
| Streptococcus dysgalactiae subsp. equisimilis strain NCTC11555 | rpoB | DNA-directed RNA polymerase beta subunit (EC 2.7.7.6) | | | antibiotic target in susceptible species | 3050121;15047531;16723576 |
| Streptococcus dysgalactiae subsp. equisimilis strain NCTC11556 | rpoB | DNA-directed RNA polymerase beta subunit (EC 2.7.7.6) | | | antibiotic target in susceptible species | 3050121;15047531;16723576 |
| Streptococcus dysgalactiae subsp. equisimilis strain NCTC11557 | rpoB | DNA-directed RNA polymerase beta subunit (EC 2.7.7.6) | | | antibiotic target in susceptible species | 3050121;15047531;16723576 |
| Streptococcus dysgalactiae subsp. equisimilis strain NCTC11564 | rpoB | DNA-directed RNA polymerase beta subunit (EC 2.7.7.6) | | | antibiotic target in susceptible species | 3050121;15047531;16723576 |
| Streptococcus dysgalactiae subsp. equisimilis strain NCTC11565 | rpoB | DNA-directed RNA polymerase beta subunit (EC 2.7.7.6) | | | antibiotic target in susceptible species | 3050121;15047531;16723576 |
| Streptococcus dysgalactiae subsp. equisimilis strain NCTC5370 | rpoB | DNA-directed RNA polymerase beta subunit (EC 2.7.7.6) | | | antibiotic target in susceptible species | 3050121;15047531;16723576 |
| Streptococcus dysgalactiae subsp. equisimilis strain NCTC5371 | rpoB | DNA-directed RNA polymerase beta subunit (EC 2.7.7.6) | | | antibiotic target in susceptible species | 3050121;15047531;16723576 |
| Streptococcus dysgalactiae subsp. equisimilis strain NCTC5969 | rpoB | DNA-directed RNA polymerase beta subunit (EC 2.7.7.6) | | | antibiotic target in susceptible species | 3050121;15047531;16723576 |
| Streptococcus dysgalactiae subsp. equisimilis strain NCTC6179 | rpoB | DNA-directed RNA polymerase beta subunit (EC 2.7.7.6) | | | antibiotic target in susceptible species | 3050121;15047531;16723576 |
| Streptococcus dysgalactiae subsp. equisimilis strain NCTC6181 | rpoB | DNA-directed RNA polymerase beta subunit (EC 2.7.7.6) | | | antibiotic target in susceptible species | 3050121;15047531;16723576 |
| Streptococcus dysgalactiae subsp. equisimilis strain NCTC6407 | rpoB | DNA-directed RNA polymerase beta subunit (EC 2.7.7.6) | | | antibiotic target in susceptible species | 3050121;15047531;16723576 |
| Streptococcus dysgalactiae subsp. equisimilis strain NCTC7136 | rpoB | DNA-directed RNA polymerase beta subunit (EC 2.7.7.6) | | | antibiotic target in susceptible species | 3050121;15047531;16723576 |
| Streptococcus dysgalactiae subsp. equisimilis strain NCTC8543 | rpoB | DNA-directed RNA polymerase beta subunit (EC 2.7.7.6) | | | antibiotic target in susceptible species | 3050121;15047531;16723576 |
| Streptococcus dysgalactiae subsp. equisimilis strain NCTC8546 | rpoB | DNA-directed RNA polymerase beta subunit (EC 2.7.7.6) | | | antibiotic target in susceptible species | 3050121;15047531;16723576 |
| Streptococcus dysgalactiae subsp. equisimilis strain NCTC9413 | rpoB | DNA-directed RNA polymerase beta subunit (EC 2.7.7.6) | | | antibiotic target in susceptible species | 3050121;15047531;16723576 |
| Streptococcus dysgalactiae subsp. equisimilis strain NCTC9414 | rpoB | DNA-directed RNA polymerase beta subunit (EC 2.7.7.6) | | | antibiotic target in susceptible species | 3050121;15047531;16723576 |
| Streptococcus dysgalactiae subsp. equisimilis strain NCTC9603 | rpoB | DNA-directed RNA polymerase beta subunit (EC 2.7.7.6) | | | antibiotic target in susceptible species | 3050121;15047531;16723576 |
| Streptococcus dysgalactiae subsp. equisimilis strain SS1575 | rpoB | DNA-directed RNA polymerase beta subunit (EC 2.7.7.6) | | | antibiotic target in susceptible species | 3050121;15047531;16723576 |
| Streptococcus dysgalactiae subsp. equisimilis strain T642 | rpoB | DNA-directed RNA polymerase beta subunit (EC 2.7.7.6) | | | antibiotic target in susceptible species | 3050121;15047531;16723576 |
| Streptococcus dysgalactiae subsp. equisimilis strain UT_4031CC | rpoB | DNA-directed RNA polymerase beta subunit (EC 2.7.7.6) | | | antibiotic target in susceptible species | 3050121;15047531;16723576 |
| Streptococcus dysgalactiae subsp. equisimilis strain UT_4231_KK | rpoB | DNA-directed RNA polymerase beta subunit (EC 2.7.7.6) | | | antibiotic target in susceptible species | 3050121;15047531;16723576 |
| Streptococcus dysgalactiae subsp. equisimilis strain UT_4234_DH | rpoB | DNA-directed RNA polymerase beta subunit (EC 2.7.7.6) | | | antibiotic target in susceptible species | 3050121;15047531;16723576 |
| Streptococcus dysgalactiae subsp. equisimilis strain UT_4241_XS | rpoB | DNA-directed RNA polymerase beta subunit (EC 2.7.7.6) | | | antibiotic target in susceptible species | 3050121;15047531;16723576 |
| Streptococcus dysgalactiae subsp. equisimilis strain UT_4242_AB | rpoB | DNA-directed RNA polymerase beta subunit (EC 2.7.7.6) | | | antibiotic target in susceptible species | 3050121;15047531;16723576 |
| Streptococcus dysgalactiae subsp. equisimilis strain UT_4255RC | rpoB | DNA-directed RNA polymerase beta subunit (EC 2.7.7.6) | | | antibiotic target in susceptible species | 3050121;15047531;16723576 |
| Streptococcus dysgalactiae subsp. equisimilis strain UT_4277_BB | rpoB | DNA-directed RNA polymerase beta subunit (EC 2.7.7.6) | | | antibiotic target in susceptible species | 3050121;15047531;16723576 |
| Streptococcus dysgalactiae subsp. equisimilis strain UT_4966_RC | rpoB | DNA-directed RNA polymerase beta subunit (EC 2.7.7.6) | | | antibiotic target in susceptible species | 3050121;15047531;16723576 |
| Streptococcus dysgalactiae subsp. equisimilis strain UT-5345 | rpoB | DNA-directed RNA polymerase beta subunit (EC 2.7.7.6) | | | antibiotic target in susceptible species | 3050121;15047531;16723576 |
| Streptococcus dysgalactiae subsp. equisimilis strain UT-5354 | rpoB | DNA-directed RNA polymerase beta subunit (EC 2.7.7.6) | | | antibiotic target in susceptible species | 3050121;15047531;16723576 |
| Streptococcus dysgalactiae subsp. equisimilis strain UT-SS1069 | rpoB | DNA-directed RNA polymerase beta subunit (EC 2.7.7.6) | | | antibiotic target in susceptible species | 3050121;15047531;16723576 |
| Streptococcus dysgalactiae subsp. equisimilis strain UT-SS957 | rpoB | DNA-directed RNA polymerase beta subunit (EC 2.7.7.6) | | | antibiotic target in susceptible species | 3050121;15047531;16723576 |
| Streptococcus dysgalactiae subsp. equisimilis strain WCHSDSE-1 | rpoB | DNA-directed RNA polymerase beta subunit (EC 2.7.7.6) | | | antibiotic target in susceptible species | 3050121;15047531;16723576 |
| Streptococcus dysgalactiae subsp. equisimilis SD SCDR1 | rpoC | DNA-directed RNA polymerase beta' subunit (EC 2.7.7.6) | | | antibiotic target in susceptible species | 16723576 |
| Streptococcus dysgalactiae subsp. equisimilis 167 | rpoC | DNA-directed RNA polymerase beta' subunit (EC 2.7.7.6) | | | antibiotic target in susceptible species | 16723576 |
| Streptococcus dysgalactiae subsp. equisimilis AC-2713 | rpoC | DNA-directed RNA polymerase beta' subunit (EC 2.7.7.6) | | | antibiotic target in susceptible species | 16723576 |
| Streptococcus dysgalactiae subsp. equisimilis AKSDE4288 | rpoC | DNA-directed RNA polymerase beta' subunit (EC 2.7.7.6) | | | antibiotic target in susceptible species | 16723576 |
| Streptococcus dysgalactiae subsp. equisimilis ATCC 12394 | rpoC | DNA-directed RNA polymerase beta' subunit (EC 2.7.7.6) | | | antibiotic target in susceptible species | 16723576 |
| Streptococcus dysgalactiae subsp. equisimilis GGS_124 | rpoC | DNA-directed RNA polymerase beta' subunit (EC 2.7.7.6) | | | antibiotic target in susceptible species | 16723576 |
| Streptococcus dysgalactiae subsp. equisimilis RE378 | rpoC | DNA-directed RNA polymerase beta' subunit (EC 2.7.7.6) | | | antibiotic target in susceptible species | 16723576 |
| Streptococcus dysgalactiae subsp. equisimilis SK1249 | rpoC | DNA-directed RNA polymerase beta' subunit (EC 2.7.7.6) | | | antibiotic target in susceptible species | 16723576 |
| Streptococcus dysgalactiae subsp. equisimilis SK1250 | rpoC | DNA-directed RNA polymerase beta' subunit (EC 2.7.7.6) | | | antibiotic target in susceptible species | 16723576 |
| Streptococcus dysgalactiae subsp. equisimilis strain ASDSE_96 | rpoC | DNA-directed RNA polymerase beta' subunit (EC 2.7.7.6) | | | antibiotic target in susceptible species | 16723576 |
| Streptococcus dysgalactiae subsp. equisimilis strain ASDSE_99 | rpoC | DNA-directed RNA polymerase beta' subunit (EC 2.7.7.6) | | | antibiotic target in susceptible species | 16723576 |
| Streptococcus dysgalactiae subsp. equisimilis strain C161L1 | rpoC | DNA-directed RNA polymerase beta' subunit (EC 2.7.7.6) | | | antibiotic target in susceptible species | 16723576 |
| Streptococcus dysgalactiae subsp. equisimilis strain KNZ01 | rpoC | DNA-directed RNA polymerase beta' subunit (EC 2.7.7.6) | | | antibiotic target in susceptible species | 16723576 |
| Streptococcus dysgalactiae subsp. equisimilis strain KNZ03 | rpoC | DNA-directed RNA polymerase beta' subunit (EC 2.7.7.6) | | | antibiotic target in susceptible species | 16723576 |
| Streptococcus dysgalactiae subsp. equisimilis strain KNZ04 | rpoC | DNA-directed RNA polymerase beta' subunit (EC 2.7.7.6) | | | antibiotic target in susceptible species | 16723576 |
| Streptococcus dysgalactiae subsp. equisimilis strain KNZ06 | rpoC | DNA-directed RNA polymerase beta' subunit (EC 2.7.7.6) | | | antibiotic target in susceptible species | 16723576 |
| Streptococcus dysgalactiae subsp. equisimilis strain KNZ07 | rpoC | DNA-directed RNA polymerase beta' subunit (EC 2.7.7.6) | | | antibiotic target in susceptible species | 16723576 |
| Streptococcus dysgalactiae subsp. equisimilis strain KNZ10 | rpoC | DNA-directed RNA polymerase beta' subunit (EC 2.7.7.6) | | | antibiotic target in susceptible species | 16723576 |
| Streptococcus dysgalactiae subsp. equisimilis strain KNZ12 | rpoC | DNA-directed RNA polymerase beta' subunit (EC 2.7.7.6) | | | antibiotic target in susceptible species | 16723576 |
| Streptococcus dysgalactiae subsp. equisimilis strain KNZ15 | rpoC | DNA-directed RNA polymerase beta' subunit (EC 2.7.7.6) | | | antibiotic target in susceptible species | 16723576 |
| Streptococcus dysgalactiae subsp. equisimilis strain KNZ16 | rpoC | DNA-directed RNA polymerase beta' subunit (EC 2.7.7.6) | | | antibiotic target in susceptible species | 16723576 |
| Streptococcus dysgalactiae subsp. equisimilis strain NCTC10321 | rpoC | DNA-directed RNA polymerase beta' subunit (EC 2.7.7.6) | | | antibiotic target in susceptible species | 16723576 |
| Streptococcus dysgalactiae subsp. equisimilis strain NCTC10321 | rpoC | DNA-directed RNA polymerase beta' subunit (EC 2.7.7.6) | | | antibiotic target in susceptible species | 16723576 |
| Streptococcus dysgalactiae subsp. equisimilis strain NCTC10321 | rpoC | DNA-directed RNA polymerase beta' subunit (EC 2.7.7.6) | | | antibiotic target in susceptible species | 16723576 |
| Streptococcus dysgalactiae subsp. equisimilis strain NCTC11554 | rpoC | DNA-directed RNA polymerase beta' subunit (EC 2.7.7.6) | | | antibiotic target in susceptible species | 16723576 |
| Streptococcus dysgalactiae subsp. equisimilis strain NCTC11555 | rpoC | DNA-directed RNA polymerase beta' subunit (EC 2.7.7.6) | | | antibiotic target in susceptible species | 16723576 |
| Streptococcus dysgalactiae subsp. equisimilis strain NCTC11556 | rpoC | DNA-directed RNA polymerase beta' subunit (EC 2.7.7.6) | | | antibiotic target in susceptible species | 16723576 |
| Streptococcus dysgalactiae subsp. equisimilis strain NCTC11557 | rpoC | DNA-directed RNA polymerase beta' subunit (EC 2.7.7.6) | | | antibiotic target in susceptible species | 16723576 |
| Streptococcus dysgalactiae subsp. equisimilis strain NCTC11564 | rpoC | DNA-directed RNA polymerase beta' subunit (EC 2.7.7.6) | | | antibiotic target in susceptible species | 16723576 |
| Streptococcus dysgalactiae subsp. equisimilis strain NCTC11565 | rpoC | DNA-directed RNA polymerase beta' subunit (EC 2.7.7.6) | | | antibiotic target in susceptible species | 16723576 |
| Streptococcus dysgalactiae subsp. equisimilis strain NCTC5370 | rpoC | DNA-directed RNA polymerase beta' subunit (EC 2.7.7.6) | | | antibiotic target in susceptible species | 16723576 |
| Streptococcus dysgalactiae subsp. equisimilis strain NCTC5371 | rpoC | DNA-directed RNA polymerase beta' subunit (EC 2.7.7.6) | | | antibiotic target in susceptible species | 16723576 |
| Streptococcus dysgalactiae subsp. equisimilis strain NCTC5969 | rpoC | DNA-directed RNA polymerase beta' subunit (EC 2.7.7.6) | | | antibiotic target in susceptible species | 16723576 |
| Streptococcus dysgalactiae subsp. equisimilis strain NCTC6179 | rpoC | DNA-directed RNA polymerase beta' subunit (EC 2.7.7.6) | | | antibiotic target in susceptible species | 16723576 |
| Streptococcus dysgalactiae subsp. equisimilis strain NCTC6181 | rpoC | DNA-directed RNA polymerase beta' subunit (EC 2.7.7.6) | | | antibiotic target in susceptible species | 16723576 |
| Streptococcus dysgalactiae subsp. equisimilis strain NCTC6407 | rpoC | DNA-directed RNA polymerase beta' subunit (EC 2.7.7.6) | | | antibiotic target in susceptible species | 16723576 |
| Streptococcus dysgalactiae subsp. equisimilis strain NCTC7136 | rpoC | DNA-directed RNA polymerase beta' subunit (EC 2.7.7.6) | | | antibiotic target in susceptible species | 16723576 |
| Streptococcus dysgalactiae subsp. equisimilis strain NCTC8543 | rpoC | DNA-directed RNA polymerase beta' subunit (EC 2.7.7.6) | | | antibiotic target in susceptible species | 16723576 |
| Streptococcus dysgalactiae subsp. equisimilis strain NCTC8546 | rpoC | DNA-directed RNA polymerase beta' subunit (EC 2.7.7.6) | | | antibiotic target in susceptible species | 16723576 |
| Streptococcus dysgalactiae subsp. equisimilis strain NCTC9413 | rpoC | DNA-directed RNA polymerase beta' subunit (EC 2.7.7.6) | | | antibiotic target in susceptible species | 16723576 |
| Streptococcus dysgalactiae subsp. equisimilis strain NCTC9414 | rpoC | DNA-directed RNA polymerase beta' subunit (EC 2.7.7.6) | | | antibiotic target in susceptible species | 16723576 |
| Streptococcus dysgalactiae subsp. equisimilis strain NCTC9603 | rpoC | DNA-directed RNA polymerase beta' subunit (EC 2.7.7.6) | | | antibiotic target in susceptible species | 16723576 |
| Streptococcus dysgalactiae subsp. equisimilis strain SS1575 | rpoC | DNA-directed RNA polymerase beta' subunit (EC 2.7.7.6) | | | antibiotic target in susceptible species | 16723576 |
| Streptococcus dysgalactiae subsp. equisimilis strain T642 | rpoC | DNA-directed RNA polymerase beta' subunit (EC 2.7.7.6) | | | antibiotic target in susceptible species | 16723576 |
| Streptococcus dysgalactiae subsp. equisimilis strain UT_4031CC | rpoC | DNA-directed RNA polymerase beta' subunit (EC 2.7.7.6) | | | antibiotic target in susceptible species | 16723576 |
| Streptococcus dysgalactiae subsp. equisimilis strain UT_4231_KK | rpoC | DNA-directed RNA polymerase beta' subunit (EC 2.7.7.6) | | | antibiotic target in susceptible species | 16723576 |
| Streptococcus dysgalactiae subsp. equisimilis strain UT_4234_DH | rpoC | DNA-directed RNA polymerase beta' subunit (EC 2.7.7.6) | | | antibiotic target in susceptible species | 16723576 |
| Streptococcus dysgalactiae subsp. equisimilis strain UT_4241_XS | rpoC | DNA-directed RNA polymerase beta' subunit (EC 2.7.7.6) | | | antibiotic target in susceptible species | 16723576 |
| Streptococcus dysgalactiae subsp. equisimilis strain UT_4242_AB | rpoC | DNA-directed RNA polymerase beta' subunit (EC 2.7.7.6) | | | antibiotic target in susceptible species | 16723576 |
| Streptococcus dysgalactiae subsp. equisimilis strain UT_4255RC | rpoC | DNA-directed RNA polymerase beta' subunit (EC 2.7.7.6) | | | antibiotic target in susceptible species | 16723576 |
| Streptococcus dysgalactiae subsp. equisimilis strain UT_4277_BB | rpoC | DNA-directed RNA polymerase beta' subunit (EC 2.7.7.6) | | | antibiotic target in susceptible species | 16723576 |
| Streptococcus dysgalactiae subsp. equisimilis strain UT_4966_RC | rpoC | DNA-directed RNA polymerase beta' subunit (EC 2.7.7.6) | | | antibiotic target in susceptible species | 16723576 |
| Streptococcus dysgalactiae subsp. equisimilis strain UT-5345 | rpoC | DNA-directed RNA polymerase beta' subunit (EC 2.7.7.6) | | | antibiotic target in susceptible species | 16723576 |
| Streptococcus dysgalactiae subsp. equisimilis strain UT-5354 | rpoC | DNA-directed RNA polymerase beta' subunit (EC 2.7.7.6) | | | antibiotic target in susceptible species | 16723576 |
| Streptococcus dysgalactiae subsp. equisimilis strain UT-SS1069 | rpoC | DNA-directed RNA polymerase beta' subunit (EC 2.7.7.6) | | | antibiotic target in susceptible species | 16723576 |
| Streptococcus dysgalactiae subsp. equisimilis strain UT-SS957 | rpoC | DNA-directed RNA polymerase beta' subunit (EC 2.7.7.6) | | | antibiotic target in susceptible species | 16723576 |
| Streptococcus dysgalactiae subsp. equisimilis strain WCHSDSE-1 | rpoC | DNA-directed RNA polymerase beta' subunit (EC 2.7.7.6) | | | antibiotic target in susceptible species | 16723576 |
| Streptococcus dysgalactiae subsp. equisimilis SD SCDR1 | S10p | SSU ribosomal protein S10p (S20e) | | | antibiotic target in susceptible species | 26124155 |
| Streptococcus dysgalactiae subsp. equisimilis 167 | rpsJ | SSU ribosomal protein S10p (S20e) | | | antibiotic target in susceptible species | 26124155 |
| Streptococcus dysgalactiae subsp. equisimilis AC-2713 | rpsJ | SSU ribosomal protein S10p (S20e) | | | antibiotic target in susceptible species | 26124155 |
| Streptococcus dysgalactiae subsp. equisimilis AKSDE4288 | rpsJ | SSU ribosomal protein S10p (S20e) | | | antibiotic target in susceptible species | 26124155 |
| Streptococcus dysgalactiae subsp. equisimilis ATCC 12394 | rpsJ | SSU ribosomal protein S10p (S20e) | | | antibiotic target in susceptible species | 26124155 |
| Streptococcus dysgalactiae subsp. equisimilis GGS_124 | rpsJ | SSU ribosomal protein S10p (S20e) | | | antibiotic target in susceptible species | 26124155 |
| Streptococcus dysgalactiae subsp. equisimilis RE378 | rpsJ | SSU ribosomal protein S10p (S20e) | | | antibiotic target in susceptible species | 26124155 |
| Streptococcus dysgalactiae subsp. equisimilis SK1249 | rpsJ | SSU ribosomal protein S10p (S20e) | | | antibiotic target in susceptible species | 26124155 |
| Streptococcus dysgalactiae subsp. equisimilis SK1250 | rpsJ | SSU ribosomal protein S10p (S20e) | | | antibiotic target in susceptible species | 26124155 |
| Streptococcus dysgalactiae subsp. equisimilis strain ASDSE_96 | rpsJ | SSU ribosomal protein S10p (S20e) | | | antibiotic target in susceptible species | 26124155 |
| Streptococcus dysgalactiae subsp. equisimilis strain ASDSE_99 | rpsJ | SSU ribosomal protein S10p (S20e) | | | antibiotic target in susceptible species | 26124155 |
| Streptococcus dysgalactiae subsp. equisimilis strain C161L1 | rpsJ | SSU ribosomal protein S10p (S20e) | | | antibiotic target in susceptible species | 26124155 |
| Streptococcus dysgalactiae subsp. equisimilis strain KNZ01 | S10p | SSU ribosomal protein S10p (S20e) | | | antibiotic target in susceptible species | 26124155 |
| Streptococcus dysgalactiae subsp. equisimilis strain KNZ03 | S10p | SSU ribosomal protein S10p (S20e) | | | antibiotic target in susceptible species | 26124155 |
| Streptococcus dysgalactiae subsp. equisimilis strain KNZ04 | S10p | SSU ribosomal protein S10p (S20e) | | | antibiotic target in susceptible species | 26124155 |
| Streptococcus dysgalactiae subsp. equisimilis strain KNZ06 | S10p | SSU ribosomal protein S10p (S20e) | | | antibiotic target in susceptible species | 26124155 |
| Streptococcus dysgalactiae subsp. equisimilis strain KNZ07 | S10p | SSU ribosomal protein S10p (S20e) | | | antibiotic target in susceptible species | 26124155 |
| Streptococcus dysgalactiae subsp. equisimilis strain KNZ10 | S10p | SSU ribosomal protein S10p (S20e) | | | antibiotic target in susceptible species | 26124155 |
| Streptococcus dysgalactiae subsp. equisimilis strain KNZ12 | S10p | SSU ribosomal protein S10p (S20e) | | | antibiotic target in susceptible species | 26124155 |
| Streptococcus dysgalactiae subsp. equisimilis strain KNZ15 | S10p | SSU ribosomal protein S10p (S20e) | | | antibiotic target in susceptible species | 26124155 |
| Streptococcus dysgalactiae subsp. equisimilis strain KNZ16 | S10p | SSU ribosomal protein S10p (S20e) | | | antibiotic target in susceptible species | 26124155 |
| Streptococcus dysgalactiae subsp. equisimilis strain NCTC10321 | S10p | SSU ribosomal protein S10p (S20e) | | | antibiotic target in susceptible species | 26124155 |
| Streptococcus dysgalactiae subsp. equisimilis strain NCTC11554 | S10p | SSU ribosomal protein S10p (S20e) | | | antibiotic target in susceptible species | 26124155 |
| Streptococcus dysgalactiae subsp. equisimilis strain NCTC11555 | S10p | SSU ribosomal protein S10p (S20e) | | | antibiotic target in susceptible species | 26124155 |
| Streptococcus dysgalactiae subsp. equisimilis strain NCTC11556 | S10p | SSU ribosomal protein S10p (S20e) | | | antibiotic target in susceptible species | 26124155 |
| Streptococcus dysgalactiae subsp. equisimilis strain NCTC11557 | S10p | SSU ribosomal protein S10p (S20e) | | | antibiotic target in susceptible species | 26124155 |
| Streptococcus dysgalactiae subsp. equisimilis strain NCTC11564 | S10p | SSU ribosomal protein S10p (S20e) | | | antibiotic target in susceptible species | 26124155 |
| Streptococcus dysgalactiae subsp. equisimilis strain NCTC11565 | S10p | SSU ribosomal protein S10p (S20e) | | | antibiotic target in susceptible species | 26124155 |
| Streptococcus dysgalactiae subsp. equisimilis strain NCTC5370 | rpsJ | SSU ribosomal protein S10p (S20e) | | | antibiotic target in susceptible species | 26124155 |
| Streptococcus dysgalactiae subsp. equisimilis strain NCTC5371 | rpsJ | SSU ribosomal protein S10p (S20e) | | | antibiotic target in susceptible species | 26124155 |
| Streptococcus dysgalactiae subsp. equisimilis strain NCTC5969 | S10p | SSU ribosomal protein S10p (S20e) | | | antibiotic target in susceptible species | 26124155 |
| Streptococcus dysgalactiae subsp. equisimilis strain NCTC6179 | rpsJ | SSU ribosomal protein S10p (S20e) | | | antibiotic target in susceptible species | 26124155 |
| Streptococcus dysgalactiae subsp. equisimilis strain NCTC6181 | S10p | SSU ribosomal protein S10p (S20e) | | | antibiotic target in susceptible species | 26124155 |
| Streptococcus dysgalactiae subsp. equisimilis strain NCTC6407 | S10p | SSU ribosomal protein S10p (S20e) | | | antibiotic target in susceptible species | 26124155 |
| Streptococcus dysgalactiae subsp. equisimilis strain NCTC7136 | rpsJ | SSU ribosomal protein S10p (S20e) | | | antibiotic target in susceptible species | 26124155 |
| Streptococcus dysgalactiae subsp. equisimilis strain NCTC8543 | S10p | SSU ribosomal protein S10p (S20e) | | | antibiotic target in susceptible species | 26124155 |
| Streptococcus dysgalactiae subsp. equisimilis strain NCTC8546 | S10p | SSU ribosomal protein S10p (S20e) | | | antibiotic target in susceptible species | 26124155 |
| Streptococcus dysgalactiae subsp. equisimilis strain NCTC9413 | S10p | SSU ribosomal protein S10p (S20e) | | | antibiotic target in susceptible species | 26124155 |
| Streptococcus dysgalactiae subsp. equisimilis strain NCTC9414 | rpsJ | SSU ribosomal protein S10p (S20e) | | | antibiotic target in susceptible species | 26124155 |
| Streptococcus dysgalactiae subsp. equisimilis strain NCTC9603 | S10p | SSU ribosomal protein S10p (S20e) | | | antibiotic target in susceptible species | 26124155 |
| Streptococcus dysgalactiae subsp. equisimilis strain SS1575 | rpsJ | SSU ribosomal protein S10p (S20e) | | | antibiotic target in susceptible species | 26124155 |
| Streptococcus dysgalactiae subsp. equisimilis strain T642 | rpsJ | SSU ribosomal protein S10p (S20e) | | | antibiotic target in susceptible species | 26124155 |
| Streptococcus dysgalactiae subsp. equisimilis strain UT_4031CC | rpsJ | SSU ribosomal protein S10p (S20e) | | | antibiotic target in susceptible species | 26124155 |
| Streptococcus dysgalactiae subsp. equisimilis strain UT_4231_KK | rpsJ | SSU ribosomal protein S10p (S20e) | | | antibiotic target in susceptible species | 26124155 |
| Streptococcus dysgalactiae subsp. equisimilis strain UT_4234_DH | rpsJ | SSU ribosomal protein S10p (S20e) | | | antibiotic target in susceptible species | 26124155 |
| Streptococcus dysgalactiae subsp. equisimilis strain UT_4241_XS | rpsJ | SSU ribosomal protein S10p (S20e) | | | antibiotic target in susceptible species | 26124155 |
| Streptococcus dysgalactiae subsp. equisimilis strain UT_4242_AB | rpsJ | SSU ribosomal protein S10p (S20e) | | | antibiotic target in susceptible species | 26124155 |
| Streptococcus dysgalactiae subsp. equisimilis strain UT_4255RC | rpsJ | SSU ribosomal protein S10p (S20e) | | | antibiotic target in susceptible species | 26124155 |
| Streptococcus dysgalactiae subsp. equisimilis strain UT_4277_BB | rpsJ | SSU ribosomal protein S10p (S20e) | | | antibiotic target in susceptible species | 26124155 |
| Streptococcus dysgalactiae subsp. equisimilis strain UT_4966_RC | rpsJ | SSU ribosomal protein S10p (S20e) | | | antibiotic target in susceptible species | 26124155 |
| Streptococcus dysgalactiae subsp. equisimilis strain UT-5345 | rpsJ | SSU ribosomal protein S10p (S20e) | | | antibiotic target in susceptible species | 26124155 |
| Streptococcus dysgalactiae subsp. equisimilis strain UT-5354 | rpsJ | SSU ribosomal protein S10p (S20e) | | | antibiotic target in susceptible species | 26124155 |
| Streptococcus dysgalactiae subsp. equisimilis strain UT-SS1069 | rpsJ | SSU ribosomal protein S10p (S20e) | | | antibiotic target in susceptible species | 26124155 |
| Streptococcus dysgalactiae subsp. equisimilis strain UT-SS957 | rpsJ | SSU ribosomal protein S10p (S20e) | | | antibiotic target in susceptible species | 26124155 |
| Streptococcus dysgalactiae subsp. equisimilis strain WCHSDSE-1 | rpsJ | SSU ribosomal protein S10p (S20e) | | | antibiotic target in susceptible species | 26124155 |
| Streptococcus dysgalactiae subsp. equisimilis SD SCDR1 | S12p | SSU ribosomal protein S12p (S23e) | | | antibiotic target in susceptible species | 7934937 |
| Streptococcus dysgalactiae subsp. equisimilis 167 | rpsL | SSU ribosomal protein S12p (S23e) | | | antibiotic target in susceptible species | 7934937 |
| Streptococcus dysgalactiae subsp. equisimilis AC-2713 | rpsL | SSU ribosomal protein S12p (S23e) | | | antibiotic target in susceptible species | 7934937 |
| Streptococcus dysgalactiae subsp. equisimilis AKSDE4288 | S12p | SSU ribosomal protein S12p (S23e) | | | antibiotic target in susceptible species | 7934937 |
| Streptococcus dysgalactiae subsp. equisimilis ATCC 12394 | rpsL | SSU ribosomal protein S12p (S23e) | | | antibiotic target in susceptible species | 7934937 |
| Streptococcus dysgalactiae subsp. equisimilis GGS_124 | rpsL | SSU ribosomal protein S12p (S23e) | | | antibiotic target in susceptible species | 7934937 |
| Streptococcus dysgalactiae subsp. equisimilis RE378 | rpsL | SSU ribosomal protein S12p (S23e) | | | antibiotic target in susceptible species | 7934937 |
| Streptococcus dysgalactiae subsp. equisimilis SK1249 | rpsL | SSU ribosomal protein S12p (S23e) | | | antibiotic target in susceptible species | 7934937 |
| Streptococcus dysgalactiae subsp. equisimilis SK1250 | rpsL | SSU ribosomal protein S12p (S23e) | | | antibiotic target in susceptible species | 7934937 |
| Streptococcus dysgalactiae subsp. equisimilis strain ASDSE_96 | S12p | SSU ribosomal protein S12p (S23e) | | | antibiotic target in susceptible species | 7934937 |
| Streptococcus dysgalactiae subsp. equisimilis strain ASDSE_99 | S12p | SSU ribosomal protein S12p (S23e) | | | antibiotic target in susceptible species | 7934937 |
| Streptococcus dysgalactiae subsp. equisimilis strain C161L1 | S12p | SSU ribosomal protein S12p (S23e) | | | antibiotic target in susceptible species | 7934937 |
| Streptococcus dysgalactiae subsp. equisimilis strain KNZ01 | S12p | SSU ribosomal protein S12p (S23e) | | | antibiotic target in susceptible species | 7934937 |
| Streptococcus dysgalactiae subsp. equisimilis strain KNZ03 | S12p | SSU ribosomal protein S12p (S23e) | | | antibiotic target in susceptible species | 7934937 |
| Streptococcus dysgalactiae subsp. equisimilis strain KNZ04 | S12p | SSU ribosomal protein S12p (S23e) | | | antibiotic target in susceptible species | 7934937 |
| Streptococcus dysgalactiae subsp. equisimilis strain KNZ06 | S12p | SSU ribosomal protein S12p (S23e) | | | antibiotic target in susceptible species | 7934937 |
| Streptococcus dysgalactiae subsp. equisimilis strain KNZ07 | S12p | SSU ribosomal protein S12p (S23e) | | | antibiotic target in susceptible species | 7934937 |
| Streptococcus dysgalactiae subsp. equisimilis strain KNZ10 | S12p | SSU ribosomal protein S12p (S23e) | | | antibiotic target in susceptible species | 7934937 |
| Streptococcus dysgalactiae subsp. equisimilis strain KNZ12 | S12p | SSU ribosomal protein S12p (S23e) | | | antibiotic target in susceptible species | 7934937 |
| Streptococcus dysgalactiae subsp. equisimilis strain KNZ15 | S12p | SSU ribosomal protein S12p (S23e) | | | antibiotic target in susceptible species | 7934937 |
| Streptococcus dysgalactiae subsp. equisimilis strain KNZ16 | S12p | SSU ribosomal protein S12p (S23e) | | | antibiotic target in susceptible species | 7934937 |
| Streptococcus dysgalactiae subsp. equisimilis strain NCTC10321 | S12p | SSU ribosomal protein S12p (S23e) | | | antibiotic target in susceptible species | 7934937 |
| Streptococcus dysgalactiae subsp. equisimilis strain NCTC11554 | S12p | SSU ribosomal protein S12p (S23e) | | | antibiotic target in susceptible species | 7934937 |
| Streptococcus dysgalactiae subsp. equisimilis strain NCTC11555 | S12p | SSU ribosomal protein S12p (S23e) | | | antibiotic target in susceptible species | 7934937 |
| Streptococcus dysgalactiae subsp. equisimilis strain NCTC11556 | S12p | SSU ribosomal protein S12p (S23e) | | | antibiotic target in susceptible species | 7934937 |
| Streptococcus dysgalactiae subsp. equisimilis strain NCTC11557 | S12p | SSU ribosomal protein S12p (S23e) | | | antibiotic target in susceptible species | 7934937 |
| Streptococcus dysgalactiae subsp. equisimilis strain NCTC11564 | S12p | SSU ribosomal protein S12p (S23e) | | | antibiotic target in susceptible species | 7934937 |
| Streptococcus dysgalactiae subsp. equisimilis strain NCTC11565 | S12p | SSU ribosomal protein S12p (S23e) | | | antibiotic target in susceptible species | 7934937 |
| Streptococcus dysgalactiae subsp. equisimilis strain NCTC5370 | rpsL | SSU ribosomal protein S12p (S23e) | | | antibiotic target in susceptible species | 7934937 |
| Streptococcus dysgalactiae subsp. equisimilis strain NCTC5371 | rpsL | SSU ribosomal protein S12p (S23e) | | | antibiotic target in susceptible species | 7934937 |
| Streptococcus dysgalactiae subsp. equisimilis strain NCTC5969 | S12p | SSU ribosomal protein S12p (S23e) | | | antibiotic target in susceptible species | 7934937 |
| Streptococcus dysgalactiae subsp. equisimilis strain NCTC6179 | rpsL | SSU ribosomal protein S12p (S23e) | | | antibiotic target in susceptible species | 7934937 |
| Streptococcus dysgalactiae subsp. equisimilis strain NCTC6181 | S12p | SSU ribosomal protein S12p (S23e) | | | antibiotic target in susceptible species | 7934937 |
| Streptococcus dysgalactiae subsp. equisimilis strain NCTC6407 | S12p | SSU ribosomal protein S12p (S23e) | | | antibiotic target in susceptible species | 7934937 |
| Streptococcus dysgalactiae subsp. equisimilis strain NCTC7136 | rpsL | SSU ribosomal protein S12p (S23e) | | | antibiotic target in susceptible species | 7934937 |
| Streptococcus dysgalactiae subsp. equisimilis strain NCTC8543 | S12p | SSU ribosomal protein S12p (S23e) | | | antibiotic target in susceptible species | 7934937 |
| Streptococcus dysgalactiae subsp. equisimilis strain NCTC8546 | S12p | SSU ribosomal protein S12p (S23e) | | | antibiotic target in susceptible species | 7934937 |
| Streptococcus dysgalactiae subsp. equisimilis strain NCTC9413 | S12p | SSU ribosomal protein S12p (S23e) | | | antibiotic target in susceptible species | 7934937 |
| Streptococcus dysgalactiae subsp. equisimilis strain NCTC9414 | rpsL | SSU ribosomal protein S12p (S23e) | | | antibiotic target in susceptible species | 7934937 |
| Streptococcus dysgalactiae subsp. equisimilis strain NCTC9603 | S12p | SSU ribosomal protein S12p (S23e) | | | antibiotic target in susceptible species | 7934937 |
| Streptococcus dysgalactiae subsp. equisimilis strain SS1575 | S12p | SSU ribosomal protein S12p (S23e) | | | antibiotic target in susceptible species | 7934937 |
| Streptococcus dysgalactiae subsp. equisimilis strain T642 | S12p | SSU ribosomal protein S12p (S23e) | | | antibiotic target in susceptible species | 7934937 |
| Streptococcus dysgalactiae subsp. equisimilis strain UT_4031CC | S12p | SSU ribosomal protein S12p (S23e) | | | antibiotic target in susceptible species | 7934937 |
| Streptococcus dysgalactiae subsp. equisimilis strain UT_4231_KK | S12p | SSU ribosomal protein S12p (S23e) | | | antibiotic target in susceptible species | 7934937 |
| Streptococcus dysgalactiae subsp. equisimilis strain UT_4234_DH | S12p | SSU ribosomal protein S12p (S23e) | | | antibiotic target in susceptible species | 7934937 |
| Streptococcus dysgalactiae subsp. equisimilis strain UT_4241_XS | S12p | SSU ribosomal protein S12p (S23e) | | | antibiotic target in susceptible species | 7934937 |
| Streptococcus dysgalactiae subsp. equisimilis strain UT_4242_AB | S12p | SSU ribosomal protein S12p (S23e) | | | antibiotic target in susceptible species | 7934937 |
| Streptococcus dysgalactiae subsp. equisimilis strain UT_4255RC | S12p | SSU ribosomal protein S12p (S23e) | | | antibiotic target in susceptible species | 7934937 |
| Streptococcus dysgalactiae subsp. equisimilis strain UT_4277_BB | S12p | SSU ribosomal protein S12p (S23e) | | | antibiotic target in susceptible species | 7934937 |
| Streptococcus dysgalactiae subsp. equisimilis strain UT_4966_RC | S12p | SSU ribosomal protein S12p (S23e) | | | antibiotic target in susceptible species | 7934937 |
| Streptococcus dysgalactiae subsp. equisimilis strain UT-5345 | S12p | SSU ribosomal protein S12p (S23e) | | | antibiotic target in susceptible species | 7934937 |
| Streptococcus dysgalactiae subsp. equisimilis strain UT-5354 | S12p | SSU ribosomal protein S12p (S23e) | | | antibiotic target in susceptible species | 7934937 |
| Streptococcus dysgalactiae subsp. equisimilis strain UT-SS1069 | S12p | SSU ribosomal protein S12p (S23e) | | | antibiotic target in susceptible species | 7934937 |
| Streptococcus dysgalactiae subsp. equisimilis strain UT-SS957 | S12p | SSU ribosomal protein S12p (S23e) | | | antibiotic target in susceptible species | 7934937 |
| Streptococcus dysgalactiae subsp. equisimilis strain WCHSDSE-1 | S12p | SSU ribosomal protein S12p (S23e) | | | antibiotic target in susceptible species | 7934937 |
| Streptococcus dysgalactiae subsp. equisimilis AC-2713 |  | ABC-F type ribosomal protection protein => Msr(D) | | | efflux pump conferring antibiotic resistance |  |
| Streptococcus dysgalactiae subsp. equisimilis AC-2713 |  | ABC-F type ribosomal protection protein => Msr(D) | | |  |  |
| Streptococcus dysgalactiae subsp. equisimilis AC-2713 |  | ABC-F type ribosomal protection protein => Msr(D) | | | antibiotic target protection protein | 16223938 |
| Streptococcus dysgalactiae subsp. equisimilis AC-2713 |  | Macrolide resistance, MFS efflux pump => Mef(A) | | |  |  |
| Streptococcus dysgalactiae subsp. equisimilis AC-2713 |  | Macrolide resistance, MFS efflux pump => Mef(A) | | | efflux pump conferring antibiotic resistance |  |
| Streptococcus dysgalactiae subsp. equisimilis AC-2713 |  | Macrolide resistance, MFS efflux pump => Mef(A) | | | efflux pump conferring antibiotic resistance | 10952626 |
| Streptococcus dysgalactiae subsp. equisimilis AKSDE4288 |  | Tetracycline resistance, ribosomal protection type => Tet(O) | | |  |  |
| Streptococcus dysgalactiae subsp. equisimilis AKSDE4288 |  | Tetracycline resistance, ribosomal protection type => Tet(O) | | | antibiotic target protection protein | 11389850;3245693;9449261 12936983 |
| Streptococcus dysgalactiae subsp. equisimilis AKSDE4288 |  | Tetracycline resistance, ribosomal protection type => Tet(O) | | | antibiotic target protection protein;tetracycline resistance gene |  |
| Streptococcus dysgalactiae subsp. equisimilis SK1250 |  | Tetracycline resistance, ribosomal protection type => Tet(O) | | | antibiotic target protection protein;tetracycline resistance gene |  |
| Streptococcus dysgalactiae subsp. equisimilis SK1250 |  | Tetracycline resistance, ribosomal protection type => Tet(O) | | |  |  |
| Streptococcus dysgalactiae subsp. equisimilis SK1250 |  | Tetracycline resistance, ribosomal protection type => Tet(O) | | | antibiotic target protection protein | 11389850;3245693;9449261 12936983 |
| Streptococcus dysgalactiae subsp. equisimilis strain ASDSE_96 |  | Tetracycline resistance, ribosomal protection type => Tet(M) | | | antibiotic target protection protein | 1993661;8226667 |
| Streptococcus dysgalactiae subsp. equisimilis strain ASDSE_96 |  | Tetracycline resistance, ribosomal protection type => Tet(M) | | |  |  |
| Streptococcus dysgalactiae subsp. equisimilis strain ASDSE_96 |  | Tetracycline resistance, ribosomal protection type => Tet(M) | | | antibiotic target protection protein | 1993661;8226667 |
| Streptococcus dysgalactiae subsp. equisimilis strain ASDSE_96 |  | Tetracycline resistance, ribosomal protection type => Tet(M) | | | antibiotic target protection protein;tetracycline resistance gene |  |
| Streptococcus dysgalactiae subsp. equisimilis strain ASDSE_96 |  | Tetracycline resistance, ribosomal protection type => Tet(M) | | |  |  |
| Streptococcus dysgalactiae subsp. equisimilis strain ASDSE_96 |  | Tetracycline resistance, ribosomal protection type => Tet(M) | | | antibiotic target protection protein;tetracycline resistance gene |  |
| Streptococcus dysgalactiae subsp. equisimilis strain ASDSE_99 |  | Tetracycline resistance, ribosomal protection type => Tet(M) | | | antibiotic target protection protein;tetracycline resistance gene |  |
| Streptococcus dysgalactiae subsp. equisimilis strain ASDSE_99 |  | Tetracycline resistance, ribosomal protection type => Tet(M) | | |  |  |
| Streptococcus dysgalactiae subsp. equisimilis strain ASDSE_99 |  | Tetracycline resistance, ribosomal protection type => Tet(M) | | | antibiotic target protection protein | 1993661;8226667 |
| Streptococcus dysgalactiae subsp. equisimilis strain C161L1 |  | DNA topoisomerase IV subunit B (EC 5.99.1.3) | | | ARO:3000457,ARO:1000001 |  |
| Streptococcus dysgalactiae subsp. equisimilis strain C161L1 |  | Phosphate regulon transcriptional regulatory protein PhoB (SphR) | | | ARO:1000001,ARO:3000834 |  |
| Streptococcus dysgalactiae subsp. equisimilis strain KNZ04 | Tet(M) | Tetracycline resistance, ribosomal protection type => Tet(M) | | | antibiotic target protection protein | 1993661;8226667 |
| Streptococcus dysgalactiae subsp. equisimilis strain KNZ04 | tetM | Tetracycline resistance, ribosomal protection type => Tet(M) | | |  |  |
| Streptococcus dysgalactiae subsp. equisimilis strain KNZ04 | tetM | Tetracycline resistance, ribosomal protection type => Tet(M) | | | antibiotic target protection protein,tetracycline resistance gene |  |
| Streptococcus dysgalactiae subsp. equisimilis strain KNZ06 | parC | DNA topoisomerase IV subunit A (EC 5.99.1.3) | | | antibiotic resistant gene variant or mutant,fluoroquinolone resistance gene,gene involved in self resistance to antibiotic |  |
| Streptococcus dysgalactiae subsp. equisimilis strain KNZ06 | tetM | Tetracycline resistance, ribosomal protection type => Tet(M) | | |  |  |
| Streptococcus dysgalactiae subsp. equisimilis strain KNZ06 | Tet(M) | Tetracycline resistance, ribosomal protection type => Tet(M) | | | antibiotic target protection protein | 1993661;8226667 |
| Streptococcus dysgalactiae subsp. equisimilis strain KNZ06 | tetM | Tetracycline resistance, ribosomal protection type => Tet(M) | | | antibiotic target protection protein,tetracycline resistance gene |  |
| Streptococcus dysgalactiae subsp. equisimilis strain NCTC11557 | tetM | Tetracycline resistance, ribosomal protection type => Tet(M) | | | antibiotic target protection protein,tetracycline resistance gene |  |
| Streptococcus dysgalactiae subsp. equisimilis strain NCTC11557 | tetM | Tetracycline resistance, ribosomal protection type => Tet(M) | | |  |  |
| Streptococcus dysgalactiae subsp. equisimilis strain NCTC11557 | Tet(M) | Tetracycline resistance, ribosomal protection type => Tet(M) | | | antibiotic target protection protein | 1993661;8226667 |
| Streptococcus dysgalactiae subsp. equisimilis strain T642 |  | DNA topoisomerase IV subunit B (EC 5.99.1.3) | | | ARO:3000457,ARO:1000001 |  |
| Streptococcus dysgalactiae subsp. equisimilis strain T642 |  | Phosphate regulon transcriptional regulatory protein PhoB (SphR) | | | ARO:1000001,ARO:3000834 |  |
| Streptococcus dysgalactiae subsp. equisimilis strain UT_4234_DH |  | Tetracycline resistance, ribosomal protection type => Tet(M) | | | antibiotic target protection protein | 1993661;8226667 |
| Streptococcus dysgalactiae subsp. equisimilis strain UT_4234_DH |  | Tetracycline resistance, ribosomal protection type => Tet(M) | | |  |  |
| Streptococcus dysgalactiae subsp. equisimilis strain UT_4234_DH |  | Tetracycline resistance, ribosomal protection type => Tet(M) | | | antibiotic target protection protein;tetracycline resistance gene |  |
| Streptococcus dysgalactiae subsp. equisimilis strain UT_4242_AB |  | Tetracycline resistance, ribosomal protection type => Tet(M) | | |  |  |
| Streptococcus dysgalactiae subsp. equisimilis strain UT_4242_AB |  | Tetracycline resistance, ribosomal protection type => Tet(M) | | | antibiotic target protection protein | 1993661;8226667 |
| Streptococcus dysgalactiae subsp. equisimilis strain UT_4242_AB |  | Tetracycline resistance, ribosomal protection type => Tet(M) | | | antibiotic target protection protein;tetracycline resistance gene |  |
| Streptococcus dysgalactiae subsp. equisimilis strain UT_4242_AB |  | Tetracycline resistance, ribosomal protection type => Tet(M) | | | antibiotic target protection protein | 1993661;8226667 |
| Streptococcus dysgalactiae subsp. equisimilis strain UT_4242_AB |  | Tetracycline resistance, ribosomal protection type => Tet(M) | | |  |  |
| Streptococcus dysgalactiae subsp. equisimilis strain UT_4242_AB |  | Tetracycline resistance, ribosomal protection type => Tet(M) | | | antibiotic target protection protein;tetracycline resistance gene |  |
