## Supplemental File 4 for "Comprehensive comparative genomics analysis for the emerging human pathogen *Streptococcus dysgalactiae* subsp. *equisimilis* (SDSE): A case study and Pan-subspecies genomic analysis"

Supplementary Table 4: Major virulence factors (VF) genes annotated in SDSE genomes and corresponding products.

| Genome Name | Gene | Product | PubMed | Subject Coverage | Query Coverage | Identity | E-value |
| --- | --- | --- | --- | --- | --- | --- | --- |
| Streptococcus dysgalactiae subsp. equisimilis 167 | perR | Peroxide stress regulator PerR, FUR family | 12183543 | 100 | 100 | 94 | 5e-83 |
| Streptococcus dysgalactiae subsp. equisimilis AC-2713 | perR | Peroxide stress regulator PerR, FUR family | 12183543 | 100 | 100 | 94 | 7e-84 |
| Streptococcus dysgalactiae subsp. equisimilis AKSDE4288 | perR | Peroxide stress regulator PerR, FUR family | 12183543 | 100 | 100 | 94 | 7e-84 |
| Streptococcus dysgalactiae subsp. equisimilis ATCC 12394 | perR | Peroxide stress regulator PerR, FUR family | 12183543 | 100 | 100 | 94 | 6e-83 |
| Streptococcus dysgalactiae subsp. equisimilis GGS_124 | perR | Peroxide stress regulator PerR, FUR family | 12183543 | 100 | 100 | 94 | 7e-84 |
| Streptococcus dysgalactiae subsp. equisimilis RE378 | perR | Peroxide stress regulator PerR, FUR family | 12183543 | 100 | 100 | 94 | 7e-84 |
| Streptococcus dysgalactiae subsp. equisimilis SD SCDR1 | perR | Peroxide stress regulator PerR, FUR family | 12183543 | 100 | 100 | 93 | 5e-83 |
| Streptococcus dysgalactiae subsp. equisimilis SK1249 | perR | Peroxide stress regulator PerR, FUR family | 12183543 | 52 | 98 | 91 | 4e-39 |
| Streptococcus dysgalactiae subsp. equisimilis SK1250 | perR | Peroxide stress regulator PerR, FUR family | 12183543 | 100 | 100 | 93 | 5e-83 |
| Streptococcus dysgalactiae subsp. equisimilis strain ASDSE_96 | perR | Peroxide stress regulator PerR, FUR family | 12183543 | 100 | 100 | 94 | 7e-84 |
| Streptococcus dysgalactiae subsp. equisimilis strain ASDSE_99 | perR | Peroxide stress regulator PerR, FUR family | 12183543 | 100 | 100 | 94 | 7e-84 |
| Streptococcus dysgalactiae subsp. equisimilis strain C161L1 | perR | Peroxide stress regulator PerR, FUR family | 12183543 | 100 | 100 | 94 | 2e-110 |
| Streptococcus dysgalactiae subsp. equisimilis strain KNZ01 | perR | Peroxide stress regulator PerR, FUR family | 12183543 | 100 | 100 | 94 | 7e-84 |
| Streptococcus dysgalactiae subsp. equisimilis strain KNZ03 | perR | Peroxide stress regulator PerR, FUR family | 12183543 | 100 | 100 | 94 | 6e-83 |
| Streptococcus dysgalactiae subsp. equisimilis strain KNZ04 | perR | Peroxide stress regulator PerR, FUR family | 12183543 | 100 | 100 | 94 | 6e-83 |
| Streptococcus dysgalactiae subsp. equisimilis strain KNZ06 | perR | Peroxide stress regulator PerR, FUR family | 12183543 | 100 | 100 | 94 | 7e-84 |
| Streptococcus dysgalactiae subsp. equisimilis strain KNZ07 | perR | Peroxide stress regulator PerR, FUR family | 12183543 | 100 | 100 | 94 | 7e-84 |
| Streptococcus dysgalactiae subsp. equisimilis strain KNZ10 | perR | Peroxide stress regulator PerR, FUR family | 12183543 | 100 | 100 | 94 | 7e-84 |
| Streptococcus dysgalactiae subsp. equisimilis strain KNZ12 | perR | Peroxide stress regulator PerR, FUR family | 12183543 | 100 | 100 | 94 | 7e-84 |
| Streptococcus dysgalactiae subsp. equisimilis strain KNZ15 | perR | Peroxide stress regulator PerR, FUR family | 12183543 | 100 | 100 | 94 | 7e-84 |
| Streptococcus dysgalactiae subsp. equisimilis strain KNZ16 | perR | Peroxide stress regulator PerR, FUR family | 12183543 | 100 | 100 | 94 | 6e-83 |
| Streptococcus dysgalactiae subsp. equisimilis strain NCTC10321 | perR | Peroxide stress regulator PerR, FUR family | 12183543 | 100 | 100 | 94 | 2e-83 |
| Streptococcus dysgalactiae subsp. equisimilis strain NCTC11554 | perR | Peroxide stress regulator PerR, FUR family | 12183543 | 100 | 100 | 94 | 6e-83 |
| Streptococcus dysgalactiae subsp. equisimilis strain NCTC11555 | perR | Peroxide stress regulator PerR, FUR family | 12183543 | 100 | 100 | 94 | 2e-83 |
| Streptococcus dysgalactiae subsp. equisimilis strain NCTC11556 | perR | Peroxide stress regulator PerR, FUR family | 12183543 | 100 | 100 | 93 | 5e-83 |
| Streptococcus dysgalactiae subsp. equisimilis strain NCTC11557 | perR | Peroxide stress regulator PerR, FUR family | 12183543 | 100 | 100 | 94 | 7e-84 |
| Streptococcus dysgalactiae subsp. equisimilis strain NCTC11564 | perR | Peroxide stress regulator PerR, FUR family | 12183543 | 100 | 100 | 94 | 7e-84 |
| Streptococcus dysgalactiae subsp. equisimilis strain NCTC5370 | perR | Peroxide stress regulator PerR, FUR family | 12183543 | 100 | 100 | 94 | 5e-83 |
| Streptococcus dysgalactiae subsp. equisimilis strain NCTC5371 | perR | Peroxide stress regulator PerR, FUR family | 12183543 | 100 | 100 | 94 | 7e-84 |
| Streptococcus dysgalactiae subsp. equisimilis strain NCTC6179 | perR | Peroxide stress regulator PerR, FUR family | 12183543 | 100 | 98 | 94 | 7e-84 |
| Streptococcus dysgalactiae subsp. equisimilis strain NCTC6181 | perR | Peroxide stress regulator PerR, FUR family | 12183543 | 100 | 98 | 94 | 7e-84 |
| Streptococcus dysgalactiae subsp. equisimilis strain NCTC6407 | perR | Peroxide stress regulator PerR, FUR family | 12183543 | 100 | 100 | 94 | 2e-83 |
| Streptococcus dysgalactiae subsp. equisimilis strain NCTC7136 | perR | Peroxide stress regulator PerR, FUR family | 12183543 | 100 | 100 | 94 | 7e-84 |
| Streptococcus dysgalactiae subsp. equisimilis strain NCTC8543 | perR | Peroxide stress regulator PerR, FUR family | 12183543 | 100 | 100 | 93 | 1e-82 |
| Streptococcus dysgalactiae subsp. equisimilis strain NCTC8546 | perR | Peroxide stress regulator PerR, FUR family | 12183543 | 100 | 100 | 94 | 7e-84 |
| Streptococcus dysgalactiae subsp. equisimilis strain NCTC9413 | perR | Peroxide stress regulator PerR, FUR family | 12183543 | 100 | 100 | 94 | 7e-84 |
| Streptococcus dysgalactiae subsp. equisimilis strain NCTC9414 | perR | Peroxide stress regulator PerR, FUR family | 12183543 | 100 | 100 | 94 | 7e-84 |
| Streptococcus dysgalactiae subsp. equisimilis strain NCTC9603 | perR | Peroxide stress regulator PerR, FUR family | 12183543 | 100 | 100 | 94 | 7e-84 |
| Streptococcus dysgalactiae subsp. equisimilis strain SS1575 | perR | Peroxide stress regulator PerR, FUR family | 12183543 | 100 | 100 | 94 | 7e-84 |
| Streptococcus dysgalactiae subsp. equisimilis strain T642 | perR | Peroxide stress regulator PerR, FUR family | 12183543 | 100 | 100 | 94 | 0 |
| Streptococcus dysgalactiae subsp. equisimilis strain UT_4031CC | perR | Peroxide stress regulator PerR, FUR family | 12183543 | 100 | 100 | 94 | 7e-84 |
| Streptococcus dysgalactiae subsp. equisimilis strain UT_4231_KK | perR | Peroxide stress regulator PerR, FUR family | 12183543 | 100 | 100 | 94 | 5e-83 |
| Streptococcus dysgalactiae subsp. equisimilis strain UT_4234_DH | perR | Peroxide stress regulator PerR, FUR family | 12183543 | 100 | 100 | 94 | 7e-84 |
| Streptococcus dysgalactiae subsp. equisimilis strain UT_4241_XS | perR | Peroxide stress regulator PerR, FUR family | 12183543 | 100 | 100 | 94 | 5e-83 |
| Streptococcus dysgalactiae subsp. equisimilis strain UT_4242_AB | perR | Peroxide stress regulator PerR, FUR family | 12183543 | 100 | 100 | 93 | 5e-83 |
| Streptococcus dysgalactiae subsp. equisimilis strain UT_4255RC | perR | Peroxide stress regulator PerR, FUR family | 12183543 | 100 | 100 | 94 | 7e-84 |
| Streptococcus dysgalactiae subsp. equisimilis strain UT_4277_BB | perR | Peroxide stress regulator PerR, FUR family | 12183543 | 100 | 100 | 94 | 7e-84 |
| Streptococcus dysgalactiae subsp. equisimilis strain UT_4966_RC | perR | Peroxide stress regulator PerR, FUR family | 12183543 | 100 | 100 | 94 | 5e-83 |
| Streptococcus dysgalactiae subsp. equisimilis strain UT-5345 | perR | Peroxide stress regulator PerR, FUR family | 12183543 | 100 | 100 | 94 | 1e-83 |
| Streptococcus dysgalactiae subsp. equisimilis strain UT-5354 | perR | Peroxide stress regulator PerR, FUR family | 12183543 | 100 | 100 | 94 | 2e-83 |
| Streptococcus dysgalactiae subsp. equisimilis strain UT-SS1069 | perR | Peroxide stress regulator PerR, FUR family | 12183543 | 100 | 100 | 94 | 5e-83 |
| Streptococcus dysgalactiae subsp. equisimilis strain UT-SS957 | perR | Peroxide stress regulator PerR, FUR family | 12183543 | 100 | 100 | 93 | 5e-83 |
| Streptococcus dysgalactiae subsp. equisimilis strain WCHSDSE-1 | perR | Peroxide stress regulator PerR, FUR family | 12183543 | 100 | 100 | 93 | 5e-83 |
| Streptococcus dysgalactiae subsp. equisimilis 167 | purB | Adenylosuccinate lyase (EC 4.3.2.2) @ SAICAR lyase (EC 4.3.2.2) | 11359563 | 98 | 99 | 94 | 1e-236 |
| Streptococcus dysgalactiae subsp. equisimilis AC-2713 | purB | Adenylosuccinate lyase (EC 4.3.2.2) @ SAICAR lyase (EC 4.3.2.2) | 11359563 | 99 | 100 | 94 | 1e-237 |
| Streptococcus dysgalactiae subsp. equisimilis AKSDE4288 | purB | Adenylosuccinate lyase (EC 4.3.2.2) @ SAICAR lyase (EC 4.3.2.2) | 11359563 | 99 | 100 | 94 | 1e-237 |
| Streptococcus dysgalactiae subsp. equisimilis ATCC 12394 | purB | Adenylosuccinate lyase (EC 4.3.2.2) @ SAICAR lyase (EC 4.3.2.2) | 11359563 | 99 | 100 | 94 | 1e-237 |
| Streptococcus dysgalactiae subsp. equisimilis GGS_124 | purB | Adenylosuccinate lyase (EC 4.3.2.2) @ SAICAR lyase (EC 4.3.2.2) | 11359563 | 99 | 100 | 94 | 1e-237 |
| Streptococcus dysgalactiae subsp. equisimilis RE378 | purB | Adenylosuccinate lyase (EC 4.3.2.2) @ SAICAR lyase (EC 4.3.2.2) | 11359563 | 98 | 99 | 93 | 1e-235 |
| Streptococcus dysgalactiae subsp. equisimilis SD SCDR1 | purB | Adenylosuccinate lyase (EC 4.3.2.2) @ SAICAR lyase (EC 4.3.2.2) | 11359563 | 99 | 100 | 94 | 1e-237 |
| Streptococcus dysgalactiae subsp. equisimilis SK1249 | purB | Adenylosuccinate lyase (EC 4.3.2.2) @ SAICAR lyase (EC 4.3.2.2) | 11359563 | 99 | 100 | 94 | 1e-237 |
| Streptococcus dysgalactiae subsp. equisimilis SK1250 | purB | Adenylosuccinate lyase (EC 4.3.2.2) @ SAICAR lyase (EC 4.3.2.2) | 11359563 | 99 | 100 | 94 | 1e-237 |
| Streptococcus dysgalactiae subsp. equisimilis strain ASDSE_96 | purB | Adenylosuccinate lyase (EC 4.3.2.2) @ SAICAR lyase (EC 4.3.2.2) | 11359563 | 99 | 100 | 94 | 1e-237 |
| Streptococcus dysgalactiae subsp. equisimilis strain ASDSE_99 | purB | Adenylosuccinate lyase (EC 4.3.2.2) @ SAICAR lyase (EC 4.3.2.2) | 11359563 | 99 | 100 | 93 | 1e-237 |
| Streptococcus dysgalactiae subsp. equisimilis strain C161L1 | purB | Adenylosuccinate lyase (EC 4.3.2.2) @ SAICAR lyase (EC 4.3.2.2) | 11359563 | 99 | 100 | 93 | 0 |
| Streptococcus dysgalactiae subsp. equisimilis strain KNZ01 | purB | Adenylosuccinate lyase (EC 4.3.2.2) @ SAICAR lyase (EC 4.3.2.2) | 11359563 | 98 | 99 | 94 | 1e-236 |
| Streptococcus dysgalactiae subsp. equisimilis strain KNZ03 | purB | Adenylosuccinate lyase (EC 4.3.2.2) @ SAICAR lyase (EC 4.3.2.2) | 11359563 | 99 | 100 | 94 | 1e-237 |
| Streptococcus dysgalactiae subsp. equisimilis strain KNZ04 | purB | Adenylosuccinate lyase (EC 4.3.2.2) @ SAICAR lyase (EC 4.3.2.2) | 11359563 | 99 | 100 | 94 | 1e-237 |
| Streptococcus dysgalactiae subsp. equisimilis strain KNZ06 | purB | Adenylosuccinate lyase (EC 4.3.2.2) @ SAICAR lyase (EC 4.3.2.2) | 11359563 | 99 | 100 | 94 | 1e-237 |
| Streptococcus dysgalactiae subsp. equisimilis strain KNZ07 | purB | Adenylosuccinate lyase (EC 4.3.2.2) @ SAICAR lyase (EC 4.3.2.2) | 11359563 | 98 | 99 | 94 | 1e-236 |
| Streptococcus dysgalactiae subsp. equisimilis strain KNZ10 | purB | Adenylosuccinate lyase (EC 4.3.2.2) @ SAICAR lyase (EC 4.3.2.2) | 11359563 | 98 | 99 | 94 | 1e-236 |
| Streptococcus dysgalactiae subsp. equisimilis strain KNZ12 | purB | Adenylosuccinate lyase (EC 4.3.2.2) @ SAICAR lyase (EC 4.3.2.2) | 11359563 | 98 | 99 | 94 | 1e-236 |
| Streptococcus dysgalactiae subsp. equisimilis strain KNZ15 | purB | Adenylosuccinate lyase (EC 4.3.2.2) @ SAICAR lyase (EC 4.3.2.2) | 11359563 | 98 | 99 | 94 | 1e-236 |
| Streptococcus dysgalactiae subsp. equisimilis strain KNZ16 | purB | Adenylosuccinate lyase (EC 4.3.2.2) @ SAICAR lyase (EC 4.3.2.2) | 11359563 | 99 | 100 | 94 | 1e-237 |
| Streptococcus dysgalactiae subsp. equisimilis strain NCTC10321 | purB | Adenylosuccinate lyase (EC 4.3.2.2) @ SAICAR lyase (EC 4.3.2.2) | 11359563 | 98 | 99 | 94 | 1e-236 |
| Streptococcus dysgalactiae subsp. equisimilis strain NCTC11554 | purB | Adenylosuccinate lyase (EC 4.3.2.2) @ SAICAR lyase (EC 4.3.2.2) | 11359563 | 99 | 100 | 94 | 1e-237 |
| Streptococcus dysgalactiae subsp. equisimilis strain NCTC11555 | purB | Adenylosuccinate lyase (EC 4.3.2.2) @ SAICAR lyase (EC 4.3.2.2) | 11359563 | 97 | 100 | 94 | 1e-232 |
| Streptococcus dysgalactiae subsp. equisimilis strain NCTC11556 | purB | Adenylosuccinate lyase (EC 4.3.2.2) @ SAICAR lyase (EC 4.3.2.2) | 11359563 | 99 | 100 | 94 | 1e-237 |
| Streptococcus dysgalactiae subsp. equisimilis strain NCTC11557 | purB | Adenylosuccinate lyase (EC 4.3.2.2) @ SAICAR lyase (EC 4.3.2.2) | 11359563 | 99 | 100 | 93 | 1e-237 |
| Streptococcus dysgalactiae subsp. equisimilis strain NCTC11564 | purB | Adenylosuccinate lyase (EC 4.3.2.2) @ SAICAR lyase (EC 4.3.2.2) | 11359563 | 99 | 100 | 93 | 1e-237 |
| Streptococcus dysgalactiae subsp. equisimilis strain NCTC5370 | purB | Adenylosuccinate lyase (EC 4.3.2.2) @ SAICAR lyase (EC 4.3.2.2) | 11359563 | 98 | 99 | 94 | 1e-236 |
| Streptococcus dysgalactiae subsp. equisimilis strain NCTC5371 | purB | Adenylosuccinate lyase (EC 4.3.2.2) @ SAICAR lyase (EC 4.3.2.2) | 11359563 | 99 | 100 | 94 | 1e-237 |
| Streptococcus dysgalactiae subsp. equisimilis strain NCTC6179 | purB | Adenylosuccinate lyase (EC 4.3.2.2) @ SAICAR lyase (EC 4.3.2.2) | 11359563 | 98 | 99 | 93 | 1e-235 |
| Streptococcus dysgalactiae subsp. equisimilis strain NCTC6181 | purB | Adenylosuccinate lyase (EC 4.3.2.2) @ SAICAR lyase (EC 4.3.2.2) | 11359563 | 98 | 99 | 93 | 1e-235 |
| Streptococcus dysgalactiae subsp. equisimilis strain NCTC6407 | purB | Adenylosuccinate lyase (EC 4.3.2.2) @ SAICAR lyase (EC 4.3.2.2) | 11359563 | 98 | 99 | 94 | 1e-236 |
| Streptococcus dysgalactiae subsp. equisimilis strain NCTC7136 | purB | Adenylosuccinate lyase (EC 4.3.2.2) @ SAICAR lyase (EC 4.3.2.2) | 11359563 | 99 | 100 | 94 | 1e-237 |
| Streptococcus dysgalactiae subsp. equisimilis strain NCTC8543 | purB | Adenylosuccinate lyase (EC 4.3.2.2) @ SAICAR lyase (EC 4.3.2.2) | 11359563 | 98 | 99 | 94 | 1e-236 |
| Streptococcus dysgalactiae subsp. equisimilis strain NCTC8546 | purB | Adenylosuccinate lyase (EC 4.3.2.2) @ SAICAR lyase (EC 4.3.2.2) | 11359563 | 98 | 99 | 94 | 1e-236 |
| Streptococcus dysgalactiae subsp. equisimilis strain NCTC9413 | purB | Adenylosuccinate lyase (EC 4.3.2.2) @ SAICAR lyase (EC 4.3.2.2) | 11359563 | 98 | 99 | 94 | 1e-236 |
| Streptococcus dysgalactiae subsp. equisimilis strain NCTC9414 | purB | Adenylosuccinate lyase (EC 4.3.2.2) @ SAICAR lyase (EC 4.3.2.2) | 11359563 | 99 | 100 | 94 | 1e-237 |
| Streptococcus dysgalactiae subsp. equisimilis strain NCTC9603 | purB | Adenylosuccinate lyase (EC 4.3.2.2) @ SAICAR lyase (EC 4.3.2.2) | 11359563 | 99 | 100 | 93 | 1e-237 |
| Streptococcus dysgalactiae subsp. equisimilis strain SS1575 | purB | Adenylosuccinate lyase (EC 4.3.2.2) @ SAICAR lyase (EC 4.3.2.2) | 11359563 | 99 | 100 | 94 | 1e-237 |
| Streptococcus dysgalactiae subsp. equisimilis strain T642 | purB | Adenylosuccinate lyase (EC 4.3.2.2) @ SAICAR lyase (EC 4.3.2.2) | 11359563 | 99 | 100 | 94 | 0 |
| Streptococcus dysgalactiae subsp. equisimilis strain UT_4031CC | purB | Adenylosuccinate lyase (EC 4.3.2.2) @ SAICAR lyase (EC 4.3.2.2) | 11359563 | 98 | 99 | 94 | 1e-236 |
| Streptococcus dysgalactiae subsp. equisimilis strain UT_4231_KK | purB | Adenylosuccinate lyase (EC 4.3.2.2) @ SAICAR lyase (EC 4.3.2.2) | 11359563 | 98 | 99 | 94 | 1e-236 |
| Streptococcus dysgalactiae subsp. equisimilis strain UT_4234_DH | purB | Adenylosuccinate lyase (EC 4.3.2.2) @ SAICAR lyase (EC 4.3.2.2) | 11359563 | 99 | 100 | 93 | 1e-237 |
| Streptococcus dysgalactiae subsp. equisimilis strain UT_4241_XS | purB | Adenylosuccinate lyase (EC 4.3.2.2) @ SAICAR lyase (EC 4.3.2.2) | 11359563 | 98 | 99 | 94 | 1e-236 |
| Streptococcus dysgalactiae subsp. equisimilis strain UT_4242_AB | purB | Adenylosuccinate lyase (EC 4.3.2.2) @ SAICAR lyase (EC 4.3.2.2) | 11359563 | 99 | 100 | 94 | 1e-237 |
| Streptococcus dysgalactiae subsp. equisimilis strain UT_4255RC | purB | Adenylosuccinate lyase (EC 4.3.2.2) @ SAICAR lyase (EC 4.3.2.2) | 11359563 | 99 | 100 | 93 | 1e-237 |
| Streptococcus dysgalactiae subsp. equisimilis strain UT_4277_BB | purB | Adenylosuccinate lyase (EC 4.3.2.2) @ SAICAR lyase (EC 4.3.2.2) | 11359563 | 98 | 99 | 94 | 1e-236 |
| Streptococcus dysgalactiae subsp. equisimilis strain UT_4966_RC | purB | Adenylosuccinate lyase (EC 4.3.2.2) @ SAICAR lyase (EC 4.3.2.2) | 11359563 | 98 | 99 | 94 | 1e-236 |
| Streptococcus dysgalactiae subsp. equisimilis strain UT-5345 | purB | Adenylosuccinate lyase (EC 4.3.2.2) @ SAICAR lyase (EC 4.3.2.2) | 11359563 | 99 | 100 | 94 | 1e-237 |
| Streptococcus dysgalactiae subsp. equisimilis strain UT-5354 | purB | Adenylosuccinate lyase (EC 4.3.2.2) @ SAICAR lyase (EC 4.3.2.2) | 11359563 | 98 | 99 | 94 | 1e-236 |
| Streptococcus dysgalactiae subsp. equisimilis strain UT-SS1069 | purB | Adenylosuccinate lyase (EC 4.3.2.2) @ SAICAR lyase (EC 4.3.2.2) | 11359563 | 98 | 99 | 94 | 1e-236 |
| Streptococcus dysgalactiae subsp. equisimilis strain UT-SS957 | purB | Adenylosuccinate lyase (EC 4.3.2.2) @ SAICAR lyase (EC 4.3.2.2) | 11359563 | 99 | 100 | 94 | 1e-237 |
| Streptococcus dysgalactiae subsp. equisimilis strain WCHSDSE-1 | purB | Adenylosuccinate lyase (EC 4.3.2.2) @ SAICAR lyase (EC 4.3.2.2) | 11359563 | 99 | 100 | 94 | 1e-237 |
| Streptococcus dysgalactiae subsp. equisimilis 167 | csoS1A | Ethanolamine utilization protein similar to PduA/PduJ | 17307944 | 88 | 88 | 85 | 4e-35 |
| Streptococcus dysgalactiae subsp. equisimilis ATCC 12394 | csoS1A | Ethanolamine utilization protein similar to PduA/PduJ | 17307944 | 88 | 88 | 85 | 4e-35 |
| Streptococcus dysgalactiae subsp. equisimilis RE378 | csoS1A | Ethanolamine utilization protein similar to PduA/PduJ | 17307944 | 88 | 88 | 85 | 4e-35 |
| Streptococcus dysgalactiae subsp. equisimilis SK1249 | csoS1A | Ethanolamine utilization protein similar to PduA/PduJ | 17307944 | 88 | 88 | 85 | 4e-35 |
| Streptococcus dysgalactiae subsp. equisimilis strain ASDSE_96 | csoS1A | Ethanolamine utilization protein similar to PduA/PduJ | 17307944 | 88 | 89 | 85 | 4e-35 |
| Streptococcus dysgalactiae subsp. equisimilis strain C161L1 | csoS1A | Ethanolamine utilization protein similar to PduA/PduJ | 17307944 | 95 | 96 | 80 | 3e-41 |
| Streptococcus dysgalactiae subsp. equisimilis strain KNZ01 | EF1623 | Ethanolamine utilization protein similar to PduA/PduJ | 17307944 | 88 | 89 | 85 | 4e-35 |
| Streptococcus dysgalactiae subsp. equisimilis strain KNZ03 | EF1623 | Ethanolamine utilization protein similar to PduA/PduJ | 17307944 | 88 | 89 | 85 | 4e-35 |
| Streptococcus dysgalactiae subsp. equisimilis strain KNZ04 | EF1623 | Ethanolamine utilization protein similar to PduA/PduJ | 17307944 | 88 | 89 | 85 | 4e-35 |
| Streptococcus dysgalactiae subsp. equisimilis strain KNZ07 | EF1623 | Ethanolamine utilization protein similar to PduA/PduJ | 17307944 | 88 | 89 | 85 | 4e-35 |
| Streptococcus dysgalactiae subsp. equisimilis strain KNZ10 | EF1623 | Ethanolamine utilization protein similar to PduA/PduJ | 17307944 | 88 | 89 | 85 | 4e-35 |
| Streptococcus dysgalactiae subsp. equisimilis strain KNZ12 | EF1623 | Ethanolamine utilization protein similar to PduA/PduJ | 17307944 | 88 | 89 | 85 | 4e-35 |
| Streptococcus dysgalactiae subsp. equisimilis strain KNZ15 | EF1623 | Ethanolamine utilization protein similar to PduA/PduJ | 17307944 | 88 | 89 | 85 | 4e-35 |
| Streptococcus dysgalactiae subsp. equisimilis strain KNZ16 | csoS1A | Ethanolamine utilization protein similar to PduA/PduJ | 17307944 | 88 | 89 | 85 | 4e-35 |
| Streptococcus dysgalactiae subsp. equisimilis strain NCTC11554 | EF1623 | Ethanolamine utilization protein similar to PduA/PduJ | 17307944 | 88 | 89 | 85 | 4e-35 |
| Streptococcus dysgalactiae subsp. equisimilis strain NCTC11564 | EF1623 | Ethanolamine utilization protein similar to PduA/PduJ | 17307944 | 88 | 89 | 85 | 4e-35 |
| Streptococcus dysgalactiae subsp. equisimilis strain NCTC5370 | EF1623 | Ethanolamine utilization protein similar to PduA/PduJ | 17307944 | 88 | 89 | 85 | 4e-35 |
| Streptococcus dysgalactiae subsp. equisimilis strain NCTC6179 | EF1623 | Ethanolamine utilization protein similar to PduA/PduJ | 17307944 | 88 | 89 | 85 | 4e-35 |
| Streptococcus dysgalactiae subsp. equisimilis strain NCTC6181 | EF1623 | Ethanolamine utilization protein similar to PduA/PduJ | 17307944 | 88 | 89 | 85 | 4e-35 |
| Streptococcus dysgalactiae subsp. equisimilis strain NCTC8546 | EF1623 | Ethanolamine utilization protein similar to PduA/PduJ | 17307944 | 88 | 89 | 85 | 4e-35 |
| Streptococcus dysgalactiae subsp. equisimilis strain NCTC9413 | EF1623 | Ethanolamine utilization protein similar to PduA/PduJ | 17307944 | 88 | 89 | 85 | 4e-35 |
| Streptococcus dysgalactiae subsp. equisimilis strain SS1575 | EF1623 | Ethanolamine utilization protein similar to PduA/PduJ | 17307944 | 88 | 89 | 85 | 4e-35 |
| Streptococcus dysgalactiae subsp. equisimilis strain UT_4031CC | EF1623 | Ethanolamine utilization protein similar to PduA/PduJ | 17307944 | 88 | 89 | 85 | 4e-35 |
| Streptococcus dysgalactiae subsp. equisimilis strain UT_4231_KK | EF1623 | Ethanolamine utilization protein similar to PduA/PduJ | 17307944 | 88 | 89 | 85 | 4e-35 |
| Streptococcus dysgalactiae subsp. equisimilis strain UT_4241_XS | EF1623 | Ethanolamine utilization protein similar to PduA/PduJ | 17307944 | 88 | 89 | 85 | 4e-35 |
| Streptococcus dysgalactiae subsp. equisimilis strain UT_4966_RC | EF1623 | Ethanolamine utilization protein similar to PduA/PduJ | 17307944 | 88 | 89 | 85 | 4e-35 |
| Streptococcus dysgalactiae subsp. equisimilis strain UT-5345 | EF1623 | Ethanolamine utilization protein similar to PduA/PduJ | 17307944 | 88 | 89 | 85 | 4e-35 |
| Streptococcus dysgalactiae subsp. equisimilis strain UT-SS1069 | EF1623 | Ethanolamine utilization protein similar to PduA/PduJ | 17307944 | 88 | 89 | 85 | 4e-35 |
| Streptococcus dysgalactiae subsp. equisimilis AC-2713 | speG | Exotoxin, phage associated @ Streptococcal pyrogenic exotoxin G (SpeG) | 23380619 | 100 | 100 | 80 | 1e-105 |
| Streptococcus dysgalactiae subsp. equisimilis AKSDE4288 | speg | Exotoxin, phage associated @ Streptococcal pyrogenic exotoxin G (SpeG) | 23380619 | 100 | 100 | 80 | 1e-105 |
| Streptococcus dysgalactiae subsp. equisimilis GGS_124 | spegg4 | Exotoxin, phage associated @ Streptococcal pyrogenic exotoxin G (SpeG) | 23380619 | 100 | 100 | 80 | 1e-105 |
| Streptococcus dysgalactiae subsp. equisimilis SD SCDR1 | speg | Exotoxin, phage associated @ Streptococcal pyrogenic exotoxin G (SpeG) | 23380619 | 100 | 100 | 80 | 1e-106 |
| Streptococcus dysgalactiae subsp. equisimilis SK1250 | speg | Exotoxin, phage associated @ Streptococcal pyrogenic exotoxin G (SpeG) | 23380619 | 100 | 100 | 80 | 1e-106 |
| Streptococcus dysgalactiae subsp. equisimilis strain ASDSE_99 | speg | Exotoxin, phage associated @ Streptococcal pyrogenic exotoxin G (SpeG) | 23380619 | 100 | 100 | 80 | 1e-105 |
| Streptococcus dysgalactiae subsp. equisimilis strain C161L1 | speg | Exotoxin, phage associated @ Streptococcal pyrogenic exotoxin G (SpeG) | 23380619 | 100 | 100 | 80 | 9e-136 |
| Streptococcus dysgalactiae subsp. equisimilis strain NCTC10321 | speg | Exotoxin, phage associated @ Streptococcal pyrogenic exotoxin G (SpeG) | 23380619 | 100 | 100 | 81 | 1e-107 |
| Streptococcus dysgalactiae subsp. equisimilis strain NCTC11556 | speg | Exotoxin, phage associated @ Streptococcal pyrogenic exotoxin G (SpeG) | 23380619 | 100 | 100 | 80 | 1e-106 |
| Streptococcus dysgalactiae subsp. equisimilis strain NCTC11557 | speg | Exotoxin, phage associated @ Streptococcal pyrogenic exotoxin G (SpeG) | 23380619 | 100 | 100 | 80 | 1e-106 |
| Streptococcus dysgalactiae subsp. equisimilis strain NCTC5370 | speg | Exotoxin, phage associated @ Streptococcal pyrogenic exotoxin G (SpeG) | 23380619 | 100 | 100 | 80 | 1e-106 |
| Streptococcus dysgalactiae subsp. equisimilis strain NCTC5371 | speg | Exotoxin, phage associated @ Streptococcal pyrogenic exotoxin G (SpeG) | 23380619 | 85 | 96 | 80 | 4e-88 |
| Streptococcus dysgalactiae subsp. equisimilis strain NCTC6407 | speg | Exotoxin, phage associated @ Streptococcal pyrogenic exotoxin G (SpeG) | 23380619 | 100 | 100 | 81 | 1e-107 |
| Streptococcus dysgalactiae subsp. equisimilis strain NCTC7136 | speg | Exotoxin, phage associated @ Streptococcal pyrogenic exotoxin G (SpeG) | 23380619 | 100 | 100 | 80 | 1e-105 |
| Streptococcus dysgalactiae subsp. equisimilis strain NCTC9414 | speg | Exotoxin, phage associated @ Streptococcal pyrogenic exotoxin G (SpeG) | 23380619 | 85 | 96 | 80 | 4e-88 |
| Streptococcus dysgalactiae subsp. equisimilis strain NCTC9603 | speg | Exotoxin, phage associated @ Streptococcal pyrogenic exotoxin G (SpeG) | 23380619 | 100 | 100 | 80 | 1e-105 |
| Streptococcus dysgalactiae subsp. equisimilis strain T642 | speg | Exotoxin, phage associated @ Streptococcal pyrogenic exotoxin G (SpeG) | 23380619 | 100 | 100 | 80 | 1e-135 |
| Streptococcus dysgalactiae subsp. equisimilis strain UT_4234_DH | speg | Exotoxin, phage associated @ Streptococcal pyrogenic exotoxin G (SpeG) | 23380619 | 100 | 100 | 80 | 1e-106 |
| Streptococcus dysgalactiae subsp. equisimilis strain UT_4241_XS | speg | Exotoxin, phage associated @ Streptococcal pyrogenic exotoxin G (SpeG) | 23380619 | 100 | 100 | 80 | 1e-106 |
| Streptococcus dysgalactiae subsp. equisimilis strain UT_4242_AB | speg | Exotoxin, phage associated @ Streptococcal pyrogenic exotoxin G (SpeG) | 23380619 | 100 | 100 | 80 | 1e-106 |
| Streptococcus dysgalactiae subsp. equisimilis strain UT_4255RC | speg | Exotoxin, phage associated @ Streptococcal pyrogenic exotoxin G (SpeG) | 23380619 | 100 | 100 | 80 | 1e-106 |
| Streptococcus dysgalactiae subsp. equisimilis strain UT_4966_RC | speg | Exotoxin, phage associated @ Streptococcal pyrogenic exotoxin G (SpeG) | 23380619 | 100 | 100 | 80 | 1e-106 |
| Streptococcus dysgalactiae subsp. equisimilis strain UT-SS1069 | speg | Exotoxin, phage associated @ Streptococcal pyrogenic exotoxin G (SpeG) | 23380619 | 100 | 100 | 80 | 1e-106 |
| Streptococcus dysgalactiae subsp. equisimilis strain UT-SS957 | speg | Exotoxin, phage associated @ Streptococcal pyrogenic exotoxin G (SpeG) | 23380619 | 85 | 96 | 80 | 4e-88 |
| Streptococcus dysgalactiae subsp. equisimilis strain WCHSDSE-1 | speg | Exotoxin, phage associated @ Streptococcal pyrogenic exotoxin G (SpeG) | 23380619 | 100 | 100 | 80 | 1e-106 |
| Streptococcus dysgalactiae subsp. equisimilis 167 | fbp | Fibronectin/fibrinogen-binding protein | 11580843 | 99 | 99 | 93 | 1e-301 |
| Streptococcus dysgalactiae subsp. equisimilis AC-2713 | flpA | Fibronectin/fibrinogen-binding protein | 11580843 | 99 | 99 | 93 | 1e-302 |
| Streptococcus dysgalactiae subsp. equisimilis AKSDE4288 | fbp54 | Fibronectin/fibrinogen-binding protein | 11580843 | 99 | 99 | 93 | 1e-302 |
| Streptococcus dysgalactiae subsp. equisimilis ATCC 12394 | fbp54 | Fibronectin/fibrinogen-binding protein | 11580843 | 99 | 99 | 93 | 1e-302 |
| Streptococcus dysgalactiae subsp. equisimilis GGS_124 | fbp | Fibronectin/fibrinogen-binding protein | 11580843 | 99 | 99 | 93 | 1e-302 |
| Streptococcus dysgalactiae subsp. equisimilis RE378 | fbp | Fibronectin/fibrinogen-binding protein | 11580843 | 99 | 99 | 93 | 1e-302 |
| Streptococcus dysgalactiae subsp. equisimilis SD SCDR1 | fbp54 | Fibronectin/fibrinogen-binding protein | 11580843 | 99 | 99 | 93 | 1e-302 |
| Streptococcus dysgalactiae subsp. equisimilis SK1249 | fbp54 | Fibronectin/fibrinogen-binding protein | 11580843 | 93 | 97 | 94 | 1e-283 |
| Streptococcus dysgalactiae subsp. equisimilis SK1250 | fbp54 | Fibronectin/fibrinogen-binding protein | 11580843 | 99 | 99 | 93 | 1e-302 |
| Streptococcus dysgalactiae subsp. equisimilis strain ASDSE_96 | fbp54 | Fibronectin/fibrinogen-binding protein | 11580843 | 99 | 99 | 93 | 1e-301 |
| Streptococcus dysgalactiae subsp. equisimilis strain ASDSE_99 | fbp54 | Fibronectin/fibrinogen-binding protein | 11580843 | 99 | 99 | 93 | 1e-302 |
| Streptococcus dysgalactiae subsp. equisimilis strain C161L1 | fbp54 | Fibronectin/fibrinogen-binding protein | 11580843 | 99 | 99 | 93 | 0 |
| Streptococcus dysgalactiae subsp. equisimilis strain KNZ01 | fbp54 | Fibronectin/fibrinogen-binding protein | 11580843 | 99 | 94 | 93 | 1e-302 |
| Streptococcus dysgalactiae subsp. equisimilis strain KNZ03 | fbp54 | Fibronectin/fibrinogen-binding protein | 11580843 | 99 | 99 | 93 | 1e-302 |
| Streptococcus dysgalactiae subsp. equisimilis strain KNZ04 | fbp54 | Fibronectin/fibrinogen-binding protein | 11580843 | 99 | 99 | 93 | 1e-302 |
| Streptococcus dysgalactiae subsp. equisimilis strain KNZ06 | fbp54 | Fibronectin/fibrinogen-binding protein | 11580843 | 99 | 99 | 93 | 1e-301 |
| Streptococcus dysgalactiae subsp. equisimilis strain KNZ07 | fbp54 | Fibronectin/fibrinogen-binding protein | 11580843 | 99 | 94 | 93 | 1e-302 |
| Streptococcus dysgalactiae subsp. equisimilis strain KNZ10 | fbp54 | Fibronectin/fibrinogen-binding protein | 11580843 | 99 | 94 | 93 | 1e-302 |
| Streptococcus dysgalactiae subsp. equisimilis strain KNZ12 | fbp54 | Fibronectin/fibrinogen-binding protein | 11580843 | 99 | 94 | 93 | 1e-302 |
| Streptococcus dysgalactiae subsp. equisimilis strain KNZ15 | fbp54 | Fibronectin/fibrinogen-binding protein | 11580843 | 99 | 99 | 93 | 1e-302 |
| Streptococcus dysgalactiae subsp. equisimilis strain KNZ16 | fbp54 | Fibronectin/fibrinogen-binding protein | 11580843 | 99 | 99 | 93 | 1e-301 |
| Streptococcus dysgalactiae subsp. equisimilis strain NCTC10321 | fbp54 | Fibronectin/fibrinogen-binding protein | 11580843 | 99 | 99 | 94 | 1e-303 |
| Streptococcus dysgalactiae subsp. equisimilis strain NCTC11554 | fbp54 | Fibronectin/fibrinogen-binding protein | 11580843 | 99 | 99 | 93 | 1e-301 |
| Streptococcus dysgalactiae subsp. equisimilis strain NCTC11555 | fbp54 | Fibronectin/fibrinogen-binding protein | 11580843 | 99 | 99 | 93 | 1e-302 |
| Streptococcus dysgalactiae subsp. equisimilis strain NCTC11556 | fbp54 | Fibronectin/fibrinogen-binding protein | 11580843 | 99 | 99 | 93 | 1e-302 |
| Streptococcus dysgalactiae subsp. equisimilis strain NCTC11557 | fbp54 | Fibronectin/fibrinogen-binding protein | 11580843 | 99 | 99 | 93 | 1e-302 |
| Streptococcus dysgalactiae subsp. equisimilis strain NCTC11564 | fbp54 | Fibronectin/fibrinogen-binding protein | 11580843 | 99 | 99 | 93 | 1e-301 |
| Streptococcus dysgalactiae subsp. equisimilis strain NCTC5370 | fbp54 | Fibronectin/fibrinogen-binding protein | 11580843 | 99 | 99 | 93 | 1e-301 |
| Streptococcus dysgalactiae subsp. equisimilis strain NCTC5371 | fbp54 | Fibronectin/fibrinogen-binding protein | 11580843 | 99 | 99 | 93 | 1e-302 |
| Streptococcus dysgalactiae subsp. equisimilis strain NCTC6179 | fbp54 | Fibronectin/fibrinogen-binding protein | 11580843 | 99 | 99 | 93 | 1e-301 |
| Streptococcus dysgalactiae subsp. equisimilis strain NCTC6181 | fbp54 | Fibronectin/fibrinogen-binding protein | 11580843 | 99 | 99 | 93 | 1e-301 |
| Streptococcus dysgalactiae subsp. equisimilis strain NCTC6407 | fbp54 | Fibronectin/fibrinogen-binding protein | 11580843 | 99 | 99 | 94 | 1e-303 |
| Streptococcus dysgalactiae subsp. equisimilis strain NCTC7136 | fbp54 | Fibronectin/fibrinogen-binding protein | 11580843 | 99 | 99 | 93 | 1e-301 |
| Streptococcus dysgalactiae subsp. equisimilis strain NCTC8543 | fbp54 | Fibronectin/fibrinogen-binding protein | 11580843 | 99 | 99 | 93 | 1e-301 |
| Streptococcus dysgalactiae subsp. equisimilis strain NCTC8546 | fbp54 | Fibronectin/fibrinogen-binding protein | 11580843 | 99 | 99 | 93 | 1e-301 |
| Streptococcus dysgalactiae subsp. equisimilis strain NCTC9413 | fbp54 | Fibronectin/fibrinogen-binding protein | 11580843 | 99 | 99 | 93 | 1e-301 |
| Streptococcus dysgalactiae subsp. equisimilis strain NCTC9414 | fbp54 | Fibronectin/fibrinogen-binding protein | 11580843 | 99 | 99 | 93 | 1e-302 |
| Streptococcus dysgalactiae subsp. equisimilis strain NCTC9603 | fbp54 | Fibronectin/fibrinogen-binding protein | 11580843 | 99 | 99 | 93 | 1e-301 |
| Streptococcus dysgalactiae subsp. equisimilis strain SS1575 | fbp54 | Fibronectin/fibrinogen-binding protein | 11580843 | 99 | 99 | 93 | 1e-301 |
| Streptococcus dysgalactiae subsp. equisimilis strain T642 | fbp54 | Fibronectin/fibrinogen-binding protein | 11580843 | 99 | 99 | 93 | 0 |
| Streptococcus dysgalactiae subsp. equisimilis strain UT_4031CC | fbp54 | Fibronectin/fibrinogen-binding protein | 11580843 | 99 | 99 | 93 | 1e-301 |
| Streptococcus dysgalactiae subsp. equisimilis strain UT_4231_KK | fbp54 | Fibronectin/fibrinogen-binding protein | 11580843 | 99 | 99 | 93 | 1e-301 |
| Streptococcus dysgalactiae subsp. equisimilis strain UT_4234_DH | fbp54 | Fibronectin/fibrinogen-binding protein | 11580843 | 99 | 99 | 93 | 1e-302 |
| Streptococcus dysgalactiae subsp. equisimilis strain UT_4241_XS | fbp54 | Fibronectin/fibrinogen-binding protein | 11580843 | 99 | 99 | 93 | 1e-301 |
| Streptococcus dysgalactiae subsp. equisimilis strain UT_4242_AB | fbp54 | Fibronectin/fibrinogen-binding protein | 11580843 | 99 | 99 | 93 | 1e-302 |
| Streptococcus dysgalactiae subsp. equisimilis strain UT_4255RC | fbp54 | Fibronectin/fibrinogen-binding protein | 11580843 | 99 | 99 | 93 | 1e-302 |
| Streptococcus dysgalactiae subsp. equisimilis strain UT_4277_BB | fbp54 | Fibronectin/fibrinogen-binding protein | 11580843 | 99 | 99 | 93 | 1e-302 |
| Streptococcus dysgalactiae subsp. equisimilis strain UT_4966_RC | fbp54 | Fibronectin/fibrinogen-binding protein | 11580843 | 99 | 99 | 93 | 1e-301 |
| Streptococcus dysgalactiae subsp. equisimilis strain UT-5345 | fbp54 | Fibronectin/fibrinogen-binding protein | 11580843 | 99 | 99 | 93 | 1e-302 |
| Streptococcus dysgalactiae subsp. equisimilis strain UT-5354 | fbp54 | Fibronectin/fibrinogen-binding protein | 11580843 | 99 | 99 | 93 | 1e-301 |
| Streptococcus dysgalactiae subsp. equisimilis strain UT-SS1069 | fbp54 | Fibronectin/fibrinogen-binding protein | 11580843 | 63 | 100 | 92 | 1e-186 |
| Streptococcus dysgalactiae subsp. equisimilis strain UT-SS957 | fbp54 | Fibronectin/fibrinogen-binding protein | 11580843 | 99 | 99 | 93 | 1e-301 |
| Streptococcus dysgalactiae subsp. equisimilis strain WCHSDSE-1 | fbp54 | Fibronectin/fibrinogen-binding protein | 11580843 | 99 | 99 | 93 | 1e-302 |
| Streptococcus dysgalactiae subsp. equisimilis 167 | cpsY | Methionine biosynthesis and transport regulator MtaR, LysR family | 14594832 | 92 | 98 | 83 | 1e-135 |
| Streptococcus dysgalactiae subsp. equisimilis AC-2713 | cpsY | Methionine biosynthesis and transport regulator MtaR, LysR family | 14594832 | 92 | 98 | 82 | 1e-134 |
| Streptococcus dysgalactiae subsp. equisimilis AKSDE4288 | cpsY | Methionine biosynthesis and transport regulator MtaR, LysR family | 14594832 | 97 | 98 | 82 | 1e-142 |
| Streptococcus dysgalactiae subsp. equisimilis ATCC 12394 | cpsY | Methionine biosynthesis and transport regulator MtaR, LysR family | 14594832 | 92 | 98 | 82 | 1e-134 |
| Streptococcus dysgalactiae subsp. equisimilis GGS_124 | cpsY | Methionine biosynthesis and transport regulator MtaR, LysR family | 14594832 | 92 | 98 | 82 | 1e-134 |
| Streptococcus dysgalactiae subsp. equisimilis RE378 | cpsY | Methionine biosynthesis and transport regulator MtaR, LysR family | 14594832 | 92 | 98 | 83 | 1e-135 |
| Streptococcus dysgalactiae subsp. equisimilis SD SCDR1 | cpsY | Methionine biosynthesis and transport regulator MtaR, LysR family | 14594832 | 97 | 98 | 82 | 1e-142 |
| Streptococcus dysgalactiae subsp. equisimilis SK1249 | cpsY | Methionine biosynthesis and transport regulator MtaR, LysR family | 14594832 | 92 | 98 | 83 | 1e-135 |
| Streptococcus dysgalactiae subsp. equisimilis SK1250 | cpsY | Methionine biosynthesis and transport regulator MtaR, LysR family | 14594832 | 92 | 98 | 82 | 1e-134 |
| Streptococcus dysgalactiae subsp. equisimilis strain ASDSE_96 | cpsY | Methionine biosynthesis and transport regulator MtaR, LysR family | 14594832 | 97 | 98 | 82 | 1e-142 |
| Streptococcus dysgalactiae subsp. equisimilis strain ASDSE_99 | cpsY | Methionine biosynthesis and transport regulator MtaR, LysR family | 14594832 | 97 | 98 | 83 | 1e-143 |
| Streptococcus dysgalactiae subsp. equisimilis strain C161L1 | cpsY | Methionine biosynthesis and transport regulator MtaR, LysR family | 14594832 | 98 | 99 | 82 | 0 |
| Streptococcus dysgalactiae subsp. equisimilis strain KNZ01 | cpsY | Methionine biosynthesis and transport regulator MtaR, LysR family | 14594832 | 97 | 98 | 83 | 1e-143 |
| Streptococcus dysgalactiae subsp. equisimilis strain KNZ03 | cpsY | Methionine biosynthesis and transport regulator MtaR, LysR family | 14594832 | 97 | 98 | 82 | 1e-142 |
| Streptococcus dysgalactiae subsp. equisimilis strain KNZ04 | cpsY | Methionine biosynthesis and transport regulator MtaR, LysR family | 14594832 | 97 | 98 | 82 | 1e-142 |
| Streptococcus dysgalactiae subsp. equisimilis strain KNZ06 | cpsY | Methionine biosynthesis and transport regulator MtaR, LysR family | 14594832 | 97 | 98 | 83 | 1e-143 |
| Streptococcus dysgalactiae subsp. equisimilis strain KNZ07 | cpsY | Methionine biosynthesis and transport regulator MtaR, LysR family | 14594832 | 97 | 98 | 83 | 1e-143 |
| Streptococcus dysgalactiae subsp. equisimilis strain KNZ10 | cpsY | Methionine biosynthesis and transport regulator MtaR, LysR family | 14594832 | 97 | 98 | 83 | 1e-143 |
| Streptococcus dysgalactiae subsp. equisimilis strain KNZ12 | cpsY | Methionine biosynthesis and transport regulator MtaR, LysR family | 14594832 | 97 | 98 | 83 | 1e-143 |
| Streptococcus dysgalactiae subsp. equisimilis strain KNZ15 | cpsY | Methionine biosynthesis and transport regulator MtaR, LysR family | 14594832 | 97 | 98 | 83 | 1e-143 |
| Streptococcus dysgalactiae subsp. equisimilis strain KNZ16 | cpsY | Methionine biosynthesis and transport regulator MtaR, LysR family | 14594832 | 97 | 98 | 82 | 1e-142 |
| Streptococcus dysgalactiae subsp. equisimilis strain NCTC10321 | cpsY | Methionine biosynthesis and transport regulator MtaR, LysR family | 14594832 | 97 | 98 | 83 | 1e-142 |
| Streptococcus dysgalactiae subsp. equisimilis strain NCTC11554 | cpsY | Methionine biosynthesis and transport regulator MtaR, LysR family | 14594832 | 97 | 98 | 82 | 1e-142 |
| Streptococcus dysgalactiae subsp. equisimilis strain NCTC11555 | cpsY | Methionine biosynthesis and transport regulator MtaR, LysR family | 14594832 | 97 | 98 | 83 | 1e-143 |
| Streptococcus dysgalactiae subsp. equisimilis strain NCTC11556 | cpsY | Methionine biosynthesis and transport regulator MtaR, LysR family | 14594832 | 97 | 98 | 82 | 1e-142 |
| Streptococcus dysgalactiae subsp. equisimilis strain NCTC11557 | cpsY | Methionine biosynthesis and transport regulator MtaR, LysR family | 14594832 | 97 | 98 | 83 | 1e-143 |
| Streptococcus dysgalactiae subsp. equisimilis strain NCTC11564 | cpsY | Methionine biosynthesis and transport regulator MtaR, LysR family | 14594832 | 97 | 98 | 83 | 1e-143 |
| Streptococcus dysgalactiae subsp. equisimilis strain NCTC5370 | cpsY | Methionine biosynthesis and transport regulator MtaR, LysR family | 14594832 | 97 | 98 | 83 | 1e-143 |
| Streptococcus dysgalactiae subsp. equisimilis strain NCTC5371 | cpsY | Methionine biosynthesis and transport regulator MtaR, LysR family | 14594832 | 97 | 98 | 83 | 1e-143 |
| Streptococcus dysgalactiae subsp. equisimilis strain NCTC6179 | cpsY | Methionine biosynthesis and transport regulator MtaR, LysR family | 14594832 | 97 | 98 | 83 | 1e-143 |
| Streptococcus dysgalactiae subsp. equisimilis strain NCTC6181 | cpsY | Methionine biosynthesis and transport regulator MtaR, LysR family | 14594832 | 97 | 98 | 83 | 1e-143 |
| Streptococcus dysgalactiae subsp. equisimilis strain NCTC6407 | cpsY | Methionine biosynthesis and transport regulator MtaR, LysR family | 14594832 | 97 | 98 | 83 | 1e-142 |
| Streptococcus dysgalactiae subsp. equisimilis strain NCTC7136 | cpsY | Methionine biosynthesis and transport regulator MtaR, LysR family | 14594832 | 97 | 98 | 83 | 1e-143 |
| Streptococcus dysgalactiae subsp. equisimilis strain NCTC8543 | cpsY | Methionine biosynthesis and transport regulator MtaR, LysR family | 14594832 | 97 | 98 | 83 | 1e-143 |
| Streptococcus dysgalactiae subsp. equisimilis strain NCTC8546 | cpsY | Methionine biosynthesis and transport regulator MtaR, LysR family | 14594832 | 97 | 98 | 83 | 1e-143 |
| Streptococcus dysgalactiae subsp. equisimilis strain NCTC9413 | cpsY | Methionine biosynthesis and transport regulator MtaR, LysR family | 14594832 | 97 | 98 | 83 | 1e-143 |
| Streptococcus dysgalactiae subsp. equisimilis strain NCTC9414 | cpsY | Methionine biosynthesis and transport regulator MtaR, LysR family | 14594832 | 97 | 98 | 83 | 1e-143 |
| Streptococcus dysgalactiae subsp. equisimilis strain NCTC9603 | cpsY | Methionine biosynthesis and transport regulator MtaR, LysR family | 14594832 | 97 | 98 | 82 | 1e-142 |
| Streptococcus dysgalactiae subsp. equisimilis strain SS1575 | cpsY | Methionine biosynthesis and transport regulator MtaR, LysR family | 14594832 | 97 | 98 | 83 | 1e-143 |
| Streptococcus dysgalactiae subsp. equisimilis strain T642 | cpsY | Methionine biosynthesis and transport regulator MtaR, LysR family | 14594832 | 98 | 99 | 82 | 0 |
| Streptococcus dysgalactiae subsp. equisimilis strain UT_4031CC | cpsY | Methionine biosynthesis and transport regulator MtaR, LysR family | 14594832 | 97 | 98 | 82 | 1e-142 |
| Streptococcus dysgalactiae subsp. equisimilis strain UT_4231_KK | cpsY | Methionine biosynthesis and transport regulator MtaR, LysR family | 14594832 | 97 | 98 | 83 | 1e-143 |
| Streptococcus dysgalactiae subsp. equisimilis strain UT_4234_DH | cpsY | Methionine biosynthesis and transport regulator MtaR, LysR family | 14594832 | 97 | 98 | 83 | 1e-143 |
| Streptococcus dysgalactiae subsp. equisimilis strain UT_4241_XS | cpsY | Methionine biosynthesis and transport regulator MtaR, LysR family | 14594832 | 97 | 98 | 83 | 1e-143 |
| Streptococcus dysgalactiae subsp. equisimilis strain UT_4242_AB | cpsY | Methionine biosynthesis and transport regulator MtaR, LysR family | 14594832 | 97 | 98 | 82 | 1e-142 |
| Streptococcus dysgalactiae subsp. equisimilis strain UT_4255RC | cpsY | Methionine biosynthesis and transport regulator MtaR, LysR family | 14594832 | 97 | 98 | 83 | 1e-143 |
| Streptococcus dysgalactiae subsp. equisimilis strain UT_4277_BB | cpsY | Methionine biosynthesis and transport regulator MtaR, LysR family | 14594832 | 97 | 98 | 82 | 1e-142 |
| Streptococcus dysgalactiae subsp. equisimilis strain UT_4966_RC | cpsY | Methionine biosynthesis and transport regulator MtaR, LysR family | 14594832 | 97 | 98 | 83 | 1e-143 |
| Streptococcus dysgalactiae subsp. equisimilis strain UT-5345 | cpsY | Methionine biosynthesis and transport regulator MtaR, LysR family | 14594832 | 97 | 98 | 82 | 1e-142 |
| Streptococcus dysgalactiae subsp. equisimilis strain UT-5354 | cpsY | Methionine biosynthesis and transport regulator MtaR, LysR family | 14594832 | 97 | 98 | 83 | 1e-143 |
| Streptococcus dysgalactiae subsp. equisimilis strain UT-SS1069 | cpsY | Methionine biosynthesis and transport regulator MtaR, LysR family | 14594832 | 97 | 98 | 83 | 1e-143 |
| Streptococcus dysgalactiae subsp. equisimilis strain UT-SS957 | cpsY | Methionine biosynthesis and transport regulator MtaR, LysR family | 14594832 | 97 | 98 | 83 | 1e-143 |
| Streptococcus dysgalactiae subsp. equisimilis strain WCHSDSE-1 | cpsY | Methionine biosynthesis and transport regulator MtaR, LysR family | 14594832 | 97 | 98 | 82 | 1e-142 |
| Streptococcus dysgalactiae subsp. equisimilis 167 | nga | Nicotine adenine dinucleotide glycohydrolase (NADGH) (EC 3.2.2.5) | 20470439 | 47 | 100 | 98 | 1e-118 |
| Streptococcus dysgalactiae subsp. equisimilis AC-2713 | nga | Nicotine adenine dinucleotide glycohydrolase (NADGH) (EC 3.2.2.5) | 20470439 | 100 | 100 | 97 | 1e-254 |
| Streptococcus dysgalactiae subsp. equisimilis AKSDE4288 | nga | Nicotine adenine dinucleotide glycohydrolase (NADGH) (EC 3.2.2.5) | 20470439 | 100 | 100 | 97 | 1e-254 |
| Streptococcus dysgalactiae subsp. equisimilis ATCC 12394 | nga | Nicotine adenine dinucleotide glycohydrolase (NADGH) (EC 3.2.2.5) | 20470439 | 100 | 100 | 97 | 1e-254 |
| Streptococcus dysgalactiae subsp. equisimilis GGS_124 | nga | Nicotine adenine dinucleotide glycohydrolase (NADGH) (EC 3.2.2.5) | 20470439 | 100 | 100 | 97 | 1e-253 |
| Streptococcus dysgalactiae subsp. equisimilis RE378 | nga | Nicotine adenine dinucleotide glycohydrolase (NADGH) (EC 3.2.2.5) | 20470439 | 100 | 100 | 97 | 1e-254 |
| Streptococcus dysgalactiae subsp. equisimilis SD SCDR1 | nga | Nicotine adenine dinucleotide glycohydrolase (NADGH) (EC 3.2.2.5) | 20470439 | 100 | 100 | 97 | 1e-254 |
| Streptococcus dysgalactiae subsp. equisimilis SK1249 | nga | Nicotine adenine dinucleotide glycohydrolase (NADGH) (EC 3.2.2.5) | 20470439 | 100 | 100 | 97 | 1e-254 |
| Streptococcus dysgalactiae subsp. equisimilis SK1250 | nga | Nicotine adenine dinucleotide glycohydrolase (NADGH) (EC 3.2.2.5) | 20470439 | 100 | 100 | 97 | 1e-254 |
| Streptococcus dysgalactiae subsp. equisimilis strain ASDSE_96 | nga | Nicotine adenine dinucleotide glycohydrolase (NADGH) (EC 3.2.2.5) | 20470439 | 100 | 100 | 97 | 1e-254 |
| Streptococcus dysgalactiae subsp. equisimilis strain ASDSE_99 | nga | Nicotine adenine dinucleotide glycohydrolase (NADGH) (EC 3.2.2.5) | 20470439 | 100 | 100 | 97 | 1e-254 |
| Streptococcus dysgalactiae subsp. equisimilis strain C161L1 | nga | Nicotine adenine dinucleotide glycohydrolase (NADGH) (EC 3.2.2.5) | 20470439 | 100 | 100 | 97 | 0 |
| Streptococcus dysgalactiae subsp. equisimilis strain KNZ01 | nga | Nicotine adenine dinucleotide glycohydrolase (NADGH) (EC 3.2.2.5) | 20470439 | 100 | 100 | 97 | 1e-254 |
| Streptococcus dysgalactiae subsp. equisimilis strain KNZ03 | nga | Nicotine adenine dinucleotide glycohydrolase (NADGH) (EC 3.2.2.5) | 20470439 | 100 | 100 | 97 | 1e-254 |
| Streptococcus dysgalactiae subsp. equisimilis strain KNZ04 | nga | Nicotine adenine dinucleotide glycohydrolase (NADGH) (EC 3.2.2.5) | 20470439 | 100 | 100 | 97 | 1e-254 |
| Streptococcus dysgalactiae subsp. equisimilis strain KNZ06 | nga | Nicotine adenine dinucleotide glycohydrolase (NADGH) (EC 3.2.2.5) | 20470439 | 100 | 100 | 97 | 1e-254 |
| Streptococcus dysgalactiae subsp. equisimilis strain KNZ07 | nga | Nicotine adenine dinucleotide glycohydrolase (NADGH) (EC 3.2.2.5) | 20470439 | 100 | 100 | 97 | 1e-254 |
| Streptococcus dysgalactiae subsp. equisimilis strain KNZ10 | nga | Nicotine adenine dinucleotide glycohydrolase (NADGH) (EC 3.2.2.5) | 20470439 | 100 | 100 | 97 | 1e-254 |
| Streptococcus dysgalactiae subsp. equisimilis strain KNZ12 | nga | Nicotine adenine dinucleotide glycohydrolase (NADGH) (EC 3.2.2.5) | 20470439 | 100 | 100 | 97 | 1e-254 |
| Streptococcus dysgalactiae subsp. equisimilis strain KNZ15 | nga | Nicotine adenine dinucleotide glycohydrolase (NADGH) (EC 3.2.2.5) | 20470439 | 100 | 100 | 97 | 1e-254 |
| Streptococcus dysgalactiae subsp. equisimilis strain KNZ16 | nga | Nicotine adenine dinucleotide glycohydrolase (NADGH) (EC 3.2.2.5) | 20470439 | 100 | 100 | 97 | 1e-254 |
| Streptococcus dysgalactiae subsp. equisimilis strain NCTC11554 | nga | Nicotine adenine dinucleotide glycohydrolase (NADGH) (EC 3.2.2.5) | 20470439 | 100 | 100 | 97 | 1e-254 |
| Streptococcus dysgalactiae subsp. equisimilis strain NCTC11555 | nga | Nicotine adenine dinucleotide glycohydrolase (NADGH) (EC 3.2.2.5) | 20470439 | 100 | 100 | 97 | 1e-254 |
| Streptococcus dysgalactiae subsp. equisimilis strain NCTC11556 | nga | Nicotine adenine dinucleotide glycohydrolase (NADGH) (EC 3.2.2.5) | 20470439 | 100 | 100 | 97 | 1e-254 |
| Streptococcus dysgalactiae subsp. equisimilis strain NCTC11557 | nga | Nicotine adenine dinucleotide glycohydrolase (NADGH) (EC 3.2.2.5) | 20470439 | 100 | 100 | 97 | 1e-254 |
| Streptococcus dysgalactiae subsp. equisimilis strain NCTC11564 | nga | Nicotine adenine dinucleotide glycohydrolase (NADGH) (EC 3.2.2.5) | 20470439 | 100 | 100 | 97 | 1e-254 |
| Streptococcus dysgalactiae subsp. equisimilis strain NCTC5370 | nga | Nicotine adenine dinucleotide glycohydrolase (NADGH) (EC 3.2.2.5) | 20470439 | 100 | 100 | 97 | 1e-253 |
| Streptococcus dysgalactiae subsp. equisimilis strain NCTC5371 | nga | Nicotine adenine dinucleotide glycohydrolase (NADGH) (EC 3.2.2.5) | 20470439 | 100 | 100 | 97 | 1e-254 |
| Streptococcus dysgalactiae subsp. equisimilis strain NCTC7136 | nga | Nicotine adenine dinucleotide glycohydrolase (NADGH) (EC 3.2.2.5) | 20470439 | 100 | 100 | 97 | 1e-254 |
| Streptococcus dysgalactiae subsp. equisimilis strain NCTC8543 | nga | Nicotine adenine dinucleotide glycohydrolase (NADGH) (EC 3.2.2.5) | 20470439 | 100 | 100 | 97 | 1e-254 |
| Streptococcus dysgalactiae subsp. equisimilis strain NCTC8546 | nga | Nicotine adenine dinucleotide glycohydrolase (NADGH) (EC 3.2.2.5) | 20470439 | 100 | 100 | 97 | 1e-254 |
| Streptococcus dysgalactiae subsp. equisimilis strain NCTC9413 | nga | Nicotine adenine dinucleotide glycohydrolase (NADGH) (EC 3.2.2.5) | 20470439 | 100 | 100 | 97 | 1e-254 |
| Streptococcus dysgalactiae subsp. equisimilis strain NCTC9414 | nga | Nicotine adenine dinucleotide glycohydrolase (NADGH) (EC 3.2.2.5) | 20470439 | 100 | 100 | 97 | 1e-254 |
| Streptococcus dysgalactiae subsp. equisimilis strain NCTC9603 | nga | Nicotine adenine dinucleotide glycohydrolase (NADGH) (EC 3.2.2.5) | 20470439 | 100 | 100 | 97 | 1e-253 |
| Streptococcus dysgalactiae subsp. equisimilis strain SS1575 | nga | Nicotine adenine dinucleotide glycohydrolase (NADGH) (EC 3.2.2.5) | 20470439 | 100 | 100 | 97 | 1e-254 |
| Streptococcus dysgalactiae subsp. equisimilis strain T642 | nga | Nicotine adenine dinucleotide glycohydrolase (NADGH) (EC 3.2.2.5) | 20470439 | 100 | 100 | 97 | 0 |
| Streptococcus dysgalactiae subsp. equisimilis strain UT_4031CC | nga | Nicotine adenine dinucleotide glycohydrolase (NADGH) (EC 3.2.2.5) | 20470439 | 100 | 100 | 97 | 1e-254 |
| Streptococcus dysgalactiae subsp. equisimilis strain UT_4231_KK | nga | Nicotine adenine dinucleotide glycohydrolase (NADGH) (EC 3.2.2.5) | 20470439 | 100 | 100 | 97 | 1e-253 |
| Streptococcus dysgalactiae subsp. equisimilis strain UT_4234_DH | nga | Nicotine adenine dinucleotide glycohydrolase (NADGH) (EC 3.2.2.5) | 20470439 | 100 | 100 | 97 | 1e-254 |
| Streptococcus dysgalactiae subsp. equisimilis strain UT_4241_XS | nga | Nicotine adenine dinucleotide glycohydrolase (NADGH) (EC 3.2.2.5) | 20470439 | 100 | 100 | 97 | 1e-253 |
| Streptococcus dysgalactiae subsp. equisimilis strain UT_4242_AB | nga | Nicotine adenine dinucleotide glycohydrolase (NADGH) (EC 3.2.2.5) | 20470439 | 100 | 100 | 97 | 1e-254 |
| Streptococcus dysgalactiae subsp. equisimilis strain UT_4255RC | nga | Nicotine adenine dinucleotide glycohydrolase (NADGH) (EC 3.2.2.5) | 20470439 | 100 | 100 | 97 | 1e-254 |
| Streptococcus dysgalactiae subsp. equisimilis strain UT_4277_BB | nga | Nicotine adenine dinucleotide glycohydrolase (NADGH) (EC 3.2.2.5) | 20470439 | 100 | 100 | 97 | 1e-254 |
| Streptococcus dysgalactiae subsp. equisimilis strain UT_4966_RC | nga | Nicotine adenine dinucleotide glycohydrolase (NADGH) (EC 3.2.2.5) | 20470439 | 100 | 100 | 97 | 1e-253 |
| Streptococcus dysgalactiae subsp. equisimilis strain UT-5345 | nga | Nicotine adenine dinucleotide glycohydrolase (NADGH) (EC 3.2.2.5) | 20470439 | 100 | 100 | 97 | 1e-254 |
| Streptococcus dysgalactiae subsp. equisimilis strain UT-5354 | nga | Nicotine adenine dinucleotide glycohydrolase (NADGH) (EC 3.2.2.5) | 20470439 | 100 | 100 | 97 | 1e-254 |
| Streptococcus dysgalactiae subsp. equisimilis strain UT-SS1069 | nga | Nicotine adenine dinucleotide glycohydrolase (NADGH) (EC 3.2.2.5) | 20470439 | 100 | 100 | 97 | 1e-253 |
| Streptococcus dysgalactiae subsp. equisimilis strain UT-SS957 | nga | Nicotine adenine dinucleotide glycohydrolase (NADGH) (EC 3.2.2.5) | 20470439 | 100 | 100 | 97 | 1e-253 |
| Streptococcus dysgalactiae subsp. equisimilis strain WCHSDSE-1 | nga | Nicotine adenine dinucleotide glycohydrolase (NADGH) (EC 3.2.2.5) | 20470439 | 100 | 100 | 97 | 1e-254 |
| Streptococcus dysgalactiae subsp. equisimilis 167 | oppA | Oligopeptide ABC transporter, substrate-binding protein OppA (TC 3.A.1.5.1) | 15845494 | 100 | 100 | 96 | 0.0 |
| Streptococcus dysgalactiae subsp. equisimilis AC-2713 | oppA | Oligopeptide ABC transporter, substrate-binding protein OppA (TC 3.A.1.5.1) | 15845494 | 100 | 100 | 96 | 0.0 |
| Streptococcus dysgalactiae subsp. equisimilis AKSDE4288 | oppA | Oligopeptide ABC transporter, substrate-binding protein OppA (TC 3.A.1.5.1) | 15845494 | 92 | 92 | 96 | 0.0 |
| Streptococcus dysgalactiae subsp. equisimilis ATCC 12394 | oppA | Oligopeptide ABC transporter, substrate-binding protein OppA (TC 3.A.1.5.1) | 15845494 | 100 | 100 | 96 | 0.0 |
| Streptococcus dysgalactiae subsp. equisimilis GGS_124 | oppA | Oligopeptide ABC transporter, substrate-binding protein OppA (TC 3.A.1.5.1) | 15845494 | 100 | 100 | 96 | 0.0 |
| Streptococcus dysgalactiae subsp. equisimilis RE378 | oppA | Oligopeptide ABC transporter, substrate-binding protein OppA (TC 3.A.1.5.1) | 15845494 | 100 | 100 | 96 | 0.0 |
| Streptococcus dysgalactiae subsp. equisimilis SD SCDR1 | oppA | Oligopeptide ABC transporter, substrate-binding protein OppA (TC 3.A.1.5.1) | 15845494 | 100 | 100 | 96 | 0.0 |
| Streptococcus dysgalactiae subsp. equisimilis SK1249 | sarA | Oligopeptide ABC transporter, substrate-binding protein OppA (TC 3.A.1.5.1) | 15845494 | 100 | 100 | 96 | 0.0 |
| Streptococcus dysgalactiae subsp. equisimilis SK1250 | sarA | Oligopeptide ABC transporter, substrate-binding protein OppA (TC 3.A.1.5.1) | 15845494 | 100 | 100 | 96 | 0.0 |
| Streptococcus dysgalactiae subsp. equisimilis strain ASDSE_96 | oppA | Oligopeptide ABC transporter, substrate-binding protein OppA (TC 3.A.1.5.1) | 15845494 | 100 | 100 | 96 | 0.0 |
| Streptococcus dysgalactiae subsp. equisimilis strain ASDSE_99 | oppA | Oligopeptide ABC transporter, substrate-binding protein OppA (TC 3.A.1.5.1) | 15845494 | 100 | 100 | 96 | 0.0 |
| Streptococcus dysgalactiae subsp. equisimilis strain C161L1 | oppA | Oligopeptide ABC transporter, substrate-binding protein OppA (TC 3.A.1.5.1) | 15845494 | 100 | 100 | 96 | 0 |
| Streptococcus dysgalactiae subsp. equisimilis strain KNZ01 | oppA | Oligopeptide ABC transporter, substrate-binding protein OppA (TC 3.A.1.5.1) | 15845494 | 100 | 100 | 96 | 0.0 |
| Streptococcus dysgalactiae subsp. equisimilis strain KNZ03 | oppA | Oligopeptide ABC transporter, substrate-binding protein OppA (TC 3.A.1.5.1) | 15845494 | 100 | 100 | 96 | 0.0 |
| Streptococcus dysgalactiae subsp. equisimilis strain KNZ04 | oppA | Oligopeptide ABC transporter, substrate-binding protein OppA (TC 3.A.1.5.1) | 15845494 | 64 | 100 | 96 | 1e-242 |
| Streptococcus dysgalactiae subsp. equisimilis strain KNZ06 | oppA | Oligopeptide ABC transporter, substrate-binding protein OppA (TC 3.A.1.5.1) | 15845494 | 100 | 100 | 96 | 0.0 |
| Streptococcus dysgalactiae subsp. equisimilis strain KNZ07 | oppA | Oligopeptide ABC transporter, substrate-binding protein OppA (TC 3.A.1.5.1) | 15845494 | 100 | 100 | 96 | 0.0 |
| Streptococcus dysgalactiae subsp. equisimilis strain KNZ10 | oppA | Oligopeptide ABC transporter, substrate-binding protein OppA (TC 3.A.1.5.1) | 15845494 | 92 | 92 | 96 | 0.0 |
| Streptococcus dysgalactiae subsp. equisimilis strain KNZ12 | oppA | Oligopeptide ABC transporter, substrate-binding protein OppA (TC 3.A.1.5.1) | 15845494 | 100 | 100 | 96 | 0.0 |
| Streptococcus dysgalactiae subsp. equisimilis strain KNZ15 | oppA | Oligopeptide ABC transporter, substrate-binding protein OppA (TC 3.A.1.5.1) | 15845494 | 92 | 92 | 96 | 0.0 |
| Streptococcus dysgalactiae subsp. equisimilis strain KNZ16 | oppA | Oligopeptide ABC transporter, substrate-binding protein OppA (TC 3.A.1.5.1) | 15845494 | 100 | 100 | 96 | 0.0 |
| Streptococcus dysgalactiae subsp. equisimilis strain NCTC10321 | oppA | Oligopeptide ABC transporter, substrate-binding protein OppA (TC 3.A.1.5.1) | 15845494 | 100 | 100 | 94 | 0.0 |
| Streptococcus dysgalactiae subsp. equisimilis strain NCTC11554 | oppA | Oligopeptide ABC transporter, substrate-binding protein OppA (TC 3.A.1.5.1) | 15845494 | 100 | 100 | 96 | 0.0 |
| Streptococcus dysgalactiae subsp. equisimilis strain NCTC11555 | oppA | Oligopeptide ABC transporter, substrate-binding protein OppA (TC 3.A.1.5.1) | 15845494 | 54 | 100 | 96 | 1e-203 |
| Streptococcus dysgalactiae subsp. equisimilis strain NCTC11556 | oppA | Oligopeptide ABC transporter, substrate-binding protein OppA (TC 3.A.1.5.1) | 15845494 | 100 | 100 | 96 | 0.0 |
| Streptococcus dysgalactiae subsp. equisimilis strain NCTC11557 | oppA | Oligopeptide ABC transporter, substrate-binding protein OppA (TC 3.A.1.5.1) | 15845494 | 100 | 100 | 96 | 0.0 |
| Streptococcus dysgalactiae subsp. equisimilis strain NCTC11564 | oppA | Oligopeptide ABC transporter, substrate-binding protein OppA (TC 3.A.1.5.1) | 15845494 | 92 | 92 | 96 | 0.0 |
| Streptococcus dysgalactiae subsp. equisimilis strain NCTC5370 | oppA | Oligopeptide ABC transporter, substrate-binding protein OppA (TC 3.A.1.5.1) | 15845494 | 100 | 100 | 96 | 0.0 |
| Streptococcus dysgalactiae subsp. equisimilis strain NCTC5371 | oppA | Oligopeptide ABC transporter, substrate-binding protein OppA (TC 3.A.1.5.1) | 15845494 | 92 | 92 | 96 | 0.0 |
| Streptococcus dysgalactiae subsp. equisimilis strain NCTC6179 | oppA | Oligopeptide ABC transporter, substrate-binding protein OppA (TC 3.A.1.5.1) | 15845494 | 100 | 100 | 94 | 0.0 |
| Streptococcus dysgalactiae subsp. equisimilis strain NCTC6181 | oppA | Oligopeptide ABC transporter, substrate-binding protein OppA (TC 3.A.1.5.1) | 15845494 | 100 | 100 | 94 | 0.0 |
| Streptococcus dysgalactiae subsp. equisimilis strain NCTC6407 | oppA | Oligopeptide ABC transporter, substrate-binding protein OppA (TC 3.A.1.5.1) | 15845494 | 100 | 100 | 94 | 0.0 |
| Streptococcus dysgalactiae subsp. equisimilis strain NCTC7136 | oppA | Oligopeptide ABC transporter, substrate-binding protein OppA (TC 3.A.1.5.1) | 15845494 | 100 | 100 | 96 | 0.0 |
| Streptococcus dysgalactiae subsp. equisimilis strain NCTC8543 | oppA | Oligopeptide ABC transporter, substrate-binding protein OppA (TC 3.A.1.5.1) | 15845494 | 100 | 100 | 96 | 0.0 |
| Streptococcus dysgalactiae subsp. equisimilis strain NCTC8546 | oppA | Oligopeptide ABC transporter, substrate-binding protein OppA (TC 3.A.1.5.1) | 15845494 | 100 | 100 | 96 | 0.0 |
| Streptococcus dysgalactiae subsp. equisimilis strain NCTC9413 | oppA | Oligopeptide ABC transporter, substrate-binding protein OppA (TC 3.A.1.5.1) | 15845494 | 100 | 100 | 96 | 0.0 |
| Streptococcus dysgalactiae subsp. equisimilis strain NCTC9414 | oppA | Oligopeptide ABC transporter, substrate-binding protein OppA (TC 3.A.1.5.1) | 15845494 | 92 | 92 | 96 | 0.0 |
| Streptococcus dysgalactiae subsp. equisimilis strain NCTC9603 | oppA | Oligopeptide ABC transporter, substrate-binding protein OppA (TC 3.A.1.5.1) | 15845494 | 100 | 100 | 96 | 0.0 |
| Streptococcus dysgalactiae subsp. equisimilis strain SS1575 | oppA | Oligopeptide ABC transporter, substrate-binding protein OppA (TC 3.A.1.5.1) | 15845494 | 100 | 100 | 96 | 0.0 |
| Streptococcus dysgalactiae subsp. equisimilis strain T642 | oppA | Oligopeptide ABC transporter, substrate-binding protein OppA (TC 3.A.1.5.1) | 15845494 | 100 | 100 | 96 | 0 |
| Streptococcus dysgalactiae subsp. equisimilis strain UT_4031CC | oppA | Oligopeptide ABC transporter, substrate-binding protein OppA (TC 3.A.1.5.1) | 15845494 | 92 | 92 | 96 | 0.0 |
| Streptococcus dysgalactiae subsp. equisimilis strain UT_4231_KK | oppA | Oligopeptide ABC transporter, substrate-binding protein OppA (TC 3.A.1.5.1) | 15845494 | 100 | 100 | 96 | 0.0 |
| Streptococcus dysgalactiae subsp. equisimilis strain UT_4234_DH | oppA | Oligopeptide ABC transporter, substrate-binding protein OppA (TC 3.A.1.5.1) | 15845494 | 100 | 100 | 96 | 0.0 |
| Streptococcus dysgalactiae subsp. equisimilis strain UT_4241_XS | oppA | Oligopeptide ABC transporter, substrate-binding protein OppA (TC 3.A.1.5.1) | 15845494 | 100 | 100 | 96 | 0.0 |
| Streptococcus dysgalactiae subsp. equisimilis strain UT_4242_AB | oppA | Oligopeptide ABC transporter, substrate-binding protein OppA (TC 3.A.1.5.1) | 15845494 | 100 | 100 | 96 | 0.0 |
| Streptococcus dysgalactiae subsp. equisimilis strain UT_4255RC | oppA | Oligopeptide ABC transporter, substrate-binding protein OppA (TC 3.A.1.5.1) | 15845494 | 100 | 100 | 96 | 0.0 |
| Streptococcus dysgalactiae subsp. equisimilis strain UT_4277_BB | oppA | Oligopeptide ABC transporter, substrate-binding protein OppA (TC 3.A.1.5.1) | 15845494 | 92 | 92 | 96 | 0.0 |
| Streptococcus dysgalactiae subsp. equisimilis strain UT_4966_RC | oppA | Oligopeptide ABC transporter, substrate-binding protein OppA (TC 3.A.1.5.1) | 15845494 | 100 | 100 | 96 | 0.0 |
| Streptococcus dysgalactiae subsp. equisimilis strain UT-5345 | oppA | Oligopeptide ABC transporter, substrate-binding protein OppA (TC 3.A.1.5.1) | 15845494 | 100 | 100 | 96 | 0.0 |
| Streptococcus dysgalactiae subsp. equisimilis strain UT-5354 | oppA | Oligopeptide ABC transporter, substrate-binding protein OppA (TC 3.A.1.5.1) | 15845494 | 100 | 100 | 96 | 0.0 |
| Streptococcus dysgalactiae subsp. equisimilis strain UT-SS1069 | oppA | Oligopeptide ABC transporter, substrate-binding protein OppA (TC 3.A.1.5.1) | 15845494 | 100 | 100 | 96 | 0.0 |
| Streptococcus dysgalactiae subsp. equisimilis strain UT-SS957 | oppA | Oligopeptide ABC transporter, substrate-binding protein OppA (TC 3.A.1.5.1) | 15845494 | 77 | 100 | 97 | 1e-296 |
| Streptococcus dysgalactiae subsp. equisimilis strain WCHSDSE-1 | oppA | Oligopeptide ABC transporter, substrate-binding protein OppA (TC 3.A.1.5.1) | 15845494 | 100 | 100 | 96 | 0.0 |
| Streptococcus dysgalactiae subsp. equisimilis 167 | perR | Peroxide stress regulator PerR, FUR family | 12183543 | 100 | 100 | 94 | 5e-83 |
| Streptococcus dysgalactiae subsp. equisimilis AC-2713 | perR | Peroxide stress regulator PerR, FUR family | 12183543 | 100 | 100 | 94 | 7e-84 |
| Streptococcus dysgalactiae subsp. equisimilis AKSDE4288 | perR | Peroxide stress regulator PerR, FUR family | 12183543 | 100 | 100 | 94 | 7e-84 |
| Streptococcus dysgalactiae subsp. equisimilis ATCC 12394 | perR | Peroxide stress regulator PerR, FUR family | 12183543 | 100 | 100 | 94 | 6e-83 |
| Streptococcus dysgalactiae subsp. equisimilis GGS_124 | perR | Peroxide stress regulator PerR, FUR family | 12183543 | 100 | 100 | 94 | 7e-84 |
| Streptococcus dysgalactiae subsp. equisimilis RE378 | perR | Peroxide stress regulator PerR, FUR family | 12183543 | 100 | 100 | 94 | 7e-84 |
| Streptococcus dysgalactiae subsp. equisimilis SD SCDR1 | perR | Peroxide stress regulator PerR, FUR family | 12183543 | 100 | 100 | 93 | 5e-83 |
| Streptococcus dysgalactiae subsp. equisimilis SK1249 | perR | Peroxide stress regulator PerR, FUR family | 12183543 | 52 | 98 | 91 | 4e-39 |
| Streptococcus dysgalactiae subsp. equisimilis SK1250 | perR | Peroxide stress regulator PerR, FUR family | 12183543 | 100 | 100 | 93 | 5e-83 |
| Streptococcus dysgalactiae subsp. equisimilis strain ASDSE_96 | perR | Peroxide stress regulator PerR, FUR family | 12183543 | 100 | 100 | 94 | 7e-84 |
| Streptococcus dysgalactiae subsp. equisimilis strain ASDSE_99 | perR | Peroxide stress regulator PerR, FUR family | 12183543 | 100 | 100 | 94 | 7e-84 |
| Streptococcus dysgalactiae subsp. equisimilis strain C161L1 | perR | Peroxide stress regulator PerR, FUR family | 12183543 | 100 | 100 | 94 | 2e-110 |
| Streptococcus dysgalactiae subsp. equisimilis strain KNZ01 | perR | Peroxide stress regulator PerR, FUR family | 12183543 | 100 | 100 | 94 | 7e-84 |
| Streptococcus dysgalactiae subsp. equisimilis strain KNZ03 | perR | Peroxide stress regulator PerR, FUR family | 12183543 | 100 | 100 | 94 | 6e-83 |
| Streptococcus dysgalactiae subsp. equisimilis strain KNZ04 | perR | Peroxide stress regulator PerR, FUR family | 12183543 | 100 | 100 | 94 | 6e-83 |
| Streptococcus dysgalactiae subsp. equisimilis strain KNZ06 | perR | Peroxide stress regulator PerR, FUR family | 12183543 | 100 | 100 | 94 | 7e-84 |
| Streptococcus dysgalactiae subsp. equisimilis strain KNZ07 | perR | Peroxide stress regulator PerR, FUR family | 12183543 | 100 | 100 | 94 | 7e-84 |
| Streptococcus dysgalactiae subsp. equisimilis strain KNZ10 | perR | Peroxide stress regulator PerR, FUR family | 12183543 | 100 | 100 | 94 | 7e-84 |
| Streptococcus dysgalactiae subsp. equisimilis strain KNZ12 | perR | Peroxide stress regulator PerR, FUR family | 12183543 | 100 | 100 | 94 | 7e-84 |
| Streptococcus dysgalactiae subsp. equisimilis strain KNZ15 | perR | Peroxide stress regulator PerR, FUR family | 12183543 | 100 | 100 | 94 | 7e-84 |
| Streptococcus dysgalactiae subsp. equisimilis strain KNZ16 | perR | Peroxide stress regulator PerR, FUR family | 12183543 | 100 | 100 | 94 | 6e-83 |
| Streptococcus dysgalactiae subsp. equisimilis strain NCTC10321 | perR | Peroxide stress regulator PerR, FUR family | 12183543 | 100 | 100 | 94 | 2e-83 |
| Streptococcus dysgalactiae subsp. equisimilis strain NCTC11554 | perR | Peroxide stress regulator PerR, FUR family | 12183543 | 100 | 100 | 94 | 6e-83 |
| Streptococcus dysgalactiae subsp. equisimilis strain NCTC11555 | perR | Peroxide stress regulator PerR, FUR family | 12183543 | 100 | 100 | 94 | 2e-83 |
| Streptococcus dysgalactiae subsp. equisimilis strain NCTC11556 | perR | Peroxide stress regulator PerR, FUR family | 12183543 | 100 | 100 | 93 | 5e-83 |
| Streptococcus dysgalactiae subsp. equisimilis strain NCTC11557 | perR | Peroxide stress regulator PerR, FUR family | 12183543 | 100 | 100 | 94 | 7e-84 |
| Streptococcus dysgalactiae subsp. equisimilis strain NCTC11564 | perR | Peroxide stress regulator PerR, FUR family | 12183543 | 100 | 100 | 94 | 7e-84 |
| Streptococcus dysgalactiae subsp. equisimilis strain NCTC5370 | perR | Peroxide stress regulator PerR, FUR family | 12183543 | 100 | 100 | 94 | 5e-83 |
| Streptococcus dysgalactiae subsp. equisimilis strain NCTC5371 | perR | Peroxide stress regulator PerR, FUR family | 12183543 | 100 | 100 | 94 | 7e-84 |
| Streptococcus dysgalactiae subsp. equisimilis strain NCTC6179 | perR | Peroxide stress regulator PerR, FUR family | 12183543 | 100 | 98 | 94 | 7e-84 |
| Streptococcus dysgalactiae subsp. equisimilis strain NCTC6181 | perR | Peroxide stress regulator PerR, FUR family | 12183543 | 100 | 98 | 94 | 7e-84 |
| Streptococcus dysgalactiae subsp. equisimilis strain NCTC6407 | perR | Peroxide stress regulator PerR, FUR family | 12183543 | 100 | 100 | 94 | 2e-83 |
| Streptococcus dysgalactiae subsp. equisimilis strain NCTC7136 | perR | Peroxide stress regulator PerR, FUR family | 12183543 | 100 | 100 | 94 | 7e-84 |
| Streptococcus dysgalactiae subsp. equisimilis strain NCTC8543 | perR | Peroxide stress regulator PerR, FUR family | 12183543 | 100 | 100 | 93 | 1e-82 |
| Streptococcus dysgalactiae subsp. equisimilis strain NCTC8546 | perR | Peroxide stress regulator PerR, FUR family | 12183543 | 100 | 100 | 94 | 7e-84 |
| Streptococcus dysgalactiae subsp. equisimilis strain NCTC9413 | perR | Peroxide stress regulator PerR, FUR family | 12183543 | 100 | 100 | 94 | 7e-84 |
| Streptococcus dysgalactiae subsp. equisimilis strain NCTC9414 | perR | Peroxide stress regulator PerR, FUR family | 12183543 | 100 | 100 | 94 | 7e-84 |
| Streptococcus dysgalactiae subsp. equisimilis strain NCTC9603 | perR | Peroxide stress regulator PerR, FUR family | 12183543 | 100 | 100 | 94 | 7e-84 |
| Streptococcus dysgalactiae subsp. equisimilis strain SS1575 | perR | Peroxide stress regulator PerR, FUR family | 12183543 | 100 | 100 | 94 | 7e-84 |
| Streptococcus dysgalactiae subsp. equisimilis strain T642 | perR | Peroxide stress regulator PerR, FUR family | 12183543 | 100 | 100 | 94 | 0 |
| Streptococcus dysgalactiae subsp. equisimilis strain UT_4031CC | perR | Peroxide stress regulator PerR, FUR family | 12183543 | 100 | 100 | 94 | 7e-84 |
| Streptococcus dysgalactiae subsp. equisimilis strain UT_4231_KK | perR | Peroxide stress regulator PerR, FUR family | 12183543 | 100 | 100 | 94 | 5e-83 |
| Streptococcus dysgalactiae subsp. equisimilis strain UT_4234_DH | perR | Peroxide stress regulator PerR, FUR family | 12183543 | 100 | 100 | 94 | 7e-84 |
| Streptococcus dysgalactiae subsp. equisimilis strain UT_4241_XS | perR | Peroxide stress regulator PerR, FUR family | 12183543 | 100 | 100 | 94 | 5e-83 |
| Streptococcus dysgalactiae subsp. equisimilis strain UT_4242_AB | perR | Peroxide stress regulator PerR, FUR family | 12183543 | 100 | 100 | 93 | 5e-83 |
| Streptococcus dysgalactiae subsp. equisimilis strain UT_4255RC | perR | Peroxide stress regulator PerR, FUR family | 12183543 | 100 | 100 | 94 | 7e-84 |
| Streptococcus dysgalactiae subsp. equisimilis strain UT_4277_BB | perR | Peroxide stress regulator PerR, FUR family | 12183543 | 100 | 100 | 94 | 7e-84 |
| Streptococcus dysgalactiae subsp. equisimilis strain UT_4966_RC | perR | Peroxide stress regulator PerR, FUR family | 12183543 | 100 | 100 | 94 | 5e-83 |
| Streptococcus dysgalactiae subsp. equisimilis strain UT-5345 | perR | Peroxide stress regulator PerR, FUR family | 12183543 | 100 | 100 | 94 | 1e-83 |
| Streptococcus dysgalactiae subsp. equisimilis strain UT-5354 | perR | Peroxide stress regulator PerR, FUR family | 12183543 | 100 | 100 | 94 | 2e-83 |
| Streptococcus dysgalactiae subsp. equisimilis strain UT-SS1069 | perR | Peroxide stress regulator PerR, FUR family | 12183543 | 100 | 100 | 94 | 5e-83 |
| Streptococcus dysgalactiae subsp. equisimilis strain UT-SS957 | perR | Peroxide stress regulator PerR, FUR family | 12183543 | 100 | 100 | 93 | 5e-83 |
| Streptococcus dysgalactiae subsp. equisimilis strain WCHSDSE-1 | perR | Peroxide stress regulator PerR, FUR family | 12183543 | 100 | 100 | 93 | 5e-83 |
| Streptococcus dysgalactiae subsp. equisimilis 167 | pstB | Phosphate ABC transporter, ATP-binding protein PstB (TC 3.A.1.7.1) | 12207705 | 100 | 100 | 81 | 1e-119 |
| Streptococcus dysgalactiae subsp. equisimilis AC-2713 | pstB | Phosphate ABC transporter, ATP-binding protein PstB (TC 3.A.1.7.1) | 12207705 | 100 | 100 | 81 | 1e-119 |
| Streptococcus dysgalactiae subsp. equisimilis AKSDE4288 | SP_1396 | Phosphate ABC transporter, ATP-binding protein PstB (TC 3.A.1.7.1) | 12207705 | 100 | 100 | 81 | 1e-119 |
| Streptococcus dysgalactiae subsp. equisimilis ATCC 12394 | SP_1396 | Phosphate ABC transporter, ATP-binding protein PstB (TC 3.A.1.7.1) | 12207705 | 100 | 100 | 81 | 1e-119 |
| Streptococcus dysgalactiae subsp. equisimilis GGS_124 | pstB | Phosphate ABC transporter, ATP-binding protein PstB (TC 3.A.1.7.1) | 12207705 | 100 | 100 | 81 | 1e-119 |
| Streptococcus dysgalactiae subsp. equisimilis RE378 | pstB | Phosphate ABC transporter, ATP-binding protein PstB (TC 3.A.1.7.1) | 12207705 | 100 | 100 | 81 | 1e-119 |
| Streptococcus dysgalactiae subsp. equisimilis SD SCDR1 | SP_1396 | Phosphate ABC transporter, ATP-binding protein PstB (TC 3.A.1.7.1) | 12207705 | 100 | 100 | 81 | 1e-119 |
| Streptococcus dysgalactiae subsp. equisimilis SK1249 | pstB | Phosphate ABC transporter, ATP-binding protein PstB (TC 3.A.1.7.1) | 12207705 | 100 | 100 | 81 | 1e-119 |
| Streptococcus dysgalactiae subsp. equisimilis SK1250 | pstB_2 | Phosphate ABC transporter, ATP-binding protein PstB (TC 3.A.1.7.1) | 12207705 | 100 | 100 | 81 | 1e-119 |
| Streptococcus dysgalactiae subsp. equisimilis strain ASDSE_96 | SP_1396 | Phosphate ABC transporter, ATP-binding protein PstB (TC 3.A.1.7.1) | 12207705 | 100 | 100 | 81 | 1e-119 |
| Streptococcus dysgalactiae subsp. equisimilis strain ASDSE_99 | SP_1396 | Phosphate ABC transporter, ATP-binding protein PstB (TC 3.A.1.7.1) | 12207705 | 100 | 100 | 81 | 1e-119 |
| Streptococcus dysgalactiae subsp. equisimilis strain C161L1 | SP_1396 | Phosphate ABC transporter, ATP-binding protein PstB (TC 3.A.1.7.1) | 12207705 | 100 | 100 | 81 | 0 |
| Streptococcus dysgalactiae subsp. equisimilis strain KNZ01 | SP_1396 | Phosphate ABC transporter, ATP-binding protein PstB (TC 3.A.1.7.1) | 12207705 | 100 | 100 | 81 | 1e-119 |
| Streptococcus dysgalactiae subsp. equisimilis strain KNZ03 | SP_1396 | Phosphate ABC transporter, ATP-binding protein PstB (TC 3.A.1.7.1) | 12207705 | 100 | 100 | 81 | 1e-119 |
| Streptococcus dysgalactiae subsp. equisimilis strain KNZ04 | SP_1396 | Phosphate ABC transporter, ATP-binding protein PstB (TC 3.A.1.7.1) | 12207705 | 100 | 100 | 81 | 1e-119 |
| Streptococcus dysgalactiae subsp. equisimilis strain KNZ06 | SP_1396 | Phosphate ABC transporter, ATP-binding protein PstB (TC 3.A.1.7.1) | 12207705 | 100 | 100 | 81 | 1e-119 |
| Streptococcus dysgalactiae subsp. equisimilis strain KNZ07 | SP_1396 | Phosphate ABC transporter, ATP-binding protein PstB (TC 3.A.1.7.1) | 12207705 | 100 | 100 | 81 | 1e-119 |
| Streptococcus dysgalactiae subsp. equisimilis strain KNZ10 | SP_1396 | Phosphate ABC transporter, ATP-binding protein PstB (TC 3.A.1.7.1) | 12207705 | 100 | 100 | 81 | 1e-119 |
| Streptococcus dysgalactiae subsp. equisimilis strain KNZ12 | SP_1396 | Phosphate ABC transporter, ATP-binding protein PstB (TC 3.A.1.7.1) | 12207705 | 100 | 100 | 81 | 1e-119 |
| Streptococcus dysgalactiae subsp. equisimilis strain KNZ15 | SP_1396 | Phosphate ABC transporter, ATP-binding protein PstB (TC 3.A.1.7.1) | 12207705 | 100 | 100 | 81 | 1e-119 |
| Streptococcus dysgalactiae subsp. equisimilis strain KNZ16 | SP_1396 | Phosphate ABC transporter, ATP-binding protein PstB (TC 3.A.1.7.1) | 12207705 | 100 | 100 | 81 | 1e-119 |
| Streptococcus dysgalactiae subsp. equisimilis strain NCTC10321 | SP_1396 | Phosphate ABC transporter, ATP-binding protein PstB (TC 3.A.1.7.1) | 12207705 | 100 | 100 | 81 | 1e-118 |
| Streptococcus dysgalactiae subsp. equisimilis strain NCTC11554 | SP_1396 | Phosphate ABC transporter, ATP-binding protein PstB (TC 3.A.1.7.1) | 12207705 | 100 | 100 | 81 | 1e-119 |
| Streptococcus dysgalactiae subsp. equisimilis strain NCTC11555 | SP_1396 | Phosphate ABC transporter, ATP-binding protein PstB (TC 3.A.1.7.1) | 12207705 | 100 | 100 | 81 | 1e-119 |
| Streptococcus dysgalactiae subsp. equisimilis strain NCTC11556 | SP_1396 | Phosphate ABC transporter, ATP-binding protein PstB (TC 3.A.1.7.1) | 12207705 | 100 | 100 | 80 | 1e-118 |
| Streptococcus dysgalactiae subsp. equisimilis strain NCTC11557 | SP_1396 | Phosphate ABC transporter, ATP-binding protein PstB (TC 3.A.1.7.1) | 12207705 | 100 | 100 | 81 | 1e-119 |
| Streptococcus dysgalactiae subsp. equisimilis strain NCTC11564 | SP_1396 | Phosphate ABC transporter, ATP-binding protein PstB (TC 3.A.1.7.1) | 12207705 | 100 | 100 | 81 | 1e-119 |
| Streptococcus dysgalactiae subsp. equisimilis strain NCTC5370 | SP_1396 | Phosphate ABC transporter, ATP-binding protein PstB (TC 3.A.1.7.1) | 12207705 | 100 | 100 | 81 | 1e-119 |
| Streptococcus dysgalactiae subsp. equisimilis strain NCTC5371 | SP_1396 | Phosphate ABC transporter, ATP-binding protein PstB (TC 3.A.1.7.1) | 12207705 | 100 | 100 | 81 | 1e-119 |
| Streptococcus dysgalactiae subsp. equisimilis strain NCTC6179 | SP_1396 | Phosphate ABC transporter, ATP-binding protein PstB (TC 3.A.1.7.1) | 12207705 | 100 | 100 | 81 | 1e-119 |
| Streptococcus dysgalactiae subsp. equisimilis strain NCTC6181 | SP_1396 | Phosphate ABC transporter, ATP-binding protein PstB (TC 3.A.1.7.1) | 12207705 | 100 | 100 | 81 | 1e-119 |
| Streptococcus dysgalactiae subsp. equisimilis strain NCTC6407 | SP_1396 | Phosphate ABC transporter, ATP-binding protein PstB (TC 3.A.1.7.1) | 12207705 | 100 | 100 | 81 | 1e-118 |
| Streptococcus dysgalactiae subsp. equisimilis strain NCTC7136 | SP_1396 | Phosphate ABC transporter, ATP-binding protein PstB (TC 3.A.1.7.1) | 12207705 | 100 | 100 | 81 | 1e-119 |
| Streptococcus dysgalactiae subsp. equisimilis strain NCTC8543 | SP_1396 | Phosphate ABC transporter, ATP-binding protein PstB (TC 3.A.1.7.1) | 12207705 | 100 | 100 | 80 | 1e-118 |
| Streptococcus dysgalactiae subsp. equisimilis strain NCTC8546 | SP_1396 | Phosphate ABC transporter, ATP-binding protein PstB (TC 3.A.1.7.1) | 12207705 | 100 | 100 | 81 | 1e-119 |
| Streptococcus dysgalactiae subsp. equisimilis strain NCTC9413 | SP_1396 | Phosphate ABC transporter, ATP-binding protein PstB (TC 3.A.1.7.1) | 12207705 | 100 | 100 | 81 | 1e-119 |
| Streptococcus dysgalactiae subsp. equisimilis strain NCTC9414 | SP_1396 | Phosphate ABC transporter, ATP-binding protein PstB (TC 3.A.1.7.1) | 12207705 | 100 | 100 | 81 | 1e-119 |
| Streptococcus dysgalactiae subsp. equisimilis strain NCTC9603 | SP_1396 | Phosphate ABC transporter, ATP-binding protein PstB (TC 3.A.1.7.1) | 12207705 | 100 | 100 | 81 | 1e-119 |
| Streptococcus dysgalactiae subsp. equisimilis strain SS1575 | SP_1396 | Phosphate ABC transporter, ATP-binding protein PstB (TC 3.A.1.7.1) | 12207705 | 100 | 100 | 81 | 1e-119 |
| Streptococcus dysgalactiae subsp. equisimilis strain T642 | SP_1396 | Phosphate ABC transporter, ATP-binding protein PstB (TC 3.A.1.7.1) | 12207705 | 100 | 100 | 81 | 0 |
| Streptococcus dysgalactiae subsp. equisimilis strain UT_4031CC | SP_1396 | Phosphate ABC transporter, ATP-binding protein PstB (TC 3.A.1.7.1) | 12207705 | 100 | 100 | 81 | 1e-119 |
| Streptococcus dysgalactiae subsp. equisimilis strain UT_4231_KK | SP_1396 | Phosphate ABC transporter, ATP-binding protein PstB (TC 3.A.1.7.1) | 12207705 | 100 | 100 | 81 | 1e-119 |
| Streptococcus dysgalactiae subsp. equisimilis strain UT_4234_DH | SP_1396 | Phosphate ABC transporter, ATP-binding protein PstB (TC 3.A.1.7.1) | 12207705 | 100 | 100 | 81 | 1e-119 |
| Streptococcus dysgalactiae subsp. equisimilis strain UT_4241_XS | SP_1396 | Phosphate ABC transporter, ATP-binding protein PstB (TC 3.A.1.7.1) | 12207705 | 100 | 100 | 81 | 1e-119 |
| Streptococcus dysgalactiae subsp. equisimilis strain UT_4242_AB | SP_1396 | Phosphate ABC transporter, ATP-binding protein PstB (TC 3.A.1.7.1) | 12207705 | 100 | 100 | 80 | 1e-118 |
| Streptococcus dysgalactiae subsp. equisimilis strain UT_4255RC | SP_1396 | Phosphate ABC transporter, ATP-binding protein PstB (TC 3.A.1.7.1) | 12207705 | 100 | 100 | 81 | 1e-119 |
| Streptococcus dysgalactiae subsp. equisimilis strain UT_4277_BB | SP_1396 | Phosphate ABC transporter, ATP-binding protein PstB (TC 3.A.1.7.1) | 12207705 | 100 | 100 | 81 | 1e-119 |
| Streptococcus dysgalactiae subsp. equisimilis strain UT_4966_RC | SP_1396 | Phosphate ABC transporter, ATP-binding protein PstB (TC 3.A.1.7.1) | 12207705 | 100 | 100 | 81 | 1e-119 |
| Streptococcus dysgalactiae subsp. equisimilis strain UT-5345 | SP_1396 | Phosphate ABC transporter, ATP-binding protein PstB (TC 3.A.1.7.1) | 12207705 | 100 | 100 | 81 | 1e-119 |
| Streptococcus dysgalactiae subsp. equisimilis strain UT-5354 | SP_1396 | Phosphate ABC transporter, ATP-binding protein PstB (TC 3.A.1.7.1) | 12207705 | 100 | 100 | 81 | 1e-119 |
| Streptococcus dysgalactiae subsp. equisimilis strain UT-SS1069 | SP_1396 | Phosphate ABC transporter, ATP-binding protein PstB (TC 3.A.1.7.1) | 12207705 | 100 | 100 | 81 | 1e-119 |
| Streptococcus dysgalactiae subsp. equisimilis strain UT-SS957 | SP_1396 | Phosphate ABC transporter, ATP-binding protein PstB (TC 3.A.1.7.1) | 12207705 | 100 | 100 | 81 | 1e-119 |
| Streptococcus dysgalactiae subsp. equisimilis strain WCHSDSE-1 | SP_1396 | Phosphate ABC transporter, ATP-binding protein PstB (TC 3.A.1.7.1) | 12207705 | 100 | 100 | 81 | 1e-119 |
| Streptococcus dysgalactiae subsp. equisimilis 167 | pstA | Phosphate ABC transporter, permease protein PstA (TC 3.A.1.7.1) | 12207705 | 100 | 100 | 93 | 1e-136 |
| Streptococcus dysgalactiae subsp. equisimilis 167 | pstA | Phosphate ABC transporter, permease protein PstA (TC 3.A.1.7.1) | 12207705 | 100 | 99 | 88 | 1e-148 |
| Streptococcus dysgalactiae subsp. equisimilis AC-2713 | pstA | Phosphate ABC transporter, permease protein PstA (TC 3.A.1.7.1) | 12207705 | 100 | 99 | 88 | 1e-147 |
| Streptococcus dysgalactiae subsp. equisimilis AC-2713 | SP_1398 | Phosphate ABC transporter, permease protein PstA (TC 3.A.1.7.1) | 12207705 | 100 | 100 | 92 | 1e-135 |
| Streptococcus dysgalactiae subsp. equisimilis AKSDE4288 | SP_1398 | Phosphate ABC transporter, permease protein PstA (TC 3.A.1.7.1) | 12207705 | 100 | 99 | 88 | 1e-148 |
| Streptococcus dysgalactiae subsp. equisimilis AKSDE4288 | SP_1398 | Phosphate ABC transporter, permease protein PstA (TC 3.A.1.7.1) | 12207705 | 100 | 100 | 92 | 1e-135 |
| Streptococcus dysgalactiae subsp. equisimilis ATCC 12394 | SP_1398 | Phosphate ABC transporter, permease protein PstA (TC 3.A.1.7.1) | 12207705 | 100 | 100 | 93 | 1e-136 |
| Streptococcus dysgalactiae subsp. equisimilis ATCC 12394 | SP_1398 | Phosphate ABC transporter, permease protein PstA (TC 3.A.1.7.1) | 12207705 | 100 | 99 | 88 | 1e-148 |
| Streptococcus dysgalactiae subsp. equisimilis GGS_124 | pstA | Phosphate ABC transporter, permease protein PstA (TC 3.A.1.7.1) | 12207705 | 100 | 99 | 88 | 1e-148 |
| Streptococcus dysgalactiae subsp. equisimilis GGS_124 | pstA | Phosphate ABC transporter, permease protein PstA (TC 3.A.1.7.1) | 12207705 | 100 | 100 | 93 | 1e-136 |
| Streptococcus dysgalactiae subsp. equisimilis RE378 | pstA | Phosphate ABC transporter, permease protein PstA (TC 3.A.1.7.1) | 12207705 | 100 | 99 | 88 | 1e-148 |
| Streptococcus dysgalactiae subsp. equisimilis RE378 | pstA | Phosphate ABC transporter, permease protein PstA (TC 3.A.1.7.1) | 12207705 | 100 | 100 | 93 | 1e-136 |
| Streptococcus dysgalactiae subsp. equisimilis SD SCDR1 | SP_1398 | Phosphate ABC transporter, permease protein PstA (TC 3.A.1.7.1) | 12207705 | 100 | 99 | 88 | 1e-148 |
| Streptococcus dysgalactiae subsp. equisimilis SD SCDR1 | SP_2086 | Phosphate ABC transporter, permease protein PstA (TC 3.A.1.7.1) | 12207705 | 100 | 100 | 93 | 1e-136 |
| Streptococcus dysgalactiae subsp. equisimilis SK1249 | pstA | Phosphate ABC transporter, permease protein PstA (TC 3.A.1.7.1) | 12207705 | 100 | 99 | 88 | 1e-148 |
| Streptococcus dysgalactiae subsp. equisimilis SK1249 | pstA | Phosphate ABC transporter, permease protein PstA (TC 3.A.1.7.1) | 12207705 | 100 | 100 | 93 | 1e-135 |
| Streptococcus dysgalactiae subsp. equisimilis SK1250 | pstA_2 | Phosphate ABC transporter, permease protein PstA (TC 3.A.1.7.1) | 12207705 | 100 | 99 | 88 | 1e-148 |
| Streptococcus dysgalactiae subsp. equisimilis SK1250 | pstA_1 | Phosphate ABC transporter, permease protein PstA (TC 3.A.1.7.1) | 12207705 | 100 | 100 | 93 | 1e-136 |
| Streptococcus dysgalactiae subsp. equisimilis strain ASDSE_96 | pstA_1 | Phosphate ABC transporter, permease protein PstA (TC 3.A.1.7.1) | 12207705 | 100 | 99 | 88 | 1e-147 |
| Streptococcus dysgalactiae subsp. equisimilis strain ASDSE_96 | pstA_1 | Phosphate ABC transporter, permease protein PstA (TC 3.A.1.7.1) | 12207705 | 100 | 100 | 93 | 1e-136 |
| Streptococcus dysgalactiae subsp. equisimilis strain ASDSE_99 | pstA_1 | Phosphate ABC transporter, permease protein PstA (TC 3.A.1.7.1) | 12207705 | 100 | 100 | 92 | 1e-135 |
| Streptococcus dysgalactiae subsp. equisimilis strain ASDSE_99 | pstA_1 | Phosphate ABC transporter, permease protein PstA (TC 3.A.1.7.1) | 12207705 | 100 | 99 | 88 | 1e-147 |
| Streptococcus dysgalactiae subsp. equisimilis strain C161L1 | pstA_1 | Phosphate ABC transporter, permease protein PstA (TC 3.A.1.7.1) | 12207705 | 100 | 99 | 88 | 0 |
| Streptococcus dysgalactiae subsp. equisimilis strain C161L1 | pstA_1 | Phosphate ABC transporter, permease protein PstA (TC 3.A.1.7.1) | 12207705 | 100 | 100 | 93 | 0 |
| Streptococcus dysgalactiae subsp. equisimilis strain KNZ01 | SP_2086 | Phosphate ABC transporter, permease protein PstA (TC 3.A.1.7.1) | 12207705 | 100 | 100 | 93 | 1e-136 |
| Streptococcus dysgalactiae subsp. equisimilis strain KNZ01 | SP_1398 | Phosphate ABC transporter, permease protein PstA (TC 3.A.1.7.1) | 12207705 | 100 | 99 | 88 | 1e-148 |
| Streptococcus dysgalactiae subsp. equisimilis strain KNZ01 | SP_1398 | Phosphate ABC transporter, permease protein PstA (TC 3.A.1.7.1) | 12207705 | 100 | 99 | 88 | 1e-148 |
| Streptococcus dysgalactiae subsp. equisimilis strain KNZ01 | SP_2086 | Phosphate ABC transporter, permease protein PstA (TC 3.A.1.7.1) | 12207705 | 100 | 100 | 93 | 1e-136 |
| Streptococcus dysgalactiae subsp. equisimilis strain KNZ03 | SP_1398 | Phosphate ABC transporter, permease protein PstA (TC 3.A.1.7.1) | 12207705 | 100 | 99 | 88 | 1e-148 |
| Streptococcus dysgalactiae subsp. equisimilis strain KNZ03 | SP_1398 | Phosphate ABC transporter, permease protein PstA (TC 3.A.1.7.1) | 12207705 | 100 | 99 | 88 | 1e-148 |
| Streptococcus dysgalactiae subsp. equisimilis strain KNZ03 | SP_2086 | Phosphate ABC transporter, permease protein PstA (TC 3.A.1.7.1) | 12207705 | 100 | 100 | 93 | 1e-135 |
| Streptococcus dysgalactiae subsp. equisimilis strain KNZ03 | SP_2086 | Phosphate ABC transporter, permease protein PstA (TC 3.A.1.7.1) | 12207705 | 100 | 100 | 93 | 1e-135 |
| Streptococcus dysgalactiae subsp. equisimilis strain KNZ04 | SP_2086 | Phosphate ABC transporter, permease protein PstA (TC 3.A.1.7.1) | 12207705 | 100 | 100 | 93 | 1e-135 |
| Streptococcus dysgalactiae subsp. equisimilis strain KNZ04 | SP_1398 | Phosphate ABC transporter, permease protein PstA (TC 3.A.1.7.1) | 12207705 | 100 | 99 | 88 | 1e-148 |
| Streptococcus dysgalactiae subsp. equisimilis strain KNZ06 | SP_2086 | Phosphate ABC transporter, permease protein PstA (TC 3.A.1.7.1) | 12207705 | 100 | 100 | 93 | 1e-136 |
| Streptococcus dysgalactiae subsp. equisimilis strain KNZ06 | SP_1398 | Phosphate ABC transporter, permease protein PstA (TC 3.A.1.7.1) | 12207705 | 100 | 99 | 88 | 1e-148 |
| Streptococcus dysgalactiae subsp. equisimilis strain KNZ07 | SP_1398 | Phosphate ABC transporter, permease protein PstA (TC 3.A.1.7.1) | 12207705 | 100 | 99 | 88 | 1e-148 |
| Streptococcus dysgalactiae subsp. equisimilis strain KNZ07 | SP_2086 | Phosphate ABC transporter, permease protein PstA (TC 3.A.1.7.1) | 12207705 | 100 | 100 | 93 | 1e-136 |
| Streptococcus dysgalactiae subsp. equisimilis strain KNZ10 | SP_1398 | Phosphate ABC transporter, permease protein PstA (TC 3.A.1.7.1) | 12207705 | 100 | 99 | 88 | 1e-148 |
| Streptococcus dysgalactiae subsp. equisimilis strain KNZ10 | SP_2086 | Phosphate ABC transporter, permease protein PstA (TC 3.A.1.7.1) | 12207705 | 100 | 100 | 93 | 1e-136 |
| Streptococcus dysgalactiae subsp. equisimilis strain KNZ12 | SP_2086 | Phosphate ABC transporter, permease protein PstA (TC 3.A.1.7.1) | 12207705 | 100 | 100 | 93 | 1e-136 |
| Streptococcus dysgalactiae subsp. equisimilis strain KNZ12 | SP_1398 | Phosphate ABC transporter, permease protein PstA (TC 3.A.1.7.1) | 12207705 | 100 | 99 | 88 | 1e-148 |
| Streptococcus dysgalactiae subsp. equisimilis strain KNZ15 | SP_1398 | Phosphate ABC transporter, permease protein PstA (TC 3.A.1.7.1) | 12207705 | 100 | 99 | 88 | 1e-148 |
| Streptococcus dysgalactiae subsp. equisimilis strain KNZ15 | SP_2086 | Phosphate ABC transporter, permease protein PstA (TC 3.A.1.7.1) | 12207705 | 100 | 100 | 93 | 1e-136 |
| Streptococcus dysgalactiae subsp. equisimilis strain KNZ16 | SP_1398 | Phosphate ABC transporter, permease protein PstA (TC 3.A.1.7.1) | 12207705 | 100 | 99 | 88 | 1e-148 |
| Streptococcus dysgalactiae subsp. equisimilis strain KNZ16 | SP_2086 | Phosphate ABC transporter, permease protein PstA (TC 3.A.1.7.1) | 12207705 | 100 | 100 | 93 | 1e-136 |
| Streptococcus dysgalactiae subsp. equisimilis strain NCTC10321 | SP_1398 | Phosphate ABC transporter, permease protein PstA (TC 3.A.1.7.1) | 12207705 | 100 | 99 | 88 | 1e-148 |
| Streptococcus dysgalactiae subsp. equisimilis strain NCTC10321 | SP_2086 | Phosphate ABC transporter, permease protein PstA (TC 3.A.1.7.1) | 12207705 | 100 | 100 | 93 | 1e-135 |
| Streptococcus dysgalactiae subsp. equisimilis strain NCTC11554 | SP_1398 | Phosphate ABC transporter, permease protein PstA (TC 3.A.1.7.1) | 12207705 | 100 | 99 | 88 | 1e-148 |
| Streptococcus dysgalactiae subsp. equisimilis strain NCTC11554 | SP_2086 | Phosphate ABC transporter, permease protein PstA (TC 3.A.1.7.1) | 12207705 | 100 | 100 | 93 | 1e-136 |
| Streptococcus dysgalactiae subsp. equisimilis strain NCTC11555 | SP_2086 | Phosphate ABC transporter, permease protein PstA (TC 3.A.1.7.1) | 12207705 | 100 | 100 | 93 | 1e-136 |
| Streptococcus dysgalactiae subsp. equisimilis strain NCTC11555 | SP_1398 | Phosphate ABC transporter, permease protein PstA (TC 3.A.1.7.1) | 12207705 | 100 | 99 | 88 | 1e-148 |
| Streptococcus dysgalactiae subsp. equisimilis strain NCTC11556 | SP_2086 | Phosphate ABC transporter, permease protein PstA (TC 3.A.1.7.1) | 12207705 | 100 | 100 | 93 | 1e-135 |
| Streptococcus dysgalactiae subsp. equisimilis strain NCTC11556 | SP_1398 | Phosphate ABC transporter, permease protein PstA (TC 3.A.1.7.1) | 12207705 | 100 | 99 | 88 | 1e-148 |
| Streptococcus dysgalactiae subsp. equisimilis strain NCTC11557 | SP_1398 | Phosphate ABC transporter, permease protein PstA (TC 3.A.1.7.1) | 12207705 | 100 | 99 | 88 | 1e-147 |
| Streptococcus dysgalactiae subsp. equisimilis strain NCTC11557 | SP_2086 | Phosphate ABC transporter, permease protein PstA (TC 3.A.1.7.1) | 12207705 | 100 | 100 | 93 | 1e-136 |
| Streptococcus dysgalactiae subsp. equisimilis strain NCTC11564 | SP_1398 | Phosphate ABC transporter, permease protein PstA (TC 3.A.1.7.1) | 12207705 | 100 | 99 | 88 | 1e-148 |
| Streptococcus dysgalactiae subsp. equisimilis strain NCTC11564 | SP_2086 | Phosphate ABC transporter, permease protein PstA (TC 3.A.1.7.1) | 12207705 | 100 | 100 | 92 | 1e-135 |
| Streptococcus dysgalactiae subsp. equisimilis strain NCTC5370 | SP_2086 | Phosphate ABC transporter, permease protein PstA (TC 3.A.1.7.1) | 12207705 | 100 | 100 | 93 | 1e-136 |
| Streptococcus dysgalactiae subsp. equisimilis strain NCTC5370 | SP_1398 | Phosphate ABC transporter, permease protein PstA (TC 3.A.1.7.1) | 12207705 | 100 | 99 | 88 | 1e-148 |
| Streptococcus dysgalactiae subsp. equisimilis strain NCTC5371 | SP_2086 | Phosphate ABC transporter, permease protein PstA (TC 3.A.1.7.1) | 12207705 | 100 | 100 | 92 | 1e-135 |
| Streptococcus dysgalactiae subsp. equisimilis strain NCTC5371 | SP_1398 | Phosphate ABC transporter, permease protein PstA (TC 3.A.1.7.1) | 12207705 | 100 | 99 | 88 | 1e-148 |
| Streptococcus dysgalactiae subsp. equisimilis strain NCTC6179 | SP_1398 | Phosphate ABC transporter, permease protein PstA (TC 3.A.1.7.1) | 12207705 | 100 | 99 | 88 | 1e-147 |
| Streptococcus dysgalactiae subsp. equisimilis strain NCTC6179 | SP_2086 | Phosphate ABC transporter, permease protein PstA (TC 3.A.1.7.1) | 12207705 | 100 | 100 | 92 | 1e-135 |
| Streptococcus dysgalactiae subsp. equisimilis strain NCTC6181 | SP_2086 | Phosphate ABC transporter, permease protein PstA (TC 3.A.1.7.1) | 12207705 | 100 | 100 | 92 | 1e-135 |
| Streptococcus dysgalactiae subsp. equisimilis strain NCTC6181 | SP_1398 | Phosphate ABC transporter, permease protein PstA (TC 3.A.1.7.1) | 12207705 | 100 | 99 | 88 | 1e-147 |
| Streptococcus dysgalactiae subsp. equisimilis strain NCTC6407 | SP_2086 | Phosphate ABC transporter, permease protein PstA (TC 3.A.1.7.1) | 12207705 | 100 | 100 | 93 | 1e-135 |
| Streptococcus dysgalactiae subsp. equisimilis strain NCTC6407 | SP_1398 | Phosphate ABC transporter, permease protein PstA (TC 3.A.1.7.1) | 12207705 | 100 | 99 | 88 | 1e-148 |
| Streptococcus dysgalactiae subsp. equisimilis strain NCTC7136 | SP_2086 | Phosphate ABC transporter, permease protein PstA (TC 3.A.1.7.1) | 12207705 | 100 | 100 | 93 | 1e-136 |
| Streptococcus dysgalactiae subsp. equisimilis strain NCTC7136 | SP_1398 | Phosphate ABC transporter, permease protein PstA (TC 3.A.1.7.1) | 12207705 | 100 | 99 | 88 | 1e-148 |
| Streptococcus dysgalactiae subsp. equisimilis strain NCTC8543 | SP_1398 | Phosphate ABC transporter, permease protein PstA (TC 3.A.1.7.1) | 12207705 | 100 | 99 | 88 | 1e-148 |
| Streptococcus dysgalactiae subsp. equisimilis strain NCTC8543 | SP_2086 | Phosphate ABC transporter, permease protein PstA (TC 3.A.1.7.1) | 12207705 | 100 | 100 | 93 | 1e-135 |
| Streptococcus dysgalactiae subsp. equisimilis strain NCTC8546 | SP_1398 | Phosphate ABC transporter, permease protein PstA (TC 3.A.1.7.1) | 12207705 | 100 | 99 | 88 | 1e-148 |
| Streptococcus dysgalactiae subsp. equisimilis strain NCTC8546 | SP_2086 | Phosphate ABC transporter, permease protein PstA (TC 3.A.1.7.1) | 12207705 | 100 | 100 | 93 | 1e-136 |
| Streptococcus dysgalactiae subsp. equisimilis strain NCTC9413 | SP_1398 | Phosphate ABC transporter, permease protein PstA (TC 3.A.1.7.1) | 12207705 | 100 | 99 | 88 | 1e-148 |
| Streptococcus dysgalactiae subsp. equisimilis strain NCTC9413 | SP_2086 | Phosphate ABC transporter, permease protein PstA (TC 3.A.1.7.1) | 12207705 | 100 | 100 | 93 | 1e-136 |
| Streptococcus dysgalactiae subsp. equisimilis strain NCTC9414 | SP_2086 | Phosphate ABC transporter, permease protein PstA (TC 3.A.1.7.1) | 12207705 | 100 | 100 | 92 | 1e-135 |
| Streptococcus dysgalactiae subsp. equisimilis strain NCTC9414 | SP_1398 | Phosphate ABC transporter, permease protein PstA (TC 3.A.1.7.1) | 12207705 | 100 | 99 | 88 | 1e-148 |
| Streptococcus dysgalactiae subsp. equisimilis strain NCTC9603 | SP_2086 | Phosphate ABC transporter, permease protein PstA (TC 3.A.1.7.1) | 12207705 | 100 | 100 | 92 | 1e-135 |
| Streptococcus dysgalactiae subsp. equisimilis strain NCTC9603 | SP_1398 | Phosphate ABC transporter, permease protein PstA (TC 3.A.1.7.1) | 12207705 | 100 | 99 | 88 | 1e-148 |
| Streptococcus dysgalactiae subsp. equisimilis strain SS1575 | SP_1398 | Phosphate ABC transporter, permease protein PstA (TC 3.A.1.7.1) | 12207705 | 100 | 99 | 88 | 1e-148 |
| Streptococcus dysgalactiae subsp. equisimilis strain SS1575 | SP_2086 | Phosphate ABC transporter, permease protein PstA (TC 3.A.1.7.1) | 12207705 | 100 | 100 | 93 | 1e-136 |
| Streptococcus dysgalactiae subsp. equisimilis strain T642 | pstA_1 | Phosphate ABC transporter, permease protein PstA (TC 3.A.1.7.1) | 12207705 | 100 | 99 | 88 | 0 |
| Streptococcus dysgalactiae subsp. equisimilis strain T642 | pstA_1 | Phosphate ABC transporter, permease protein PstA (TC 3.A.1.7.1) | 12207705 | 100 | 100 | 93 | 0 |
| Streptococcus dysgalactiae subsp. equisimilis strain UT_4031CC | pstA_1 | Phosphate ABC transporter, permease protein PstA (TC 3.A.1.7.1) | 12207705 | 100 | 100 | 93 | 1e-136 |
| Streptococcus dysgalactiae subsp. equisimilis strain UT_4031CC | pstA_1 | Phosphate ABC transporter, permease protein PstA (TC 3.A.1.7.1) | 12207705 | 100 | 99 | 88 | 1e-148 |
| Streptococcus dysgalactiae subsp. equisimilis strain UT_4231_KK | pstA_1 | Phosphate ABC transporter, permease protein PstA (TC 3.A.1.7.1) | 12207705 | 100 | 100 | 93 | 1e-136 |
| Streptococcus dysgalactiae subsp. equisimilis strain UT_4231_KK | pstA_1 | Phosphate ABC transporter, permease protein PstA (TC 3.A.1.7.1) | 12207705 | 100 | 99 | 88 | 1e-148 |
| Streptococcus dysgalactiae subsp. equisimilis strain UT_4234_DH | pstA_1 | Phosphate ABC transporter, permease protein PstA (TC 3.A.1.7.1) | 12207705 | 100 | 100 | 93 | 1e-136 |
| Streptococcus dysgalactiae subsp. equisimilis strain UT_4234_DH | pstA_1 | Phosphate ABC transporter, permease protein PstA (TC 3.A.1.7.1) | 12207705 | 100 | 99 | 88 | 1e-147 |
| Streptococcus dysgalactiae subsp. equisimilis strain UT_4241_XS | pstA_1 | Phosphate ABC transporter, permease protein PstA (TC 3.A.1.7.1) | 12207705 | 100 | 99 | 88 | 1e-148 |
| Streptococcus dysgalactiae subsp. equisimilis strain UT_4241_XS | pstA_1 | Phosphate ABC transporter, permease protein PstA (TC 3.A.1.7.1) | 12207705 | 100 | 100 | 93 | 1e-136 |
| Streptococcus dysgalactiae subsp. equisimilis strain UT_4242_AB | pstA_1 | Phosphate ABC transporter, permease protein PstA (TC 3.A.1.7.1) | 12207705 | 100 | 99 | 88 | 1e-148 |
| Streptococcus dysgalactiae subsp. equisimilis strain UT_4242_AB | pstA_1 | Phosphate ABC transporter, permease protein PstA (TC 3.A.1.7.1) | 12207705 | 100 | 100 | 93 | 1e-135 |
| Streptococcus dysgalactiae subsp. equisimilis strain UT_4255RC | pstA_1 | Phosphate ABC transporter, permease protein PstA (TC 3.A.1.7.1) | 12207705 | 100 | 100 | 93 | 1e-136 |
| Streptococcus dysgalactiae subsp. equisimilis strain UT_4255RC | pstA_1 | Phosphate ABC transporter, permease protein PstA (TC 3.A.1.7.1) | 12207705 | 100 | 99 | 88 | 1e-147 |
| Streptococcus dysgalactiae subsp. equisimilis strain UT_4277_BB | pstA_1 | Phosphate ABC transporter, permease protein PstA (TC 3.A.1.7.1) | 12207705 | 100 | 100 | 93 | 1e-136 |
| Streptococcus dysgalactiae subsp. equisimilis strain UT_4277_BB | pstA_1 | Phosphate ABC transporter, permease protein PstA (TC 3.A.1.7.1) | 12207705 | 100 | 99 | 88 | 1e-148 |
| Streptococcus dysgalactiae subsp. equisimilis strain UT_4966_RC | pstA_1 | Phosphate ABC transporter, permease protein PstA (TC 3.A.1.7.1) | 12207705 | 100 | 99 | 88 | 1e-148 |
| Streptococcus dysgalactiae subsp. equisimilis strain UT_4966_RC | pstA_1 | Phosphate ABC transporter, permease protein PstA (TC 3.A.1.7.1) | 12207705 | 100 | 100 | 93 | 1e-136 |
| Streptococcus dysgalactiae subsp. equisimilis strain UT-5345 | pstA_1 | Phosphate ABC transporter, permease protein PstA (TC 3.A.1.7.1) | 12207705 | 100 | 100 | 93 | 1e-136 |
| Streptococcus dysgalactiae subsp. equisimilis strain UT-5345 | pstA_1 | Phosphate ABC transporter, permease protein PstA (TC 3.A.1.7.1) | 12207705 | 94 | 99 | 89 | 1e-140 |
| Streptococcus dysgalactiae subsp. equisimilis strain UT-5354 | pstA_1 | Phosphate ABC transporter, permease protein PstA (TC 3.A.1.7.1) | 12207705 | 100 | 100 | 93 | 1e-135 |
| Streptococcus dysgalactiae subsp. equisimilis strain UT-5354 | pstA_1 | Phosphate ABC transporter, permease protein PstA (TC 3.A.1.7.1) | 12207705 | 100 | 99 | 88 | 1e-148 |
| Streptococcus dysgalactiae subsp. equisimilis strain UT-SS1069 | pstA_1 | Phosphate ABC transporter, permease protein PstA (TC 3.A.1.7.1) | 12207705 | 100 | 100 | 93 | 1e-136 |
| Streptococcus dysgalactiae subsp. equisimilis strain UT-SS1069 | pstA_1 | Phosphate ABC transporter, permease protein PstA (TC 3.A.1.7.1) | 12207705 | 100 | 99 | 88 | 1e-148 |
| Streptococcus dysgalactiae subsp. equisimilis strain UT-SS957 | pstA_1 | Phosphate ABC transporter, permease protein PstA (TC 3.A.1.7.1) | 12207705 | 100 | 99 | 88 | 1e-148 |
| Streptococcus dysgalactiae subsp. equisimilis strain UT-SS957 | pstA_1 | Phosphate ABC transporter, permease protein PstA (TC 3.A.1.7.1) | 12207705 | 100 | 100 | 92 | 1e-135 |
| Streptococcus dysgalactiae subsp. equisimilis strain WCHSDSE-1 | pstA_1 | Phosphate ABC transporter, permease protein PstA (TC 3.A.1.7.1) | 12207705 | 100 | 99 | 88 | 1e-148 |
| Streptococcus dysgalactiae subsp. equisimilis strain WCHSDSE-1 | pstA_1 | Phosphate ABC transporter, permease protein PstA (TC 3.A.1.7.1) | 12207705 | 100 | 100 | 93 | 1e-136 |
| Streptococcus dysgalactiae subsp. equisimilis 167 | fba | Fructose-bisphosphate aldolase class II (EC 4.1.2.13) | 11679068 | 100 | 100 | 99 | 1e-166 |
| Streptococcus dysgalactiae subsp. equisimilis AC-2713 | fba | Fructose-bisphosphate aldolase class II (EC 4.1.2.13) | 11679068 | 100 | 100 | 99 | 1e-166 |
| Streptococcus dysgalactiae subsp. equisimilis AKSDE4288 | fba | Fructose-bisphosphate aldolase class II (EC 4.1.2.13) | 11679068 | 100 | 100 | 99 | 1e-166 |
| Streptococcus dysgalactiae subsp. equisimilis ATCC 12394 | fba | Fructose-bisphosphate aldolase class II (EC 4.1.2.13) | 11679068 | 100 | 100 | 99 | 1e-166 |
| Streptococcus dysgalactiae subsp. equisimilis GGS_124 | fba | Fructose-bisphosphate aldolase class II (EC 4.1.2.13) | 11679068 | 100 | 100 | 99 | 1e-166 |
| Streptococcus dysgalactiae subsp. equisimilis RE378 | fba | Fructose-bisphosphate aldolase class II (EC 4.1.2.13) | 11679068 | 100 | 100 | 99 | 1e-166 |
| Streptococcus dysgalactiae subsp. equisimilis SD SCDR1 | fba | Fructose-bisphosphate aldolase class II (EC 4.1.2.13) | 11679068 | 100 | 100 | 99 | 1e-166 |
| Streptococcus dysgalactiae subsp. equisimilis SK1249 | fba | Fructose-bisphosphate aldolase class II (EC 4.1.2.13) | 11679068 | 100 | 100 | 99 | 1e-166 |
| Streptococcus dysgalactiae subsp. equisimilis SK1250 | fba | Fructose-bisphosphate aldolase class II (EC 4.1.2.13) | 11679068 | 100 | 100 | 99 | 1e-166 |
| Streptococcus dysgalactiae subsp. equisimilis strain ASDSE_96 | fba | Fructose-bisphosphate aldolase class II (EC 4.1.2.13) | 11679068 | 100 | 100 | 99 | 1e-166 |
| Streptococcus dysgalactiae subsp. equisimilis strain ASDSE_99 | fba | Fructose-bisphosphate aldolase class II (EC 4.1.2.13) | 11679068 | 100 | 100 | 99 | 1e-166 |
| Streptococcus dysgalactiae subsp. equisimilis strain C161L1 | fba | Fructose-bisphosphate aldolase class II (EC 4.1.2.13) | 11679068 | 100 | 100 | 99 | 0 |
| Streptococcus dysgalactiae subsp. equisimilis strain KNZ01 | fba | Fructose-bisphosphate aldolase class II (EC 4.1.2.13) | 11679068 | 100 | 100 | 99 | 1e-166 |
| Streptococcus dysgalactiae subsp. equisimilis strain KNZ03 | fba | Fructose-bisphosphate aldolase class II (EC 4.1.2.13) | 11679068 | 100 | 100 | 99 | 1e-166 |
| Streptococcus dysgalactiae subsp. equisimilis strain KNZ04 | fba | Fructose-bisphosphate aldolase class II (EC 4.1.2.13) | 11679068 | 100 | 100 | 99 | 1e-166 |
| Streptococcus dysgalactiae subsp. equisimilis strain KNZ06 | fba | Fructose-bisphosphate aldolase class II (EC 4.1.2.13) | 11679068 | 100 | 100 | 99 | 1e-166 |
| Streptococcus dysgalactiae subsp. equisimilis strain KNZ07 | fba | Fructose-bisphosphate aldolase class II (EC 4.1.2.13) | 11679068 | 100 | 100 | 99 | 1e-166 |
| Streptococcus dysgalactiae subsp. equisimilis strain KNZ10 | fba | Fructose-bisphosphate aldolase class II (EC 4.1.2.13) | 11679068 | 100 | 100 | 99 | 1e-166 |
| Streptococcus dysgalactiae subsp. equisimilis strain KNZ12 | fba | Fructose-bisphosphate aldolase class II (EC 4.1.2.13) | 11679068 | 100 | 100 | 99 | 1e-166 |
| Streptococcus dysgalactiae subsp. equisimilis strain KNZ15 | fba | Fructose-bisphosphate aldolase class II (EC 4.1.2.13) | 11679068 | 100 | 100 | 99 | 1e-166 |
| Streptococcus dysgalactiae subsp. equisimilis strain KNZ16 | fba | Fructose-bisphosphate aldolase class II (EC 4.1.2.13) | 11679068 | 100 | 100 | 99 | 1e-166 |
| Streptococcus dysgalactiae subsp. equisimilis strain NCTC10321 | fba | Fructose-bisphosphate aldolase class II (EC 4.1.2.13) | 11679068 | 66 | 99 | 99 | 1e-107 |
| Streptococcus dysgalactiae subsp. equisimilis strain NCTC11554 | fba | Fructose-bisphosphate aldolase class II (EC 4.1.2.13) | 11679068 | 100 | 100 | 99 | 1e-166 |
| Streptococcus dysgalactiae subsp. equisimilis strain NCTC11555 | fba | Fructose-bisphosphate aldolase class II (EC 4.1.2.13) | 11679068 | 100 | 100 | 99 | 1e-166 |
| Streptococcus dysgalactiae subsp. equisimilis strain NCTC11556 | fba | Fructose-bisphosphate aldolase class II (EC 4.1.2.13) | 11679068 | 100 | 100 | 99 | 1e-166 |
| Streptococcus dysgalactiae subsp. equisimilis strain NCTC11557 | fba | Fructose-bisphosphate aldolase class II (EC 4.1.2.13) | 11679068 | 100 | 100 | 99 | 1e-166 |
| Streptococcus dysgalactiae subsp. equisimilis strain NCTC11564 | fba | Fructose-bisphosphate aldolase class II (EC 4.1.2.13) | 11679068 | 100 | 100 | 99 | 1e-166 |
| Streptococcus dysgalactiae subsp. equisimilis strain NCTC5370 | fba | Fructose-bisphosphate aldolase class II (EC 4.1.2.13) | 11679068 | 100 | 100 | 99 | 1e-166 |
| Streptococcus dysgalactiae subsp. equisimilis strain NCTC5371 | fba | Fructose-bisphosphate aldolase class II (EC 4.1.2.13) | 11679068 | 100 | 100 | 99 | 1e-166 |
| Streptococcus dysgalactiae subsp. equisimilis strain NCTC6179 | fba | Fructose-bisphosphate aldolase class II (EC 4.1.2.13) | 11679068 | 100 | 100 | 99 | 1e-166 |
| Streptococcus dysgalactiae subsp. equisimilis strain NCTC6181 | fba | Fructose-bisphosphate aldolase class II (EC 4.1.2.13) | 11679068 | 100 | 100 | 99 | 1e-166 |
| Streptococcus dysgalactiae subsp. equisimilis strain NCTC6407 | fba | Fructose-bisphosphate aldolase class II (EC 4.1.2.13) | 11679068 | 100 | 100 | 99 | 1e-166 |
| Streptococcus dysgalactiae subsp. equisimilis strain NCTC7136 | fba | Fructose-bisphosphate aldolase class II (EC 4.1.2.13) | 11679068 | 100 | 100 | 99 | 1e-166 |
| Streptococcus dysgalactiae subsp. equisimilis strain NCTC8543 | fba | Fructose-bisphosphate aldolase class II (EC 4.1.2.13) | 11679068 | 100 | 100 | 99 | 1e-166 |
| Streptococcus dysgalactiae subsp. equisimilis strain NCTC8546 | fba | Fructose-bisphosphate aldolase class II (EC 4.1.2.13) | 11679068 | 100 | 100 | 99 | 1e-166 |
| Streptococcus dysgalactiae subsp. equisimilis strain NCTC9413 | fba | Fructose-bisphosphate aldolase class II (EC 4.1.2.13) | 11679068 | 100 | 100 | 99 | 1e-166 |
| Streptococcus dysgalactiae subsp. equisimilis strain NCTC9414 | fba | Fructose-bisphosphate aldolase class II (EC 4.1.2.13) | 11679068 | 100 | 100 | 99 | 1e-166 |
| Streptococcus dysgalactiae subsp. equisimilis strain NCTC9603 | fba | Fructose-bisphosphate aldolase class II (EC 4.1.2.13) | 11679068 | 100 | 100 | 99 | 1e-166 |
| Streptococcus dysgalactiae subsp. equisimilis strain SS1575 | fba | Fructose-bisphosphate aldolase class II (EC 4.1.2.13) | 11679068 | 100 | 100 | 99 | 1e-166 |
| Streptococcus dysgalactiae subsp. equisimilis strain T642 | fba | Fructose-bisphosphate aldolase class II (EC 4.1.2.13) | 11679068 | 100 | 100 | 99 | 0 |
| Streptococcus dysgalactiae subsp. equisimilis strain UT_4031CC | fba | Fructose-bisphosphate aldolase class II (EC 4.1.2.13) | 11679068 | 100 | 100 | 99 | 1e-166 |
| Streptococcus dysgalactiae subsp. equisimilis strain UT_4231_KK | fba | Fructose-bisphosphate aldolase class II (EC 4.1.2.13) | 11679068 | 100 | 100 | 99 | 1e-166 |
| Streptococcus dysgalactiae subsp. equisimilis strain UT_4234_DH | fba | Fructose-bisphosphate aldolase class II (EC 4.1.2.13) | 11679068 | 100 | 100 | 99 | 1e-166 |
| Streptococcus dysgalactiae subsp. equisimilis strain UT_4241_XS | fba | Fructose-bisphosphate aldolase class II (EC 4.1.2.13) | 11679068 | 100 | 100 | 99 | 1e-166 |
| Streptococcus dysgalactiae subsp. equisimilis strain UT_4242_AB | fba | Fructose-bisphosphate aldolase class II (EC 4.1.2.13) | 11679068 | 100 | 100 | 99 | 1e-166 |
| Streptococcus dysgalactiae subsp. equisimilis strain UT_4255RC | fba | Fructose-bisphosphate aldolase class II (EC 4.1.2.13) | 11679068 | 100 | 100 | 99 | 1e-166 |
| Streptococcus dysgalactiae subsp. equisimilis strain UT_4277_BB | fba | Fructose-bisphosphate aldolase class II (EC 4.1.2.13) | 11679068 | 100 | 100 | 99 | 1e-166 |
| Streptococcus dysgalactiae subsp. equisimilis strain UT_4966_RC | fba | Fructose-bisphosphate aldolase class II (EC 4.1.2.13) | 11679068 | 100 | 100 | 99 | 1e-166 |
| Streptococcus dysgalactiae subsp. equisimilis strain UT-5345 | fba | Fructose-bisphosphate aldolase class II (EC 4.1.2.13) | 11679068 | 100 | 100 | 99 | 1e-166 |
| Streptococcus dysgalactiae subsp. equisimilis strain UT-5354 | fba | Fructose-bisphosphate aldolase class II (EC 4.1.2.13) | 11679068 | 100 | 100 | 99 | 1e-166 |
| Streptococcus dysgalactiae subsp. equisimilis strain UT-SS1069 | fba | Fructose-bisphosphate aldolase class II (EC 4.1.2.13) | 11679068 | 100 | 100 | 99 | 1e-166 |
| Streptococcus dysgalactiae subsp. equisimilis strain UT-SS957 | fba | Fructose-bisphosphate aldolase class II (EC 4.1.2.13) | 11679068 | 100 | 100 | 99 | 1e-166 |
| Streptococcus dysgalactiae subsp. equisimilis strain WCHSDSE-1 | fba | Fructose-bisphosphate aldolase class II (EC 4.1.2.13) | 11679068 | 100 | 100 | 99 | 1e-166 |
| Streptococcus dysgalactiae subsp. equisimilis 167 | glnA | Glutamine synthetase type I (EC 6.3.1.2) | 16787930 | 99 | 99 | 86 | 1e-233 |
| Streptococcus dysgalactiae subsp. equisimilis AC-2713 | glnA | Glutamine synthetase type I (EC 6.3.1.2) | 16787930 | 99 | 99 | 86 | 1e-233 |
| Streptococcus dysgalactiae subsp. equisimilis AKSDE4288 | glnA | Glutamine synthetase type I (EC 6.3.1.2) | 16787930 | 99 | 99 | 86 | 1e-233 |
| Streptococcus dysgalactiae subsp. equisimilis ATCC 12394 | glnA | Glutamine synthetase type I (EC 6.3.1.2) | 16787930 | 99 | 99 | 86 | 1e-233 |
| Streptococcus dysgalactiae subsp. equisimilis GGS_124 | glnA | Glutamine synthetase type I (EC 6.3.1.2) | 16787930 | 99 | 99 | 86 | 1e-233 |
| Streptococcus dysgalactiae subsp. equisimilis RE378 | glnA | Glutamine synthetase type I (EC 6.3.1.2) | 16787930 | 99 | 99 | 86 | 1e-233 |
| Streptococcus dysgalactiae subsp. equisimilis SD SCDR1 | glnA | Glutamine synthetase type I (EC 6.3.1.2) | 16787930 | 99 | 99 | 86 | 1e-233 |
| Streptococcus dysgalactiae subsp. equisimilis SK1249 | glnA | Glutamine synthetase type I (EC 6.3.1.2) | 16787930 | 99 | 99 | 86 | 1e-233 |
| Streptococcus dysgalactiae subsp. equisimilis SK1250 | glnA | Glutamine synthetase type I (EC 6.3.1.2) | 16787930 | 99 | 99 | 86 | 1e-233 |
| Streptococcus dysgalactiae subsp. equisimilis strain ASDSE_96 | glnA | Glutamine synthetase type I (EC 6.3.1.2) | 16787930 | 99 | 99 | 86 | 1e-233 |
| Streptococcus dysgalactiae subsp. equisimilis strain ASDSE_99 | glnA | Glutamine synthetase type I (EC 6.3.1.2) | 16787930 | 99 | 99 | 86 | 1e-233 |
| Streptococcus dysgalactiae subsp. equisimilis strain C161L1 | glnA | Glutamine synthetase type I (EC 6.3.1.2) | 16787930 | 100 | 100 | 85 | 0 |
| Streptococcus dysgalactiae subsp. equisimilis strain KNZ01 | glnA | Glutamine synthetase type I (EC 6.3.1.2) | 16787930 | 99 | 99 | 86 | 1e-233 |
| Streptococcus dysgalactiae subsp. equisimilis strain KNZ03 | glnA | Glutamine synthetase type I (EC 6.3.1.2) | 16787930 | 99 | 99 | 86 | 1e-233 |
| Streptococcus dysgalactiae subsp. equisimilis strain KNZ04 | glnA | Glutamine synthetase type I (EC 6.3.1.2) | 16787930 | 99 | 99 | 86 | 1e-233 |
| Streptococcus dysgalactiae subsp. equisimilis strain KNZ06 | glnA | Glutamine synthetase type I (EC 6.3.1.2) | 16787930 | 99 | 99 | 86 | 1e-233 |
| Streptococcus dysgalactiae subsp. equisimilis strain KNZ07 | glnA | Glutamine synthetase type I (EC 6.3.1.2) | 16787930 | 99 | 99 | 86 | 1e-233 |
| Streptococcus dysgalactiae subsp. equisimilis strain KNZ10 | glnA | Glutamine synthetase type I (EC 6.3.1.2) | 16787930 | 99 | 99 | 86 | 1e-233 |
| Streptococcus dysgalactiae subsp. equisimilis strain KNZ12 | glnA | Glutamine synthetase type I (EC 6.3.1.2) | 16787930 | 99 | 99 | 86 | 1e-233 |
| Streptococcus dysgalactiae subsp. equisimilis strain KNZ15 | glnA | Glutamine synthetase type I (EC 6.3.1.2) | 16787930 | 99 | 99 | 86 | 1e-233 |
| Streptococcus dysgalactiae subsp. equisimilis strain KNZ16 | glnA | Glutamine synthetase type I (EC 6.3.1.2) | 16787930 | 99 | 99 | 86 | 1e-233 |
| Streptococcus dysgalactiae subsp. equisimilis strain NCTC10321 | glnA | Glutamine synthetase type I (EC 6.3.1.2) | 16787930 | 99 | 99 | 86 | 1e-233 |
| Streptococcus dysgalactiae subsp. equisimilis strain NCTC11554 | glnA | Glutamine synthetase type I (EC 6.3.1.2) | 16787930 | 99 | 99 | 86 | 1e-233 |
| Streptococcus dysgalactiae subsp. equisimilis strain NCTC11555 | glnA | Glutamine synthetase type I (EC 6.3.1.2) | 16787930 | 99 | 99 | 86 | 1e-233 |
| Streptococcus dysgalactiae subsp. equisimilis strain NCTC11556 | glnA | Glutamine synthetase type I (EC 6.3.1.2) | 16787930 | 99 | 99 | 86 | 1e-233 |
| Streptococcus dysgalactiae subsp. equisimilis strain NCTC11557 | glnA | Glutamine synthetase type I (EC 6.3.1.2) | 16787930 | 99 | 99 | 86 | 1e-233 |
| Streptococcus dysgalactiae subsp. equisimilis strain NCTC11564 | glnA | Glutamine synthetase type I (EC 6.3.1.2) | 16787930 | 99 | 99 | 86 | 1e-233 |
| Streptococcus dysgalactiae subsp. equisimilis strain NCTC5370 | glnA | Glutamine synthetase type I (EC 6.3.1.2) | 16787930 | 99 | 99 | 86 | 1e-233 |
| Streptococcus dysgalactiae subsp. equisimilis strain NCTC5371 | glnA | Glutamine synthetase type I (EC 6.3.1.2) | 16787930 | 99 | 99 | 86 | 1e-233 |
| Streptococcus dysgalactiae subsp. equisimilis strain NCTC6179 | glnA | Glutamine synthetase type I (EC 6.3.1.2) | 16787930 | 99 | 99 | 86 | 1e-233 |
| Streptococcus dysgalactiae subsp. equisimilis strain NCTC6181 | glnA | Glutamine synthetase type I (EC 6.3.1.2) | 16787930 | 99 | 99 | 86 | 1e-233 |
| Streptococcus dysgalactiae subsp. equisimilis strain NCTC6407 | glnA | Glutamine synthetase type I (EC 6.3.1.2) | 16787930 | 99 | 99 | 86 | 1e-233 |
| Streptococcus dysgalactiae subsp. equisimilis strain NCTC7136 | glnA | Glutamine synthetase type I (EC 6.3.1.2) | 16787930 | 99 | 99 | 86 | 1e-233 |
| Streptococcus dysgalactiae subsp. equisimilis strain NCTC8543 | glnA | Glutamine synthetase type I (EC 6.3.1.2) | 16787930 | 99 | 99 | 86 | 1e-233 |
| Streptococcus dysgalactiae subsp. equisimilis strain NCTC8546 | glnA | Glutamine synthetase type I (EC 6.3.1.2) | 16787930 | 99 | 99 | 86 | 1e-233 |
| Streptococcus dysgalactiae subsp. equisimilis strain NCTC9413 | glnA | Glutamine synthetase type I (EC 6.3.1.2) | 16787930 | 99 | 99 | 86 | 1e-233 |
| Streptococcus dysgalactiae subsp. equisimilis strain NCTC9414 | glnA | Glutamine synthetase type I (EC 6.3.1.2) | 16787930 | 99 | 99 | 86 | 1e-233 |
| Streptococcus dysgalactiae subsp. equisimilis strain NCTC9603 | glnA | Glutamine synthetase type I (EC 6.3.1.2) | 16787930 | 99 | 99 | 86 | 1e-233 |
| Streptococcus dysgalactiae subsp. equisimilis strain SS1575 | glnA | Glutamine synthetase type I (EC 6.3.1.2) | 16787930 | 99 | 99 | 86 | 1e-233 |
| Streptococcus dysgalactiae subsp. equisimilis strain T642 | glnA | Glutamine synthetase type I (EC 6.3.1.2) | 16787930 | 100 | 100 | 85 | 0 |
| Streptococcus dysgalactiae subsp. equisimilis strain UT_4031CC | glnA | Glutamine synthetase type I (EC 6.3.1.2) | 16787930 | 99 | 99 | 85 | 1e-232 |
| Streptococcus dysgalactiae subsp. equisimilis strain UT_4231_KK | glnA | Glutamine synthetase type I (EC 6.3.1.2) | 16787930 | 99 | 99 | 86 | 1e-233 |
| Streptococcus dysgalactiae subsp. equisimilis strain UT_4234_DH | glnA | Glutamine synthetase type I (EC 6.3.1.2) | 16787930 | 99 | 99 | 86 | 1e-233 |
| Streptococcus dysgalactiae subsp. equisimilis strain UT_4241_XS | glnA | Glutamine synthetase type I (EC 6.3.1.2) | 16787930 | 99 | 99 | 86 | 1e-233 |
| Streptococcus dysgalactiae subsp. equisimilis strain UT_4242_AB | glnA | Glutamine synthetase type I (EC 6.3.1.2) | 16787930 | 99 | 99 | 86 | 1e-233 |
| Streptococcus dysgalactiae subsp. equisimilis strain UT_4255RC | glnA | Glutamine synthetase type I (EC 6.3.1.2) | 16787930 | 99 | 99 | 86 | 1e-233 |
| Streptococcus dysgalactiae subsp. equisimilis strain UT_4277_BB | glnA | Glutamine synthetase type I (EC 6.3.1.2) | 16787930 | 99 | 99 | 86 | 1e-233 |
| Streptococcus dysgalactiae subsp. equisimilis strain UT_4966_RC | glnA | Glutamine synthetase type I (EC 6.3.1.2) | 16787930 | 99 | 99 | 86 | 1e-233 |
| Streptococcus dysgalactiae subsp. equisimilis strain UT-5345 | glnA | Glutamine synthetase type I (EC 6.3.1.2) | 16787930 | 99 | 99 | 86 | 1e-233 |
| Streptococcus dysgalactiae subsp. equisimilis strain UT-5354 | glnA | Glutamine synthetase type I (EC 6.3.1.2) | 16787930 | 99 | 99 | 86 | 1e-233 |
| Streptococcus dysgalactiae subsp. equisimilis strain UT-SS1069 | glnA | Glutamine synthetase type I (EC 6.3.1.2) | 16787930 | 99 | 99 | 85 | 1e-232 |
| Streptococcus dysgalactiae subsp. equisimilis strain UT-SS957 | glnA | Glutamine synthetase type I (EC 6.3.1.2) | 16787930 | 99 | 99 | 86 | 1e-233 |
| Streptococcus dysgalactiae subsp. equisimilis strain WCHSDSE-1 | glnA | Glutamine synthetase type I (EC 6.3.1.2) | 16787930 | 99 | 99 | 86 | 1e-233 |
| Streptococcus dysgalactiae subsp. equisimilis 167 | gldA | Glycerol dehydrogenase (EC 1.1.1.6) | 12207705 | 82 | 82 | 80 | 1e-137 |
| Streptococcus dysgalactiae subsp. equisimilis AC-2713 | gldA | Glycerol dehydrogenase (EC 1.1.1.6) | 12207705 | 82 | 82 | 80 | 1e-136 |
| Streptococcus dysgalactiae subsp. equisimilis AKSDE4288 | glnA | Glycerol dehydrogenase (EC 1.1.1.6) | 12207705 | 82 | 82 | 80 | 1e-136 |
| Streptococcus dysgalactiae subsp. equisimilis ATCC 12394 | gldA | Glycerol dehydrogenase (EC 1.1.1.6) | 12207705 | 82 | 82 | 80 | 1e-137 |
| Streptococcus dysgalactiae subsp. equisimilis GGS_124 | gldA | Glycerol dehydrogenase (EC 1.1.1.6) | 12207705 | 82 | 82 | 80 | 1e-137 |
| Streptococcus dysgalactiae subsp. equisimilis RE378 | gldA | Glycerol dehydrogenase (EC 1.1.1.6) | 12207705 | 82 | 82 | 80 | 1e-137 |
| Streptococcus dysgalactiae subsp. equisimilis SD SCDR1 | gldA | Glycerol dehydrogenase (EC 1.1.1.6) | 12207705 | 82 | 82 | 80 | 1e-137 |
| Streptococcus dysgalactiae subsp. equisimilis SK1249 | gldA | Glycerol dehydrogenase (EC 1.1.1.6) | 12207705 | 82 | 82 | 80 | 1e-137 |
| Streptococcus dysgalactiae subsp. equisimilis SK1250 | gldA | Glycerol dehydrogenase (EC 1.1.1.6) | 12207705 | 82 | 82 | 80 | 1e-137 |
| Streptococcus dysgalactiae subsp. equisimilis strain ASDSE_96 | glnA | Glycerol dehydrogenase (EC 1.1.1.6) | 12207705 | 82 | 82 | 80 | 1e-137 |
| Streptococcus dysgalactiae subsp. equisimilis strain ASDSE_99 | glnA | Glycerol dehydrogenase (EC 1.1.1.6) | 12207705 | 82 | 82 | 81 | 1e-138 |
| Streptococcus dysgalactiae subsp. equisimilis strain KNZ01 | gldA | Glycerol dehydrogenase (EC 1.1.1.6) | 12207705 | 82 | 82 | 80 | 1e-137 |
| Streptococcus dysgalactiae subsp. equisimilis strain KNZ03 | gldA | Glycerol dehydrogenase (EC 1.1.1.6) | 12207705 | 82 | 82 | 80 | 1e-137 |
| Streptococcus dysgalactiae subsp. equisimilis strain KNZ04 | gldA | Glycerol dehydrogenase (EC 1.1.1.6) | 12207705 | 82 | 82 | 80 | 1e-137 |
| Streptococcus dysgalactiae subsp. equisimilis strain KNZ06 | gldA | Glycerol dehydrogenase (EC 1.1.1.6) | 12207705 | 82 | 82 | 81 | 1e-137 |
| Streptococcus dysgalactiae subsp. equisimilis strain KNZ07 | gldA | Glycerol dehydrogenase (EC 1.1.1.6) | 12207705 | 82 | 82 | 80 | 1e-137 |
| Streptococcus dysgalactiae subsp. equisimilis strain KNZ10 | gldA | Glycerol dehydrogenase (EC 1.1.1.6) | 12207705 | 82 | 82 | 80 | 1e-137 |
| Streptococcus dysgalactiae subsp. equisimilis strain KNZ12 | gldA | Glycerol dehydrogenase (EC 1.1.1.6) | 12207705 | 82 | 82 | 80 | 1e-137 |
| Streptococcus dysgalactiae subsp. equisimilis strain KNZ15 | gldA | Glycerol dehydrogenase (EC 1.1.1.6) | 12207705 | 82 | 82 | 80 | 1e-137 |
| Streptococcus dysgalactiae subsp. equisimilis strain KNZ16 | gldA | Glycerol dehydrogenase (EC 1.1.1.6) | 12207705 | 82 | 82 | 80 | 1e-137 |
| Streptococcus dysgalactiae subsp. equisimilis strain NCTC10321 | gldA | Glycerol dehydrogenase (EC 1.1.1.6) | 12207705 | 82 | 82 | 80 | 1e-137 |
| Streptococcus dysgalactiae subsp. equisimilis strain NCTC11554 | gldA | Glycerol dehydrogenase (EC 1.1.1.6) | 12207705 | 82 | 82 | 80 | 1e-137 |
| Streptococcus dysgalactiae subsp. equisimilis strain NCTC11555 | gldA | Glycerol dehydrogenase (EC 1.1.1.6) | 12207705 | 82 | 82 | 80 | 1e-137 |
| Streptococcus dysgalactiae subsp. equisimilis strain NCTC11556 | gldA | Glycerol dehydrogenase (EC 1.1.1.6) | 12207705 | 82 | 82 | 80 | 1e-137 |
| Streptococcus dysgalactiae subsp. equisimilis strain NCTC11557 | gldA | Glycerol dehydrogenase (EC 1.1.1.6) | 12207705 | 82 | 82 | 81 | 1e-138 |
| Streptococcus dysgalactiae subsp. equisimilis strain NCTC11564 | gldA | Glycerol dehydrogenase (EC 1.1.1.6) | 12207705 | 82 | 82 | 80 | 1e-137 |
| Streptococcus dysgalactiae subsp. equisimilis strain NCTC5370 | gldA | Glycerol dehydrogenase (EC 1.1.1.6) | 12207705 | 82 | 82 | 80 | 1e-137 |
| Streptococcus dysgalactiae subsp. equisimilis strain NCTC5371 | gldA | Glycerol dehydrogenase (EC 1.1.1.6) | 12207705 | 82 | 82 | 80 | 1e-137 |
| Streptococcus dysgalactiae subsp. equisimilis strain NCTC6179 | gldA | Glycerol dehydrogenase (EC 1.1.1.6) | 12207705 | 82 | 82 | 80 | 1e-137 |
| Streptococcus dysgalactiae subsp. equisimilis strain NCTC6181 | gldA | Glycerol dehydrogenase (EC 1.1.1.6) | 12207705 | 82 | 82 | 80 | 1e-137 |
| Streptococcus dysgalactiae subsp. equisimilis strain NCTC6407 | gldA | Glycerol dehydrogenase (EC 1.1.1.6) | 12207705 | 82 | 82 | 80 | 1e-137 |
| Streptococcus dysgalactiae subsp. equisimilis strain NCTC7136 | gldA | Glycerol dehydrogenase (EC 1.1.1.6) | 12207705 | 82 | 82 | 81 | 1e-137 |
| Streptococcus dysgalactiae subsp. equisimilis strain NCTC8543 | gldA | Glycerol dehydrogenase (EC 1.1.1.6) | 12207705 | 82 | 82 | 80 | 1e-137 |
| Streptococcus dysgalactiae subsp. equisimilis strain NCTC8546 | gldA | Glycerol dehydrogenase (EC 1.1.1.6) | 12207705 | 82 | 82 | 80 | 1e-137 |
| Streptococcus dysgalactiae subsp. equisimilis strain NCTC9413 | gldA | Glycerol dehydrogenase (EC 1.1.1.6) | 12207705 | 82 | 82 | 80 | 1e-137 |
| Streptococcus dysgalactiae subsp. equisimilis strain NCTC9414 | gldA | Glycerol dehydrogenase (EC 1.1.1.6) | 12207705 | 82 | 82 | 80 | 1e-137 |
| Streptococcus dysgalactiae subsp. equisimilis strain NCTC9603 | gldA | Glycerol dehydrogenase (EC 1.1.1.6) | 12207705 | 82 | 82 | 80 | 1e-137 |
| Streptococcus dysgalactiae subsp. equisimilis strain SS1575 | gldA | Glycerol dehydrogenase (EC 1.1.1.6) | 12207705 | 82 | 82 | 81 | 1e-138 |
| Streptococcus dysgalactiae subsp. equisimilis strain UT_4031CC | gldA | Glycerol dehydrogenase (EC 1.1.1.6) | 12207705 | 82 | 82 | 80 | 1e-136 |
| Streptococcus dysgalactiae subsp. equisimilis strain UT_4231_KK | gldA | Glycerol dehydrogenase (EC 1.1.1.6) | 12207705 | 82 | 82 | 80 | 1e-137 |
| Streptococcus dysgalactiae subsp. equisimilis strain UT_4234_DH | gldA | Glycerol dehydrogenase (EC 1.1.1.6) | 12207705 | 82 | 82 | 81 | 1e-138 |
| Streptococcus dysgalactiae subsp. equisimilis strain UT_4241_XS | gldA | Glycerol dehydrogenase (EC 1.1.1.6) | 12207705 | 82 | 82 | 80 | 1e-137 |
| Streptococcus dysgalactiae subsp. equisimilis strain UT_4242_AB | gldA | Glycerol dehydrogenase (EC 1.1.1.6) | 12207705 | 82 | 82 | 80 | 1e-137 |
| Streptococcus dysgalactiae subsp. equisimilis strain UT_4255RC | gldA | Glycerol dehydrogenase (EC 1.1.1.6) | 12207705 | 82 | 82 | 81 | 1e-138 |
| Streptococcus dysgalactiae subsp. equisimilis strain UT_4277_BB | gldA | Glycerol dehydrogenase (EC 1.1.1.6) | 12207705 | 82 | 82 | 80 | 1e-137 |
| Streptococcus dysgalactiae subsp. equisimilis strain UT_4966_RC | gldA | Glycerol dehydrogenase (EC 1.1.1.6) | 12207705 | 82 | 82 | 80 | 1e-137 |
| Streptococcus dysgalactiae subsp. equisimilis strain UT-5345 | gldA | Glycerol dehydrogenase (EC 1.1.1.6) | 12207705 | 82 | 82 | 80 | 1e-137 |
| Streptococcus dysgalactiae subsp. equisimilis strain UT-5354 | gldA | Glycerol dehydrogenase (EC 1.1.1.6) | 12207705 | 82 | 82 | 80 | 1e-137 |
| Streptococcus dysgalactiae subsp. equisimilis strain UT-SS1069 | gldA | Glycerol dehydrogenase (EC 1.1.1.6) | 12207705 | 82 | 82 | 80 | 1e-137 |
| Streptococcus dysgalactiae subsp. equisimilis strain UT-SS957 | gldA | Glycerol dehydrogenase (EC 1.1.1.6) | 12207705 | 82 | 82 | 80 | 1e-137 |
| Streptococcus dysgalactiae subsp. equisimilis strain WCHSDSE-1 | gldA | Glycerol dehydrogenase (EC 1.1.1.6) | 12207705 | 82 | 82 | 80 | 1e-137 |
| Streptococcus dysgalactiae subsp. equisimilis 167 | guaA | GMP synthase [glutamine-hydrolyzing], amidotransferase subunit (EC 6.3.5.2) / GMP synthase [glutamine-hydrolyzing], ATP pyrophosphatase subunit (EC 6.3.5.2) | 10998175 | 100 | 100 | 93 | 1e-287 |
| Streptococcus dysgalactiae subsp. equisimilis AC-2713 | guaA | GMP synthase [glutamine-hydrolyzing], amidotransferase subunit (EC 6.3.5.2) / GMP synthase [glutamine-hydrolyzing], ATP pyrophosphatase subunit (EC 6.3.5.2) | 10998175 | 100 | 100 | 93 | 1e-287 |
| Streptococcus dysgalactiae subsp. equisimilis AKSDE4288 | guaA | GMP synthase [glutamine-hydrolyzing], amidotransferase subunit (EC 6.3.5.2) / GMP synthase [glutamine-hydrolyzing], ATP pyrophosphatase subunit (EC 6.3.5.2) | 10998175 | 100 | 100 | 93 | 1e-287 |
| Streptococcus dysgalactiae subsp. equisimilis ATCC 12394 | guaA | GMP synthase [glutamine-hydrolyzing], amidotransferase subunit (EC 6.3.5.2) / GMP synthase [glutamine-hydrolyzing], ATP pyrophosphatase subunit (EC 6.3.5.2) | 10998175 | 100 | 100 | 93 | 1e-286 |
| Streptococcus dysgalactiae subsp. equisimilis GGS_124 | guaA | GMP synthase [glutamine-hydrolyzing], amidotransferase subunit (EC 6.3.5.2) / GMP synthase [glutamine-hydrolyzing], ATP pyrophosphatase subunit (EC 6.3.5.2) | 10998175 | 100 | 100 | 93 | 1e-286 |
| Streptococcus dysgalactiae subsp. equisimilis RE378 | guaA | GMP synthase [glutamine-hydrolyzing], amidotransferase subunit (EC 6.3.5.2) / GMP synthase [glutamine-hydrolyzing], ATP pyrophosphatase subunit (EC 6.3.5.2) | 10998175 | 100 | 100 | 93 | 1e-287 |
| Streptococcus dysgalactiae subsp. equisimilis SD SCDR1 | guaA | GMP synthase [glutamine-hydrolyzing], amidotransferase subunit (EC 6.3.5.2) / GMP synthase [glutamine-hydrolyzing], ATP pyrophosphatase subunit (EC 6.3.5.2) | 10998175 | 100 | 100 | 93 | 1e-287 |
| Streptococcus dysgalactiae subsp. equisimilis SK1249 | guaA | GMP synthase [glutamine-hydrolyzing], amidotransferase subunit (EC 6.3.5.2) / GMP synthase [glutamine-hydrolyzing], ATP pyrophosphatase subunit (EC 6.3.5.2) | 10998175 | 100 | 100 | 93 | 1e-287 |
| Streptococcus dysgalactiae subsp. equisimilis SK1250 | guaA | GMP synthase [glutamine-hydrolyzing], amidotransferase subunit (EC 6.3.5.2) / GMP synthase [glutamine-hydrolyzing], ATP pyrophosphatase subunit (EC 6.3.5.2) | 10998175 | 100 | 100 | 92 | 1e-287 |
| Streptococcus dysgalactiae subsp. equisimilis strain ASDSE_96 | guaA | GMP synthase [glutamine-hydrolyzing], amidotransferase subunit (EC 6.3.5.2) / GMP synthase [glutamine-hydrolyzing], ATP pyrophosphatase subunit (EC 6.3.5.2) | 10998175 | 100 | 100 | 92 | 1e-284 |
| Streptococcus dysgalactiae subsp. equisimilis strain ASDSE_99 | guaA | GMP synthase [glutamine-hydrolyzing], amidotransferase subunit (EC 6.3.5.2) / GMP synthase [glutamine-hydrolyzing], ATP pyrophosphatase subunit (EC 6.3.5.2) | 10998175 | 100 | 100 | 93 | 1e-288 |
| Streptococcus dysgalactiae subsp. equisimilis strain C161L1 | guaA | GMP synthase [glutamine-hydrolyzing], amidotransferase subunit (EC 6.3.5.2) / GMP synthase [glutamine-hydrolyzing], ATP pyrophosphatase subunit (EC 6.3.5.2) | 10998175 | 100 | 100 | 93 | 0 |
| Streptococcus dysgalactiae subsp. equisimilis strain KNZ01 | guaA | GMP synthase [glutamine-hydrolyzing], amidotransferase subunit (EC 6.3.5.2) / GMP synthase [glutamine-hydrolyzing], ATP pyrophosphatase subunit (EC 6.3.5.2) | 10998175 | 100 | 100 | 93 | 1e-287 |
| Streptococcus dysgalactiae subsp. equisimilis strain KNZ03 | guaA | GMP synthase [glutamine-hydrolyzing], amidotransferase subunit (EC 6.3.5.2) / GMP synthase [glutamine-hydrolyzing], ATP pyrophosphatase subunit (EC 6.3.5.2) | 10998175 | 100 | 100 | 93 | 1e-286 |
| Streptococcus dysgalactiae subsp. equisimilis strain KNZ04 | guaA | GMP synthase [glutamine-hydrolyzing], amidotransferase subunit (EC 6.3.5.2) / GMP synthase [glutamine-hydrolyzing], ATP pyrophosphatase subunit (EC 6.3.5.2) | 10998175 | 100 | 100 | 93 | 1e-286 |
| Streptococcus dysgalactiae subsp. equisimilis strain KNZ06 | guaA | GMP synthase [glutamine-hydrolyzing], amidotransferase subunit (EC 6.3.5.2) / GMP synthase [glutamine-hydrolyzing], ATP pyrophosphatase subunit (EC 6.3.5.2) | 10998175 | 100 | 100 | 93 | 1e-287 |
| Streptococcus dysgalactiae subsp. equisimilis strain KNZ07 | guaA | GMP synthase [glutamine-hydrolyzing], amidotransferase subunit (EC 6.3.5.2) / GMP synthase [glutamine-hydrolyzing], ATP pyrophosphatase subunit (EC 6.3.5.2) | 10998175 | 100 | 100 | 93 | 1e-287 |
| Streptococcus dysgalactiae subsp. equisimilis strain KNZ10 | guaA | GMP synthase [glutamine-hydrolyzing], amidotransferase subunit (EC 6.3.5.2) / GMP synthase [glutamine-hydrolyzing], ATP pyrophosphatase subunit (EC 6.3.5.2) | 10998175 | 100 | 100 | 93 | 1e-287 |
| Streptococcus dysgalactiae subsp. equisimilis strain KNZ12 | guaA | GMP synthase [glutamine-hydrolyzing], amidotransferase subunit (EC 6.3.5.2) / GMP synthase [glutamine-hydrolyzing], ATP pyrophosphatase subunit (EC 6.3.5.2) | 10998175 | 100 | 100 | 93 | 1e-287 |
| Streptococcus dysgalactiae subsp. equisimilis strain KNZ15 | guaA | GMP synthase [glutamine-hydrolyzing], amidotransferase subunit (EC 6.3.5.2) / GMP synthase [glutamine-hydrolyzing], ATP pyrophosphatase subunit (EC 6.3.5.2) | 10998175 | 100 | 100 | 93 | 1e-287 |
| Streptococcus dysgalactiae subsp. equisimilis strain KNZ16 | guaA | GMP synthase [glutamine-hydrolyzing], amidotransferase subunit (EC 6.3.5.2) / GMP synthase [glutamine-hydrolyzing], ATP pyrophosphatase subunit (EC 6.3.5.2) | 10998175 | 100 | 100 | 93 | 1e-286 |
| Streptococcus dysgalactiae subsp. equisimilis strain NCTC10321 | guaA | GMP synthase [glutamine-hydrolyzing], amidotransferase subunit (EC 6.3.5.2) / GMP synthase [glutamine-hydrolyzing], ATP pyrophosphatase subunit (EC 6.3.5.2) | 10998175 | 100 | 100 | 93 | 1e-288 |
| Streptococcus dysgalactiae subsp. equisimilis strain NCTC11554 | guaA | GMP synthase [glutamine-hydrolyzing], amidotransferase subunit (EC 6.3.5.2) / GMP synthase [glutamine-hydrolyzing], ATP pyrophosphatase subunit (EC 6.3.5.2) | 10998175 | 100 | 100 | 93 | 1e-286 |
| Streptococcus dysgalactiae subsp. equisimilis strain NCTC11555 | guaA | GMP synthase [glutamine-hydrolyzing], amidotransferase subunit (EC 6.3.5.2) / GMP synthase [glutamine-hydrolyzing], ATP pyrophosphatase subunit (EC 6.3.5.2) | 10998175 | 100 | 100 | 93 | 1e-288 |
| Streptococcus dysgalactiae subsp. equisimilis strain NCTC11556 | guaA | GMP synthase [glutamine-hydrolyzing], amidotransferase subunit (EC 6.3.5.2) / GMP synthase [glutamine-hydrolyzing], ATP pyrophosphatase subunit (EC 6.3.5.2) | 10998175 | 100 | 100 | 93 | 1e-286 |
| Streptococcus dysgalactiae subsp. equisimilis strain NCTC11557 | guaA | GMP synthase [glutamine-hydrolyzing], amidotransferase subunit (EC 6.3.5.2) / GMP synthase [glutamine-hydrolyzing], ATP pyrophosphatase subunit (EC 6.3.5.2) | 10998175 | 100 | 100 | 93 | 1e-288 |
| Streptococcus dysgalactiae subsp. equisimilis strain NCTC11564 | guaA | GMP synthase [glutamine-hydrolyzing], amidotransferase subunit (EC 6.3.5.2) / GMP synthase [glutamine-hydrolyzing], ATP pyrophosphatase subunit (EC 6.3.5.2) | 10998175 | 100 | 100 | 93 | 1e-287 |
| Streptococcus dysgalactiae subsp. equisimilis strain NCTC5370 | guaA | GMP synthase [glutamine-hydrolyzing], amidotransferase subunit (EC 6.3.5.2) / GMP synthase [glutamine-hydrolyzing], ATP pyrophosphatase subunit (EC 6.3.5.2) | 10998175 | 100 | 100 | 93 | 1e-287 |
| Streptococcus dysgalactiae subsp. equisimilis strain NCTC5371 | guaA | GMP synthase [glutamine-hydrolyzing], amidotransferase subunit (EC 6.3.5.2) / GMP synthase [glutamine-hydrolyzing], ATP pyrophosphatase subunit (EC 6.3.5.2) | 10998175 | 100 | 100 | 93 | 1e-287 |
| Streptococcus dysgalactiae subsp. equisimilis strain NCTC6179 | guaA | GMP synthase [glutamine-hydrolyzing], amidotransferase subunit (EC 6.3.5.2) / GMP synthase [glutamine-hydrolyzing], ATP pyrophosphatase subunit (EC 6.3.5.2) | 10998175 | 100 | 100 | 93 | 1e-287 |
| Streptococcus dysgalactiae subsp. equisimilis strain NCTC6181 | guaA | GMP synthase [glutamine-hydrolyzing], amidotransferase subunit (EC 6.3.5.2) / GMP synthase [glutamine-hydrolyzing], ATP pyrophosphatase subunit (EC 6.3.5.2) | 10998175 | 100 | 100 | 93 | 1e-287 |
| Streptococcus dysgalactiae subsp. equisimilis strain NCTC6407 | guaA | GMP synthase [glutamine-hydrolyzing], amidotransferase subunit (EC 6.3.5.2) / GMP synthase [glutamine-hydrolyzing], ATP pyrophosphatase subunit (EC 6.3.5.2) | 10998175 | 100 | 100 | 93 | 1e-288 |
| Streptococcus dysgalactiae subsp. equisimilis strain NCTC7136 | guaA | GMP synthase [glutamine-hydrolyzing], amidotransferase subunit (EC 6.3.5.2) / GMP synthase [glutamine-hydrolyzing], ATP pyrophosphatase subunit (EC 6.3.5.2) | 10998175 | 100 | 100 | 93 | 1e-287 |
| Streptococcus dysgalactiae subsp. equisimilis strain NCTC8543 | guaA | GMP synthase [glutamine-hydrolyzing], amidotransferase subunit (EC 6.3.5.2) / GMP synthase [glutamine-hydrolyzing], ATP pyrophosphatase subunit (EC 6.3.5.2) | 10998175 | 100 | 100 | 93 | 1e-286 |
| Streptococcus dysgalactiae subsp. equisimilis strain NCTC8546 | guaA | GMP synthase [glutamine-hydrolyzing], amidotransferase subunit (EC 6.3.5.2) / GMP synthase [glutamine-hydrolyzing], ATP pyrophosphatase subunit (EC 6.3.5.2) | 10998175 | 100 | 100 | 93 | 1e-287 |
| Streptococcus dysgalactiae subsp. equisimilis strain NCTC9413 | guaA | GMP synthase [glutamine-hydrolyzing], amidotransferase subunit (EC 6.3.5.2) / GMP synthase [glutamine-hydrolyzing], ATP pyrophosphatase subunit (EC 6.3.5.2) | 10998175 | 100 | 100 | 93 | 1e-287 |
| Streptococcus dysgalactiae subsp. equisimilis strain NCTC9414 | guaA | GMP synthase [glutamine-hydrolyzing], amidotransferase subunit (EC 6.3.5.2) / GMP synthase [glutamine-hydrolyzing], ATP pyrophosphatase subunit (EC 6.3.5.2) | 10998175 | 100 | 100 | 93 | 1e-287 |
| Streptococcus dysgalactiae subsp. equisimilis strain NCTC9603 | guaA | GMP synthase [glutamine-hydrolyzing], amidotransferase subunit (EC 6.3.5.2) / GMP synthase [glutamine-hydrolyzing], ATP pyrophosphatase subunit (EC 6.3.5.2) | 10998175 | 100 | 100 | 93 | 1e-287 |
| Streptococcus dysgalactiae subsp. equisimilis strain SS1575 | guaA | GMP synthase [glutamine-hydrolyzing], amidotransferase subunit (EC 6.3.5.2) / GMP synthase [glutamine-hydrolyzing], ATP pyrophosphatase subunit (EC 6.3.5.2) | 10998175 | 100 | 100 | 93 | 1e-287 |
| Streptococcus dysgalactiae subsp. equisimilis strain T642 | guaA | GMP synthase [glutamine-hydrolyzing], amidotransferase subunit (EC 6.3.5.2) / GMP synthase [glutamine-hydrolyzing], ATP pyrophosphatase subunit (EC 6.3.5.2) | 10998175 | 100 | 100 | 93 | 0 |
| Streptococcus dysgalactiae subsp. equisimilis strain UT_4031CC | guaA | GMP synthase [glutamine-hydrolyzing], amidotransferase subunit (EC 6.3.5.2) / GMP synthase [glutamine-hydrolyzing], ATP pyrophosphatase subunit (EC 6.3.5.2) | 10998175 | 100 | 100 | 93 | 1e-287 |
| Streptococcus dysgalactiae subsp. equisimilis strain UT_4231_KK | guaA | GMP synthase [glutamine-hydrolyzing], amidotransferase subunit (EC 6.3.5.2) / GMP synthase [glutamine-hydrolyzing], ATP pyrophosphatase subunit (EC 6.3.5.2) | 10998175 | 100 | 100 | 93 | 1e-287 |
| Streptococcus dysgalactiae subsp. equisimilis strain UT_4234_DH | guaA | GMP synthase [glutamine-hydrolyzing], amidotransferase subunit (EC 6.3.5.2) / GMP synthase [glutamine-hydrolyzing], ATP pyrophosphatase subunit (EC 6.3.5.2) | 10998175 | 100 | 100 | 93 | 1e-288 |
| Streptococcus dysgalactiae subsp. equisimilis strain UT_4241_XS | guaA | GMP synthase [glutamine-hydrolyzing], amidotransferase subunit (EC 6.3.5.2) / GMP synthase [glutamine-hydrolyzing], ATP pyrophosphatase subunit (EC 6.3.5.2) | 10998175 | 100 | 100 | 93 | 1e-287 |
| Streptococcus dysgalactiae subsp. equisimilis strain UT_4242_AB | guaA | GMP synthase [glutamine-hydrolyzing], amidotransferase subunit (EC 6.3.5.2) / GMP synthase [glutamine-hydrolyzing], ATP pyrophosphatase subunit (EC 6.3.5.2) | 10998175 | 100 | 100 | 93 | 1e-286 |
| Streptococcus dysgalactiae subsp. equisimilis strain UT_4255RC | guaA | GMP synthase [glutamine-hydrolyzing], amidotransferase subunit (EC 6.3.5.2) / GMP synthase [glutamine-hydrolyzing], ATP pyrophosphatase subunit (EC 6.3.5.2) | 10998175 | 100 | 100 | 93 | 1e-288 |
| Streptococcus dysgalactiae subsp. equisimilis strain UT_4277_BB | guaA | GMP synthase [glutamine-hydrolyzing], amidotransferase subunit (EC 6.3.5.2) / GMP synthase [glutamine-hydrolyzing], ATP pyrophosphatase subunit (EC 6.3.5.2) | 10998175 | 100 | 100 | 93 | 1e-287 |
| Streptococcus dysgalactiae subsp. equisimilis strain UT_4966_RC | guaA | GMP synthase [glutamine-hydrolyzing], amidotransferase subunit (EC 6.3.5.2) / GMP synthase [glutamine-hydrolyzing], ATP pyrophosphatase subunit (EC 6.3.5.2) | 10998175 | 100 | 100 | 92 | 1e-286 |
| Streptococcus dysgalactiae subsp. equisimilis strain UT-5345 | guaA | GMP synthase [glutamine-hydrolyzing], amidotransferase subunit (EC 6.3.5.2) / GMP synthase [glutamine-hydrolyzing], ATP pyrophosphatase subunit (EC 6.3.5.2) | 10998175 | 100 | 100 | 93 | 1e-287 |
| Streptococcus dysgalactiae subsp. equisimilis strain UT-5354 | guaA | GMP synthase [glutamine-hydrolyzing], amidotransferase subunit (EC 6.3.5.2) / GMP synthase [glutamine-hydrolyzing], ATP pyrophosphatase subunit (EC 6.3.5.2) | 10998175 | 100 | 100 | 93 | 1e-287 |
| Streptococcus dysgalactiae subsp. equisimilis strain UT-SS1069 | guaA | GMP synthase [glutamine-hydrolyzing], amidotransferase subunit (EC 6.3.5.2) / GMP synthase [glutamine-hydrolyzing], ATP pyrophosphatase subunit (EC 6.3.5.2) | 10998175 | 100 | 100 | 93 | 1e-287 |
| Streptococcus dysgalactiae subsp. equisimilis strain UT-SS957 | guaA | GMP synthase [glutamine-hydrolyzing], amidotransferase subunit (EC 6.3.5.2) / GMP synthase [glutamine-hydrolyzing], ATP pyrophosphatase subunit (EC 6.3.5.2) | 10998175 | 100 | 100 | 93 | 1e-287 |
| Streptococcus dysgalactiae subsp. equisimilis strain WCHSDSE-1 | guaA | GMP synthase [glutamine-hydrolyzing], amidotransferase subunit (EC 6.3.5.2) / GMP synthase [glutamine-hydrolyzing], ATP pyrophosphatase subunit (EC 6.3.5.2) | 10998175 | 100 | 100 | 93 | 1e-287 |
| Streptococcus dysgalactiae subsp. equisimilis 167 | gbs0632 | hypothetical protein | 19086816 | 99 | 100 | 94 | 0.0 |
| Streptococcus dysgalactiae subsp. equisimilis AC-2713 | gbs0632 | hypothetical protein | 19086816 | 99 | 100 | 93 | 0.0 |
| Streptococcus dysgalactiae subsp. equisimilis RE378 | gbs0632 | hypothetical protein | 19086816 | 99 | 100 | 94 | 0.0 |
| Streptococcus dysgalactiae subsp. equisimilis SD SCDR1 | gbs0632 | hypothetical protein | 19086816 | 99 | 100 | 95 | 0.0 |
| Streptococcus dysgalactiae subsp. equisimilis SK1250 | gbs0632 | hypothetical protein | 19086816 | 73 | 100 | 95 | 0.0 |
| Streptococcus dysgalactiae subsp. equisimilis strain ASDSE_96 | gbs0632 | hypothetical protein | 19086816 | 100 | 100 | 93 | 0.0 |
| Streptococcus dysgalactiae subsp. equisimilis strain ASDSE_99 | gbs0632 | hypothetical protein | 19086816 | 99 | 99 | 93 | 0.0 |
| Streptococcus dysgalactiae subsp. equisimilis strain KNZ01 | gbs0632 | hypothetical protein | 19086816 | 99 | 100 | 94 | 0.0 |
| Streptococcus dysgalactiae subsp. equisimilis strain KNZ03 | gbs0632 | hypothetical protein | 19086816 | 99 | 100 | 92 | 0.0 |
| Streptococcus dysgalactiae subsp. equisimilis strain KNZ04 | gbs0632 | hypothetical protein | 19086816 | 99 | 100 | 93 | 0.0 |
| Streptococcus dysgalactiae subsp. equisimilis strain KNZ06 | gbs0632 | hypothetical protein | 19086816 | 99 | 99 | 93 | 0.0 |
| Streptococcus dysgalactiae subsp. equisimilis strain KNZ07 | gbs0632 | hypothetical protein | 19086816 | 99 | 100 | 94 | 0.0 |
| Streptococcus dysgalactiae subsp. equisimilis strain KNZ10 | gbs0632 | hypothetical protein | 19086816 | 99 | 100 | 94 | 0.0 |
| Streptococcus dysgalactiae subsp. equisimilis strain KNZ12 | gbs0632 | hypothetical protein | 19086816 | 99 | 100 | 94 | 0.0 |
| Streptococcus dysgalactiae subsp. equisimilis strain KNZ15 | gbs0632 | hypothetical protein | 19086816 | 99 | 100 | 94 | 0.0 |
| Streptococcus dysgalactiae subsp. equisimilis strain NCTC10321 | gbs0632 | hypothetical protein | 19086816 | 99 | 99 | 93 | 0.0 |
| Streptococcus dysgalactiae subsp. equisimilis strain NCTC11555 | gbs0632 | hypothetical protein | 19086816 | 99 | 99 | 93 | 0.0 |
| Streptococcus dysgalactiae subsp. equisimilis strain NCTC11556 | gbs0632 | hypothetical protein | 19086816 | 100 | 100 | 94 | 0.0 |
| Streptococcus dysgalactiae subsp. equisimilis strain NCTC11557 | gbs0632 | hypothetical protein | 19086816 | 100 | 100 | 94 | 0.0 |
| Streptococcus dysgalactiae subsp. equisimilis strain NCTC11564 | gbs0632 | hypothetical protein | 19086816 | 99 | 100 | 93 | 0.0 |
| Streptococcus dysgalactiae subsp. equisimilis strain NCTC11565 | tse2 | hypothetical protein | 19086816 | 100 | 100 | 97 | 1e-86 |
| Streptococcus dysgalactiae subsp. equisimilis strain NCTC5371 | gbs0632 | hypothetical protein | 19086816 | 99 | 100 | 93 | 0.0 |
| Streptococcus dysgalactiae subsp. equisimilis strain NCTC5969 | SA1262 | hypothetical protein | 20347953 | 100 | 100 | 100 | 1e-218 |
| Streptococcus dysgalactiae subsp. equisimilis strain NCTC6407 | gbs0632 | hypothetical protein | 20347953 | 99 | 99 | 93 | 0.0 |
| Streptococcus dysgalactiae subsp. equisimilis strain NCTC7136 | gbs0632 | hypothetical protein | 20347953 | 99 | 100 | 95 | 0.0 |
| Streptococcus dysgalactiae subsp. equisimilis strain NCTC8546 | gbs0632 | hypothetical protein | 20347953 | 99 | 100 | 92 | 0.0 |
| Streptococcus dysgalactiae subsp. equisimilis strain NCTC9414 | gbs0632 | hypothetical protein | 20347953 | 99 | 100 | 93 | 0.0 |
| Streptococcus dysgalactiae subsp. equisimilis strain UT_4031CC | gbs0632 | hypothetical protein | 20347953 | 99 | 100 | 92 | 0.0 |
| Streptococcus dysgalactiae subsp. equisimilis strain UT_4231_KK | gbs0632 | hypothetical protein | 20347953 | 99 | 100 | 92 | 0.0 |
| Streptococcus dysgalactiae subsp. equisimilis strain UT_4234_DH | gbs0632 | hypothetical protein | 20347953 | 99 | 99 | 93 | 0.0 |
| Streptococcus dysgalactiae subsp. equisimilis strain UT_4241_XS | gbs0632 | hypothetical protein | 20347953 | 99 | 100 | 94 | 0.0 |
| Streptococcus dysgalactiae subsp. equisimilis strain UT_4242_AB | gbs0632 | hypothetical protein | 20347953 | 100 | 100 | 94 | 0.0 |
| Streptococcus dysgalactiae subsp. equisimilis strain UT_4255RC | gbs0632 | hypothetical protein | 20347953 | 100 | 100 | 94 | 0.0 |
| Streptococcus dysgalactiae subsp. equisimilis strain UT_4277_BB | gbs0632 | hypothetical protein | 20347953 | 100 | 100 | 94 | 0.0 |
| Streptococcus dysgalactiae subsp. equisimilis strain UT_4966_RC | gbs0632 | hypothetical protein | 20347953 | 99 | 100 | 94 | 0.0 |
| Streptococcus dysgalactiae subsp. equisimilis strain UT-5345 | gbs0632 | hypothetical protein | 20347953 | 99 | 100 | 93 | 0.0 |
| Streptococcus dysgalactiae subsp. equisimilis strain UT-SS1069 | gbs0632 | hypothetical protein | 20347953 | 99 | 100 | 94 | 0.0 |
| Streptococcus dysgalactiae subsp. equisimilis strain UT-SS957 | gbs0632 | hypothetical protein | 20347953 | 99 | 100 | 93 | 0.0 |
| Streptococcus dysgalactiae subsp. equisimilis 167 | purH | IMP cyclohydrolase (EC 3.5.4.10) / Phosphoribosylaminoimidazolecarboxamide formyltransferase (EC 2.1.2.3) | 12207705 | 99 | 99 | 84 | 1e-258 |
| Streptococcus dysgalactiae subsp. equisimilis AC-2713 | purH | IMP cyclohydrolase (EC 3.5.4.10) / Phosphoribosylaminoimidazolecarboxamide formyltransferase (EC 2.1.2.3) | 12207705 | 99 | 99 | 84 | 1e-257 |
| Streptococcus dysgalactiae subsp. equisimilis AKSDE4288 | purH | IMP cyclohydrolase (EC 3.5.4.10) / Phosphoribosylaminoimidazolecarboxamide formyltransferase (EC 2.1.2.3) | 12207705 | 99 | 99 | 84 | 1e-257 |
| Streptococcus dysgalactiae subsp. equisimilis ATCC 12394 | purH | IMP cyclohydrolase (EC 3.5.4.10) / Phosphoribosylaminoimidazolecarboxamide formyltransferase (EC 2.1.2.3) | 12207705 | 99 | 99 | 84 | 1e-258 |
| Streptococcus dysgalactiae subsp. equisimilis GGS_124 | purH | IMP cyclohydrolase (EC 3.5.4.10) / Phosphoribosylaminoimidazolecarboxamide formyltransferase (EC 2.1.2.3) | 12207705 | 99 | 99 | 84 | 1e-257 |
| Streptococcus dysgalactiae subsp. equisimilis RE378 | purH | IMP cyclohydrolase (EC 3.5.4.10) / Phosphoribosylaminoimidazolecarboxamide formyltransferase (EC 2.1.2.3) | 12207705 | 99 | 99 | 84 | 1e-257 |
| Streptococcus dysgalactiae subsp. equisimilis SD SCDR1 | purH | IMP cyclohydrolase (EC 3.5.4.10) / Phosphoribosylaminoimidazolecarboxamide formyltransferase (EC 2.1.2.3) | 12207705 | 99 | 99 | 84 | 1e-258 |
| Streptococcus dysgalactiae subsp. equisimilis SK1249 | purH | IMP cyclohydrolase (EC 3.5.4.10) / Phosphoribosylaminoimidazolecarboxamide formyltransferase (EC 2.1.2.3) | 12207705 | 99 | 99 | 84 | 1e-257 |
| Streptococcus dysgalactiae subsp. equisimilis SK1250 | purH | IMP cyclohydrolase (EC 3.5.4.10) / Phosphoribosylaminoimidazolecarboxamide formyltransferase (EC 2.1.2.3) | 12207705 | 99 | 99 | 84 | 1e-258 |
| Streptococcus dysgalactiae subsp. equisimilis strain ASDSE_96 | purH | IMP cyclohydrolase (EC 3.5.4.10) / Phosphoribosylaminoimidazolecarboxamide formyltransferase (EC 2.1.2.3) | 12207705 | 99 | 99 | 84 | 1e-258 |
| Streptococcus dysgalactiae subsp. equisimilis strain ASDSE_99 | purH | IMP cyclohydrolase (EC 3.5.4.10) / Phosphoribosylaminoimidazolecarboxamide formyltransferase (EC 2.1.2.3) | 12207705 | 99 | 99 | 84 | 1e-257 |
| Streptococcus dysgalactiae subsp. equisimilis strain C161L1 | purH | IMP cyclohydrolase (EC 3.5.4.10) / Phosphoribosylaminoimidazolecarboxamide formyltransferase (EC 2.1.2.3) | 12207705 | 99 | 99 | 84 | 0 |
| Streptococcus dysgalactiae subsp. equisimilis strain KNZ01 | purH | IMP cyclohydrolase (EC 3.5.4.10) / Phosphoribosylaminoimidazolecarboxamide formyltransferase (EC 2.1.2.3) | 12207705 | 99 | 99 | 84 | 1e-257 |
| Streptococcus dysgalactiae subsp. equisimilis strain KNZ03 | purH | IMP cyclohydrolase (EC 3.5.4.10) / Phosphoribosylaminoimidazolecarboxamide formyltransferase (EC 2.1.2.3) | 12207705 | 99 | 99 | 84 | 1e-258 |
| Streptococcus dysgalactiae subsp. equisimilis strain KNZ04 | purH | IMP cyclohydrolase (EC 3.5.4.10) / Phosphoribosylaminoimidazolecarboxamide formyltransferase (EC 2.1.2.3) | 12207705 | 99 | 99 | 84 | 1e-258 |
| Streptococcus dysgalactiae subsp. equisimilis strain KNZ06 | purH | IMP cyclohydrolase (EC 3.5.4.10) / Phosphoribosylaminoimidazolecarboxamide formyltransferase (EC 2.1.2.3) | 12207705 | 99 | 99 | 84 | 1e-258 |
| Streptococcus dysgalactiae subsp. equisimilis strain KNZ07 | purH | IMP cyclohydrolase (EC 3.5.4.10) / Phosphoribosylaminoimidazolecarboxamide formyltransferase (EC 2.1.2.3) | 12207705 | 99 | 99 | 84 | 1e-257 |
| Streptococcus dysgalactiae subsp. equisimilis strain KNZ10 | purH | IMP cyclohydrolase (EC 3.5.4.10) / Phosphoribosylaminoimidazolecarboxamide formyltransferase (EC 2.1.2.3) | 12207705 | 99 | 99 | 84 | 1e-257 |
| Streptococcus dysgalactiae subsp. equisimilis strain KNZ12 | purH | IMP cyclohydrolase (EC 3.5.4.10) / Phosphoribosylaminoimidazolecarboxamide formyltransferase (EC 2.1.2.3) | 12207705 | 99 | 99 | 84 | 1e-257 |
| Streptococcus dysgalactiae subsp. equisimilis strain KNZ15 | purH | IMP cyclohydrolase (EC 3.5.4.10) / Phosphoribosylaminoimidazolecarboxamide formyltransferase (EC 2.1.2.3) | 12207705 | 99 | 99 | 84 | 1e-257 |
| Streptococcus dysgalactiae subsp. equisimilis strain KNZ16 | purH | IMP cyclohydrolase (EC 3.5.4.10) / Phosphoribosylaminoimidazolecarboxamide formyltransferase (EC 2.1.2.3) | 12207705 | 99 | 99 | 84 | 1e-258 |
| Streptococcus dysgalactiae subsp. equisimilis strain NCTC10321 | purH | IMP cyclohydrolase (EC 3.5.4.10) / Phosphoribosylaminoimidazolecarboxamide formyltransferase (EC 2.1.2.3) | 12207705 | 99 | 99 | 84 | 1e-258 |
| Streptococcus dysgalactiae subsp. equisimilis strain NCTC11554 | purH | IMP cyclohydrolase (EC 3.5.4.10) / Phosphoribosylaminoimidazolecarboxamide formyltransferase (EC 2.1.2.3) | 12207705 | 99 | 99 | 84 | 1e-258 |
| Streptococcus dysgalactiae subsp. equisimilis strain NCTC11555 | purH | IMP cyclohydrolase (EC 3.5.4.10) / Phosphoribosylaminoimidazolecarboxamide formyltransferase (EC 2.1.2.3) | 12207705 | 99 | 99 | 84 | 1e-258 |
| Streptococcus dysgalactiae subsp. equisimilis strain NCTC11556 | purH | IMP cyclohydrolase (EC 3.5.4.10) / Phosphoribosylaminoimidazolecarboxamide formyltransferase (EC 2.1.2.3) | 12207705 | 99 | 99 | 84 | 1e-258 |
| Streptococcus dysgalactiae subsp. equisimilis strain NCTC11557 | purH | IMP cyclohydrolase (EC 3.5.4.10) / Phosphoribosylaminoimidazolecarboxamide formyltransferase (EC 2.1.2.3) | 12207705 | 99 | 99 | 84 | 1e-257 |
| Streptococcus dysgalactiae subsp. equisimilis strain NCTC11564 | purH | IMP cyclohydrolase (EC 3.5.4.10) / Phosphoribosylaminoimidazolecarboxamide formyltransferase (EC 2.1.2.3) | 12207705 | 99 | 99 | 84 | 1e-258 |
| Streptococcus dysgalactiae subsp. equisimilis strain NCTC11565 | purH | IMP cyclohydrolase (EC 3.5.4.10) / Phosphoribosylaminoimidazolecarboxamide formyltransferase (EC 2.1.2.3) | 14641575 | 100 | 100 | 99 | 1e-308 |
| Streptococcus dysgalactiae subsp. equisimilis strain NCTC5370 | purH | IMP cyclohydrolase (EC 3.5.4.10) / Phosphoribosylaminoimidazolecarboxamide formyltransferase (EC 2.1.2.3) | 12207705 | 99 | 99 | 84 | 1e-258 |
| Streptococcus dysgalactiae subsp. equisimilis strain NCTC5371 | purH | IMP cyclohydrolase (EC 3.5.4.10) / Phosphoribosylaminoimidazolecarboxamide formyltransferase (EC 2.1.2.3) | 12207705 | 99 | 99 | 84 | 1e-258 |
| Streptococcus dysgalactiae subsp. equisimilis strain NCTC6179 | purH | IMP cyclohydrolase (EC 3.5.4.10) / Phosphoribosylaminoimidazolecarboxamide formyltransferase (EC 2.1.2.3) | 12207705 | 99 | 99 | 85 | 1e-258 |
| Streptococcus dysgalactiae subsp. equisimilis strain NCTC6181 | purH | IMP cyclohydrolase (EC 3.5.4.10) / Phosphoribosylaminoimidazolecarboxamide formyltransferase (EC 2.1.2.3) | 12207705 | 99 | 99 | 85 | 1e-258 |
| Streptococcus dysgalactiae subsp. equisimilis strain NCTC6407 | purH | IMP cyclohydrolase (EC 3.5.4.10) / Phosphoribosylaminoimidazolecarboxamide formyltransferase (EC 2.1.2.3) | 12207705 | 99 | 99 | 84 | 1e-258 |
| Streptococcus dysgalactiae subsp. equisimilis strain NCTC7136 | purH | IMP cyclohydrolase (EC 3.5.4.10) / Phosphoribosylaminoimidazolecarboxamide formyltransferase (EC 2.1.2.3) | 12207705 | 99 | 99 | 84 | 1e-258 |
| Streptococcus dysgalactiae subsp. equisimilis strain NCTC8543 | purH | IMP cyclohydrolase (EC 3.5.4.10) / Phosphoribosylaminoimidazolecarboxamide formyltransferase (EC 2.1.2.3) | 12207705 | 99 | 99 | 84 | 1e-257 |
| Streptococcus dysgalactiae subsp. equisimilis strain NCTC8546 | purH | IMP cyclohydrolase (EC 3.5.4.10) / Phosphoribosylaminoimidazolecarboxamide formyltransferase (EC 2.1.2.3) | 12207705 | 99 | 99 | 84 | 1e-258 |
| Streptococcus dysgalactiae subsp. equisimilis strain NCTC9413 | purH | IMP cyclohydrolase (EC 3.5.4.10) / Phosphoribosylaminoimidazolecarboxamide formyltransferase (EC 2.1.2.3) | 12207705 | 99 | 99 | 84 | 1e-258 |
| Streptococcus dysgalactiae subsp. equisimilis strain NCTC9414 | purH | IMP cyclohydrolase (EC 3.5.4.10) / Phosphoribosylaminoimidazolecarboxamide formyltransferase (EC 2.1.2.3) | 12207705 | 99 | 99 | 84 | 1e-258 |
| Streptococcus dysgalactiae subsp. equisimilis strain NCTC9603 | purH | IMP cyclohydrolase (EC 3.5.4.10) / Phosphoribosylaminoimidazolecarboxamide formyltransferase (EC 2.1.2.3) | 12207705 | 99 | 99 | 84 | 1e-258 |
| Streptococcus dysgalactiae subsp. equisimilis strain SS1575 | purH | IMP cyclohydrolase (EC 3.5.4.10) / Phosphoribosylaminoimidazolecarboxamide formyltransferase (EC 2.1.2.3) | 12207705 | 99 | 99 | 84 | 1e-258 |
| Streptococcus dysgalactiae subsp. equisimilis strain T642 | purH | IMP cyclohydrolase (EC 3.5.4.10) / Phosphoribosylaminoimidazolecarboxamide formyltransferase (EC 2.1.2.3) | 12207705 | 99 | 99 | 84 | 0.0 |
| Streptococcus dysgalactiae subsp. equisimilis strain UT_4031CC | purH | IMP cyclohydrolase (EC 3.5.4.10) / Phosphoribosylaminoimidazolecarboxamide formyltransferase (EC 2.1.2.3) | 12207705 | 99 | 99 | 84 | 1e-258 |
| Streptococcus dysgalactiae subsp. equisimilis strain UT_4231_KK | purH | IMP cyclohydrolase (EC 3.5.4.10) / Phosphoribosylaminoimidazolecarboxamide formyltransferase (EC 2.1.2.3) | 12207705 | 99 | 99 | 84 | 1e-258 |
| Streptococcus dysgalactiae subsp. equisimilis strain UT_4234_DH | purH | IMP cyclohydrolase (EC 3.5.4.10) / Phosphoribosylaminoimidazolecarboxamide formyltransferase (EC 2.1.2.3) | 12207705 | 99 | 99 | 84 | 1e-257 |
| Streptococcus dysgalactiae subsp. equisimilis strain UT_4241_XS | purH | IMP cyclohydrolase (EC 3.5.4.10) / Phosphoribosylaminoimidazolecarboxamide formyltransferase (EC 2.1.2.3) | 12207705 | 99 | 99 | 84 | 1e-258 |
| Streptococcus dysgalactiae subsp. equisimilis strain UT_4242_AB | purH | IMP cyclohydrolase (EC 3.5.4.10) / Phosphoribosylaminoimidazolecarboxamide formyltransferase (EC 2.1.2.3) | 12207705 | 99 | 99 | 84 | 1e-258 |
| Streptococcus dysgalactiae subsp. equisimilis strain UT_4255RC | purH | IMP cyclohydrolase (EC 3.5.4.10) / Phosphoribosylaminoimidazolecarboxamide formyltransferase (EC 2.1.2.3) | 12207705 | 99 | 99 | 84 | 1e-257 |
| Streptococcus dysgalactiae subsp. equisimilis strain UT_4277_BB | purH | IMP cyclohydrolase (EC 3.5.4.10) / Phosphoribosylaminoimidazolecarboxamide formyltransferase (EC 2.1.2.3) | 12207705 | 99 | 99 | 84 | 1e-258 |
| Streptococcus dysgalactiae subsp. equisimilis strain UT_4966_RC | purH | IMP cyclohydrolase (EC 3.5.4.10) / Phosphoribosylaminoimidazolecarboxamide formyltransferase (EC 2.1.2.3) | 12207705 | 99 | 99 | 84 | 1e-258 |
| Streptococcus dysgalactiae subsp. equisimilis strain UT-5345 | purH | IMP cyclohydrolase (EC 3.5.4.10) / Phosphoribosylaminoimidazolecarboxamide formyltransferase (EC 2.1.2.3) | 12207705 | 99 | 99 | 84 | 1e-258 |
| Streptococcus dysgalactiae subsp. equisimilis strain UT-5354 | purH | IMP cyclohydrolase (EC 3.5.4.10) / Phosphoribosylaminoimidazolecarboxamide formyltransferase (EC 2.1.2.3) | 12207705 | 99 | 99 | 84 | 1e-257 |
| Streptococcus dysgalactiae subsp. equisimilis strain UT-SS1069 | purH | IMP cyclohydrolase (EC 3.5.4.10) / Phosphoribosylaminoimidazolecarboxamide formyltransferase (EC 2.1.2.3) | 12207705 | 99 | 99 | 84 | 1e-258 |
| Streptococcus dysgalactiae subsp. equisimilis strain UT-SS957 | purH | IMP cyclohydrolase (EC 3.5.4.10) / Phosphoribosylaminoimidazolecarboxamide formyltransferase (EC 2.1.2.3) | 12207705 | 99 | 99 | 84 | 1e-257 |
| Streptococcus dysgalactiae subsp. equisimilis strain WCHSDSE-1 | purH | IMP cyclohydrolase (EC 3.5.4.10) / Phosphoribosylaminoimidazolecarboxamide formyltransferase (EC 2.1.2.3) | 12207705 | 99 | 99 | 84 | 1e-258 |
| Streptococcus dysgalactiae subsp. equisimilis 167 | lmb | Laminin-binding surface protein | 17400016 | 100 | 100 | 98 | 1e-172 |
| Streptococcus dysgalactiae subsp. equisimilis 167 | lmb | Laminin-binding surface protein | 17400016 | 100 | 100 | 99 | 1e-174 |
| Streptococcus dysgalactiae subsp. equisimilis AC-2713 | psaA | Laminin-binding surface protein | 17400016 | 100 | 100 | 98 | 1e-171 |
| Streptococcus dysgalactiae subsp. equisimilis AC-2713 | psaA | Laminin-binding surface protein | 17400016 | 100 | 100 | 99 | 1e-173 |
| Streptococcus dysgalactiae subsp. equisimilis AKSDE4288 | lmb | Laminin-binding surface protein | 17400016 | 100 | 100 | 99 | 1e-173 |
| Streptococcus dysgalactiae subsp. equisimilis AKSDE4288 | lmb | Laminin-binding surface protein | 17400016 | 100 | 100 | 98 | 1e-171 |
| Streptococcus dysgalactiae subsp. equisimilis ATCC 12394 | lmb | Laminin-binding surface protein | 17400016 | 100 | 100 | 98 | 1e-173 |
| Streptococcus dysgalactiae subsp. equisimilis ATCC 12394 | lmb | Laminin-binding surface protein | 17400016 | 100 | 100 | 97 | 1e-171 |
| Streptococcus dysgalactiae subsp. equisimilis GGS_124 | lmb | Laminin-binding surface protein | 17400016 | 100 | 100 | 98 | 1e-171 |
| Streptococcus dysgalactiae subsp. equisimilis GGS_124 | lmb | Laminin-binding surface protein | 17400016 | 100 | 100 | 99 | 1e-174 |
| Streptococcus dysgalactiae subsp. equisimilis RE378 | lmb | Laminin-binding surface protein | 17400016 | 100 | 100 | 99 | 1e-174 |
| Streptococcus dysgalactiae subsp. equisimilis RE378 | lmb | Laminin-binding surface protein | 17400016 | 100 | 100 | 98 | 1e-172 |
| Streptococcus dysgalactiae subsp. equisimilis SD SCDR1 | lmb | Laminin-binding surface protein | 17400016 | 100 | 100 | 99 | 1e-175 |
| Streptococcus dysgalactiae subsp. equisimilis SD SCDR1 | lmb | Laminin-binding surface protein | 17400016 | 100 | 100 | 98 | 1e-172 |
| Streptococcus dysgalactiae subsp. equisimilis SK1249 | lmb | Laminin-binding surface protein | 17400016 | 100 | 100 | 97 | 1e-171 |
| Streptococcus dysgalactiae subsp. equisimilis SK1249 | lmb | Laminin-binding surface protein | 17400016 | 100 | 100 | 98 | 1e-173 |
| Streptococcus dysgalactiae subsp. equisimilis SK1250 | lmb | Laminin-binding surface protein | 17400016 | 100 | 100 | 100 | 1e-175 |
| Streptococcus dysgalactiae subsp. equisimilis SK1250 | lmb | Laminin-binding surface protein | 17400016 | 100 | 100 | 99 | 1e-173 |
| Streptococcus dysgalactiae subsp. equisimilis strain ASDSE_96 | lmb | Laminin-binding surface protein | 17400016 | 100 | 100 | 100 | 1e-175 |
| Streptococcus dysgalactiae subsp. equisimilis strain ASDSE_96 | lmb | Laminin-binding surface protein | 17400016 | 100 | 100 | 99 | 1e-173 |
| Streptococcus dysgalactiae subsp. equisimilis strain ASDSE_99 | lmb | Laminin-binding surface protein | 17400016 | 100 | 100 | 99 | 1e-173 |
| Streptococcus dysgalactiae subsp. equisimilis strain ASDSE_99 | lmb | Laminin-binding surface protein | 17400016 | 100 | 100 | 98 | 1e-171 |
| Streptococcus dysgalactiae subsp. equisimilis strain C161L1 | lmb | Laminin-binding surface protein | 17400016 | 100 | 100 | 99 | 0.0 |
| Streptococcus dysgalactiae subsp. equisimilis strain C161L1 | lmb | Laminin-binding surface protein | 17400016 | 100 | 100 | 98 | 0.0 |
| Streptococcus dysgalactiae subsp. equisimilis strain KNZ01 | lmb | Laminin-binding surface protein | 17400016 | 100 | 100 | 99 | 1e-174 |
| Streptococcus dysgalactiae subsp. equisimilis strain KNZ01 | lmb | Laminin-binding surface protein | 17400016 | 100 | 100 | 98 | 1e-172 |
| Streptococcus dysgalactiae subsp. equisimilis strain KNZ01 | lmb | Laminin-binding surface protein | 17400016 | 100 | 100 | 98 | 1e-172 |
| Streptococcus dysgalactiae subsp. equisimilis strain KNZ01 | lmb | Laminin-binding surface protein | 17400016 | 100 | 100 | 99 | 1e-174 |
| Streptococcus dysgalactiae subsp. equisimilis strain KNZ03 | lmb | Laminin-binding surface protein | 17400016 | 100 | 100 | 97 | 1e-171 |
| Streptococcus dysgalactiae subsp. equisimilis strain KNZ03 | lmb | Laminin-binding surface protein | 17400016 | 100 | 100 | 97 | 1e-171 |
| Streptococcus dysgalactiae subsp. equisimilis strain KNZ03 | lmb | Laminin-binding surface protein | 17400016 | 100 | 100 | 98 | 1e-173 |
| Streptococcus dysgalactiae subsp. equisimilis strain KNZ03 | lmb | Laminin-binding surface protein | 17400016 | 100 | 100 | 98 | 1e-173 |
| Streptococcus dysgalactiae subsp. equisimilis strain KNZ04 | lmb | Laminin-binding surface protein | 17400016 | 100 | 100 | 97 | 1e-171 |
| Streptococcus dysgalactiae subsp. equisimilis strain KNZ04 | lmb | Laminin-binding surface protein | 17400016 | 100 | 100 | 98 | 1e-173 |
| Streptococcus dysgalactiae subsp. equisimilis strain KNZ06 | lmb | Laminin-binding surface protein | 17400016 | 100 | 100 | 99 | 1e-174 |
| Streptococcus dysgalactiae subsp. equisimilis strain KNZ06 | lmb | Laminin-binding surface protein | 17400016 | 100 | 100 | 98 | 1e-171 |
| Streptococcus dysgalactiae subsp. equisimilis strain KNZ07 | lmb | Laminin-binding surface protein | 17400016 | 100 | 100 | 99 | 1e-174 |
| Streptococcus dysgalactiae subsp. equisimilis strain KNZ07 | lmb | Laminin-binding surface protein | 17400016 | 100 | 100 | 98 | 1e-172 |
| Streptococcus dysgalactiae subsp. equisimilis strain KNZ10 | lmb | Laminin-binding surface protein | 17400016 | 100 | 100 | 99 | 1e-174 |
| Streptococcus dysgalactiae subsp. equisimilis strain KNZ10 | lmb | Laminin-binding surface protein | 17400016 | 100 | 100 | 98 | 1e-172 |
| Streptococcus dysgalactiae subsp. equisimilis strain KNZ12 | lmb | Laminin-binding surface protein | 17400016 | 100 | 100 | 98 | 1e-172 |
| Streptococcus dysgalactiae subsp. equisimilis strain KNZ12 | lmb | Laminin-binding surface protein | 17400016 | 100 | 100 | 99 | 1e-174 |
| Streptococcus dysgalactiae subsp. equisimilis strain KNZ15 | lmb | Laminin-binding surface protein | 17400016 | 100 | 100 | 99 | 1e-174 |
| Streptococcus dysgalactiae subsp. equisimilis strain KNZ15 | lmb | Laminin-binding surface protein | 17400016 | 100 | 100 | 98 | 1e-171 |
| Streptococcus dysgalactiae subsp. equisimilis strain KNZ16 | lmb | Laminin-binding surface protein | 17400016 | 100 | 100 | 97 | 1e-171 |
| Streptococcus dysgalactiae subsp. equisimilis strain KNZ16 | lmb | Laminin-binding surface protein | 17400016 | 100 | 100 | 98 | 1e-173 |
| Streptococcus dysgalactiae subsp. equisimilis strain NCTC11554 | lmb | Laminin-binding surface protein | 17400016 | 100 | 100 | 98 | 1e-173 |
| Streptococcus dysgalactiae subsp. equisimilis strain NCTC11554 | lmb | Laminin-binding surface protein | 17400016 | 100 | 100 | 97 | 1e-171 |
| Streptococcus dysgalactiae subsp. equisimilis strain NCTC11555 | lmb | Laminin-binding surface protein | 17400016 | 100 | 100 | 99 | 1e-173 |
| Streptococcus dysgalactiae subsp. equisimilis strain NCTC11555 | lmb | Laminin-binding surface protein | 17400016 | 100 | 100 | 98 | 1e-171 |
| Streptococcus dysgalactiae subsp. equisimilis strain NCTC11556 | lmb | Laminin-binding surface protein | 17400016 | 100 | 100 | 98 | 1e-172 |
| Streptococcus dysgalactiae subsp. equisimilis strain NCTC11556 | lmb | Laminin-binding surface protein | 17400016 | 100 | 100 | 99 | 1e-174 |
| Streptococcus dysgalactiae subsp. equisimilis strain NCTC11557 | lmb | Laminin-binding surface protein | 17400016 | 100 | 100 | 99 | 1e-173 |
| Streptococcus dysgalactiae subsp. equisimilis strain NCTC11557 | lmb | Laminin-binding surface protein | 17400016 | 100 | 100 | 98 | 1e-171 |
| Streptococcus dysgalactiae subsp. equisimilis strain NCTC11564 | lmb | Laminin-binding surface protein | 17400016 | 100 | 100 | 98 | 1e-171 |
| Streptococcus dysgalactiae subsp. equisimilis strain NCTC11564 | lmb | Laminin-binding surface protein | 17400016 | 100 | 100 | 99 | 1e-174 |
| Streptococcus dysgalactiae subsp. equisimilis strain NCTC5370 | lmb | Laminin-binding surface protein | 17400016 | 100 | 100 | 99 | 1e-174 |
| Streptococcus dysgalactiae subsp. equisimilis strain NCTC5370 | lmb | Laminin-binding surface protein | 17400016 | 100 | 100 | 98 | 1e-172 |
| Streptococcus dysgalactiae subsp. equisimilis strain NCTC5371 | lmb | Laminin-binding surface protein | 17400016 | 100 | 100 | 99 | 1e-174 |
| Streptococcus dysgalactiae subsp. equisimilis strain NCTC5371 | lmb | Laminin-binding surface protein | 17400016 | 100 | 100 | 98 | 1e-171 |
| Streptococcus dysgalactiae subsp. equisimilis strain NCTC6179 | lmb | Laminin-binding surface protein | 17400016 | 100 | 100 | 83 | 1e-147 |
| Streptococcus dysgalactiae subsp. equisimilis strain NCTC6179 | lmb | Laminin-binding surface protein | 17400016 | 100 | 100 | 84 | 1e-148 |
| Streptococcus dysgalactiae subsp. equisimilis strain NCTC6181 | lmb | Laminin-binding surface protein | 17400016 | 100 | 100 | 83 | 1e-147 |
| Streptococcus dysgalactiae subsp. equisimilis strain NCTC6181 | lmb | Laminin-binding surface protein | 17400016 | 100 | 100 | 84 | 1e-148 |
| Streptococcus dysgalactiae subsp. equisimilis strain NCTC7136 | lmb | Laminin-binding surface protein | 17400016 | 100 | 100 | 98 | 1e-171 |
| Streptococcus dysgalactiae subsp. equisimilis strain NCTC7136 | lmb | Laminin-binding surface protein | 17400016 | 100 | 100 | 99 | 1e-174 |
| Streptococcus dysgalactiae subsp. equisimilis strain NCTC8543 | lmb | Laminin-binding surface protein | 17400016 | 100 | 100 | 98 | 1e-172 |
| Streptococcus dysgalactiae subsp. equisimilis strain NCTC8543 | lmb | Laminin-binding surface protein | 17400016 | 100 | 100 | 99 | 1e-174 |
| Streptococcus dysgalactiae subsp. equisimilis strain NCTC8546 | lmb | Laminin-binding surface protein | 17400016 | 100 | 100 | 99 | 1e-174 |
| Streptococcus dysgalactiae subsp. equisimilis strain NCTC8546 | lmb | Laminin-binding surface protein | 17400016 | 100 | 100 | 98 | 1e-171 |
| Streptococcus dysgalactiae subsp. equisimilis strain NCTC9413 | lmb | Laminin-binding surface protein | 17400016 | 100 | 100 | 99 | 1e-174 |
| Streptococcus dysgalactiae subsp. equisimilis strain NCTC9413 | lmb | Laminin-binding surface protein | 17400016 | 100 | 100 | 98 | 1e-171 |
| Streptococcus dysgalactiae subsp. equisimilis strain NCTC9414 | lmb | Laminin-binding surface protein | 17400016 | 100 | 100 | 99 | 1e-174 |
| Streptococcus dysgalactiae subsp. equisimilis strain NCTC9414 | lmb | Laminin-binding surface protein | 17400016 | 100 | 100 | 98 | 1e-171 |
| Streptococcus dysgalactiae subsp. equisimilis strain NCTC9603 | lmb | Laminin-binding surface protein | 17400016 | 100 | 100 | 98 | 1e-171 |
| Streptococcus dysgalactiae subsp. equisimilis strain NCTC9603 | lmb | Laminin-binding surface protein | 17400016 | 100 | 100 | 99 | 1e-173 |
| Streptococcus dysgalactiae subsp. equisimilis strain SS1575 | lmb | Laminin-binding surface protein | 17400016 | 100 | 100 | 100 | 1e-175 |
| Streptococcus dysgalactiae subsp. equisimilis strain SS1575 | lmb | Laminin-binding surface protein | 17400016 | 100 | 100 | 99 | 1e-173 |
| Streptococcus dysgalactiae subsp. equisimilis strain T642 | lmb | Laminin-binding surface protein | 17400016 | 100 | 100 | 99 | 0.0 |
| Streptococcus dysgalactiae subsp. equisimilis strain T642 | lmb | Laminin-binding surface protein | 17400016 | 100 | 100 | 98 | 0.0 |
| Streptococcus dysgalactiae subsp. equisimilis strain UT_4031CC | lmb | Laminin-binding surface protein | 17400016 | 100 | 100 | 99 | 1e-173 |
| Streptococcus dysgalactiae subsp. equisimilis strain UT_4031CC | lmb | Laminin-binding surface protein | 17400016 | 100 | 100 | 100 | 1e-175 |
| Streptococcus dysgalactiae subsp. equisimilis strain UT_4231_KK | lmb | Laminin-binding surface protein | 17400016 | 100 | 100 | 99 | 1e-173 |
| Streptococcus dysgalactiae subsp. equisimilis strain UT_4231_KK | lmb | Laminin-binding surface protein | 17400016 | 100 | 100 | 98 | 1e-170 |
| Streptococcus dysgalactiae subsp. equisimilis strain UT_4234_DH | lmb | Laminin-binding surface protein | 17400016 | 100 | 100 | 99 | 1e-173 |
| Streptococcus dysgalactiae subsp. equisimilis strain UT_4234_DH | lmb | Laminin-binding surface protein | 17400016 | 100 | 100 | 98 | 1e-171 |
| Streptococcus dysgalactiae subsp. equisimilis strain UT_4241_XS | lmb | Laminin-binding surface protein | 17400016 | 100 | 100 | 98 | 1e-172 |
| Streptococcus dysgalactiae subsp. equisimilis strain UT_4241_XS | lmb | Laminin-binding surface protein | 17400016 | 100 | 100 | 99 | 1e-174 |
| Streptococcus dysgalactiae subsp. equisimilis strain UT_4242_AB | lmb | Laminin-binding surface protein | 17400016 | 100 | 100 | 98 | 1e-172 |
| Streptococcus dysgalactiae subsp. equisimilis strain UT_4242_AB | lmb | Laminin-binding surface protein | 17400016 | 100 | 100 | 99 | 1e-174 |
| Streptococcus dysgalactiae subsp. equisimilis strain UT_4255RC | lmb | Laminin-binding surface protein | 17400016 | 100 | 100 | 99 | 1e-173 |
| Streptococcus dysgalactiae subsp. equisimilis strain UT_4255RC | lmb | Laminin-binding surface protein | 17400016 | 100 | 100 | 98 | 1e-171 |
| Streptococcus dysgalactiae subsp. equisimilis strain UT_4277_BB | lmb | Laminin-binding surface protein | 17400016 | 100 | 100 | 100 | 1e-175 |
| Streptococcus dysgalactiae subsp. equisimilis strain UT_4277_BB | lmb | Laminin-binding surface protein | 17400016 | 100 | 100 | 99 | 1e-173 |
| Streptococcus dysgalactiae subsp. equisimilis strain UT_4966_RC | lmb | Laminin-binding surface protein | 17400016 | 100 | 100 | 99 | 1e-174 |
| Streptococcus dysgalactiae subsp. equisimilis strain UT_4966_RC | lmb | Laminin-binding surface protein | 17400016 | 100 | 100 | 98 | 1e-172 |
| Streptococcus dysgalactiae subsp. equisimilis strain UT-5345 | lmb | Laminin-binding surface protein | 17400016 | 100 | 100 | 99 | 1e-173 |
| Streptococcus dysgalactiae subsp. equisimilis strain UT-5345 | lmb | Laminin-binding surface protein | 17400016 | 100 | 100 | 98 | 1e-171 |
| Streptococcus dysgalactiae subsp. equisimilis strain UT-5354 | lmb | Laminin-binding surface protein | 17400016 | 97 | 100 | 99 | 1e-170 |
| Streptococcus dysgalactiae subsp. equisimilis strain UT-5354 | lmb | Laminin-binding surface protein | 17400016 | 97 | 100 | 98 | 1e-169 |
| Streptococcus dysgalactiae subsp. equisimilis strain UT-SS1069 | lmb | Laminin-binding surface protein | 17400016 | 100 | 100 | 98 | 1e-172 |
| Streptococcus dysgalactiae subsp. equisimilis strain UT-SS1069 | lmb | Laminin-binding surface protein | 17400016 | 100 | 100 | 99 | 1e-174 |
| Streptococcus dysgalactiae subsp. equisimilis strain UT-SS957 | lmb | Laminin-binding surface protein | 17400016 | 100 | 100 | 98 | 1e-171 |
| Streptococcus dysgalactiae subsp. equisimilis strain UT-SS957 | lmb | Laminin-binding surface protein | 17400016 | 100 | 100 | 99 | 1e-173 |
| Streptococcus dysgalactiae subsp. equisimilis strain WCHSDSE-1 | lmb | Laminin-binding surface protein | 17400016 | 100 | 100 | 99 | 1e-174 |
| Streptococcus dysgalactiae subsp. equisimilis strain WCHSDSE-1 | lmb | Laminin-binding surface protein | 17400016 | 100 | 100 | 99 | 1e-174 |
| Streptococcus dysgalactiae subsp. equisimilis 167 | leuS | Leucyl-tRNA synthetase (EC 6.1.1.4) | 12207705 | 100 | 100 | 90 | 0.0 |
| Streptococcus dysgalactiae subsp. equisimilis AC-2713 | leuS | Leucyl-tRNA synthetase (EC 6.1.1.4) | 12207705 | 100 | 100 | 90 | 0.0 |
| Streptococcus dysgalactiae subsp. equisimilis AKSDE4288 | leuS | Leucyl-tRNA synthetase (EC 6.1.1.4) | 12207705 | 100 | 100 | 89 | 0.0 |
| Streptococcus dysgalactiae subsp. equisimilis ATCC 12394 | leuS | Leucyl-tRNA synthetase (EC 6.1.1.4) | 12207705 | 100 | 100 | 90 | 0.0 |
| Streptococcus dysgalactiae subsp. equisimilis GGS_124 | leuS | Leucyl-tRNA synthetase (EC 6.1.1.4) | 12207705 | 100 | 100 | 90 | 0.0 |
| Streptococcus dysgalactiae subsp. equisimilis RE378 | leuS | Leucyl-tRNA synthetase (EC 6.1.1.4) | 12207705 | 100 | 100 | 90 | 0.0 |
| Streptococcus dysgalactiae subsp. equisimilis SD SCDR1 | leuS | Leucyl-tRNA synthetase (EC 6.1.1.4) | 12207705 | 100 | 100 | 90 | 0.0 |
| Streptococcus dysgalactiae subsp. equisimilis SK1249 | leuS | Leucyl-tRNA synthetase (EC 6.1.1.4) | 12207705 | 100 | 100 | 89 | 0.0 |
| Streptococcus dysgalactiae subsp. equisimilis SK1250 | leuS | Leucyl-tRNA synthetase (EC 6.1.1.4) | 12207705 | 100 | 100 | 90 | 0.0 |
| Streptococcus dysgalactiae subsp. equisimilis strain ASDSE_96 | leuS | Leucyl-tRNA synthetase (EC 6.1.1.4) | 12207705 | 100 | 100 | 89 | 0.0 |
| Streptococcus dysgalactiae subsp. equisimilis strain ASDSE_99 | leuS | Leucyl-tRNA synthetase (EC 6.1.1.4) | 12207705 | 100 | 100 | 90 | 0.0 |
| Streptococcus dysgalactiae subsp. equisimilis strain C161L1 | leuS | Leucyl-tRNA synthetase (EC 6.1.1.4) | 12207705 | 100 | 100 | 90 | 0 |
| Streptococcus dysgalactiae subsp. equisimilis strain KNZ01 | leuS | Leucyl-tRNA synthetase (EC 6.1.1.4) | 12207705 | 100 | 100 | 90 | 0.0 |
| Streptococcus dysgalactiae subsp. equisimilis strain KNZ03 | leuS | Leucyl-tRNA synthetase (EC 6.1.1.4) | 12207705 | 100 | 100 | 90 | 0.0 |
| Streptococcus dysgalactiae subsp. equisimilis strain KNZ04 | leuS | Leucyl-tRNA synthetase (EC 6.1.1.4) | 12207705 | 100 | 100 | 90 | 0.0 |
| Streptococcus dysgalactiae subsp. equisimilis strain KNZ06 | leuS | Leucyl-tRNA synthetase (EC 6.1.1.4) | 12207705 | 100 | 100 | 90 | 0.0 |
| Streptococcus dysgalactiae subsp. equisimilis strain KNZ07 | leuS | Leucyl-tRNA synthetase (EC 6.1.1.4) | 12207705 | 100 | 100 | 90 | 0.0 |
| Streptococcus dysgalactiae subsp. equisimilis strain KNZ10 | leuS | Leucyl-tRNA synthetase (EC 6.1.1.4) | 12207705 | 100 | 100 | 90 | 0.0 |
| Streptococcus dysgalactiae subsp. equisimilis strain KNZ12 | leuS | Leucyl-tRNA synthetase (EC 6.1.1.4) | 12207705 | 100 | 100 | 90 | 0.0 |
| Streptococcus dysgalactiae subsp. equisimilis strain KNZ15 | leuS | Leucyl-tRNA synthetase (EC 6.1.1.4) | 12207705 | 100 | 100 | 90 | 0.0 |
| Streptococcus dysgalactiae subsp. equisimilis strain KNZ16 | leuS | Leucyl-tRNA synthetase (EC 6.1.1.4) | 12207705 | 100 | 100 | 90 | 0.0 |
| Streptococcus dysgalactiae subsp. equisimilis strain NCTC10321 | leuS | Leucyl-tRNA synthetase (EC 6.1.1.4) | 12207705 | 100 | 100 | 90 | 0.0 |
| Streptococcus dysgalactiae subsp. equisimilis strain NCTC11554 | leuS | Leucyl-tRNA synthetase (EC 6.1.1.4) | 12207705 | 100 | 100 | 90 | 0.0 |
| Streptococcus dysgalactiae subsp. equisimilis strain NCTC11555 | leuS | Leucyl-tRNA synthetase (EC 6.1.1.4) | 12207705 | 100 | 100 | 89 | 0.0 |
| Streptococcus dysgalactiae subsp. equisimilis strain NCTC11556 | leuS | Leucyl-tRNA synthetase (EC 6.1.1.4) | 12207705 | 100 | 100 | 90 | 0.0 |
| Streptococcus dysgalactiae subsp. equisimilis strain NCTC11557 | leuS | Leucyl-tRNA synthetase (EC 6.1.1.4) | 12207705 | 100 | 100 | 90 | 0.0 |
| Streptococcus dysgalactiae subsp. equisimilis strain NCTC11564 | leuS | Leucyl-tRNA synthetase (EC 6.1.1.4) | 12207705 | 100 | 100 | 90 | 0.0 |
| Streptococcus dysgalactiae subsp. equisimilis strain NCTC5370 | leuS | Leucyl-tRNA synthetase (EC 6.1.1.4) | 12207705 | 100 | 100 | 90 | 0.0 |
| Streptococcus dysgalactiae subsp. equisimilis strain NCTC5371 | leuS | Leucyl-tRNA synthetase (EC 6.1.1.4) | 12207705 | 100 | 100 | 90 | 0.0 |
| Streptococcus dysgalactiae subsp. equisimilis strain NCTC6179 | leuS | Leucyl-tRNA synthetase (EC 6.1.1.4) | 12207705 | 100 | 100 | 88 | 0.0 |
| Streptococcus dysgalactiae subsp. equisimilis strain NCTC6181 | leuS | Leucyl-tRNA synthetase (EC 6.1.1.4) | 12207705 | 100 | 100 | 88 | 0.0 |
| Streptococcus dysgalactiae subsp. equisimilis strain NCTC6407 | leuS | Leucyl-tRNA synthetase (EC 6.1.1.4) | 12207705 | 100 | 100 | 90 | 0.0 |
| Streptococcus dysgalactiae subsp. equisimilis strain NCTC7136 | leuS | Leucyl-tRNA synthetase (EC 6.1.1.4) | 12207705 | 100 | 100 | 90 | 0.0 |
| Streptococcus dysgalactiae subsp. equisimilis strain NCTC8543 | leuS | Leucyl-tRNA synthetase (EC 6.1.1.4) | 12207705 | 100 | 100 | 90 | 0.0 |
| Streptococcus dysgalactiae subsp. equisimilis strain NCTC8546 | leuS | Leucyl-tRNA synthetase (EC 6.1.1.4) | 12207705 | 100 | 100 | 89 | 0.0 |
| Streptococcus dysgalactiae subsp. equisimilis strain NCTC9413 | leuS | Leucyl-tRNA synthetase (EC 6.1.1.4) | 12207705 | 100 | 100 | 89 | 0.0 |
| Streptococcus dysgalactiae subsp. equisimilis strain NCTC9414 | leuS | Leucyl-tRNA synthetase (EC 6.1.1.4) | 12207705 | 100 | 100 | 90 | 0.0 |
| Streptococcus dysgalactiae subsp. equisimilis strain NCTC9603 | leuS | Leucyl-tRNA synthetase (EC 6.1.1.4) | 12207705 | 100 | 100 | 90 | 0.0 |
| Streptococcus dysgalactiae subsp. equisimilis strain SS1575 | leuS | Leucyl-tRNA synthetase (EC 6.1.1.4) | 12207705 | 100 | 100 | 90 | 0.0 |
| Streptococcus dysgalactiae subsp. equisimilis strain T642 | leuS | Leucyl-tRNA synthetase (EC 6.1.1.4) | 12207705 | 100 | 100 | 89 | 0.0 |
| Streptococcus dysgalactiae subsp. equisimilis strain UT_4031CC | leuS | Leucyl-tRNA synthetase (EC 6.1.1.4) | 12207705 | 100 | 100 | 90 | 0.0 |
| Streptococcus dysgalactiae subsp. equisimilis strain UT_4231_KK | leuS | Leucyl-tRNA synthetase (EC 6.1.1.4) | 12207705 | 100 | 100 | 90 | 0.0 |
| Streptococcus dysgalactiae subsp. equisimilis strain UT_4234_DH | leuS | Leucyl-tRNA synthetase (EC 6.1.1.4) | 12207705 | 100 | 100 | 90 | 0.0 |
| Streptococcus dysgalactiae subsp. equisimilis strain UT_4241_XS | leuS | Leucyl-tRNA synthetase (EC 6.1.1.4) | 12207705 | 100 | 100 | 90 | 0.0 |
| Streptococcus dysgalactiae subsp. equisimilis strain UT_4242_AB | leuS | Leucyl-tRNA synthetase (EC 6.1.1.4) | 12207705 | 100 | 100 | 90 | 0.0 |
| Streptococcus dysgalactiae subsp. equisimilis strain UT_4255RC | leuS | Leucyl-tRNA synthetase (EC 6.1.1.4) | 12207705 | 100 | 100 | 90 | 0.0 |
| Streptococcus dysgalactiae subsp. equisimilis strain UT_4277_BB | leuS | Leucyl-tRNA synthetase (EC 6.1.1.4) | 12207705 | 100 | 100 | 90 | 0.0 |
| Streptococcus dysgalactiae subsp. equisimilis strain UT_4966_RC | leuS | Leucyl-tRNA synthetase (EC 6.1.1.4) | 12207705 | 100 | 100 | 90 | 0.0 |
| Streptococcus dysgalactiae subsp. equisimilis strain UT-5345 | leuS | Leucyl-tRNA synthetase (EC 6.1.1.4) | 12207705 | 100 | 100 | 90 | 0.0 |
| Streptococcus dysgalactiae subsp. equisimilis strain UT-5354 | leuS | Leucyl-tRNA synthetase (EC 6.1.1.4) | 12207705 | 100 | 100 | 90 | 0.0 |
| Streptococcus dysgalactiae subsp. equisimilis strain UT-SS1069 | leuS | Leucyl-tRNA synthetase (EC 6.1.1.4) | 12207705 | 61 | 100 | 88 | 1e-277 |
| Streptococcus dysgalactiae subsp. equisimilis strain UT-SS957 | leuS | Leucyl-tRNA synthetase (EC 6.1.1.4) | 12207705 | 100 | 100 | 90 | 0.0 |
| Streptococcus dysgalactiae subsp. equisimilis strain WCHSDSE-1 | leuS | Leucyl-tRNA synthetase (EC 6.1.1.4) | 12207705 | 100 | 100 | 90 | 0.0 |
| Streptococcus dysgalactiae subsp. equisimilis 167 | lsp | Lipoprotein signal peptidase (EC 3.4.23.36) | 19398546 | 63 | 98 | 84 | 7e-44 |
| Streptococcus dysgalactiae subsp. equisimilis AC-2713 | lsp | Lipoprotein signal peptidase (EC 3.4.23.36) | 19398546 | 100 | 100 | 83 | 3e-71 |
| Streptococcus dysgalactiae subsp. equisimilis AKSDE4288 | lsp | Lipoprotein signal peptidase (EC 3.4.23.36) | 19398546 | 100 | 100 | 83 | 3e-71 |
| Streptococcus dysgalactiae subsp. equisimilis ATCC 12394 | lsp | Lipoprotein signal peptidase (EC 3.4.23.36) | 19398546 | 100 | 100 | 83 | 3e-71 |
| Streptococcus dysgalactiae subsp. equisimilis GGS_124 | lsp | Lipoprotein signal peptidase (EC 3.4.23.36) | 19398546 | 100 | 100 | 83 | 3e-71 |
| Streptococcus dysgalactiae subsp. equisimilis RE378 | lsp | Lipoprotein signal peptidase (EC 3.4.23.36) | 19398546 | 100 | 100 | 83 | 3e-71 |
| Streptococcus dysgalactiae subsp. equisimilis SD SCDR1 | lsp | Lipoprotein signal peptidase (EC 3.4.23.36) | 19398546 | 100 | 100 | 83 | 3e-71 |
| Streptococcus dysgalactiae subsp. equisimilis SK1249 | lspA | Lipoprotein signal peptidase (EC 3.4.23.36) | 19398546 | 100 | 100 | 83 | 3e-71 |
| Streptococcus dysgalactiae subsp. equisimilis SK1250 | lspA | Lipoprotein signal peptidase (EC 3.4.23.36) | 19398546 | 100 | 100 | 98 | 2e-81 |
| Streptococcus dysgalactiae subsp. equisimilis strain ASDSE_96 | lsp | Lipoprotein signal peptidase (EC 3.4.23.36) | 19398546 | 100 | 100 | 83 | 3e-71 |
| Streptococcus dysgalactiae subsp. equisimilis strain ASDSE_99 | lsp | Lipoprotein signal peptidase (EC 3.4.23.36) | 19398546 | 100 | 100 | 99 | 9e-82 |
| Streptococcus dysgalactiae subsp. equisimilis strain C161L1 | lsp | Lipoprotein signal peptidase (EC 3.4.23.36) | 19398546 | 100 | 100 | 83 | 0 |
| Streptococcus dysgalactiae subsp. equisimilis strain KNZ01 | lsp | Lipoprotein signal peptidase (EC 3.4.23.36) | 19398546 | 100 | 100 | 83 | 3e-71 |
| Streptococcus dysgalactiae subsp. equisimilis strain KNZ03 | lsp | Lipoprotein signal peptidase (EC 3.4.23.36) | 19398546 | 100 | 100 | 83 | 3e-71 |
| Streptococcus dysgalactiae subsp. equisimilis strain KNZ04 | lsp | Lipoprotein signal peptidase (EC 3.4.23.36) | 19398546 | 100 | 100 | 83 | 3e-71 |
| Streptococcus dysgalactiae subsp. equisimilis strain KNZ06 | lsp | Lipoprotein signal peptidase (EC 3.4.23.36) | 19398546 | 100 | 100 | 83 | 3e-71 |
| Streptococcus dysgalactiae subsp. equisimilis strain KNZ07 | lsp | Lipoprotein signal peptidase (EC 3.4.23.36) | 19398546 | 100 | 100 | 83 | 3e-71 |
| Streptococcus dysgalactiae subsp. equisimilis strain KNZ10 | lsp | Lipoprotein signal peptidase (EC 3.4.23.36) | 19398546 | 100 | 100 | 83 | 3e-71 |
| Streptococcus dysgalactiae subsp. equisimilis strain KNZ12 | lsp | Lipoprotein signal peptidase (EC 3.4.23.36) | 19398546 | 100 | 100 | 83 | 3e-71 |
| Streptococcus dysgalactiae subsp. equisimilis strain KNZ15 | lsp | Lipoprotein signal peptidase (EC 3.4.23.36) | 19398546 | 100 | 100 | 83 | 3e-71 |
| Streptococcus dysgalactiae subsp. equisimilis strain KNZ16 | lsp | Lipoprotein signal peptidase (EC 3.4.23.36) | 19398546 | 100 | 100 | 83 | 3e-71 |
| Streptococcus dysgalactiae subsp. equisimilis strain NCTC10321 | lsp | Lipoprotein signal peptidase (EC 3.4.23.36) | 19398546 | 100 | 100 | 82 | 1e-70 |
| Streptococcus dysgalactiae subsp. equisimilis strain NCTC11554 | lsp | Lipoprotein signal peptidase (EC 3.4.23.36) | 19398546 | 100 | 100 | 83 | 3e-71 |
| Streptococcus dysgalactiae subsp. equisimilis strain NCTC11555 | lsp | Lipoprotein signal peptidase (EC 3.4.23.36) | 19398546 | 100 | 100 | 99 | 9e-82 |
| Streptococcus dysgalactiae subsp. equisimilis strain NCTC11556 | lsp | Lipoprotein signal peptidase (EC 3.4.23.36) | 19398546 | 100 | 100 | 83 | 3e-71 |
| Streptococcus dysgalactiae subsp. equisimilis strain NCTC11557 | lsp | Lipoprotein signal peptidase (EC 3.4.23.36) | 19398546 | 100 | 100 | 99 | 9e-82 |
| Streptococcus dysgalactiae subsp. equisimilis strain NCTC11564 | lsp | Lipoprotein signal peptidase (EC 3.4.23.36) | 19398546 | 100 | 100 | 83 | 3e-71 |
| Streptococcus dysgalactiae subsp. equisimilis strain NCTC5370 | lsp | Lipoprotein signal peptidase (EC 3.4.23.36) | 19398546 | 100 | 100 | 83 | 3e-71 |
| Streptococcus dysgalactiae subsp. equisimilis strain NCTC5371 | lsp | Lipoprotein signal peptidase (EC 3.4.23.36) | 19398546 | 100 | 100 | 83 | 3e-71 |
| Streptococcus dysgalactiae subsp. equisimilis strain NCTC6179 | lsp | Lipoprotein signal peptidase (EC 3.4.23.36) | 19398546 | 100 | 100 | 84 | 6e-72 |
| Streptococcus dysgalactiae subsp. equisimilis strain NCTC6181 | lsp | Lipoprotein signal peptidase (EC 3.4.23.36) | 19398546 | 100 | 100 | 84 | 6e-72 |
| Streptococcus dysgalactiae subsp. equisimilis strain NCTC6407 | lsp | Lipoprotein signal peptidase (EC 3.4.23.36) | 19398546 | 100 | 100 | 82 | 1e-70 |
| Streptococcus dysgalactiae subsp. equisimilis strain NCTC7136 | lsp | Lipoprotein signal peptidase (EC 3.4.23.36) | 19398546 | 100 | 100 | 83 | 3e-71 |
| Streptococcus dysgalactiae subsp. equisimilis strain NCTC8543 | lsp | Lipoprotein signal peptidase (EC 3.4.23.36) | 19398546 | 100 | 100 | 83 | 3e-71 |
| Streptococcus dysgalactiae subsp. equisimilis strain NCTC8546 | lsp | Lipoprotein signal peptidase (EC 3.4.23.36) | 19398546 | 100 | 100 | 83 | 3e-71 |
| Streptococcus dysgalactiae subsp. equisimilis strain NCTC9413 | lsp | Lipoprotein signal peptidase (EC 3.4.23.36) | 19398546 | 100 | 100 | 83 | 3e-71 |
| Streptococcus dysgalactiae subsp. equisimilis strain NCTC9414 | lsp | Lipoprotein signal peptidase (EC 3.4.23.36) | 19398546 | 100 | 100 | 83 | 3e-71 |
| Streptococcus dysgalactiae subsp. equisimilis strain NCTC9603 | lsp | Lipoprotein signal peptidase (EC 3.4.23.36) | 19398546 | 100 | 100 | 83 | 3e-71 |
| Streptococcus dysgalactiae subsp. equisimilis strain SS1575 | lsp | Lipoprotein signal peptidase (EC 3.4.23.36) | 19398546 | 100 | 100 | 83 | 3e-71 |
| Streptococcus dysgalactiae subsp. equisimilis strain T642 | lsp | Lipoprotein signal peptidase (EC 3.4.23.36) | 19398546 | 100 | 100 | 82 | 8e-91 |
| Streptococcus dysgalactiae subsp. equisimilis strain UT_4031CC | lsp | Lipoprotein signal peptidase (EC 3.4.23.36) | 19398546 | 100 | 100 | 83 | 3e-71 |
| Streptococcus dysgalactiae subsp. equisimilis strain UT_4231_KK | lsp | Lipoprotein signal peptidase (EC 3.4.23.36) | 19398546 | 100 | 100 | 83 | 3e-71 |
| Streptococcus dysgalactiae subsp. equisimilis strain UT_4234_DH | lsp | Lipoprotein signal peptidase (EC 3.4.23.36) | 19398546 | 100 | 100 | 99 | 9e-82 |
| Streptococcus dysgalactiae subsp. equisimilis strain UT_4241_XS | lsp | Lipoprotein signal peptidase (EC 3.4.23.36) | 19398546 | 100 | 100 | 83 | 3e-71 |
| Streptococcus dysgalactiae subsp. equisimilis strain UT_4242_AB | lsp | Lipoprotein signal peptidase (EC 3.4.23.36) | 19398546 | 100 | 100 | 83 | 3e-71 |
| Streptococcus dysgalactiae subsp. equisimilis strain UT_4255RC | lsp | Lipoprotein signal peptidase (EC 3.4.23.36) | 19398546 | 100 | 100 | 99 | 9e-82 |
| Streptococcus dysgalactiae subsp. equisimilis strain UT_4277_BB | lsp | Lipoprotein signal peptidase (EC 3.4.23.36) | 19398546 | 100 | 100 | 83 | 3e-71 |
| Streptococcus dysgalactiae subsp. equisimilis strain UT_4966_RC | lsp | Lipoprotein signal peptidase (EC 3.4.23.36) | 19398546 | 100 | 100 | 83 | 3e-71 |
| Streptococcus dysgalactiae subsp. equisimilis strain UT-5345 | lsp | Lipoprotein signal peptidase (EC 3.4.23.36) | 19398546 | 100 | 100 | 83 | 3e-71 |
| Streptococcus dysgalactiae subsp. equisimilis strain UT-5354 | lsp | Lipoprotein signal peptidase (EC 3.4.23.36) | 19398546 | 100 | 100 | 98 | 2e-81 |
| Streptococcus dysgalactiae subsp. equisimilis strain UT-SS1069 | lsp | Lipoprotein signal peptidase (EC 3.4.23.36) | 19398546 | 100 | 100 | 83 | 3e-71 |
| Streptococcus dysgalactiae subsp. equisimilis strain UT-SS957 | lsp | Lipoprotein signal peptidase (EC 3.4.23.36) | 19398546 | 100 | 100 | 83 | 3e-71 |
| Streptococcus dysgalactiae subsp. equisimilis strain WCHSDSE-1 | lsp | Lipoprotein signal peptidase (EC 3.4.23.36) | 19398546 | 100 | 100 | 83 | 3e-71 |
| Streptococcus dysgalactiae subsp. equisimilis 167 | atmB | Methionine ABC transporter substrate-binding protein | 19223485 | 98 | 98 | 95 | 1e-150 |
| Streptococcus dysgalactiae subsp. equisimilis AC-2713 | atmB | Methionine ABC transporter substrate-binding protein | 19223485 | 98 | 98 | 95 | 1e-150 |
| Streptococcus dysgalactiae subsp. equisimilis AKSDE4288 | atmB | Methionine ABC transporter substrate-binding protein | 19223485 | 98 | 98 | 95 | 1e-150 |
| Streptococcus dysgalactiae subsp. equisimilis ATCC 12394 | atmB | Methionine ABC transporter substrate-binding protein | 19223485 | 98 | 98 | 95 | 1e-149 |
| Streptococcus dysgalactiae subsp. equisimilis GGS_124 | atmB | Methionine ABC transporter substrate-binding protein | 19223485 | 98 | 98 | 95 | 1e-150 |
| Streptococcus dysgalactiae subsp. equisimilis RE378 | metQ | Methionine ABC transporter substrate-binding protein | 19223485 | 98 | 98 | 95 | 1e-150 |
| Streptococcus dysgalactiae subsp. equisimilis SD SCDR1 | atmB | Methionine ABC transporter substrate-binding protein | 19223485 | 98 | 98 | 95 | 1e-149 |
| Streptococcus dysgalactiae subsp. equisimilis SK1249 | atmB | Methionine ABC transporter substrate-binding protein | 19223485 | 92 | 99 | 95 | 1e-141 |
| Streptococcus dysgalactiae subsp. equisimilis SK1250 | atmB | Methionine ABC transporter substrate-binding protein | 19223485 | 98 | 98 | 95 | 1e-149 |
| Streptococcus dysgalactiae subsp. equisimilis strain ASDSE_96 | atmB | Methionine ABC transporter substrate-binding protein | 19223485 | 98 | 98 | 95 | 1e-150 |
| Streptococcus dysgalactiae subsp. equisimilis strain ASDSE_99 | atmB | Methionine ABC transporter substrate-binding protein | 19223485 | 98 | 98 | 95 | 1e-149 |
| Streptococcus dysgalactiae subsp. equisimilis strain C161L1 | atmB | Methionine ABC transporter substrate-binding protein | 19223485 | 100 | 100 | 94 | 0 |
| Streptococcus dysgalactiae subsp. equisimilis strain KNZ01 | atmB | Methionine ABC transporter substrate-binding protein | 19223485 | 98 | 98 | 95 | 1e-150 |
| Streptococcus dysgalactiae subsp. equisimilis strain KNZ03 | atmB | Methionine ABC transporter substrate-binding protein | 19223485 | 98 | 98 | 95 | 1e-149 |
| Streptococcus dysgalactiae subsp. equisimilis strain KNZ04 | atmB | Methionine ABC transporter substrate-binding protein | 19223485 | 98 | 98 | 95 | 1e-149 |
| Streptococcus dysgalactiae subsp. equisimilis strain KNZ06 | atmB | Methionine ABC transporter substrate-binding protein | 19223485 | 98 | 98 | 95 | 1e-149 |
| Streptococcus dysgalactiae subsp. equisimilis strain KNZ07 | atmB | Methionine ABC transporter substrate-binding protein | 19223485 | 98 | 98 | 95 | 1e-150 |
| Streptococcus dysgalactiae subsp. equisimilis strain KNZ10 | atmB | Methionine ABC transporter substrate-binding protein | 19223485 | 98 | 98 | 95 | 1e-150 |
| Streptococcus dysgalactiae subsp. equisimilis strain KNZ12 | atmB | Methionine ABC transporter substrate-binding protein | 19223485 | 98 | 98 | 95 | 1e-150 |
| Streptococcus dysgalactiae subsp. equisimilis strain KNZ15 | atmB | Methionine ABC transporter substrate-binding protein | 19223485 | 98 | 98 | 95 | 1e-150 |
| Streptococcus dysgalactiae subsp. equisimilis strain KNZ16 | atmB | Methionine ABC transporter substrate-binding protein | 19223485 | 98 | 98 | 95 | 1e-149 |
| Streptococcus dysgalactiae subsp. equisimilis strain NCTC10321 | atmB | Methionine ABC transporter substrate-binding protein | 19223485 | 98 | 98 | 95 | 1e-150 |
| Streptococcus dysgalactiae subsp. equisimilis strain NCTC11554 | atmB | Methionine ABC transporter substrate-binding protein | 19223485 | 98 | 98 | 95 | 1e-149 |
| Streptococcus dysgalactiae subsp. equisimilis strain NCTC11555 | atmB | Methionine ABC transporter substrate-binding protein | 19223485 | 98 | 98 | 95 | 1e-149 |
| Streptococcus dysgalactiae subsp. equisimilis strain NCTC11556 | atmB | Methionine ABC transporter substrate-binding protein | 19223485 | 98 | 98 | 95 | 1e-149 |
| Streptococcus dysgalactiae subsp. equisimilis strain NCTC11557 | atmB | Methionine ABC transporter substrate-binding protein | 19223485 | 98 | 98 | 95 | 1e-149 |
| Streptococcus dysgalactiae subsp. equisimilis strain NCTC11564 | atmB | Methionine ABC transporter substrate-binding protein | 19223485 | 98 | 98 | 95 | 1e-150 |
| Streptococcus dysgalactiae subsp. equisimilis strain NCTC5370 | atmB | Methionine ABC transporter substrate-binding protein | 19223485 | 98 | 98 | 95 | 1e-150 |
| Streptococcus dysgalactiae subsp. equisimilis strain NCTC5371 | atmB | Methionine ABC transporter substrate-binding protein | 19223485 | 98 | 98 | 95 | 1e-150 |
| Streptococcus dysgalactiae subsp. equisimilis strain NCTC6179 | atmB | Methionine ABC transporter substrate-binding protein | 19223485 | 98 | 98 | 95 | 1e-150 |
| Streptococcus dysgalactiae subsp. equisimilis strain NCTC6181 | atmB | Methionine ABC transporter substrate-binding protein | 19223485 | 98 | 98 | 95 | 1e-150 |
| Streptococcus dysgalactiae subsp. equisimilis strain NCTC6407 | atmB | Methionine ABC transporter substrate-binding protein | 19223485 | 98 | 98 | 95 | 1e-150 |
| Streptococcus dysgalactiae subsp. equisimilis strain NCTC7136 | atmB | Methionine ABC transporter substrate-binding protein | 19223485 | 98 | 98 | 95 | 1e-150 |
| Streptococcus dysgalactiae subsp. equisimilis strain NCTC8543 | atmB | Methionine ABC transporter substrate-binding protein | 19223485 | 98 | 98 | 95 | 1e-150 |
| Streptococcus dysgalactiae subsp. equisimilis strain NCTC8546 | atmB | Methionine ABC transporter substrate-binding protein | 19223485 | 98 | 98 | 94 | 1e-148 |
| Streptococcus dysgalactiae subsp. equisimilis strain NCTC9413 | atmB | Methionine ABC transporter substrate-binding protein | 19223485 | 98 | 98 | 94 | 1e-148 |
| Streptococcus dysgalactiae subsp. equisimilis strain NCTC9414 | atmB | Methionine ABC transporter substrate-binding protein | 19223485 | 98 | 98 | 95 | 1e-150 |
| Streptococcus dysgalactiae subsp. equisimilis strain NCTC9603 | atmB | Methionine ABC transporter substrate-binding protein | 19223485 | 98 | 98 | 95 | 1e-150 |
| Streptococcus dysgalactiae subsp. equisimilis strain SS1575 | atmB | Methionine ABC transporter substrate-binding protein | 19223485 | 98 | 98 | 95 | 1e-150 |
| Streptococcus dysgalactiae subsp. equisimilis strain T642 | atmB | Methionine ABC transporter substrate-binding protein | 19223485 | 100 | 100 | 95 | 0 |
| Streptococcus dysgalactiae subsp. equisimilis strain UT_4031CC | atmB | Methionine ABC transporter substrate-binding protein | 19223485 | 98 | 98 | 95 | 1e-150 |
| Streptococcus dysgalactiae subsp. equisimilis strain UT_4231_KK | atmB | Methionine ABC transporter substrate-binding protein | 19223485 | 98 | 98 | 95 | 1e-150 |
| Streptococcus dysgalactiae subsp. equisimilis strain UT_4234_DH | atmB | Methionine ABC transporter substrate-binding protein | 19223485 | 98 | 98 | 95 | 1e-149 |
| Streptococcus dysgalactiae subsp. equisimilis strain UT_4241_XS | atmB | Methionine ABC transporter substrate-binding protein | 19223485 | 98 | 98 | 95 | 1e-150 |
| Streptococcus dysgalactiae subsp. equisimilis strain UT_4242_AB | atmB | Methionine ABC transporter substrate-binding protein | 19223485 | 98 | 98 | 95 | 1e-149 |
| Streptococcus dysgalactiae subsp. equisimilis strain UT_4255RC | atmB | Methionine ABC transporter substrate-binding protein | 19223485 | 98 | 98 | 95 | 1e-149 |
| Streptococcus dysgalactiae subsp. equisimilis strain UT_4277_BB | atmB | Methionine ABC transporter substrate-binding protein | 19223485 | 98 | 98 | 95 | 1e-150 |
| Streptococcus dysgalactiae subsp. equisimilis strain UT_4966_RC | atmB | Methionine ABC transporter substrate-binding protein | 19223485 | 98 | 98 | 95 | 1e-150 |
| Streptococcus dysgalactiae subsp. equisimilis strain UT-5345 | atmB | Methionine ABC transporter substrate-binding protein | 19223485 | 98 | 98 | 95 | 1e-149 |
| Streptococcus dysgalactiae subsp. equisimilis strain UT-5354 | atmB | Methionine ABC transporter substrate-binding protein | 19223485 | 98 | 98 | 95 | 1e-150 |
| Streptococcus dysgalactiae subsp. equisimilis strain UT-SS1069 | atmB | Methionine ABC transporter substrate-binding protein | 19223485 | 98 | 98 | 95 | 1e-150 |
| Streptococcus dysgalactiae subsp. equisimilis strain UT-SS957 | atmB | Methionine ABC transporter substrate-binding protein | 19223485 | 98 | 98 | 95 | 1e-150 |
| Streptococcus dysgalactiae subsp. equisimilis strain WCHSDSE-1 | atmB | Methionine ABC transporter substrate-binding protein | 19223485 | 98 | 98 | 95 | 1e-149 |
| Streptococcus dysgalactiae subsp. equisimilis 167 | deoB | Phosphopentomutase (EC 5.4.2.7) | 12207705 | 99 | 99 | 86 | 1e-205 |
| Streptococcus dysgalactiae subsp. equisimilis AC-2713 | deoB | Phosphopentomutase (EC 5.4.2.7) | 12207705 | 99 | 99 | 86 | 1e-205 |
| Streptococcus dysgalactiae subsp. equisimilis AKSDE4288 | SP_0829 | Phosphopentomutase (EC 5.4.2.7) | 12207705 | 99 | 99 | 86 | 1e-205 |
| Streptococcus dysgalactiae subsp. equisimilis ATCC 12394 | SP_0829 | Phosphopentomutase (EC 5.4.2.7) | 12207705 | 99 | 99 | 86 | 1e-205 |
| Streptococcus dysgalactiae subsp. equisimilis GGS_124 | deoB | Phosphopentomutase (EC 5.4.2.7) | 12207705 | 99 | 99 | 86 | 1e-205 |
| Streptococcus dysgalactiae subsp. equisimilis RE378 | deoB | Phosphopentomutase (EC 5.4.2.7) | 12207705 | 99 | 99 | 86 | 1e-205 |
| Streptococcus dysgalactiae subsp. equisimilis SD SCDR1 | SP_0829 | Phosphopentomutase (EC 5.4.2.7) | 12207705 | 99 | 99 | 86 | 1e-205 |
| Streptococcus dysgalactiae subsp. equisimilis SK1249 | deoB | Phosphopentomutase (EC 5.4.2.7) | 12207705 | 99 | 99 | 86 | 1e-205 |
| Streptococcus dysgalactiae subsp. equisimilis SK1250 | deoB | Phosphopentomutase (EC 5.4.2.7) | 12207705 | 99 | 99 | 86 | 1e-205 |
| Streptococcus dysgalactiae subsp. equisimilis strain ASDSE_96 | SP_0829 | Phosphopentomutase (EC 5.4.2.7) | 12207705 | 99 | 99 | 86 | 1e-205 |
| Streptococcus dysgalactiae subsp. equisimilis strain ASDSE_99 | SP_0829 | Phosphopentomutase (EC 5.4.2.7) | 12207705 | 99 | 99 | 86 | 1e-205 |
| Streptococcus dysgalactiae subsp. equisimilis strain C161L1 | SP_0829 | Phosphopentomutase (EC 5.4.2.7) | 12207705 | 99 | 99 | 86 | 0 |
| Streptococcus dysgalactiae subsp. equisimilis strain KNZ01 | SP_0829 | Phosphopentomutase (EC 5.4.2.7) | 12207705 | 99 | 99 | 86 | 1e-205 |
| Streptococcus dysgalactiae subsp. equisimilis strain KNZ03 | SP_0829 | Phosphopentomutase (EC 5.4.2.7) | 12207705 | 99 | 99 | 86 | 1e-205 |
| Streptococcus dysgalactiae subsp. equisimilis strain KNZ04 | SP_0829 | Phosphopentomutase (EC 5.4.2.7) | 12207705 | 99 | 99 | 86 | 1e-205 |
| Streptococcus dysgalactiae subsp. equisimilis strain KNZ06 | SP_0829 | Phosphopentomutase (EC 5.4.2.7) | 12207705 | 99 | 99 | 86 | 1e-205 |
| Streptococcus dysgalactiae subsp. equisimilis strain KNZ07 | SP_0829 | Phosphopentomutase (EC 5.4.2.7) | 12207705 | 99 | 99 | 86 | 1e-205 |
| Streptococcus dysgalactiae subsp. equisimilis strain KNZ10 | SP_0829 | Phosphopentomutase (EC 5.4.2.7) | 12207705 | 99 | 99 | 86 | 1e-205 |
| Streptococcus dysgalactiae subsp. equisimilis strain KNZ12 | SP_0829 | Phosphopentomutase (EC 5.4.2.7) | 12207705 | 99 | 99 | 86 | 1e-205 |
| Streptococcus dysgalactiae subsp. equisimilis strain KNZ15 | SP_0829 | Phosphopentomutase (EC 5.4.2.7) | 12207705 | 99 | 99 | 86 | 1e-205 |
| Streptococcus dysgalactiae subsp. equisimilis strain KNZ16 | SP_0829 | Phosphopentomutase (EC 5.4.2.7) | 12207705 | 99 | 99 | 86 | 1e-205 |
| Streptococcus dysgalactiae subsp. equisimilis strain NCTC10321 | SP_0829 | Phosphopentomutase (EC 5.4.2.7) | 12207705 | 99 | 99 | 86 | 1e-205 |
| Streptococcus dysgalactiae subsp. equisimilis strain NCTC11554 | SP_0829 | Phosphopentomutase (EC 5.4.2.7) | 12207705 | 99 | 99 | 86 | 1e-205 |
| Streptococcus dysgalactiae subsp. equisimilis strain NCTC11555 | SP_0829 | Phosphopentomutase (EC 5.4.2.7) | 12207705 | 99 | 99 | 86 | 1e-205 |
| Streptococcus dysgalactiae subsp. equisimilis strain NCTC11556 | SP_0829 | Phosphopentomutase (EC 5.4.2.7) | 12207705 | 99 | 99 | 85 | 1e-203 |
| Streptococcus dysgalactiae subsp. equisimilis strain NCTC11557 | SP_0829 | Phosphopentomutase (EC 5.4.2.7) | 12207705 | 99 | 99 | 86 | 1e-205 |
| Streptococcus dysgalactiae subsp. equisimilis strain NCTC11564 | SP_0829 | Phosphopentomutase (EC 5.4.2.7) | 12207705 | 99 | 99 | 86 | 1e-205 |
| Streptococcus dysgalactiae subsp. equisimilis strain NCTC5370 | SP_0829 | Phosphopentomutase (EC 5.4.2.7) | 12207705 | 99 | 99 | 86 | 1e-205 |
| Streptococcus dysgalactiae subsp. equisimilis strain NCTC5371 | SP_0829 | Phosphopentomutase (EC 5.4.2.7) | 12207705 | 99 | 99 | 86 | 1e-205 |
| Streptococcus dysgalactiae subsp. equisimilis strain NCTC6179 | SP_0829 | Phosphopentomutase (EC 5.4.2.7) | 12207705 | 99 | 99 | 85 | 1e-203 |
| Streptococcus dysgalactiae subsp. equisimilis strain NCTC6181 | SP_0829 | Phosphopentomutase (EC 5.4.2.7) | 12207705 | 99 | 99 | 85 | 1e-203 |
| Streptococcus dysgalactiae subsp. equisimilis strain NCTC6407 | SP_0829 | Phosphopentomutase (EC 5.4.2.7) | 12207705 | 99 | 99 | 86 | 1e-205 |
| Streptococcus dysgalactiae subsp. equisimilis strain NCTC7136 | SP_0829 | Phosphopentomutase (EC 5.4.2.7) | 12207705 | 99 | 99 | 86 | 1e-205 |
| Streptococcus dysgalactiae subsp. equisimilis strain NCTC8543 | SP_0829 | Phosphopentomutase (EC 5.4.2.7) | 12207705 | 99 | 99 | 86 | 1e-204 |
| Streptococcus dysgalactiae subsp. equisimilis strain NCTC8546 | SP_0829 | Phosphopentomutase (EC 5.4.2.7) | 12207705 | 99 | 99 | 86 | 1e-204 |
| Streptococcus dysgalactiae subsp. equisimilis strain NCTC9413 | SP_0829 | Phosphopentomutase (EC 5.4.2.7) | 12207705 | 99 | 99 | 86 | 1e-205 |
| Streptococcus dysgalactiae subsp. equisimilis strain NCTC9414 | SP_0829 | Phosphopentomutase (EC 5.4.2.7) | 12207705 | 99 | 99 | 86 | 1e-205 |
| Streptococcus dysgalactiae subsp. equisimilis strain NCTC9603 | SP_0829 | Phosphopentomutase (EC 5.4.2.7) | 12207705 | 99 | 99 | 86 | 1e-205 |
| Streptococcus dysgalactiae subsp. equisimilis strain SS1575 | SP_0829 | Phosphopentomutase (EC 5.4.2.7) | 12207705 | 99 | 99 | 86 | 1e-205 |
| Streptococcus dysgalactiae subsp. equisimilis strain T642 | SP_0829 | Phosphopentomutase (EC 5.4.2.7) | 12207705 | 99 | 99 | 86 | 0 |
| Streptococcus dysgalactiae subsp. equisimilis strain UT_4031CC | SP_0829 | Phosphopentomutase (EC 5.4.2.7) | 12207705 | 99 | 99 | 86 | 1e-205 |
| Streptococcus dysgalactiae subsp. equisimilis strain UT_4231_KK | SP_0829 | Phosphopentomutase (EC 5.4.2.7) | 12207705 | 99 | 99 | 86 | 1e-205 |
| Streptococcus dysgalactiae subsp. equisimilis strain UT_4234_DH | SP_0829 | Phosphopentomutase (EC 5.4.2.7) | 12207705 | 99 | 99 | 86 | 1e-204 |
| Streptococcus dysgalactiae subsp. equisimilis strain UT_4241_XS | SP_0829 | Phosphopentomutase (EC 5.4.2.7) | 12207705 | 99 | 99 | 86 | 1e-204 |
| Streptococcus dysgalactiae subsp. equisimilis strain UT_4242_AB | SP_0829 | Phosphopentomutase (EC 5.4.2.7) | 12207705 | 99 | 99 | 86 | 1e-203 |
| Streptococcus dysgalactiae subsp. equisimilis strain UT_4255RC | SP_0829 | Phosphopentomutase (EC 5.4.2.7) | 12207705 | 99 | 99 | 86 | 1e-204 |
| Streptococcus dysgalactiae subsp. equisimilis strain UT_4277_BB | SP_0829 | Phosphopentomutase (EC 5.4.2.7) | 12207705 | 99 | 99 | 86 | 1e-204 |
| Streptococcus dysgalactiae subsp. equisimilis strain UT_4966_RC | SP_0829 | Phosphopentomutase (EC 5.4.2.7) | 12207705 | 99 | 99 | 86 | 1e-204 |
| Streptococcus dysgalactiae subsp. equisimilis strain UT-5345 | SP_0829 | Phosphopentomutase (EC 5.4.2.7) | 12207705 | 99 | 99 | 86 | 1e-205 |
| Streptococcus dysgalactiae subsp. equisimilis strain UT-5354 | SP_0829 | Phosphopentomutase (EC 5.4.2.7) | 12207705 | 99 | 99 | 86 | 1e-204 |
| Streptococcus dysgalactiae subsp. equisimilis strain UT-SS1069 | SP_0829 | Phosphopentomutase (EC 5.4.2.7) | 12207705 | 99 | 99 | 86 | 1e-205 |
| Streptococcus dysgalactiae subsp. equisimilis strain UT-SS957 | SP_0829 | Phosphopentomutase (EC 5.4.2.7) | 12207705 | 99 | 99 | 86 | 1e-205 |
| Streptococcus dysgalactiae subsp. equisimilis strain WCHSDSE-1 | SP_0829 | Phosphopentomutase (EC 5.4.2.7) | 12207705 | 99 | 99 | 86 | 1e-205 |
| Streptococcus dysgalactiae subsp. equisimilis 167 | lgt | Prolipoprotein diacylglyceryl transferase | 17015455 | 99 | 99 | 82 | 1e-129 |
| Streptococcus dysgalactiae subsp. equisimilis AC-2713 | lgt | Prolipoprotein diacylglyceryl transferase | 17015455 | 99 | 99 | 81 | 1e-128 |
| Streptococcus dysgalactiae subsp. equisimilis AKSDE4288 | lgt | Prolipoprotein diacylglyceryl transferase | 17015455 | 99 | 99 | 82 | 1e-129 |
| Streptococcus dysgalactiae subsp. equisimilis ATCC 12394 | lgt | Prolipoprotein diacylglyceryl transferase | 17015455 | 99 | 99 | 81 | 1e-129 |
| Streptococcus dysgalactiae subsp. equisimilis GGS_124 | lgt | Prolipoprotein diacylglyceryl transferase | 17015455 | 99 | 99 | 82 | 1e-129 |
| Streptococcus dysgalactiae subsp. equisimilis RE378 | lgt | Prolipoprotein diacylglyceryl transferase | 17015455 | 99 | 99 | 82 | 1e-129 |
| Streptococcus dysgalactiae subsp. equisimilis SD SCDR1 | lgt | Prolipoprotein diacylglyceryl transferase | 17015455 | 99 | 99 | 82 | 1e-129 |
| Streptococcus dysgalactiae subsp. equisimilis SK1249 | lgt | Prolipoprotein diacylglyceryl transferase | 17015455 | 99 | 99 | 82 | 1e-129 |
| Streptococcus dysgalactiae subsp. equisimilis SK1250 | lgt | Prolipoprotein diacylglyceryl transferase | 17015455 | 99 | 99 | 82 | 1e-129 |
| Streptococcus dysgalactiae subsp. equisimilis strain ASDSE_96 | lgt | Prolipoprotein diacylglyceryl transferase | 17015455 | 99 | 99 | 82 | 1e-129 |
| Streptococcus dysgalactiae subsp. equisimilis strain ASDSE_99 | lgt | Prolipoprotein diacylglyceryl transferase | 17015455 | 99 | 99 | 82 | 1e-129 |
| Streptococcus dysgalactiae subsp. equisimilis strain C161L1 | lgt | Prolipoprotein diacylglyceryl transferase | 17015455 | 100 | 100 | 81 | 0 |
| Streptococcus dysgalactiae subsp. equisimilis strain KNZ01 | lgt | Prolipoprotein diacylglyceryl transferase | 17015455 | 99 | 99 | 82 | 1e-129 |
| Streptococcus dysgalactiae subsp. equisimilis strain KNZ03 | lgt | Prolipoprotein diacylglyceryl transferase | 17015455 | 99 | 99 | 81 | 1e-129 |
| Streptococcus dysgalactiae subsp. equisimilis strain KNZ04 | lgt | Prolipoprotein diacylglyceryl transferase | 17015455 | 99 | 99 | 81 | 1e-129 |
| Streptococcus dysgalactiae subsp. equisimilis strain KNZ06 | lgt | Prolipoprotein diacylglyceryl transferase | 17015455 | 99 | 99 | 82 | 1e-129 |
| Streptococcus dysgalactiae subsp. equisimilis strain KNZ07 | lgt | Prolipoprotein diacylglyceryl transferase | 17015455 | 99 | 99 | 82 | 1e-129 |
| Streptococcus dysgalactiae subsp. equisimilis strain KNZ10 | lgt | Prolipoprotein diacylglyceryl transferase | 17015455 | 99 | 99 | 82 | 1e-129 |
| Streptococcus dysgalactiae subsp. equisimilis strain KNZ12 | lgt | Prolipoprotein diacylglyceryl transferase | 17015455 | 99 | 99 | 82 | 1e-129 |
| Streptococcus dysgalactiae subsp. equisimilis strain KNZ15 | lgt | Prolipoprotein diacylglyceryl transferase | 17015455 | 99 | 99 | 82 | 1e-129 |
| Streptococcus dysgalactiae subsp. equisimilis strain KNZ16 | lgt | Prolipoprotein diacylglyceryl transferase | 17015455 | 99 | 99 | 81 | 1e-129 |
| Streptococcus dysgalactiae subsp. equisimilis strain NCTC10321 | lgt | Prolipoprotein diacylglyceryl transferase | 17015455 | 99 | 99 | 82 | 1e-129 |
| Streptococcus dysgalactiae subsp. equisimilis strain NCTC11554 | lgt | Prolipoprotein diacylglyceryl transferase | 17015455 | 99 | 99 | 81 | 1e-129 |
| Streptococcus dysgalactiae subsp. equisimilis strain NCTC11555 | lgt | Prolipoprotein diacylglyceryl transferase | 17015455 | 99 | 99 | 82 | 1e-129 |
| Streptococcus dysgalactiae subsp. equisimilis strain NCTC11556 | lgt | Prolipoprotein diacylglyceryl transferase | 17015455 | 99 | 99 | 81 | 1e-129 |
| Streptococcus dysgalactiae subsp. equisimilis strain NCTC11557 | lgt | Prolipoprotein diacylglyceryl transferase | 17015455 | 99 | 99 | 82 | 1e-129 |
| Streptococcus dysgalactiae subsp. equisimilis strain NCTC11564 | lgt | Prolipoprotein diacylglyceryl transferase | 17015455 | 99 | 99 | 82 | 1e-129 |
| Streptococcus dysgalactiae subsp. equisimilis strain NCTC5370 | lgt | Prolipoprotein diacylglyceryl transferase | 17015455 | 99 | 99 | 81 | 1e-129 |
| Streptococcus dysgalactiae subsp. equisimilis strain NCTC5371 | lgt | Prolipoprotein diacylglyceryl transferase | 17015455 | 99 | 99 | 81 | 1e-129 |
| Streptococcus dysgalactiae subsp. equisimilis strain NCTC6179 | lgt | Prolipoprotein diacylglyceryl transferase | 17015455 | 99 | 99 | 81 | 1e-128 |
| Streptococcus dysgalactiae subsp. equisimilis strain NCTC6181 | lgt | Prolipoprotein diacylglyceryl transferase | 17015455 | 99 | 99 | 81 | 1e-128 |
| Streptococcus dysgalactiae subsp. equisimilis strain NCTC6407 | lgt | Prolipoprotein diacylglyceryl transferase | 17015455 | 99 | 99 | 82 | 1e-129 |
| Streptococcus dysgalactiae subsp. equisimilis strain NCTC7136 | lgt | Prolipoprotein diacylglyceryl transferase | 17015455 | 99 | 99 | 82 | 1e-129 |
| Streptococcus dysgalactiae subsp. equisimilis strain NCTC8543 | lgt | Prolipoprotein diacylglyceryl transferase | 17015455 | 99 | 99 | 81 | 1e-129 |
| Streptococcus dysgalactiae subsp. equisimilis strain NCTC8546 | lgt | Prolipoprotein diacylglyceryl transferase | 17015455 | 99 | 99 | 82 | 1e-129 |
| Streptococcus dysgalactiae subsp. equisimilis strain NCTC9413 | lgt | Prolipoprotein diacylglyceryl transferase | 17015455 | 99 | 99 | 82 | 1e-129 |
| Streptococcus dysgalactiae subsp. equisimilis strain NCTC9414 | lgt | Prolipoprotein diacylglyceryl transferase | 17015455 | 99 | 99 | 81 | 1e-129 |
| Streptococcus dysgalactiae subsp. equisimilis strain NCTC9603 | lgt | Prolipoprotein diacylglyceryl transferase | 17015455 | 99 | 99 | 82 | 1e-129 |
| Streptococcus dysgalactiae subsp. equisimilis strain SS1575 | lgt | Prolipoprotein diacylglyceryl transferase | 17015455 | 99 | 99 | 82 | 1e-129 |
| Streptococcus dysgalactiae subsp. equisimilis strain T642 | lgt | Prolipoprotein diacylglyceryl transferase | 17015455 | 100 | 100 | 81 | 0 |
| Streptococcus dysgalactiae subsp. equisimilis strain UT_4031CC | lgt | Prolipoprotein diacylglyceryl transferase | 17015455 | 99 | 99 | 82 | 1e-129 |
| Streptococcus dysgalactiae subsp. equisimilis strain UT_4231_KK | lgt | Prolipoprotein diacylglyceryl transferase | 17015455 | 99 | 99 | 82 | 1e-129 |
| Streptococcus dysgalactiae subsp. equisimilis strain UT_4234_DH | lgt | Prolipoprotein diacylglyceryl transferase | 17015455 | 99 | 99 | 82 | 1e-129 |
| Streptococcus dysgalactiae subsp. equisimilis strain UT_4241_XS | lgt | Prolipoprotein diacylglyceryl transferase | 17015455 | 99 | 99 | 82 | 1e-129 |
| Streptococcus dysgalactiae subsp. equisimilis strain UT_4242_AB | lgt | Prolipoprotein diacylglyceryl transferase | 17015455 | 99 | 99 | 81 | 1e-129 |
| Streptococcus dysgalactiae subsp. equisimilis strain UT_4255RC | lgt | Prolipoprotein diacylglyceryl transferase | 17015455 | 99 | 99 | 82 | 1e-129 |
| Streptococcus dysgalactiae subsp. equisimilis strain UT_4277_BB | lgt | Prolipoprotein diacylglyceryl transferase | 17015455 | 99 | 99 | 82 | 1e-129 |
| Streptococcus dysgalactiae subsp. equisimilis strain UT_4966_RC | lgt | Prolipoprotein diacylglyceryl transferase | 17015455 | 99 | 99 | 82 | 1e-129 |
| Streptococcus dysgalactiae subsp. equisimilis strain UT-5345 | lgt | Prolipoprotein diacylglyceryl transferase | 17015455 | 99 | 99 | 82 | 1e-129 |
| Streptococcus dysgalactiae subsp. equisimilis strain UT-5354 | lgt | Prolipoprotein diacylglyceryl transferase | 17015455 | 99 | 99 | 81 | 1e-129 |
| Streptococcus dysgalactiae subsp. equisimilis strain UT-SS1069 | lgt | Prolipoprotein diacylglyceryl transferase | 17015455 | 99 | 99 | 82 | 1e-129 |
| Streptococcus dysgalactiae subsp. equisimilis strain UT-SS957 | lgt | Prolipoprotein diacylglyceryl transferase | 17015455 | 99 | 99 | 82 | 1e-129 |
| Streptococcus dysgalactiae subsp. equisimilis strain WCHSDSE-1 | lgt | Prolipoprotein diacylglyceryl transferase | 17015455 | 99 | 99 | 82 | 1e-129 |
| Streptococcus dysgalactiae subsp. equisimilis 167 | SP_0251 | Pyruvate formate-lyase (EC 2.3.1.54) | 12207705 | 92 | 93 | 87 | 0.0 |
| Streptococcus dysgalactiae subsp. equisimilis AC-2713 | pflD | Pyruvate formate-lyase (EC 2.3.1.54) | 12207705 | 92 | 93 | 87 | 0.0 |
| Streptococcus dysgalactiae subsp. equisimilis AKSDE4288 | SP_0251 | Pyruvate formate-lyase (EC 2.3.1.54) | 12207705 | 92 | 93 | 87 | 0.0 |
| Streptococcus dysgalactiae subsp. equisimilis ATCC 12394 | SP_0251 | Pyruvate formate-lyase (EC 2.3.1.54) | 12207705 | 92 | 93 | 87 | 0.0 |
| Streptococcus dysgalactiae subsp. equisimilis GGS_124 | SP_0251 | Pyruvate formate-lyase (EC 2.3.1.54) | 12207705 | 92 | 93 | 87 | 0.0 |
| Streptococcus dysgalactiae subsp. equisimilis RE378 | pflD | Pyruvate formate-lyase (EC 2.3.1.54) | 12207705 | 92 | 93 | 87 | 0.0 |
| Streptococcus dysgalactiae subsp. equisimilis SD SCDR1 | SP_0251 | Pyruvate formate-lyase (EC 2.3.1.54) | 12207705 | 92 | 93 | 86 | 0.0 |
| Streptococcus dysgalactiae subsp. equisimilis SK1249 | SP_0251 | Pyruvate formate-lyase (EC 2.3.1.54) | 12207705 | 92 | 93 | 87 | 0.0 |
| Streptococcus dysgalactiae subsp. equisimilis SK1250 | SP_0251 | Pyruvate formate-lyase (EC 2.3.1.54) | 12207705 | 92 | 93 | 86 | 0.0 |
| Streptococcus dysgalactiae subsp. equisimilis strain ASDSE_96 | SP_0251 | Pyruvate formate-lyase (EC 2.3.1.54) | 12207705 | 92 | 93 | 87 | 0.0 |
| Streptococcus dysgalactiae subsp. equisimilis strain ASDSE_99 | SP_0251 | Pyruvate formate-lyase (EC 2.3.1.54) | 12207705 | 92 | 93 | 87 | 0.0 |
| Streptococcus dysgalactiae subsp. equisimilis strain C161L1 | SP_0251 | Pyruvate formate-lyase (EC 2.3.1.54) | 12207705 | 99 | 99 | 85 | 0 |
| Streptococcus dysgalactiae subsp. equisimilis strain KNZ01 | SP_0251 | Pyruvate formate-lyase (EC 2.3.1.54) | 12207705 | 92 | 93 | 87 | 0.0 |
| Streptococcus dysgalactiae subsp. equisimilis strain KNZ03 | SP_0251 | Pyruvate formate-lyase (EC 2.3.1.54) | 12207705 | 92 | 93 | 87 | 0.0 |
| Streptococcus dysgalactiae subsp. equisimilis strain KNZ04 | SP_0251 | Pyruvate formate-lyase (EC 2.3.1.54) | 12207705 | 92 | 93 | 87 | 0.0 |
| Streptococcus dysgalactiae subsp. equisimilis strain KNZ06 | SP_0251 | Pyruvate formate-lyase (EC 2.3.1.54) | 12207705 | 92 | 93 | 87 | 0.0 |
| Streptococcus dysgalactiae subsp. equisimilis strain KNZ07 | SP_0251 | Pyruvate formate-lyase (EC 2.3.1.54) | 12207705 | 92 | 93 | 87 | 0.0 |
| Streptococcus dysgalactiae subsp. equisimilis strain KNZ10 | SP_0251 | Pyruvate formate-lyase (EC 2.3.1.54) | 12207705 | 92 | 93 | 87 | 0.0 |
| Streptococcus dysgalactiae subsp. equisimilis strain KNZ12 | SP_0251 | Pyruvate formate-lyase (EC 2.3.1.54) | 12207705 | 92 | 93 | 87 | 0.0 |
| Streptococcus dysgalactiae subsp. equisimilis strain KNZ15 | SP_0251 | Pyruvate formate-lyase (EC 2.3.1.54) | 12207705 | 92 | 93 | 87 | 0.0 |
| Streptococcus dysgalactiae subsp. equisimilis strain KNZ16 | SP_0251 | Pyruvate formate-lyase (EC 2.3.1.54) | 12207705 | 92 | 93 | 87 | 0.0 |
| Streptococcus dysgalactiae subsp. equisimilis strain NCTC10321 | SP_0251 | Pyruvate formate-lyase (EC 2.3.1.54) | 12207705 | 92 | 93 | 86 | 0.0 |
| Streptococcus dysgalactiae subsp. equisimilis strain NCTC11554 | SP_0251 | Pyruvate formate-lyase (EC 2.3.1.54) | 12207705 | 92 | 93 | 87 | 0.0 |
| Streptococcus dysgalactiae subsp. equisimilis strain NCTC11555 | SP_0251 | Pyruvate formate-lyase (EC 2.3.1.54) | 12207705 | 92 | 93 | 87 | 0.0 |
| Streptococcus dysgalactiae subsp. equisimilis strain NCTC11556 | SP_0251 | Pyruvate formate-lyase (EC 2.3.1.54) | 12207705 | 92 | 93 | 87 | 0.0 |
| Streptococcus dysgalactiae subsp. equisimilis strain NCTC11557 | SP_0251 | Pyruvate formate-lyase (EC 2.3.1.54) | 12207705 | 92 | 93 | 87 | 0.0 |
| Streptococcus dysgalactiae subsp. equisimilis strain NCTC11564 | SP_0251 | Pyruvate formate-lyase (EC 2.3.1.54) | 12207705 | 92 | 93 | 87 | 0.0 |
| Streptococcus dysgalactiae subsp. equisimilis strain NCTC5370 | SP_0251 | Pyruvate formate-lyase (EC 2.3.1.54) | 12207705 | 92 | 93 | 87 | 0.0 |
| Streptococcus dysgalactiae subsp. equisimilis strain NCTC5371 | SP_0251 | Pyruvate formate-lyase (EC 2.3.1.54) | 12207705 | 92 | 93 | 87 | 0.0 |
| Streptococcus dysgalactiae subsp. equisimilis strain NCTC6179 | SP_0251 | Pyruvate formate-lyase (EC 2.3.1.54) | 12207705 | 92 | 93 | 87 | 0.0 |
| Streptococcus dysgalactiae subsp. equisimilis strain NCTC6181 | SP_0251 | Pyruvate formate-lyase (EC 2.3.1.54) | 12207705 | 92 | 93 | 87 | 0.0 |
| Streptococcus dysgalactiae subsp. equisimilis strain NCTC6407 | SP_0251 | Pyruvate formate-lyase (EC 2.3.1.54) | 12207705 | 92 | 93 | 86 | 0.0 |
| Streptococcus dysgalactiae subsp. equisimilis strain NCTC7136 | SP_0251 | Pyruvate formate-lyase (EC 2.3.1.54) | 12207705 | 92 | 93 | 87 | 0.0 |
| Streptococcus dysgalactiae subsp. equisimilis strain NCTC8543 | SP_0251 | Pyruvate formate-lyase (EC 2.3.1.54) | 12207705 | 92 | 93 | 86 | 0.0 |
| Streptococcus dysgalactiae subsp. equisimilis strain NCTC8546 | SP_0251 | Pyruvate formate-lyase (EC 2.3.1.54) | 12207705 | 92 | 93 | 86 | 0.0 |
| Streptococcus dysgalactiae subsp. equisimilis strain NCTC9413 | SP_0251 | Pyruvate formate-lyase (EC 2.3.1.54) | 12207705 | 92 | 93 | 87 | 0.0 |
| Streptococcus dysgalactiae subsp. equisimilis strain NCTC9414 | SP_0251 | Pyruvate formate-lyase (EC 2.3.1.54) | 12207705 | 92 | 93 | 87 | 0.0 |
| Streptococcus dysgalactiae subsp. equisimilis strain NCTC9603 | SP_0251 | Pyruvate formate-lyase (EC 2.3.1.54) | 12207705 | 92 | 93 | 87 | 0.0 |
| Streptococcus dysgalactiae subsp. equisimilis strain SS1575 | SP_0251 | Pyruvate formate-lyase (EC 2.3.1.54) | 12207705 | 92 | 93 | 86 | 0.0 |
| Streptococcus dysgalactiae subsp. equisimilis strain T642 | SP_0251 | Pyruvate formate-lyase (EC 2.3.1.54) | 12207705 | 99 | 99 | 85 | 0.0 |
| Streptococcus dysgalactiae subsp. equisimilis strain UT_4031CC | SP_0251 | Pyruvate formate-lyase (EC 2.3.1.54) | 12207705 | 92 | 93 | 86 | 0.0 |
| Streptococcus dysgalactiae subsp. equisimilis strain UT_4231_KK | SP_0251 | Pyruvate formate-lyase (EC 2.3.1.54) | 12207705 | 92 | 93 | 86 | 0.0 |
| Streptococcus dysgalactiae subsp. equisimilis strain UT_4234_DH | SP_0251 | Pyruvate formate-lyase (EC 2.3.1.54) | 12207705 | 92 | 93 | 87 | 0.0 |
| Streptococcus dysgalactiae subsp. equisimilis strain UT_4241_XS | SP_0251 | Pyruvate formate-lyase (EC 2.3.1.54) | 12207705 | 92 | 93 | 87 | 0.0 |
| Streptococcus dysgalactiae subsp. equisimilis strain UT_4242_AB | SP_0251 | Pyruvate formate-lyase (EC 2.3.1.54) | 12207705 | 92 | 93 | 87 | 0.0 |
| Streptococcus dysgalactiae subsp. equisimilis strain UT_4255RC | SP_0251 | Pyruvate formate-lyase (EC 2.3.1.54) | 12207705 | 92 | 93 | 87 | 0.0 |
| Streptococcus dysgalactiae subsp. equisimilis strain UT_4277_BB | SP_0251 | Pyruvate formate-lyase (EC 2.3.1.54) | 12207705 | 92 | 93 | 87 | 0.0 |
| Streptococcus dysgalactiae subsp. equisimilis strain UT_4966_RC | SP_0251 | Pyruvate formate-lyase (EC 2.3.1.54) | 12207705 | 92 | 93 | 87 | 0.0 |
| Streptococcus dysgalactiae subsp. equisimilis strain UT-5345 | SP_0251 | Pyruvate formate-lyase (EC 2.3.1.54) | 12207705 | 92 | 93 | 86 | 0.0 |
| Streptococcus dysgalactiae subsp. equisimilis strain UT-5354 | SP_0251 | Pyruvate formate-lyase (EC 2.3.1.54) | 12207705 | 92 | 93 | 87 | 0.0 |
| Streptococcus dysgalactiae subsp. equisimilis strain UT-SS1069 | SP_0251 | Pyruvate formate-lyase (EC 2.3.1.54) | 12207705 | 92 | 93 | 87 | 0.0 |
| Streptococcus dysgalactiae subsp. equisimilis strain UT-SS957 | SP_0251 | Pyruvate formate-lyase (EC 2.3.1.54) | 12207705 | 92 | 93 | 87 | 0.0 |
| Streptococcus dysgalactiae subsp. equisimilis strain WCHSDSE-1 | SP_0251 | Pyruvate formate-lyase (EC 2.3.1.54) | 12207705 | 92 | 93 | 86 | 0.0 |
| Streptococcus dysgalactiae subsp. equisimilis 167 | SP_0095 | Rhodanese domain protein UPF0176, Firmicutes subgroup | 12207705 | 65 | 87 | 90 | 1e-116 |
| Streptococcus dysgalactiae subsp. equisimilis AC-2713 | SP_0095 | Rhodanese domain protein UPF0176, Firmicutes subgroup | 12207705 | 90 | 90 | 87 | 1e-161 |
| Streptococcus dysgalactiae subsp. equisimilis AKSDE4288 | SP_0095 | Rhodanese domain protein UPF0176, Firmicutes subgroup | 12207705 | 90 | 90 | 87 | 1e-161 |
| Streptococcus dysgalactiae subsp. equisimilis ATCC 12394 | SP_0095 | Rhodanese domain protein UPF0176, Firmicutes subgroup | 12207705 | 90 | 90 | 87 | 1e-161 |
| Streptococcus dysgalactiae subsp. equisimilis GGS_124 | SP_0095 | Rhodanese domain protein UPF0176, Firmicutes subgroup | 12207705 | 90 | 90 | 87 | 1e-161 |
| Streptococcus dysgalactiae subsp. equisimilis RE378 | SP_0095 | Rhodanese domain protein UPF0176, Firmicutes subgroup | 12207705 | 90 | 90 | 87 | 1e-161 |
| Streptococcus dysgalactiae subsp. equisimilis SD SCDR1 | SP_0095 | Rhodanese domain protein UPF0176, Firmicutes subgroup | 12207705 | 90 | 90 | 87 | 1e-161 |
| Streptococcus dysgalactiae subsp. equisimilis SK1249 | SP_0095 | Rhodanese domain protein UPF0176, Firmicutes subgroup | 12207705 | 90 | 90 | 87 | 1e-161 |
| Streptococcus dysgalactiae subsp. equisimilis SK1250 | SP_0095 | Rhodanese domain protein UPF0176, Firmicutes subgroup | 12207705 | 90 | 90 | 87 | 1e-161 |
| Streptococcus dysgalactiae subsp. equisimilis strain ASDSE_96 | SP_0095 | Rhodanese domain protein UPF0176, Firmicutes subgroup | 12207705 | 90 | 90 | 87 | 1e-161 |
| Streptococcus dysgalactiae subsp. equisimilis strain ASDSE_99 | SP_0095 | Rhodanese domain protein UPF0176, Firmicutes subgroup | 12207705 | 90 | 90 | 87 | 1e-161 |
| Streptococcus dysgalactiae subsp. equisimilis strain C161L1 | SP_0095 | Rhodanese domain protein UPF0176, Firmicutes subgroup | 12207705 | 98 | 98 | 83 | 0 |
| Streptococcus dysgalactiae subsp. equisimilis strain KNZ01 | SP_0095 | Rhodanese domain protein UPF0176, Firmicutes subgroup | 12207705 | 90 | 90 | 87 | 1e-161 |
| Streptococcus dysgalactiae subsp. equisimilis strain KNZ03 | SP_0095 | Rhodanese domain protein UPF0176, Firmicutes subgroup | 12207705 | 90 | 90 | 87 | 1e-161 |
| Streptococcus dysgalactiae subsp. equisimilis strain KNZ04 | SP_0095 | Rhodanese domain protein UPF0176, Firmicutes subgroup | 12207705 | 90 | 90 | 87 | 1e-161 |
| Streptococcus dysgalactiae subsp. equisimilis strain KNZ06 | SP_0095 | Rhodanese domain protein UPF0176, Firmicutes subgroup | 12207705 | 90 | 90 | 87 | 1e-161 |
| Streptococcus dysgalactiae subsp. equisimilis strain KNZ07 | SP_0095 | Rhodanese domain protein UPF0176, Firmicutes subgroup | 12207705 | 90 | 90 | 87 | 1e-161 |
| Streptococcus dysgalactiae subsp. equisimilis strain KNZ10 | SP_0095 | Rhodanese domain protein UPF0176, Firmicutes subgroup | 12207705 | 90 | 90 | 87 | 1e-161 |
| Streptococcus dysgalactiae subsp. equisimilis strain KNZ12 | SP_0095 | Rhodanese domain protein UPF0176, Firmicutes subgroup | 12207705 | 90 | 90 | 87 | 1e-161 |
| Streptococcus dysgalactiae subsp. equisimilis strain KNZ15 | SP_0095 | Rhodanese domain protein UPF0176, Firmicutes subgroup | 12207705 | 90 | 90 | 87 | 1e-161 |
| Streptococcus dysgalactiae subsp. equisimilis strain KNZ16 | SP_0095 | Rhodanese domain protein UPF0176, Firmicutes subgroup | 12207705 | 90 | 90 | 87 | 1e-161 |
| Streptococcus dysgalactiae subsp. equisimilis strain NCTC10321 | SP_0095 | Rhodanese domain protein UPF0176, Firmicutes subgroup | 12207705 | 96 | 96 | 84 | 1e-166 |
| Streptococcus dysgalactiae subsp. equisimilis strain NCTC11554 | SP_0095 | Rhodanese domain protein UPF0176, Firmicutes subgroup | 12207705 | 90 | 90 | 87 | 1e-161 |
| Streptococcus dysgalactiae subsp. equisimilis strain NCTC11555 | SP_0095 | Rhodanese domain protein UPF0176, Firmicutes subgroup | 12207705 | 90 | 90 | 87 | 1e-161 |
| Streptococcus dysgalactiae subsp. equisimilis strain NCTC11556 | SP_0095 | Rhodanese domain protein UPF0176, Firmicutes subgroup | 12207705 | 90 | 90 | 86 | 1e-160 |
| Streptococcus dysgalactiae subsp. equisimilis strain NCTC11557 | SP_0095 | Rhodanese domain protein UPF0176, Firmicutes subgroup | 12207705 | 90 | 90 | 87 | 1e-161 |
| Streptococcus dysgalactiae subsp. equisimilis strain NCTC11564 | SP_0095 | Rhodanese domain protein UPF0176, Firmicutes subgroup | 12207705 | 90 | 90 | 87 | 1e-161 |
| Streptococcus dysgalactiae subsp. equisimilis strain NCTC5370 | SP_0095 | Rhodanese domain protein UPF0176, Firmicutes subgroup | 12207705 | 90 | 90 | 87 | 1e-161 |
| Streptococcus dysgalactiae subsp. equisimilis strain NCTC5371 | SP_0095 | Rhodanese domain protein UPF0176, Firmicutes subgroup | 12207705 | 90 | 90 | 87 | 1e-161 |
| Streptococcus dysgalactiae subsp. equisimilis strain NCTC6179 | SP_0095 | Rhodanese domain protein UPF0176, Firmicutes subgroup | 12207705 | 96 | 96 | 84 | 1e-166 |
| Streptococcus dysgalactiae subsp. equisimilis strain NCTC6181 | SP_0095 | Rhodanese domain protein UPF0176, Firmicutes subgroup | 12207705 | 96 | 96 | 84 | 1e-166 |
| Streptococcus dysgalactiae subsp. equisimilis strain NCTC6407 | SP_0095 | Rhodanese domain protein UPF0176, Firmicutes subgroup | 12207705 | 96 | 96 | 84 | 1e-166 |
| Streptococcus dysgalactiae subsp. equisimilis strain NCTC7136 | SP_0095 | Rhodanese domain protein UPF0176, Firmicutes subgroup | 12207705 | 90 | 90 | 87 | 1e-161 |
| Streptococcus dysgalactiae subsp. equisimilis strain NCTC8543 | SP_0095 | Rhodanese domain protein UPF0176, Firmicutes subgroup | 12207705 | 90 | 90 | 87 | 1e-161 |
| Streptococcus dysgalactiae subsp. equisimilis strain NCTC8546 | SP_0095 | Rhodanese domain protein UPF0176, Firmicutes subgroup | 12207705 | 90 | 90 | 87 | 1e-161 |
| Streptococcus dysgalactiae subsp. equisimilis strain NCTC9413 | SP_0095 | Rhodanese domain protein UPF0176, Firmicutes subgroup | 12207705 | 90 | 90 | 87 | 1e-161 |
| Streptococcus dysgalactiae subsp. equisimilis strain NCTC9414 | SP_0095 | Rhodanese domain protein UPF0176, Firmicutes subgroup | 12207705 | 90 | 90 | 87 | 1e-161 |
| Streptococcus dysgalactiae subsp. equisimilis strain NCTC9603 | SP_0095 | Rhodanese domain protein UPF0176, Firmicutes subgroup | 12207705 | 90 | 90 | 87 | 1e-161 |
| Streptococcus dysgalactiae subsp. equisimilis strain SS1575 | SP_0095 | Rhodanese domain protein UPF0176, Firmicutes subgroup | 12207705 | 90 | 90 | 86 | 1e-160 |
| Streptococcus dysgalactiae subsp. equisimilis strain T642 | SP_0095 | Rhodanese domain protein UPF0176, Firmicutes subgroup | 12207705 | 98 | 98 | 83 | 0 |
| Streptococcus dysgalactiae subsp. equisimilis strain UT_4031CC | SP_0095 | Rhodanese domain protein UPF0176, Firmicutes subgroup | 12207705 | 90 | 90 | 87 | 1e-161 |
| Streptococcus dysgalactiae subsp. equisimilis strain UT_4231_KK | SP_0095 | Rhodanese domain protein UPF0176, Firmicutes subgroup | 12207705 | 90 | 90 | 87 | 1e-161 |
| Streptococcus dysgalactiae subsp. equisimilis strain UT_4234_DH | SP_0095 | Rhodanese domain protein UPF0176, Firmicutes subgroup | 12207705 | 90 | 90 | 87 | 1e-161 |
| Streptococcus dysgalactiae subsp. equisimilis strain UT_4241_XS | SP_0095 | Rhodanese domain protein UPF0176, Firmicutes subgroup | 12207705 | 90 | 90 | 87 | 1e-161 |
| Streptococcus dysgalactiae subsp. equisimilis strain UT_4242_AB | SP_0095 | Rhodanese domain protein UPF0176, Firmicutes subgroup | 12207705 | 90 | 90 | 86 | 1e-160 |
| Streptococcus dysgalactiae subsp. equisimilis strain UT_4255RC | SP_0095 | Rhodanese domain protein UPF0176, Firmicutes subgroup | 12207705 | 90 | 90 | 87 | 1e-161 |
| Streptococcus dysgalactiae subsp. equisimilis strain UT_4277_BB | SP_0095 | Rhodanese domain protein UPF0176, Firmicutes subgroup | 12207705 | 90 | 90 | 87 | 1e-161 |
| Streptococcus dysgalactiae subsp. equisimilis strain UT_4966_RC | SP_0095 | Rhodanese domain protein UPF0176, Firmicutes subgroup | 12207705 | 90 | 90 | 87 | 1e-161 |
| Streptococcus dysgalactiae subsp. equisimilis strain UT-5345 | SP_0095 | Rhodanese domain protein UPF0176, Firmicutes subgroup | 12207705 | 90 | 90 | 87 | 1e-161 |
| Streptococcus dysgalactiae subsp. equisimilis strain UT-5354 | SP_0095 | Rhodanese domain protein UPF0176, Firmicutes subgroup | 12207705 | 90 | 90 | 87 | 1e-161 |
| Streptococcus dysgalactiae subsp. equisimilis strain UT-SS1069 | SP_0095 | Rhodanese domain protein UPF0176, Firmicutes subgroup | 12207705 | 90 | 90 | 87 | 1e-161 |
| Streptococcus dysgalactiae subsp. equisimilis strain UT-SS957 | SP_0095 | Rhodanese domain protein UPF0176, Firmicutes subgroup | 12207705 | 90 | 90 | 87 | 1e-161 |
| Streptococcus dysgalactiae subsp. equisimilis strain WCHSDSE-1 | SP_0095 | Rhodanese domain protein UPF0176, Firmicutes subgroup | 12207705 | 90 | 90 | 87 | 1e-161 |
| Streptococcus dysgalactiae subsp. equisimilis 167 | SP_0121 | Ribonuclease J1 (endonuclease and 5' exonuclease) | 12207705 | 28 | 98 | 87 | 7e-78 |
| Streptococcus dysgalactiae subsp. equisimilis AC-2713 | SP_0121 | Ribonuclease J1 (endonuclease and 5' exonuclease) | 12207705 | 98 | 98 | 90 | 1e-297 |
| Streptococcus dysgalactiae subsp. equisimilis AKSDE4288 | SP_0121 | Ribonuclease J1 (endonuclease and 5' exonuclease) | 12207705 | 98 | 98 | 90 | 1e-297 |
| Streptococcus dysgalactiae subsp. equisimilis ATCC 12394 | SP_0121 | Ribonuclease J1 (endonuclease and 5' exonuclease) | 12207705 | 98 | 98 | 90 | 1e-297 |
| Streptococcus dysgalactiae subsp. equisimilis GGS_124 | SP_0121 | Ribonuclease J1 (endonuclease and 5' exonuclease) | 12207705 | 98 | 98 | 90 | 1e-297 |
| Streptococcus dysgalactiae subsp. equisimilis RE378 | ykqC | Ribonuclease J1 (endonuclease and 5' exonuclease) | 12207705 | 98 | 98 | 90 | 1e-297 |
| Streptococcus dysgalactiae subsp. equisimilis SD SCDR1 | SP_0121 | Ribonuclease J1 (endonuclease and 5' exonuclease) | 12207705 | 98 | 98 | 90 | 1e-297 |
| Streptococcus dysgalactiae subsp. equisimilis SK1249 | rnjA | Ribonuclease J1 (endonuclease and 5' exonuclease) | 12207705 | 98 | 98 | 90 | 1e-297 |
| Streptococcus dysgalactiae subsp. equisimilis SK1250 | SP_0121 | Ribonuclease J1 (endonuclease and 5' exonuclease) | 12207705 | 98 | 98 | 90 | 1e-297 |
| Streptococcus dysgalactiae subsp. equisimilis strain ASDSE_96 | SP_0121 | Ribonuclease J1 (endonuclease and 5' exonuclease) | 12207705 | 98 | 98 | 90 | 1e-297 |
| Streptococcus dysgalactiae subsp. equisimilis strain ASDSE_99 | SP_0121 | Ribonuclease J1 (endonuclease and 5' exonuclease) | 12207705 | 98 | 98 | 90 | 1e-297 |
| Streptococcus dysgalactiae subsp. equisimilis strain C161L1 | SP_0121 | Ribonuclease J1 (endonuclease and 5' exonuclease) | 12207705 | 99 | 99 | 89 | 0 |
| Streptococcus dysgalactiae subsp. equisimilis strain KNZ01 | SP_0121 | Ribonuclease J1 (endonuclease and 5' exonuclease) | 12207705 | 98 | 98 | 90 | 1e-297 |
| Streptococcus dysgalactiae subsp. equisimilis strain KNZ03 | SP_0121 | Ribonuclease J1 (endonuclease and 5' exonuclease) | 12207705 | 98 | 98 | 90 | 1e-297 |
| Streptococcus dysgalactiae subsp. equisimilis strain KNZ04 | SP_0121 | Ribonuclease J1 (endonuclease and 5' exonuclease) | 12207705 | 98 | 98 | 90 | 1e-297 |
| Streptococcus dysgalactiae subsp. equisimilis strain KNZ06 | SP_0121 | Ribonuclease J1 (endonuclease and 5' exonuclease) | 12207705 | 98 | 98 | 90 | 1e-297 |
| Streptococcus dysgalactiae subsp. equisimilis strain KNZ07 | SP_0121 | Ribonuclease J1 (endonuclease and 5' exonuclease) | 12207705 | 98 | 98 | 90 | 1e-297 |
| Streptococcus dysgalactiae subsp. equisimilis strain KNZ10 | SP_0121 | Ribonuclease J1 (endonuclease and 5' exonuclease) | 12207705 | 98 | 98 | 90 | 1e-297 |
| Streptococcus dysgalactiae subsp. equisimilis strain KNZ12 | SP_0121 | Ribonuclease J1 (endonuclease and 5' exonuclease) | 12207705 | 98 | 98 | 90 | 1e-297 |
| Streptococcus dysgalactiae subsp. equisimilis strain KNZ15 | SP_0121 | Ribonuclease J1 (endonuclease and 5' exonuclease) | 12207705 | 98 | 98 | 90 | 1e-297 |
| Streptococcus dysgalactiae subsp. equisimilis strain KNZ16 | SP_0121 | Ribonuclease J1 (endonuclease and 5' exonuclease) | 12207705 | 98 | 98 | 90 | 1e-297 |
| Streptococcus dysgalactiae subsp. equisimilis strain NCTC10321 | SP_0121 | Ribonuclease J1 (endonuclease and 5' exonuclease) | 12207705 | 98 | 98 | 90 | 1e-297 |
| Streptococcus dysgalactiae subsp. equisimilis strain NCTC11554 | SP_0121 | Ribonuclease J1 (endonuclease and 5' exonuclease) | 12207705 | 98 | 98 | 90 | 1e-297 |
| Streptococcus dysgalactiae subsp. equisimilis strain NCTC11555 | SP_0121 | Ribonuclease J1 (endonuclease and 5' exonuclease) | 12207705 | 98 | 98 | 90 | 1e-297 |
| Streptococcus dysgalactiae subsp. equisimilis strain NCTC11556 | SP_0121 | Ribonuclease J1 (endonuclease and 5' exonuclease) | 12207705 | 98 | 98 | 90 | 1e-297 |
| Streptococcus dysgalactiae subsp. equisimilis strain NCTC11557 | SP_0121 | Ribonuclease J1 (endonuclease and 5' exonuclease) | 12207705 | 98 | 98 | 90 | 1e-297 |
| Streptococcus dysgalactiae subsp. equisimilis strain NCTC11564 | SP_0121 | Ribonuclease J1 (endonuclease and 5' exonuclease) | 12207705 | 98 | 98 | 90 | 1e-297 |
| Streptococcus dysgalactiae subsp. equisimilis strain NCTC5370 | SP_0121 | Ribonuclease J1 (endonuclease and 5' exonuclease) | 12207705 | 98 | 98 | 90 | 1e-297 |
| Streptococcus dysgalactiae subsp. equisimilis strain NCTC5371 | SP_0121 | Ribonuclease J1 (endonuclease and 5' exonuclease) | 12207705 | 98 | 98 | 90 | 1e-297 |
| Streptococcus dysgalactiae subsp. equisimilis strain NCTC6179 | SP_0121 | Ribonuclease J1 (endonuclease and 5' exonuclease) | 12207705 | 98 | 98 | 90 | 1e-297 |
| Streptococcus dysgalactiae subsp. equisimilis strain NCTC6181 | SP_0121 | Ribonuclease J1 (endonuclease and 5' exonuclease) | 12207705 | 98 | 98 | 90 | 1e-297 |
| Streptococcus dysgalactiae subsp. equisimilis strain NCTC6407 | SP_0121 | Ribonuclease J1 (endonuclease and 5' exonuclease) | 12207705 | 98 | 98 | 90 | 1e-297 |
| Streptococcus dysgalactiae subsp. equisimilis strain NCTC7136 | SP_0121 | Ribonuclease J1 (endonuclease and 5' exonuclease) | 12207705 | 98 | 98 | 90 | 1e-297 |
| Streptococcus dysgalactiae subsp. equisimilis strain NCTC8543 | SP_0121 | Ribonuclease J1 (endonuclease and 5' exonuclease) | 12207705 | 98 | 98 | 90 | 1e-297 |
| Streptococcus dysgalactiae subsp. equisimilis strain NCTC8546 | SP_0121 | Ribonuclease J1 (endonuclease and 5' exonuclease) | 12207705 | 98 | 98 | 90 | 1e-297 |
| Streptococcus dysgalactiae subsp. equisimilis strain NCTC9413 | SP_0121 | Ribonuclease J1 (endonuclease and 5' exonuclease) | 12207705 | 98 | 98 | 90 | 1e-297 |
| Streptococcus dysgalactiae subsp. equisimilis strain NCTC9414 | SP_0121 | Ribonuclease J1 (endonuclease and 5' exonuclease) | 12207705 | 98 | 98 | 90 | 1e-297 |
| Streptococcus dysgalactiae subsp. equisimilis strain NCTC9603 | SP_0121 | Ribonuclease J1 (endonuclease and 5' exonuclease) | 12207705 | 98 | 98 | 90 | 1e-296 |
| Streptococcus dysgalactiae subsp. equisimilis strain SS1575 | SP_0121 | Ribonuclease J1 (endonuclease and 5' exonuclease) | 12207705 | 98 | 98 | 90 | 1e-297 |
| Streptococcus dysgalactiae subsp. equisimilis strain T642 | SP_0121 | Ribonuclease J1 (endonuclease and 5' exonuclease) | 12207705 | 99 | 99 | 89 | 0 |
| Streptococcus dysgalactiae subsp. equisimilis strain UT_4031CC | SP_0121 | Ribonuclease J1 (endonuclease and 5' exonuclease) | 12207705 | 98 | 98 | 90 | 1e-297 |
| Streptococcus dysgalactiae subsp. equisimilis strain UT_4231_KK | SP_0121 | Ribonuclease J1 (endonuclease and 5' exonuclease) | 12207705 | 98 | 98 | 90 | 1e-297 |
| Streptococcus dysgalactiae subsp. equisimilis strain UT_4234_DH | SP_0121 | Ribonuclease J1 (endonuclease and 5' exonuclease) | 12207705 | 98 | 98 | 90 | 1e-297 |
| Streptococcus dysgalactiae subsp. equisimilis strain UT_4241_XS | SP_0121 | Ribonuclease J1 (endonuclease and 5' exonuclease) | 12207705 | 98 | 98 | 90 | 1e-297 |
| Streptococcus dysgalactiae subsp. equisimilis strain UT_4242_AB | SP_0121 | Ribonuclease J1 (endonuclease and 5' exonuclease) | 12207705 | 98 | 98 | 90 | 1e-297 |
| Streptococcus dysgalactiae subsp. equisimilis strain UT_4255RC | SP_0121 | Ribonuclease J1 (endonuclease and 5' exonuclease) | 12207705 | 98 | 98 | 90 | 1e-297 |
| Streptococcus dysgalactiae subsp. equisimilis strain UT_4277_BB | SP_0121 | Ribonuclease J1 (endonuclease and 5' exonuclease) | 12207705 | 98 | 98 | 90 | 1e-297 |
| Streptococcus dysgalactiae subsp. equisimilis strain UT_4966_RC | SP_0121 | Ribonuclease J1 (endonuclease and 5' exonuclease) | 12207705 | 98 | 98 | 90 | 1e-297 |
| Streptococcus dysgalactiae subsp. equisimilis strain UT-5345 | SP_0121 | Ribonuclease J1 (endonuclease and 5' exonuclease) | 12207705 | 98 | 98 | 90 | 1e-297 |
| Streptococcus dysgalactiae subsp. equisimilis strain UT-5354 | SP_0121 | Ribonuclease J1 (endonuclease and 5' exonuclease) | 12207705 | 98 | 98 | 90 | 1e-297 |
| Streptococcus dysgalactiae subsp. equisimilis strain UT-SS1069 | SP_0121 | Ribonuclease J1 (endonuclease and 5' exonuclease) | 12207705 | 98 | 98 | 90 | 1e-297 |
| Streptococcus dysgalactiae subsp. equisimilis strain UT-SS957 | SP_0121 | Ribonuclease J1 (endonuclease and 5' exonuclease) | 12207705 | 98 | 98 | 90 | 1e-297 |
| Streptococcus dysgalactiae subsp. equisimilis strain WCHSDSE-1 | SP_0121 | Ribonuclease J1 (endonuclease and 5' exonuclease) | 12207705 | 98 | 98 | 90 | 1e-296 |
| Streptococcus dysgalactiae subsp. equisimilis 167 | SPy_1633 | Ribonuclease Y | 20385762 | 92 | 99 | 99 | 1e-273 |
| Streptococcus dysgalactiae subsp. equisimilis AC-2713 | SPy_1633 | Ribonuclease Y | 20385762 | 100 | 100 | 99 | 1e-297 |
| Streptococcus dysgalactiae subsp. equisimilis AKSDE4288 | SPy_1633 | Ribonuclease Y | 20385762 | 100 | 100 | 99 | 1e-297 |
| Streptococcus dysgalactiae subsp. equisimilis ATCC 12394 | SPy_1633 | Ribonuclease Y | 20385762 | 100 | 100 | 98 | 1e-296 |
| Streptococcus dysgalactiae subsp. equisimilis GGS_124 | SPy_1633 | Ribonuclease Y | 20385762 | 100 | 100 | 99 | 1e-297 |
| Streptococcus dysgalactiae subsp. equisimilis RE378 | SPy_1633 | Ribonuclease Y | 20385762 | 100 | 100 | 99 | 1e-297 |
| Streptococcus dysgalactiae subsp. equisimilis SD SCDR1 | SPy_1633 | Ribonuclease Y | 20385762 | 100 | 100 | 99 | 1e-297 |
| Streptococcus dysgalactiae subsp. equisimilis SK1249 | SPy_1633 | Ribonuclease Y | 20385762 | 100 | 100 | 99 | 1e-297 |
| Streptococcus dysgalactiae subsp. equisimilis SK1250 | SPy_1633 | Ribonuclease Y | 20385762 | 100 | 100 | 99 | 1e-297 |
| Streptococcus dysgalactiae subsp. equisimilis strain ASDSE_96 | SPy_1633 | Ribonuclease Y | 20385762 | 100 | 100 | 99 | 1e-297 |
| Streptococcus dysgalactiae subsp. equisimilis strain ASDSE_99 | SPy_1633 | Ribonuclease Y | 20385762 | 100 | 100 | 99 | 1e-297 |
| Streptococcus dysgalactiae subsp. equisimilis strain C161L1 | SPy_1633 | Ribonuclease Y | 20385762 | 100 | 100 | 99 | 0 |
| Streptococcus dysgalactiae subsp. equisimilis strain KNZ01 | SPy_1633 | Ribonuclease Y | 20385762 | 100 | 100 | 98 | 1e-296 |
| Streptococcus dysgalactiae subsp. equisimilis strain KNZ03 | SPy_1633 | Ribonuclease Y | 20385762 | 100 | 100 | 98 | 1e-296 |
| Streptococcus dysgalactiae subsp. equisimilis strain KNZ04 | SPy_1633 | Ribonuclease Y | 20385762 | 100 | 100 | 98 | 1e-296 |
| Streptococcus dysgalactiae subsp. equisimilis strain KNZ06 | SPy_1633 | Ribonuclease Y | 20385762 | 100 | 100 | 99 | 1e-297 |
| Streptococcus dysgalactiae subsp. equisimilis strain KNZ07 | SPy_1633 | Ribonuclease Y | 20385762 | 100 | 100 | 99 | 1e-297 |
| Streptococcus dysgalactiae subsp. equisimilis strain KNZ10 | SPy_1633 | Ribonuclease Y | 20385762 | 100 | 100 | 99 | 1e-297 |
| Streptococcus dysgalactiae subsp. equisimilis strain KNZ12 | SPy_1633 | Ribonuclease Y | 20385762 | 100 | 100 | 99 | 1e-297 |
| Streptococcus dysgalactiae subsp. equisimilis strain KNZ15 | SPy_1633 | Ribonuclease Y | 20385762 | 100 | 100 | 99 | 1e-297 |
| Streptococcus dysgalactiae subsp. equisimilis strain KNZ16 | SPy_1633 | Ribonuclease Y | 20385762 | 100 | 100 | 98 | 1e-296 |
| Streptococcus dysgalactiae subsp. equisimilis strain NCTC10321 | SPy_1633 | Ribonuclease Y | 20385762 | 100 | 100 | 98 | 1e-295 |
| Streptococcus dysgalactiae subsp. equisimilis strain NCTC11554 | SPy_1633 | Ribonuclease Y | 20385762 | 100 | 100 | 98 | 1e-296 |
| Streptococcus dysgalactiae subsp. equisimilis strain NCTC11555 | SPy_1633 | Ribonuclease Y | 20385762 | 100 | 100 | 99 | 1e-297 |
| Streptococcus dysgalactiae subsp. equisimilis strain NCTC11556 | SPy_1633 | Ribonuclease Y | 20385762 | 100 | 100 | 98 | 1e-296 |
| Streptococcus dysgalactiae subsp. equisimilis strain NCTC11557 | SPy_1633 | Ribonuclease Y | 20385762 | 100 | 100 | 99 | 1e-297 |
| Streptococcus dysgalactiae subsp. equisimilis strain NCTC11564 | SPy_1633 | Ribonuclease Y | 20385762 | 100 | 100 | 99 | 1e-297 |
| Streptococcus dysgalactiae subsp. equisimilis strain NCTC5370 | SPy_1633 | Ribonuclease Y | 20385762 | 100 | 100 | 99 | 1e-297 |
| Streptococcus dysgalactiae subsp. equisimilis strain NCTC5371 | SPy_1633 | Ribonuclease Y | 20385762 | 100 | 100 | 99 | 1e-297 |
| Streptococcus dysgalactiae subsp. equisimilis strain NCTC6179 | SPy_1633 | Ribonuclease Y | 20385762 | 100 | 100 | 98 | 1e-295 |
| Streptococcus dysgalactiae subsp. equisimilis strain NCTC6181 | SPy_1633 | Ribonuclease Y | 20385762 | 100 | 100 | 98 | 1e-295 |
| Streptococcus dysgalactiae subsp. equisimilis strain NCTC6407 | SPy_1633 | Ribonuclease Y | 20385762 | 100 | 100 | 98 | 1e-295 |
| Streptococcus dysgalactiae subsp. equisimilis strain NCTC7136 | SPy_1633 | Ribonuclease Y | 20385762 | 100 | 100 | 99 | 1e-297 |
| Streptococcus dysgalactiae subsp. equisimilis strain NCTC8543 | SPy_1633 | Ribonuclease Y | 20385762 | 100 | 100 | 99 | 1e-297 |
| Streptococcus dysgalactiae subsp. equisimilis strain NCTC8546 | SPy_1633 | Ribonuclease Y | 20385762 | 100 | 100 | 99 | 1e-297 |
| Streptococcus dysgalactiae subsp. equisimilis strain NCTC9413 | SPy_1633 | Ribonuclease Y | 20385762 | 100 | 100 | 99 | 1e-297 |
| Streptococcus dysgalactiae subsp. equisimilis strain NCTC9414 | SPy_1633 | Ribonuclease Y | 20385762 | 100 | 100 | 99 | 1e-297 |
| Streptococcus dysgalactiae subsp. equisimilis strain NCTC9603 | SPy_1633 | Ribonuclease Y | 20385762 | 100 | 100 | 99 | 1e-297 |
| Streptococcus dysgalactiae subsp. equisimilis strain SS1575 | SPy_1633 | Ribonuclease Y | 20385762 | 100 | 100 | 99 | 1e-297 |
| Streptococcus dysgalactiae subsp. equisimilis strain T642 | SPy_1633 | Ribonuclease Y | 20385762 | 100 | 100 | 99 | 0 |
| Streptococcus dysgalactiae subsp. equisimilis strain UT_4031CC | SPy_1633 | Ribonuclease Y | 20385762 | 100 | 100 | 99 | 1e-297 |
| Streptococcus dysgalactiae subsp. equisimilis strain UT_4231_KK | SPy_1633 | Ribonuclease Y | 20385762 | 100 | 100 | 99 | 1e-297 |
| Streptococcus dysgalactiae subsp. equisimilis strain UT_4234_DH | SPy_1633 | Ribonuclease Y | 20385762 | 100 | 100 | 99 | 1e-297 |
| Streptococcus dysgalactiae subsp. equisimilis strain UT_4241_XS | SPy_1633 | Ribonuclease Y | 20385762 | 100 | 100 | 99 | 1e-297 |
| Streptococcus dysgalactiae subsp. equisimilis strain UT_4242_AB | SPy_1633 | Ribonuclease Y | 20385762 | 100 | 100 | 98 | 1e-296 |
| Streptococcus dysgalactiae subsp. equisimilis strain UT_4255RC | SPy_1633 | Ribonuclease Y | 20385762 | 100 | 100 | 99 | 1e-297 |
| Streptococcus dysgalactiae subsp. equisimilis strain UT_4277_BB | SPy_1633 | Ribonuclease Y | 20385762 | 100 | 100 | 99 | 1e-297 |
| Streptococcus dysgalactiae subsp. equisimilis strain UT_4966_RC | SPy_1633 | Ribonuclease Y | 20385762 | 100 | 100 | 99 | 1e-297 |
| Streptococcus dysgalactiae subsp. equisimilis strain UT-5345 | SPy_1633 | Ribonuclease Y | 20385762 | 100 | 100 | 99 | 1e-297 |
| Streptococcus dysgalactiae subsp. equisimilis strain UT-5354 | SPy_1633 | Ribonuclease Y | 20385762 | 89 | 98 | 98 | 1e-265 |
| Streptococcus dysgalactiae subsp. equisimilis strain UT-SS1069 | SPy_1633 | Ribonuclease Y | 20385762 | 100 | 100 | 99 | 1e-297 |
| Streptococcus dysgalactiae subsp. equisimilis strain UT-SS957 | SPy_1633 | Ribonuclease Y | 20385762 | 100 | 100 | 99 | 1e-297 |
| Streptococcus dysgalactiae subsp. equisimilis strain WCHSDSE-1 | SPy_1633 | Ribonuclease Y | 20385762 | 100 | 100 | 99 | 1e-297 |
| Streptococcus dysgalactiae subsp. equisimilis AC-2713 | ccrB | Site-specific recombinase | 12207705 | 99 | 99 | 90 | 1e-299 |
| Streptococcus dysgalactiae subsp. equisimilis AKSDE4288 | SP_1040 | Site-specific recombinase | 12207705 | 93 | 99 | 90 | 1e-281 |
| Streptococcus dysgalactiae subsp. equisimilis ATCC 12394 | SP_1040 | Site-specific recombinase | 12207705 | 99 | 99 | 90 | 1e-299 |
| Streptococcus dysgalactiae subsp. equisimilis RE378 | SP_1040 | Site-specific recombinase | 12207705 | 99 | 99 | 90 | 1e-299 |
| Streptococcus dysgalactiae subsp. equisimilis SK1250 | SP_1040 | Site-specific recombinase | 12207705 | 93 | 98 | 90 | 1e-281 |
| Streptococcus dysgalactiae subsp. equisimilis strain KNZ01 | SP_1040 | Site-specific recombinase | 12207705 | 99 | 99 | 90 | 1e-299 |
| Streptococcus dysgalactiae subsp. equisimilis strain KNZ03 | SP_1040 | Site-specific recombinase | 12207705 | 98 | 100 | 90 | 1e-295 |
| Streptococcus dysgalactiae subsp. equisimilis strain KNZ04 | SP_1040 | Site-specific recombinase | 12207705 | 99 | 99 | 90 | 1e-299 |
| Streptococcus dysgalactiae subsp. equisimilis strain KNZ06 | SP_1040 | Site-specific recombinase | 12207705 | 49 | 97 | 90 | 1e-143 |
| Streptococcus dysgalactiae subsp. equisimilis strain KNZ07 | SP_1040 | Site-specific recombinase | 12207705 | 85 | 100 | 90 | 1e-257 |
| Streptococcus dysgalactiae subsp. equisimilis strain KNZ10 | SP_1040 | Site-specific recombinase | 12207705 | 99 | 99 | 90 | 1e-299 |
| Streptococcus dysgalactiae subsp. equisimilis strain KNZ12 | SP_1040 | Site-specific recombinase | 12207705 | 99 | 99 | 90 | 1e-299 |
| Streptococcus dysgalactiae subsp. equisimilis strain KNZ15 | SP_1040 | Site-specific recombinase | 12207705 | 99 | 99 | 90 | 1e-299 |
| Streptococcus dysgalactiae subsp. equisimilis strain KNZ16 | SP_1040 | Site-specific recombinase | 12207705 | 99 | 99 | 90 | 1e-299 |
| Streptococcus dysgalactiae subsp. equisimilis strain NCTC11554 | SP_1040 | Site-specific recombinase | 12207705 | 99 | 99 | 90 | 1e-299 |
| Streptococcus dysgalactiae subsp. equisimilis strain NCTC11555 | SP_1040 | Site-specific recombinase | 12207705 | 99 | 99 | 94 | 1e-309 |
| Streptococcus dysgalactiae subsp. equisimilis strain NCTC11556 | SP_1040 | Site-specific recombinase | 12207705 | 99 | 99 | 90 | 1e-299 |
| Streptococcus dysgalactiae subsp. equisimilis strain NCTC11557 | SP_1040 | Site-specific recombinase | 12207705 | 99 | 99 | 96 | 1e-314 |
| Streptococcus dysgalactiae subsp. equisimilis strain NCTC11564 | SP_1040 | Site-specific recombinase | 12207705 | 99 | 98 | 96 | 1e-313 |
| Streptococcus dysgalactiae subsp. equisimilis strain T642 | SP_1040 | Site-specific recombinase | 12207705 | 99 | 99 | 90 | 0.0 |
| Streptococcus dysgalactiae subsp. equisimilis strain UT_4242_AB | SP_1040 | Site-specific recombinase | 12207705 | 99 | 99 | 90 | 1e-299 |
| Streptococcus dysgalactiae subsp. equisimilis strain UT_4255RC | SP_1040 | Site-specific recombinase | 12207705 | 99 | 99 | 96 | 1e-314 |
| Streptococcus dysgalactiae subsp. equisimilis strain UT-5345 | SP_1040 | Site-specific recombinase | 12207705 | 98 | 98 | 91 | 1e-298 |
| Streptococcus dysgalactiae subsp. equisimilis 167 | gbs0630 | Sortase A, LPXTG specific | 16824100; 25576026 | 98 | 98 | 89 | 1e-154 |
| Streptococcus dysgalactiae subsp. equisimilis 167 | gbs0631 | Sortase A, LPXTG specific | 16824100; 25576026 | 100 | 100 | 97 | 1e-154 |
| Streptococcus dysgalactiae subsp. equisimilis AC-2713 | gbs0630 | Sortase A, LPXTG specific | 16824100; 25576026 | 100 | 100 | 97 | 1e-154 |
| Streptococcus dysgalactiae subsp. equisimilis AC-2713 | gbs0631 | Sortase A, LPXTG specific | 16824100; 25576026 | 98 | 98 | 90 | 1e-155 |
| Streptococcus dysgalactiae subsp. equisimilis RE378 | srtC.4 | Sortase A, LPXTG specific | 16824100; 25576026 | 100 | 100 | 90 | 1e-158 |
| Streptococcus dysgalactiae subsp. equisimilis RE378 | srtC.3 | Sortase A, LPXTG specific | 16824100; 25576026 | 100 | 100 | 96 | 1e-153 |
| Streptococcus dysgalactiae subsp. equisimilis SD SCDR1 | gbs0630 | Sortase A, LPXTG specific | 16824100; 25576026 | 98 | 98 | 90 | 1e-155 |
| Streptococcus dysgalactiae subsp. equisimilis SD SCDR1 | gbs0631 | Sortase A, LPXTG specific | 16824100; 25576026 | 100 | 100 | 98 | 1e-155 |
| Streptococcus dysgalactiae subsp. equisimilis SK1250 | gbs0630 | Sortase A, LPXTG specific | 16824100; 25576026 | 100 | 100 | 98 | 1e-155 |
| Streptococcus dysgalactiae subsp. equisimilis SK1250 | gbs0631 | Sortase A, LPXTG specific | 16824100; 25576026 | 100 | 100 | 90 | 1e-158 |
| Streptococcus dysgalactiae subsp. equisimilis strain ASDSE_96 | gbs0630 | Sortase A, LPXTG specific | 16824100; 25576026 | 98 | 98 | 90 | 1e-157 |
| Streptococcus dysgalactiae subsp. equisimilis strain ASDSE_96 | gbs0631 | Sortase A, LPXTG specific | 16824100; 25576026 | 100 | 100 | 86 | 1e-140 |
| Streptococcus dysgalactiae subsp. equisimilis strain ASDSE_99 | gbs0630 | Sortase A, LPXTG specific | 16824100; 25576026 | 100 | 100 | 97 | 1e-155 |
| Streptococcus dysgalactiae subsp. equisimilis strain ASDSE_99 | gbs0631 | Sortase A, LPXTG specific | 16824100; 25576026 | 98 | 98 | 91 | 1e-156 |
| Streptococcus dysgalactiae subsp. equisimilis strain KNZ01 | gbs0631 | Sortase A, LPXTG specific | 16824100; 25576026 | 100 | 100 | 96 | 1e-153 |
| Streptococcus dysgalactiae subsp. equisimilis strain KNZ01 | gbs0630 | Sortase A, LPXTG specific | 16824100; 25576026 | 100 | 100 | 90 | 1e-158 |
| Streptococcus dysgalactiae subsp. equisimilis strain KNZ01 | gbs0630 | Sortase A, LPXTG specific | 16824100; 25576026 | 100 | 100 | 90 | 1e-158 |
| Streptococcus dysgalactiae subsp. equisimilis strain KNZ01 | gbs0631 | Sortase A, LPXTG specific | 16824100; 25576026 | 100 | 100 | 96 | 1e-153 |
| Streptococcus dysgalactiae subsp. equisimilis strain KNZ03 | gbs0631 | Sortase A, LPXTG specific | 16824100; 25576026 | 100 | 100 | 86 | 1e-139 |
| Streptococcus dysgalactiae subsp. equisimilis strain KNZ03 | gbs0631 | Sortase A, LPXTG specific | 16824100; 25576026 | 100 | 100 | 86 | 1e-139 |
| Streptococcus dysgalactiae subsp. equisimilis strain KNZ03 | gbs0630 | Sortase A, LPXTG specific | 16824100; 25576026 | 98 | 98 | 91 | 1e-157 |
| Streptococcus dysgalactiae subsp. equisimilis strain KNZ03 | gbs0630 | Sortase A, LPXTG specific | 16824100; 25576026 | 98 | 98 | 91 | 1e-157 |
| Streptococcus dysgalactiae subsp. equisimilis strain KNZ04 | gbs0631 | Sortase A, LPXTG specific | 16824100; 25576026 | 100 | 100 | 97 | 1e-154 |
| Streptococcus dysgalactiae subsp. equisimilis strain KNZ04 | gbs0630 | Sortase A, LPXTG specific | 16824100; 25576026 | 98 | 98 | 90 | 1e-155 |
| Streptococcus dysgalactiae subsp. equisimilis strain KNZ06 | gbs0631 | Sortase A, LPXTG specific | 16824100; 25576026 | 100 | 100 | 86 | 1e-140 |
| Streptococcus dysgalactiae subsp. equisimilis strain KNZ06 | gbs0630 | Sortase A, LPXTG specific | 16824100; 25576026 | 98 | 98 | 90 | 1e-156 |
| Streptococcus dysgalactiae subsp. equisimilis strain KNZ07 | gbs0631 | Sortase A, LPXTG specific | 16824100; 25576026 | 100 | 100 | 96 | 1e-153 |
| Streptococcus dysgalactiae subsp. equisimilis strain KNZ07 | gbs0630 | Sortase A, LPXTG specific | 16824100; 25576026 | 100 | 100 | 90 | 1e-158 |
| Streptococcus dysgalactiae subsp. equisimilis strain KNZ10 | gbs0630 | Sortase A, LPXTG specific | 16824100; 25576026 | 100 | 100 | 90 | 1e-158 |
| Streptococcus dysgalactiae subsp. equisimilis strain KNZ10 | gbs0631 | Sortase A, LPXTG specific | 16824100; 25576026 | 100 | 100 | 96 | 1e-153 |
| Streptococcus dysgalactiae subsp. equisimilis strain KNZ12 | gbs0631 | Sortase A, LPXTG specific | 16824100; 25576026 | 100 | 100 | 96 | 1e-153 |
| Streptococcus dysgalactiae subsp. equisimilis strain KNZ12 | gbs0630 | Sortase A, LPXTG specific | 16824100; 25576026 | 100 | 100 | 90 | 1e-158 |
| Streptococcus dysgalactiae subsp. equisimilis strain KNZ15 | gbs0630 | Sortase A, LPXTG specific | 16824100; 25576026 | 100 | 100 | 90 | 1e-158 |
| Streptococcus dysgalactiae subsp. equisimilis strain KNZ15 | gbs0631 | Sortase A, LPXTG specific | 16824100; 25576026 | 100 | 100 | 96 | 1e-153 |
| Streptococcus dysgalactiae subsp. equisimilis strain NCTC10321 | gbs0631 | Sortase A, LPXTG specific | 16824100; 25576026 | 100 | 100 | 97 | 1e-154 |
| Streptococcus dysgalactiae subsp. equisimilis strain NCTC10321 | gbs0630 | Sortase A, LPXTG specific | 16824100; 25576026 | 98 | 98 | 90 | 1e-156 |
| Streptococcus dysgalactiae subsp. equisimilis strain NCTC11555 | gbs0630 | Sortase A, LPXTG specific | 16824100; 25576026 | 98 | 98 | 91 | 1e-157 |
| Streptococcus dysgalactiae subsp. equisimilis strain NCTC11555 | gbs0631 | Sortase A, LPXTG specific | 16824100; 25576026 | 100 | 100 | 86 | 1e-139 |
| Streptococcus dysgalactiae subsp. equisimilis strain NCTC11556 | gbs0630 | Sortase A, LPXTG specific | 16824100; 25576026 | 98 | 98 | 90 | 1e-157 |
| Streptococcus dysgalactiae subsp. equisimilis strain NCTC11556 | gbs0631 | Sortase A, LPXTG specific | 16824100; 25576026 | 100 | 100 | 86 | 1e-140 |
| Streptococcus dysgalactiae subsp. equisimilis strain NCTC11557 | gbs0631 | Sortase A, LPXTG specific | 16824100; 25576026 | 100 | 100 | 97 | 1e-154 |
| Streptococcus dysgalactiae subsp. equisimilis strain NCTC11557 | gbs0630 | Sortase A, LPXTG specific | 16824100; 25576026 | 98 | 98 | 90 | 1e-155 |
| Streptococcus dysgalactiae subsp. equisimilis strain NCTC11564 | gbs0630 | Sortase A, LPXTG specific | 16824100; 25576026 | 98 | 98 | 90 | 1e-156 |
| Streptococcus dysgalactiae subsp. equisimilis strain NCTC11564 | gbs0631 | Sortase A, LPXTG specific | 16824100; 25576026 | 100 | 100 | 97 | 1e-155 |
| Streptococcus dysgalactiae subsp. equisimilis strain NCTC5371 | gbs0631 | Sortase A, LPXTG specific | 16824100; 25576026 | 100 | 100 | 97 | 1e-155 |
| Streptococcus dysgalactiae subsp. equisimilis strain NCTC5371 | gbs0630 | Sortase A, LPXTG specific | 16824100; 25576026 | 98 | 98 | 90 | 1e-156 |
| Streptococcus dysgalactiae subsp. equisimilis strain NCTC6407 | gbs0630 | Sortase A, LPXTG specific | 16824100; 25576026 | 98 | 98 | 90 | 1e-156 |
| Streptococcus dysgalactiae subsp. equisimilis strain NCTC6407 | gbs0631 | Sortase A, LPXTG specific | 16824100; 25576026 | 100 | 100 | 97 | 1e-154 |
| Streptococcus dysgalactiae subsp. equisimilis strain NCTC7136 | gbs0630 | Sortase A, LPXTG specific | 16824100; 25576026 | 98 | 98 | 90 | 1e-156 |
| Streptococcus dysgalactiae subsp. equisimilis strain NCTC7136 | gbs0631 | Sortase A, LPXTG specific | 16824100; 25576026 | 100 | 100 | 96 | 1e-154 |
| Streptococcus dysgalactiae subsp. equisimilis strain NCTC8546 | gbs0631 | Sortase A, LPXTG specific | 16824100; 25576026 | 100 | 100 | 91 | 1e-145 |
| Streptococcus dysgalactiae subsp. equisimilis strain NCTC8546 | gbs0630 | Sortase A, LPXTG specific | 16824100; 25576026 | 98 | 98 | 90 | 1e-155 |
| Streptococcus dysgalactiae subsp. equisimilis strain NCTC9414 | gbs0631 | Sortase A, LPXTG specific | 16824100; 25576026 | 100 | 100 | 97 | 1e-155 |
| Streptococcus dysgalactiae subsp. equisimilis strain NCTC9414 | gbs0630 | Sortase A, LPXTG specific | 16824100; 25576026 | 98 | 98 | 90 | 1e-156 |
| Streptococcus dysgalactiae subsp. equisimilis strain UT_4031CC | gbs0630 | Sortase A, LPXTG specific | 16824100; 25576026 | 98 | 98 | 90 | 1e-156 |
| Streptococcus dysgalactiae subsp. equisimilis strain UT_4031CC | gbs0631 | Sortase A, LPXTG specific | 16824100; 25576026 | 100 | 100 | 97 | 1e-154 |
| Streptococcus dysgalactiae subsp. equisimilis strain UT_4231_KK | gbs0630 | Sortase A, LPXTG specific | 16824100; 25576026 | 100 | 100 | 91 | 1e-145 |
| Streptococcus dysgalactiae subsp. equisimilis strain UT_4231_KK | gbs0631 | Sortase A, LPXTG specific | 16824100; 25576026 | 98 | 98 | 90 | 1e-155 |
| Streptococcus dysgalactiae subsp. equisimilis strain UT_4234_DH | gbs0630 | Sortase A, LPXTG specific | 16824100; 25576026 | 98 | 98 | 91 | 1e-156 |
| Streptococcus dysgalactiae subsp. equisimilis strain UT_4234_DH | gbs0631 | Sortase A, LPXTG specific | 16824100; 25576026 | 100 | 100 | 97 | 1e-155 |
| Streptococcus dysgalactiae subsp. equisimilis strain UT_4241_XS | gbs0630 | Sortase A, LPXTG specific | 16824100; 25576026 | 100 | 100 | 97 | 1e-154 |
| Streptococcus dysgalactiae subsp. equisimilis strain UT_4241_XS | gbs0631 | Sortase A, LPXTG specific | 16824100; 25576026 | 98 | 98 | 89 | 1e-153 |
| Streptococcus dysgalactiae subsp. equisimilis strain UT_4242_AB | gbs0630 | Sortase A, LPXTG specific | 16824100; 25576026 | 100 | 100 | 86 | 1e-140 |
| Streptococcus dysgalactiae subsp. equisimilis strain UT_4242_AB | gbs0631 | Sortase A, LPXTG specific | 16824100; 25576026 | 98 | 98 | 90 | 1e-157 |
| Streptococcus dysgalactiae subsp. equisimilis strain UT_4255RC | gbs0630 | Sortase A, LPXTG specific | 16824100; 25576026 | 100 | 100 | 97 | 1e-154 |
| Streptococcus dysgalactiae subsp. equisimilis strain UT_4255RC | gbs0631 | Sortase A, LPXTG specific | 16824100; 25576026 | 98 | 98 | 90 | 1e-155 |
| Streptococcus dysgalactiae subsp. equisimilis strain UT_4277_BB | gbs0630 | Sortase A, LPXTG specific | 16824100; 25576026 | 98 | 98 | 90 | 1e-157 |
| Streptococcus dysgalactiae subsp. equisimilis strain UT_4277_BB | gbs0631 | Sortase A, LPXTG specific | 16824100; 25576026 | 100 | 100 | 86 | 1e-140 |
| Streptococcus dysgalactiae subsp. equisimilis strain UT_4966_RC | gbs0630 | Sortase A, LPXTG specific | 16824100; 25576026 | 98 | 98 | 89 | 1e-154 |
| Streptococcus dysgalactiae subsp. equisimilis strain UT_4966_RC | gbs0631 | Sortase A, LPXTG specific | 16824100; 25576026 | 100 | 100 | 97 | 1e-154 |
| Streptococcus dysgalactiae subsp. equisimilis strain UT-5345 | gbs0630 | Sortase A, LPXTG specific | 16824100; 25576026 | 100 | 100 | 97 | 1e-154 |
| Streptococcus dysgalactiae subsp. equisimilis strain UT-5345 | gbs0631 | Sortase A, LPXTG specific | 16824100; 25576026 | 98 | 98 | 90 | 1e-156 |
| Streptococcus dysgalactiae subsp. equisimilis strain UT-SS1069 | gbs0630 | Sortase A, LPXTG specific | 16824100; 25576026 | 100 | 100 | 97 | 1e-154 |
| Streptococcus dysgalactiae subsp. equisimilis strain UT-SS1069 | gbs0631 | Sortase A, LPXTG specific | 16824100; 25576026 | 98 | 98 | 89 | 1e-154 |
| Streptococcus dysgalactiae subsp. equisimilis strain UT-SS957 | gbs0630 | Sortase A, LPXTG specific | 16824100; 25576026 | 98 | 98 | 90 | 1e-156 |
| Streptococcus dysgalactiae subsp. equisimilis strain UT-SS957 | gbs0631 | Sortase A, LPXTG specific | 16824100; 25576026 | 100 | 100 | 97 | 1e-155 |
| Streptococcus dysgalactiae subsp. equisimilis 167 | luxS | S-ribosylhomocysteine lyase (EC 4.4.1.21) @ Autoinducer-2 production protein LuxS | 15102809 | 100 | 100 | 83 | 1e-75 |
| Streptococcus dysgalactiae subsp. equisimilis AC-2713 | luxS | S-ribosylhomocysteine lyase (EC 4.4.1.21) @ Autoinducer-2 production protein LuxS | 15102809 | 100 | 100 | 83 | 1e-75 |
| Streptococcus dysgalactiae subsp. equisimilis AKSDE4288 | luxS | S-ribosylhomocysteine lyase (EC 4.4.1.21) @ Autoinducer-2 production protein LuxS | 15102809 | 100 | 100 | 83 | 1e-75 |
| Streptococcus dysgalactiae subsp. equisimilis ATCC 12394 | luxS | S-ribosylhomocysteine lyase (EC 4.4.1.21) @ Autoinducer-2 production protein LuxS | 15102809 | 100 | 100 | 83 | 2e-75 |
| Streptococcus dysgalactiae subsp. equisimilis GGS_124 | luxS | S-ribosylhomocysteine lyase (EC 4.4.1.21) @ Autoinducer-2 production protein LuxS | 15102809 | 100 | 100 | 83 | 1e-75 |
| Streptococcus dysgalactiae subsp. equisimilis RE378 | luxS | S-ribosylhomocysteine lyase (EC 4.4.1.21) @ Autoinducer-2 production protein LuxS | 15102809 | 100 | 100 | 83 | 8e-75 |
| Streptococcus dysgalactiae subsp. equisimilis SD SCDR1 | luxS | S-ribosylhomocysteine lyase (EC 4.4.1.21) @ Autoinducer-2 production protein LuxS | 15102809 | 100 | 100 | 83 | 1e-75 |
| Streptococcus dysgalactiae subsp. equisimilis SK1249 | luxS | S-ribosylhomocysteine lyase (EC 4.4.1.21) @ Autoinducer-2 production protein LuxS | 15102809 | 100 | 100 | 83 | 2e-75 |
| Streptococcus dysgalactiae subsp. equisimilis SK1250 | luxS | S-ribosylhomocysteine lyase (EC 4.4.1.21) @ Autoinducer-2 production protein LuxS | 15102809 | 100 | 100 | 83 | 1e-75 |
| Streptococcus dysgalactiae subsp. equisimilis strain ASDSE_96 | luxS | S-ribosylhomocysteine lyase (EC 4.4.1.21) @ Autoinducer-2 production protein LuxS | 15102809 | 100 | 100 | 83 | 2e-75 |
| Streptococcus dysgalactiae subsp. equisimilis strain ASDSE_99 | luxS | S-ribosylhomocysteine lyase (EC 4.4.1.21) @ Autoinducer-2 production protein LuxS | 15102809 | 100 | 100 | 83 | 1e-75 |
| Streptococcus dysgalactiae subsp. equisimilis strain C161L1 | luxS | S-ribosylhomocysteine lyase (EC 4.4.1.21) @ Autoinducer-2 production protein LuxS | 15102809 | 100 | 100 | 83 | 0 |
| Streptococcus dysgalactiae subsp. equisimilis strain KNZ01 | luxS | S-ribosylhomocysteine lyase (EC 4.4.1.21) @ Autoinducer-2 production protein LuxS | 15102809 | 100 | 100 | 83 | 8e-75 |
| Streptococcus dysgalactiae subsp. equisimilis strain KNZ03 | luxS | S-ribosylhomocysteine lyase (EC 4.4.1.21) @ Autoinducer-2 production protein LuxS | 15102809 | 100 | 100 | 83 | 2e-75 |
| Streptococcus dysgalactiae subsp. equisimilis strain KNZ04 | luxS | S-ribosylhomocysteine lyase (EC 4.4.1.21) @ Autoinducer-2 production protein LuxS | 15102809 | 100 | 100 | 83 | 2e-75 |
| Streptococcus dysgalactiae subsp. equisimilis strain KNZ06 | luxS | S-ribosylhomocysteine lyase (EC 4.4.1.21) @ Autoinducer-2 production protein LuxS | 15102809 | 100 | 100 | 83 | 2e-75 |
| Streptococcus dysgalactiae subsp. equisimilis strain KNZ07 | luxS | S-ribosylhomocysteine lyase (EC 4.4.1.21) @ Autoinducer-2 production protein LuxS | 15102809 | 100 | 100 | 83 | 8e-75 |
| Streptococcus dysgalactiae subsp. equisimilis strain KNZ10 | luxS | S-ribosylhomocysteine lyase (EC 4.4.1.21) @ Autoinducer-2 production protein LuxS | 15102809 | 100 | 100 | 83 | 8e-75 |
| Streptococcus dysgalactiae subsp. equisimilis strain KNZ12 | luxS | S-ribosylhomocysteine lyase (EC 4.4.1.21) @ Autoinducer-2 production protein LuxS | 15102809 | 100 | 100 | 83 | 8e-75 |
| Streptococcus dysgalactiae subsp. equisimilis strain KNZ15 | luxS | S-ribosylhomocysteine lyase (EC 4.4.1.21) @ Autoinducer-2 production protein LuxS | 15102809 | 100 | 100 | 83 | 8e-75 |
| Streptococcus dysgalactiae subsp. equisimilis strain KNZ16 | luxS | S-ribosylhomocysteine lyase (EC 4.4.1.21) @ Autoinducer-2 production protein LuxS | 15102809 | 100 | 100 | 83 | 2e-75 |
| Streptococcus dysgalactiae subsp. equisimilis strain NCTC10321 | luxS | S-ribosylhomocysteine lyase (EC 4.4.1.21) @ Autoinducer-2 production protein LuxS | 15102809 | 100 | 100 | 83 | 2e-75 |
| Streptococcus dysgalactiae subsp. equisimilis strain NCTC11554 | luxS | S-ribosylhomocysteine lyase (EC 4.4.1.21) @ Autoinducer-2 production protein LuxS | 15102809 | 100 | 100 | 83 | 2e-75 |
| Streptococcus dysgalactiae subsp. equisimilis strain NCTC11555 | luxS | S-ribosylhomocysteine lyase (EC 4.4.1.21) @ Autoinducer-2 production protein LuxS | 15102809 | 100 | 100 | 83 | 1e-75 |
| Streptococcus dysgalactiae subsp. equisimilis strain NCTC11556 | luxS | S-ribosylhomocysteine lyase (EC 4.4.1.21) @ Autoinducer-2 production protein LuxS | 15102809 | 100 | 100 | 83 | 2e-75 |
| Streptococcus dysgalactiae subsp. equisimilis strain NCTC11557 | luxS | S-ribosylhomocysteine lyase (EC 4.4.1.21) @ Autoinducer-2 production protein LuxS | 15102809 | 100 | 100 | 83 | 1e-75 |
| Streptococcus dysgalactiae subsp. equisimilis strain NCTC11564 | luxS | S-ribosylhomocysteine lyase (EC 4.4.1.21) @ Autoinducer-2 production protein LuxS | 15102809 | 100 | 100 | 83 | 2e-75 |
| Streptococcus dysgalactiae subsp. equisimilis strain NCTC5370 | luxS | S-ribosylhomocysteine lyase (EC 4.4.1.21) @ Autoinducer-2 production protein LuxS | 15102809 | 100 | 100 | 83 | 1e-75 |
| Streptococcus dysgalactiae subsp. equisimilis strain NCTC5371 | luxS | S-ribosylhomocysteine lyase (EC 4.4.1.21) @ Autoinducer-2 production protein LuxS | 15102809 | 100 | 100 | 83 | 1e-75 |
| Streptococcus dysgalactiae subsp. equisimilis strain NCTC6179 | luxS | S-ribosylhomocysteine lyase (EC 4.4.1.21) @ Autoinducer-2 production protein LuxS | 15102809 | 100 | 100 | 83 | 3e-75 |
| Streptococcus dysgalactiae subsp. equisimilis strain NCTC6181 | luxS | S-ribosylhomocysteine lyase (EC 4.4.1.21) @ Autoinducer-2 production protein LuxS | 15102809 | 100 | 100 | 83 | 3e-75 |
| Streptococcus dysgalactiae subsp. equisimilis strain NCTC6407 | luxS | S-ribosylhomocysteine lyase (EC 4.4.1.21) @ Autoinducer-2 production protein LuxS | 15102809 | 100 | 100 | 83 | 2e-75 |
| Streptococcus dysgalactiae subsp. equisimilis strain NCTC7136 | luxS | S-ribosylhomocysteine lyase (EC 4.4.1.21) @ Autoinducer-2 production protein LuxS | 15102809 | 100 | 100 | 83 | 2e-75 |
| Streptococcus dysgalactiae subsp. equisimilis strain NCTC8543 | luxS | S-ribosylhomocysteine lyase (EC 4.4.1.21) @ Autoinducer-2 production protein LuxS | 15102809 | 100 | 100 | 83 | 1e-75 |
| Streptococcus dysgalactiae subsp. equisimilis strain NCTC8546 | luxS | S-ribosylhomocysteine lyase (EC 4.4.1.21) @ Autoinducer-2 production protein LuxS | 15102809 | 100 | 100 | 83 | 1e-75 |
| Streptococcus dysgalactiae subsp. equisimilis strain NCTC9413 | luxS | S-ribosylhomocysteine lyase (EC 4.4.1.21) @ Autoinducer-2 production protein LuxS | 15102809 | 100 | 100 | 83 | 1e-75 |
| Streptococcus dysgalactiae subsp. equisimilis strain NCTC9414 | luxS | S-ribosylhomocysteine lyase (EC 4.4.1.21) @ Autoinducer-2 production protein LuxS | 15102809 | 100 | 100 | 83 | 1e-75 |
| Streptococcus dysgalactiae subsp. equisimilis strain NCTC9603 | luxS | S-ribosylhomocysteine lyase (EC 4.4.1.21) @ Autoinducer-2 production protein LuxS | 15102809 | 100 | 100 | 83 | 8e-75 |
| Streptococcus dysgalactiae subsp. equisimilis strain SS1575 | luxS | S-ribosylhomocysteine lyase (EC 4.4.1.21) @ Autoinducer-2 production protein LuxS | 15102809 | 100 | 100 | 83 | 1e-75 |
| Streptococcus dysgalactiae subsp. equisimilis strain T642 | luxS | S-ribosylhomocysteine lyase (EC 4.4.1.21) @ Autoinducer-2 production protein LuxS | 15102809 | 100 | 100 | 83 | 0 |
| Streptococcus dysgalactiae subsp. equisimilis strain UT_4031CC | luxS | S-ribosylhomocysteine lyase (EC 4.4.1.21) @ Autoinducer-2 production protein LuxS | 15102809 | 100 | 100 | 83 | 1e-75 |
| Streptococcus dysgalactiae subsp. equisimilis strain UT_4231_KK | luxS | S-ribosylhomocysteine lyase (EC 4.4.1.21) @ Autoinducer-2 production protein LuxS | 15102809 | 100 | 100 | 83 | 1e-75 |
| Streptococcus dysgalactiae subsp. equisimilis strain UT_4234_DH | luxS | S-ribosylhomocysteine lyase (EC 4.4.1.21) @ Autoinducer-2 production protein LuxS | 15102809 | 100 | 100 | 83 | 1e-75 |
| Streptococcus dysgalactiae subsp. equisimilis strain UT_4241_XS | luxS | S-ribosylhomocysteine lyase (EC 4.4.1.21) @ Autoinducer-2 production protein LuxS | 15102809 | 100 | 100 | 83 | 1e-75 |
| Streptococcus dysgalactiae subsp. equisimilis strain UT_4242_AB | luxS | S-ribosylhomocysteine lyase (EC 4.4.1.21) @ Autoinducer-2 production protein LuxS | 15102809 | 100 | 100 | 83 | 2e-75 |
| Streptococcus dysgalactiae subsp. equisimilis strain UT_4255RC | luxS | S-ribosylhomocysteine lyase (EC 4.4.1.21) @ Autoinducer-2 production protein LuxS | 15102809 | 100 | 100 | 83 | 1e-75 |
| Streptococcus dysgalactiae subsp. equisimilis strain UT_4277_BB | luxS | S-ribosylhomocysteine lyase (EC 4.4.1.21) @ Autoinducer-2 production protein LuxS | 15102809 | 100 | 100 | 83 | 1e-75 |
| Streptococcus dysgalactiae subsp. equisimilis strain UT_4966_RC | luxS | S-ribosylhomocysteine lyase (EC 4.4.1.21) @ Autoinducer-2 production protein LuxS | 15102809 | 100 | 100 | 83 | 1e-75 |
| Streptococcus dysgalactiae subsp. equisimilis strain UT-5345 | luxS | S-ribosylhomocysteine lyase (EC 4.4.1.21) @ Autoinducer-2 production protein LuxS | 15102809 | 100 | 100 | 83 | 1e-75 |
| Streptococcus dysgalactiae subsp. equisimilis strain UT-5354 | luxS | S-ribosylhomocysteine lyase (EC 4.4.1.21) @ Autoinducer-2 production protein LuxS | 15102809 | 100 | 100 | 83 | 1e-75 |
| Streptococcus dysgalactiae subsp. equisimilis strain UT-SS1069 | luxS | S-ribosylhomocysteine lyase (EC 4.4.1.21) @ Autoinducer-2 production protein LuxS | 15102809 | 100 | 100 | 83 | 1e-75 |
| Streptococcus dysgalactiae subsp. equisimilis strain UT-SS957 | luxS | S-ribosylhomocysteine lyase (EC 4.4.1.21) @ Autoinducer-2 production protein LuxS | 15102809 | 100 | 100 | 83 | 1e-75 |
| Streptococcus dysgalactiae subsp. equisimilis strain WCHSDSE-1 | luxS | S-ribosylhomocysteine lyase (EC 4.4.1.21) @ Autoinducer-2 production protein LuxS | 15102809 | 100 | 100 | 83 | 1e-75 |
| Streptococcus dysgalactiae subsp. equisimilis 167 | ska | Streptokinase | 10678902 | 52 | 99 | 89 | 1e-118 |
| Streptococcus dysgalactiae subsp. equisimilis 167 | ska | Streptokinase | 10678902 | 51 | 99 | 87 | 1e-116 |
| Streptococcus dysgalactiae subsp. equisimilis 167 | ska | Streptokinase | 10678902 | 43 | 98 | 90 | 8e-94 |
| Streptococcus dysgalactiae subsp. equisimilis 167 | ska | Streptokinase | 10678902 | 44 | 100 | 86 | 2e-92 |
| Streptococcus dysgalactiae subsp. equisimilis AC-2713 | ska | Streptokinase | 10678902 | 100 | 100 | 89 | 1e-235 |
| Streptococcus dysgalactiae subsp. equisimilis AKSDE4288 | ska | Streptokinase | 10678902 | 100 | 100 | 89 | 1e-235 |
| Streptococcus dysgalactiae subsp. equisimilis ATCC 12394 | ska | Streptokinase | 10678902 | 100 | 100 | 89 | 1e-233 |
| Streptococcus dysgalactiae subsp. equisimilis ATCC 12394 | ska | Streptokinase | 10678902 | 100 | 100 | 85 | 1e-219 |
| Streptococcus dysgalactiae subsp. equisimilis GGS_124 | ska | Streptokinase | 10678902 | 100 | 100 | 85 | 1e-220 |
| Streptococcus dysgalactiae subsp. equisimilis GGS_124 | ska | Streptokinase | 10678902 | 100 | 100 | 88 | 1e-230 |
| Streptococcus dysgalactiae subsp. equisimilis RE378 | ska | Streptokinase | 10678902 | 100 | 100 | 87 | 1e-227 |
| Streptococcus dysgalactiae subsp. equisimilis RE378 | ska | Streptokinase | 10678902 | 100 | 100 | 83 | 1e-214 |
| Streptococcus dysgalactiae subsp. equisimilis SD SCDR1 | ska | Streptokinase | 10678902 | 100 | 100 | 84 | 1e-215 |
| Streptococcus dysgalactiae subsp. equisimilis SD SCDR1 | ska | Streptokinase | 10678902 | 100 | 100 | 87 | 1e-229 |
| Streptococcus dysgalactiae subsp. equisimilis SK1249 | ska | Streptokinase | 10678902 | 27 | 99 | 88 | 9e-59 |
| Streptococcus dysgalactiae subsp. equisimilis SK1249 | ska | Streptokinase | 10678902 | 27 | 99 | 97 | 2e-65 |
| Streptococcus dysgalactiae subsp. equisimilis SK1249 | ska | Streptokinase | 10678902 | 64 | 98 | 91 | 1e-147 |
| Streptococcus dysgalactiae subsp. equisimilis SK1249 | ska | Streptokinase | 10678902 | 64 | 98 | 80 | 1e-125 |
| Streptococcus dysgalactiae subsp. equisimilis SK1250 | skc | Streptokinase | 10678902 | 100 | 100 | 84 | 1e-215 |
| Streptococcus dysgalactiae subsp. equisimilis SK1250 | skc | Streptokinase | 10678902 | 100 | 100 | 87 | 1e-229 |
| Streptococcus dysgalactiae subsp. equisimilis strain ASDSE_96 | ska | Streptokinase | 10678902 | 100 | 100 | 89 | 1e-233 |
| Streptococcus dysgalactiae subsp. equisimilis strain ASDSE_99 | ska | Streptokinase | 10678902 | 100 | 100 | 84 | 1e-216 |
| Streptococcus dysgalactiae subsp. equisimilis strain ASDSE_99 | ska | Streptokinase | 10678902 | 100 | 100 | 87 | 1e-228 |
| Streptococcus dysgalactiae subsp. equisimilis strain C161L1 | ska | Streptokinase | 10678902 | 100 | 100 | 86 | 0 |
| Streptococcus dysgalactiae subsp. equisimilis strain C161L1 | ska | Streptokinase | 10678902 | 100 | 100 | 88 | 0 |
| Streptococcus dysgalactiae subsp. equisimilis strain KNZ01 | ska | Streptokinase | 10678902 | 100 | 100 | 87 | 1e-227 |
| Streptococcus dysgalactiae subsp. equisimilis strain KNZ01 | ska | Streptokinase | 10678902 | 100 | 100 | 83 | 1e-214 |
| Streptococcus dysgalactiae subsp. equisimilis strain KNZ01 | ska | Streptokinase | 10678902 | 100 | 100 | 87 | 1e-227 |
| Streptococcus dysgalactiae subsp. equisimilis strain KNZ01 | ska | Streptokinase | 10678902 | 100 | 100 | 83 | 1e-214 |
| Streptococcus dysgalactiae subsp. equisimilis strain KNZ03 | ska | Streptokinase | 10678902 | 100 | 100 | 89 | 1e-235 |
| Streptococcus dysgalactiae subsp. equisimilis strain KNZ03 | ska | Streptokinase | 10678902 | 100 | 100 | 89 | 1e-235 |
| Streptococcus dysgalactiae subsp. equisimilis strain KNZ04 | ska | Streptokinase | 10678902 | 100 | 100 | 89 | 1e-235 |
| Streptococcus dysgalactiae subsp. equisimilis strain KNZ06 | ska | Streptokinase | 10678902 | 100 | 100 | 87 | 1e-227 |
| Streptococcus dysgalactiae subsp. equisimilis strain KNZ06 | ska | Streptokinase | 10678902 | 100 | 100 | 83 | 1e-214 |
| Streptococcus dysgalactiae subsp. equisimilis strain KNZ07 | ska | Streptokinase | 10678902 | 100 | 100 | 87 | 1e-227 |
| Streptococcus dysgalactiae subsp. equisimilis strain KNZ07 | ska | Streptokinase | 10678902 | 100 | 100 | 83 | 1e-214 |
| Streptococcus dysgalactiae subsp. equisimilis strain KNZ10 | ska | Streptokinase | 10678902 | 100 | 100 | 89 | 1e-235 |
| Streptococcus dysgalactiae subsp. equisimilis strain KNZ12 | ska | Streptokinase | 10678902 | 100 | 100 | 83 | 1e-214 |
| Streptococcus dysgalactiae subsp. equisimilis strain KNZ12 | ska | Streptokinase | 10678902 | 100 | 100 | 87 | 1e-227 |
| Streptococcus dysgalactiae subsp. equisimilis strain KNZ15 | ska | Streptokinase | 10678902 | 100 | 100 | 83 | 1e-214 |
| Streptococcus dysgalactiae subsp. equisimilis strain KNZ15 | ska | Streptokinase | 10678902 | 100 | 100 | 87 | 1e-227 |
| Streptococcus dysgalactiae subsp. equisimilis strain KNZ16 | ska | Streptokinase | 10678902 | 100 | 100 | 85 | 1e-219 |
| Streptococcus dysgalactiae subsp. equisimilis strain KNZ16 | ska | Streptokinase | 10678902 | 100 | 100 | 89 | 1e-233 |
| Streptococcus dysgalactiae subsp. equisimilis strain NCTC11554 | ska | Streptokinase | 10678902 | 100 | 100 | 85 | 1e-219 |
| Streptococcus dysgalactiae subsp. equisimilis strain NCTC11554 | ska | Streptokinase | 10678902 | 100 | 100 | 89 | 1e-233 |
| Streptococcus dysgalactiae subsp. equisimilis strain NCTC11555 | ska | Streptokinase | 10678902 | 100 | 100 | 84 | 1e-217 |
| Streptococcus dysgalactiae subsp. equisimilis strain NCTC11555 | ska | Streptokinase | 10678902 | 100 | 100 | 87 | 1e-228 |
| Streptococcus dysgalactiae subsp. equisimilis strain NCTC11556 | ska | Streptokinase | 10678902 | 100 | 100 | 85 | 1e-220 |
| Streptococcus dysgalactiae subsp. equisimilis strain NCTC11556 | ska | Streptokinase | 10678902 | 100 | 100 | 88 | 1e-230 |
| Streptococcus dysgalactiae subsp. equisimilis strain NCTC11557 | ska | Streptokinase | 10678902 | 100 | 100 | 87 | 1e-228 |
| Streptococcus dysgalactiae subsp. equisimilis strain NCTC11557 | ska | Streptokinase | 10678902 | 100 | 100 | 84 | 1e-216 |
| Streptococcus dysgalactiae subsp. equisimilis strain NCTC11564 | ska | Streptokinase | 10678902 | 100 | 100 | 83 | 1e-216 |
| Streptococcus dysgalactiae subsp. equisimilis strain NCTC11564 | ska | Streptokinase | 10678902 | 100 | 100 | 87 | 1e-229 |
| Streptococcus dysgalactiae subsp. equisimilis strain NCTC5370 | ska | Streptokinase | 10678902 | 100 | 100 | 87 | 1e-229 |
| Streptococcus dysgalactiae subsp. equisimilis strain NCTC5371 | ska | Streptokinase | 10678902 | 100 | 100 | 85 | 1e-219 |
| Streptococcus dysgalactiae subsp. equisimilis strain NCTC5371 | ska | Streptokinase | 10678902 | 100 | 100 | 89 | 1e-234 |
| Streptococcus dysgalactiae subsp. equisimilis strain NCTC7136 | ska | Streptokinase | 10678902 | 100 | 100 | 88 | 1e-232 |
| Streptococcus dysgalactiae subsp. equisimilis strain NCTC8543 | ska | Streptokinase | 10678902 | 97 | 97 | 85 | 1e-212 |
| Streptococcus dysgalactiae subsp. equisimilis strain NCTC8543 | ska | Streptokinase | 10678902 | 97 | 97 | 90 | 1e-229 |
| Streptococcus dysgalactiae subsp. equisimilis strain NCTC8546 | ska | Streptokinase | 10678902 | 100 | 100 | 88 | 1e-231 |
| Streptococcus dysgalactiae subsp. equisimilis strain NCTC9413 | ska | Streptokinase | 10678902 | 97 | 99 | 89 | 1e-227 |
| Streptococcus dysgalactiae subsp. equisimilis strain NCTC9413 | ska | Streptokinase | 10678902 | 98 | 100 | 85 | 1e-215 |
| Streptococcus dysgalactiae subsp. equisimilis strain NCTC9414 | ska | Streptokinase | 10678902 | 100 | 100 | 85 | 1e-219 |
| Streptococcus dysgalactiae subsp. equisimilis strain NCTC9414 | ska | Streptokinase | 10678902 | 100 | 100 | 89 | 1e-234 |
| Streptococcus dysgalactiae subsp. equisimilis strain NCTC9603 | ska | Streptokinase | 10678902 | 100 | 100 | 84 | 1e-216 |
| Streptococcus dysgalactiae subsp. equisimilis strain NCTC9603 | ska | Streptokinase | 10678902 | 100 | 100 | 87 | 1e-230 |
| Streptococcus dysgalactiae subsp. equisimilis strain SS1575 | ska | Streptokinase | 10678902 | 100 | 100 | 85 | 1e-219 |
| Streptococcus dysgalactiae subsp. equisimilis strain SS1575 | ska | Streptokinase | 10678902 | 100 | 100 | 89 | 1e-233 |
| Streptococcus dysgalactiae subsp. equisimilis strain T642 | ska | Streptokinase | 10678902 | 100 | 100 | 85 | 0.0 |
| Streptococcus dysgalactiae subsp. equisimilis strain T642 | ska | Streptokinase | 10678902 | 100 | 100 | 88 | 0.0 |
| Streptococcus dysgalactiae subsp. equisimilis strain UT_4031CC | ska | Streptokinase | 10678902 | 100 | 100 | 88 | 1e-232 |
| Streptococcus dysgalactiae subsp. equisimilis strain UT_4231_KK | ska | Streptokinase | 10678902 | 100 | 100 | 89 | 1e-233 |
| Streptococcus dysgalactiae subsp. equisimilis strain UT_4234_DH | ska | Streptokinase | 10678902 | 100 | 100 | 84 | 1e-216 |
| Streptococcus dysgalactiae subsp. equisimilis strain UT_4234_DH | ska | Streptokinase | 10678902 | 100 | 100 | 87 | 1e-228 |
| Streptococcus dysgalactiae subsp. equisimilis strain UT_4241_XS | ska | Streptokinase | 10678902 | 100 | 100 | 86 | 1e-221 |
| Streptococcus dysgalactiae subsp. equisimilis strain UT_4241_XS | ska | Streptokinase | 10678902 | 100 | 100 | 88 | 1e-231 |
| Streptococcus dysgalactiae subsp. equisimilis strain UT_4242_AB | ska | Streptokinase | 10678902 | 100 | 100 | 87 | 1e-230 |
| Streptococcus dysgalactiae subsp. equisimilis strain UT_4242_AB | ska | Streptokinase | 10678902 | 100 | 100 | 85 | 1e-220 |
| Streptococcus dysgalactiae subsp. equisimilis strain UT_4255RC | ska | Streptokinase | 10678902 | 100 | 100 | 84 | 1e-216 |
| Streptococcus dysgalactiae subsp. equisimilis strain UT_4255RC | ska | Streptokinase | 10678902 | 100 | 100 | 87 | 1e-228 |
| Streptococcus dysgalactiae subsp. equisimilis strain UT_4277_BB | ska | Streptokinase | 10678902 | 100 | 100 | 86 | 1e-221 |
| Streptococcus dysgalactiae subsp. equisimilis strain UT_4277_BB | ska | Streptokinase | 10678902 | 100 | 100 | 88 | 1e-231 |
| Streptococcus dysgalactiae subsp. equisimilis strain UT_4966_RC | ska | Streptokinase | 10678902 | 100 | 100 | 86 | 1e-221 |
| Streptococcus dysgalactiae subsp. equisimilis strain UT_4966_RC | ska | Streptokinase | 10678902 | 100 | 100 | 88 | 1e-231 |
| Streptococcus dysgalactiae subsp. equisimilis strain UT-5345 | ska | Streptokinase | 10678902 | 100 | 100 | 88 | 1e-231 |
| Streptococcus dysgalactiae subsp. equisimilis strain UT-5354 | ska | Streptokinase | 10678902 | 100 | 100 | 88 | 1e-232 |
| Streptococcus dysgalactiae subsp. equisimilis strain UT-5354 | ska | Streptokinase | 10678902 | 100 | 100 | 86 | 1e-221 |
| Streptococcus dysgalactiae subsp. equisimilis strain UT-SS1069 | ska | Streptokinase | 10678902 | 86 | 97 | 85 | 1e-188 |
| Streptococcus dysgalactiae subsp. equisimilis strain UT-SS1069 | ska | Streptokinase | 10678902 | 6 | 93 | 86 | 2e-05 |
| Streptococcus dysgalactiae subsp. equisimilis strain UT-SS1069 | ska | Streptokinase | 10678902 | 86 | 97 | 91 | 1e-206 |
| Streptococcus dysgalactiae subsp. equisimilis strain UT-SS1069 | ska | Streptokinase | 10678902 | 7 | 100 | 90 | 6e-07 |
| Streptococcus dysgalactiae subsp. equisimilis strain UT-SS957 | ska | Streptokinase | 10678902 | 100 | 100 | 87 | 1e-230 |
| Streptococcus dysgalactiae subsp. equisimilis strain WCHSDSE-1 | ska | Streptokinase | 10678902 | 100 | 100 | 88 | 1e-232 |
| Streptococcus dysgalactiae subsp. equisimilis 167 | sagA | Streptolysin S precursor (SagA) | 9529097 | 100 | 100 | 88 | 1e-22 |
| Streptococcus dysgalactiae subsp. equisimilis AKSDE4288 | sagA | Streptolysin S precursor (SagA) | 9529097 | 100 | 100 | 88 | 1e-22 |
| Streptococcus dysgalactiae subsp. equisimilis GGS_124 | sagA | Streptolysin S precursor (SagA) | 9529097 | 100 | 86 | 88 | 1e-22 |
| Streptococcus dysgalactiae subsp. equisimilis SD SCDR1 | sagA | Streptolysin S precursor (SagA) | 9529097 | 100 | 100 | 88 | 1e-22 |
| Streptococcus dysgalactiae subsp. equisimilis SK1249 | sagA | Streptolysin S precursor (SagA) | 9529097 | 100 | 100 | 88 | 1e-22 |
| Streptococcus dysgalactiae subsp. equisimilis strain ASDSE_96 | sagA | Streptolysin S precursor (SagA) | 9529097 | 100 | 100 | 88 | 1e-22 |
| Streptococcus dysgalactiae subsp. equisimilis strain ASDSE_99 | sagA | Streptolysin S precursor (SagA) | 9529097 | 100 | 100 | 88 | 1e-22 |
| Streptococcus dysgalactiae subsp. equisimilis strain C161L1 | sagA | Streptolysin S precursor (SagA) | 11801184 | 100 | 100 | 88 | 0.000000000000099999998245167 |
| Streptococcus dysgalactiae subsp. equisimilis strain KNZ01 | sagA | Streptolysin S precursor (SagA) | 9529097 | 100 | 100 | 88 | 1e-22 |
| Streptococcus dysgalactiae subsp. equisimilis strain KNZ03 | sagA | Streptolysin S precursor (SagA) | 9529097 | 100 | 100 | 88 | 1e-22 |
| Streptococcus dysgalactiae subsp. equisimilis strain KNZ04 | sagA | Streptolysin S precursor (SagA) | 9529097 | 100 | 100 | 88 | 1e-22 |
| Streptococcus dysgalactiae subsp. equisimilis strain KNZ06 | sagA | Streptolysin S precursor (SagA) | 9529097 | 100 | 100 | 88 | 1e-22 |
| Streptococcus dysgalactiae subsp. equisimilis strain KNZ07 | sagA | Streptolysin S precursor (SagA) | 9529097 | 100 | 100 | 88 | 1e-22 |
| Streptococcus dysgalactiae subsp. equisimilis strain KNZ10 | sagA | Streptolysin S precursor (SagA) | 9529097 | 100 | 100 | 88 | 1e-22 |
| Streptococcus dysgalactiae subsp. equisimilis strain KNZ12 | sagA | Streptolysin S precursor (SagA) | 9529097 | 100 | 100 | 88 | 1e-22 |
| Streptococcus dysgalactiae subsp. equisimilis strain KNZ15 | sagA | Streptolysin S precursor (SagA) | 9529097 | 100 | 100 | 88 | 1e-22 |
| Streptococcus dysgalactiae subsp. equisimilis strain KNZ16 | sagA | Streptolysin S precursor (SagA) | 9529097 | 100 | 100 | 88 | 1e-22 |
| Streptococcus dysgalactiae subsp. equisimilis strain NCTC10321 | sagA | Streptolysin S precursor (SagA) | 9529097 | 100 | 100 | 88 | 1e-22 |
| Streptococcus dysgalactiae subsp. equisimilis strain NCTC11554 | sagA | Streptolysin S precursor (SagA) | 9529097 | 100 | 100 | 88 | 1e-22 |
| Streptococcus dysgalactiae subsp. equisimilis strain NCTC11555 | sagA | Streptolysin S precursor (SagA) | 9529097 | 100 | 100 | 88 | 1e-22 |
| Streptococcus dysgalactiae subsp. equisimilis strain NCTC11556 | sagA | Streptolysin S precursor (SagA) | 9529097 | 100 | 100 | 88 | 1e-22 |
| Streptococcus dysgalactiae subsp. equisimilis strain NCTC11557 | sagA | Streptolysin S precursor (SagA) | 9529097 | 100 | 100 | 88 | 1e-22 |
| Streptococcus dysgalactiae subsp. equisimilis strain NCTC11564 | sagA | Streptolysin S precursor (SagA) | 9529097 | 100 | 100 | 88 | 1e-22 |
| Streptococcus dysgalactiae subsp. equisimilis strain NCTC5370 | sagA | Streptolysin S precursor (SagA) | 9529097 | 100 | 100 | 88 | 1e-22 |
| Streptococcus dysgalactiae subsp. equisimilis strain NCTC5371 | sagA | Streptolysin S precursor (SagA) | 9529097 | 100 | 100 | 88 | 1e-22 |
| Streptococcus dysgalactiae subsp. equisimilis strain NCTC6407 | sagA | Streptolysin S precursor (SagA) | 9529097 | 100 | 100 | 88 | 1e-22 |
| Streptococcus dysgalactiae subsp. equisimilis strain NCTC7136 | sagA | Streptolysin S precursor (SagA) | 9529097 | 100 | 100 | 88 | 1e-22 |
| Streptococcus dysgalactiae subsp. equisimilis strain NCTC8543 | sagA | Streptolysin S precursor (SagA) | 9529097 | 100 | 100 | 88 | 1e-22 |
| Streptococcus dysgalactiae subsp. equisimilis strain NCTC8546 | sagA | Streptolysin S precursor (SagA) | 9529097 | 100 | 100 | 88 | 1e-22 |
| Streptococcus dysgalactiae subsp. equisimilis strain NCTC9413 | sagA | Streptolysin S precursor (SagA) | 9529097 | 100 | 100 | 88 | 1e-22 |
| Streptococcus dysgalactiae subsp. equisimilis strain NCTC9414 | sagA | Streptolysin S precursor (SagA) | 9529097 | 100 | 100 | 88 | 1e-22 |
| Streptococcus dysgalactiae subsp. equisimilis strain NCTC9603 | sagA | Streptolysin S precursor (SagA) | 9529097 | 100 | 100 | 88 | 1e-22 |
| Streptococcus dysgalactiae subsp. equisimilis strain SS1575 | sagA | Streptolysin S precursor (SagA) | 9529097 | 100 | 100 | 88 | 1e-22 |
| Streptococcus dysgalactiae subsp. equisimilis strain T642 | sagA | Streptolysin S precursor (SagA) |  | 100 | 100 | 88 | 0.0000000000000599999989471002 |
| Streptococcus dysgalactiae subsp. equisimilis strain UT_4031CC | sagA | Streptolysin S precursor (SagA) | 9529097 | 100 | 100 | 88 | 1e-22 |
| Streptococcus dysgalactiae subsp. equisimilis strain UT_4234_DH | sagA | Streptolysin S precursor (SagA) | 9529097 | 100 | 100 | 88 | 1e-22 |
| Streptococcus dysgalactiae subsp. equisimilis strain UT_4241_XS | sagA | Streptolysin S precursor (SagA) | 9529097 | 100 | 100 | 88 | 1e-22 |
| Streptococcus dysgalactiae subsp. equisimilis strain UT_4242_AB | sagA | Streptolysin S precursor (SagA) | 9529097 | 100 | 100 | 88 | 1e-22 |
| Streptococcus dysgalactiae subsp. equisimilis strain UT_4255RC | sagA | Streptolysin S precursor (SagA) | 9529097 | 100 | 100 | 88 | 1e-22 |
| Streptococcus dysgalactiae subsp. equisimilis strain UT_4277_BB | sagA | Streptolysin S precursor (SagA) | 9529097 | 100 | 100 | 88 | 1e-22 |
| Streptococcus dysgalactiae subsp. equisimilis strain UT_4966_RC | sagA | Streptolysin S precursor (SagA) | 9529097 | 100 | 100 | 88 | 1e-22 |
| Streptococcus dysgalactiae subsp. equisimilis strain UT-5345 | sagA | Streptolysin S precursor (SagA) | 9529097 | 100 | 100 | 88 | 1e-22 |
| Streptococcus dysgalactiae subsp. equisimilis strain UT-5354 | sagA | Streptolysin S precursor (SagA) | 9529097 | 100 | 56 | 88 | 1e-22 |
| Streptococcus dysgalactiae subsp. equisimilis strain UT-SS1069 | sagA | Streptolysin S precursor (SagA) | 9529097 | 100 | 100 | 88 | 1e-22 |
| Streptococcus dysgalactiae subsp. equisimilis strain UT-SS957 | sagA | Streptolysin S precursor (SagA) | 9529097 | 100 | 100 | 88 | 1e-22 |
| Streptococcus dysgalactiae subsp. equisimilis strain WCHSDSE-1 | sagA | Streptolysin S precursor (SagA) | 9529097 | 100 | 100 | 88 | 1e-22 |
| Streptococcus dysgalactiae subsp. equisimilis 167 | slo | Thiol-activated cytolysin => streptolysin O | 11035749 | 63 | 100 | 99 | 1e-211 |
| Streptococcus dysgalactiae subsp. equisimilis 167 | slo | Thiol-activated cytolysin => streptolysin O | 11035749 | 63 | 100 | 99 | 1e-211 |
| Streptococcus dysgalactiae subsp. equisimilis AC-2713 | slo | Thiol-activated cytolysin => streptolysin O | 11035749 | 100 | 99 | 98 | 0.0 |
| Streptococcus dysgalactiae subsp. equisimilis AC-2713 | slo | Thiol-activated cytolysin => streptolysin O | 11035749 | 100 | 99 | 98 | 0.0 |
| Streptococcus dysgalactiae subsp. equisimilis AKSDE4288 | slo | Thiol-activated cytolysin => streptolysin O | 11035749 | 100 | 100 | 98 | 0.0 |
| Streptococcus dysgalactiae subsp. equisimilis AKSDE4288 | slo | Thiol-activated cytolysin => streptolysin O | 11035749 | 100 | 100 | 98 | 0.0 |
| Streptococcus dysgalactiae subsp. equisimilis ATCC 12394 | slo | Thiol-activated cytolysin => streptolysin O | 11035749 | 100 | 99 | 98 | 0.0 |
| Streptococcus dysgalactiae subsp. equisimilis ATCC 12394 | slo | Thiol-activated cytolysin => streptolysin O | 11035749 | 100 | 99 | 98 | 0.0 |
| Streptococcus dysgalactiae subsp. equisimilis GGS_124 | slo | Thiol-activated cytolysin => streptolysin O | 11035749 | 100 | 99 | 98 | 0.0 |
| Streptococcus dysgalactiae subsp. equisimilis GGS_124 | slo | Thiol-activated cytolysin => streptolysin O | 11035749 | 100 | 99 | 98 | 0.0 |
| Streptococcus dysgalactiae subsp. equisimilis RE378 | slo | Thiol-activated cytolysin => streptolysin O | 11035749 | 100 | 99 | 98 | 0.0 |
| Streptococcus dysgalactiae subsp. equisimilis RE378 | slo | Thiol-activated cytolysin => streptolysin O | 11035749 | 100 | 99 | 98 | 0.0 |
| Streptococcus dysgalactiae subsp. equisimilis SD SCDR1 | slo | Thiol-activated cytolysin => streptolysin O | 11035749 | 100 | 100 | 98 | 0.0 |
| Streptococcus dysgalactiae subsp. equisimilis SD SCDR1 | slo | Thiol-activated cytolysin => streptolysin O | 11035749 | 100 | 100 | 98 | 0.0 |
| Streptococcus dysgalactiae subsp. equisimilis SK1249 | pfo | Thiol-activated cytolysin => streptolysin O | 11035749 | 100 | 99 | 98 | 0.0 |
| Streptococcus dysgalactiae subsp. equisimilis SK1249 | pfo | Thiol-activated cytolysin => streptolysin O | 11035749 | 100 | 99 | 98 | 0.0 |
| Streptococcus dysgalactiae subsp. equisimilis SK1250 | pfo | Thiol-activated cytolysin => streptolysin O | 11035749 | 63 | 100 | 99 | 1e-211 |
| Streptococcus dysgalactiae subsp. equisimilis SK1250 | pfo | Thiol-activated cytolysin => streptolysin O | 11035749 | 63 | 100 | 99 | 1e-211 |
| Streptococcus dysgalactiae subsp. equisimilis strain ASDSE_96 | slo | Thiol-activated cytolysin => streptolysin O | 11035749 | 100 | 100 | 98 | 0.0 |
| Streptococcus dysgalactiae subsp. equisimilis strain ASDSE_96 | slo | Thiol-activated cytolysin => streptolysin O | 11035749 | 100 | 100 | 98 | 0.0 |
| Streptococcus dysgalactiae subsp. equisimilis strain ASDSE_99 | slo | Thiol-activated cytolysin => streptolysin O | 11035749 | 100 | 100 | 99 | 0.0 |
| Streptococcus dysgalactiae subsp. equisimilis strain ASDSE_99 | slo | Thiol-activated cytolysin => streptolysin O | 11035749 | 100 | 100 | 99 | 0.0 |
| Streptococcus dysgalactiae subsp. equisimilis strain C161L1 | slo | Thiol-activated cytolysin => streptolysin O | 11035749 | 100 | 100 | 98 | 0 |
| Streptococcus dysgalactiae subsp. equisimilis strain C161L1 | slo | Thiol-activated cytolysin => streptolysin O | 11035749 | 100 | 100 | 98 | 0 |
| Streptococcus dysgalactiae subsp. equisimilis strain KNZ01 | slo | Thiol-activated cytolysin => streptolysin O | 11035749 | 100 | 100 | 98 | 0.0 |
| Streptococcus dysgalactiae subsp. equisimilis strain KNZ01 | slo | Thiol-activated cytolysin => streptolysin O | 11035749 | 100 | 100 | 98 | 0.0 |
| Streptococcus dysgalactiae subsp. equisimilis strain KNZ01 | slo | Thiol-activated cytolysin => streptolysin O | 11035749 | 100 | 100 | 98 | 0.0 |
| Streptococcus dysgalactiae subsp. equisimilis strain KNZ01 | slo | Thiol-activated cytolysin => streptolysin O | 11035749 | 100 | 100 | 98 | 0.0 |
| Streptococcus dysgalactiae subsp. equisimilis strain KNZ03 | slo | Thiol-activated cytolysin => streptolysin O | 11035749 | 100 | 100 | 98 | 0.0 |
| Streptococcus dysgalactiae subsp. equisimilis strain KNZ03 | slo | Thiol-activated cytolysin => streptolysin O | 11035749 | 100 | 100 | 98 | 0.0 |
| Streptococcus dysgalactiae subsp. equisimilis strain KNZ03 | slo | Thiol-activated cytolysin => streptolysin O | 11035749 | 100 | 100 | 98 | 0.0 |
| Streptococcus dysgalactiae subsp. equisimilis strain KNZ03 | slo | Thiol-activated cytolysin => streptolysin O | 11035749 | 100 | 100 | 98 | 0.0 |
| Streptococcus dysgalactiae subsp. equisimilis strain KNZ04 | slo | Thiol-activated cytolysin => streptolysin O | 11035749 | 100 | 100 | 98 | 0.0 |
| Streptococcus dysgalactiae subsp. equisimilis strain KNZ04 | slo | Thiol-activated cytolysin => streptolysin O | 11035749 | 100 | 100 | 98 | 0.0 |
| Streptococcus dysgalactiae subsp. equisimilis strain KNZ06 | slo | Thiol-activated cytolysin => streptolysin O | 11035749 | 100 | 100 | 98 | 0.0 |
| Streptococcus dysgalactiae subsp. equisimilis strain KNZ06 | slo | Thiol-activated cytolysin => streptolysin O | 11035749 | 100 | 100 | 98 | 0.0 |
| Streptococcus dysgalactiae subsp. equisimilis strain KNZ07 | slo | Thiol-activated cytolysin => streptolysin O | 11035749 | 100 | 100 | 98 | 0.0 |
| Streptococcus dysgalactiae subsp. equisimilis strain KNZ07 | slo | Thiol-activated cytolysin => streptolysin O | 11035749 | 100 | 100 | 98 | 0.0 |
| Streptococcus dysgalactiae subsp. equisimilis strain KNZ10 | slo | Thiol-activated cytolysin => streptolysin O | 11035749 | 100 | 100 | 98 | 0.0 |
| Streptococcus dysgalactiae subsp. equisimilis strain KNZ10 | slo | Thiol-activated cytolysin => streptolysin O | 11035749 | 100 | 100 | 98 | 0.0 |
| Streptococcus dysgalactiae subsp. equisimilis strain KNZ12 | slo | Thiol-activated cytolysin => streptolysin O | 11035749 | 100 | 100 | 98 | 0.0 |
| Streptococcus dysgalactiae subsp. equisimilis strain KNZ12 | slo | Thiol-activated cytolysin => streptolysin O | 11035749 | 100 | 100 | 98 | 0.0 |
| Streptococcus dysgalactiae subsp. equisimilis strain KNZ15 | slo | Thiol-activated cytolysin => streptolysin O | 11035749 | 100 | 100 | 98 | 0.0 |
| Streptococcus dysgalactiae subsp. equisimilis strain KNZ15 | slo | Thiol-activated cytolysin => streptolysin O | 11035749 | 100 | 100 | 98 | 0.0 |
| Streptococcus dysgalactiae subsp. equisimilis strain KNZ16 | slo | Thiol-activated cytolysin => streptolysin O | 11035749 | 100 | 100 | 98 | 0.0 |
| Streptococcus dysgalactiae subsp. equisimilis strain KNZ16 | slo | Thiol-activated cytolysin => streptolysin O | 11035749 | 100 | 100 | 98 | 0.0 |
| Streptococcus dysgalactiae subsp. equisimilis strain NCTC11554 | slo | Thiol-activated cytolysin => streptolysin O | 11035749 | 100 | 100 | 98 | 0.0 |
| Streptococcus dysgalactiae subsp. equisimilis strain NCTC11554 | slo | Thiol-activated cytolysin => streptolysin O | 11035749 | 100 | 100 | 98 | 0.0 |
| Streptococcus dysgalactiae subsp. equisimilis strain NCTC11555 | slo | Thiol-activated cytolysin => streptolysin O | 11035749 | 100 | 100 | 99 | 0.0 |
| Streptococcus dysgalactiae subsp. equisimilis strain NCTC11555 | slo | Thiol-activated cytolysin => streptolysin O | 11035749 | 100 | 100 | 99 | 0.0 |
| Streptococcus dysgalactiae subsp. equisimilis strain NCTC11556 | slo | Thiol-activated cytolysin => streptolysin O | 11035749 | 100 | 100 | 98 | 0.0 |
| Streptococcus dysgalactiae subsp. equisimilis strain NCTC11556 | slo | Thiol-activated cytolysin => streptolysin O | 11035749 | 100 | 100 | 98 | 0.0 |
| Streptococcus dysgalactiae subsp. equisimilis strain NCTC11557 | slo | Thiol-activated cytolysin => streptolysin O | 11035749 | 100 | 100 | 99 | 0.0 |
| Streptococcus dysgalactiae subsp. equisimilis strain NCTC11557 | slo | Thiol-activated cytolysin => streptolysin O | 11035749 | 100 | 100 | 99 | 0.0 |
| Streptococcus dysgalactiae subsp. equisimilis strain NCTC11564 | slo | Thiol-activated cytolysin => streptolysin O | 11035749 | 100 | 100 | 98 | 0.0 |
| Streptococcus dysgalactiae subsp. equisimilis strain NCTC11564 | slo | Thiol-activated cytolysin => streptolysin O | 11035749 | 100 | 100 | 98 | 0.0 |
| Streptococcus dysgalactiae subsp. equisimilis strain NCTC5370 | slo | Thiol-activated cytolysin => streptolysin O | 11035749 | 100 | 100 | 98 | 0.0 |
| Streptococcus dysgalactiae subsp. equisimilis strain NCTC5370 | slo | Thiol-activated cytolysin => streptolysin O | 11035749 | 100 | 100 | 98 | 0.0 |
| Streptococcus dysgalactiae subsp. equisimilis strain NCTC5371 | slo | Thiol-activated cytolysin => streptolysin O | 11035749 | 100 | 100 | 98 | 0.0 |
| Streptococcus dysgalactiae subsp. equisimilis strain NCTC5371 | slo | Thiol-activated cytolysin => streptolysin O | 11035749 | 100 | 100 | 98 | 0.0 |
| Streptococcus dysgalactiae subsp. equisimilis strain NCTC7136 | slo | Thiol-activated cytolysin => streptolysin O | 11035749 | 100 | 100 | 98 | 0.0 |
| Streptococcus dysgalactiae subsp. equisimilis strain NCTC7136 | slo | Thiol-activated cytolysin => streptolysin O | 11035749 | 100 | 100 | 98 | 0.0 |
| Streptococcus dysgalactiae subsp. equisimilis strain NCTC8543 | slo | Thiol-activated cytolysin => streptolysin O | 11035749 | 100 | 100 | 98 | 0.0 |
| Streptococcus dysgalactiae subsp. equisimilis strain NCTC8543 | slo | Thiol-activated cytolysin => streptolysin O | 11035749 | 100 | 100 | 98 | 0.0 |
| Streptococcus dysgalactiae subsp. equisimilis strain NCTC8546 | slo | Thiol-activated cytolysin => streptolysin O | 11035749 | 100 | 100 | 98 | 0.0 |
| Streptococcus dysgalactiae subsp. equisimilis strain NCTC8546 | slo | Thiol-activated cytolysin => streptolysin O | 11035749 | 100 | 100 | 98 | 0.0 |
| Streptococcus dysgalactiae subsp. equisimilis strain NCTC9413 | slo | Thiol-activated cytolysin => streptolysin O | 11035749 | 100 | 100 | 98 | 0.0 |
| Streptococcus dysgalactiae subsp. equisimilis strain NCTC9413 | slo | Thiol-activated cytolysin => streptolysin O | 11035749 | 100 | 100 | 98 | 0.0 |
| Streptococcus dysgalactiae subsp. equisimilis strain NCTC9414 | slo | Thiol-activated cytolysin => streptolysin O | 11035749 | 100 | 100 | 98 | 0.0 |
| Streptococcus dysgalactiae subsp. equisimilis strain NCTC9414 | slo | Thiol-activated cytolysin => streptolysin O | 11035749 | 100 | 100 | 98 | 0.0 |
| Streptococcus dysgalactiae subsp. equisimilis strain NCTC9603 | slo | Thiol-activated cytolysin => streptolysin O | 11035749 | 100 | 100 | 98 | 0.0 |
| Streptococcus dysgalactiae subsp. equisimilis strain NCTC9603 | slo | Thiol-activated cytolysin => streptolysin O | 11035749 | 100 | 100 | 98 | 0.0 |
| Streptococcus dysgalactiae subsp. equisimilis strain SS1575 | slo | Thiol-activated cytolysin => streptolysin O | 11035749 | 100 | 100 | 98 | 0.0 |
| Streptococcus dysgalactiae subsp. equisimilis strain SS1575 | slo | Thiol-activated cytolysin => streptolysin O | 11035749 | 100 | 100 | 98 | 0.0 |
| Streptococcus dysgalactiae subsp. equisimilis strain T642 | slo | Thiol-activated cytolysin => streptolysin O | 11035749 | 100 | 100 | 98 | 0 |
| Streptococcus dysgalactiae subsp. equisimilis strain T642 | slo | Thiol-activated cytolysin => streptolysin O | 11035749 | 100 | 100 | 98 | 0 |
| Streptococcus dysgalactiae subsp. equisimilis strain UT_4031CC | slo | Thiol-activated cytolysin => streptolysin O | 11035749 | 100 | 100 | 98 | 0.0 |
| Streptococcus dysgalactiae subsp. equisimilis strain UT_4031CC | slo | Thiol-activated cytolysin => streptolysin O | 11035749 | 100 | 100 | 98 | 0.0 |
| Streptococcus dysgalactiae subsp. equisimilis strain UT_4231_KK | slo | Thiol-activated cytolysin => streptolysin O | 11035749 | 100 | 100 | 98 | 0.0 |
| Streptococcus dysgalactiae subsp. equisimilis strain UT_4231_KK | slo | Thiol-activated cytolysin => streptolysin O | 11035749 | 100 | 100 | 98 | 0.0 |
| Streptococcus dysgalactiae subsp. equisimilis strain UT_4234_DH | slo | Thiol-activated cytolysin => streptolysin O | 11035749 | 100 | 100 | 99 | 0.0 |
| Streptococcus dysgalactiae subsp. equisimilis strain UT_4234_DH | slo | Thiol-activated cytolysin => streptolysin O | 11035749 | 100 | 100 | 99 | 0.0 |
| Streptococcus dysgalactiae subsp. equisimilis strain UT_4241_XS | slo | Thiol-activated cytolysin => streptolysin O | 11035749 | 100 | 100 | 98 | 0.0 |
| Streptococcus dysgalactiae subsp. equisimilis strain UT_4241_XS | slo | Thiol-activated cytolysin => streptolysin O | 11035749 | 100 | 100 | 98 | 0.0 |
| Streptococcus dysgalactiae subsp. equisimilis strain UT_4242_AB | slo | Thiol-activated cytolysin => streptolysin O | 11035749 | 100 | 100 | 98 | 0.0 |
| Streptococcus dysgalactiae subsp. equisimilis strain UT_4242_AB | slo | Thiol-activated cytolysin => streptolysin O | 11035749 | 100 | 100 | 98 | 0.0 |
| Streptococcus dysgalactiae subsp. equisimilis strain UT_4255RC | slo | Thiol-activated cytolysin => streptolysin O | 11035749 | 100 | 100 | 99 | 0.0 |
| Streptococcus dysgalactiae subsp. equisimilis strain UT_4255RC | slo | Thiol-activated cytolysin => streptolysin O | 11035749 | 100 | 100 | 99 | 0.0 |
| Streptococcus dysgalactiae subsp. equisimilis strain UT_4277_BB | slo | Thiol-activated cytolysin => streptolysin O | 11035749 | 100 | 100 | 98 | 0.0 |
| Streptococcus dysgalactiae subsp. equisimilis strain UT_4277_BB | slo | Thiol-activated cytolysin => streptolysin O | 11035749 | 100 | 100 | 98 | 0.0 |
| Streptococcus dysgalactiae subsp. equisimilis strain UT_4966_RC | slo | Thiol-activated cytolysin => streptolysin O | 11035749 | 100 | 100 | 98 | 0.0 |
| Streptococcus dysgalactiae subsp. equisimilis strain UT_4966_RC | slo | Thiol-activated cytolysin => streptolysin O | 11035749 | 100 | 100 | 98 | 0.0 |
| Streptococcus dysgalactiae subsp. equisimilis strain UT-5345 | slo | Thiol-activated cytolysin => streptolysin O | 11035749 | 88 | 99 | 99 | 1e-295 |
| Streptococcus dysgalactiae subsp. equisimilis strain UT-5345 | slo | Thiol-activated cytolysin => streptolysin O | 11035749 | 88 | 99 | 99 | 1e-295 |
| Streptococcus dysgalactiae subsp. equisimilis strain UT-5354 | slo | Thiol-activated cytolysin => streptolysin O | 11035749 | 100 | 100 | 98 | 0.0 |
| Streptococcus dysgalactiae subsp. equisimilis strain UT-5354 | slo | Thiol-activated cytolysin => streptolysin O | 11035749 | 100 | 100 | 98 | 0.0 |
| Streptococcus dysgalactiae subsp. equisimilis strain UT-SS1069 | slo | Thiol-activated cytolysin => streptolysin O | 11035749 | 100 | 100 | 98 | 0.0 |
| Streptococcus dysgalactiae subsp. equisimilis strain UT-SS1069 | slo | Thiol-activated cytolysin => streptolysin O | 11035749 | 100 | 100 | 98 | 0.0 |
| Streptococcus dysgalactiae subsp. equisimilis strain UT-SS957 | slo | Thiol-activated cytolysin => streptolysin O | 11035749 | 100 | 100 | 98 | 0.0 |
| Streptococcus dysgalactiae subsp. equisimilis strain UT-SS957 | slo | Thiol-activated cytolysin => streptolysin O | 11035749 | 100 | 100 | 98 | 0.0 |
| Streptococcus dysgalactiae subsp. equisimilis strain WCHSDSE-1 | slo | Thiol-activated cytolysin => streptolysin O | 11035749 | 100 | 100 | 98 | 0.0 |
| Streptococcus dysgalactiae subsp. equisimilis strain WCHSDSE-1 | slo | Thiol-activated cytolysin => streptolysin O | 11035749 | 100 | 100 | 98 | 0.0 |
| Streptococcus dysgalactiae subsp. equisimilis 167 | lepA | Translation elongation factor LepA | 10998175 | 86 | 90 | 96 | 1e-294 |
| Streptococcus dysgalactiae subsp. equisimilis AC-2713 | lepA | Translation elongation factor LepA | 10998175 | 100 | 100 | 96 | 0.0 |
| Streptococcus dysgalactiae subsp. equisimilis AKSDE4288 | lepA | Translation elongation factor LepA | 10998175 | 100 | 100 | 96 | 0.0 |
| Streptococcus dysgalactiae subsp. equisimilis ATCC 12394 | lepA | Translation elongation factor LepA | 10998175 | 100 | 100 | 96 | 0.0 |
| Streptococcus dysgalactiae subsp. equisimilis GGS_124 | lepA | Translation elongation factor LepA | 10998175 | 100 | 100 | 96 | 0.0 |
| Streptococcus dysgalactiae subsp. equisimilis RE378 | lepA | Translation elongation factor LepA | 10998175 | 100 | 100 | 96 | 0.0 |
| Streptococcus dysgalactiae subsp. equisimilis SD SCDR1 | lepA | Translation elongation factor LepA | 10998175 | 100 | 100 | 96 | 0.0 |
| Streptococcus dysgalactiae subsp. equisimilis SK1249 | lepA | Translation elongation factor LepA | 10998175 | 100 | 100 | 96 | 0.0 |
| Streptococcus dysgalactiae subsp. equisimilis SK1250 | lepA | Translation elongation factor LepA | 10998175 | 86 | 90 | 96 | 1e-293 |
| Streptococcus dysgalactiae subsp. equisimilis strain ASDSE_96 | lepA | Translation elongation factor LepA | 10998175 | 100 | 100 | 96 | 0.0 |
| Streptococcus dysgalactiae subsp. equisimilis strain ASDSE_99 | lepA | Translation elongation factor LepA | 10998175 | 100 | 100 | 96 | 0.0 |
| Streptococcus dysgalactiae subsp. equisimilis strain C161L1 | lepA | Translation elongation factor LepA | 10998175 | 100 | 100 | 96 | 0 |
| Streptococcus dysgalactiae subsp. equisimilis strain KNZ01 | lepA | Translation elongation factor LepA | 10998175 | 100 | 100 | 96 | 0.0 |
| Streptococcus dysgalactiae subsp. equisimilis strain KNZ03 | lepA | Translation elongation factor LepA | 10998175 | 100 | 100 | 96 | 0.0 |
| Streptococcus dysgalactiae subsp. equisimilis strain KNZ04 | lepA | Translation elongation factor LepA | 10998175 | 100 | 100 | 96 | 0.0 |
| Streptococcus dysgalactiae subsp. equisimilis strain KNZ06 | lepA | Translation elongation factor LepA | 10998175 | 100 | 100 | 96 | 0.0 |
| Streptococcus dysgalactiae subsp. equisimilis strain KNZ07 | lepA | Translation elongation factor LepA | 10998175 | 100 | 100 | 96 | 0.0 |
| Streptococcus dysgalactiae subsp. equisimilis strain KNZ10 | lepA | Translation elongation factor LepA | 10998175 | 100 | 100 | 96 | 0.0 |
| Streptococcus dysgalactiae subsp. equisimilis strain KNZ12 | lepA | Translation elongation factor LepA | 10998175 | 100 | 100 | 96 | 0.0 |
| Streptococcus dysgalactiae subsp. equisimilis strain KNZ15 | lepA | Translation elongation factor LepA | 10998175 | 100 | 100 | 96 | 0.0 |
| Streptococcus dysgalactiae subsp. equisimilis strain KNZ16 | lepA | Translation elongation factor LepA | 10998175 | 100 | 100 | 96 | 0.0 |
| Streptococcus dysgalactiae subsp. equisimilis strain NCTC10321 | lepA | Translation elongation factor LepA | 10998175 | 100 | 100 | 96 | 0.0 |
| Streptococcus dysgalactiae subsp. equisimilis strain NCTC11554 | lepA | Translation elongation factor LepA | 10998175 | 100 | 100 | 96 | 0.0 |
| Streptococcus dysgalactiae subsp. equisimilis strain NCTC11555 | lepA | Translation elongation factor LepA | 10998175 | 100 | 100 | 96 | 0.0 |
| Streptococcus dysgalactiae subsp. equisimilis strain NCTC11556 | lepA | Translation elongation factor LepA | 10998175 | 100 | 100 | 96 | 0.0 |
| Streptococcus dysgalactiae subsp. equisimilis strain NCTC11557 | lepA | Translation elongation factor LepA | 10998175 | 100 | 100 | 96 | 0.0 |
| Streptococcus dysgalactiae subsp. equisimilis strain NCTC11564 | lepA | Translation elongation factor LepA | 10998175 | 100 | 100 | 96 | 0.0 |
| Streptococcus dysgalactiae subsp. equisimilis strain NCTC5370 | lepA | Translation elongation factor LepA | 10998175 | 100 | 100 | 96 | 0.0 |
| Streptococcus dysgalactiae subsp. equisimilis strain NCTC5371 | lepA | Translation elongation factor LepA | 10998175 | 100 | 100 | 96 | 0.0 |
| Streptococcus dysgalactiae subsp. equisimilis strain NCTC6179 | lepA | Translation elongation factor LepA | 10998175 | 100 | 100 | 96 | 0.0 |
| Streptococcus dysgalactiae subsp. equisimilis strain NCTC6181 | lepA | Translation elongation factor LepA | 10998175 | 100 | 100 | 96 | 0.0 |
| Streptococcus dysgalactiae subsp. equisimilis strain NCTC6407 | lepA | Translation elongation factor LepA | 10998175 | 100 | 100 | 96 | 0.0 |
| Streptococcus dysgalactiae subsp. equisimilis strain NCTC7136 | lepA | Translation elongation factor LepA | 10998175 | 100 | 100 | 96 | 0.0 |
| Streptococcus dysgalactiae subsp. equisimilis strain NCTC8543 | lepA | Translation elongation factor LepA | 10998175 | 100 | 100 | 96 | 0.0 |
| Streptococcus dysgalactiae subsp. equisimilis strain NCTC8546 | lepA | Translation elongation factor LepA | 10998175 | 100 | 100 | 96 | 0.0 |
| Streptococcus dysgalactiae subsp. equisimilis strain NCTC9413 | lepA | Translation elongation factor LepA | 10998175 | 100 | 100 | 96 | 0.0 |
| Streptococcus dysgalactiae subsp. equisimilis strain NCTC9414 | lepA | Translation elongation factor LepA | 10998175 | 100 | 100 | 96 | 0.0 |
| Streptococcus dysgalactiae subsp. equisimilis strain NCTC9603 | lepA | Translation elongation factor LepA | 10998175 | 100 | 100 | 96 | 0.0 |
| Streptococcus dysgalactiae subsp. equisimilis strain SS1575 | lepA | Translation elongation factor LepA | 10998175 | 100 | 100 | 96 | 0.0 |
| Streptococcus dysgalactiae subsp. equisimilis strain T642 | lepA | Translation elongation factor LepA | 10998175 | 100 | 100 | 96 | 0.0 |
| Streptococcus dysgalactiae subsp. equisimilis strain UT_4031CC | lepA | Translation elongation factor LepA | 10998175 | 100 | 100 | 96 | 0.0 |
| Streptococcus dysgalactiae subsp. equisimilis strain UT_4231_KK | lepA | Translation elongation factor LepA | 10998175 | 100 | 100 | 96 | 0.0 |
| Streptococcus dysgalactiae subsp. equisimilis strain UT_4234_DH | lepA | Translation elongation factor LepA | 10998175 | 100 | 100 | 96 | 0.0 |
| Streptococcus dysgalactiae subsp. equisimilis strain UT_4241_XS | lepA | Translation elongation factor LepA | 10998175 | 100 | 100 | 96 | 0.0 |
| Streptococcus dysgalactiae subsp. equisimilis strain UT_4242_AB | lepA | Translation elongation factor LepA | 10998175 | 100 | 100 | 96 | 0.0 |
| Streptococcus dysgalactiae subsp. equisimilis strain UT_4255RC | lepA | Translation elongation factor LepA | 10998175 | 100 | 100 | 96 | 0.0 |
| Streptococcus dysgalactiae subsp. equisimilis strain UT_4277_BB | lepA | Translation elongation factor LepA | 10998175 | 100 | 100 | 96 | 0.0 |
| Streptococcus dysgalactiae subsp. equisimilis strain UT_4966_RC | lepA | Translation elongation factor LepA | 10998175 | 100 | 100 | 96 | 0.0 |
| Streptococcus dysgalactiae subsp. equisimilis strain UT-5345 | lepA | Translation elongation factor LepA | 10998175 | 100 | 100 | 96 | 0.0 |
| Streptococcus dysgalactiae subsp. equisimilis strain UT-5354 | lepA | Translation elongation factor LepA | 10998175 | 100 | 100 | 96 | 0.0 |
| Streptococcus dysgalactiae subsp. equisimilis strain UT-SS1069 | lepA | Translation elongation factor LepA | 10998175 | 100 | 100 | 96 | 0.0 |
| Streptococcus dysgalactiae subsp. equisimilis strain UT-SS957 | lepA | Translation elongation factor LepA | 10998175 | 100 | 100 | 96 | 0.0 |
| Streptococcus dysgalactiae subsp. equisimilis strain WCHSDSE-1 | lepA | Translation elongation factor LepA | 10998175 | 100 | 100 | 96 | 0.0 |
| Streptococcus dysgalactiae subsp. equisimilis 167 | covS | Transmembrane histidine kinase CsrS | 20507231 | 99 | 98 | 93 | 1e-245 |
| Streptococcus dysgalactiae subsp. equisimilis AC-2713 | covS | Transmembrane histidine kinase CsrS | 20507231 | 99 | 90 | 93 | 1e-245 |
| Streptococcus dysgalactiae subsp. equisimilis AKSDE4288 | covS | Transmembrane histidine kinase CsrS | 20507231 | 99 | 90 | 93 | 1e-245 |
| Streptococcus dysgalactiae subsp. equisimilis ATCC 12394 | covS | Transmembrane histidine kinase CsrS | 20507231 | 99 | 90 | 93 | 1e-245 |
| Streptococcus dysgalactiae subsp. equisimilis GGS_124 | covS | Transmembrane histidine kinase CsrS | 20507231 | 99 | 90 | 93 | 1e-245 |
| Streptococcus dysgalactiae subsp. equisimilis RE378 | covS | Transmembrane histidine kinase CsrS | 20507231 | 99 | 90 | 93 | 1e-245 |
| Streptococcus dysgalactiae subsp. equisimilis SD SCDR1 | covS | Transmembrane histidine kinase CsrS | 20507231 | 99 | 90 | 93 | 1e-245 |
| Streptococcus dysgalactiae subsp. equisimilis SK1249 | covS | Transmembrane histidine kinase CsrS | 20507231 | 99 | 90 | 93 | 1e-245 |
| Streptococcus dysgalactiae subsp. equisimilis SK1250 | covS | Transmembrane histidine kinase CsrS | 20507231 | 91 | 99 | 93 | 1e-227 |
| Streptococcus dysgalactiae subsp. equisimilis strain ASDSE_96 | covS | Transmembrane histidine kinase CsrS | 20507231 | 99 | 90 | 93 | 1e-245 |
| Streptococcus dysgalactiae subsp. equisimilis strain ASDSE_99 | covS | Transmembrane histidine kinase CsrS | 20507231 | 99 | 90 | 93 | 1e-244 |
| Streptococcus dysgalactiae subsp. equisimilis strain C161L1 | covS | Transmembrane histidine kinase CsrS | 20507231 | 100 | 90 | 92 | 0 |
| Streptococcus dysgalactiae subsp. equisimilis strain KNZ01 | covS | Transmembrane histidine kinase CsrS | 20507231 | 99 | 90 | 93 | 1e-245 |
| Streptococcus dysgalactiae subsp. equisimilis strain KNZ03 | covS | Transmembrane histidine kinase CsrS | 20507231 | 99 | 90 | 93 | 1e-245 |
| Streptococcus dysgalactiae subsp. equisimilis strain KNZ04 | covS | Transmembrane histidine kinase CsrS | 20507231 | 99 | 90 | 93 | 1e-245 |
| Streptococcus dysgalactiae subsp. equisimilis strain KNZ06 | covS | Transmembrane histidine kinase CsrS | 20507231 | 99 | 90 | 93 | 1e-244 |
| Streptococcus dysgalactiae subsp. equisimilis strain KNZ07 | covS | Transmembrane histidine kinase CsrS | 20507231 | 99 | 90 | 93 | 1e-245 |
| Streptococcus dysgalactiae subsp. equisimilis strain KNZ10 | covS | Transmembrane histidine kinase CsrS | 20507231 | 99 | 90 | 93 | 1e-245 |
| Streptococcus dysgalactiae subsp. equisimilis strain KNZ12 | covS | Transmembrane histidine kinase CsrS | 20507231 | 99 | 90 | 93 | 1e-245 |
| Streptococcus dysgalactiae subsp. equisimilis strain KNZ15 | covS | Transmembrane histidine kinase CsrS | 20507231 | 99 | 90 | 93 | 1e-245 |
| Streptococcus dysgalactiae subsp. equisimilis strain KNZ16 | covS | Transmembrane histidine kinase CsrS | 20507231 | 99 | 90 | 93 | 1e-245 |
| Streptococcus dysgalactiae subsp. equisimilis strain NCTC10321 | covS | Transmembrane histidine kinase CsrS | 20507231 | 99 | 90 | 92 | 1e-246 |
| Streptococcus dysgalactiae subsp. equisimilis strain NCTC11554 | covS | Transmembrane histidine kinase CsrS | 20507231 | 99 | 90 | 93 | 1e-245 |
| Streptococcus dysgalactiae subsp. equisimilis strain NCTC11555 | covS | Transmembrane histidine kinase CsrS | 20507231 | 99 | 90 | 93 | 1e-245 |
| Streptococcus dysgalactiae subsp. equisimilis strain NCTC11556 | covS | Transmembrane histidine kinase CsrS | 20507231 | 99 | 90 | 92 | 1e-245 |
| Streptococcus dysgalactiae subsp. equisimilis strain NCTC11557 | covS | Transmembrane histidine kinase CsrS | 20507231 | 99 | 90 | 93 | 1e-245 |
| Streptococcus dysgalactiae subsp. equisimilis strain NCTC11564 | covS | Transmembrane histidine kinase CsrS | 20507231 | 99 | 90 | 93 | 1e-245 |
| Streptococcus dysgalactiae subsp. equisimilis strain NCTC5370 | covS | Transmembrane histidine kinase CsrS | 20507231 | 99 | 90 | 93 | 1e-245 |
| Streptococcus dysgalactiae subsp. equisimilis strain NCTC5371 | covS | Transmembrane histidine kinase CsrS | 20507231 | 99 | 90 | 92 | 1e-244 |
| Streptococcus dysgalactiae subsp. equisimilis strain NCTC6179 | covS | Transmembrane histidine kinase CsrS | 20507231 | 99 | 90 | 92 | 1e-245 |
| Streptococcus dysgalactiae subsp. equisimilis strain NCTC6181 | covS | Transmembrane histidine kinase CsrS | 20507231 | 99 | 90 | 92 | 1e-245 |
| Streptococcus dysgalactiae subsp. equisimilis strain NCTC6407 | covS | Transmembrane histidine kinase CsrS | 20507231 | 99 | 90 | 93 | 1e-246 |
| Streptococcus dysgalactiae subsp. equisimilis strain NCTC7136 | covS | Transmembrane histidine kinase CsrS | 20507231 | 99 | 90 | 93 | 1e-245 |
| Streptococcus dysgalactiae subsp. equisimilis strain NCTC8543 | covS | Transmembrane histidine kinase CsrS | 20507231 | 99 | 90 | 93 | 1e-245 |
| Streptococcus dysgalactiae subsp. equisimilis strain NCTC8546 | covS | Transmembrane histidine kinase CsrS | 20507231 | 99 | 90 | 93 | 1e-245 |
| Streptococcus dysgalactiae subsp. equisimilis strain NCTC9413 | covS | Transmembrane histidine kinase CsrS | 20507231 | 99 | 90 | 93 | 1e-245 |
| Streptococcus dysgalactiae subsp. equisimilis strain NCTC9414 | covS | Transmembrane histidine kinase CsrS | 20507231 | 99 | 90 | 93 | 1e-245 |
| Streptococcus dysgalactiae subsp. equisimilis strain NCTC9603 | covS | Transmembrane histidine kinase CsrS | 20507231 | 99 | 90 | 93 | 1e-245 |
| Streptococcus dysgalactiae subsp. equisimilis strain SS1575 | covS | Transmembrane histidine kinase CsrS | 20507231 | 99 | 90 | 93 | 1e-245 |
| Streptococcus dysgalactiae subsp. equisimilis strain T642 | covS | Transmembrane histidine kinase CsrS | 20507231 | 100 | 90 | 92 | 0 |
| Streptococcus dysgalactiae subsp. equisimilis strain UT_4031CC | covS | Transmembrane histidine kinase CsrS | 20507231 | 99 | 90 | 93 | 1e-245 |
| Streptococcus dysgalactiae subsp. equisimilis strain UT_4231_KK | covS | Transmembrane histidine kinase CsrS | 20507231 | 99 | 90 | 93 | 1e-245 |
| Streptococcus dysgalactiae subsp. equisimilis strain UT_4234_DH | covS | Transmembrane histidine kinase CsrS | 20507231 | 99 | 90 | 93 | 1e-245 |
| Streptococcus dysgalactiae subsp. equisimilis strain UT_4241_XS | covS | Transmembrane histidine kinase CsrS | 20507231 | 99 | 90 | 93 | 1e-245 |
| Streptococcus dysgalactiae subsp. equisimilis strain UT_4242_AB | covS | Transmembrane histidine kinase CsrS | 20507231 | 99 | 90 | 92 | 1e-245 |
| Streptococcus dysgalactiae subsp. equisimilis strain UT_4255RC | covS | Transmembrane histidine kinase CsrS | 20507231 | 99 | 90 | 92 | 1e-243 |
| Streptococcus dysgalactiae subsp. equisimilis strain UT_4277_BB | covS | Transmembrane histidine kinase CsrS | 20507231 | 99 | 90 | 93 | 1e-245 |
| Streptococcus dysgalactiae subsp. equisimilis strain UT_4966_RC | covS | Transmembrane histidine kinase CsrS | 20507231 | 99 | 90 | 93 | 1e-245 |
| Streptococcus dysgalactiae subsp. equisimilis strain UT-5345 | covS | Transmembrane histidine kinase CsrS | 20507231 | 99 | 90 | 93 | 1e-245 |
| Streptococcus dysgalactiae subsp. equisimilis strain UT-5354 | covS | Transmembrane histidine kinase CsrS | 20507231 | 99 | 90 | 93 | 1e-245 |
| Streptococcus dysgalactiae subsp. equisimilis strain UT-SS1069 | covS | Transmembrane histidine kinase CsrS | 20507231 | 99 | 90 | 93 | 1e-245 |
| Streptococcus dysgalactiae subsp. equisimilis strain UT-SS957 | covS | Transmembrane histidine kinase CsrS | 20507231 | 99 | 90 | 93 | 1e-245 |
| Streptococcus dysgalactiae subsp. equisimilis strain WCHSDSE-1 | covS | Transmembrane histidine kinase CsrS | 20507231 | 99 | 90 | 93 | 1e-245 |
| Streptococcus dysgalactiae subsp. equisimilis 167 | gidA | tRNA-5-carboxymethylaminomethyl-2-thiouridine(34) synthesis protein MnmG | 18426891 | 99 | 99 | 98 | 0.0 |
| Streptococcus dysgalactiae subsp. equisimilis AC-2713 | gidA | tRNA-5-carboxymethylaminomethyl-2-thiouridine(34) synthesis protein MnmG | 18426891 | 99 | 99 | 98 | 0.0 |
| Streptococcus dysgalactiae subsp. equisimilis AKSDE4288 | gidA | tRNA-5-carboxymethylaminomethyl-2-thiouridine(34) synthesis protein MnmG | 18426891 | 99 | 99 | 98 | 0.0 |
| Streptococcus dysgalactiae subsp. equisimilis ATCC 12394 | gidA | tRNA-5-carboxymethylaminomethyl-2-thiouridine(34) synthesis protein MnmG | 18426891 | 99 | 99 | 98 | 0.0 |
| Streptococcus dysgalactiae subsp. equisimilis GGS_124 | gidA | tRNA-5-carboxymethylaminomethyl-2-thiouridine(34) synthesis protein MnmG | 18426891 | 99 | 99 | 98 | 0.0 |
| Streptococcus dysgalactiae subsp. equisimilis RE378 | gidA | tRNA-5-carboxymethylaminomethyl-2-thiouridine(34) synthesis protein MnmG | 18426891 | 90 | 99 | 98 | 0.0 |
| Streptococcus dysgalactiae subsp. equisimilis SD SCDR1 | gidA | tRNA-5-carboxymethylaminomethyl-2-thiouridine(34) synthesis protein MnmG | 18426891 | 99 | 99 | 98 | 0.0 |
| Streptococcus dysgalactiae subsp. equisimilis SK1249 | gidA | tRNA-5-carboxymethylaminomethyl-2-thiouridine(34) synthesis protein MnmG | 18426891 | 99 | 99 | 98 | 0.0 |
| Streptococcus dysgalactiae subsp. equisimilis SK1250 | gidA | tRNA-5-carboxymethylaminomethyl-2-thiouridine(34) synthesis protein MnmG | 18426891 | 99 | 99 | 98 | 0.0 |
| Streptococcus dysgalactiae subsp. equisimilis strain ASDSE_96 | gidA | tRNA-5-carboxymethylaminomethyl-2-thiouridine(34) synthesis protein MnmG | 18426891 | 99 | 99 | 98 | 0.0 |
| Streptococcus dysgalactiae subsp. equisimilis strain ASDSE_99 | gidA | tRNA-5-carboxymethylaminomethyl-2-thiouridine(34) synthesis protein MnmG | 18426891 | 99 | 99 | 98 | 0.0 |
| Streptococcus dysgalactiae subsp. equisimilis strain C161L1 | gidA | tRNA-5-carboxymethylaminomethyl-2-thiouridine(34) synthesis protein MnmG | 18426891 | 99 | 99 | 98 | 0 |
| Streptococcus dysgalactiae subsp. equisimilis strain KNZ01 | gidA | tRNA-5-carboxymethylaminomethyl-2-thiouridine(34) synthesis protein MnmG | 18426891 | 99 | 99 | 98 | 0.0 |
| Streptococcus dysgalactiae subsp. equisimilis strain KNZ03 | gidA | tRNA-5-carboxymethylaminomethyl-2-thiouridine(34) synthesis protein MnmG | 18426891 | 99 | 99 | 98 | 0.0 |
| Streptococcus dysgalactiae subsp. equisimilis strain KNZ04 | gidA | tRNA-5-carboxymethylaminomethyl-2-thiouridine(34) synthesis protein MnmG | 18426891 | 99 | 99 | 98 | 0.0 |
| Streptococcus dysgalactiae subsp. equisimilis strain KNZ06 | gidA | tRNA-5-carboxymethylaminomethyl-2-thiouridine(34) synthesis protein MnmG | 18426891 | 99 | 99 | 98 | 0.0 |
| Streptococcus dysgalactiae subsp. equisimilis strain KNZ07 | gidA | tRNA-5-carboxymethylaminomethyl-2-thiouridine(34) synthesis protein MnmG | 18426891 | 99 | 99 | 98 | 0.0 |
| Streptococcus dysgalactiae subsp. equisimilis strain KNZ10 | gidA | tRNA-5-carboxymethylaminomethyl-2-thiouridine(34) synthesis protein MnmG | 18426891 | 99 | 99 | 98 | 0.0 |
| Streptococcus dysgalactiae subsp. equisimilis strain KNZ12 | gidA | tRNA-5-carboxymethylaminomethyl-2-thiouridine(34) synthesis protein MnmG | 18426891 | 99 | 99 | 98 | 0.0 |
| Streptococcus dysgalactiae subsp. equisimilis strain KNZ15 | gidA | tRNA-5-carboxymethylaminomethyl-2-thiouridine(34) synthesis protein MnmG | 18426891 | 99 | 99 | 98 | 0.0 |
| Streptococcus dysgalactiae subsp. equisimilis strain KNZ16 | gidA | tRNA-5-carboxymethylaminomethyl-2-thiouridine(34) synthesis protein MnmG | 18426891 | 99 | 99 | 98 | 0.0 |
| Streptococcus dysgalactiae subsp. equisimilis strain NCTC10321 | gidA | tRNA-5-carboxymethylaminomethyl-2-thiouridine(34) synthesis protein MnmG | 18426891 | 99 | 99 | 98 | 0.0 |
| Streptococcus dysgalactiae subsp. equisimilis strain NCTC11554 | gidA | tRNA-5-carboxymethylaminomethyl-2-thiouridine(34) synthesis protein MnmG | 18426891 | 99 | 99 | 98 | 0.0 |
| Streptococcus dysgalactiae subsp. equisimilis strain NCTC11555 | gidA | tRNA-5-carboxymethylaminomethyl-2-thiouridine(34) synthesis protein MnmG | 18426891 | 99 | 99 | 98 | 0.0 |
| Streptococcus dysgalactiae subsp. equisimilis strain NCTC11556 | gidA | tRNA-5-carboxymethylaminomethyl-2-thiouridine(34) synthesis protein MnmG | 18426891 | 99 | 99 | 98 | 0.0 |
| Streptococcus dysgalactiae subsp. equisimilis strain NCTC11557 | gidA | tRNA-5-carboxymethylaminomethyl-2-thiouridine(34) synthesis protein MnmG | 18426891 | 99 | 99 | 98 | 0.0 |
| Streptococcus dysgalactiae subsp. equisimilis strain NCTC11564 | gidA | tRNA-5-carboxymethylaminomethyl-2-thiouridine(34) synthesis protein MnmG | 18426891 | 99 | 99 | 98 | 0.0 |
| Streptococcus dysgalactiae subsp. equisimilis strain NCTC5370 | gidA | tRNA-5-carboxymethylaminomethyl-2-thiouridine(34) synthesis protein MnmG | 18426891 | 99 | 99 | 98 | 0.0 |
| Streptococcus dysgalactiae subsp. equisimilis strain NCTC5371 | gidA | tRNA-5-carboxymethylaminomethyl-2-thiouridine(34) synthesis protein MnmG | 18426891 | 99 | 99 | 98 | 0.0 |
| Streptococcus dysgalactiae subsp. equisimilis strain NCTC6179 | gidA | tRNA-5-carboxymethylaminomethyl-2-thiouridine(34) synthesis protein MnmG | 18426891 | 99 | 99 | 98 | 0.0 |
| Streptococcus dysgalactiae subsp. equisimilis strain NCTC6181 | gidA | tRNA-5-carboxymethylaminomethyl-2-thiouridine(34) synthesis protein MnmG | 18426891 | 99 | 99 | 98 | 0.0 |
| Streptococcus dysgalactiae subsp. equisimilis strain NCTC6407 | gidA | tRNA-5-carboxymethylaminomethyl-2-thiouridine(34) synthesis protein MnmG | 18426891 | 99 | 99 | 98 | 0.0 |
| Streptococcus dysgalactiae subsp. equisimilis strain NCTC7136 | gidA | tRNA-5-carboxymethylaminomethyl-2-thiouridine(34) synthesis protein MnmG | 18426891 | 99 | 99 | 98 | 0.0 |
| Streptococcus dysgalactiae subsp. equisimilis strain NCTC8543 | gidA | tRNA-5-carboxymethylaminomethyl-2-thiouridine(34) synthesis protein MnmG | 18426891 | 99 | 99 | 98 | 0.0 |
| Streptococcus dysgalactiae subsp. equisimilis strain NCTC8546 | gidA | tRNA-5-carboxymethylaminomethyl-2-thiouridine(34) synthesis protein MnmG | 18426891 | 99 | 99 | 98 | 0.0 |
| Streptococcus dysgalactiae subsp. equisimilis strain NCTC9413 | gidA | tRNA-5-carboxymethylaminomethyl-2-thiouridine(34) synthesis protein MnmG | 18426891 | 99 | 99 | 98 | 0.0 |
| Streptococcus dysgalactiae subsp. equisimilis strain NCTC9414 | gidA | tRNA-5-carboxymethylaminomethyl-2-thiouridine(34) synthesis protein MnmG | 18426891 | 99 | 99 | 98 | 0.0 |
| Streptococcus dysgalactiae subsp. equisimilis strain NCTC9603 | gidA | tRNA-5-carboxymethylaminomethyl-2-thiouridine(34) synthesis protein MnmG | 18426891 | 99 | 99 | 98 | 0.0 |
| Streptococcus dysgalactiae subsp. equisimilis strain SS1575 | gidA | tRNA-5-carboxymethylaminomethyl-2-thiouridine(34) synthesis protein MnmG | 18426891 | 99 | 99 | 98 | 0.0 |
| Streptococcus dysgalactiae subsp. equisimilis strain T642 | gidA | tRNA-5-carboxymethylaminomethyl-2-thiouridine(34) synthesis protein MnmG | 18426891 | 99 | 99 | 98 | 0 |
| Streptococcus dysgalactiae subsp. equisimilis strain UT_4031CC | gidA | tRNA-5-carboxymethylaminomethyl-2-thiouridine(34) synthesis protein MnmG | 18426891 | 99 | 99 | 98 | 0.0 |
| Streptococcus dysgalactiae subsp. equisimilis strain UT_4231_KK | gidA | tRNA-5-carboxymethylaminomethyl-2-thiouridine(34) synthesis protein MnmG | 18426891 | 99 | 99 | 98 | 0.0 |
| Streptococcus dysgalactiae subsp. equisimilis strain UT_4234_DH | gidA | tRNA-5-carboxymethylaminomethyl-2-thiouridine(34) synthesis protein MnmG | 18426891 | 99 | 99 | 98 | 0.0 |
| Streptococcus dysgalactiae subsp. equisimilis strain UT_4241_XS | gidA | tRNA-5-carboxymethylaminomethyl-2-thiouridine(34) synthesis protein MnmG | 18426891 | 99 | 99 | 98 | 0.0 |
| Streptococcus dysgalactiae subsp. equisimilis strain UT_4242_AB | gidA | tRNA-5-carboxymethylaminomethyl-2-thiouridine(34) synthesis protein MnmG | 18426891 | 99 | 99 | 98 | 0.0 |
| Streptococcus dysgalactiae subsp. equisimilis strain UT_4255RC | gidA | tRNA-5-carboxymethylaminomethyl-2-thiouridine(34) synthesis protein MnmG | 18426891 | 99 | 99 | 98 | 0.0 |
| Streptococcus dysgalactiae subsp. equisimilis strain UT_4277_BB | gidA | tRNA-5-carboxymethylaminomethyl-2-thiouridine(34) synthesis protein MnmG | 18426891 | 99 | 99 | 98 | 0.0 |
| Streptococcus dysgalactiae subsp. equisimilis strain UT_4966_RC | gidA | tRNA-5-carboxymethylaminomethyl-2-thiouridine(34) synthesis protein MnmG | 18426891 | 99 | 99 | 98 | 0.0 |
| Streptococcus dysgalactiae subsp. equisimilis strain UT-5345 | gidA | tRNA-5-carboxymethylaminomethyl-2-thiouridine(34) synthesis protein MnmG | 18426891 | 99 | 99 | 98 | 0.0 |
| Streptococcus dysgalactiae subsp. equisimilis strain UT-5354 | gidA | tRNA-5-carboxymethylaminomethyl-2-thiouridine(34) synthesis protein MnmG | 18426891 | 99 | 99 | 98 | 0.0 |
| Streptococcus dysgalactiae subsp. equisimilis strain UT-SS1069 | gidA | tRNA-5-carboxymethylaminomethyl-2-thiouridine(34) synthesis protein MnmG | 18426891 | 99 | 99 | 98 | 0.0 |
| Streptococcus dysgalactiae subsp. equisimilis strain UT-SS957 | gidA | tRNA-5-carboxymethylaminomethyl-2-thiouridine(34) synthesis protein MnmG | 18426891 | 99 | 99 | 98 | 0.0 |
| Streptococcus dysgalactiae subsp. equisimilis strain WCHSDSE-1 | gidA | tRNA-5-carboxymethylaminomethyl-2-thiouridine(34) synthesis protein MnmG | 18426891 | 99 | 99 | 98 | 0.0 |
| Streptococcus dysgalactiae subsp. equisimilis 167 | vicK | Two-component sensor kinase SA14-24 | 19088917 | 100 | 100 | 93 | 1e-242 |
| Streptococcus dysgalactiae subsp. equisimilis AC-2713 | vicK | Two-component sensor kinase SA14-24 | 19088917 | 100 | 100 | 93 | 1e-242 |
| Streptococcus dysgalactiae subsp. equisimilis AKSDE4288 | vicK | Two-component sensor kinase SA14-24 | 19088917 | 100 | 100 | 93 | 1e-242 |
| Streptococcus dysgalactiae subsp. equisimilis ATCC 12394 | vicK | Two-component sensor kinase SA14-24 | 19088917 | 100 | 100 | 93 | 1e-242 |
| Streptococcus dysgalactiae subsp. equisimilis GGS_124 | vicK | Two-component sensor kinase SA14-24 | 19088917 | 100 | 100 | 92 | 1e-241 |
| Streptococcus dysgalactiae subsp. equisimilis RE378 | vicK | Two-component sensor kinase SA14-24 | 19088917 | 84 | 100 | 92 | 1e-200 |
| Streptococcus dysgalactiae subsp. equisimilis SD SCDR1 | vicK | Two-component sensor kinase SA14-24 | 19088917 | 100 | 100 | 93 | 1e-242 |
| Streptococcus dysgalactiae subsp. equisimilis SK1249 | vicK | Two-component sensor kinase SA14-24 | 19088917 | 100 | 100 | 92 | 1e-241 |
| Streptococcus dysgalactiae subsp. equisimilis SK1250 | vicK | Two-component sensor kinase SA14-24 | 19088917 | 100 | 100 | 93 | 1e-242 |
| Streptococcus dysgalactiae subsp. equisimilis strain ASDSE_96 | vicK | Two-component sensor kinase SA14-24 | 19088917 | 100 | 100 | 93 | 1e-243 |
| Streptococcus dysgalactiae subsp. equisimilis strain ASDSE_99 | vicK | Two-component sensor kinase SA14-24 | 19088917 | 100 | 100 | 93 | 1e-242 |
| Streptococcus dysgalactiae subsp. equisimilis strain C161L1 | vicK | Two-component sensor kinase SA14-24 | 19088917 | 100 | 100 | 93 | 0 |
| Streptococcus dysgalactiae subsp. equisimilis strain KNZ01 | vicK | Two-component sensor kinase SA14-24 | 19088917 | 100 | 100 | 93 | 1e-242 |
| Streptococcus dysgalactiae subsp. equisimilis strain KNZ03 | vicK | Two-component sensor kinase SA14-24 | 19088917 | 100 | 100 | 93 | 1e-242 |
| Streptococcus dysgalactiae subsp. equisimilis strain KNZ04 | vicK | Two-component sensor kinase SA14-24 | 19088917 | 100 | 100 | 93 | 1e-242 |
| Streptococcus dysgalactiae subsp. equisimilis strain KNZ06 | vicK | Two-component sensor kinase SA14-24 | 19088917 | 100 | 100 | 93 | 1e-242 |
| Streptococcus dysgalactiae subsp. equisimilis strain KNZ07 | vicK | Two-component sensor kinase SA14-24 | 19088917 | 100 | 100 | 93 | 1e-242 |
| Streptococcus dysgalactiae subsp. equisimilis strain KNZ10 | vicK | Two-component sensor kinase SA14-24 | 19088917 | 100 | 100 | 93 | 1e-242 |
| Streptococcus dysgalactiae subsp. equisimilis strain KNZ12 | vicK | Two-component sensor kinase SA14-24 | 19088917 | 100 | 100 | 93 | 1e-242 |
| Streptococcus dysgalactiae subsp. equisimilis strain KNZ15 | vicK | Two-component sensor kinase SA14-24 | 19088917 | 100 | 100 | 93 | 1e-242 |
| Streptococcus dysgalactiae subsp. equisimilis strain KNZ16 | vicK | Two-component sensor kinase SA14-24 | 19088917 | 100 | 100 | 93 | 1e-242 |
| Streptococcus dysgalactiae subsp. equisimilis strain NCTC10321 | vicK | Two-component sensor kinase SA14-24 | 19088917 | 100 | 100 | 93 | 1e-243 |
| Streptococcus dysgalactiae subsp. equisimilis strain NCTC11554 | vicK | Two-component sensor kinase SA14-24 | 19088917 | 100 | 100 | 93 | 1e-242 |
| Streptococcus dysgalactiae subsp. equisimilis strain NCTC11555 | vicK | Two-component sensor kinase SA14-24 | 19088917 | 100 | 100 | 92 | 1e-242 |
| Streptococcus dysgalactiae subsp. equisimilis strain NCTC11556 | vicK | Two-component sensor kinase SA14-24 | 19088917 | 100 | 100 | 93 | 1e-242 |
| Streptococcus dysgalactiae subsp. equisimilis strain NCTC11557 | vicK | Two-component sensor kinase SA14-24 | 19088917 | 100 | 100 | 93 | 1e-242 |
| Streptococcus dysgalactiae subsp. equisimilis strain NCTC11564 | vicK | Two-component sensor kinase SA14-24 | 19088917 | 100 | 100 | 93 | 1e-242 |
| Streptococcus dysgalactiae subsp. equisimilis strain NCTC5370 | vicK | Two-component sensor kinase SA14-24 | 19088917 | 100 | 100 | 93 | 1e-242 |
| Streptococcus dysgalactiae subsp. equisimilis strain NCTC5371 | vicK | Two-component sensor kinase SA14-24 | 19088917 | 100 | 100 | 93 | 1e-242 |
| Streptococcus dysgalactiae subsp. equisimilis strain NCTC6179 | vicK | Two-component sensor kinase SA14-24 | 19088917 | 100 | 100 | 93 | 1e-243 |
| Streptococcus dysgalactiae subsp. equisimilis strain NCTC6181 | vicK | Two-component sensor kinase SA14-24 | 19088917 | 100 | 100 | 93 | 1e-243 |
| Streptococcus dysgalactiae subsp. equisimilis strain NCTC6407 | vicK | Two-component sensor kinase SA14-24 | 19088917 | 100 | 100 | 93 | 1e-243 |
| Streptococcus dysgalactiae subsp. equisimilis strain NCTC7136 | vicK | Two-component sensor kinase SA14-24 | 19088917 | 100 | 100 | 92 | 1e-241 |
| Streptococcus dysgalactiae subsp. equisimilis strain NCTC8543 | vicK | Two-component sensor kinase SA14-24 | 19088917 | 100 | 100 | 93 | 1e-242 |
| Streptococcus dysgalactiae subsp. equisimilis strain NCTC8546 | vicK | Two-component sensor kinase SA14-24 | 19088917 | 100 | 100 | 93 | 1e-242 |
| Streptococcus dysgalactiae subsp. equisimilis strain NCTC9413 | vicK | Two-component sensor kinase SA14-24 | 19088917 | 100 | 100 | 93 | 1e-242 |
| Streptococcus dysgalactiae subsp. equisimilis strain NCTC9414 | vicK | Two-component sensor kinase SA14-24 | 19088917 | 100 | 100 | 93 | 1e-242 |
| Streptococcus dysgalactiae subsp. equisimilis strain NCTC9603 | vicK | Two-component sensor kinase SA14-24 | 19088917 | 100 | 100 | 93 | 1e-242 |
| Streptococcus dysgalactiae subsp. equisimilis strain SS1575 | vicK | Two-component sensor kinase SA14-24 | 19088917 | 100 | 100 | 93 | 1e-242 |
| Streptococcus dysgalactiae subsp. equisimilis strain T642 | vicK | Two-component sensor kinase SA14-24 | 19088917 | 100 | 100 | 92 | 0.0 |
| Streptococcus dysgalactiae subsp. equisimilis strain UT_4031CC | vicK | Two-component sensor kinase SA14-24 | 19088917 | 100 | 100 | 93 | 1e-242 |
| Streptococcus dysgalactiae subsp. equisimilis strain UT_4231_KK | vicK | Two-component sensor kinase SA14-24 | 19088917 | 100 | 100 | 93 | 1e-242 |
| Streptococcus dysgalactiae subsp. equisimilis strain UT_4234_DH | vicK | Two-component sensor kinase SA14-24 | 19088917 | 100 | 100 | 93 | 1e-242 |
| Streptococcus dysgalactiae subsp. equisimilis strain UT_4241_XS | vicK | Two-component sensor kinase SA14-24 | 19088917 | 100 | 100 | 93 | 1e-242 |
| Streptococcus dysgalactiae subsp. equisimilis strain UT_4242_AB | vicK | Two-component sensor kinase SA14-24 | 19088917 | 100 | 100 | 93 | 1e-242 |
| Streptococcus dysgalactiae subsp. equisimilis strain UT_4255RC | vicK | Two-component sensor kinase SA14-24 | 19088917 | 100 | 100 | 93 | 1e-242 |
| Streptococcus dysgalactiae subsp. equisimilis strain UT_4277_BB | vicK | Two-component sensor kinase SA14-24 | 19088917 | 100 | 100 | 93 | 1e-242 |
| Streptococcus dysgalactiae subsp. equisimilis strain UT_4966_RC | vicK | Two-component sensor kinase SA14-24 | 19088917 | 100 | 100 | 93 | 1e-242 |
| Streptococcus dysgalactiae subsp. equisimilis strain UT-5345 | vicK | Two-component sensor kinase SA14-24 | 19088917 | 100 | 100 | 93 | 1e-242 |
| Streptococcus dysgalactiae subsp. equisimilis strain UT-5354 | vicK | Two-component sensor kinase SA14-24 | 19088917 | 100 | 100 | 93 | 1e-242 |
| Streptococcus dysgalactiae subsp. equisimilis strain UT-SS1069 | vicK | Two-component sensor kinase SA14-24 | 19088917 | 100 | 100 | 93 | 1e-242 |
| Streptococcus dysgalactiae subsp. equisimilis strain UT-SS957 | vicK | Two-component sensor kinase SA14-24 | 19088917 | 100 | 100 | 93 | 1e-242 |
| Streptococcus dysgalactiae subsp. equisimilis strain WCHSDSE-1 | vicK | Two-component sensor kinase SA14-24 | 19088917 | 100 | 100 | 93 | 1e-242 |
| Streptococcus dysgalactiae subsp. equisimilis 167 | hasC | UTP--glucose-1-phosphate uridylyltransferase (EC 2.7.7.9) | 12679262 | 97 | 99 | 98 | 1e-165 |
| Streptococcus dysgalactiae subsp. equisimilis AC-2713 | hasC | UTP--glucose-1-phosphate uridylyltransferase (EC 2.7.7.9) | 12679262 | 97 | 99 | 98 | 1e-165 |
| Streptococcus dysgalactiae subsp. equisimilis AKSDE4288 | hasC | UTP--glucose-1-phosphate uridylyltransferase (EC 2.7.7.9) | 12679262 | 97 | 99 | 98 | 1e-165 |
| Streptococcus dysgalactiae subsp. equisimilis ATCC 12394 | hasC | UTP--glucose-1-phosphate uridylyltransferase (EC 2.7.7.9) | 12679262 | 97 | 99 | 98 | 1e-164 |
| Streptococcus dysgalactiae subsp. equisimilis GGS_124 | hasC | UTP--glucose-1-phosphate uridylyltransferase (EC 2.7.7.9) | 12679262 | 97 | 99 | 98 | 1e-164 |
| Streptococcus dysgalactiae subsp. equisimilis RE378 | hasC | UTP--glucose-1-phosphate uridylyltransferase (EC 2.7.7.9) | 12679262 | 97 | 99 | 98 | 1e-165 |
| Streptococcus dysgalactiae subsp. equisimilis SD SCDR1 | hasC | UTP--glucose-1-phosphate uridylyltransferase (EC 2.7.7.9) | 12679262 | 97 | 99 | 98 | 1e-165 |
| Streptococcus dysgalactiae subsp. equisimilis SK1249 | galU | UTP--glucose-1-phosphate uridylyltransferase (EC 2.7.7.9) | 15213167 | 97 | 99 | 98 | 1e-164 |
| Streptococcus dysgalactiae subsp. equisimilis SK1250 | galU | UTP--glucose-1-phosphate uridylyltransferase (EC 2.7.7.9) | 15213167 | 97 | 99 | 98 | 1e-165 |
| Streptococcus dysgalactiae subsp. equisimilis strain ASDSE_96 | hasC | UTP--glucose-1-phosphate uridylyltransferase (EC 2.7.7.9) | 12679262 | 97 | 99 | 98 | 1e-165 |
| Streptococcus dysgalactiae subsp. equisimilis strain ASDSE_99 | hasC | UTP--glucose-1-phosphate uridylyltransferase (EC 2.7.7.9) | 12679262 | 97 | 99 | 98 | 1e-165 |
| Streptococcus dysgalactiae subsp. equisimilis strain C161L1 | hasC | UTP--glucose-1-phosphate uridylyltransferase (EC 2.7.7.9) | 12679262 | 98 | 99 | 98 | 0 |
| Streptococcus dysgalactiae subsp. equisimilis strain KNZ01 | hasC | UTP--glucose-1-phosphate uridylyltransferase (EC 2.7.7.9) | 12679262 | 97 | 99 | 98 | 1e-165 |
| Streptococcus dysgalactiae subsp. equisimilis strain KNZ01 | hasC | UTP--glucose-1-phosphate uridylyltransferase (EC 2.7.7.9) | 12679262 | 97 | 99 | 98 | 1e-165 |
| Streptococcus dysgalactiae subsp. equisimilis strain KNZ03 | hasC | UTP--glucose-1-phosphate uridylyltransferase (EC 2.7.7.9) | 12679262 | 97 | 99 | 98 | 1e-164 |
| Streptococcus dysgalactiae subsp. equisimilis strain KNZ03 | hasC | UTP--glucose-1-phosphate uridylyltransferase (EC 2.7.7.9) | 12679262 | 97 | 99 | 98 | 1e-164 |
| Streptococcus dysgalactiae subsp. equisimilis strain KNZ04 | hasC | UTP--glucose-1-phosphate uridylyltransferase (EC 2.7.7.9) | 12679262 | 97 | 99 | 98 | 1e-164 |
| Streptococcus dysgalactiae subsp. equisimilis strain KNZ06 | hasC | UTP--glucose-1-phosphate uridylyltransferase (EC 2.7.7.9) | 12679262 | 97 | 99 | 98 | 1e-165 |
| Streptococcus dysgalactiae subsp. equisimilis strain KNZ07 | hasC | UTP--glucose-1-phosphate uridylyltransferase (EC 2.7.7.9) | 12679262 | 97 | 99 | 98 | 1e-165 |
| Streptococcus dysgalactiae subsp. equisimilis strain KNZ10 | hasC | UTP--glucose-1-phosphate uridylyltransferase (EC 2.7.7.9) | 12679262 | 97 | 99 | 98 | 1e-165 |
| Streptococcus dysgalactiae subsp. equisimilis strain KNZ12 | hasC | UTP--glucose-1-phosphate uridylyltransferase (EC 2.7.7.9) | 12679262 | 97 | 99 | 98 | 1e-165 |
| Streptococcus dysgalactiae subsp. equisimilis strain KNZ15 | hasC | UTP--glucose-1-phosphate uridylyltransferase (EC 2.7.7.9) | 12679262 | 97 | 99 | 98 | 1e-165 |
| Streptococcus dysgalactiae subsp. equisimilis strain KNZ16 | hasC | UTP--glucose-1-phosphate uridylyltransferase (EC 2.7.7.9) | 12679262 | 97 | 99 | 98 | 1e-164 |
| Streptococcus dysgalactiae subsp. equisimilis strain NCTC10321 | hasC | UTP--glucose-1-phosphate uridylyltransferase (EC 2.7.7.9) | 12679262 | 97 | 99 | 98 | 1e-165 |
| Streptococcus dysgalactiae subsp. equisimilis strain NCTC11554 | hasC | UTP--glucose-1-phosphate uridylyltransferase (EC 2.7.7.9) | 12679262 | 97 | 99 | 98 | 1e-164 |
| Streptococcus dysgalactiae subsp. equisimilis strain NCTC11555 | hasC | UTP--glucose-1-phosphate uridylyltransferase (EC 2.7.7.9) | 12679262 | 97 | 99 | 98 | 1e-165 |
| Streptococcus dysgalactiae subsp. equisimilis strain NCTC11556 | hasC | UTP--glucose-1-phosphate uridylyltransferase (EC 2.7.7.9) | 12679262 | 97 | 99 | 98 | 1e-165 |
| Streptococcus dysgalactiae subsp. equisimilis strain NCTC11557 | hasC | UTP--glucose-1-phosphate uridylyltransferase (EC 2.7.7.9) | 12679262 | 97 | 99 | 98 | 1e-165 |
| Streptococcus dysgalactiae subsp. equisimilis strain NCTC11564 | hasC | UTP--glucose-1-phosphate uridylyltransferase (EC 2.7.7.9) | 12679262 | 97 | 99 | 98 | 1e-165 |
| Streptococcus dysgalactiae subsp. equisimilis strain NCTC11565 | galU | UTP--glucose-1-phosphate uridylyltransferase (EC 2.7.7.9) | 15213167 | 100 | 100 | 100 | 1e-161 |
| Streptococcus dysgalactiae subsp. equisimilis strain NCTC5370 | hasC | UTP--glucose-1-phosphate uridylyltransferase (EC 2.7.7.9) | 12679262 | 97 | 99 | 98 | 1e-165 |
| Streptococcus dysgalactiae subsp. equisimilis strain NCTC5371 | hasC | UTP--glucose-1-phosphate uridylyltransferase (EC 2.7.7.9) | 12679262 | 97 | 99 | 98 | 1e-165 |
| Streptococcus dysgalactiae subsp. equisimilis strain NCTC6179 | hasC | UTP--glucose-1-phosphate uridylyltransferase (EC 2.7.7.9) | 12679262 | 97 | 98 | 98 | 1e-164 |
| Streptococcus dysgalactiae subsp. equisimilis strain NCTC6181 | hasC | UTP--glucose-1-phosphate uridylyltransferase (EC 2.7.7.9) | 12679262 | 97 | 98 | 98 | 1e-164 |
| Streptococcus dysgalactiae subsp. equisimilis strain NCTC6407 | hasC | UTP--glucose-1-phosphate uridylyltransferase (EC 2.7.7.9) | 12679262 | 97 | 99 | 98 | 1e-165 |
| Streptococcus dysgalactiae subsp. equisimilis strain NCTC7136 | hasC | UTP--glucose-1-phosphate uridylyltransferase (EC 2.7.7.9) | 12679262 | 97 | 99 | 98 | 1e-165 |
| Streptococcus dysgalactiae subsp. equisimilis strain NCTC8543 | hasC | UTP--glucose-1-phosphate uridylyltransferase (EC 2.7.7.9) | 12679262 | 97 | 99 | 98 | 1e-164 |
| Streptococcus dysgalactiae subsp. equisimilis strain NCTC8546 | hasC | UTP--glucose-1-phosphate uridylyltransferase (EC 2.7.7.9) | 12679262 | 97 | 99 | 98 | 1e-165 |
| Streptococcus dysgalactiae subsp. equisimilis strain NCTC9413 | hasC | UTP--glucose-1-phosphate uridylyltransferase (EC 2.7.7.9) | 12679262 | 97 | 99 | 98 | 1e-165 |
| Streptococcus dysgalactiae subsp. equisimilis strain NCTC9414 | hasC | UTP--glucose-1-phosphate uridylyltransferase (EC 2.7.7.9) | 12679262 | 97 | 99 | 98 | 1e-165 |
| Streptococcus dysgalactiae subsp. equisimilis strain NCTC9603 | hasC | UTP--glucose-1-phosphate uridylyltransferase (EC 2.7.7.9) | 12679262 | 97 | 99 | 98 | 1e-165 |
| Streptococcus dysgalactiae subsp. equisimilis strain SS1575 | hasC | UTP--glucose-1-phosphate uridylyltransferase (EC 2.7.7.9) | 12679262 | 97 | 99 | 98 | 1e-165 |
| Streptococcus dysgalactiae subsp. equisimilis strain T642 | hasC | UTP--glucose-1-phosphate uridylyltransferase (EC 2.7.7.9) | 12679262 | 98 | 99 | 98 | 0 |
| Streptococcus dysgalactiae subsp. equisimilis strain UT_4031CC | hasC | UTP--glucose-1-phosphate uridylyltransferase (EC 2.7.7.9) | 12679262 | 97 | 99 | 98 | 1e-165 |
| Streptococcus dysgalactiae subsp. equisimilis strain UT_4231_KK | hasC | UTP--glucose-1-phosphate uridylyltransferase (EC 2.7.7.9) | 12679262 | 97 | 99 | 98 | 1e-165 |
| Streptococcus dysgalactiae subsp. equisimilis strain UT_4234_DH | hasC | UTP--glucose-1-phosphate uridylyltransferase (EC 2.7.7.9) | 12679262 | 97 | 99 | 98 | 1e-165 |
| Streptococcus dysgalactiae subsp. equisimilis strain UT_4241_XS | hasC | UTP--glucose-1-phosphate uridylyltransferase (EC 2.7.7.9) | 12679262 | 97 | 99 | 98 | 1e-165 |
| Streptococcus dysgalactiae subsp. equisimilis strain UT_4242_AB | hasC | UTP--glucose-1-phosphate uridylyltransferase (EC 2.7.7.9) | 12679262 | 97 | 99 | 98 | 1e-165 |
| Streptococcus dysgalactiae subsp. equisimilis strain UT_4255RC | hasC | UTP--glucose-1-phosphate uridylyltransferase (EC 2.7.7.9) | 12679262 | 97 | 99 | 98 | 1e-165 |
| Streptococcus dysgalactiae subsp. equisimilis strain UT_4277_BB | hasC | UTP--glucose-1-phosphate uridylyltransferase (EC 2.7.7.9) | 12679262 | 97 | 99 | 98 | 1e-165 |
| Streptococcus dysgalactiae subsp. equisimilis strain UT_4966_RC | hasC | UTP--glucose-1-phosphate uridylyltransferase (EC 2.7.7.9) | 12679262 | 97 | 99 | 98 | 1e-165 |
| Streptococcus dysgalactiae subsp. equisimilis strain UT-5345 | hasC | UTP--glucose-1-phosphate uridylyltransferase (EC 2.7.7.9) | 12679262 | 97 | 99 | 98 | 1e-165 |
| Streptococcus dysgalactiae subsp. equisimilis strain UT-5354 | hasC | UTP--glucose-1-phosphate uridylyltransferase (EC 2.7.7.9) | 12679262 | 97 | 99 | 98 | 1e-165 |
| Streptococcus dysgalactiae subsp. equisimilis strain UT-SS1069 | hasC | UTP--glucose-1-phosphate uridylyltransferase (EC 2.7.7.9) | 12679262 | 97 | 99 | 98 | 1e-165 |
| Streptococcus dysgalactiae subsp. equisimilis strain UT-SS957 | hasC | UTP--glucose-1-phosphate uridylyltransferase (EC 2.7.7.9) | 12679262 | 97 | 99 | 98 | 1e-165 |
| Streptococcus dysgalactiae subsp. equisimilis strain WCHSDSE-1 | hasC | UTP--glucose-1-phosphate uridylyltransferase (EC 2.7.7.9) | 12679262 | 97 | 99 | 98 | 1e-165 |
| Streptococcus dysgalactiae subsp. equisimilis 167 | csoS1A | Ethanolamine utilization polyhedral-body-like protein EutM | 17307944 | 90 | 95 | 85 | 8e-36 |
| Streptococcus dysgalactiae subsp. equisimilis ATCC 12394 | csoS1A | Ethanolamine utilization polyhedral-body-like protein EutM | 17307944 | 88 | 92 | 85 | 4e-35 |
| Streptococcus dysgalactiae subsp. equisimilis RE378 | csoS1A | Ethanolamine utilization polyhedral-body-like protein EutM | 17307944 | 88 | 92 | 85 | 4e-35 |
| Streptococcus dysgalactiae subsp. equisimilis SK1249 | csoS1A | Ethanolamine utilization polyhedral-body-like protein EutM | 17307944 | 88 | 92 | 85 | 4e-35 |
| Streptococcus dysgalactiae subsp. equisimilis strain ASDSE_96 | csoS1A | Ethanolamine utilization polyhedral-body-like protein EutM | 17307944 | 88 | 92 | 85 | 4e-35 |
| Streptococcus dysgalactiae subsp. equisimilis strain C161L1 | csoS1A | Ethanolamine utilization polyhedral-body-like protein EutM | 17307944 | 92 | 97 | 84 | 5.999939634699569e-41 |
| Streptococcus dysgalactiae subsp. equisimilis strain KNZ01 | EF1623 | Ethanolamine utilization polyhedral-body-like protein EutM | 17307944 | 83 | 95 | 86 | 2e-32 |
| Streptococcus dysgalactiae subsp. equisimilis strain KNZ04 | EF1623 | Ethanolamine utilization polyhedral-body-like protein EutM | 17307944 | 88 | 92 | 85 | 4e-35 |
| Streptococcus dysgalactiae subsp. equisimilis strain KNZ07 | EF1623 | Ethanolamine utilization polyhedral-body-like protein EutM | 17307944 | 88 | 92 | 85 | 4e-35 |
| Streptococcus dysgalactiae subsp. equisimilis strain KNZ10 | EF1623 | Ethanolamine utilization polyhedral-body-like protein EutM | 17307944 | 90 | 95 | 85 | 8e-36 |
| Streptococcus dysgalactiae subsp. equisimilis strain KNZ12 | EF1623 | Ethanolamine utilization polyhedral-body-like protein EutM | 17307944 | 88 | 92 | 85 | 4e-35 |
| Streptococcus dysgalactiae subsp. equisimilis strain KNZ15 | EF1623 | Ethanolamine utilization polyhedral-body-like protein EutM | 17307944 | 88 | 96 | 85 | 4e-35 |
| Streptococcus dysgalactiae subsp. equisimilis strain KNZ16 | EF1623 | Ethanolamine utilization polyhedral-body-like protein EutM | 17307944 | 88 | 92 | 85 | 4e-35 |
| Streptococcus dysgalactiae subsp. equisimilis strain NCTC11554 | EF1623 | Ethanolamine utilization polyhedral-body-like protein EutM | 17307944 | 88 | 92 | 85 | 4e-35 |
| Streptococcus dysgalactiae subsp. equisimilis strain NCTC11564 | EF1623 | Ethanolamine utilization polyhedral-body-like protein EutM | 17307944 | 90 | 95 | 86 | 4e-36 |
| Streptococcus dysgalactiae subsp. equisimilis strain NCTC5370 | EF1623 | Ethanolamine utilization polyhedral-body-like protein EutM | 17307944 | 88 | 92 | 85 | 4e-35 |
| Streptococcus dysgalactiae subsp. equisimilis strain NCTC6179 | EF1623 | Ethanolamine utilization polyhedral-body-like protein EutM | 17307944 | 88 | 92 | 85 | 4e-35 |
| Streptococcus dysgalactiae subsp. equisimilis strain NCTC6181 | EF1623 | Ethanolamine utilization polyhedral-body-like protein EutM | 17307944 | 88 | 92 | 85 | 4e-35 |
| Streptococcus dysgalactiae subsp. equisimilis strain NCTC8546 | EF1623 | Ethanolamine utilization polyhedral-body-like protein EutM | 17307944 | 88 | 92 | 85 | 4e-35 |
| Streptococcus dysgalactiae subsp. equisimilis strain NCTC9413 | EF1623 | Ethanolamine utilization polyhedral-body-like protein EutM | 17307944 | 90 | 95 | 86 | 4e-36 |
| Streptococcus dysgalactiae subsp. equisimilis strain SS1575 | EF1623 | Ethanolamine utilization polyhedral-body-like protein EutM | 17307944 | 78 | 100 | 88 | 3e-31 |
| Streptococcus dysgalactiae subsp. equisimilis strain UT_4031CC | csoS1A | Ethanolamine utilization polyhedral-body-like protein EutM | 17307944 | 88 | 92 | 85 | 4e-35 |
| Streptococcus dysgalactiae subsp. equisimilis strain UT_4231_KK | csoS1A | Ethanolamine utilization polyhedral-body-like protein EutM | 17307944 | 90 | 95 | 86 | 4e-36 |
| Streptococcus dysgalactiae subsp. equisimilis strain UT_4241_XS | csoS1A | Ethanolamine utilization polyhedral-body-like protein EutM | 17307944 | 90 | 95 | 85 | 8e-36 |
| Streptococcus dysgalactiae subsp. equisimilis strain UT_4966_RC | csoS1A | Ethanolamine utilization polyhedral-body-like protein EutM | 17307944 | 90 | 95 | 86 | 4e-36 |
| Streptococcus dysgalactiae subsp. equisimilis strain UT-5345 | csoS1A | Ethanolamine utilization polyhedral-body-like protein EutM | 17307944 | 88 | 92 | 85 | 4e-35 |
| Streptococcus dysgalactiae subsp. equisimilis strain UT-SS1069 | csoS1A | Ethanolamine utilization polyhedral-body-like protein EutM | 17307944 | 87 | 91 | 85 | 1e-34 |
| Streptococcus dysgalactiae subsp. equisimilis 167 | SPy_1718 | Esterase (EC 3.1.1.-) | 19805529 | 100 | 100 | 93 | 1e-187 |
| Streptococcus dysgalactiae subsp. equisimilis AC-2713 | BCHE | Esterase (EC 3.1.1.-) | 19805529 | 100 | 100 | 94 | 1e-187 |
| Streptococcus dysgalactiae subsp. equisimilis AKSDE4288 | SPy_1718 | Esterase (EC 3.1.1.-) | 19805529 | 100 | 100 | 94 | 1e-187 |
| Streptococcus dysgalactiae subsp. equisimilis ATCC 12394 | SPy_1718 | Esterase (EC 3.1.1.-) | 19805529 | 100 | 100 | 94 | 1e-187 |
| Streptococcus dysgalactiae subsp. equisimilis GGS_124 | SPy_1718 | Esterase (EC 3.1.1.-) | 19805529 | 100 | 100 | 94 | 1e-187 |
| Streptococcus dysgalactiae subsp. equisimilis RE378 | SPy_1718 | Esterase (EC 3.1.1.-) | 19805529 | 100 | 100 | 94 | 1e-187 |
| Streptococcus dysgalactiae subsp. equisimilis SD SCDR1 | SPy_1718 | Esterase (EC 3.1.1.-) | 19805529 | 100 | 100 | 94 | 1e-187 |
| Streptococcus dysgalactiae subsp. equisimilis SK1249 | SPy_1718 | Esterase (EC 3.1.1.-) | 19805529 | 100 | 100 | 94 | 1e-187 |
| Streptococcus dysgalactiae subsp. equisimilis SK1250 | SPy_1718 | Esterase (EC 3.1.1.-) | 19805529 | 100 | 100 | 93 | 1e-187 |
| Streptococcus dysgalactiae subsp. equisimilis strain ASDSE_96 | SPy_1718 | Esterase (EC 3.1.1.-) | 19805529 | 100 | 100 | 93 | 1e-187 |
| Streptococcus dysgalactiae subsp. equisimilis strain ASDSE_99 | SPy_1718 | Esterase (EC 3.1.1.-) | 19805529 | 100 | 100 | 93 | 1e-187 |
| Streptococcus dysgalactiae subsp. equisimilis strain C161L1 | SPy_1718 | Esterase (EC 3.1.1.-) | 19805529 | 100 | 100 | 94 | 0 |
| Streptococcus dysgalactiae subsp. equisimilis strain KNZ01 | SPy_1718 | Esterase (EC 3.1.1.-) | 19805529 | 100 | 100 | 94 | 1e-187 |
| Streptococcus dysgalactiae subsp. equisimilis strain KNZ03 | SPy_1718 | Esterase (EC 3.1.1.-) | 19805529 | 100 | 100 | 94 | 1e-187 |
| Streptococcus dysgalactiae subsp. equisimilis strain KNZ04 | SPy_1718 | Esterase (EC 3.1.1.-) | 19805529 | 100 | 100 | 94 | 1e-187 |
| Streptococcus dysgalactiae subsp. equisimilis strain KNZ06 | SPy_1718 | Esterase (EC 3.1.1.-) | 19805529 | 100 | 100 | 93 | 1e-187 |
| Streptococcus dysgalactiae subsp. equisimilis strain KNZ07 | SPy_1718 | Esterase (EC 3.1.1.-) | 19805529 | 100 | 100 | 94 | 1e-187 |
| Streptococcus dysgalactiae subsp. equisimilis strain KNZ10 | SPy_1718 | Esterase (EC 3.1.1.-) | 19805529 | 100 | 100 | 94 | 1e-187 |
| Streptococcus dysgalactiae subsp. equisimilis strain KNZ12 | SPy_1718 | Esterase (EC 3.1.1.-) | 19805529 | 100 | 100 | 94 | 1e-187 |
| Streptococcus dysgalactiae subsp. equisimilis strain KNZ15 | SPy_1718 | Esterase (EC 3.1.1.-) | 19805529 | 100 | 100 | 94 | 1e-187 |
| Streptococcus dysgalactiae subsp. equisimilis strain KNZ16 | SPy_1718 | Esterase (EC 3.1.1.-) | 19805529 | 100 | 100 | 94 | 1e-187 |
| Streptococcus dysgalactiae subsp. equisimilis strain NCTC10321 | SPy_1718 | Esterase (EC 3.1.1.-) | 19805529 | 100 | 100 | 85 | 1e-170 |
| Streptococcus dysgalactiae subsp. equisimilis strain NCTC11554 | SPy_1718 | Esterase (EC 3.1.1.-) | 19805529 | 100 | 100 | 94 | 1e-187 |
| Streptococcus dysgalactiae subsp. equisimilis strain NCTC11555 | SPy_1718 | Esterase (EC 3.1.1.-) | 19805529 | 100 | 100 | 93 | 1e-187 |
| Streptococcus dysgalactiae subsp. equisimilis strain NCTC11556 | SPy_1718 | Esterase (EC 3.1.1.-) | 19805529 | 100 | 100 | 94 | 1e-187 |
| Streptococcus dysgalactiae subsp. equisimilis strain NCTC11557 | SPy_1718 | Esterase (EC 3.1.1.-) | 19805529 | 100 | 100 | 94 | 1e-187 |
| Streptococcus dysgalactiae subsp. equisimilis strain NCTC11564 | SPy_1718 | Esterase (EC 3.1.1.-) | 19805529 | 100 | 100 | 94 | 1e-187 |
| Streptococcus dysgalactiae subsp. equisimilis strain NCTC5370 | SPy_1718 | Esterase (EC 3.1.1.-) | 19805529 | 100 | 100 | 93 | 1e-187 |
| Streptococcus dysgalactiae subsp. equisimilis strain NCTC5371 | SPy_1718 | Esterase (EC 3.1.1.-) | 19805529 | 100 | 100 | 94 | 1e-187 |
| Streptococcus dysgalactiae subsp. equisimilis strain NCTC6179 | SPy_1718 | Esterase (EC 3.1.1.-) | 19805529 | 100 | 100 | 85 | 1e-172 |
| Streptococcus dysgalactiae subsp. equisimilis strain NCTC6181 | SPy_1718 | Esterase (EC 3.1.1.-) | 19805529 | 100 | 100 | 85 | 1e-172 |
| Streptococcus dysgalactiae subsp. equisimilis strain NCTC6407 | SPy_1718 | Esterase (EC 3.1.1.-) | 19805529 | 100 | 100 | 85 | 1e-170 |
| Streptococcus dysgalactiae subsp. equisimilis strain NCTC7136 | SPy_1718 | Esterase (EC 3.1.1.-) | 19805529 | 100 | 100 | 93 | 1e-187 |
| Streptococcus dysgalactiae subsp. equisimilis strain NCTC8543 | SPy_1718 | Esterase (EC 3.1.1.-) | 19805529 | 100 | 100 | 94 | 1e-187 |
| Streptococcus dysgalactiae subsp. equisimilis strain NCTC8546 | SPy_1718 | Esterase (EC 3.1.1.-) | 19805529 | 100 | 100 | 94 | 1e-187 |
| Streptococcus dysgalactiae subsp. equisimilis strain NCTC9413 | SPy_1718 | Esterase (EC 3.1.1.-) | 19805529 | 100 | 100 | 94 | 1e-187 |
| Streptococcus dysgalactiae subsp. equisimilis strain NCTC9414 | SPy_1718 | Esterase (EC 3.1.1.-) | 19805529 | 100 | 100 | 94 | 1e-187 |
| Streptococcus dysgalactiae subsp. equisimilis strain NCTC9603 | SPy_1718 | Esterase (EC 3.1.1.-) | 19805529 | 100 | 100 | 93 | 1e-187 |
| Streptococcus dysgalactiae subsp. equisimilis strain SS1575 | SPy_1718 | Esterase (EC 3.1.1.-) | 19805529 | 100 | 100 | 93 | 1e-187 |
| Streptococcus dysgalactiae subsp. equisimilis strain T642 | SPy_1718 | Esterase (EC 3.1.1.-) | 19805529 | 100 | 100 | 93 | 0 |
| Streptococcus dysgalactiae subsp. equisimilis strain UT_4031CC | SPy_1718 | Esterase (EC 3.1.1.-) | 19805529 | 100 | 100 | 93 | 1e-187 |
| Streptococcus dysgalactiae subsp. equisimilis strain UT_4231_KK | SPy_1718 | Esterase (EC 3.1.1.-) | 19805529 | 100 | 100 | 94 | 1e-187 |
| Streptococcus dysgalactiae subsp. equisimilis strain UT_4234_DH | SPy_1718 | Esterase (EC 3.1.1.-) | 19805529 | 100 | 100 | 93 | 1e-187 |
| Streptococcus dysgalactiae subsp. equisimilis strain UT_4241_XS | SPy_1718 | Esterase (EC 3.1.1.-) | 19805529 | 100 | 100 | 93 | 1e-187 |
| Streptococcus dysgalactiae subsp. equisimilis strain UT_4242_AB | SPy_1718 | Esterase (EC 3.1.1.-) | 19805529 | 100 | 100 | 94 | 1e-187 |
| Streptococcus dysgalactiae subsp. equisimilis strain UT_4255RC | SPy_1718 | Esterase (EC 3.1.1.-) | 19805529 | 100 | 100 | 93 | 1e-187 |
| Streptococcus dysgalactiae subsp. equisimilis strain UT_4277_BB | SPy_1718 | Esterase (EC 3.1.1.-) | 19805529 | 100 | 100 | 93 | 1e-187 |
| Streptococcus dysgalactiae subsp. equisimilis strain UT_4966_RC | SPy_1718 | Esterase (EC 3.1.1.-) | 19805529 | 100 | 100 | 93 | 1e-187 |
| Streptococcus dysgalactiae subsp. equisimilis strain UT-5345 | SPy_1718 | Esterase (EC 3.1.1.-) | 19805529 | 100 | 100 | 93 | 1e-187 |
| Streptococcus dysgalactiae subsp. equisimilis strain UT-5354 | SPy_1718 | Esterase (EC 3.1.1.-) | 19805529 | 100 | 100 | 93 | 1e-187 |
| Streptococcus dysgalactiae subsp. equisimilis strain UT-SS1069 | SPy_1718 | Esterase (EC 3.1.1.-) | 19805529 | 100 | 100 | 93 | 1e-187 |
| Streptococcus dysgalactiae subsp. equisimilis strain UT-SS957 | SPy_1718 | Esterase (EC 3.1.1.-) | 19805529 | 100 | 100 | 93 | 1e-187 |
| Streptococcus dysgalactiae subsp. equisimilis strain WCHSDSE-1 | SPy_1718 | Esterase (EC 3.1.1.-) | 19805529 | 100 | 100 | 94 | 1e-187 |
| Streptococcus dysgalactiae subsp. equisimilis strain ASDSE_99 | SP_0939 | DUF1706 domain-containing protein | 12207705 | 98 | 94 | 84 | 4e-83 |
| Streptococcus dysgalactiae subsp. equisimilis strain KNZ01 | SP_0939 | DUF1706 domain-containing protein | 12207705 | 96 | 96 | 84 | 2e-80 |
| Streptococcus dysgalactiae subsp. equisimilis strain KNZ06 | SP_0939 | DUF1706 domain-containing protein | 12207705 | 96 | 96 | 84 | 2e-80 |
| Streptococcus dysgalactiae subsp. equisimilis strain KNZ12 | SP_0939 | DUF1706 domain-containing protein | 12207705 | 96 | 96 | 84 | 2e-80 |
| Streptococcus dysgalactiae subsp. equisimilis strain NCTC11555 | SP_0939 | DUF1706 domain-containing protein | 12207705 | 96 | 96 | 84 | 2e-80 |
| Streptococcus dysgalactiae subsp. equisimilis strain NCTC11556 | SP_0939 | DUF1706 domain-containing protein | 12207705 | 96 | 96 | 84 | 2e-80 |
| Streptococcus dysgalactiae subsp. equisimilis strain NCTC11557 | SP_0939 | DUF1706 domain-containing protein | 12207705 | 100 | 100 | 86 | 5e-86 |
| Streptococcus dysgalactiae subsp. equisimilis strain NCTC11564 | SP_0939 | DUF1706 domain-containing protein | 12207705 | 100 | 100 | 83 | 5e-83 |
| Streptococcus dysgalactiae subsp. equisimilis strain NCTC9413 | SP_0939 | DUF1706 domain-containing protein | 12207705 | 100 | 100 | 94 | 1e-93 |
| Streptococcus dysgalactiae subsp. equisimilis strain T642 | SP_0939 | DUF1706 domain-containing protein | 12207705 | 100 | 100 | 95 | 3e-119 |
| Streptococcus dysgalactiae subsp. equisimilis strain UT_4234_DH | SP_0939 | DUF1706 domain-containing protein | 12207705 | 96 | 96 | 84 | 2e-80 |
| Streptococcus dysgalactiae subsp. equisimilis strain UT_4242_AB | SP_0939 | DUF1706 domain-containing protein | 12207705 | 96 | 96 | 84 | 2e-80 |
| Streptococcus dysgalactiae subsp. equisimilis strain UT_4277_BB | SP_0939 | DUF1706 domain-containing protein | 12207705 | 100 | 100 | 94 | 3e-93 |
| Streptococcus dysgalactiae subsp. equisimilis strain UT-5345 | SP_0939 | DUF1706 domain-containing protein | 12207705 | 96 | 96 | 84 | 2e-80 |
| Streptococcus dysgalactiae subsp. equisimilis strain UT-5354 | SP_0939 | DUF1706 domain-containing protein | 12207705 | 100 | 100 | 83 | 1e-82 |
| Streptococcus dysgalactiae subsp. equisimilis AC-2713 | rpoE | DNA-directed RNA polymerase delta subunit (EC 2.7.7.6) | 16513739 | 87 | 86 | 87 | 5e-83 |
| Streptococcus dysgalactiae subsp. equisimilis AKSDE4288 | rpoE | DNA-directed RNA polymerase delta subunit (EC 2.7.7.6) | 16513739 | 87 | 86 | 87 | 5e-83 |
| Streptococcus dysgalactiae subsp. equisimilis ATCC 12394 | rpoE | DNA-directed RNA polymerase delta subunit (EC 2.7.7.6) | 16513739 | 87 | 86 | 87 | 5e-83 |
| Streptococcus dysgalactiae subsp. equisimilis GGS_124 | rpoE | DNA-directed RNA polymerase delta subunit (EC 2.7.7.6) | 16513739 | 87 | 86 | 87 | 5e-83 |
| Streptococcus dysgalactiae subsp. equisimilis RE378 | rpoE | DNA-directed RNA polymerase delta subunit (EC 2.7.7.6) | 16513739 | 87 | 86 | 87 | 5e-83 |
| Streptococcus dysgalactiae subsp. equisimilis SD SCDR1 | rpoE | DNA-directed RNA polymerase delta subunit (EC 2.7.7.6) | 16513739 | 87 | 86 | 87 | 5e-83 |
| Streptococcus dysgalactiae subsp. equisimilis SK1249 | rpoE | DNA-directed RNA polymerase delta subunit (EC 2.7.7.6) | 16513739 | 87 | 86 | 87 | 5e-83 |
| Streptococcus dysgalactiae subsp. equisimilis SK1250 | rpoE | DNA-directed RNA polymerase delta subunit (EC 2.7.7.6) | 16513739 | 87 | 86 | 87 | 5e-83 |
| Streptococcus dysgalactiae subsp. equisimilis strain ASDSE_96 | rpoE | DNA-directed RNA polymerase delta subunit (EC 2.7.7.6) | 16513739 | 87 | 86 | 87 | 5e-83 |
| Streptococcus dysgalactiae subsp. equisimilis strain ASDSE_99 | rpoE | DNA-directed RNA polymerase delta subunit (EC 2.7.7.6) | 16513739 | 87 | 86 | 87 | 3e-83 |
| Streptococcus dysgalactiae subsp. equisimilis strain C161L1 | rpoE | DNA-directed RNA polymerase delta subunit (EC 2.7.7.6) | 16513739 | 89 | 89 | 85 | 0 |
| Streptococcus dysgalactiae subsp. equisimilis strain KNZ01 | rpoE | DNA-directed RNA polymerase delta subunit (EC 2.7.7.6) | 16513739 | 87 | 86 | 87 | 5e-83 |
| Streptococcus dysgalactiae subsp. equisimilis strain KNZ03 | rpoE | DNA-directed RNA polymerase delta subunit (EC 2.7.7.6) | 16513739 | 87 | 86 | 87 | 5e-83 |
| Streptococcus dysgalactiae subsp. equisimilis strain KNZ04 | rpoE | DNA-directed RNA polymerase delta subunit (EC 2.7.7.6) | 16513739 | 87 | 86 | 87 | 5e-83 |
| Streptococcus dysgalactiae subsp. equisimilis strain KNZ06 | rpoE | DNA-directed RNA polymerase delta subunit (EC 2.7.7.6) | 16513739 | 87 | 86 | 87 | 5e-83 |
| Streptococcus dysgalactiae subsp. equisimilis strain KNZ07 | rpoE | DNA-directed RNA polymerase delta subunit (EC 2.7.7.6) | 16513739 | 87 | 86 | 87 | 5e-83 |
| Streptococcus dysgalactiae subsp. equisimilis strain KNZ10 | rpoE | DNA-directed RNA polymerase delta subunit (EC 2.7.7.6) | 16513739 | 87 | 86 | 87 | 5e-83 |
| Streptococcus dysgalactiae subsp. equisimilis strain KNZ12 | rpoE | DNA-directed RNA polymerase delta subunit (EC 2.7.7.6) | 16513739 | 87 | 86 | 87 | 5e-83 |
| Streptococcus dysgalactiae subsp. equisimilis strain KNZ15 | rpoE | DNA-directed RNA polymerase delta subunit (EC 2.7.7.6) | 16513739 | 87 | 86 | 87 | 5e-83 |
| Streptococcus dysgalactiae subsp. equisimilis strain KNZ16 | rpoE | DNA-directed RNA polymerase delta subunit (EC 2.7.7.6) | 16513739 | 87 | 86 | 87 | 5e-83 |
| Streptococcus dysgalactiae subsp. equisimilis strain NCTC10321 | rpoE | DNA-directed RNA polymerase delta subunit (EC 2.7.7.6) | 16513739 | 87 | 86 | 87 | 5e-83 |
| Streptococcus dysgalactiae subsp. equisimilis strain NCTC11554 | rpoE | DNA-directed RNA polymerase delta subunit (EC 2.7.7.6) | 16513739 | 87 | 86 | 87 | 5e-83 |
| Streptococcus dysgalactiae subsp. equisimilis strain NCTC11555 | rpoE | DNA-directed RNA polymerase delta subunit (EC 2.7.7.6) | 16513739 | 87 | 86 | 87 | 3e-83 |
| Streptococcus dysgalactiae subsp. equisimilis strain NCTC11556 | rpoE | DNA-directed RNA polymerase delta subunit (EC 2.7.7.6) | 16513739 | 87 | 86 | 87 | 5e-83 |
| Streptococcus dysgalactiae subsp. equisimilis strain NCTC11557 | rpoE | DNA-directed RNA polymerase delta subunit (EC 2.7.7.6) | 16513739 | 87 | 86 | 87 | 3e-83 |
| Streptococcus dysgalactiae subsp. equisimilis strain NCTC11564 | rpoE | DNA-directed RNA polymerase delta subunit (EC 2.7.7.6) | 16513739 | 87 | 86 | 87 | 5e-83 |
| Streptococcus dysgalactiae subsp. equisimilis strain NCTC5370 | rpoE | DNA-directed RNA polymerase delta subunit (EC 2.7.7.6) | 16513739 | 87 | 86 | 87 | 5e-83 |
| Streptococcus dysgalactiae subsp. equisimilis strain NCTC5371 | rpoE | DNA-directed RNA polymerase delta subunit (EC 2.7.7.6) | 16513739 | 87 | 86 | 87 | 5e-83 |
| Streptococcus dysgalactiae subsp. equisimilis strain NCTC6179 | rpoE | DNA-directed RNA polymerase delta subunit (EC 2.7.7.6) | 16513739 | 87 | 88 | 87 | 5e-83 |
| Streptococcus dysgalactiae subsp. equisimilis strain NCTC6181 | rpoE | DNA-directed RNA polymerase delta subunit (EC 2.7.7.6) | 16513739 | 87 | 88 | 87 | 5e-83 |
| Streptococcus dysgalactiae subsp. equisimilis strain NCTC6407 | rpoE | DNA-directed RNA polymerase delta subunit (EC 2.7.7.6) | 16513739 | 87 | 86 | 87 | 5e-83 |
| Streptococcus dysgalactiae subsp. equisimilis strain NCTC7136 | rpoE | DNA-directed RNA polymerase delta subunit (EC 2.7.7.6) | 16513739 | 87 | 86 | 87 | 5e-83 |
| Streptococcus dysgalactiae subsp. equisimilis strain NCTC8543 | rpoE | DNA-directed RNA polymerase delta subunit (EC 2.7.7.6) | 16513739 | 87 | 86 | 87 | 5e-83 |
| Streptococcus dysgalactiae subsp. equisimilis strain NCTC8546 | rpoE | DNA-directed RNA polymerase delta subunit (EC 2.7.7.6) | 16513739 | 87 | 86 | 87 | 5e-83 |
| Streptococcus dysgalactiae subsp. equisimilis strain NCTC9413 | rpoE | DNA-directed RNA polymerase delta subunit (EC 2.7.7.6) | 16513739 | 87 | 86 | 87 | 5e-83 |
| Streptococcus dysgalactiae subsp. equisimilis strain NCTC9414 | rpoE | DNA-directed RNA polymerase delta subunit (EC 2.7.7.6) | 16513739 | 87 | 86 | 87 | 5e-83 |
| Streptococcus dysgalactiae subsp. equisimilis strain NCTC9603 | rpoE | DNA-directed RNA polymerase delta subunit (EC 2.7.7.6) | 16513739 | 87 | 86 | 87 | 5e-83 |
| Streptococcus dysgalactiae subsp. equisimilis strain SS1575 | rpoE | DNA-directed RNA polymerase delta subunit (EC 2.7.7.6) | 16513739 | 87 | 86 | 87 | 5e-83 |
| Streptococcus dysgalactiae subsp. equisimilis strain T642 | rpoE | DNA-directed RNA polymerase delta subunit (EC 2.7.7.6) | 16513739 | 89 | 89 | 85 | 0 |
| Streptococcus dysgalactiae subsp. equisimilis strain UT_4031CC | rpoE | DNA-directed RNA polymerase delta subunit (EC 2.7.7.6) | 16513739 | 87 | 86 | 87 | 5e-83 |
| Streptococcus dysgalactiae subsp. equisimilis strain UT_4231_KK | rpoE | DNA-directed RNA polymerase delta subunit (EC 2.7.7.6) | 16513739 | 87 | 86 | 87 | 5e-83 |
| Streptococcus dysgalactiae subsp. equisimilis strain UT_4234_DH | rpoE | DNA-directed RNA polymerase delta subunit (EC 2.7.7.6) | 16513739 | 87 | 86 | 87 | 3e-83 |
| Streptococcus dysgalactiae subsp. equisimilis strain UT_4241_XS | rpoE | DNA-directed RNA polymerase delta subunit (EC 2.7.7.6) | 16513739 | 87 | 86 | 87 | 5e-83 |
| Streptococcus dysgalactiae subsp. equisimilis strain UT_4242_AB | rpoE | DNA-directed RNA polymerase delta subunit (EC 2.7.7.6) | 16513739 | 87 | 86 | 87 | 5e-83 |
| Streptococcus dysgalactiae subsp. equisimilis strain UT_4255RC | rpoE | DNA-directed RNA polymerase delta subunit (EC 2.7.7.6) | 16513739 | 87 | 86 | 87 | 3e-83 |
| Streptococcus dysgalactiae subsp. equisimilis strain UT_4277_BB | rpoE | DNA-directed RNA polymerase delta subunit (EC 2.7.7.6) | 16513739 | 87 | 86 | 87 | 5e-83 |
| Streptococcus dysgalactiae subsp. equisimilis strain UT_4966_RC | rpoE | DNA-directed RNA polymerase delta subunit (EC 2.7.7.6) | 16513739 | 87 | 86 | 87 | 5e-83 |
| Streptococcus dysgalactiae subsp. equisimilis strain UT-5345 | rpoE | DNA-directed RNA polymerase delta subunit (EC 2.7.7.6) | 16513739 | 87 | 86 | 87 | 5e-83 |
| Streptococcus dysgalactiae subsp. equisimilis strain UT-5354 | rpoE | DNA-directed RNA polymerase delta subunit (EC 2.7.7.6) | 16513739 | 87 | 86 | 87 | 5e-83 |
| Streptococcus dysgalactiae subsp. equisimilis strain UT-SS1069 | rpoE | DNA-directed RNA polymerase delta subunit (EC 2.7.7.6) | 16513739 | 87 | 86 | 87 | 5e-83 |
| Streptococcus dysgalactiae subsp. equisimilis strain UT-SS957 | rpoE | DNA-directed RNA polymerase delta subunit (EC 2.7.7.6) | 16513739 | 87 | 86 | 87 | 5e-83 |
| Streptococcus dysgalactiae subsp. equisimilis strain WCHSDSE-1 | rpoE | DNA-directed RNA polymerase delta subunit (EC 2.7.7.6) | 16513739 | 87 | 86 | 87 | 5e-83 |
| Streptococcus dysgalactiae subsp. equisimilis 167 | dltA | D-alanine--poly(phosphoribitol) ligase subunit 1 (EC 6.1.1.13) | 16166534 | 100 | 100 | 90 | 1e-275 |
| Streptococcus dysgalactiae subsp. equisimilis AC-2713 | dltA | D-alanine--poly(phosphoribitol) ligase subunit 1 (EC 6.1.1.13) | 16166534 | 100 | 100 | 90 | 1e-275 |
| Streptococcus dysgalactiae subsp. equisimilis AKSDE4288 | dltA | D-alanine--poly(phosphoribitol) ligase subunit 1 (EC 6.1.1.13) | 16166534 | 100 | 100 | 90 | 1e-275 |
| Streptococcus dysgalactiae subsp. equisimilis ATCC 12394 | dltA | D-alanine--poly(phosphoribitol) ligase subunit 1 (EC 6.1.1.13) | 16166534 | 100 | 100 | 90 | 1e-274 |
| Streptococcus dysgalactiae subsp. equisimilis GGS_124 | dltA | D-alanine--poly(phosphoribitol) ligase subunit 1 (EC 6.1.1.13) | 16166534 | 100 | 100 | 90 | 1e-275 |
| Streptococcus dysgalactiae subsp. equisimilis RE378 | dltA | D-alanine--poly(phosphoribitol) ligase subunit 1 (EC 6.1.1.13) | 16166534 | 100 | 100 | 90 | 1e-275 |
| Streptococcus dysgalactiae subsp. equisimilis SD SCDR1 | dltA | D-alanine--poly(phosphoribitol) ligase subunit 1 (EC 6.1.1.13) | 16166534 | 100 | 100 | 90 | 1e-275 |
| Streptococcus dysgalactiae subsp. equisimilis SK1249 | dltA | D-alanine--poly(phosphoribitol) ligase subunit 1 (EC 6.1.1.13) | 16166534 | 100 | 100 | 90 | 1e-274 |
| Streptococcus dysgalactiae subsp. equisimilis SK1250 | dltA | D-alanine--poly(phosphoribitol) ligase subunit 1 (EC 6.1.1.13) | 16166534 | 100 | 100 | 90 | 1e-275 |
| Streptococcus dysgalactiae subsp. equisimilis strain ASDSE_96 | dltA | D-alanine--poly(phosphoribitol) ligase subunit 1 (EC 6.1.1.13) | 16166534 | 100 | 100 | 90 | 1e-274 |
| Streptococcus dysgalactiae subsp. equisimilis strain ASDSE_99 | dltA | D-alanine--poly(phosphoribitol) ligase subunit 1 (EC 6.1.1.13) | 16166534 | 100 | 100 | 90 | 1e-275 |
| Streptococcus dysgalactiae subsp. equisimilis strain C161L1 | dltA | D-alanine--poly(phosphoribitol) ligase subunit 1 (EC 6.1.1.13) | 16166534 | 100 | 100 | 90 | 0 |
| Streptococcus dysgalactiae subsp. equisimilis strain KNZ01 | dltA | D-alanine--poly(phosphoribitol) ligase subunit 1 (EC 6.1.1.13) | 16166534 | 100 | 100 | 90 | 1e-275 |
| Streptococcus dysgalactiae subsp. equisimilis strain KNZ03 | dltA | D-alanine--poly(phosphoribitol) ligase subunit 1 (EC 6.1.1.13) | 16166534 | 100 | 100 | 90 | 1e-274 |
| Streptococcus dysgalactiae subsp. equisimilis strain KNZ04 | dltA | D-alanine--poly(phosphoribitol) ligase subunit 1 (EC 6.1.1.13) | 16166534 | 100 | 100 | 90 | 1e-274 |
| Streptococcus dysgalactiae subsp. equisimilis strain KNZ06 | dltA | D-alanine--poly(phosphoribitol) ligase subunit 1 (EC 6.1.1.13) | 16166534 | 100 | 100 | 90 | 1e-275 |
| Streptococcus dysgalactiae subsp. equisimilis strain KNZ07 | dltA | D-alanine--poly(phosphoribitol) ligase subunit 1 (EC 6.1.1.13) | 16166534 | 100 | 100 | 90 | 1e-275 |
| Streptococcus dysgalactiae subsp. equisimilis strain KNZ10 | dltA | D-alanine--poly(phosphoribitol) ligase subunit 1 (EC 6.1.1.13) | 16166534 | 100 | 100 | 90 | 1e-275 |
| Streptococcus dysgalactiae subsp. equisimilis strain KNZ12 | dltA | D-alanine--poly(phosphoribitol) ligase subunit 1 (EC 6.1.1.13) | 16166534 | 100 | 100 | 90 | 1e-275 |
| Streptococcus dysgalactiae subsp. equisimilis strain KNZ15 | dltA | D-alanine--poly(phosphoribitol) ligase subunit 1 (EC 6.1.1.13) | 16166534 | 100 | 100 | 90 | 1e-275 |
| Streptococcus dysgalactiae subsp. equisimilis strain KNZ16 | dltA | D-alanine--poly(phosphoribitol) ligase subunit 1 (EC 6.1.1.13) | 16166534 | 100 | 100 | 90 | 1e-274 |
| Streptococcus dysgalactiae subsp. equisimilis strain NCTC10321 | dltA | D-alanine--poly(phosphoribitol) ligase subunit 1 (EC 6.1.1.13) | 16166534 | 100 | 100 | 90 | 1e-275 |
| Streptococcus dysgalactiae subsp. equisimilis strain NCTC11554 | dltA | D-alanine--poly(phosphoribitol) ligase subunit 1 (EC 6.1.1.13) | 16166534 | 100 | 100 | 90 | 1e-274 |
| Streptococcus dysgalactiae subsp. equisimilis strain NCTC11555 | dltA | D-alanine--poly(phosphoribitol) ligase subunit 1 (EC 6.1.1.13) | 16166534 | 100 | 100 | 90 | 1e-275 |
| Streptococcus dysgalactiae subsp. equisimilis strain NCTC11556 | dltA | D-alanine--poly(phosphoribitol) ligase subunit 1 (EC 6.1.1.13) | 16166534 | 100 | 100 | 90 | 1e-274 |
| Streptococcus dysgalactiae subsp. equisimilis strain NCTC11557 | dltA | D-alanine--poly(phosphoribitol) ligase subunit 1 (EC 6.1.1.13) | 16166534 | 100 | 100 | 90 | 1e-275 |
| Streptococcus dysgalactiae subsp. equisimilis strain NCTC11564 | dltA | D-alanine--poly(phosphoribitol) ligase subunit 1 (EC 6.1.1.13) | 16166534 | 100 | 100 | 90 | 1e-275 |
| Streptococcus dysgalactiae subsp. equisimilis strain NCTC5370 | dltA | D-alanine--poly(phosphoribitol) ligase subunit 1 (EC 6.1.1.13) | 16166534 | 100 | 100 | 90 | 1e-275 |
| Streptococcus dysgalactiae subsp. equisimilis strain NCTC5371 | dltA | D-alanine--poly(phosphoribitol) ligase subunit 1 (EC 6.1.1.13) | 16166534 | 100 | 100 | 90 | 1e-274 |
| Streptococcus dysgalactiae subsp. equisimilis strain NCTC6179 | dltA | D-alanine--poly(phosphoribitol) ligase subunit 1 (EC 6.1.1.13) | 16166534 | 100 | 100 | 89 | 1e-273 |
| Streptococcus dysgalactiae subsp. equisimilis strain NCTC6181 | dltA | D-alanine--poly(phosphoribitol) ligase subunit 1 (EC 6.1.1.13) | 16166534 | 100 | 100 | 89 | 1e-273 |
| Streptococcus dysgalactiae subsp. equisimilis strain NCTC6407 | dltA | D-alanine--poly(phosphoribitol) ligase subunit 1 (EC 6.1.1.13) | 16166534 | 100 | 100 | 90 | 1e-275 |
| Streptococcus dysgalactiae subsp. equisimilis strain NCTC7136 | dltA | D-alanine--poly(phosphoribitol) ligase subunit 1 (EC 6.1.1.13) | 16166534 | 100 | 100 | 90 | 1e-274 |
| Streptococcus dysgalactiae subsp. equisimilis strain NCTC8543 | dltA | D-alanine--poly(phosphoribitol) ligase subunit 1 (EC 6.1.1.13) | 16166534 | 100 | 100 | 90 | 1e-275 |
| Streptococcus dysgalactiae subsp. equisimilis strain NCTC8546 | dltA | D-alanine--poly(phosphoribitol) ligase subunit 1 (EC 6.1.1.13) | 16166534 | 100 | 100 | 90 | 1e-275 |
| Streptococcus dysgalactiae subsp. equisimilis strain NCTC9413 | dltA | D-alanine--poly(phosphoribitol) ligase subunit 1 (EC 6.1.1.13) | 16166534 | 100 | 100 | 90 | 1e-275 |
| Streptococcus dysgalactiae subsp. equisimilis strain NCTC9414 | dltA | D-alanine--poly(phosphoribitol) ligase subunit 1 (EC 6.1.1.13) | 16166534 | 100 | 100 | 90 | 1e-274 |
| Streptococcus dysgalactiae subsp. equisimilis strain NCTC9603 | dltA | D-alanine--poly(phosphoribitol) ligase subunit 1 (EC 6.1.1.13) | 16166534 | 100 | 100 | 90 | 1e-275 |
| Streptococcus dysgalactiae subsp. equisimilis strain SS1575 | dltA | D-alanine--poly(phosphoribitol) ligase subunit 1 (EC 6.1.1.13) | 16166534 | 100 | 100 | 90 | 1e-274 |
| Streptococcus dysgalactiae subsp. equisimilis strain T642 | dltA | D-alanine--poly(phosphoribitol) ligase subunit 1 (EC 6.1.1.13) | 16166534 | 100 | 100 | 90 | 0.0 |
| Streptococcus dysgalactiae subsp. equisimilis strain UT_4031CC | dltA | D-alanine--poly(phosphoribitol) ligase subunit 1 (EC 6.1.1.13) | 16166534 | 100 | 100 | 90 | 1e-275 |
| Streptococcus dysgalactiae subsp. equisimilis strain UT_4231_KK | dltA | D-alanine--poly(phosphoribitol) ligase subunit 1 (EC 6.1.1.13) | 16166534 | 100 | 100 | 90 | 1e-275 |
| Streptococcus dysgalactiae subsp. equisimilis strain UT_4234_DH | dltA | D-alanine--poly(phosphoribitol) ligase subunit 1 (EC 6.1.1.13) | 16166534 | 100 | 100 | 90 | 1e-275 |
| Streptococcus dysgalactiae subsp. equisimilis strain UT_4241_XS | dltA | D-alanine--poly(phosphoribitol) ligase subunit 1 (EC 6.1.1.13) | 16166534 | 100 | 100 | 90 | 1e-275 |
| Streptococcus dysgalactiae subsp. equisimilis strain UT_4242_AB | dltA | D-alanine--poly(phosphoribitol) ligase subunit 1 (EC 6.1.1.13) | 16166534 | 100 | 100 | 90 | 1e-274 |
| Streptococcus dysgalactiae subsp. equisimilis strain UT_4255RC | dltA | D-alanine--poly(phosphoribitol) ligase subunit 1 (EC 6.1.1.13) | 16166534 | 100 | 100 | 90 | 1e-275 |
| Streptococcus dysgalactiae subsp. equisimilis strain UT_4277_BB | dltA | D-alanine--poly(phosphoribitol) ligase subunit 1 (EC 6.1.1.13) | 16166534 | 100 | 100 | 90 | 1e-275 |
| Streptococcus dysgalactiae subsp. equisimilis strain UT_4966_RC | dltA | D-alanine--poly(phosphoribitol) ligase subunit 1 (EC 6.1.1.13) | 16166534 | 100 | 100 | 90 | 1e-275 |
| Streptococcus dysgalactiae subsp. equisimilis strain UT-5345 | dltA | D-alanine--poly(phosphoribitol) ligase subunit 1 (EC 6.1.1.13) | 16166534 | 100 | 100 | 90 | 1e-275 |
| Streptococcus dysgalactiae subsp. equisimilis strain UT-5354 | dltA | D-alanine--poly(phosphoribitol) ligase subunit 1 (EC 6.1.1.13) | 16166534 | 100 | 100 | 90 | 1e-275 |
| Streptococcus dysgalactiae subsp. equisimilis strain UT-SS1069 | dltA | D-alanine--poly(phosphoribitol) ligase subunit 1 (EC 6.1.1.13) | 16166534 | 100 | 100 | 90 | 1e-275 |
| Streptococcus dysgalactiae subsp. equisimilis strain UT-SS957 | dltA | D-alanine--poly(phosphoribitol) ligase subunit 1 (EC 6.1.1.13) | 16166534 | 100 | 100 | 90 | 1e-275 |
| Streptococcus dysgalactiae subsp. equisimilis strain WCHSDSE-1 | dltA | D-alanine--poly(phosphoribitol) ligase subunit 1 (EC 6.1.1.13) | 16166534 | 100 | 100 | 90 | 1e-275 |
| Streptococcus dysgalactiae subsp. equisimilis 167 | cydA | Cytochrome d ubiquinol oxidase subunit I (EC 1.10.3.-) | 15813741 | 95 | 95 | 81 | 1e-230 |
| Streptococcus dysgalactiae subsp. equisimilis AC-2713 | cydA | Cytochrome d ubiquinol oxidase subunit I (EC 1.10.3.-) | 15813741 | 95 | 95 | 81 | 1e-230 |
| Streptococcus dysgalactiae subsp. equisimilis AKSDE4288 | cydA | Cytochrome d ubiquinol oxidase subunit I (EC 1.10.3.-) | 15813741 | 95 | 95 | 81 | 1e-230 |
| Streptococcus dysgalactiae subsp. equisimilis ATCC 12394 | cydA | Cytochrome d ubiquinol oxidase subunit I (EC 1.10.3.-) | 15813741 | 95 | 95 | 81 | 1e-230 |
| Streptococcus dysgalactiae subsp. equisimilis GGS_124 | cydA | Cytochrome d ubiquinol oxidase subunit I (EC 1.10.3.-) | 15813741 | 95 | 95 | 81 | 1e-230 |
| Streptococcus dysgalactiae subsp. equisimilis RE378 | cydA | Cytochrome d ubiquinol oxidase subunit I (EC 1.10.3.-) | 15813741 | 95 | 95 | 81 | 1e-230 |
| Streptococcus dysgalactiae subsp. equisimilis SD SCDR1 | cydA | Cytochrome d ubiquinol oxidase subunit I (EC 1.10.3.-) | 15813741 | 95 | 95 | 81 | 1e-230 |
| Streptococcus dysgalactiae subsp. equisimilis SK1249 | cydA | Cytochrome d ubiquinol oxidase subunit I (EC 1.10.3.-) | 15813741 | 95 | 95 | 81 | 1e-230 |
| Streptococcus dysgalactiae subsp. equisimilis SK1250 | cydA | Cytochrome d ubiquinol oxidase subunit I (EC 1.10.3.-) | 15813741 | 95 | 95 | 81 | 1e-230 |
| Streptococcus dysgalactiae subsp. equisimilis strain ASDSE_96 | cydA | Cytochrome d ubiquinol oxidase subunit I (EC 1.10.3.-) | 15813741 | 95 | 95 | 81 | 1e-230 |
| Streptococcus dysgalactiae subsp. equisimilis strain ASDSE_99 | cydA | Cytochrome d ubiquinol oxidase subunit I (EC 1.10.3.-) | 15813741 | 95 | 95 | 81 | 1e-230 |
| Streptococcus dysgalactiae subsp. equisimilis strain C161L1 | cydA | Cytochrome d ubiquinol oxidase subunit I (EC 1.10.3.-) | 15813741 | 100 | 100 | 80 | 0 |
| Streptococcus dysgalactiae subsp. equisimilis strain KNZ01 | cydA | Cytochrome d ubiquinol oxidase subunit I (EC 1.10.3.-) | 15813741 | 95 | 95 | 81 | 1e-230 |
| Streptococcus dysgalactiae subsp. equisimilis strain KNZ03 | cydA | Cytochrome d ubiquinol oxidase subunit I (EC 1.10.3.-) | 15813741 | 95 | 95 | 81 | 1e-230 |
| Streptococcus dysgalactiae subsp. equisimilis strain KNZ04 | cydA | Cytochrome d ubiquinol oxidase subunit I (EC 1.10.3.-) | 15813741 | 95 | 95 | 81 | 1e-230 |
| Streptococcus dysgalactiae subsp. equisimilis strain KNZ06 | cydA | Cytochrome d ubiquinol oxidase subunit I (EC 1.10.3.-) | 15813741 | 95 | 95 | 81 | 1e-229 |
| Streptococcus dysgalactiae subsp. equisimilis strain KNZ07 | cydA | Cytochrome d ubiquinol oxidase subunit I (EC 1.10.3.-) | 15813741 | 95 | 95 | 81 | 1e-230 |
| Streptococcus dysgalactiae subsp. equisimilis strain KNZ10 | cydA | Cytochrome d ubiquinol oxidase subunit I (EC 1.10.3.-) | 15813741 | 95 | 95 | 81 | 1e-230 |
| Streptococcus dysgalactiae subsp. equisimilis strain KNZ12 | cydA | Cytochrome d ubiquinol oxidase subunit I (EC 1.10.3.-) | 15813741 | 95 | 95 | 81 | 1e-230 |
| Streptococcus dysgalactiae subsp. equisimilis strain KNZ15 | cydA | Cytochrome d ubiquinol oxidase subunit I (EC 1.10.3.-) | 15813741 | 95 | 95 | 81 | 1e-230 |
| Streptococcus dysgalactiae subsp. equisimilis strain KNZ16 | cydA | Cytochrome d ubiquinol oxidase subunit I (EC 1.10.3.-) | 15813741 | 95 | 95 | 81 | 1e-230 |
| Streptococcus dysgalactiae subsp. equisimilis strain NCTC10321 | cydA | Cytochrome d ubiquinol oxidase subunit I (EC 1.10.3.-) | 15813741 | 95 | 95 | 81 | 1e-229 |
| Streptococcus dysgalactiae subsp. equisimilis strain NCTC11554 | cydA | Cytochrome d ubiquinol oxidase subunit I (EC 1.10.3.-) | 15813741 | 95 | 95 | 81 | 1e-230 |
| Streptococcus dysgalactiae subsp. equisimilis strain NCTC11555 | cydA | Cytochrome d ubiquinol oxidase subunit I (EC 1.10.3.-) | 15813741 | 95 | 95 | 81 | 1e-230 |
| Streptococcus dysgalactiae subsp. equisimilis strain NCTC11556 | cydA | Cytochrome d ubiquinol oxidase subunit I (EC 1.10.3.-) | 15813741 | 95 | 95 | 81 | 1e-230 |
| Streptococcus dysgalactiae subsp. equisimilis strain NCTC11557 | cydA | Cytochrome d ubiquinol oxidase subunit I (EC 1.10.3.-) | 15813741 | 95 | 95 | 81 | 1e-230 |
| Streptococcus dysgalactiae subsp. equisimilis strain NCTC11564 | cydA | Cytochrome d ubiquinol oxidase subunit I (EC 1.10.3.-) | 15813741 | 95 | 95 | 81 | 1e-230 |
| Streptococcus dysgalactiae subsp. equisimilis strain NCTC5370 | cydA | Cytochrome d ubiquinol oxidase subunit I (EC 1.10.3.-) | 15813741 | 95 | 95 | 81 | 1e-230 |
| Streptococcus dysgalactiae subsp. equisimilis strain NCTC5371 | cydA | Cytochrome d ubiquinol oxidase subunit I (EC 1.10.3.-) | 15813741 | 95 | 95 | 81 | 1e-230 |
| Streptococcus dysgalactiae subsp. equisimilis strain NCTC6179 | cydA | Cytochrome d ubiquinol oxidase subunit I (EC 1.10.3.-) | 15813741 | 95 | 95 | 81 | 1e-230 |
| Streptococcus dysgalactiae subsp. equisimilis strain NCTC6181 | cydA | Cytochrome d ubiquinol oxidase subunit I (EC 1.10.3.-) | 15813741 | 95 | 95 | 81 | 1e-230 |
| Streptococcus dysgalactiae subsp. equisimilis strain NCTC6407 | cydA | Cytochrome d ubiquinol oxidase subunit I (EC 1.10.3.-) | 15813741 | 95 | 95 | 81 | 1e-229 |
| Streptococcus dysgalactiae subsp. equisimilis strain NCTC7136 | cydA | Cytochrome d ubiquinol oxidase subunit I (EC 1.10.3.-) | 15813741 | 95 | 95 | 81 | 1e-228 |
| Streptococcus dysgalactiae subsp. equisimilis strain NCTC8543 | cydA | Cytochrome d ubiquinol oxidase subunit I (EC 1.10.3.-) | 15813741 | 95 | 95 | 81 | 1e-230 |
| Streptococcus dysgalactiae subsp. equisimilis strain NCTC8546 | cydA | Cytochrome d ubiquinol oxidase subunit I (EC 1.10.3.-) | 15813741 | 95 | 95 | 81 | 1e-230 |
| Streptococcus dysgalactiae subsp. equisimilis strain NCTC9413 | cydA | Cytochrome d ubiquinol oxidase subunit I (EC 1.10.3.-) | 15813741 | 95 | 95 | 81 | 1e-230 |
| Streptococcus dysgalactiae subsp. equisimilis strain NCTC9414 | cydA | Cytochrome d ubiquinol oxidase subunit I (EC 1.10.3.-) | 15813741 | 95 | 95 | 81 | 1e-230 |
| Streptococcus dysgalactiae subsp. equisimilis strain NCTC9603 | cydA | Cytochrome d ubiquinol oxidase subunit I (EC 1.10.3.-) | 15813741 | 95 | 95 | 81 | 1e-230 |
| Streptococcus dysgalactiae subsp. equisimilis strain SS1575 | cydA | Cytochrome d ubiquinol oxidase subunit I (EC 1.10.3.-) | 15813741 | 95 | 95 | 81 | 1e-230 |
| Streptococcus dysgalactiae subsp. equisimilis strain T642 | cydA | Cytochrome d ubiquinol oxidase subunit I (EC 1.10.3.-) | 15813741 | 100 | 100 | 80 | 0.0 |
| Streptococcus dysgalactiae subsp. equisimilis strain UT_4031CC | cydA | Cytochrome d ubiquinol oxidase subunit I (EC 1.10.3.-) | 15813741 | 95 | 95 | 81 | 1e-229 |
| Streptococcus dysgalactiae subsp. equisimilis strain UT_4231_KK | cydA | Cytochrome d ubiquinol oxidase subunit I (EC 1.10.3.-) | 15813741 | 95 | 95 | 81 | 1e-229 |
| Streptococcus dysgalactiae subsp. equisimilis strain UT_4234_DH | cydA | Cytochrome d ubiquinol oxidase subunit I (EC 1.10.3.-) | 15813741 | 95 | 95 | 81 | 1e-230 |
| Streptococcus dysgalactiae subsp. equisimilis strain UT_4241_XS | cydA | Cytochrome d ubiquinol oxidase subunit I (EC 1.10.3.-) | 15813741 | 95 | 95 | 81 | 1e-230 |
| Streptococcus dysgalactiae subsp. equisimilis strain UT_4242_AB | cydA | Cytochrome d ubiquinol oxidase subunit I (EC 1.10.3.-) | 15813741 | 95 | 95 | 81 | 1e-230 |
| Streptococcus dysgalactiae subsp. equisimilis strain UT_4255RC | cydA | Cytochrome d ubiquinol oxidase subunit I (EC 1.10.3.-) | 15813741 | 95 | 95 | 81 | 1e-230 |
| Streptococcus dysgalactiae subsp. equisimilis strain UT_4277_BB | cydA | Cytochrome d ubiquinol oxidase subunit I (EC 1.10.3.-) | 15813741 | 95 | 95 | 81 | 1e-229 |
| Streptococcus dysgalactiae subsp. equisimilis strain UT_4966_RC | cydA | Cytochrome d ubiquinol oxidase subunit I (EC 1.10.3.-) | 15813741 | 95 | 95 | 81 | 1e-230 |
| Streptococcus dysgalactiae subsp. equisimilis strain UT-5345 | cydA | Cytochrome d ubiquinol oxidase subunit I (EC 1.10.3.-) | 15813741 | 95 | 95 | 81 | 1e-230 |
| Streptococcus dysgalactiae subsp. equisimilis strain UT-5354 | cydA | Cytochrome d ubiquinol oxidase subunit I (EC 1.10.3.-) | 15813741 | 95 | 95 | 81 | 1e-230 |
| Streptococcus dysgalactiae subsp. equisimilis strain UT-SS1069 | cydA | Cytochrome d ubiquinol oxidase subunit I (EC 1.10.3.-) | 15813741 | 95 | 95 | 81 | 1e-230 |
| Streptococcus dysgalactiae subsp. equisimilis strain UT-SS957 | cydA | Cytochrome d ubiquinol oxidase subunit I (EC 1.10.3.-) | 15813741 | 95 | 95 | 81 | 1e-230 |
| Streptococcus dysgalactiae subsp. equisimilis strain WCHSDSE-1 | cydA | Cytochrome d ubiquinol oxidase subunit I (EC 1.10.3.-) | 15813741 | 95 | 95 | 81 | 1e-230 |
| Streptococcus dysgalactiae subsp. equisimilis 167 | pyrG | CTP synthase (EC 6.3.4.2) | 12207705 | 99 | 99 | 90 | 1e-284 |
| Streptococcus dysgalactiae subsp. equisimilis AC-2713 | pyrG | CTP synthase (EC 6.3.4.2) | 12207705 | 99 | 99 | 90 | 1e-284 |
| Streptococcus dysgalactiae subsp. equisimilis AKSDE4288 | SP_0494 | CTP synthase (EC 6.3.4.2) | 12207705 | 99 | 99 | 90 | 1e-284 |
| Streptococcus dysgalactiae subsp. equisimilis ATCC 12394 | pyrG | CTP synthase (EC 6.3.4.2) | 12207705 | 99 | 99 | 90 | 1e-284 |
| Streptococcus dysgalactiae subsp. equisimilis GGS_124 | pyrG | CTP synthase (EC 6.3.4.2) | 12207705 | 99 | 99 | 90 | 1e-284 |
| Streptococcus dysgalactiae subsp. equisimilis RE378 | pyrG | CTP synthase (EC 6.3.4.2) | 12207705 | 99 | 99 | 90 | 1e-284 |
| Streptococcus dysgalactiae subsp. equisimilis SD SCDR1 | SP_0494 | CTP synthase (EC 6.3.4.2) | 12207705 | 99 | 99 | 90 | 1e-284 |
| Streptococcus dysgalactiae subsp. equisimilis SK1249 | pyrG | CTP synthase (EC 6.3.4.2) | 12207705 | 99 | 99 | 90 | 1e-284 |
| Streptococcus dysgalactiae subsp. equisimilis SK1250 | pyrG | CTP synthase (EC 6.3.4.2) | 12207705 | 99 | 99 | 90 | 1e-284 |
| Streptococcus dysgalactiae subsp. equisimilis strain ASDSE_96 | SP_0494 | CTP synthase (EC 6.3.4.2) | 12207705 | 99 | 99 | 90 | 1e-284 |
| Streptococcus dysgalactiae subsp. equisimilis strain ASDSE_99 | SP_0494 | CTP synthase (EC 6.3.4.2) | 12207705 | 99 | 99 | 90 | 1e-284 |
| Streptococcus dysgalactiae subsp. equisimilis strain C161L1 | SP_0494 | CTP synthase (EC 6.3.4.2) | 12207705 | 99 | 99 | 90 | 0.0 |
| Streptococcus dysgalactiae subsp. equisimilis strain KNZ01 | SP_0494 | CTP synthase (EC 6.3.4.2) | 12207705 | 99 | 99 | 90 | 1e-284 |
| Streptococcus dysgalactiae subsp. equisimilis strain KNZ03 | SP_0494 | CTP synthase (EC 6.3.4.2) | 12207705 | 99 | 99 | 90 | 1e-284 |
| Streptococcus dysgalactiae subsp. equisimilis strain KNZ04 | SP_0494 | CTP synthase (EC 6.3.4.2) | 12207705 | 99 | 99 | 90 | 1e-284 |
| Streptococcus dysgalactiae subsp. equisimilis strain KNZ06 | SP_0494 | CTP synthase (EC 6.3.4.2) | 12207705 | 99 | 99 | 90 | 1e-284 |
| Streptococcus dysgalactiae subsp. equisimilis strain KNZ07 | SP_0494 | CTP synthase (EC 6.3.4.2) | 12207705 | 99 | 99 | 90 | 1e-284 |
| Streptococcus dysgalactiae subsp. equisimilis strain KNZ10 | SP_0494 | CTP synthase (EC 6.3.4.2) | 12207705 | 99 | 99 | 90 | 1e-284 |
| Streptococcus dysgalactiae subsp. equisimilis strain KNZ12 | SP_0494 | CTP synthase (EC 6.3.4.2) | 12207705 | 99 | 99 | 90 | 1e-284 |
| Streptococcus dysgalactiae subsp. equisimilis strain KNZ15 | SP_0494 | CTP synthase (EC 6.3.4.2) | 12207705 | 99 | 99 | 90 | 1e-284 |
| Streptococcus dysgalactiae subsp. equisimilis strain KNZ16 | SP_0494 | CTP synthase (EC 6.3.4.2) | 12207705 | 99 | 99 | 90 | 1e-284 |
| Streptococcus dysgalactiae subsp. equisimilis strain NCTC10321 | SP_0494 | CTP synthase (EC 6.3.4.2) | 12207705 | 99 | 99 | 90 | 1e-284 |
| Streptococcus dysgalactiae subsp. equisimilis strain NCTC11554 | SP_0494 | CTP synthase (EC 6.3.4.2) | 12207705 | 99 | 99 | 90 | 1e-284 |
| Streptococcus dysgalactiae subsp. equisimilis strain NCTC11555 | SP_0494 | CTP synthase (EC 6.3.4.2) | 12207705 | 99 | 99 | 90 | 1e-284 |
| Streptococcus dysgalactiae subsp. equisimilis strain NCTC11556 | SP_0494 | CTP synthase (EC 6.3.4.2) | 12207705 | 99 | 99 | 90 | 1e-284 |
| Streptococcus dysgalactiae subsp. equisimilis strain NCTC11557 | SP_0494 | CTP synthase (EC 6.3.4.2) | 12207705 | 99 | 99 | 90 | 1e-284 |
| Streptococcus dysgalactiae subsp. equisimilis strain NCTC11564 | SP_0494 | CTP synthase (EC 6.3.4.2) | 12207705 | 99 | 99 | 90 | 1e-284 |
| Streptococcus dysgalactiae subsp. equisimilis strain NCTC5370 | SP_0494 | CTP synthase (EC 6.3.4.2) | 12207705 | 99 | 99 | 90 | 1e-284 |
| Streptococcus dysgalactiae subsp. equisimilis strain NCTC5371 | SP_0494 | CTP synthase (EC 6.3.4.2) | 12207705 | 99 | 99 | 90 | 1e-284 |
| Streptococcus dysgalactiae subsp. equisimilis strain NCTC6179 | SP_0494 | CTP synthase (EC 6.3.4.2) | 12207705 | 99 | 99 | 90 | 1e-283 |
| Streptococcus dysgalactiae subsp. equisimilis strain NCTC6181 | SP_0494 | CTP synthase (EC 6.3.4.2) | 12207705 | 99 | 99 | 90 | 1e-283 |
| Streptococcus dysgalactiae subsp. equisimilis strain NCTC6407 | SP_0494 | CTP synthase (EC 6.3.4.2) | 12207705 | 99 | 99 | 90 | 1e-284 |
| Streptococcus dysgalactiae subsp. equisimilis strain NCTC7136 | SP_0494 | CTP synthase (EC 6.3.4.2) | 12207705 | 99 | 99 | 90 | 1e-284 |
| Streptococcus dysgalactiae subsp. equisimilis strain NCTC8543 | SP_0494 | CTP synthase (EC 6.3.4.2) | 12207705 | 99 | 99 | 90 | 1e-284 |
| Streptococcus dysgalactiae subsp. equisimilis strain NCTC8546 | SP_0494 | CTP synthase (EC 6.3.4.2) | 12207705 | 99 | 99 | 90 | 1e-284 |
| Streptococcus dysgalactiae subsp. equisimilis strain NCTC9413 | SP_0494 | CTP synthase (EC 6.3.4.2) | 12207705 | 99 | 99 | 90 | 1e-284 |
| Streptococcus dysgalactiae subsp. equisimilis strain NCTC9414 | SP_0494 | CTP synthase (EC 6.3.4.2) | 12207705 | 99 | 99 | 90 | 1e-284 |
| Streptococcus dysgalactiae subsp. equisimilis strain NCTC9603 | SP_0494 | CTP synthase (EC 6.3.4.2) | 12207705 | 99 | 99 | 90 | 1e-284 |
| Streptococcus dysgalactiae subsp. equisimilis strain SS1575 | SP_0494 | CTP synthase (EC 6.3.4.2) | 12207705 | 99 | 99 | 90 | 1e-284 |
| Streptococcus dysgalactiae subsp. equisimilis strain T642 | SP_0494 | CTP synthase (EC 6.3.4.2) | 12207705 | 99 | 99 | 90 | 0 |
| Streptococcus dysgalactiae subsp. equisimilis strain UT_4031CC | SP_0494 | CTP synthase (EC 6.3.4.2) | 12207705 | 99 | 99 | 90 | 1e-284 |
| Streptococcus dysgalactiae subsp. equisimilis strain UT_4231_KK | SP_0494 | CTP synthase (EC 6.3.4.2) | 12207705 | 99 | 99 | 90 | 1e-284 |
| Streptococcus dysgalactiae subsp. equisimilis strain UT_4234_DH | SP_0494 | CTP synthase (EC 6.3.4.2) | 12207705 | 99 | 99 | 90 | 1e-284 |
| Streptococcus dysgalactiae subsp. equisimilis strain UT_4241_XS | SP_0494 | CTP synthase (EC 6.3.4.2) | 12207705 | 99 | 99 | 90 | 1e-284 |
| Streptococcus dysgalactiae subsp. equisimilis strain UT_4242_AB | SP_0494 | CTP synthase (EC 6.3.4.2) | 12207705 | 99 | 99 | 90 | 1e-284 |
| Streptococcus dysgalactiae subsp. equisimilis strain UT_4255RC | SP_0494 | CTP synthase (EC 6.3.4.2) | 12207705 | 99 | 99 | 90 | 1e-284 |
| Streptococcus dysgalactiae subsp. equisimilis strain UT_4277_BB | SP_0494 | CTP synthase (EC 6.3.4.2) | 12207705 | 99 | 99 | 90 | 1e-283 |
| Streptococcus dysgalactiae subsp. equisimilis strain UT_4966_RC | SP_0494 | CTP synthase (EC 6.3.4.2) | 12207705 | 99 | 99 | 90 | 1e-284 |
| Streptococcus dysgalactiae subsp. equisimilis strain UT-5345 | SP_0494 | CTP synthase (EC 6.3.4.2) | 12207705 | 99 | 99 | 90 | 1e-284 |
| Streptococcus dysgalactiae subsp. equisimilis strain UT-5354 | SP_0494 | CTP synthase (EC 6.3.4.2) | 12207705 | 41 | 96 | 96 | 1e-119 |
| Streptococcus dysgalactiae subsp. equisimilis strain UT-SS1069 | SP_0494 | CTP synthase (EC 6.3.4.2) | 12207705 | 99 | 99 | 90 | 1e-284 |
| Streptococcus dysgalactiae subsp. equisimilis strain UT-SS957 | SP_0494 | CTP synthase (EC 6.3.4.2) | 12207705 | 99 | 99 | 90 | 1e-284 |
| Streptococcus dysgalactiae subsp. equisimilis strain WCHSDSE-1 | SP_0494 | CTP synthase (EC 6.3.4.2) | 12207705 | 99 | 99 | 90 | 1e-284 |
| Streptococcus dysgalactiae subsp. equisimilis 167 | SpyM3_0013 | Cationic amino acid transporter - APC Superfamily | 19223485 | 99 | 99 | 83 | 1e-227 |
| Streptococcus dysgalactiae subsp. equisimilis AC-2713 | SpyM3_0013 | Cationic amino acid transporter - APC Superfamily | 19223485 | 99 | 99 | 83 | 1e-227 |
| Streptococcus dysgalactiae subsp. equisimilis AKSDE4288 | SpyM3_0013 | Cationic amino acid transporter - APC Superfamily | 19223485 | 99 | 99 | 83 | 1e-227 |
| Streptococcus dysgalactiae subsp. equisimilis ATCC 12394 | SpyM3_0013 | Cationic amino acid transporter - APC Superfamily | 19223485 | 99 | 99 | 83 | 1e-227 |
| Streptococcus dysgalactiae subsp. equisimilis GGS_124 | SpyM3_0013 | Cationic amino acid transporter - APC Superfamily | 19223485 | 99 | 99 | 83 | 1e-227 |
| Streptococcus dysgalactiae subsp. equisimilis RE378 | SpyM3_0013 | Cationic amino acid transporter - APC Superfamily | 19223485 | 99 | 99 | 83 | 1e-227 |
| Streptococcus dysgalactiae subsp. equisimilis SD SCDR1 | SpyM3_0013 | Cationic amino acid transporter - APC Superfamily | 19223485 | 99 | 99 | 83 | 1e-227 |
| Streptococcus dysgalactiae subsp. equisimilis SK1249 | SpyM3_0013 | Cationic amino acid transporter - APC Superfamily | 19223485 | 99 | 99 | 83 | 1e-227 |
| Streptococcus dysgalactiae subsp. equisimilis SK1250 | SpyM3_0013 | Cationic amino acid transporter - APC Superfamily | 19223485 | 99 | 99 | 83 | 1e-227 |
| Streptococcus dysgalactiae subsp. equisimilis strain ASDSE_96 | SpyM3_0013 | Cationic amino acid transporter - APC Superfamily | 19223485 | 99 | 99 | 83 | 1e-227 |
| Streptococcus dysgalactiae subsp. equisimilis strain ASDSE_99 | SpyM3_0013 | Cationic amino acid transporter - APC Superfamily | 19223485 | 99 | 99 | 82 | 1e-227 |
| Streptococcus dysgalactiae subsp. equisimilis strain C161L1 | SpyM3_0013 | Cationic amino acid transporter - APC Superfamily | 19223485 | 100 | 100 | 82 | 0 |
| Streptococcus dysgalactiae subsp. equisimilis strain KNZ01 | SpyM3_0013 | Cationic amino acid transporter - APC Superfamily | 19223485 | 99 | 99 | 83 | 1e-227 |
| Streptococcus dysgalactiae subsp. equisimilis strain KNZ03 | SpyM3_0013 | Cationic amino acid transporter - APC Superfamily | 19223485 | 99 | 99 | 83 | 1e-227 |
| Streptococcus dysgalactiae subsp. equisimilis strain KNZ04 | SpyM3_0013 | Cationic amino acid transporter - APC Superfamily | 19223485 | 99 | 99 | 83 | 1e-227 |
| Streptococcus dysgalactiae subsp. equisimilis strain KNZ06 | SpyM3_0013 | Cationic amino acid transporter - APC Superfamily | 19223485 | 82 | 97 | 85 | 1e-188 |
| Streptococcus dysgalactiae subsp. equisimilis strain KNZ07 | SpyM3_0013 | Cationic amino acid transporter - APC Superfamily | 19223485 | 99 | 99 | 83 | 1e-227 |
| Streptococcus dysgalactiae subsp. equisimilis strain KNZ10 | SpyM3_0013 | Cationic amino acid transporter - APC Superfamily | 19223485 | 99 | 99 | 83 | 1e-227 |
| Streptococcus dysgalactiae subsp. equisimilis strain KNZ12 | SpyM3_0013 | Cationic amino acid transporter - APC Superfamily | 19223485 | 99 | 99 | 83 | 1e-227 |
| Streptococcus dysgalactiae subsp. equisimilis strain KNZ15 | SpyM3_0013 | Cationic amino acid transporter - APC Superfamily | 19223485 | 99 | 99 | 83 | 1e-227 |
| Streptococcus dysgalactiae subsp. equisimilis strain KNZ16 | SpyM3_0013 | Cationic amino acid transporter - APC Superfamily | 19223485 | 99 | 99 | 83 | 1e-227 |
| Streptococcus dysgalactiae subsp. equisimilis strain NCTC10321 | SpyM3_0013 | Cationic amino acid transporter - APC Superfamily | 19223485 | 99 | 98 | 83 | 1e-228 |
| Streptococcus dysgalactiae subsp. equisimilis strain NCTC11554 | SpyM3_0013 | Cationic amino acid transporter - APC Superfamily | 19223485 | 99 | 99 | 83 | 1e-227 |
| Streptococcus dysgalactiae subsp. equisimilis strain NCTC11555 | SpyM3_0013 | Cationic amino acid transporter - APC Superfamily | 19223485 | 99 | 99 | 83 | 1e-228 |
| Streptococcus dysgalactiae subsp. equisimilis strain NCTC11556 | SpyM3_0013 | Cationic amino acid transporter - APC Superfamily | 19223485 | 99 | 99 | 83 | 1e-227 |
| Streptococcus dysgalactiae subsp. equisimilis strain NCTC11557 | SpyM3_0013 | Cationic amino acid transporter - APC Superfamily | 19223485 | 99 | 99 | 82 | 1e-227 |
| Streptococcus dysgalactiae subsp. equisimilis strain NCTC11564 | SpyM3_0013 | Cationic amino acid transporter - APC Superfamily | 19223485 | 99 | 99 | 83 | 1e-227 |
| Streptococcus dysgalactiae subsp. equisimilis strain NCTC5370 | SpyM3_0013 | Cationic amino acid transporter - APC Superfamily | 19223485 | 99 | 99 | 83 | 1e-227 |
| Streptococcus dysgalactiae subsp. equisimilis strain NCTC5371 | SpyM3_0013 | Cationic amino acid transporter - APC Superfamily | 19223485 | 99 | 99 | 83 | 1e-227 |
| Streptococcus dysgalactiae subsp. equisimilis strain NCTC6179 | SpyM3_0013 | Cationic amino acid transporter - APC Superfamily | 19223485 | 99 | 99 | 83 | 1e-227 |
| Streptococcus dysgalactiae subsp. equisimilis strain NCTC6181 | SpyM3_0013 | Cationic amino acid transporter - APC Superfamily | 19223485 | 99 | 99 | 83 | 1e-227 |
| Streptococcus dysgalactiae subsp. equisimilis strain NCTC6407 | SpyM3_0013 | Cationic amino acid transporter - APC Superfamily | 19223485 | 99 | 98 | 83 | 1e-228 |
| Streptococcus dysgalactiae subsp. equisimilis strain NCTC7136 | SpyM3_0013 | Cationic amino acid transporter - APC Superfamily | 19223485 | 99 | 99 | 83 | 1e-227 |
| Streptococcus dysgalactiae subsp. equisimilis strain NCTC8543 | SpyM3_0013 | Cationic amino acid transporter - APC Superfamily | 19223485 | 99 | 99 | 82 | 1e-227 |
| Streptococcus dysgalactiae subsp. equisimilis strain NCTC8546 | SpyM3_0013 | Cationic amino acid transporter - APC Superfamily | 19223485 | 99 | 99 | 83 | 1e-227 |
| Streptococcus dysgalactiae subsp. equisimilis strain NCTC9413 | SpyM3_0013 | Cationic amino acid transporter - APC Superfamily | 19223485 | 99 | 99 | 83 | 1e-227 |
| Streptococcus dysgalactiae subsp. equisimilis strain NCTC9414 | SpyM3_0013 | Cationic amino acid transporter - APC Superfamily | 19223485 | 99 | 99 | 83 | 1e-227 |
| Streptococcus dysgalactiae subsp. equisimilis strain NCTC9603 | SpyM3_0013 | Cationic amino acid transporter - APC Superfamily | 19223485 | 99 | 99 | 83 | 1e-227 |
| Streptococcus dysgalactiae subsp. equisimilis strain SS1575 | SpyM3_0013 | Cationic amino acid transporter - APC Superfamily | 19223485 | 99 | 99 | 83 | 1e-227 |
| Streptococcus dysgalactiae subsp. equisimilis strain T642 | SpyM3_0013 | Cationic amino acid transporter - APC Superfamily | 19223485 | 100 | 100 | 82 | 0 |
| Streptococcus dysgalactiae subsp. equisimilis strain UT_4031CC | SpyM3_0013 | Cationic amino acid transporter - APC Superfamily | 19223485 | 99 | 99 | 82 | 1e-227 |
| Streptococcus dysgalactiae subsp. equisimilis strain UT_4231_KK | SpyM3_0013 | Cationic amino acid transporter - APC Superfamily | 19223485 | 99 | 99 | 83 | 1e-227 |
| Streptococcus dysgalactiae subsp. equisimilis strain UT_4234_DH | SpyM3_0013 | Cationic amino acid transporter - APC Superfamily | 19223485 | 99 | 99 | 82 | 1e-227 |
| Streptococcus dysgalactiae subsp. equisimilis strain UT_4241_XS | SpyM3_0013 | Cationic amino acid transporter - APC Superfamily | 19223485 | 99 | 99 | 83 | 1e-227 |
| Streptococcus dysgalactiae subsp. equisimilis strain UT_4242_AB | SpyM3_0013 | Cationic amino acid transporter - APC Superfamily | 19223485 | 99 | 99 | 83 | 1e-227 |
| Streptococcus dysgalactiae subsp. equisimilis strain UT_4255RC | SpyM3_0013 | Cationic amino acid transporter - APC Superfamily | 19223485 | 99 | 99 | 82 | 1e-227 |
| Streptococcus dysgalactiae subsp. equisimilis strain UT_4277_BB | SpyM3_0013 | Cationic amino acid transporter - APC Superfamily | 19223485 | 99 | 99 | 82 | 1e-227 |
| Streptococcus dysgalactiae subsp. equisimilis strain UT_4966_RC | SpyM3_0013 | Cationic amino acid transporter - APC Superfamily | 19223485 | 99 | 99 | 83 | 1e-227 |
| Streptococcus dysgalactiae subsp. equisimilis strain UT-5345 | SpyM3_0013 | Cationic amino acid transporter - APC Superfamily | 19223485 | 99 | 99 | 83 | 1e-227 |
| Streptococcus dysgalactiae subsp. equisimilis strain UT-5354 | SpyM3_0013 | Cationic amino acid transporter - APC Superfamily | 19223485 | 99 | 99 | 82 | 1e-227 |
| Streptococcus dysgalactiae subsp. equisimilis strain UT-SS1069 | SpyM3_0013 | Cationic amino acid transporter - APC Superfamily | 19223485 | 99 | 99 | 83 | 1e-227 |
| Streptococcus dysgalactiae subsp. equisimilis strain UT-SS957 | SpyM3_0013 | Cationic amino acid transporter - APC Superfamily | 19223485 | 99 | 99 | 83 | 1e-227 |
| Streptococcus dysgalactiae subsp. equisimilis strain WCHSDSE-1 | SpyM3_0013 | Cationic amino acid transporter - APC Superfamily | 19223485 | 99 | 99 | 83 | 1e-227 |
| Streptococcus dysgalactiae subsp. equisimilis 167 | ccpA | Catabolite control protein A | 20333240 | 100 | 100 | 98 | 1e-185 |
| Streptococcus dysgalactiae subsp. equisimilis AC-2713 | ccpA | Catabolite control protein A | 20333240 | 100 | 100 | 99 | 1e-185 |
| Streptococcus dysgalactiae subsp. equisimilis AKSDE4288 | ccpA | Catabolite control protein A | 20333240 | 100 | 100 | 98 | 1e-185 |
| Streptococcus dysgalactiae subsp. equisimilis ATCC 12394 | ccpA | Catabolite control protein A | 20333240 | 100 | 100 | 98 | 1e-185 |
| Streptococcus dysgalactiae subsp. equisimilis GGS_124 | ccpA | Catabolite control protein A | 20333240 | 100 | 100 | 98 | 1e-185 |
| Streptococcus dysgalactiae subsp. equisimilis RE378 | ccpA | Catabolite control protein A | 20333240 | 100 | 100 | 98 | 1e-185 |
| Streptococcus dysgalactiae subsp. equisimilis SD SCDR1 | ccpA | Catabolite control protein A | 20333240 | 100 | 100 | 98 | 1e-185 |
| Streptococcus dysgalactiae subsp. equisimilis SK1249 | ccpA | Catabolite control protein A | 20333240 | 24 | 98 | 100 | 1e-37 |
| Streptococcus dysgalactiae subsp. equisimilis SK1250 | ccpA | Catabolite control protein A | 20333240 | 100 | 100 | 98 | 1e-185 |
| Streptococcus dysgalactiae subsp. equisimilis strain ASDSE_96 | ccpA | Catabolite control protein A | 20333240 | 100 | 100 | 98 | 1e-185 |
| Streptococcus dysgalactiae subsp. equisimilis strain ASDSE_99 | ccpA | Catabolite control protein A | 20333240 | 100 | 100 | 98 | 1e-185 |
| Streptococcus dysgalactiae subsp. equisimilis strain C161L1 | ccpA | Catabolite control protein A | 20333240 | 100 | 100 | 98 | 0 |
| Streptococcus dysgalactiae subsp. equisimilis strain KNZ01 | ccpA | Catabolite control protein A | 20333240 | 100 | 100 | 98 | 1e-185 |
| Streptococcus dysgalactiae subsp. equisimilis strain KNZ03 | ccpA | Catabolite control protein A | 20333240 | 100 | 100 | 98 | 1e-185 |
| Streptococcus dysgalactiae subsp. equisimilis strain KNZ04 | ccpA | Catabolite control protein A | 20333240 | 100 | 100 | 98 | 1e-185 |
| Streptococcus dysgalactiae subsp. equisimilis strain KNZ06 | ccpA | Catabolite control protein A | 20333240 | 100 | 100 | 98 | 1e-185 |
| Streptococcus dysgalactiae subsp. equisimilis strain KNZ07 | ccpA | Catabolite control protein A | 20333240 | 100 | 100 | 98 | 1e-185 |
| Streptococcus dysgalactiae subsp. equisimilis strain KNZ10 | ccpA | Catabolite control protein A | 20333240 | 100 | 100 | 98 | 1e-185 |
| Streptococcus dysgalactiae subsp. equisimilis strain KNZ12 | ccpA | Catabolite control protein A | 20333240 | 100 | 100 | 98 | 1e-185 |
| Streptococcus dysgalactiae subsp. equisimilis strain KNZ15 | ccpA | Catabolite control protein A | 20333240 | 100 | 100 | 98 | 1e-185 |
| Streptococcus dysgalactiae subsp. equisimilis strain KNZ16 | ccpA | Catabolite control protein A | 20333240 | 100 | 100 | 98 | 1e-185 |
| Streptococcus dysgalactiae subsp. equisimilis strain NCTC10321 | ccpA | Catabolite control protein A | 20333240 | 100 | 100 | 98 | 1e-185 |
| Streptococcus dysgalactiae subsp. equisimilis strain NCTC11554 | ccpA | Catabolite control protein A | 20333240 | 100 | 100 | 98 | 1e-185 |
| Streptococcus dysgalactiae subsp. equisimilis strain NCTC11555 | ccpA | Catabolite control protein A | 20333240 | 100 | 100 | 98 | 1e-185 |
| Streptococcus dysgalactiae subsp. equisimilis strain NCTC11556 | ccpA | Catabolite control protein A | 20333240 | 100 | 100 | 98 | 1e-185 |
| Streptococcus dysgalactiae subsp. equisimilis strain NCTC11557 | ccpA | Catabolite control protein A | 20333240 | 100 | 100 | 98 | 1e-185 |
| Streptococcus dysgalactiae subsp. equisimilis strain NCTC11564 | ccpA | Catabolite control protein A | 20333240 | 100 | 100 | 98 | 1e-185 |
| Streptococcus dysgalactiae subsp. equisimilis strain NCTC5370 | ccpA | Catabolite control protein A | 20333240 | 100 | 100 | 98 | 1e-185 |
| Streptococcus dysgalactiae subsp. equisimilis strain NCTC5371 | ccpA | Catabolite control protein A | 20333240 | 100 | 100 | 98 | 1e-185 |
| Streptococcus dysgalactiae subsp. equisimilis strain NCTC5969 | ccpA | Catabolite control protein A | 20525824 | 100 | 100 | 99 | 1e-184 |
| Streptococcus dysgalactiae subsp. equisimilis strain NCTC6179 | ccpA | Catabolite control protein A | 20333240 | 100 | 100 | 98 | 1e-185 |
| Streptococcus dysgalactiae subsp. equisimilis strain NCTC6181 | ccpA | Catabolite control protein A | 20333240 | 100 | 100 | 98 | 1e-185 |
| Streptococcus dysgalactiae subsp. equisimilis strain NCTC6407 | ccpA | Catabolite control protein A | 20333240 | 100 | 100 | 98 | 1e-185 |
| Streptococcus dysgalactiae subsp. equisimilis strain NCTC7136 | ccpA | Catabolite control protein A | 20333240 | 100 | 100 | 98 | 1e-185 |
| Streptococcus dysgalactiae subsp. equisimilis strain NCTC8543 | ccpA | Catabolite control protein A | 20333240 | 100 | 100 | 98 | 1e-185 |
| Streptococcus dysgalactiae subsp. equisimilis strain NCTC8546 | ccpA | Catabolite control protein A | 20333240 | 100 | 100 | 98 | 1e-185 |
| Streptococcus dysgalactiae subsp. equisimilis strain NCTC9413 | ccpA | Catabolite control protein A | 20333240 | 100 | 100 | 98 | 1e-185 |
| Streptococcus dysgalactiae subsp. equisimilis strain NCTC9414 | ccpA | Catabolite control protein A | 20333240 | 100 | 100 | 98 | 1e-185 |
| Streptococcus dysgalactiae subsp. equisimilis strain NCTC9603 | ccpA | Catabolite control protein A | 20333240 | 100 | 100 | 98 | 1e-185 |
| Streptococcus dysgalactiae subsp. equisimilis strain SS1575 | ccpA | Catabolite control protein A | 20333240 | 100 | 100 | 98 | 1e-185 |
| Streptococcus dysgalactiae subsp. equisimilis strain T642 | ccpA | Catabolite control protein A | 20333240 | 100 | 100 | 98 | 0 |
| Streptococcus dysgalactiae subsp. equisimilis strain UT_4031CC | ccpA | Catabolite control protein A | 20333240 | 100 | 100 | 98 | 1e-185 |
| Streptococcus dysgalactiae subsp. equisimilis strain UT_4231_KK | ccpA | Catabolite control protein A | 20333240 | 100 | 100 | 98 | 1e-185 |
| Streptococcus dysgalactiae subsp. equisimilis strain UT_4234_DH | ccpA | Catabolite control protein A | 20333240 | 100 | 100 | 98 | 1e-185 |
| Streptococcus dysgalactiae subsp. equisimilis strain UT_4241_XS | ccpA | Catabolite control protein A | 20333240 | 100 | 100 | 98 | 1e-185 |
| Streptococcus dysgalactiae subsp. equisimilis strain UT_4242_AB | ccpA | Catabolite control protein A | 20333240 | 100 | 100 | 98 | 1e-185 |
| Streptococcus dysgalactiae subsp. equisimilis strain UT_4255RC | ccpA | Catabolite control protein A | 20333240 | 100 | 100 | 98 | 1e-185 |
| Streptococcus dysgalactiae subsp. equisimilis strain UT_4277_BB | ccpA | Catabolite control protein A | 20333240 | 100 | 100 | 98 | 1e-185 |
| Streptococcus dysgalactiae subsp. equisimilis strain UT_4966_RC | ccpA | Catabolite control protein A | 20333240 | 100 | 100 | 98 | 1e-185 |
| Streptococcus dysgalactiae subsp. equisimilis strain UT-5345 | ccpA | Catabolite control protein A | 20333240 | 100 | 100 | 98 | 1e-185 |
| Streptococcus dysgalactiae subsp. equisimilis strain UT-5354 | ccpA | Catabolite control protein A | 20333240 | 100 | 100 | 98 | 1e-185 |
| Streptococcus dysgalactiae subsp. equisimilis strain UT-SS1069 | ccpA | Catabolite control protein A | 20333240 | 66 | 100 | 98 | 1e-119 |
| Streptococcus dysgalactiae subsp. equisimilis strain UT-SS957 | ccpA | Catabolite control protein A | 20333240 | 100 | 100 | 98 | 1e-185 |
| Streptococcus dysgalactiae subsp. equisimilis strain WCHSDSE-1 | ccpA | Catabolite control protein A | 20333240 | 100 | 100 | 98 | 1e-185 |
| Streptococcus dysgalactiae subsp. equisimilis 167 | scpB | C5a peptidase (EC 3.4.21.-) | 11953377; 9119483 | 65 | 96 | 99 | 0.0 |
| Streptococcus dysgalactiae subsp. equisimilis 167 | scpB | C5a peptidase (EC 3.4.21.-) | 11953377; 9119483 | 27 | 99 | 99 | 1e-179 |
| Streptococcus dysgalactiae subsp. equisimilis 167 | scpB | C5a peptidase (EC 3.4.21.-) | 11953377; 9119483 | 27 | 99 | 99 | 1e-179 |
| Streptococcus dysgalactiae subsp. equisimilis 167 | scpB | C5a peptidase (EC 3.4.21.-) | 11953377; 9119483 | 65 | 96 | 99 | 0.0 |
| Streptococcus dysgalactiae subsp. equisimilis AC-2713 | scpA | C5a peptidase (EC 3.4.21.-) | 11953377; 9119483 | 94 | 92 | 97 | 0.0 |
| Streptococcus dysgalactiae subsp. equisimilis AC-2713 | scpA | C5a peptidase (EC 3.4.21.-) | 11953377; 9119483 | 99 | 99 | 97 | 0.0 |
| Streptococcus dysgalactiae subsp. equisimilis AKSDE4288 | scpA | C5a peptidase (EC 3.4.21.-) | 11953377; 9119483 | 93 | 94 | 97 | 0.0 |
| Streptococcus dysgalactiae subsp. equisimilis AKSDE4288 | scpA | C5a peptidase (EC 3.4.21.-) | 11953377; 9119483 | 100 | 100 | 97 | 0.0 |
| Streptococcus dysgalactiae subsp. equisimilis ATCC 12394 | scpA | C5a peptidase (EC 3.4.21.-) | 11953377; 9119483 | 76 | 95 | 99 | 0.0 |
| Streptococcus dysgalactiae subsp. equisimilis ATCC 12394 | scpA | C5a peptidase (EC 3.4.21.-) | 11953377; 9119483 | 76 | 95 | 99 | 0.0 |
| Streptococcus dysgalactiae subsp. equisimilis ATCC 12394 | scpB | C5a peptidase (EC 3.4.21.-) | 11953377; 9119483 | 17 | 99 | 96 | 1e-111 |
| Streptococcus dysgalactiae subsp. equisimilis ATCC 12394 | scpB | C5a peptidase (EC 3.4.21.-) | 11953377; 9119483 | 17 | 99 | 96 | 1e-111 |
| Streptococcus dysgalactiae subsp. equisimilis GGS_124 | scpB | C5a peptidase (EC 3.4.21.-) | 11953377; 9119483 | 100 | 100 | 99 | 0.0 |
| Streptococcus dysgalactiae subsp. equisimilis GGS_124 | scpB | C5a peptidase (EC 3.4.21.-) | 11953377; 9119483 | 100 | 100 | 99 | 0.0 |
| Streptococcus dysgalactiae subsp. equisimilis RE378 | scpB | C5a peptidase (EC 3.4.21.-) | 11953377; 9119483 | 100 | 100 | 97 | 0.0 |
| Streptococcus dysgalactiae subsp. equisimilis RE378 | scpB | C5a peptidase (EC 3.4.21.-) | 11953377; 9119483 | 93 | 94 | 97 | 0.0 |
| Streptococcus dysgalactiae subsp. equisimilis SD SCDR1 | scpA | C5a peptidase (EC 3.4.21.-) | 11953377; 9119483 | 99 | 99 | 97 | 0.0 |
| Streptococcus dysgalactiae subsp. equisimilis SD SCDR1 | scpA | C5a peptidase (EC 3.4.21.-) | 11953377; 9119483 | 94 | 92 | 97 | 0.0 |
| Streptococcus dysgalactiae subsp. equisimilis SK1249 | scpA | C5a peptidase (EC 3.4.21.-) | 11953377; 9119483 | 100 | 100 | 98 | 0.0 |
| Streptococcus dysgalactiae subsp. equisimilis SK1249 | scpA | C5a peptidase (EC 3.4.21.-) | 11953377; 9119483 | 100 | 100 | 98 | 0.0 |
| Streptococcus dysgalactiae subsp. equisimilis SK1250 | scpA | C5a peptidase (EC 3.4.21.-) | 11953377; 9119483 | 100 | 100 | 97 | 0.0 |
| Streptococcus dysgalactiae subsp. equisimilis SK1250 | scpA | C5a peptidase (EC 3.4.21.-) | 11953377; 9119483 | 93 | 94 | 97 | 0.0 |
| Streptococcus dysgalactiae subsp. equisimilis strain ASDSE_96 | scpA | C5a peptidase (EC 3.4.21.-) | 11953377; 9119483 | 99 | 99 | 98 | 0.0 |
| Streptococcus dysgalactiae subsp. equisimilis strain ASDSE_96 | scpA | C5a peptidase (EC 3.4.21.-) | 11953377; 9119483 | 99 | 99 | 98 | 0.0 |
| Streptococcus dysgalactiae subsp. equisimilis strain ASDSE_99 | scpB | C5a peptidase (EC 3.4.21.-) | 11953377; 9119483 | 100 | 100 | 99 | 0.0 |
| Streptococcus dysgalactiae subsp. equisimilis strain ASDSE_99 | scpB | C5a peptidase (EC 3.4.21.-) | 11953377; 9119483 | 100 | 100 | 99 | 0.0 |
| Streptococcus dysgalactiae subsp. equisimilis strain C161L1 | scpB | C5a peptidase (EC 3.4.21.-) | 11953377; 9119483 | 100 | 100 | 99 | 0 |
| Streptococcus dysgalactiae subsp. equisimilis strain C161L1 | scpB | C5a peptidase (EC 3.4.21.-) | 11953377; 9119483 | 100 | 100 | 99 | 0 |
| Streptococcus dysgalactiae subsp. equisimilis strain KNZ01 | scpA | C5a peptidase (EC 3.4.21.-) | 11953377; 9119483 | 93 | 94 | 97 | 0.0 |
| Streptococcus dysgalactiae subsp. equisimilis strain KNZ01 | scpA | C5a peptidase (EC 3.4.21.-) | 11953377; 9119483 | 93 | 94 | 97 | 0.0 |
| Streptococcus dysgalactiae subsp. equisimilis strain KNZ01 | scpA | C5a peptidase (EC 3.4.21.-) | 11953377; 9119483 | 100 | 100 | 97 | 0.0 |
| Streptococcus dysgalactiae subsp. equisimilis strain KNZ01 | scpA | C5a peptidase (EC 3.4.21.-) | 11953377; 9119483 | 100 | 100 | 97 | 0.0 |
| Streptococcus dysgalactiae subsp. equisimilis strain KNZ03 | scpB | C5a peptidase (EC 3.4.21.-) | 11953377; 9119483 | 100 | 100 | 99 | 0.0 |
| Streptococcus dysgalactiae subsp. equisimilis strain KNZ03 | scpB | C5a peptidase (EC 3.4.21.-) | 11953377; 9119483 | 100 | 100 | 99 | 0.0 |
| Streptococcus dysgalactiae subsp. equisimilis strain KNZ03 | scpB | C5a peptidase (EC 3.4.21.-) | 11953377; 9119483 | 100 | 100 | 99 | 0.0 |
| Streptococcus dysgalactiae subsp. equisimilis strain KNZ03 | scpB | C5a peptidase (EC 3.4.21.-) | 11953377; 9119483 | 100 | 100 | 99 | 0.0 |
| Streptococcus dysgalactiae subsp. equisimilis strain KNZ04 | scpB | C5a peptidase (EC 3.4.21.-) | 11953377; 9119483 | 100 | 100 | 99 | 0.0 |
| Streptococcus dysgalactiae subsp. equisimilis strain KNZ04 | scpB | C5a peptidase (EC 3.4.21.-) | 11953377; 9119483 | 100 | 100 | 99 | 0.0 |
| Streptococcus dysgalactiae subsp. equisimilis strain KNZ06 | scpB | C5a peptidase (EC 3.4.21.-) | 11953377; 9119483 | 90 | 100 | 99 | 0.0 |
| Streptococcus dysgalactiae subsp. equisimilis strain KNZ06 | scpB | C5a peptidase (EC 3.4.21.-) | 11953377; 9119483 | 90 | 100 | 99 | 0.0 |
| Streptococcus dysgalactiae subsp. equisimilis strain KNZ06 | scpB | C5a peptidase (EC 3.4.21.-) | 11953377; 9119483 | 6 | 100 | 100 | 2e-37 |
| Streptococcus dysgalactiae subsp. equisimilis strain KNZ06 | scpB | C5a peptidase (EC 3.4.21.-) | 11953377; 9119483 | 6 | 100 | 100 | 2e-37 |
| Streptococcus dysgalactiae subsp. equisimilis strain KNZ07 | scpA | C5a peptidase (EC 3.4.21.-) | 11953377; 9119483 | 100 | 100 | 97 | 0.0 |
| Streptococcus dysgalactiae subsp. equisimilis strain KNZ07 | scpA | C5a peptidase (EC 3.4.21.-) | 11953377; 9119483 | 93 | 94 | 97 | 0.0 |
| Streptococcus dysgalactiae subsp. equisimilis strain KNZ10 | scpA | C5a peptidase (EC 3.4.21.-) | 11953377; 9119483 | 93 | 94 | 97 | 0.0 |
| Streptococcus dysgalactiae subsp. equisimilis strain KNZ10 | scpA | C5a peptidase (EC 3.4.21.-) | 11953377; 9119483 | 100 | 100 | 97 | 0.0 |
| Streptococcus dysgalactiae subsp. equisimilis strain KNZ12 | scpA | C5a peptidase (EC 3.4.21.-) | 11953377; 9119483 | 100 | 100 | 97 | 0.0 |
| Streptococcus dysgalactiae subsp. equisimilis strain KNZ12 | scpA | C5a peptidase (EC 3.4.21.-) | 11953377; 9119483 | 93 | 94 | 97 | 0.0 |
| Streptococcus dysgalactiae subsp. equisimilis strain KNZ15 | scpA | C5a peptidase (EC 3.4.21.-) | 11953377; 9119483 | 100 | 100 | 98 | 0.0 |
| Streptococcus dysgalactiae subsp. equisimilis strain KNZ15 | scpA | C5a peptidase (EC 3.4.21.-) | 11953377; 9119483 | 93 | 94 | 98 | 0.0 |
| Streptococcus dysgalactiae subsp. equisimilis strain KNZ16 | scpB | C5a peptidase (EC 3.4.21.-) | 11953377; 9119483 | 100 | 100 | 98 | 0.0 |
| Streptococcus dysgalactiae subsp. equisimilis strain KNZ16 | scpB | C5a peptidase (EC 3.4.21.-) | 11953377; 9119483 | 100 | 100 | 98 | 0.0 |
| Streptococcus dysgalactiae subsp. equisimilis strain NCTC11554 | scpB | C5a peptidase (EC 3.4.21.-) | 11953377; 9119483 | 100 | 100 | 98 | 0.0 |
| Streptococcus dysgalactiae subsp. equisimilis strain NCTC11554 | scpB | C5a peptidase (EC 3.4.21.-) | 11953377; 9119483 | 100 | 100 | 98 | 0.0 |
| Streptococcus dysgalactiae subsp. equisimilis strain NCTC11555 | scpA | C5a peptidase (EC 3.4.21.-) | 11953377; 9119483 | 100 | 100 | 98 | 0.0 |
| Streptococcus dysgalactiae subsp. equisimilis strain NCTC11555 | scpB | C5a peptidase (EC 3.4.21.-) | 11953377; 9119483 | 100 | 100 | 98 | 0.0 |
| Streptococcus dysgalactiae subsp. equisimilis strain NCTC11556 | scpA | C5a peptidase (EC 3.4.21.-) | 11953377; 9119483 | 94 | 92 | 97 | 0.0 |
| Streptococcus dysgalactiae subsp. equisimilis strain NCTC11556 | scpA | C5a peptidase (EC 3.4.21.-) | 11953377; 9119483 | 99 | 99 | 97 | 0.0 |
| Streptococcus dysgalactiae subsp. equisimilis strain NCTC11557 | scpA | C5a peptidase (EC 3.4.21.-) | 11953377; 9119483 | 100 | 100 | 97 | 0.0 |
| Streptococcus dysgalactiae subsp. equisimilis strain NCTC11557 | scpA | C5a peptidase (EC 3.4.21.-) | 11953377; 9119483 | 93 | 94 | 97 | 0.0 |
| Streptococcus dysgalactiae subsp. equisimilis strain NCTC11564 | scpA | C5a peptidase (EC 3.4.21.-) | 11953377; 9119483 | 99 | 99 | 97 | 0.0 |
| Streptococcus dysgalactiae subsp. equisimilis strain NCTC11564 | scpA | C5a peptidase (EC 3.4.21.-) | 11953377; 9119483 | 99 | 99 | 97 | 0.0 |
| Streptococcus dysgalactiae subsp. equisimilis strain NCTC5370 | scpA | C5a peptidase (EC 3.4.21.-) | 11953377; 9119483 | 93 | 94 | 97 | 0.0 |
| Streptococcus dysgalactiae subsp. equisimilis strain NCTC5370 | scpA | C5a peptidase (EC 3.4.21.-) | 11953377; 9119483 | 100 | 100 | 97 | 0.0 |
| Streptococcus dysgalactiae subsp. equisimilis strain NCTC5371 | scpA | C5a peptidase (EC 3.4.21.-) | 11953377; 9119483 | 93 | 94 | 98 | 0.0 |
| Streptococcus dysgalactiae subsp. equisimilis strain NCTC5371 | scpA | C5a peptidase (EC 3.4.21.-) | 11953377; 9119483 | 100 | 100 | 98 | 0.0 |
| Streptococcus dysgalactiae subsp. equisimilis strain NCTC7136 | scpA | C5a peptidase (EC 3.4.21.-) | 11953377; 9119483 | 99 | 99 | 98 | 0.0 |
| Streptococcus dysgalactiae subsp. equisimilis strain NCTC7136 | scpA | C5a peptidase (EC 3.4.21.-) | 11953377; 9119483 | 99 | 99 | 98 | 0.0 |
| Streptococcus dysgalactiae subsp. equisimilis strain NCTC8543 | scpA | C5a peptidase (EC 3.4.21.-) | 11953377; 9119483 | 99 | 99 | 97 | 0.0 |
| Streptococcus dysgalactiae subsp. equisimilis strain NCTC8543 | scpA | C5a peptidase (EC 3.4.21.-) | 11953377; 9119483 | 94 | 92 | 97 | 0.0 |
| Streptococcus dysgalactiae subsp. equisimilis strain NCTC8546 | scpA | C5a peptidase (EC 3.4.21.-) | 11953377; 9119483 | 93 | 94 | 97 | 0.0 |
| Streptococcus dysgalactiae subsp. equisimilis strain NCTC8546 | scpA | C5a peptidase (EC 3.4.21.-) | 11953377; 9119483 | 100 | 100 | 97 | 0.0 |
| Streptococcus dysgalactiae subsp. equisimilis strain NCTC9413 | scpA | C5a peptidase (EC 3.4.21.-) | 11953377; 9119483 | 99 | 99 | 97 | 0.0 |
| Streptococcus dysgalactiae subsp. equisimilis strain NCTC9413 | scpA | C5a peptidase (EC 3.4.21.-) | 11953377; 9119483 | 99 | 99 | 97 | 0.0 |
| Streptococcus dysgalactiae subsp. equisimilis strain NCTC9414 | scpA | C5a peptidase (EC 3.4.21.-) | 11953377; 9119483 | 100 | 100 | 98 | 0.0 |
| Streptococcus dysgalactiae subsp. equisimilis strain NCTC9414 | scpA | C5a peptidase (EC 3.4.21.-) | 11953377; 9119483 | 93 | 94 | 98 | 0.0 |
| Streptococcus dysgalactiae subsp. equisimilis strain NCTC9603 | scpB | C5a peptidase (EC 3.4.21.-) | 11953377; 9119483 | 100 | 100 | 99 | 0.0 |
| Streptococcus dysgalactiae subsp. equisimilis strain NCTC9603 | scpB | C5a peptidase (EC 3.4.21.-) | 11953377; 9119483 | 100 | 100 | 99 | 0.0 |
| Streptococcus dysgalactiae subsp. equisimilis strain SS1575 | scpB | C5a peptidase (EC 3.4.21.-) | 11953377; 9119483 | 100 | 100 | 99 | 0.0 |
| Streptococcus dysgalactiae subsp. equisimilis strain SS1575 | scpB | C5a peptidase (EC 3.4.21.-) | 11953377; 9119483 | 100 | 100 | 99 | 0.0 |
| Streptococcus dysgalactiae subsp. equisimilis strain T642 | scpA | C5a peptidase (EC 3.4.21.-) | 11953377; 9119483 | 100 | 100 | 97 | 0.0 |
| Streptococcus dysgalactiae subsp. equisimilis strain T642 | scpA | C5a peptidase (EC 3.4.21.-) | 11953377; 9119483 | 100 | 100 | 97 | 0.0 |
| Streptococcus dysgalactiae subsp. equisimilis strain UT_4031CC | scpA | C5a peptidase (EC 3.4.21.-) | 11953377; 9119483 | 99 | 99 | 97 | 0.0 |
| Streptococcus dysgalactiae subsp. equisimilis strain UT_4031CC | scpA | C5a peptidase (EC 3.4.21.-) | 11953377; 9119483 | 99 | 99 | 97 | 0.0 |
| Streptococcus dysgalactiae subsp. equisimilis strain UT_4231_KK | scpA | C5a peptidase (EC 3.4.21.-) | 11953377; 9119483 | 100 | 100 | 99 | 0.0 |
| Streptococcus dysgalactiae subsp. equisimilis strain UT_4231_KK | scpA | C5a peptidase (EC 3.4.21.-) | 11953377; 9119483 | 100 | 100 | 99 | 0.0 |
| Streptococcus dysgalactiae subsp. equisimilis strain UT_4234_DH | scpA | C5a peptidase (EC 3.4.21.-) | 11953377; 9119483 | 93 | 94 | 97 | 0.0 |
| Streptococcus dysgalactiae subsp. equisimilis strain UT_4234_DH | scpA | C5a peptidase (EC 3.4.21.-) | 11953377; 9119483 | 100 | 100 | 97 | 0.0 |
| Streptococcus dysgalactiae subsp. equisimilis strain UT_4241_XS | scpA | C5a peptidase (EC 3.4.21.-) | 11953377; 9119483 | 100 | 100 | 99 | 0.0 |
| Streptococcus dysgalactiae subsp. equisimilis strain UT_4241_XS | scpA | C5a peptidase (EC 3.4.21.-) | 11953377; 9119483 | 100 | 100 | 99 | 0.0 |
| Streptococcus dysgalactiae subsp. equisimilis strain UT_4242_AB | scpA | C5a peptidase (EC 3.4.21.-) | 11953377; 9119483 | 94 | 92 | 97 | 0.0 |
| Streptococcus dysgalactiae subsp. equisimilis strain UT_4242_AB | scpA | C5a peptidase (EC 3.4.21.-) | 11953377; 9119483 | 99 | 99 | 97 | 0.0 |
| Streptococcus dysgalactiae subsp. equisimilis strain UT_4255RC | scpA | C5a peptidase (EC 3.4.21.-) | 11953377; 9119483 | 100 | 100 | 97 | 0.0 |
| Streptococcus dysgalactiae subsp. equisimilis strain UT_4255RC | scpA | C5a peptidase (EC 3.4.21.-) | 11953377; 9119483 | 93 | 94 | 97 | 0.0 |
| Streptococcus dysgalactiae subsp. equisimilis strain UT_4277_BB | scpA | C5a peptidase (EC 3.4.21.-) | 11953377; 9119483 | 99 | 99 | 98 | 0.0 |
| Streptococcus dysgalactiae subsp. equisimilis strain UT_4277_BB | scpA | C5a peptidase (EC 3.4.21.-) | 11953377; 9119483 | 99 | 99 | 98 | 0.0 |
| Streptococcus dysgalactiae subsp. equisimilis strain UT_4966_RC | scpA | C5a peptidase (EC 3.4.21.-) | 11953377; 9119483 | 100 | 100 | 99 | 0.0 |
| Streptococcus dysgalactiae subsp. equisimilis strain UT_4966_RC | scpA | C5a peptidase (EC 3.4.21.-) | 11953377; 9119483 | 100 | 100 | 99 | 0.0 |
| Streptococcus dysgalactiae subsp. equisimilis strain UT-5345 | scpA | C5a peptidase (EC 3.4.21.-) | 11953377; 9119483 | 100 | 100 | 97 | 0.0 |
| Streptococcus dysgalactiae subsp. equisimilis strain UT-5345 | scpA | C5a peptidase (EC 3.4.21.-) | 11953377; 9119483 | 93 | 91 | 99 | 0.0 |
| Streptococcus dysgalactiae subsp. equisimilis strain UT-5354 | scpA | C5a peptidase (EC 3.4.21.-) | 11953377; 9119483 | 92 | 93 | 99 | 0.0 |
| Streptococcus dysgalactiae subsp. equisimilis strain UT-5354 | scpA | C5a peptidase (EC 3.4.21.-) | 11953377; 9119483 | 92 | 93 | 99 | 0.0 |
| Streptococcus dysgalactiae subsp. equisimilis strain UT-SS1069 | scpA | C5a peptidase (EC 3.4.21.-) | 11953377; 9119483 | 47 | 100 | 99 | 0.0 |
| Streptococcus dysgalactiae subsp. equisimilis strain UT-SS1069 | scpA | C5a peptidase (EC 3.4.21.-) | 11953377; 9119483 | 52 | 100 | 99 | 0.0 |
| Streptococcus dysgalactiae subsp. equisimilis strain UT-SS1069 | scpA | C5a peptidase (EC 3.4.21.-) | 11953377; 9119483 | 52 | 100 | 99 | 0.0 |
| Streptococcus dysgalactiae subsp. equisimilis strain UT-SS1069 | scpA | C5a peptidase (EC 3.4.21.-) | 11953377; 9119483 | 47 | 100 | 99 | 0.0 |
| Streptococcus dysgalactiae subsp. equisimilis strain UT-SS957 | scpA | C5a peptidase (EC 3.4.21.-) | 11953377; 9119483 | 100 | 100 | 97 | 0.0 |
| Streptococcus dysgalactiae subsp. equisimilis strain UT-SS957 | scpA | C5a peptidase (EC 3.4.21.-) | 11953377; 9119483 | 93 | 94 | 97 | 0.0 |
| Streptococcus dysgalactiae subsp. equisimilis strain WCHSDSE-1 | scpA | C5a peptidase (EC 3.4.21.-) | 11953377; 9119483 | 100 | 100 | 99 | 0.0 |
| Streptococcus dysgalactiae subsp. equisimilis strain WCHSDSE-1 | scpA | C5a peptidase (EC 3.4.21.-) | 11953377; 9119483 | 100 | 100 | 99 | 0.0 |
| Streptococcus dysgalactiae subsp. equisimilis 167 | bcaT | Branched-chain amino acid aminotransferase (EC 2.6.1.42) | 12207705 | 100 | 99 | 82 | 1e-169 |
| Streptococcus dysgalactiae subsp. equisimilis AC-2713 | bcaT | Branched-chain amino acid aminotransferase (EC 2.6.1.42) | 12207705 | 100 | 99 | 82 | 1e-169 |
| Streptococcus dysgalactiae subsp. equisimilis AKSDE4288 | SP_0856 | Branched-chain amino acid aminotransferase (EC 2.6.1.42) | 12207705 | 100 | 100 | 82 | 1e-169 |
| Streptococcus dysgalactiae subsp. equisimilis ATCC 12394 | SP_0856 | Branched-chain amino acid aminotransferase (EC 2.6.1.42) | 12207705 | 100 | 99 | 82 | 1e-169 |
| Streptococcus dysgalactiae subsp. equisimilis GGS_124 | bcaT | Branched-chain amino acid aminotransferase (EC 2.6.1.42) | 12207705 | 100 | 99 | 82 | 1e-169 |
| Streptococcus dysgalactiae subsp. equisimilis RE378 | bcaT | Branched-chain amino acid aminotransferase (EC 2.6.1.42) | 12207705 | 100 | 99 | 82 | 1e-169 |
| Streptococcus dysgalactiae subsp. equisimilis SD SCDR1 | SP_0856 | Branched-chain amino acid aminotransferase (EC 2.6.1.42) | 12207705 | 100 | 100 | 81 | 1e-168 |
| Streptococcus dysgalactiae subsp. equisimilis SK1249 | ilvE | Branched-chain amino acid aminotransferase (EC 2.6.1.42) | 12207705 | 100 | 99 | 82 | 1e-169 |
| Streptococcus dysgalactiae subsp. equisimilis SK1250 | ilvE | Branched-chain amino acid aminotransferase (EC 2.6.1.42) | 12207705 | 100 | 99 | 82 | 1e-169 |
| Streptococcus dysgalactiae subsp. equisimilis strain ASDSE_96 | SP_0856 | Branched-chain amino acid aminotransferase (EC 2.6.1.42) | 12207705 | 100 | 100 | 82 | 1e-169 |
| Streptococcus dysgalactiae subsp. equisimilis strain ASDSE_99 | SP_0856 | Branched-chain amino acid aminotransferase (EC 2.6.1.42) | 12207705 | 100 | 100 | 82 | 1e-169 |
| Streptococcus dysgalactiae subsp. equisimilis strain C161L1 | SP_0856 | Branched-chain amino acid aminotransferase (EC 2.6.1.42) | 12207705 | 100 | 100 | 82 | 0 |
| Streptococcus dysgalactiae subsp. equisimilis strain KNZ01 | SP_0856 | Branched-chain amino acid aminotransferase (EC 2.6.1.42) | 12207705 | 100 | 100 | 82 | 1e-169 |
| Streptococcus dysgalactiae subsp. equisimilis strain KNZ03 | SP_0856 | Branched-chain amino acid aminotransferase (EC 2.6.1.42) | 12207705 | 100 | 100 | 82 | 1e-169 |
| Streptococcus dysgalactiae subsp. equisimilis strain KNZ04 | SP_0856 | Branched-chain amino acid aminotransferase (EC 2.6.1.42) | 12207705 | 100 | 100 | 82 | 1e-169 |
| Streptococcus dysgalactiae subsp. equisimilis strain KNZ06 | SP_0856 | Branched-chain amino acid aminotransferase (EC 2.6.1.42) | 12207705 | 100 | 100 | 82 | 1e-169 |
| Streptococcus dysgalactiae subsp. equisimilis strain KNZ07 | SP_0856 | Branched-chain amino acid aminotransferase (EC 2.6.1.42) | 12207705 | 100 | 100 | 82 | 1e-169 |
| Streptococcus dysgalactiae subsp. equisimilis strain KNZ10 | SP_0856 | Branched-chain amino acid aminotransferase (EC 2.6.1.42) | 12207705 | 100 | 100 | 82 | 1e-169 |
| Streptococcus dysgalactiae subsp. equisimilis strain KNZ12 | SP_0856 | Branched-chain amino acid aminotransferase (EC 2.6.1.42) | 12207705 | 100 | 100 | 82 | 1e-169 |
| Streptococcus dysgalactiae subsp. equisimilis strain KNZ15 | SP_0856 | Branched-chain amino acid aminotransferase (EC 2.6.1.42) | 12207705 | 100 | 100 | 82 | 1e-169 |
| Streptococcus dysgalactiae subsp. equisimilis strain KNZ16 | SP_0856 | Branched-chain amino acid aminotransferase (EC 2.6.1.42) | 12207705 | 100 | 100 | 82 | 1e-169 |
| Streptococcus dysgalactiae subsp. equisimilis strain NCTC10321 | SP_0856 | Branched-chain amino acid aminotransferase (EC 2.6.1.42) | 12207705 | 100 | 100 | 82 | 1e-169 |
| Streptococcus dysgalactiae subsp. equisimilis strain NCTC11554 | SP_0856 | Branched-chain amino acid aminotransferase (EC 2.6.1.42) | 12207705 | 100 | 100 | 82 | 1e-169 |
| Streptococcus dysgalactiae subsp. equisimilis strain NCTC11555 | SP_0856 | Branched-chain amino acid aminotransferase (EC 2.6.1.42) | 12207705 | 100 | 100 | 82 | 1e-169 |
| Streptococcus dysgalactiae subsp. equisimilis strain NCTC11556 | SP_0856 | Branched-chain amino acid aminotransferase (EC 2.6.1.42) | 12207705 | 100 | 100 | 82 | 1e-169 |
| Streptococcus dysgalactiae subsp. equisimilis strain NCTC11557 | SP_0856 | Branched-chain amino acid aminotransferase (EC 2.6.1.42) | 12207705 | 100 | 100 | 82 | 1e-169 |
| Streptococcus dysgalactiae subsp. equisimilis strain NCTC11564 | SP_0856 | Branched-chain amino acid aminotransferase (EC 2.6.1.42) | 12207705 | 100 | 100 | 81 | 1e-168 |
| Streptococcus dysgalactiae subsp. equisimilis strain NCTC5370 | SP_0856 | Branched-chain amino acid aminotransferase (EC 2.6.1.42) | 12207705 | 100 | 100 | 82 | 1e-169 |
| Streptococcus dysgalactiae subsp. equisimilis strain NCTC5371 | SP_0856 | Branched-chain amino acid aminotransferase (EC 2.6.1.42) | 12207705 | 100 | 100 | 82 | 1e-169 |
| Streptococcus dysgalactiae subsp. equisimilis strain NCTC6179 | SP_0856 | Branched-chain amino acid aminotransferase (EC 2.6.1.42) | 12207705 | 100 | 100 | 82 | 1e-169 |
| Streptococcus dysgalactiae subsp. equisimilis strain NCTC6181 | SP_0856 | Branched-chain amino acid aminotransferase (EC 2.6.1.42) | 12207705 | 100 | 100 | 82 | 1e-169 |
| Streptococcus dysgalactiae subsp. equisimilis strain NCTC6407 | SP_0856 | Branched-chain amino acid aminotransferase (EC 2.6.1.42) | 12207705 | 100 | 100 | 82 | 1e-169 |
| Streptococcus dysgalactiae subsp. equisimilis strain NCTC7136 | SP_0856 | Branched-chain amino acid aminotransferase (EC 2.6.1.42) | 12207705 | 100 | 100 | 82 | 1e-169 |
| Streptococcus dysgalactiae subsp. equisimilis strain NCTC8543 | SP_0856 | Branched-chain amino acid aminotransferase (EC 2.6.1.42) | 12207705 | 100 | 100 | 82 | 1e-169 |
| Streptococcus dysgalactiae subsp. equisimilis strain NCTC8546 | SP_0856 | Branched-chain amino acid aminotransferase (EC 2.6.1.42) | 12207705 | 100 | 100 | 82 | 1e-169 |
| Streptococcus dysgalactiae subsp. equisimilis strain NCTC9413 | SP_0856 | Branched-chain amino acid aminotransferase (EC 2.6.1.42) | 12207705 | 100 | 100 | 82 | 1e-169 |
| Streptococcus dysgalactiae subsp. equisimilis strain NCTC9414 | SP_0856 | Branched-chain amino acid aminotransferase (EC 2.6.1.42) | 12207705 | 100 | 100 | 82 | 1e-169 |
| Streptococcus dysgalactiae subsp. equisimilis strain NCTC9603 | SP_0856 | Branched-chain amino acid aminotransferase (EC 2.6.1.42) | 12207705 | 100 | 100 | 82 | 1e-169 |
| Streptococcus dysgalactiae subsp. equisimilis strain SS1575 | SP_0856 | Branched-chain amino acid aminotransferase (EC 2.6.1.42) | 12207705 | 100 | 100 | 81 | 1e-168 |
| Streptococcus dysgalactiae subsp. equisimilis strain T642 | SP_0856 | Branched-chain amino acid aminotransferase (EC 2.6.1.42) | 12207705 | 100 | 100 | 82 | 0 |
| Streptococcus dysgalactiae subsp. equisimilis strain UT_4031CC | SP_0856 | Branched-chain amino acid aminotransferase (EC 2.6.1.42) | 12207705 | 100 | 100 | 82 | 1e-169 |
| Streptococcus dysgalactiae subsp. equisimilis strain UT_4231_KK | SP_0856 | Branched-chain amino acid aminotransferase (EC 2.6.1.42) | 12207705 | 100 | 100 | 82 | 1e-169 |
| Streptococcus dysgalactiae subsp. equisimilis strain UT_4234_DH | SP_0856 | Branched-chain amino acid aminotransferase (EC 2.6.1.42) | 12207705 | 100 | 100 | 82 | 1e-169 |
| Streptococcus dysgalactiae subsp. equisimilis strain UT_4241_XS | SP_0856 | Branched-chain amino acid aminotransferase (EC 2.6.1.42) | 12207705 | 100 | 100 | 82 | 1e-169 |
| Streptococcus dysgalactiae subsp. equisimilis strain UT_4242_AB | SP_0856 | Branched-chain amino acid aminotransferase (EC 2.6.1.42) | 12207705 | 100 | 100 | 82 | 1e-169 |
| Streptococcus dysgalactiae subsp. equisimilis strain UT_4255RC | SP_0856 | Branched-chain amino acid aminotransferase (EC 2.6.1.42) | 12207705 | 100 | 100 | 82 | 1e-169 |
| Streptococcus dysgalactiae subsp. equisimilis strain UT_4277_BB | SP_0856 | Branched-chain amino acid aminotransferase (EC 2.6.1.42) | 12207705 | 100 | 100 | 82 | 1e-169 |
| Streptococcus dysgalactiae subsp. equisimilis strain UT_4966_RC | SP_0856 | Branched-chain amino acid aminotransferase (EC 2.6.1.42) | 12207705 | 100 | 100 | 82 | 1e-169 |
| Streptococcus dysgalactiae subsp. equisimilis strain UT-5345 | SP_0856 | Branched-chain amino acid aminotransferase (EC 2.6.1.42) | 12207705 | 100 | 100 | 82 | 1e-169 |
| Streptococcus dysgalactiae subsp. equisimilis strain UT-5354 | SP_0856 | Branched-chain amino acid aminotransferase (EC 2.6.1.42) | 12207705 | 100 | 100 | 82 | 1e-169 |
| Streptococcus dysgalactiae subsp. equisimilis strain UT-SS1069 | SP_0856 | Branched-chain amino acid aminotransferase (EC 2.6.1.42) | 12207705 | 100 | 100 | 82 | 1e-169 |
| Streptococcus dysgalactiae subsp. equisimilis strain UT-SS957 | SP_0856 | Branched-chain amino acid aminotransferase (EC 2.6.1.42) | 12207705 | 100 | 100 | 82 | 1e-169 |
| Streptococcus dysgalactiae subsp. equisimilis strain WCHSDSE-1 | SP_0856 | Branched-chain amino acid aminotransferase (EC 2.6.1.42) | 12207705 | 100 | 100 | 82 | 1e-169 |
| Streptococcus dysgalactiae subsp. equisimilis 167 | clpP | ATP-dependent Clp protease proteolytic subunit ClpP (EC 3.4.21.92) | 15664911 | 99 | 99 | 90 | 3e-97 |
| Streptococcus dysgalactiae subsp. equisimilis AC-2713 | clpP | ATP-dependent Clp protease proteolytic subunit ClpP (EC 3.4.21.92) | 15664911 | 99 | 99 | 90 | 3e-97 |
| Streptococcus dysgalactiae subsp. equisimilis AKSDE4288 | clpP | ATP-dependent Clp protease proteolytic subunit ClpP (EC 3.4.21.92) | 15664911 | 99 | 99 | 90 | 3e-97 |
| Streptococcus dysgalactiae subsp. equisimilis ATCC 12394 | clpP | ATP-dependent Clp protease proteolytic subunit ClpP (EC 3.4.21.92) | 15664911 | 99 | 99 | 90 | 3e-97 |
| Streptococcus dysgalactiae subsp. equisimilis GGS_124 | clpP | ATP-dependent Clp protease proteolytic subunit ClpP (EC 3.4.21.92) | 15664911 | 99 | 99 | 90 | 3e-97 |
| Streptococcus dysgalactiae subsp. equisimilis RE378 | clpP | ATP-dependent Clp protease proteolytic subunit ClpP (EC 3.4.21.92) | 15664911 | 99 | 99 | 90 | 3e-97 |
| Streptococcus dysgalactiae subsp. equisimilis SD SCDR1 | clpP | ATP-dependent Clp protease proteolytic subunit ClpP (EC 3.4.21.92) | 15664911 | 99 | 99 | 90 | 3e-97 |
| Streptococcus dysgalactiae subsp. equisimilis SK1249 | clpP | ATP-dependent Clp protease proteolytic subunit ClpP (EC 3.4.21.92) | 15664911 | 99 | 99 | 90 | 3e-97 |
| Streptococcus dysgalactiae subsp. equisimilis SK1250 | clpP | ATP-dependent Clp protease proteolytic subunit ClpP (EC 3.4.21.92) | 15664911 | 99 | 99 | 90 | 3e-97 |
| Streptococcus dysgalactiae subsp. equisimilis strain ASDSE_96 | clpP | ATP-dependent Clp protease proteolytic subunit ClpP (EC 3.4.21.92) | 15664911 | 99 | 99 | 90 | 3e-97 |
| Streptococcus dysgalactiae subsp. equisimilis strain ASDSE_99 | clpP | ATP-dependent Clp protease proteolytic subunit ClpP (EC 3.4.21.92) | 15664911 | 99 | 99 | 90 | 3e-97 |
| Streptococcus dysgalactiae subsp. equisimilis strain C161L1 | clpP | ATP-dependent Clp protease proteolytic subunit ClpP (EC 3.4.21.92) | 15664911 | 99 | 99 | 90 | 0 |
| Streptococcus dysgalactiae subsp. equisimilis strain KNZ01 | clpP | ATP-dependent Clp protease proteolytic subunit ClpP (EC 3.4.21.92) | 15664911 | 99 | 99 | 90 | 3e-97 |
| Streptococcus dysgalactiae subsp. equisimilis strain KNZ03 | clpP | ATP-dependent Clp protease proteolytic subunit ClpP (EC 3.4.21.92) | 15664911 | 99 | 99 | 90 | 3e-97 |
| Streptococcus dysgalactiae subsp. equisimilis strain KNZ04 | clpP | ATP-dependent Clp protease proteolytic subunit ClpP (EC 3.4.21.92) | 15664911 | 99 | 99 | 90 | 3e-97 |
| Streptococcus dysgalactiae subsp. equisimilis strain KNZ06 | clpP | ATP-dependent Clp protease proteolytic subunit ClpP (EC 3.4.21.92) | 15664911 | 99 | 99 | 90 | 3e-97 |
| Streptococcus dysgalactiae subsp. equisimilis strain KNZ07 | clpP | ATP-dependent Clp protease proteolytic subunit ClpP (EC 3.4.21.92) | 15664911 | 99 | 99 | 90 | 3e-97 |
| Streptococcus dysgalactiae subsp. equisimilis strain KNZ10 | clpP | ATP-dependent Clp protease proteolytic subunit ClpP (EC 3.4.21.92) | 15664911 | 99 | 99 | 90 | 3e-97 |
| Streptococcus dysgalactiae subsp. equisimilis strain KNZ12 | clpP | ATP-dependent Clp protease proteolytic subunit ClpP (EC 3.4.21.92) | 15664911 | 99 | 99 | 90 | 3e-97 |
| Streptococcus dysgalactiae subsp. equisimilis strain KNZ15 | clpP | ATP-dependent Clp protease proteolytic subunit ClpP (EC 3.4.21.92) | 15664911 | 99 | 99 | 90 | 3e-97 |
| Streptococcus dysgalactiae subsp. equisimilis strain KNZ16 | clpP | ATP-dependent Clp protease proteolytic subunit ClpP (EC 3.4.21.92) | 15664911 | 99 | 99 | 90 | 3e-97 |
| Streptococcus dysgalactiae subsp. equisimilis strain NCTC10321 | clpP | ATP-dependent Clp protease proteolytic subunit ClpP (EC 3.4.21.92) | 15664911 | 99 | 99 | 90 | 3e-97 |
| Streptococcus dysgalactiae subsp. equisimilis strain NCTC11554 | clpP | ATP-dependent Clp protease proteolytic subunit ClpP (EC 3.4.21.92) | 15664911 | 99 | 99 | 90 | 3e-97 |
| Streptococcus dysgalactiae subsp. equisimilis strain NCTC11555 | clpP | ATP-dependent Clp protease proteolytic subunit ClpP (EC 3.4.21.92) | 15664911 | 99 | 99 | 90 | 3e-97 |
| Streptococcus dysgalactiae subsp. equisimilis strain NCTC11556 | clpP | ATP-dependent Clp protease proteolytic subunit ClpP (EC 3.4.21.92) | 15664911 | 99 | 99 | 90 | 3e-97 |
| Streptococcus dysgalactiae subsp. equisimilis strain NCTC11557 | clpP | ATP-dependent Clp protease proteolytic subunit ClpP (EC 3.4.21.92) | 15664911 | 99 | 99 | 90 | 3e-97 |
| Streptococcus dysgalactiae subsp. equisimilis strain NCTC11564 | clpP | ATP-dependent Clp protease proteolytic subunit ClpP (EC 3.4.21.92) | 15664911 | 99 | 99 | 90 | 3e-97 |
| Streptococcus dysgalactiae subsp. equisimilis strain NCTC5370 | clpP | ATP-dependent Clp protease proteolytic subunit ClpP (EC 3.4.21.92) | 15664911 | 99 | 99 | 90 | 3e-97 |
| Streptococcus dysgalactiae subsp. equisimilis strain NCTC5371 | clpP | ATP-dependent Clp protease proteolytic subunit ClpP (EC 3.4.21.92) | 15664911 | 99 | 99 | 90 | 3e-97 |
| Streptococcus dysgalactiae subsp. equisimilis strain NCTC5969 | clpP | ATP-dependent Clp protease proteolytic subunit ClpP (EC 3.4.21.92) | 10760131 | 94 | 95 | 80 | 4e-82 |
| Streptococcus dysgalactiae subsp. equisimilis strain NCTC6179 | clpP | ATP-dependent Clp protease proteolytic subunit ClpP (EC 3.4.21.92) | 15664911 | 99 | 99 | 90 | 3e-97 |
| Streptococcus dysgalactiae subsp. equisimilis strain NCTC6181 | clpP | ATP-dependent Clp protease proteolytic subunit ClpP (EC 3.4.21.92) | 15664911 | 99 | 99 | 90 | 3e-97 |
| Streptococcus dysgalactiae subsp. equisimilis strain NCTC6407 | clpP | ATP-dependent Clp protease proteolytic subunit ClpP (EC 3.4.21.92) | 15664911 | 99 | 99 | 90 | 3e-97 |
| Streptococcus dysgalactiae subsp. equisimilis strain NCTC7136 | clpP | ATP-dependent Clp protease proteolytic subunit ClpP (EC 3.4.21.92) | 15664911 | 99 | 99 | 90 | 3e-97 |
| Streptococcus dysgalactiae subsp. equisimilis strain NCTC8543 | clpP | ATP-dependent Clp protease proteolytic subunit ClpP (EC 3.4.21.92) | 15664911 | 99 | 99 | 90 | 3e-97 |
| Streptococcus dysgalactiae subsp. equisimilis strain NCTC8546 | clpP | ATP-dependent Clp protease proteolytic subunit ClpP (EC 3.4.21.92) | 15664911 | 99 | 99 | 90 | 3e-97 |
| Streptococcus dysgalactiae subsp. equisimilis strain NCTC9413 | clpP | ATP-dependent Clp protease proteolytic subunit ClpP (EC 3.4.21.92) | 15664911 | 99 | 99 | 90 | 3e-97 |
| Streptococcus dysgalactiae subsp. equisimilis strain NCTC9414 | clpP | ATP-dependent Clp protease proteolytic subunit ClpP (EC 3.4.21.92) | 15664911 | 99 | 99 | 90 | 3e-97 |
| Streptococcus dysgalactiae subsp. equisimilis strain NCTC9603 | clpP | ATP-dependent Clp protease proteolytic subunit ClpP (EC 3.4.21.92) | 15664911 | 99 | 99 | 90 | 3e-97 |
| Streptococcus dysgalactiae subsp. equisimilis strain SS1575 | clpP | ATP-dependent Clp protease proteolytic subunit ClpP (EC 3.4.21.92) | 15664911 | 99 | 99 | 90 | 3e-97 |
| Streptococcus dysgalactiae subsp. equisimilis strain T642 | clpP | ATP-dependent Clp protease proteolytic subunit ClpP (EC 3.4.21.92) | 15664911 | 99 | 99 | 90 | 0 |
| Streptococcus dysgalactiae subsp. equisimilis strain UT_4031CC | clpP | ATP-dependent Clp protease proteolytic subunit ClpP (EC 3.4.21.92) | 15664911 | 99 | 99 | 90 | 3e-97 |
| Streptococcus dysgalactiae subsp. equisimilis strain UT_4231_KK | clpP | ATP-dependent Clp protease proteolytic subunit ClpP (EC 3.4.21.92) | 15664911 | 99 | 99 | 90 | 3e-97 |
| Streptococcus dysgalactiae subsp. equisimilis strain UT_4234_DH | clpP | ATP-dependent Clp protease proteolytic subunit ClpP (EC 3.4.21.92) | 15664911 | 99 | 99 | 90 | 3e-97 |
| Streptococcus dysgalactiae subsp. equisimilis strain UT_4241_XS | clpP | ATP-dependent Clp protease proteolytic subunit ClpP (EC 3.4.21.92) | 15664911 | 99 | 99 | 90 | 3e-97 |
| Streptococcus dysgalactiae subsp. equisimilis strain UT_4242_AB | clpP | ATP-dependent Clp protease proteolytic subunit ClpP (EC 3.4.21.92) | 15664911 | 99 | 99 | 90 | 3e-97 |
| Streptococcus dysgalactiae subsp. equisimilis strain UT_4255RC | clpP | ATP-dependent Clp protease proteolytic subunit ClpP (EC 3.4.21.92) | 15664911 | 99 | 99 | 90 | 3e-97 |
| Streptococcus dysgalactiae subsp. equisimilis strain UT_4277_BB | clpP | ATP-dependent Clp protease proteolytic subunit ClpP (EC 3.4.21.92) | 15664911 | 99 | 99 | 90 | 3e-97 |
| Streptococcus dysgalactiae subsp. equisimilis strain UT_4966_RC | clpP | ATP-dependent Clp protease proteolytic subunit ClpP (EC 3.4.21.92) | 15664911 | 99 | 99 | 90 | 3e-97 |
| Streptococcus dysgalactiae subsp. equisimilis strain UT-5345 | clpP | ATP-dependent Clp protease proteolytic subunit ClpP (EC 3.4.21.92) | 15664911 | 99 | 99 | 90 | 3e-97 |
| Streptococcus dysgalactiae subsp. equisimilis strain UT-5354 | clpP | ATP-dependent Clp protease proteolytic subunit ClpP (EC 3.4.21.92) | 15664911 | 99 | 99 | 90 | 3e-97 |
| Streptococcus dysgalactiae subsp. equisimilis strain UT-SS1069 | clpP | ATP-dependent Clp protease proteolytic subunit ClpP (EC 3.4.21.92) | 15664911 | 99 | 99 | 90 | 3e-97 |
| Streptococcus dysgalactiae subsp. equisimilis strain UT-SS957 | clpP | ATP-dependent Clp protease proteolytic subunit ClpP (EC 3.4.21.92) | 15664911 | 99 | 99 | 90 | 3e-97 |
| Streptococcus dysgalactiae subsp. equisimilis strain WCHSDSE-1 | clpP | ATP-dependent Clp protease proteolytic subunit ClpP (EC 3.4.21.92) | 15664911 | 99 | 99 | 90 | 3e-97 |
| Streptococcus dysgalactiae subsp. equisimilis 167 | asnA | Aspartate--ammonia ligase (EC 6.3.1.1) | 12207705 | 100 | 100 | 87 | 1e-173 |
| Streptococcus dysgalactiae subsp. equisimilis AC-2713 | asnA | Aspartate--ammonia ligase (EC 6.3.1.1) | 12207705 | 100 | 100 | 87 | 1e-173 |
| Streptococcus dysgalactiae subsp. equisimilis AKSDE4288 | SP_1970 | Aspartate--ammonia ligase (EC 6.3.1.1) | 12207705 | 100 | 100 | 87 | 1e-173 |
| Streptococcus dysgalactiae subsp. equisimilis ATCC 12394 | SP_1970 | Aspartate--ammonia ligase (EC 6.3.1.1) | 12207705 | 100 | 100 | 87 | 1e-172 |
| Streptococcus dysgalactiae subsp. equisimilis GGS_124 | asnA | Aspartate--ammonia ligase (EC 6.3.1.1) | 12207705 | 100 | 100 | 87 | 1e-172 |
| Streptococcus dysgalactiae subsp. equisimilis RE378 | asnA | Aspartate--ammonia ligase (EC 6.3.1.1) | 12207705 | 100 | 100 | 87 | 1e-173 |
| Streptococcus dysgalactiae subsp. equisimilis SD SCDR1 | SP_1970 | Aspartate--ammonia ligase (EC 6.3.1.1) | 12207705 | 100 | 100 | 87 | 1e-173 |
| Streptococcus dysgalactiae subsp. equisimilis SK1249 | asnA | Aspartate--ammonia ligase (EC 6.3.1.1) | 12207705 | 100 | 100 | 87 | 1e-172 |
| Streptococcus dysgalactiae subsp. equisimilis SK1250 | asnA | Aspartate--ammonia ligase (EC 6.3.1.1) | 12207705 | 100 | 100 | 87 | 1e-173 |
| Streptococcus dysgalactiae subsp. equisimilis strain ASDSE_96 | SP_1970 | Aspartate--ammonia ligase (EC 6.3.1.1) | 12207705 | 100 | 100 | 87 | 1e-173 |
| Streptococcus dysgalactiae subsp. equisimilis strain ASDSE_99 | SP_1970 | Aspartate--ammonia ligase (EC 6.3.1.1) | 12207705 | 100 | 100 | 87 | 1e-171 |
| Streptococcus dysgalactiae subsp. equisimilis strain C161L1 | SP_1970 | Aspartate--ammonia ligase (EC 6.3.1.1) | 12207705 | 100 | 100 | 87 | 0 |
| Streptococcus dysgalactiae subsp. equisimilis strain KNZ01 | SP_1970 | Aspartate--ammonia ligase (EC 6.3.1.1) | 12207705 | 100 | 100 | 87 | 1e-173 |
| Streptococcus dysgalactiae subsp. equisimilis strain KNZ03 | SP_1970 | Aspartate--ammonia ligase (EC 6.3.1.1) | 12207705 | 100 | 100 | 87 | 1e-171 |
| Streptococcus dysgalactiae subsp. equisimilis strain KNZ04 | SP_1970 | Aspartate--ammonia ligase (EC 6.3.1.1) | 12207705 | 100 | 100 | 87 | 1e-171 |
| Streptococcus dysgalactiae subsp. equisimilis strain KNZ06 | SP_1970 | Aspartate--ammonia ligase (EC 6.3.1.1) | 12207705 | 100 | 100 | 87 | 1e-173 |
| Streptococcus dysgalactiae subsp. equisimilis strain KNZ07 | SP_1970 | Aspartate--ammonia ligase (EC 6.3.1.1) | 12207705 | 100 | 100 | 87 | 1e-173 |
| Streptococcus dysgalactiae subsp. equisimilis strain KNZ10 | SP_1970 | Aspartate--ammonia ligase (EC 6.3.1.1) | 12207705 | 100 | 100 | 87 | 1e-173 |
| Streptococcus dysgalactiae subsp. equisimilis strain KNZ12 | SP_1970 | Aspartate--ammonia ligase (EC 6.3.1.1) | 12207705 | 100 | 100 | 87 | 1e-173 |
| Streptococcus dysgalactiae subsp. equisimilis strain KNZ15 | SP_1970 | Aspartate--ammonia ligase (EC 6.3.1.1) | 12207705 | 100 | 100 | 87 | 1e-173 |
| Streptococcus dysgalactiae subsp. equisimilis strain KNZ16 | SP_1970 | Aspartate--ammonia ligase (EC 6.3.1.1) | 12207705 | 100 | 100 | 87 | 1e-172 |
| Streptococcus dysgalactiae subsp. equisimilis strain NCTC10321 | SP_1970 | Aspartate--ammonia ligase (EC 6.3.1.1) | 12207705 | 100 | 100 | 87 | 1e-173 |
| Streptococcus dysgalactiae subsp. equisimilis strain NCTC11554 | SP_1970 | Aspartate--ammonia ligase (EC 6.3.1.1) | 12207705 | 100 | 100 | 87 | 1e-172 |
| Streptococcus dysgalactiae subsp. equisimilis strain NCTC11555 | SP_1970 | Aspartate--ammonia ligase (EC 6.3.1.1) | 12207705 | 100 | 100 | 87 | 1e-173 |
| Streptococcus dysgalactiae subsp. equisimilis strain NCTC11556 | SP_1970 | Aspartate--ammonia ligase (EC 6.3.1.1) | 12207705 | 100 | 100 | 87 | 1e-171 |
| Streptococcus dysgalactiae subsp. equisimilis strain NCTC11557 | SP_1970 | Aspartate--ammonia ligase (EC 6.3.1.1) | 12207705 | 100 | 100 | 87 | 1e-171 |
| Streptococcus dysgalactiae subsp. equisimilis strain NCTC11564 | SP_1970 | Aspartate--ammonia ligase (EC 6.3.1.1) | 12207705 | 100 | 100 | 87 | 1e-173 |
| Streptococcus dysgalactiae subsp. equisimilis strain NCTC5370 | SP_1970 | Aspartate--ammonia ligase (EC 6.3.1.1) | 12207705 | 100 | 100 | 87 | 1e-173 |
| Streptococcus dysgalactiae subsp. equisimilis strain NCTC5371 | SP_1970 | Aspartate--ammonia ligase (EC 6.3.1.1) | 12207705 | 100 | 100 | 87 | 1e-173 |
| Streptococcus dysgalactiae subsp. equisimilis strain NCTC6179 | SP_1970 | Aspartate--ammonia ligase (EC 6.3.1.1) | 12207705 | 100 | 100 | 87 | 1e-173 |
| Streptococcus dysgalactiae subsp. equisimilis strain NCTC6181 | SP_1970 | Aspartate--ammonia ligase (EC 6.3.1.1) | 12207705 | 100 | 100 | 87 | 1e-173 |
| Streptococcus dysgalactiae subsp. equisimilis strain NCTC6407 | SP_1970 | Aspartate--ammonia ligase (EC 6.3.1.1) | 12207705 | 100 | 100 | 87 | 1e-173 |
| Streptococcus dysgalactiae subsp. equisimilis strain NCTC7136 | SP_1970 | Aspartate--ammonia ligase (EC 6.3.1.1) | 12207705 | 100 | 100 | 87 | 1e-173 |
| Streptococcus dysgalactiae subsp. equisimilis strain NCTC8543 | SP_1970 | Aspartate--ammonia ligase (EC 6.3.1.1) | 12207705 | 100 | 100 | 87 | 1e-172 |
| Streptococcus dysgalactiae subsp. equisimilis strain NCTC8546 | SP_1970 | Aspartate--ammonia ligase (EC 6.3.1.1) | 12207705 | 100 | 100 | 87 | 1e-173 |
| Streptococcus dysgalactiae subsp. equisimilis strain NCTC9413 | SP_1970 | Aspartate--ammonia ligase (EC 6.3.1.1) | 12207705 | 100 | 100 | 87 | 1e-173 |
| Streptococcus dysgalactiae subsp. equisimilis strain NCTC9414 | SP_1970 | Aspartate--ammonia ligase (EC 6.3.1.1) | 12207705 | 100 | 100 | 87 | 1e-173 |
| Streptococcus dysgalactiae subsp. equisimilis strain NCTC9603 | SP_1970 | Aspartate--ammonia ligase (EC 6.3.1.1) | 12207705 | 100 | 100 | 87 | 1e-173 |
| Streptococcus dysgalactiae subsp. equisimilis strain SS1575 | SP_1970 | Aspartate--ammonia ligase (EC 6.3.1.1) | 12207705 | 100 | 100 | 87 | 1e-173 |
| Streptococcus dysgalactiae subsp. equisimilis strain T642 | SP_1970 | Aspartate--ammonia ligase (EC 6.3.1.1) | 12207705 | 100 | 100 | 87 | 0 |
| Streptococcus dysgalactiae subsp. equisimilis strain UT_4031CC | SP_1970 | Aspartate--ammonia ligase (EC 6.3.1.1) | 12207705 | 100 | 100 | 87 | 1e-173 |
| Streptococcus dysgalactiae subsp. equisimilis strain UT_4231_KK | SP_1970 | Aspartate--ammonia ligase (EC 6.3.1.1) | 12207705 | 100 | 100 | 87 | 1e-173 |
| Streptococcus dysgalactiae subsp. equisimilis strain UT_4234_DH | SP_1970 | Aspartate--ammonia ligase (EC 6.3.1.1) | 12207705 | 100 | 100 | 87 | 1e-171 |
| Streptococcus dysgalactiae subsp. equisimilis strain UT_4241_XS | SP_1970 | Aspartate--ammonia ligase (EC 6.3.1.1) | 12207705 | 100 | 100 | 87 | 1e-173 |
| Streptococcus dysgalactiae subsp. equisimilis strain UT_4242_AB | SP_1970 | Aspartate--ammonia ligase (EC 6.3.1.1) | 12207705 | 100 | 100 | 87 | 1e-171 |
| Streptococcus dysgalactiae subsp. equisimilis strain UT_4255RC | SP_1970 | Aspartate--ammonia ligase (EC 6.3.1.1) | 12207705 | 100 | 100 | 87 | 1e-171 |
| Streptococcus dysgalactiae subsp. equisimilis strain UT_4277_BB | SP_1970 | Aspartate--ammonia ligase (EC 6.3.1.1) | 12207705 | 100 | 100 | 87 | 1e-173 |
| Streptococcus dysgalactiae subsp. equisimilis strain UT_4966_RC | SP_1970 | Aspartate--ammonia ligase (EC 6.3.1.1) | 12207705 | 100 | 100 | 87 | 1e-173 |
| Streptococcus dysgalactiae subsp. equisimilis strain UT-5345 | SP_1970 | Aspartate--ammonia ligase (EC 6.3.1.1) | 12207705 | 100 | 100 | 87 | 1e-173 |
| Streptococcus dysgalactiae subsp. equisimilis strain UT-5354 | SP_1970 | Aspartate--ammonia ligase (EC 6.3.1.1) | 12207705 | 100 | 100 | 87 | 1e-173 |
| Streptococcus dysgalactiae subsp. equisimilis strain UT-SS1069 | SP_1970 | Aspartate--ammonia ligase (EC 6.3.1.1) | 12207705 | 100 | 100 | 87 | 1e-173 |
| Streptococcus dysgalactiae subsp. equisimilis strain UT-SS957 | SP_1970 | Aspartate--ammonia ligase (EC 6.3.1.1) | 12207705 | 100 | 100 | 87 | 1e-173 |
| Streptococcus dysgalactiae subsp. equisimilis strain WCHSDSE-1 | SP_1970 | Aspartate--ammonia ligase (EC 6.3.1.1) | 12207705 | 100 | 100 | 87 | 1e-173 |
| Streptococcus dysgalactiae subsp. equisimilis 167 | idnO | 2-dehydro-3-deoxy-D-gluconate 5-dehydrogenase (EC 1.1.1.127) @ 2-deoxy-D-gluconate 3-dehydrogenase (EC 1.1.1.125) | 12207705 | 81 | 96 | 87 | 1e-108 |
| Streptococcus dysgalactiae subsp. equisimilis AC-2713 | idnO | 2-dehydro-3-deoxy-D-gluconate 5-dehydrogenase (EC 1.1.1.127) @ 2-deoxy-D-gluconate 3-dehydrogenase (EC 1.1.1.125) | 12207705 | 95 | 98 | 88 | 1e-130 |
| Streptococcus dysgalactiae subsp. equisimilis AKSDE4288 | SP_0320 | 2-dehydro-3-deoxy-D-gluconate 5-dehydrogenase (EC 1.1.1.127) @ 2-deoxy-D-gluconate 3-dehydrogenase (EC 1.1.1.125) | 12207705 | 95 | 98 | 88 | 1e-130 |
| Streptococcus dysgalactiae subsp. equisimilis ATCC 12394 | SP_0320 | 2-dehydro-3-deoxy-D-gluconate 5-dehydrogenase (EC 1.1.1.127) @ 2-deoxy-D-gluconate 3-dehydrogenase (EC 1.1.1.125) | 12207705 | 95 | 98 | 88 | 1e-130 |
| Streptococcus dysgalactiae subsp. equisimilis GGS_124 | idnO | 2-dehydro-3-deoxy-D-gluconate 5-dehydrogenase (EC 1.1.1.127) @ 2-deoxy-D-gluconate 3-dehydrogenase (EC 1.1.1.125) | 12207705 | 95 | 98 | 88 | 1e-130 |
| Streptococcus dysgalactiae subsp. equisimilis RE378 | idnO | 2-dehydro-3-deoxy-D-gluconate 5-dehydrogenase (EC 1.1.1.127) @ 2-deoxy-D-gluconate 3-dehydrogenase (EC 1.1.1.125) | 12207705 | 95 | 98 | 88 | 1e-130 |
| Streptococcus dysgalactiae subsp. equisimilis SD SCDR1 | SP_0320 | 2-dehydro-3-deoxy-D-gluconate 5-dehydrogenase (EC 1.1.1.127) @ 2-deoxy-D-gluconate 3-dehydrogenase (EC 1.1.1.125) | 12207705 | 95 | 98 | 88 | 1e-130 |
| Streptococcus dysgalactiae subsp. equisimilis SK1249 | SP_0320 | 2-dehydro-3-deoxy-D-gluconate 5-dehydrogenase (EC 1.1.1.127) @ 2-deoxy-D-gluconate 3-dehydrogenase (EC 1.1.1.125) | 12207705 | 77 | 96 | 86 | 1e-101 |
| Streptococcus dysgalactiae subsp. equisimilis SK1250 | SP_0320 | 2-dehydro-3-deoxy-D-gluconate 5-dehydrogenase (EC 1.1.1.127) @ 2-deoxy-D-gluconate 3-dehydrogenase (EC 1.1.1.125) | 12207705 | 95 | 98 | 88 | 1e-130 |
| Streptococcus dysgalactiae subsp. equisimilis strain ASDSE_96 | SP_0320 | 2-dehydro-3-deoxy-D-gluconate 5-dehydrogenase (EC 1.1.1.127) @ 2-deoxy-D-gluconate 3-dehydrogenase (EC 1.1.1.125) | 12207705 | 95 | 98 | 88 | 1e-130 |
| Streptococcus dysgalactiae subsp. equisimilis strain ASDSE_99 | SP_0320 | 2-dehydro-3-deoxy-D-gluconate 5-dehydrogenase (EC 1.1.1.127) @ 2-deoxy-D-gluconate 3-dehydrogenase (EC 1.1.1.125) | 12207705 | 95 | 98 | 88 | 1e-131 |
| Streptococcus dysgalactiae subsp. equisimilis strain C161L1 | SP_0320 | 2-dehydro-3-deoxy-D-gluconate 5-dehydrogenase (EC 1.1.1.127) @ 2-deoxy-D-gluconate 3-dehydrogenase (EC 1.1.1.125) | 12207705 | 95 | 98 | 88 | 0 |
| Streptococcus dysgalactiae subsp. equisimilis strain KNZ01 | SP_0320 | 2-dehydro-3-deoxy-D-gluconate 5-dehydrogenase (EC 1.1.1.127) @ 2-deoxy-D-gluconate 3-dehydrogenase (EC 1.1.1.125) | 12207705 | 95 | 98 | 88 | 1e-130 |
| Streptococcus dysgalactiae subsp. equisimilis strain KNZ03 | SP_0320 | 2-dehydro-3-deoxy-D-gluconate 5-dehydrogenase (EC 1.1.1.127) @ 2-deoxy-D-gluconate 3-dehydrogenase (EC 1.1.1.125) | 12207705 | 95 | 98 | 88 | 1e-130 |
| Streptococcus dysgalactiae subsp. equisimilis strain KNZ04 | SP_0320 | 2-dehydro-3-deoxy-D-gluconate 5-dehydrogenase (EC 1.1.1.127) @ 2-deoxy-D-gluconate 3-dehydrogenase (EC 1.1.1.125) | 12207705 | 95 | 98 | 88 | 1e-130 |
| Streptococcus dysgalactiae subsp. equisimilis strain KNZ06 | SP_0320 | 2-dehydro-3-deoxy-D-gluconate 5-dehydrogenase (EC 1.1.1.127) @ 2-deoxy-D-gluconate 3-dehydrogenase (EC 1.1.1.125) | 12207705 | 95 | 98 | 88 | 1e-130 |
| Streptococcus dysgalactiae subsp. equisimilis strain KNZ07 | SP_0320 | 2-dehydro-3-deoxy-D-gluconate 5-dehydrogenase (EC 1.1.1.127) @ 2-deoxy-D-gluconate 3-dehydrogenase (EC 1.1.1.125) | 12207705 | 95 | 98 | 88 | 1e-130 |
| Streptococcus dysgalactiae subsp. equisimilis strain KNZ10 | SP_0320 | 2-dehydro-3-deoxy-D-gluconate 5-dehydrogenase (EC 1.1.1.127) @ 2-deoxy-D-gluconate 3-dehydrogenase (EC 1.1.1.125) | 12207705 | 95 | 98 | 88 | 1e-130 |
| Streptococcus dysgalactiae subsp. equisimilis strain KNZ12 | SP_0320 | 2-dehydro-3-deoxy-D-gluconate 5-dehydrogenase (EC 1.1.1.127) @ 2-deoxy-D-gluconate 3-dehydrogenase (EC 1.1.1.125) | 12207705 | 95 | 98 | 88 | 1e-130 |
| Streptococcus dysgalactiae subsp. equisimilis strain KNZ15 | SP_0320 | 2-dehydro-3-deoxy-D-gluconate 5-dehydrogenase (EC 1.1.1.127) @ 2-deoxy-D-gluconate 3-dehydrogenase (EC 1.1.1.125) | 12207705 | 95 | 98 | 88 | 1e-130 |
| Streptococcus dysgalactiae subsp. equisimilis strain KNZ16 | SP_0320 | 2-dehydro-3-deoxy-D-gluconate 5-dehydrogenase (EC 1.1.1.127) @ 2-deoxy-D-gluconate 3-dehydrogenase (EC 1.1.1.125) | 12207705 | 95 | 98 | 88 | 1e-130 |
| Streptococcus dysgalactiae subsp. equisimilis strain NCTC10321 | SP_0320 | 2-dehydro-3-deoxy-D-gluconate 5-dehydrogenase (EC 1.1.1.127) @ 2-deoxy-D-gluconate 3-dehydrogenase (EC 1.1.1.125) | 12207705 | 95 | 98 | 88 | 1e-131 |
| Streptococcus dysgalactiae subsp. equisimilis strain NCTC11554 | SP_0320 | 2-dehydro-3-deoxy-D-gluconate 5-dehydrogenase (EC 1.1.1.127) @ 2-deoxy-D-gluconate 3-dehydrogenase (EC 1.1.1.125) | 12207705 | 95 | 98 | 88 | 1e-130 |
| Streptococcus dysgalactiae subsp. equisimilis strain NCTC11555 | SP_0320 | 2-dehydro-3-deoxy-D-gluconate 5-dehydrogenase (EC 1.1.1.127) @ 2-deoxy-D-gluconate 3-dehydrogenase (EC 1.1.1.125) | 12207705 | 95 | 98 | 88 | 1e-131 |
| Streptococcus dysgalactiae subsp. equisimilis strain NCTC11556 | SP_0320 | 2-dehydro-3-deoxy-D-gluconate 5-dehydrogenase (EC 1.1.1.127) @ 2-deoxy-D-gluconate 3-dehydrogenase (EC 1.1.1.125) | 12207705 | 95 | 98 | 88 | 1e-130 |
| Streptococcus dysgalactiae subsp. equisimilis strain NCTC11557 | SP_0320 | 2-dehydro-3-deoxy-D-gluconate 5-dehydrogenase (EC 1.1.1.127) @ 2-deoxy-D-gluconate 3-dehydrogenase (EC 1.1.1.125) | 12207705 | 95 | 98 | 88 | 1e-131 |
| Streptococcus dysgalactiae subsp. equisimilis strain NCTC11564 | SP_0320 | 2-dehydro-3-deoxy-D-gluconate 5-dehydrogenase (EC 1.1.1.127) @ 2-deoxy-D-gluconate 3-dehydrogenase (EC 1.1.1.125) | 12207705 | 95 | 98 | 88 | 1e-130 |
| Streptococcus dysgalactiae subsp. equisimilis strain NCTC5370 | SP_0320 | 2-dehydro-3-deoxy-D-gluconate 5-dehydrogenase (EC 1.1.1.127) @ 2-deoxy-D-gluconate 3-dehydrogenase (EC 1.1.1.125) | 12207705 | 95 | 98 | 88 | 1e-130 |
| Streptococcus dysgalactiae subsp. equisimilis strain NCTC5371 | SP_0320 | 2-dehydro-3-deoxy-D-gluconate 5-dehydrogenase (EC 1.1.1.127) @ 2-deoxy-D-gluconate 3-dehydrogenase (EC 1.1.1.125) | 12207705 | 95 | 98 | 88 | 1e-130 |
| Streptococcus dysgalactiae subsp. equisimilis strain NCTC6179 | SP_0320 | 2-dehydro-3-deoxy-D-gluconate 5-dehydrogenase (EC 1.1.1.127) @ 2-deoxy-D-gluconate 3-dehydrogenase (EC 1.1.1.125) | 12207705 | 95 | 98 | 88 | 1e-131 |
| Streptococcus dysgalactiae subsp. equisimilis strain NCTC6181 | SP_0320 | 2-dehydro-3-deoxy-D-gluconate 5-dehydrogenase (EC 1.1.1.127) @ 2-deoxy-D-gluconate 3-dehydrogenase (EC 1.1.1.125) | 12207705 | 95 | 98 | 88 | 1e-131 |
| Streptococcus dysgalactiae subsp. equisimilis strain NCTC6407 | SP_0320 | 2-dehydro-3-deoxy-D-gluconate 5-dehydrogenase (EC 1.1.1.127) @ 2-deoxy-D-gluconate 3-dehydrogenase (EC 1.1.1.125) | 12207705 | 95 | 98 | 88 | 1e-131 |
| Streptococcus dysgalactiae subsp. equisimilis strain NCTC7136 | SP_0320 | 2-dehydro-3-deoxy-D-gluconate 5-dehydrogenase (EC 1.1.1.127) @ 2-deoxy-D-gluconate 3-dehydrogenase (EC 1.1.1.125) | 12207705 | 95 | 98 | 88 | 1e-130 |
| Streptococcus dysgalactiae subsp. equisimilis strain NCTC8543 | SP_0320 | 2-dehydro-3-deoxy-D-gluconate 5-dehydrogenase (EC 1.1.1.127) @ 2-deoxy-D-gluconate 3-dehydrogenase (EC 1.1.1.125) | 12207705 | 95 | 98 | 88 | 1e-130 |
| Streptococcus dysgalactiae subsp. equisimilis strain NCTC8546 | SP_0320 | 2-dehydro-3-deoxy-D-gluconate 5-dehydrogenase (EC 1.1.1.127) @ 2-deoxy-D-gluconate 3-dehydrogenase (EC 1.1.1.125) | 12207705 | 95 | 98 | 88 | 1e-130 |
| Streptococcus dysgalactiae subsp. equisimilis strain NCTC9413 | SP_0320 | 2-dehydro-3-deoxy-D-gluconate 5-dehydrogenase (EC 1.1.1.127) @ 2-deoxy-D-gluconate 3-dehydrogenase (EC 1.1.1.125) | 12207705 | 95 | 98 | 88 | 1e-130 |
| Streptococcus dysgalactiae subsp. equisimilis strain NCTC9414 | SP_0320 | 2-dehydro-3-deoxy-D-gluconate 5-dehydrogenase (EC 1.1.1.127) @ 2-deoxy-D-gluconate 3-dehydrogenase (EC 1.1.1.125) | 12207705 | 95 | 98 | 88 | 1e-130 |
| Streptococcus dysgalactiae subsp. equisimilis strain NCTC9603 | SP_0320 | 2-dehydro-3-deoxy-D-gluconate 5-dehydrogenase (EC 1.1.1.127) @ 2-deoxy-D-gluconate 3-dehydrogenase (EC 1.1.1.125) | 12207705 | 95 | 98 | 88 | 1e-130 |
| Streptococcus dysgalactiae subsp. equisimilis strain SS1575 | SP_0320 | 2-dehydro-3-deoxy-D-gluconate 5-dehydrogenase (EC 1.1.1.127) @ 2-deoxy-D-gluconate 3-dehydrogenase (EC 1.1.1.125) | 12207705 | 95 | 98 | 88 | 1e-130 |
| Streptococcus dysgalactiae subsp. equisimilis strain T642 | SP_0320 | 2-dehydro-3-deoxy-D-gluconate 5-dehydrogenase (EC 1.1.1.127) @ 2-deoxy-D-gluconate 3-dehydrogenase (EC 1.1.1.125) | 12207705 | 95 | 98 | 88 | 0 |
| Streptococcus dysgalactiae subsp. equisimilis strain UT_4031CC | SP_0320 | 2-dehydro-3-deoxy-D-gluconate 5-dehydrogenase (EC 1.1.1.127) @ 2-deoxy-D-gluconate 3-dehydrogenase (EC 1.1.1.125) | 12207705 | 95 | 98 | 88 | 1e-130 |
| Streptococcus dysgalactiae subsp. equisimilis strain UT_4231_KK | SP_0320 | 2-dehydro-3-deoxy-D-gluconate 5-dehydrogenase (EC 1.1.1.127) @ 2-deoxy-D-gluconate 3-dehydrogenase (EC 1.1.1.125) | 12207705 | 95 | 98 | 88 | 1e-130 |
| Streptococcus dysgalactiae subsp. equisimilis strain UT_4234_DH | SP_0320 | 2-dehydro-3-deoxy-D-gluconate 5-dehydrogenase (EC 1.1.1.127) @ 2-deoxy-D-gluconate 3-dehydrogenase (EC 1.1.1.125) | 12207705 | 95 | 98 | 88 | 1e-131 |
| Streptococcus dysgalactiae subsp. equisimilis strain UT_4241_XS | SP_0320 | 2-dehydro-3-deoxy-D-gluconate 5-dehydrogenase (EC 1.1.1.127) @ 2-deoxy-D-gluconate 3-dehydrogenase (EC 1.1.1.125) | 12207705 | 95 | 98 | 88 | 1e-130 |
| Streptococcus dysgalactiae subsp. equisimilis strain UT_4242_AB | SP_0320 | 2-dehydro-3-deoxy-D-gluconate 5-dehydrogenase (EC 1.1.1.127) @ 2-deoxy-D-gluconate 3-dehydrogenase (EC 1.1.1.125) | 12207705 | 95 | 98 | 88 | 1e-130 |
| Streptococcus dysgalactiae subsp. equisimilis strain UT_4255RC | SP_0320 | 2-dehydro-3-deoxy-D-gluconate 5-dehydrogenase (EC 1.1.1.127) @ 2-deoxy-D-gluconate 3-dehydrogenase (EC 1.1.1.125) | 12207705 | 95 | 98 | 88 | 1e-131 |
| Streptococcus dysgalactiae subsp. equisimilis strain UT_4277_BB | SP_0320 | 2-dehydro-3-deoxy-D-gluconate 5-dehydrogenase (EC 1.1.1.127) @ 2-deoxy-D-gluconate 3-dehydrogenase (EC 1.1.1.125) | 12207705 | 95 | 98 | 88 | 1e-130 |
| Streptococcus dysgalactiae subsp. equisimilis strain UT_4966_RC | SP_0320 | 2-dehydro-3-deoxy-D-gluconate 5-dehydrogenase (EC 1.1.1.127) @ 2-deoxy-D-gluconate 3-dehydrogenase (EC 1.1.1.125) | 12207705 | 95 | 98 | 88 | 1e-130 |
| Streptococcus dysgalactiae subsp. equisimilis strain UT-5345 | SP_0320 | 2-dehydro-3-deoxy-D-gluconate 5-dehydrogenase (EC 1.1.1.127) @ 2-deoxy-D-gluconate 3-dehydrogenase (EC 1.1.1.125) | 12207705 | 95 | 98 | 88 | 1e-130 |
| Streptococcus dysgalactiae subsp. equisimilis strain UT-5354 | SP_0320 | 2-dehydro-3-deoxy-D-gluconate 5-dehydrogenase (EC 1.1.1.127) @ 2-deoxy-D-gluconate 3-dehydrogenase (EC 1.1.1.125) | 12207705 | 95 | 98 | 88 | 1e-130 |
| Streptococcus dysgalactiae subsp. equisimilis strain UT-SS1069 | SP_0320 | 2-dehydro-3-deoxy-D-gluconate 5-dehydrogenase (EC 1.1.1.127) @ 2-deoxy-D-gluconate 3-dehydrogenase (EC 1.1.1.125) | 12207705 | 95 | 98 | 88 | 1e-130 |
| Streptococcus dysgalactiae subsp. equisimilis strain UT-SS957 | SP_0320 | 2-dehydro-3-deoxy-D-gluconate 5-dehydrogenase (EC 1.1.1.127) @ 2-deoxy-D-gluconate 3-dehydrogenase (EC 1.1.1.125) | 12207705 | 95 | 98 | 88 | 1e-131 |
| Streptococcus dysgalactiae subsp. equisimilis strain WCHSDSE-1 | SP_0320 | 2-dehydro-3-deoxy-D-gluconate 5-dehydrogenase (EC 1.1.1.127) @ 2-deoxy-D-gluconate 3-dehydrogenase (EC 1.1.1.125) | 12207705 | 95 | 98 | 88 | 1e-130 |
