## Supplemental File 5 for "Comprehensive comparative genomics analysis for the emerging human pathogen *Streptococcus dysgalactiae* subsp. *equisimilis* (SDSE): A case study and Pan-subspecies genomic analysis"

**Supplementary Table 5:** Drug target genes/products annotated in SDSE genomes and proposed drug.

| Strain | Gene | Product/Target | Drug | Subject Coverage | Query Coverage | Identity | E-value |
| --- | --- | --- | --- | --- | --- | --- | --- |
| Streptococcus dysgalactiae subsp. equisimilis SD SCDR1 | def | Peptide deformylase (EC 3.5.1.88) | 2-[(Formyl-Hydroxy-Amino)-Methyl]-Heptanoic Acid [1-(2-Hydroxymethyl-Pyrrolidine-1-Carbonyl)-2-Methyl-Propyl]-Amide | 100 | 100 | 88 | 1e-101 |
| Streptococcus dysgalactiae subsp. equisimilis 167 | def | Peptide deformylase (EC 3.5.1.88) | 2-[(Formyl-Hydroxy-Amino)-Methyl]-Heptanoic Acid [1-(2-Hydroxymethyl-Pyrrolidine-1-Carbonyl)-2-Methyl-Propyl]-Amide | 100 | 100 | 88 | 1e-101 |
| Streptococcus dysgalactiae subsp. equisimilis AC-2713 | def | Peptide deformylase (EC 3.5.1.88) | 2-[(Formyl-Hydroxy-Amino)-Methyl]-Heptanoic Acid [1-(2-Hydroxymethyl-Pyrrolidine-1-Carbonyl)-2-Methyl-Propyl]-Amide | 100 | 100 | 88 | 1e-101 |
| Streptococcus dysgalactiae subsp. equisimilis AKSDE4288 | def | Peptide deformylase (EC 3.5.1.88) | 2-[(Formyl-Hydroxy-Amino)-Methyl]-Heptanoic Acid [1-(2-Hydroxymethyl-Pyrrolidine-1-Carbonyl)-2-Methyl-Propyl]-Amide | 100 | 100 | 88 | 1e-101 |
| Streptococcus dysgalactiae subsp. equisimilis ATCC 12394 | def | Peptide deformylase (EC 3.5.1.88) | 2-[(Formyl-Hydroxy-Amino)-Methyl]-Heptanoic Acid [1-(2-Hydroxymethyl-Pyrrolidine-1-Carbonyl)-2-Methyl-Propyl]-Amide | 100 | 100 | 88 | 1e-101 |
| Streptococcus dysgalactiae subsp. equisimilis GGS_124 | def | Peptide deformylase (EC 3.5.1.88) | 2-[(Formyl-Hydroxy-Amino)-Methyl]-Heptanoic Acid [1-(2-Hydroxymethyl-Pyrrolidine-1-Carbonyl)-2-Methyl-Propyl]-Amide | 100 | 100 | 88 | 1e-100 |
| Streptococcus dysgalactiae subsp. equisimilis RE378 | def | Peptide deformylase (EC 3.5.1.88) | 2-[(Formyl-Hydroxy-Amino)-Methyl]-Heptanoic Acid [1-(2-Hydroxymethyl-Pyrrolidine-1-Carbonyl)-2-Methyl-Propyl]-Amide | 100 | 100 | 88 | 1e-101 |
| Streptococcus dysgalactiae subsp. equisimilis SK1249 | def | Peptide deformylase (EC 3.5.1.88) | 2-[(Formyl-Hydroxy-Amino)-Methyl]-Heptanoic Acid [1-(2-Hydroxymethyl-Pyrrolidine-1-Carbonyl)-2-Methyl-Propyl]-Amide | 100 | 100 | 88 | 1e-101 |
| Streptococcus dysgalactiae subsp. equisimilis SK1250 | def_1 | Peptide deformylase (EC 3.5.1.88) | 2-[(Formyl-Hydroxy-Amino)-Methyl]-Heptanoic Acid [1-(2-Hydroxymethyl-Pyrrolidine-1-Carbonyl)-2-Methyl-Propyl]-Amide | 100 | 100 | 88 | 1e-101 |
| Streptococcus dysgalactiae subsp. equisimilis strain ASDSE_96 | def | Peptide deformylase (EC 3.5.1.88) | 2-[(Formyl-Hydroxy-Amino)-Methyl]-Heptanoic Acid [1-(2-Hydroxymethyl-Pyrrolidine-1-Carbonyl)-2-Methyl-Propyl]-Amide | 100 | 100 | 88 | 1e-101 |
| Streptococcus dysgalactiae subsp. equisimilis strain ASDSE_99 | def | Peptide deformylase (EC 3.5.1.88) | 2-[(Formyl-Hydroxy-Amino)-Methyl]-Heptanoic Acid [1-(2-Hydroxymethyl-Pyrrolidine-1-Carbonyl)-2-Methyl-Propyl]-Amide | 100 | 100 | 88 | 1e-101 |
| Streptococcus dysgalactiae subsp. equisimilis strain C161L1 | def | Peptide deformylase (EC 3.5.1.88) | 2-[(Formyl-Hydroxy-Amino)-Methyl]-Heptanoic Acid [1-(2-Hydroxymethyl-Pyrrolidine-1-Carbonyl)-2-Methyl-Propyl]-Amide | 100 | 100 | 88 | 0 |
| Streptococcus dysgalactiae subsp. equisimilis strain KNZ01 | def | Peptide deformylase (EC 3.5.1.88) | 2-[(Formyl-Hydroxy-Amino)-Methyl]-Heptanoic Acid [1-(2-Hydroxymethyl-Pyrrolidine-1-Carbonyl)-2-Methyl-Propyl]-Amide | 100 | 100 | 88 | 1e-101 |
| Streptococcus dysgalactiae subsp. equisimilis strain KNZ03 | def | Peptide deformylase (EC 3.5.1.88) | 2-[(Formyl-Hydroxy-Amino)-Methyl]-Heptanoic Acid [1-(2-Hydroxymethyl-Pyrrolidine-1-Carbonyl)-2-Methyl-Propyl]-Amide | 100 | 100 | 88 | 1e-101 |
| Streptococcus dysgalactiae subsp. equisimilis strain KNZ04 | def | Peptide deformylase (EC 3.5.1.88) | 2-[(Formyl-Hydroxy-Amino)-Methyl]-Heptanoic Acid [1-(2-Hydroxymethyl-Pyrrolidine-1-Carbonyl)-2-Methyl-Propyl]-Amide | 100 | 100 | 88 | 1e-101 |
| Streptococcus dysgalactiae subsp. equisimilis strain KNZ06 | def | Peptide deformylase (EC 3.5.1.88) | 2-[(Formyl-Hydroxy-Amino)-Methyl]-Heptanoic Acid [1-(2-Hydroxymethyl-Pyrrolidine-1-Carbonyl)-2-Methyl-Propyl]-Amide | 100 | 100 | 88 | 1e-101 |
| Streptococcus dysgalactiae subsp. equisimilis strain KNZ07 | def | Peptide deformylase (EC 3.5.1.88) | 2-[(Formyl-Hydroxy-Amino)-Methyl]-Heptanoic Acid [1-(2-Hydroxymethyl-Pyrrolidine-1-Carbonyl)-2-Methyl-Propyl]-Amide | 100 | 100 | 88 | 1e-101 |
| Streptococcus dysgalactiae subsp. equisimilis strain KNZ10 | def | Peptide deformylase (EC 3.5.1.88) | 2-[(Formyl-Hydroxy-Amino)-Methyl]-Heptanoic Acid [1-(2-Hydroxymethyl-Pyrrolidine-1-Carbonyl)-2-Methyl-Propyl]-Amide | 100 | 100 | 88 | 1e-101 |
| Streptococcus dysgalactiae subsp. equisimilis strain KNZ12 | def | Peptide deformylase (EC 3.5.1.88) | 2-[(Formyl-Hydroxy-Amino)-Methyl]-Heptanoic Acid [1-(2-Hydroxymethyl-Pyrrolidine-1-Carbonyl)-2-Methyl-Propyl]-Amide | 100 | 100 | 88 | 1e-101 |
| Streptococcus dysgalactiae subsp. equisimilis strain KNZ15 | def | Peptide deformylase (EC 3.5.1.88) | 2-[(Formyl-Hydroxy-Amino)-Methyl]-Heptanoic Acid [1-(2-Hydroxymethyl-Pyrrolidine-1-Carbonyl)-2-Methyl-Propyl]-Amide | 100 | 100 | 88 | 1e-101 |
| Streptococcus dysgalactiae subsp. equisimilis strain KNZ16 | def | Peptide deformylase (EC 3.5.1.88) | 2-[(Formyl-Hydroxy-Amino)-Methyl]-Heptanoic Acid [1-(2-Hydroxymethyl-Pyrrolidine-1-Carbonyl)-2-Methyl-Propyl]-Amide | 100 | 100 | 88 | 1e-101 |
| Streptococcus dysgalactiae subsp. equisimilis strain NCTC10321 | def | Peptide deformylase (EC 3.5.1.88) | 2-[(Formyl-Hydroxy-Amino)-Methyl]-Heptanoic Acid [1-(2-Hydroxymethyl-Pyrrolidine-1-Carbonyl)-2-Methyl-Propyl]-Amide | 100 | 100 | 88 | 1e-101 |
| Streptococcus dysgalactiae subsp. equisimilis strain NCTC11554 | def | Peptide deformylase (EC 3.5.1.88) | 2-[(Formyl-Hydroxy-Amino)-Methyl]-Heptanoic Acid [1-(2-Hydroxymethyl-Pyrrolidine-1-Carbonyl)-2-Methyl-Propyl]-Amide | 100 | 100 | 88 | 1e-101 |
| Streptococcus dysgalactiae subsp. equisimilis strain NCTC11555 | def | Peptide deformylase (EC 3.5.1.88) | 2-[(Formyl-Hydroxy-Amino)-Methyl]-Heptanoic Acid [1-(2-Hydroxymethyl-Pyrrolidine-1-Carbonyl)-2-Methyl-Propyl]-Amide | 100 | 100 | 88 | 1e-101 |
| Streptococcus dysgalactiae subsp. equisimilis strain NCTC11556 | def | Peptide deformylase (EC 3.5.1.88) | 2-[(Formyl-Hydroxy-Amino)-Methyl]-Heptanoic Acid [1-(2-Hydroxymethyl-Pyrrolidine-1-Carbonyl)-2-Methyl-Propyl]-Amide | 100 | 100 | 88 | 1e-101 |
| Streptococcus dysgalactiae subsp. equisimilis strain NCTC11557 | def | Peptide deformylase (EC 3.5.1.88) | 2-[(Formyl-Hydroxy-Amino)-Methyl]-Heptanoic Acid [1-(2-Hydroxymethyl-Pyrrolidine-1-Carbonyl)-2-Methyl-Propyl]-Amide | 100 | 100 | 88 | 1e-101 |
| Streptococcus dysgalactiae subsp. equisimilis strain NCTC11564 | def | Peptide deformylase (EC 3.5.1.88) | 2-[(Formyl-Hydroxy-Amino)-Methyl]-Heptanoic Acid [1-(2-Hydroxymethyl-Pyrrolidine-1-Carbonyl)-2-Methyl-Propyl]-Amide | 100 | 100 | 88 | 1e-101 |
| Streptococcus dysgalactiae subsp. equisimilis strain NCTC5370 | def | Peptide deformylase (EC 3.5.1.88) | 2-[(Formyl-Hydroxy-Amino)-Methyl]-Heptanoic Acid [1-(2-Hydroxymethyl-Pyrrolidine-1-Carbonyl)-2-Methyl-Propyl]-Amide | 100 | 100 | 88 | 1e-101 |
| Streptococcus dysgalactiae subsp. equisimilis strain NCTC5371 | def | Peptide deformylase (EC 3.5.1.88) | 2-[(Formyl-Hydroxy-Amino)-Methyl]-Heptanoic Acid [1-(2-Hydroxymethyl-Pyrrolidine-1-Carbonyl)-2-Methyl-Propyl]-Amide | 100 | 100 | 88 | 1e-101 |
| Streptococcus dysgalactiae subsp. equisimilis strain NCTC6179 | def | Peptide deformylase (EC 3.5.1.88) | 2-[(Formyl-Hydroxy-Amino)-Methyl]-Heptanoic Acid [1-(2-Hydroxymethyl-Pyrrolidine-1-Carbonyl)-2-Methyl-Propyl]-Amide | 100 | 100 | 88 | 1e-101 |
| Streptococcus dysgalactiae subsp. equisimilis strain NCTC6181 | def | Peptide deformylase (EC 3.5.1.88) | 2-[(Formyl-Hydroxy-Amino)-Methyl]-Heptanoic Acid [1-(2-Hydroxymethyl-Pyrrolidine-1-Carbonyl)-2-Methyl-Propyl]-Amide | 100 | 100 | 88 | 1e-101 |
| Streptococcus dysgalactiae subsp. equisimilis strain NCTC6407 | def | Peptide deformylase (EC 3.5.1.88) | 2-[(Formyl-Hydroxy-Amino)-Methyl]-Heptanoic Acid [1-(2-Hydroxymethyl-Pyrrolidine-1-Carbonyl)-2-Methyl-Propyl]-Amide | 100 | 100 | 88 | 1e-101 |
| Streptococcus dysgalactiae subsp. equisimilis strain NCTC7136 | def | Peptide deformylase (EC 3.5.1.88) | 2-[(Formyl-Hydroxy-Amino)-Methyl]-Heptanoic Acid [1-(2-Hydroxymethyl-Pyrrolidine-1-Carbonyl)-2-Methyl-Propyl]-Amide | 100 | 100 | 88 | 1e-100 |
| Streptococcus dysgalactiae subsp. equisimilis strain NCTC8543 | def | Peptide deformylase (EC 3.5.1.88) | 2-[(Formyl-Hydroxy-Amino)-Methyl]-Heptanoic Acid [1-(2-Hydroxymethyl-Pyrrolidine-1-Carbonyl)-2-Methyl-Propyl]-Amide | 100 | 100 | 88 | 1e-101 |
| Streptococcus dysgalactiae subsp. equisimilis strain NCTC8546 | def | Peptide deformylase (EC 3.5.1.88) | 2-[(Formyl-Hydroxy-Amino)-Methyl]-Heptanoic Acid [1-(2-Hydroxymethyl-Pyrrolidine-1-Carbonyl)-2-Methyl-Propyl]-Amide | 100 | 100 | 88 | 1e-101 |
| Streptococcus dysgalactiae subsp. equisimilis strain NCTC9413 | def | Peptide deformylase (EC 3.5.1.88) | 2-[(Formyl-Hydroxy-Amino)-Methyl]-Heptanoic Acid [1-(2-Hydroxymethyl-Pyrrolidine-1-Carbonyl)-2-Methyl-Propyl]-Amide | 100 | 100 | 88 | 1e-101 |
| Streptococcus dysgalactiae subsp. equisimilis strain NCTC9414 | def | Peptide deformylase (EC 3.5.1.88) | 2-[(Formyl-Hydroxy-Amino)-Methyl]-Heptanoic Acid [1-(2-Hydroxymethyl-Pyrrolidine-1-Carbonyl)-2-Methyl-Propyl]-Amide | 100 | 100 | 88 | 1e-101 |
| Streptococcus dysgalactiae subsp. equisimilis strain NCTC9603 | def | Peptide deformylase (EC 3.5.1.88) | 2-[(Formyl-Hydroxy-Amino)-Methyl]-Heptanoic Acid [1-(2-Hydroxymethyl-Pyrrolidine-1-Carbonyl)-2-Methyl-Propyl]-Amide | 100 | 100 | 88 | 1e-101 |
| Streptococcus dysgalactiae subsp. equisimilis strain SS1575 | def | Peptide deformylase (EC 3.5.1.88) | 2-[(Formyl-Hydroxy-Amino)-Methyl]-Heptanoic Acid [1-(2-Hydroxymethyl-Pyrrolidine-1-Carbonyl)-2-Methyl-Propyl]-Amide | 100 | 100 | 88 | 1e-101 |
| Streptococcus dysgalactiae subsp. equisimilis strain T642 | def | Peptide deformylase (EC 3.5.1.88) | 2-[(Formyl-Hydroxy-Amino)-Methyl]-Heptanoic Acid [1-(2-Hydroxymethyl-Pyrrolidine-1-Carbonyl)-2-Methyl-Propyl]-Amide | 100 | 100 | 88 | 0 |
| Streptococcus dysgalactiae subsp. equisimilis strain UT_4031CC | def | Peptide deformylase (EC 3.5.1.88) | 2-[(Formyl-Hydroxy-Amino)-Methyl]-Heptanoic Acid [1-(2-Hydroxymethyl-Pyrrolidine-1-Carbonyl)-2-Methyl-Propyl]-Amide | 100 | 100 | 88 | 1e-101 |
| Streptococcus dysgalactiae subsp. equisimilis strain UT_4231_KK | def | Peptide deformylase (EC 3.5.1.88) | 2-[(Formyl-Hydroxy-Amino)-Methyl]-Heptanoic Acid [1-(2-Hydroxymethyl-Pyrrolidine-1-Carbonyl)-2-Methyl-Propyl]-Amide | 100 | 100 | 88 | 1e-101 |
| Streptococcus dysgalactiae subsp. equisimilis strain UT_4234_DH | def | Peptide deformylase (EC 3.5.1.88) | 2-[(Formyl-Hydroxy-Amino)-Methyl]-Heptanoic Acid [1-(2-Hydroxymethyl-Pyrrolidine-1-Carbonyl)-2-Methyl-Propyl]-Amide | 100 | 100 | 88 | 1e-101 |
| Streptococcus dysgalactiae subsp. equisimilis strain UT_4241_XS | def | Peptide deformylase (EC 3.5.1.88) | 2-[(Formyl-Hydroxy-Amino)-Methyl]-Heptanoic Acid [1-(2-Hydroxymethyl-Pyrrolidine-1-Carbonyl)-2-Methyl-Propyl]-Amide | 100 | 100 | 88 | 1e-101 |
| Streptococcus dysgalactiae subsp. equisimilis strain UT_4242_AB | def | Peptide deformylase (EC 3.5.1.88) | 2-[(Formyl-Hydroxy-Amino)-Methyl]-Heptanoic Acid [1-(2-Hydroxymethyl-Pyrrolidine-1-Carbonyl)-2-Methyl-Propyl]-Amide | 100 | 100 | 88 | 1e-101 |
| Streptococcus dysgalactiae subsp. equisimilis strain UT_4255RC | def | Peptide deformylase (EC 3.5.1.88) | 2-[(Formyl-Hydroxy-Amino)-Methyl]-Heptanoic Acid [1-(2-Hydroxymethyl-Pyrrolidine-1-Carbonyl)-2-Methyl-Propyl]-Amide | 100 | 100 | 88 | 1e-101 |
| Streptococcus dysgalactiae subsp. equisimilis strain UT_4277_BB | def | Peptide deformylase (EC 3.5.1.88) | 2-[(Formyl-Hydroxy-Amino)-Methyl]-Heptanoic Acid [1-(2-Hydroxymethyl-Pyrrolidine-1-Carbonyl)-2-Methyl-Propyl]-Amide | 100 | 100 | 88 | 1e-101 |
| Streptococcus dysgalactiae subsp. equisimilis strain UT_4966_RC | def | Peptide deformylase (EC 3.5.1.88) | 2-[(Formyl-Hydroxy-Amino)-Methyl]-Heptanoic Acid [1-(2-Hydroxymethyl-Pyrrolidine-1-Carbonyl)-2-Methyl-Propyl]-Amide | 100 | 100 | 88 | 1e-101 |
| Streptococcus dysgalactiae subsp. equisimilis strain UT-5345 | def | Peptide deformylase (EC 3.5.1.88) | 2-[(Formyl-Hydroxy-Amino)-Methyl]-Heptanoic Acid [1-(2-Hydroxymethyl-Pyrrolidine-1-Carbonyl)-2-Methyl-Propyl]-Amide | 100 | 100 | 88 | 1e-101 |
| Streptococcus dysgalactiae subsp. equisimilis strain UT-5354 | def | Peptide deformylase (EC 3.5.1.88) | 2-[(Formyl-Hydroxy-Amino)-Methyl]-Heptanoic Acid [1-(2-Hydroxymethyl-Pyrrolidine-1-Carbonyl)-2-Methyl-Propyl]-Amide | 100 | 100 | 88 | 1e-101 |
| Streptococcus dysgalactiae subsp. equisimilis strain UT-SS1069 | def | Peptide deformylase (EC 3.5.1.88) | 2-[(Formyl-Hydroxy-Amino)-Methyl]-Heptanoic Acid [1-(2-Hydroxymethyl-Pyrrolidine-1-Carbonyl)-2-Methyl-Propyl]-Amide | 100 | 100 | 88 | 1e-101 |
| Streptococcus dysgalactiae subsp. equisimilis strain UT-SS957 | def | Peptide deformylase (EC 3.5.1.88) | 2-[(Formyl-Hydroxy-Amino)-Methyl]-Heptanoic Acid [1-(2-Hydroxymethyl-Pyrrolidine-1-Carbonyl)-2-Methyl-Propyl]-Amide | 100 | 100 | 88 | 1e-101 |
| Streptococcus dysgalactiae subsp. equisimilis strain WCHSDSE-1 | def | Peptide deformylase (EC 3.5.1.88) | 2-[(Formyl-Hydroxy-Amino)-Methyl]-Heptanoic Acid [1-(2-Hydroxymethyl-Pyrrolidine-1-Carbonyl)-2-Methyl-Propyl]-Amide | 100 | 100 | 88 | 1e-101 |
| Streptococcus dysgalactiae subsp. equisimilis SD SCDR1 | gyrB | DNA gyrase subunit B (EC 5.99.1.3) | Gatifloxacin | 98 | 98 | 87 | 0.0 |
| Streptococcus dysgalactiae subsp. equisimilis 167 | gyrB | DNA gyrase subunit B (EC 5.99.1.3) | Gatifloxacin | 98 | 98 | 87 | 0.0 |
| Streptococcus dysgalactiae subsp. equisimilis AC-2713 | gyrB | DNA gyrase subunit B (EC 5.99.1.3) | Gatifloxacin | 98 | 98 | 87 | 0.0 |
| Streptococcus dysgalactiae subsp. equisimilis AKSDE4288 | gyrB | DNA gyrase subunit B (EC 5.99.1.3) | Gatifloxacin | 98 | 98 | 87 | 0.0 |
| Streptococcus dysgalactiae subsp. equisimilis ATCC 12394 | gyrB | DNA gyrase subunit B (EC 5.99.1.3) | Gatifloxacin | 15 | 98 | 86 | 7e-46 |
| Streptococcus dysgalactiae subsp. equisimilis GGS_124 | gyrB | DNA gyrase subunit B (EC 5.99.1.3) | Gatifloxacin | 98 | 98 | 87 | 0.0 |
| Streptococcus dysgalactiae subsp. equisimilis RE378 | gyrB | DNA gyrase subunit B (EC 5.99.1.3) | Gatifloxacin | 98 | 98 | 87 | 0.0 |
| Streptococcus dysgalactiae subsp. equisimilis SK1249 | gyrB | DNA gyrase subunit B (EC 5.99.1.3) | Gatifloxacin | 89 | 99 | 87 | 1e-302 |
| Streptococcus dysgalactiae subsp. equisimilis SK1250 | gyrB | DNA gyrase subunit B (EC 5.99.1.3) | Gatifloxacin | 98 | 98 | 87 | 0.0 |
| Streptococcus dysgalactiae subsp. equisimilis strain ASDSE_96 | gyrB | DNA gyrase subunit B (EC 5.99.1.3) | Gatifloxacin | 98 | 98 | 87 | 0.0 |
| Streptococcus dysgalactiae subsp. equisimilis strain ASDSE_99 | gyrB | DNA gyrase subunit B (EC 5.99.1.3) | Gatifloxacin | 98 | 98 | 87 | 0.0 |
| Streptococcus dysgalactiae subsp. equisimilis strain C161L1 | gyrB | DNA gyrase subunit B (EC 5.99.1.3) | Gatifloxacin | 100 | 100 | 86 | 0 |
| Streptococcus dysgalactiae subsp. equisimilis strain KNZ01 | gyrB | DNA gyrase subunit B (EC 5.99.1.3) | Gatifloxacin | 98 | 98 | 87 | 0.0 |
| Streptococcus dysgalactiae subsp. equisimilis strain KNZ03 | gyrB | DNA gyrase subunit B (EC 5.99.1.3) | Gatifloxacin | 98 | 98 | 87 | 0.0 |
| Streptococcus dysgalactiae subsp. equisimilis strain KNZ04 | gyrB | DNA gyrase subunit B (EC 5.99.1.3) | Gatifloxacin | 98 | 98 | 87 | 0.0 |
| Streptococcus dysgalactiae subsp. equisimilis strain KNZ06 | gyrB | DNA gyrase subunit B (EC 5.99.1.3) | Gatifloxacin | 98 | 98 | 87 | 0.0 |
| Streptococcus dysgalactiae subsp. equisimilis strain KNZ07 | gyrB | DNA gyrase subunit B (EC 5.99.1.3) | Gatifloxacin | 98 | 98 | 87 | 0.0 |
| Streptococcus dysgalactiae subsp. equisimilis strain KNZ10 | gyrB | DNA gyrase subunit B (EC 5.99.1.3) | Gatifloxacin | 98 | 98 | 87 | 0.0 |
| Streptococcus dysgalactiae subsp. equisimilis strain KNZ12 | gyrB | DNA gyrase subunit B (EC 5.99.1.3) | Gatifloxacin | 98 | 98 | 87 | 0.0 |
| Streptococcus dysgalactiae subsp. equisimilis strain KNZ15 | gyrB | DNA gyrase subunit B (EC 5.99.1.3) | Gatifloxacin | 98 | 98 | 87 | 0.0 |
| Streptococcus dysgalactiae subsp. equisimilis strain KNZ16 | gyrB | DNA gyrase subunit B (EC 5.99.1.3) | Gatifloxacin | 98 | 98 | 87 | 0.0 |
| Streptococcus dysgalactiae subsp. equisimilis strain NCTC10321 | gyrB | DNA gyrase subunit B (EC 5.99.1.3) | Gatifloxacin | 98 | 98 | 87 | 0.0 |
| Streptococcus dysgalactiae subsp. equisimilis strain NCTC11554 | gyrB | DNA gyrase subunit B (EC 5.99.1.3) | Gatifloxacin | 98 | 98 | 87 | 0.0 |
| Streptococcus dysgalactiae subsp. equisimilis strain NCTC11555 | gyrB | DNA gyrase subunit B (EC 5.99.1.3) | Gatifloxacin | 98 | 98 | 87 | 0.0 |
| Streptococcus dysgalactiae subsp. equisimilis strain NCTC11556 | gyrB | DNA gyrase subunit B (EC 5.99.1.3) | Gatifloxacin | 98 | 98 | 87 | 0.0 |
| Streptococcus dysgalactiae subsp. equisimilis strain NCTC11557 | gyrB | DNA gyrase subunit B (EC 5.99.1.3) | Gatifloxacin | 98 | 98 | 87 | 0.0 |
| Streptococcus dysgalactiae subsp. equisimilis strain NCTC11564 | gyrB | DNA gyrase subunit B (EC 5.99.1.3) | Gatifloxacin | 98 | 98 | 87 | 0.0 |
| Streptococcus dysgalactiae subsp. equisimilis strain NCTC5370 | gyrB | DNA gyrase subunit B (EC 5.99.1.3) | Gatifloxacin | 98 | 98 | 87 | 0.0 |
| Streptococcus dysgalactiae subsp. equisimilis strain NCTC5371 | gyrB | DNA gyrase subunit B (EC 5.99.1.3) | Gatifloxacin | 98 | 98 | 87 | 0.0 |
| Streptococcus dysgalactiae subsp. equisimilis strain NCTC6179 | gyrB | DNA gyrase subunit B (EC 5.99.1.3) | Gatifloxacin | 98 | 98 | 87 | 0.0 |
| Streptococcus dysgalactiae subsp. equisimilis strain NCTC6181 | gyrB | DNA gyrase subunit B (EC 5.99.1.3) | Gatifloxacin | 98 | 98 | 87 | 0.0 |
| Streptococcus dysgalactiae subsp. equisimilis strain NCTC6407 | gyrB | DNA gyrase subunit B (EC 5.99.1.3) | Gatifloxacin | 98 | 98 | 87 | 0.0 |
| Streptococcus dysgalactiae subsp. equisimilis strain NCTC7136 | gyrB | DNA gyrase subunit B (EC 5.99.1.3) | Gatifloxacin | 98 | 98 | 87 | 0.0 |
| Streptococcus dysgalactiae subsp. equisimilis strain NCTC8543 | gyrB | DNA gyrase subunit B (EC 5.99.1.3) | Gatifloxacin | 98 | 98 | 87 | 0.0 |
| Streptococcus dysgalactiae subsp. equisimilis strain NCTC8546 | gyrB | DNA gyrase subunit B (EC 5.99.1.3) | Gatifloxacin | 98 | 98 | 87 | 0.0 |
| Streptococcus dysgalactiae subsp. equisimilis strain NCTC9413 | gyrB | DNA gyrase subunit B (EC 5.99.1.3) | Gatifloxacin | 98 | 98 | 87 | 0.0 |
| Streptococcus dysgalactiae subsp. equisimilis strain NCTC9414 | gyrB | DNA gyrase subunit B (EC 5.99.1.3) | Gatifloxacin | 98 | 98 | 87 | 0.0 |
| Streptococcus dysgalactiae subsp. equisimilis strain NCTC9603 | gyrB | DNA gyrase subunit B (EC 5.99.1.3) | Gatifloxacin | 98 | 98 | 87 | 0.0 |
| Streptococcus dysgalactiae subsp. equisimilis strain SS1575 | gyrB | DNA gyrase subunit B (EC 5.99.1.3) | Gatifloxacin | 98 | 98 | 87 | 0.0 |
| Streptococcus dysgalactiae subsp. equisimilis strain T642 | gyrB | DNA gyrase subunit B (EC 5.99.1.3) | Gatifloxacin | 100 | 100 | 86 | 0 |
| Streptococcus dysgalactiae subsp. equisimilis strain UT_4031CC | gyrB | DNA gyrase subunit B (EC 5.99.1.3) | Gatifloxacin | 98 | 98 | 87 | 0.0 |
| Streptococcus dysgalactiae subsp. equisimilis strain UT_4231_KK | gyrB | DNA gyrase subunit B (EC 5.99.1.3) | Gatifloxacin | 98 | 98 | 87 | 0.0 |
| Streptococcus dysgalactiae subsp. equisimilis strain UT_4234_DH | gyrB | DNA gyrase subunit B (EC 5.99.1.3) | Gatifloxacin | 98 | 98 | 87 | 0.0 |
| Streptococcus dysgalactiae subsp. equisimilis strain UT_4241_XS | gyrB | DNA gyrase subunit B (EC 5.99.1.3) | Gatifloxacin | 98 | 98 | 87 | 0.0 |
| Streptococcus dysgalactiae subsp. equisimilis strain UT_4242_AB | gyrB | DNA gyrase subunit B (EC 5.99.1.3) | Gatifloxacin | 98 | 98 | 87 | 0.0 |
| Streptococcus dysgalactiae subsp. equisimilis strain UT_4255RC | gyrB | DNA gyrase subunit B (EC 5.99.1.3) | Gatifloxacin | 98 | 98 | 87 | 0.0 |
| Streptococcus dysgalactiae subsp. equisimilis strain UT_4277_BB | gyrB | DNA gyrase subunit B (EC 5.99.1.3) | Gatifloxacin | 98 | 98 | 87 | 0.0 |
| Streptococcus dysgalactiae subsp. equisimilis strain UT_4966_RC | gyrB | DNA gyrase subunit B (EC 5.99.1.3) | Gatifloxacin | 98 | 98 | 87 | 0.0 |
| Streptococcus dysgalactiae subsp. equisimilis strain UT-5345 | gyrB | DNA gyrase subunit B (EC 5.99.1.3) | Gatifloxacin | 98 | 98 | 87 | 0.0 |
| Streptococcus dysgalactiae subsp. equisimilis strain UT-5354 | gyrB | DNA gyrase subunit B (EC 5.99.1.3) | Gatifloxacin | 98 | 98 | 87 | 0.0 |
| Streptococcus dysgalactiae subsp. equisimilis strain UT-SS1069 | gyrB | DNA gyrase subunit B (EC 5.99.1.3) | Gatifloxacin | 62 | 98 | 88 | 1e-211 |
| Streptococcus dysgalactiae subsp. equisimilis strain UT-SS957 | gyrB | DNA gyrase subunit B (EC 5.99.1.3) | Gatifloxacin | 98 | 98 | 87 | 0.0 |
| Streptococcus dysgalactiae subsp. equisimilis strain WCHSDSE-1 | gyrB | DNA gyrase subunit B (EC 5.99.1.3) | Gatifloxacin | 98 | 98 | 87 | 0.0 |
| Streptococcus dysgalactiae subsp. equisimilis 167 | lacG | 6-phospho-beta-galactosidase (EC 3.2.1.85) | beta-D-galactose 6-phosphate | 99 | 99 | 91 | 1e-269 |
| Streptococcus dysgalactiae subsp. equisimilis RE378 | lacG | 6-phospho-beta-galactosidase (EC 3.2.1.85) | beta-D-galactose 6-phosphate | 99 | 99 | 91 | 1e-269 |
| Streptococcus dysgalactiae subsp. equisimilis SK1250 | lacG | 6-phospho-beta-galactosidase (EC 3.2.1.85) | beta-D-galactose 6-phosphate | 99 | 99 | 91 | 1e-269 |
| Streptococcus dysgalactiae subsp. equisimilis strain ASDSE_96 | lacG | 6-phospho-beta-galactosidase (EC 3.2.1.85) | beta-D-galactose 6-phosphate | 99 | 99 | 91 | 1e-269 |
| Streptococcus dysgalactiae subsp. equisimilis strain C161L1 | lacG | 6-phospho-beta-galactosidase (EC 3.2.1.85) | beta-D-galactose 6-phosphate | 100 | 100 | 91 | 0 |
| Streptococcus dysgalactiae subsp. equisimilis strain KNZ01 | lacG | 6-phospho-beta-galactosidase (EC 3.2.1.85) | beta-D-galactose 6-phosphate | 99 | 99 | 91 | 1e-269 |
| Streptococcus dysgalactiae subsp. equisimilis strain KNZ03 | lacG | 6-phospho-beta-galactosidase (EC 3.2.1.85) | beta-D-galactose 6-phosphate | 99 | 99 | 91 | 1e-269 |
| Streptococcus dysgalactiae subsp. equisimilis strain KNZ04 | lacG | 6-phospho-beta-galactosidase (EC 3.2.1.85) | beta-D-galactose 6-phosphate | 99 | 99 | 91 | 1e-269 |
| Streptococcus dysgalactiae subsp. equisimilis strain KNZ07 | lacG | 6-phospho-beta-galactosidase (EC 3.2.1.85) | beta-D-galactose 6-phosphate | 99 | 99 | 91 | 1e-269 |
| Streptococcus dysgalactiae subsp. equisimilis strain KNZ10 | lacG | 6-phospho-beta-galactosidase (EC 3.2.1.85) | beta-D-galactose 6-phosphate | 99 | 99 | 91 | 1e-269 |
| Streptococcus dysgalactiae subsp. equisimilis strain KNZ12 | lacG | 6-phospho-beta-galactosidase (EC 3.2.1.85) | beta-D-galactose 6-phosphate | 99 | 99 | 91 | 1e-269 |
| Streptococcus dysgalactiae subsp. equisimilis strain KNZ15 | lacG | 6-phospho-beta-galactosidase (EC 3.2.1.85) | beta-D-galactose 6-phosphate | 99 | 99 | 91 | 1e-269 |
| Streptococcus dysgalactiae subsp. equisimilis strain KNZ16 | lacG | 6-phospho-beta-galactosidase (EC 3.2.1.85) | beta-D-galactose 6-phosphate | 99 | 99 | 91 | 1e-269 |
| Streptococcus dysgalactiae subsp. equisimilis strain NCTC5370 | lacG | 6-phospho-beta-galactosidase (EC 3.2.1.85) | beta-D-galactose 6-phosphate | 99 | 99 | 91 | 1e-269 |
| Streptococcus dysgalactiae subsp. equisimilis strain NCTC5371 | lacG | 6-phospho-beta-galactosidase (EC 3.2.1.85) | beta-D-galactose 6-phosphate | 99 | 99 | 91 | 1e-269 |
| Streptococcus dysgalactiae subsp. equisimilis strain NCTC8546 | lacG | 6-phospho-beta-galactosidase (EC 3.2.1.85) | beta-D-galactose 6-phosphate | 99 | 99 | 91 | 1e-269 |
| Streptococcus dysgalactiae subsp. equisimilis strain NCTC9414 | lacG | 6-phospho-beta-galactosidase (EC 3.2.1.85) | beta-D-galactose 6-phosphate | 99 | 99 | 91 | 1e-269 |
| Streptococcus dysgalactiae subsp. equisimilis strain UT_4231_KK | lacG | 6-phospho-beta-galactosidase (EC 3.2.1.85) | beta-D-galactose 6-phosphate | 99 | 99 | 91 | 1e-269 |
| Streptococcus dysgalactiae subsp. equisimilis strain UT_4241_XS | lacG | 6-phospho-beta-galactosidase (EC 3.2.1.85) | beta-D-galactose 6-phosphate | 99 | 99 | 91 | 1e-269 |
| Streptococcus dysgalactiae subsp. equisimilis strain UT_4277_BB | lacG | 6-phospho-beta-galactosidase (EC 3.2.1.85) | beta-D-galactose 6-phosphate | 99 | 99 | 92 | 1e-271 |
| Streptococcus dysgalactiae subsp. equisimilis strain UT_4966_RC | lacG | 6-phospho-beta-galactosidase (EC 3.2.1.85) | beta-D-galactose 6-phosphate | 99 | 99 | 91 | 1e-269 |
| Streptococcus dysgalactiae subsp. equisimilis strain UT-SS1069 | lacG | 6-phospho-beta-galactosidase (EC 3.2.1.85) | beta-D-galactose 6-phosphate | 99 | 99 | 91 | 1e-269 |
| Streptococcus dysgalactiae subsp. equisimilis strain WCHSDSE-1 | lacG | 6-phospho-beta-galactosidase (EC 3.2.1.85) | beta-D-galactose 6-phosphate | 99 | 99 | 91 | 1e-269 |
| Streptococcus dysgalactiae subsp. equisimilis SD SCDR1 | murI | Glutamate racemase (EC 5.1.1.3) | (4S)-4-(2-NAPHTHYLMETHYL)-D-GLUTAMIC ACID | 100 | 100 | 85 | 1e-131 |
| Streptococcus dysgalactiae subsp. equisimilis 167 | glr | Glutamate racemase (EC 5.1.1.3) | (4S)-4-(2-NAPHTHYLMETHYL)-D-GLUTAMIC ACID | 100 | 100 | 85 | 1e-131 |
| Streptococcus dysgalactiae subsp. equisimilis AC-2713 | glr | Glutamate racemase (EC 5.1.1.3) | (4S)-4-(2-NAPHTHYLMETHYL)-D-GLUTAMIC ACID | 100 | 100 | 85 | 1e-131 |
| Streptococcus dysgalactiae subsp. equisimilis AKSDE4288 | murI | Glutamate racemase (EC 5.1.1.3) | (4S)-4-(2-NAPHTHYLMETHYL)-D-GLUTAMIC ACID | 100 | 100 | 85 | 1e-131 |
| Streptococcus dysgalactiae subsp. equisimilis ATCC 12394 | murI | Glutamate racemase (EC 5.1.1.3) | (4S)-4-(2-NAPHTHYLMETHYL)-D-GLUTAMIC ACID | 100 | 100 | 85 | 1e-131 |
| Streptococcus dysgalactiae subsp. equisimilis GGS_124 | glr | Glutamate racemase (EC 5.1.1.3) | (4S)-4-(2-NAPHTHYLMETHYL)-D-GLUTAMIC ACID | 100 | 100 | 85 | 1e-131 |
| Streptococcus dysgalactiae subsp. equisimilis RE378 | glr | Glutamate racemase (EC 5.1.1.3) | (4S)-4-(2-NAPHTHYLMETHYL)-D-GLUTAMIC ACID | 100 | 100 | 85 | 1e-131 |
| Streptococcus dysgalactiae subsp. equisimilis SK1249 | murI | Glutamate racemase (EC 5.1.1.3) | (4S)-4-(2-NAPHTHYLMETHYL)-D-GLUTAMIC ACID | 100 | 100 | 85 | 1e-131 |
| Streptococcus dysgalactiae subsp. equisimilis SK1250 | murI | Glutamate racemase (EC 5.1.1.3) | (4S)-4-(2-NAPHTHYLMETHYL)-D-GLUTAMIC ACID | 100 | 100 | 85 | 1e-131 |
| Streptococcus dysgalactiae subsp. equisimilis strain ASDSE_96 | murI | Glutamate racemase (EC 5.1.1.3) | (4S)-4-(2-NAPHTHYLMETHYL)-D-GLUTAMIC ACID | 100 | 100 | 85 | 1e-131 |
| Streptococcus dysgalactiae subsp. equisimilis strain ASDSE_99 | murI | Glutamate racemase (EC 5.1.1.3) | (4S)-4-(2-NAPHTHYLMETHYL)-D-GLUTAMIC ACID | 100 | 100 | 85 | 1e-131 |
| Streptococcus dysgalactiae subsp. equisimilis strain C161L1 | murI | Glutamate racemase (EC 5.1.1.3) | (4S)-4-(2-NAPHTHYLMETHYL)-D-GLUTAMIC ACID | 100 | 100 | 85 | 0 |
| Streptococcus dysgalactiae subsp. equisimilis strain KNZ01 | murI | Glutamate racemase (EC 5.1.1.3) | (4S)-4-(2-NAPHTHYLMETHYL)-D-GLUTAMIC ACID | 100 | 100 | 85 | 1e-131 |
| Streptococcus dysgalactiae subsp. equisimilis strain KNZ03 | murI | Glutamate racemase (EC 5.1.1.3) | (4S)-4-(2-NAPHTHYLMETHYL)-D-GLUTAMIC ACID | 100 | 100 | 85 | 1e-131 |
| Streptococcus dysgalactiae subsp. equisimilis strain KNZ04 | murI | Glutamate racemase (EC 5.1.1.3) | (4S)-4-(2-NAPHTHYLMETHYL)-D-GLUTAMIC ACID | 100 | 100 | 85 | 1e-131 |
| Streptococcus dysgalactiae subsp. equisimilis strain KNZ06 | murI | Glutamate racemase (EC 5.1.1.3) | (4S)-4-(2-NAPHTHYLMETHYL)-D-GLUTAMIC ACID | 100 | 100 | 85 | 1e-131 |
| Streptococcus dysgalactiae subsp. equisimilis strain KNZ07 | murI | Glutamate racemase (EC 5.1.1.3) | (4S)-4-(2-NAPHTHYLMETHYL)-D-GLUTAMIC ACID | 100 | 100 | 85 | 1e-131 |
| Streptococcus dysgalactiae subsp. equisimilis strain KNZ10 | murI | Glutamate racemase (EC 5.1.1.3) | (4S)-4-(2-NAPHTHYLMETHYL)-D-GLUTAMIC ACID | 100 | 100 | 85 | 1e-131 |
| Streptococcus dysgalactiae subsp. equisimilis strain KNZ12 | murI | Glutamate racemase (EC 5.1.1.3) | (4S)-4-(2-NAPHTHYLMETHYL)-D-GLUTAMIC ACID | 100 | 100 | 85 | 1e-131 |
| Streptococcus dysgalactiae subsp. equisimilis strain KNZ15 | murI | Glutamate racemase (EC 5.1.1.3) | (4S)-4-(2-NAPHTHYLMETHYL)-D-GLUTAMIC ACID | 100 | 100 | 85 | 1e-131 |
| Streptococcus dysgalactiae subsp. equisimilis strain KNZ16 | murI | Glutamate racemase (EC 5.1.1.3) | (4S)-4-(2-NAPHTHYLMETHYL)-D-GLUTAMIC ACID | 100 | 100 | 85 | 1e-131 |
| Streptococcus dysgalactiae subsp. equisimilis strain NCTC10321 | murI | Glutamate racemase (EC 5.1.1.3) | (4S)-4-(2-NAPHTHYLMETHYL)-D-GLUTAMIC ACID | 100 | 100 | 85 | 1e-131 |
| Streptococcus dysgalactiae subsp. equisimilis strain NCTC11554 | murI | Glutamate racemase (EC 5.1.1.3) | (4S)-4-(2-NAPHTHYLMETHYL)-D-GLUTAMIC ACID | 100 | 100 | 85 | 1e-131 |
| Streptococcus dysgalactiae subsp. equisimilis strain NCTC11555 | murI | Glutamate racemase (EC 5.1.1.3) | (4S)-4-(2-NAPHTHYLMETHYL)-D-GLUTAMIC ACID | 100 | 100 | 85 | 1e-131 |
| Streptococcus dysgalactiae subsp. equisimilis strain NCTC11556 | murI | Glutamate racemase (EC 5.1.1.3) | (4S)-4-(2-NAPHTHYLMETHYL)-D-GLUTAMIC ACID | 100 | 100 | 85 | 1e-131 |
| Streptococcus dysgalactiae subsp. equisimilis strain NCTC11557 | murI | Glutamate racemase (EC 5.1.1.3) | (4S)-4-(2-NAPHTHYLMETHYL)-D-GLUTAMIC ACID | 100 | 100 | 85 | 1e-131 |
| Streptococcus dysgalactiae subsp. equisimilis strain NCTC11564 | murI | Glutamate racemase (EC 5.1.1.3) | (4S)-4-(2-NAPHTHYLMETHYL)-D-GLUTAMIC ACID | 100 | 100 | 85 | 1e-131 |
| Streptococcus dysgalactiae subsp. equisimilis strain NCTC5370 | murI | Glutamate racemase (EC 5.1.1.3) | (4S)-4-(2-NAPHTHYLMETHYL)-D-GLUTAMIC ACID | 100 | 100 | 85 | 1e-131 |
| Streptococcus dysgalactiae subsp. equisimilis strain NCTC5371 | murI | Glutamate racemase (EC 5.1.1.3) | (4S)-4-(2-NAPHTHYLMETHYL)-D-GLUTAMIC ACID | 100 | 100 | 85 | 1e-131 |
| Streptococcus dysgalactiae subsp. equisimilis strain NCTC6179 | murI | Glutamate racemase (EC 5.1.1.3) | (4S)-4-(2-NAPHTHYLMETHYL)-D-GLUTAMIC ACID | 100 | 100 | 85 | 1e-131 |
| Streptococcus dysgalactiae subsp. equisimilis strain NCTC6181 | murI | Glutamate racemase (EC 5.1.1.3) | (4S)-4-(2-NAPHTHYLMETHYL)-D-GLUTAMIC ACID | 100 | 100 | 85 | 1e-131 |
| Streptococcus dysgalactiae subsp. equisimilis strain NCTC6407 | murI | Glutamate racemase (EC 5.1.1.3) | (4S)-4-(2-NAPHTHYLMETHYL)-D-GLUTAMIC ACID | 100 | 100 | 85 | 1e-131 |
| Streptococcus dysgalactiae subsp. equisimilis strain NCTC7136 | murI | Glutamate racemase (EC 5.1.1.3) | (4S)-4-(2-NAPHTHYLMETHYL)-D-GLUTAMIC ACID | 100 | 100 | 85 | 1e-131 |
| Streptococcus dysgalactiae subsp. equisimilis strain NCTC8543 | murI | Glutamate racemase (EC 5.1.1.3) | (4S)-4-(2-NAPHTHYLMETHYL)-D-GLUTAMIC ACID | 100 | 100 | 85 | 1e-131 |
| Streptococcus dysgalactiae subsp. equisimilis strain NCTC8546 | murI | Glutamate racemase (EC 5.1.1.3) | (4S)-4-(2-NAPHTHYLMETHYL)-D-GLUTAMIC ACID | 100 | 100 | 85 | 1e-131 |
| Streptococcus dysgalactiae subsp. equisimilis strain NCTC9413 | murI | Glutamate racemase (EC 5.1.1.3) | (4S)-4-(2-NAPHTHYLMETHYL)-D-GLUTAMIC ACID | 100 | 100 | 85 | 1e-131 |
| Streptococcus dysgalactiae subsp. equisimilis strain NCTC9414 | murI | Glutamate racemase (EC 5.1.1.3) | (4S)-4-(2-NAPHTHYLMETHYL)-D-GLUTAMIC ACID | 100 | 100 | 85 | 1e-131 |
| Streptococcus dysgalactiae subsp. equisimilis strain NCTC9603 | murI | Glutamate racemase (EC 5.1.1.3) | (4S)-4-(2-NAPHTHYLMETHYL)-D-GLUTAMIC ACID | 100 | 100 | 85 | 1e-131 |
| Streptococcus dysgalactiae subsp. equisimilis strain SS1575 | murI | Glutamate racemase (EC 5.1.1.3) | (4S)-4-(2-NAPHTHYLMETHYL)-D-GLUTAMIC ACID | 100 | 100 | 85 | 1e-131 |
| Streptococcus dysgalactiae subsp. equisimilis strain T642 | murI | Glutamate racemase (EC 5.1.1.3) | (4S)-4-(2-NAPHTHYLMETHYL)-D-GLUTAMIC ACID | 100 | 100 | 85 | 3e-172 |
| Streptococcus dysgalactiae subsp. equisimilis strain UT_4031CC | murI | Glutamate racemase (EC 5.1.1.3) | (4S)-4-(2-NAPHTHYLMETHYL)-D-GLUTAMIC ACID | 100 | 100 | 85 | 1e-130 |
| Streptococcus dysgalactiae subsp. equisimilis strain UT_4231_KK | murI | Glutamate racemase (EC 5.1.1.3) | (4S)-4-(2-NAPHTHYLMETHYL)-D-GLUTAMIC ACID | 100 | 100 | 85 | 1e-131 |
| Streptococcus dysgalactiae subsp. equisimilis strain UT_4234_DH | murI | Glutamate racemase (EC 5.1.1.3) | (4S)-4-(2-NAPHTHYLMETHYL)-D-GLUTAMIC ACID | 100 | 100 | 85 | 1e-131 |
| Streptococcus dysgalactiae subsp. equisimilis strain UT_4241_XS | murI | Glutamate racemase (EC 5.1.1.3) | (4S)-4-(2-NAPHTHYLMETHYL)-D-GLUTAMIC ACID | 100 | 100 | 85 | 1e-131 |
| Streptococcus dysgalactiae subsp. equisimilis strain UT_4242_AB | murI | Glutamate racemase (EC 5.1.1.3) | (4S)-4-(2-NAPHTHYLMETHYL)-D-GLUTAMIC ACID | 100 | 100 | 85 | 1e-131 |
| Streptococcus dysgalactiae subsp. equisimilis strain UT_4255RC | murI | Glutamate racemase (EC 5.1.1.3) | (4S)-4-(2-NAPHTHYLMETHYL)-D-GLUTAMIC ACID | 100 | 100 | 85 | 1e-131 |
| Streptococcus dysgalactiae subsp. equisimilis strain UT_4277_BB | murI | Glutamate racemase (EC 5.1.1.3) | (4S)-4-(2-NAPHTHYLMETHYL)-D-GLUTAMIC ACID | 100 | 100 | 85 | 1e-131 |
| Streptococcus dysgalactiae subsp. equisimilis strain UT_4966_RC | murI | Glutamate racemase (EC 5.1.1.3) | (4S)-4-(2-NAPHTHYLMETHYL)-D-GLUTAMIC ACID | 100 | 100 | 85 | 1e-131 |
| Streptococcus dysgalactiae subsp. equisimilis strain UT-5345 | murI | Glutamate racemase (EC 5.1.1.3) | (4S)-4-(2-NAPHTHYLMETHYL)-D-GLUTAMIC ACID | 100 | 100 | 85 | 1e-131 |
| Streptococcus dysgalactiae subsp. equisimilis strain UT-5354 | murI | Glutamate racemase (EC 5.1.1.3) | (4S)-4-(2-NAPHTHYLMETHYL)-D-GLUTAMIC ACID | 100 | 100 | 85 | 1e-130 |
| Streptococcus dysgalactiae subsp. equisimilis strain UT-SS1069 | murI | Glutamate racemase (EC 5.1.1.3) | (4S)-4-(2-NAPHTHYLMETHYL)-D-GLUTAMIC ACID | 100 | 100 | 85 | 1e-131 |
| Streptococcus dysgalactiae subsp. equisimilis strain UT-SS957 | murI | Glutamate racemase (EC 5.1.1.3) | (4S)-4-(2-NAPHTHYLMETHYL)-D-GLUTAMIC ACID | 100 | 100 | 85 | 1e-131 |
| Streptococcus dysgalactiae subsp. equisimilis strain WCHSDSE-1 | murI | Glutamate racemase (EC 5.1.1.3) | (4S)-4-(2-NAPHTHYLMETHYL)-D-GLUTAMIC ACID | 100 | 100 | 85 | 1e-131 |
| Streptococcus dysgalactiae subsp. equisimilis SD SCDR1 | parE | DNA topoisomerase IV subunit B (EC 5.99.1.3) | Gatifloxacin | 100 | 100 | 88 | 0.0 |
| Streptococcus dysgalactiae subsp. equisimilis 167 | parE | DNA topoisomerase IV subunit B (EC 5.99.1.3) | Gatifloxacin | 100 | 100 | 88 | 0.0 |
| Streptococcus dysgalactiae subsp. equisimilis AC-2713 | gyrB | DNA topoisomerase IV subunit B (EC 5.99.1.3) | Gatifloxacin | 100 | 100 | 88 | 0.0 |
| Streptococcus dysgalactiae subsp. equisimilis AKSDE4288 | parE | DNA topoisomerase IV subunit B (EC 5.99.1.3) | Gatifloxacin | 100 | 100 | 88 | 0.0 |
| Streptococcus dysgalactiae subsp. equisimilis ATCC 12394 | parE | DNA topoisomerase IV subunit B (EC 5.99.1.3) | Gatifloxacin | 100 | 100 | 88 | 0.0 |
| Streptococcus dysgalactiae subsp. equisimilis GGS_124 | parE | DNA topoisomerase IV subunit B (EC 5.99.1.3) | Gatifloxacin | 100 | 100 | 88 | 0.0 |
| Streptococcus dysgalactiae subsp. equisimilis RE378 | parE | DNA topoisomerase IV subunit B (EC 5.99.1.3) | Gatifloxacin | 100 | 100 | 88 | 0.0 |
| Streptococcus dysgalactiae subsp. equisimilis SK1249 | parE | DNA topoisomerase IV subunit B (EC 5.99.1.3) | Gatifloxacin | 100 | 100 | 88 | 0.0 |
| Streptococcus dysgalactiae subsp. equisimilis SK1250 | parE | DNA topoisomerase IV subunit B (EC 5.99.1.3) | Gatifloxacin | 100 | 100 | 88 | 0.0 |
| Streptococcus dysgalactiae subsp. equisimilis strain ASDSE_96 | parE | DNA topoisomerase IV subunit B (EC 5.99.1.3) | Gatifloxacin | 100 | 100 | 88 | 0.0 |
| Streptococcus dysgalactiae subsp. equisimilis strain ASDSE_99 | parE | DNA topoisomerase IV subunit B (EC 5.99.1.3) | Gatifloxacin | 100 | 100 | 88 | 0.0 |
| Streptococcus dysgalactiae subsp. equisimilis strain C161L1 | parE | DNA topoisomerase IV subunit B (EC 5.99.1.3) | Gatifloxacin | 100 | 100 | 88 | 0 |
| Streptococcus dysgalactiae subsp. equisimilis strain KNZ01 | parE | DNA topoisomerase IV subunit B (EC 5.99.1.3) | Gatifloxacin | 100 | 100 | 88 | 0.0 |
| Streptococcus dysgalactiae subsp. equisimilis strain KNZ03 | parE | DNA topoisomerase IV subunit B (EC 5.99.1.3) | Gatifloxacin | 100 | 100 | 88 | 0.0 |
| Streptococcus dysgalactiae subsp. equisimilis strain KNZ04 | parE | DNA topoisomerase IV subunit B (EC 5.99.1.3) | Gatifloxacin | 100 | 100 | 88 | 0.0 |
| Streptococcus dysgalactiae subsp. equisimilis strain KNZ07 | parE | DNA topoisomerase IV subunit B (EC 5.99.1.3) | Gatifloxacin | 100 | 100 | 88 | 0.0 |
| Streptococcus dysgalactiae subsp. equisimilis strain KNZ10 | parE | DNA topoisomerase IV subunit B (EC 5.99.1.3) | Gatifloxacin | 100 | 100 | 88 | 0.0 |
| Streptococcus dysgalactiae subsp. equisimilis strain KNZ12 | parE | DNA topoisomerase IV subunit B (EC 5.99.1.3) | Gatifloxacin | 100 | 100 | 88 | 0.0 |
| Streptococcus dysgalactiae subsp. equisimilis strain KNZ15 | parE | DNA topoisomerase IV subunit B (EC 5.99.1.3) | Gatifloxacin | 100 | 100 | 88 | 0.0 |
| Streptococcus dysgalactiae subsp. equisimilis strain KNZ16 | parE | DNA topoisomerase IV subunit B (EC 5.99.1.3) | Gatifloxacin | 100 | 100 | 88 | 0.0 |
| Streptococcus dysgalactiae subsp. equisimilis strain NCTC10321 | parE | DNA topoisomerase IV subunit B (EC 5.99.1.3) | Gatifloxacin | 100 | 100 | 88 | 0.0 |
| Streptococcus dysgalactiae subsp. equisimilis strain NCTC11554 | parE | DNA topoisomerase IV subunit B (EC 5.99.1.3) | Gatifloxacin | 100 | 100 | 88 | 0.0 |
| Streptococcus dysgalactiae subsp. equisimilis strain NCTC11555 | parE | DNA topoisomerase IV subunit B (EC 5.99.1.3) | Gatifloxacin | 100 | 100 | 88 | 0.0 |
| Streptococcus dysgalactiae subsp. equisimilis strain NCTC11556 | parE | DNA topoisomerase IV subunit B (EC 5.99.1.3) | Gatifloxacin | 100 | 100 | 88 | 0.0 |
| Streptococcus dysgalactiae subsp. equisimilis strain NCTC11557 | parE | DNA topoisomerase IV subunit B (EC 5.99.1.3) | Gatifloxacin | 100 | 100 | 88 | 0.0 |
| Streptococcus dysgalactiae subsp. equisimilis strain NCTC11564 | parE | DNA topoisomerase IV subunit B (EC 5.99.1.3) | Gatifloxacin | 100 | 100 | 88 | 0.0 |
| Streptococcus dysgalactiae subsp. equisimilis strain NCTC5370 | parE | DNA topoisomerase IV subunit B (EC 5.99.1.3) | Gatifloxacin | 100 | 100 | 88 | 0.0 |
| Streptococcus dysgalactiae subsp. equisimilis strain NCTC5371 | parE | DNA topoisomerase IV subunit B (EC 5.99.1.3) | Gatifloxacin | 100 | 100 | 88 | 0.0 |
| Streptococcus dysgalactiae subsp. equisimilis strain NCTC6179 | parE | DNA topoisomerase IV subunit B (EC 5.99.1.3) | Gatifloxacin | 100 | 100 | 88 | 0.0 |
| Streptococcus dysgalactiae subsp. equisimilis strain NCTC6181 | parE | DNA topoisomerase IV subunit B (EC 5.99.1.3) | Gatifloxacin | 100 | 100 | 88 | 0.0 |
| Streptococcus dysgalactiae subsp. equisimilis strain NCTC6407 | parE | DNA topoisomerase IV subunit B (EC 5.99.1.3) | Gatifloxacin | 100 | 100 | 88 | 0.0 |
| Streptococcus dysgalactiae subsp. equisimilis strain NCTC8543 | parE | DNA topoisomerase IV subunit B (EC 5.99.1.3) | Gatifloxacin | 100 | 100 | 88 | 0.0 |
| Streptococcus dysgalactiae subsp. equisimilis strain NCTC8546 | parE | DNA topoisomerase IV subunit B (EC 5.99.1.3) | Gatifloxacin | 100 | 100 | 88 | 0.0 |
| Streptococcus dysgalactiae subsp. equisimilis strain NCTC9413 | parE | DNA topoisomerase IV subunit B (EC 5.99.1.3) | Gatifloxacin | 100 | 100 | 88 | 0.0 |
| Streptococcus dysgalactiae subsp. equisimilis strain NCTC9414 | parE | DNA topoisomerase IV subunit B (EC 5.99.1.3) | Gatifloxacin | 100 | 100 | 88 | 0.0 |
| Streptococcus dysgalactiae subsp. equisimilis strain SS1575 | parE | DNA topoisomerase IV subunit B (EC 5.99.1.3) | Gatifloxacin | 100 | 100 | 88 | 0.0 |
| Streptococcus dysgalactiae subsp. equisimilis strain T642 | parE | DNA topoisomerase IV subunit B (EC 5.99.1.3) | Gatifloxacin | 100 | 100 | 87 | 0.0 |
| Streptococcus dysgalactiae subsp. equisimilis strain UT_4031CC | parE | DNA topoisomerase IV subunit B (EC 5.99.1.3) | Gatifloxacin | 100 | 100 | 88 | 0.0 |
| Streptococcus dysgalactiae subsp. equisimilis strain UT_4231_KK | parE | DNA topoisomerase IV subunit B (EC 5.99.1.3) | Gatifloxacin | 100 | 100 | 88 | 0.0 |
| Streptococcus dysgalactiae subsp. equisimilis strain UT_4234_DH | parE | DNA topoisomerase IV subunit B (EC 5.99.1.3) | Gatifloxacin | 100 | 100 | 88 | 0.0 |
| Streptococcus dysgalactiae subsp. equisimilis strain UT_4241_XS | parE | DNA topoisomerase IV subunit B (EC 5.99.1.3) | Gatifloxacin | 100 | 100 | 88 | 0.0 |
| Streptococcus dysgalactiae subsp. equisimilis strain UT_4242_AB | parE | DNA topoisomerase IV subunit B (EC 5.99.1.3) | Gatifloxacin | 100 | 100 | 88 | 0.0 |
| Streptococcus dysgalactiae subsp. equisimilis strain UT_4255RC | parE | DNA topoisomerase IV subunit B (EC 5.99.1.3) | Gatifloxacin | 100 | 100 | 88 | 0.0 |
| Streptococcus dysgalactiae subsp. equisimilis strain UT_4277_BB | parE | DNA topoisomerase IV subunit B (EC 5.99.1.3) | Gatifloxacin | 100 | 100 | 88 | 0.0 |
| Streptococcus dysgalactiae subsp. equisimilis strain UT_4966_RC | parE | DNA topoisomerase IV subunit B (EC 5.99.1.3) | Gatifloxacin | 100 | 100 | 88 | 0.0 |
| Streptococcus dysgalactiae subsp. equisimilis strain UT-5345 | gyrB | DNA topoisomerase IV subunit B (EC 5.99.1.3) | Gatifloxacin | 100 | 100 | 88 | 0.0 |
| Streptococcus dysgalactiae subsp. equisimilis strain UT-5354 | gyrB | DNA topoisomerase IV subunit B (EC 5.99.1.3) | Gatifloxacin | 100 | 100 | 88 | 0.0 |
| Streptococcus dysgalactiae subsp. equisimilis strain UT-SS1069 | gyrB | DNA topoisomerase IV subunit B (EC 5.99.1.3) | Gatifloxacin | 100 | 100 | 88 | 0.0 |
| Streptococcus dysgalactiae subsp. equisimilis strain UT-SS957 | gyrB | DNA topoisomerase IV subunit B (EC 5.99.1.3) | Gatifloxacin | 100 | 100 | 88 | 0.0 |
| Streptococcus dysgalactiae subsp. equisimilis strain WCHSDSE-1 | parE | DNA topoisomerase IV subunit B (EC 5.99.1.3) | Gatifloxacin | 100 | 100 | 88 | 0.0 |
| Streptococcus dysgalactiae subsp. equisimilis SD SCDR1 | relA | Guanosine-3',5'-bis(diphosphate) 3'-pyrophosphohydrolase (EC 3.1.7.2) / GTP pyrophosphokinase (EC 2.7.6.5), (p)ppGpp synthetase II | Guanosine 5'-Diphosphate 2':3'-Cyclic Monophosphate;Guanosine-5'-Diphosphate | 100 | 100 | 99 | 0.0 |
| Streptococcus dysgalactiae subsp. equisimilis 167 | relA | Guanosine-3',5'-bis(diphosphate) 3'-pyrophosphohydrolase (EC 3.1.7.2) / GTP pyrophosphokinase (EC 2.7.6.5), (p)ppGpp synthetase II | Guanosine 5'-Diphosphate 2':3'-Cyclic Monophosphate;Guanosine-5'-Diphosphate | 100 | 100 | 99 | 0.0 |
| Streptococcus dysgalactiae subsp. equisimilis AC-2713 | relA | Guanosine-3',5'-bis(diphosphate) 3'-pyrophosphohydrolase (EC 3.1.7.2) / GTP pyrophosphokinase (EC 2.7.6.5), (p)ppGpp synthetase II | Guanosine 5'-Diphosphate 2':3'-Cyclic Monophosphate;Guanosine-5'-Diphosphate | 100 | 100 | 99 | 0.0 |
| Streptococcus dysgalactiae subsp. equisimilis AKSDE4288 | relA | Guanosine-3',5'-bis(diphosphate) 3'-pyrophosphohydrolase (EC 3.1.7.2) / GTP pyrophosphokinase (EC 2.7.6.5), (p)ppGpp synthetase II | Guanosine 5'-Diphosphate 2':3'-Cyclic Monophosphate;Guanosine-5'-Diphosphate | 100 | 100 | 99 | 0.0 |
| Streptococcus dysgalactiae subsp. equisimilis ATCC 12394 | relA | Guanosine-3',5'-bis(diphosphate) 3'-pyrophosphohydrolase (EC 3.1.7.2) / GTP pyrophosphokinase (EC 2.7.6.5), (p)ppGpp synthetase II | Guanosine 5'-Diphosphate 2':3'-Cyclic Monophosphate;Guanosine-5'-Diphosphate | 100 | 100 | 99 | 0.0 |
| Streptococcus dysgalactiae subsp. equisimilis GGS_124 | relA | Guanosine-3',5'-bis(diphosphate) 3'-pyrophosphohydrolase (EC 3.1.7.2) / GTP pyrophosphokinase (EC 2.7.6.5), (p)ppGpp synthetase II | Guanosine 5'-Diphosphate 2':3'-Cyclic Monophosphate;Guanosine-5'-Diphosphate | 100 | 100 | 99 | 0.0 |
| Streptococcus dysgalactiae subsp. equisimilis RE378 | relA | Guanosine-3',5'-bis(diphosphate) 3'-pyrophosphohydrolase (EC 3.1.7.2) / GTP pyrophosphokinase (EC 2.7.6.5), (p)ppGpp synthetase II | Guanosine 5'-Diphosphate 2':3'-Cyclic Monophosphate;Guanosine-5'-Diphosphate | 100 | 100 | 99 | 0.0 |
| Streptococcus dysgalactiae subsp. equisimilis SK1249 | relA | Guanosine-3',5'-bis(diphosphate) 3'-pyrophosphohydrolase (EC 3.1.7.2) / GTP pyrophosphokinase (EC 2.7.6.5), (p)ppGpp synthetase II | Guanosine 5'-Diphosphate 2':3'-Cyclic Monophosphate;Guanosine-5'-Diphosphate | 100 | 100 | 99 | 0.0 |
| Streptococcus dysgalactiae subsp. equisimilis SK1250 | relA | Guanosine-3',5'-bis(diphosphate) 3'-pyrophosphohydrolase (EC 3.1.7.2) / GTP pyrophosphokinase (EC 2.7.6.5), (p)ppGpp synthetase II | Guanosine 5'-Diphosphate 2':3'-Cyclic Monophosphate;Guanosine-5'-Diphosphate | 100 | 100 | 99 | 0.0 |
| Streptococcus dysgalactiae subsp. equisimilis strain ASDSE_96 | relA | Guanosine-3',5'-bis(diphosphate) 3'-pyrophosphohydrolase (EC 3.1.7.2) / GTP pyrophosphokinase (EC 2.7.6.5), (p)ppGpp synthetase II | Guanosine 5'-Diphosphate 2':3'-Cyclic Monophosphate;Guanosine-5'-Diphosphate | 100 | 100 | 99 | 0.0 |
| Streptococcus dysgalactiae subsp. equisimilis strain ASDSE_99 | relA | Guanosine-3',5'-bis(diphosphate) 3'-pyrophosphohydrolase (EC 3.1.7.2) / GTP pyrophosphokinase (EC 2.7.6.5), (p)ppGpp synthetase II | Guanosine 5'-Diphosphate 2':3'-Cyclic Monophosphate;Guanosine-5'-Diphosphate | 100 | 100 | 99 | 0.0 |
| Streptococcus dysgalactiae subsp. equisimilis strain C161L1 | relA | Guanosine-3',5'-bis(diphosphate) 3'-pyrophosphohydrolase (EC 3.1.7.2) / GTP pyrophosphokinase (EC 2.7.6.5), (p)ppGpp synthetase II | Guanosine 5'-Diphosphate 2':3'-Cyclic Monophosphate;Guanosine-5'-Diphosphate | 100 | 100 | 99 | 0 |
| Streptococcus dysgalactiae subsp. equisimilis strain KNZ01 | relA | Guanosine-3',5'-bis(diphosphate) 3'-pyrophosphohydrolase (EC 3.1.7.2) / GTP pyrophosphokinase (EC 2.7.6.5), (p)ppGpp synthetase II | Guanosine 5'-Diphosphate 2':3'-Cyclic Monophosphate;Guanosine-5'-Diphosphate | 100 | 100 | 99 | 0.0 |
| Streptococcus dysgalactiae subsp. equisimilis strain KNZ03 | relA | Guanosine-3',5'-bis(diphosphate) 3'-pyrophosphohydrolase (EC 3.1.7.2) / GTP pyrophosphokinase (EC 2.7.6.5), (p)ppGpp synthetase II | Guanosine 5'-Diphosphate 2':3'-Cyclic Monophosphate;Guanosine-5'-Diphosphate | 100 | 100 | 99 | 0.0 |
| Streptococcus dysgalactiae subsp. equisimilis strain KNZ04 | relA | Guanosine-3',5'-bis(diphosphate) 3'-pyrophosphohydrolase (EC 3.1.7.2) / GTP pyrophosphokinase (EC 2.7.6.5), (p)ppGpp synthetase II | Guanosine 5'-Diphosphate 2':3'-Cyclic Monophosphate;Guanosine-5'-Diphosphate | 100 | 100 | 99 | 0.0 |
| Streptococcus dysgalactiae subsp. equisimilis strain KNZ06 | relA | Guanosine-3',5'-bis(diphosphate) 3'-pyrophosphohydrolase (EC 3.1.7.2) / GTP pyrophosphokinase (EC 2.7.6.5), (p)ppGpp synthetase II | Guanosine 5'-Diphosphate 2':3'-Cyclic Monophosphate;Guanosine-5'-Diphosphate | 100 | 100 | 99 | 0.0 |
| Streptococcus dysgalactiae subsp. equisimilis strain KNZ07 | relA | Guanosine-3',5'-bis(diphosphate) 3'-pyrophosphohydrolase (EC 3.1.7.2) / GTP pyrophosphokinase (EC 2.7.6.5), (p)ppGpp synthetase II | Guanosine 5'-Diphosphate 2':3'-Cyclic Monophosphate;Guanosine-5'-Diphosphate | 100 | 100 | 99 | 0.0 |
| Streptococcus dysgalactiae subsp. equisimilis strain KNZ10 | relA | Guanosine-3',5'-bis(diphosphate) 3'-pyrophosphohydrolase (EC 3.1.7.2) / GTP pyrophosphokinase (EC 2.7.6.5), (p)ppGpp synthetase II | Guanosine 5'-Diphosphate 2':3'-Cyclic Monophosphate;Guanosine-5'-Diphosphate | 100 | 100 | 99 | 0.0 |
| Streptococcus dysgalactiae subsp. equisimilis strain KNZ12 | relA | Guanosine-3',5'-bis(diphosphate) 3'-pyrophosphohydrolase (EC 3.1.7.2) / GTP pyrophosphokinase (EC 2.7.6.5), (p)ppGpp synthetase II | Guanosine 5'-Diphosphate 2':3'-Cyclic Monophosphate;Guanosine-5'-Diphosphate | 100 | 100 | 99 | 0.0 |
| Streptococcus dysgalactiae subsp. equisimilis strain KNZ15 | relA | Guanosine-3',5'-bis(diphosphate) 3'-pyrophosphohydrolase (EC 3.1.7.2) / GTP pyrophosphokinase (EC 2.7.6.5), (p)ppGpp synthetase II | Guanosine 5'-Diphosphate 2':3'-Cyclic Monophosphate;Guanosine-5'-Diphosphate | 100 | 100 | 99 | 0.0 |
| Streptococcus dysgalactiae subsp. equisimilis strain KNZ16 | relA | Guanosine-3',5'-bis(diphosphate) 3'-pyrophosphohydrolase (EC 3.1.7.2) / GTP pyrophosphokinase (EC 2.7.6.5), (p)ppGpp synthetase II | Guanosine 5'-Diphosphate 2':3'-Cyclic Monophosphate;Guanosine-5'-Diphosphate | 88 | 73 | 99 | 0.0 |
| Streptococcus dysgalactiae subsp. equisimilis strain NCTC10321 | relA | Guanosine-3',5'-bis(diphosphate) 3'-pyrophosphohydrolase (EC 3.1.7.2) / GTP pyrophosphokinase (EC 2.7.6.5), (p)ppGpp synthetase II | Guanosine 5'-Diphosphate 2':3'-Cyclic Monophosphate;Guanosine-5'-Diphosphate | 100 | 100 | 97 | 0.0 |
| Streptococcus dysgalactiae subsp. equisimilis strain NCTC11554 | relA | Guanosine-3',5'-bis(diphosphate) 3'-pyrophosphohydrolase (EC 3.1.7.2) / GTP pyrophosphokinase (EC 2.7.6.5), (p)ppGpp synthetase II | Guanosine 5'-Diphosphate 2':3'-Cyclic Monophosphate;Guanosine-5'-Diphosphate | 100 | 100 | 99 | 0.0 |
| Streptococcus dysgalactiae subsp. equisimilis strain NCTC11555 | relA | Guanosine-3',5'-bis(diphosphate) 3'-pyrophosphohydrolase (EC 3.1.7.2) / GTP pyrophosphokinase (EC 2.7.6.5), (p)ppGpp synthetase II | Guanosine 5'-Diphosphate 2':3'-Cyclic Monophosphate;Guanosine-5'-Diphosphate | 100 | 100 | 99 | 0.0 |
| Streptococcus dysgalactiae subsp. equisimilis strain NCTC11556 | relA | Guanosine-3',5'-bis(diphosphate) 3'-pyrophosphohydrolase (EC 3.1.7.2) / GTP pyrophosphokinase (EC 2.7.6.5), (p)ppGpp synthetase II | Guanosine 5'-Diphosphate 2':3'-Cyclic Monophosphate;Guanosine-5'-Diphosphate | 100 | 100 | 99 | 0.0 |
| Streptococcus dysgalactiae subsp. equisimilis strain NCTC11557 | relA | Guanosine-3',5'-bis(diphosphate) 3'-pyrophosphohydrolase (EC 3.1.7.2) / GTP pyrophosphokinase (EC 2.7.6.5), (p)ppGpp synthetase II | Guanosine 5'-Diphosphate 2':3'-Cyclic Monophosphate;Guanosine-5'-Diphosphate | 100 | 100 | 99 | 0.0 |
| Streptococcus dysgalactiae subsp. equisimilis strain NCTC11564 | relA | Guanosine-3',5'-bis(diphosphate) 3'-pyrophosphohydrolase (EC 3.1.7.2) / GTP pyrophosphokinase (EC 2.7.6.5), (p)ppGpp synthetase II | Guanosine 5'-Diphosphate 2':3'-Cyclic Monophosphate;Guanosine-5'-Diphosphate | 100 | 100 | 99 | 0.0 |
| Streptococcus dysgalactiae subsp. equisimilis strain NCTC5370 | relA | Guanosine-3',5'-bis(diphosphate) 3'-pyrophosphohydrolase (EC 3.1.7.2) / GTP pyrophosphokinase (EC 2.7.6.5), (p)ppGpp synthetase II | Guanosine 5'-Diphosphate 2':3'-Cyclic Monophosphate;Guanosine-5'-Diphosphate | 100 | 100 | 99 | 0.0 |
| Streptococcus dysgalactiae subsp. equisimilis strain NCTC5371 | relA | Guanosine-3',5'-bis(diphosphate) 3'-pyrophosphohydrolase (EC 3.1.7.2) / GTP pyrophosphokinase (EC 2.7.6.5), (p)ppGpp synthetase II | Guanosine 5'-Diphosphate 2':3'-Cyclic Monophosphate;Guanosine-5'-Diphosphate | 100 | 100 | 99 | 0.0 |
| Streptococcus dysgalactiae subsp. equisimilis strain NCTC6179 | relA | Guanosine-3',5'-bis(diphosphate) 3'-pyrophosphohydrolase (EC 3.1.7.2) / GTP pyrophosphokinase (EC 2.7.6.5), (p)ppGpp synthetase II | Guanosine 5'-Diphosphate 2':3'-Cyclic Monophosphate;Guanosine-5'-Diphosphate | 90 | 95 | 99 | 0.0 |
| Streptococcus dysgalactiae subsp. equisimilis strain NCTC6181 | relA | Guanosine-3',5'-bis(diphosphate) 3'-pyrophosphohydrolase (EC 3.1.7.2) / GTP pyrophosphokinase (EC 2.7.6.5), (p)ppGpp synthetase II | Guanosine 5'-Diphosphate 2':3'-Cyclic Monophosphate;Guanosine-5'-Diphosphate | 100 | 100 | 99 | 0.0 |
| Streptococcus dysgalactiae subsp. equisimilis strain NCTC6407 | relA | Guanosine-3',5'-bis(diphosphate) 3'-pyrophosphohydrolase (EC 3.1.7.2) / GTP pyrophosphokinase (EC 2.7.6.5), (p)ppGpp synthetase II | Guanosine 5'-Diphosphate 2':3'-Cyclic Monophosphate;Guanosine-5'-Diphosphate | 100 | 100 | 97 | 0.0 |
| Streptococcus dysgalactiae subsp. equisimilis strain NCTC7136 | relA | Guanosine-3',5'-bis(diphosphate) 3'-pyrophosphohydrolase (EC 3.1.7.2) / GTP pyrophosphokinase (EC 2.7.6.5), (p)ppGpp synthetase II | Guanosine 5'-Diphosphate 2':3'-Cyclic Monophosphate;Guanosine-5'-Diphosphate | 100 | 100 | 99 | 0.0 |
| Streptococcus dysgalactiae subsp. equisimilis strain NCTC8543 | relA | Guanosine-3',5'-bis(diphosphate) 3'-pyrophosphohydrolase (EC 3.1.7.2) / GTP pyrophosphokinase (EC 2.7.6.5), (p)ppGpp synthetase II | Guanosine 5'-Diphosphate 2':3'-Cyclic Monophosphate;Guanosine-5'-Diphosphate | 100 | 100 | 99 | 0.0 |
| Streptococcus dysgalactiae subsp. equisimilis strain NCTC8546 | relA | Guanosine-3',5'-bis(diphosphate) 3'-pyrophosphohydrolase (EC 3.1.7.2) / GTP pyrophosphokinase (EC 2.7.6.5), (p)ppGpp synthetase II | Guanosine 5'-Diphosphate 2':3'-Cyclic Monophosphate;Guanosine-5'-Diphosphate | 100 | 100 | 100 | 0.0 |
| Streptococcus dysgalactiae subsp. equisimilis strain NCTC9413 | relA | Guanosine-3',5'-bis(diphosphate) 3'-pyrophosphohydrolase (EC 3.1.7.2) / GTP pyrophosphokinase (EC 2.7.6.5), (p)ppGpp synthetase II | Guanosine 5'-Diphosphate 2':3'-Cyclic Monophosphate;Guanosine-5'-Diphosphate | 100 | 100 | 99 | 0.0 |
| Streptococcus dysgalactiae subsp. equisimilis strain NCTC9414 | relA | Guanosine-3',5'-bis(diphosphate) 3'-pyrophosphohydrolase (EC 3.1.7.2) / GTP pyrophosphokinase (EC 2.7.6.5), (p)ppGpp synthetase II | Guanosine 5'-Diphosphate 2':3'-Cyclic Monophosphate;Guanosine-5'-Diphosphate | 100 | 100 | 99 | 0.0 |
| Streptococcus dysgalactiae subsp. equisimilis strain NCTC9603 | relA | Guanosine-3',5'-bis(diphosphate) 3'-pyrophosphohydrolase (EC 3.1.7.2) / GTP pyrophosphokinase (EC 2.7.6.5), (p)ppGpp synthetase II | Guanosine 5'-Diphosphate 2':3'-Cyclic Monophosphate;Guanosine-5'-Diphosphate | 100 | 100 | 99 | 0.0 |
| Streptococcus dysgalactiae subsp. equisimilis strain SS1575 | relA | Guanosine-3',5'-bis(diphosphate) 3'-pyrophosphohydrolase (EC 3.1.7.2) / GTP pyrophosphokinase (EC 2.7.6.5), (p)ppGpp synthetase II | Guanosine 5'-Diphosphate 2':3'-Cyclic Monophosphate;Guanosine-5'-Diphosphate | 100 | 100 | 99 | 0.0 |
| Streptococcus dysgalactiae subsp. equisimilis strain T642 | relA | Guanosine-3',5'-bis(diphosphate) 3'-pyrophosphohydrolase (EC 3.1.7.2) / GTP pyrophosphokinase (EC 2.7.6.5), (p)ppGpp synthetase II | Guanosine 5'-Diphosphate 2':3'-Cyclic Monophosphate;Guanosine-5'-Diphosphate | 100 | 100 | 99 | 0.0 |
| Streptococcus dysgalactiae subsp. equisimilis strain UT_4031CC | relA | Guanosine-3',5'-bis(diphosphate) 3'-pyrophosphohydrolase (EC 3.1.7.2) / GTP pyrophosphokinase (EC 2.7.6.5), (p)ppGpp synthetase II | Guanosine 5'-Diphosphate 2':3'-Cyclic Monophosphate;Guanosine-5'-Diphosphate | 100 | 100 | 99 | 0.0 |
| Streptococcus dysgalactiae subsp. equisimilis strain UT_4231_KK | relA | Guanosine-3',5'-bis(diphosphate) 3'-pyrophosphohydrolase (EC 3.1.7.2) / GTP pyrophosphokinase (EC 2.7.6.5), (p)ppGpp synthetase II | Guanosine 5'-Diphosphate 2':3'-Cyclic Monophosphate;Guanosine-5'-Diphosphate | 100 | 100 | 99 | 0.0 |
| Streptococcus dysgalactiae subsp. equisimilis strain UT_4234_DH | relA | Guanosine-3',5'-bis(diphosphate) 3'-pyrophosphohydrolase (EC 3.1.7.2) / GTP pyrophosphokinase (EC 2.7.6.5), (p)ppGpp synthetase II | Guanosine 5'-Diphosphate 2':3'-Cyclic Monophosphate;Guanosine-5'-Diphosphate | 100 | 100 | 99 | 0.0 |
| Streptococcus dysgalactiae subsp. equisimilis strain UT_4241_XS | relA | Guanosine-3',5'-bis(diphosphate) 3'-pyrophosphohydrolase (EC 3.1.7.2) / GTP pyrophosphokinase (EC 2.7.6.5), (p)ppGpp synthetase II | Guanosine 5'-Diphosphate 2':3'-Cyclic Monophosphate;Guanosine-5'-Diphosphate | 100 | 100 | 99 | 0.0 |
| Streptococcus dysgalactiae subsp. equisimilis strain UT_4242_AB | relA | Guanosine-3',5'-bis(diphosphate) 3'-pyrophosphohydrolase (EC 3.1.7.2) / GTP pyrophosphokinase (EC 2.7.6.5), (p)ppGpp synthetase II | Guanosine 5'-Diphosphate 2':3'-Cyclic Monophosphate;Guanosine-5'-Diphosphate | 100 | 100 | 99 | 0.0 |
| Streptococcus dysgalactiae subsp. equisimilis strain UT_4255RC | relA | Guanosine-3',5'-bis(diphosphate) 3'-pyrophosphohydrolase (EC 3.1.7.2) / GTP pyrophosphokinase (EC 2.7.6.5), (p)ppGpp synthetase II | Guanosine 5'-Diphosphate 2':3'-Cyclic Monophosphate;Guanosine-5'-Diphosphate | 100 | 100 | 99 | 0.0 |
| Streptococcus dysgalactiae subsp. equisimilis strain UT_4277_BB | relA | Guanosine-3',5'-bis(diphosphate) 3'-pyrophosphohydrolase (EC 3.1.7.2) / GTP pyrophosphokinase (EC 2.7.6.5), (p)ppGpp synthetase II | Guanosine 5'-Diphosphate 2':3'-Cyclic Monophosphate;Guanosine-5'-Diphosphate | 100 | 100 | 99 | 0.0 |
| Streptococcus dysgalactiae subsp. equisimilis strain UT_4966_RC | relA | Guanosine-3',5'-bis(diphosphate) 3'-pyrophosphohydrolase (EC 3.1.7.2) / GTP pyrophosphokinase (EC 2.7.6.5), (p)ppGpp synthetase II | Guanosine 5'-Diphosphate 2':3'-Cyclic Monophosphate;Guanosine-5'-Diphosphate | 100 | 100 | 99 | 0.0 |
| Streptococcus dysgalactiae subsp. equisimilis strain UT-5345 | relA | Guanosine-3',5'-bis(diphosphate) 3'-pyrophosphohydrolase (EC 3.1.7.2) / GTP pyrophosphokinase (EC 2.7.6.5), (p)ppGpp synthetase II | Guanosine 5'-Diphosphate 2':3'-Cyclic Monophosphate;Guanosine-5'-Diphosphate | 100 | 100 | 99 | 0.0 |
| Streptococcus dysgalactiae subsp. equisimilis strain UT-5354 | relA | Guanosine-3',5'-bis(diphosphate) 3'-pyrophosphohydrolase (EC 3.1.7.2) / GTP pyrophosphokinase (EC 2.7.6.5), (p)ppGpp synthetase II | Guanosine 5'-Diphosphate 2':3'-Cyclic Monophosphate;Guanosine-5'-Diphosphate | 100 | 100 | 99 | 0.0 |
| Streptococcus dysgalactiae subsp. equisimilis strain UT-SS1069 | relA | Guanosine-3',5'-bis(diphosphate) 3'-pyrophosphohydrolase (EC 3.1.7.2) / GTP pyrophosphokinase (EC 2.7.6.5), (p)ppGpp synthetase II | Guanosine 5'-Diphosphate 2':3'-Cyclic Monophosphate;Guanosine-5'-Diphosphate | 100 | 100 | 99 | 0.0 |
| Streptococcus dysgalactiae subsp. equisimilis strain UT-SS957 | relA | Guanosine-3',5'-bis(diphosphate) 3'-pyrophosphohydrolase (EC 3.1.7.2) / GTP pyrophosphokinase (EC 2.7.6.5), (p)ppGpp synthetase II | Guanosine 5'-Diphosphate 2':3'-Cyclic Monophosphate;Guanosine-5'-Diphosphate | 100 | 100 | 99 | 0.0 |
| Streptococcus dysgalactiae subsp. equisimilis strain WCHSDSE-1 | relA | Guanosine-3',5'-bis(diphosphate) 3'-pyrophosphohydrolase (EC 3.1.7.2) / GTP pyrophosphokinase (EC 2.7.6.5), (p)ppGpp synthetase II | Guanosine 5'-Diphosphate 2':3'-Cyclic Monophosphate;Guanosine-5'-Diphosphate | 100 | 100 | 99 | 0.0 |
| Streptococcus dysgalactiae subsp. equisimilis SD SCDR1 | rmlB | dTDP-glucose 4,6-dehydratase (EC 4.2.1.46) | Thymidine-5'-Diphospho-Beta-D-Xylose;2'deoxy-Thymidine-5'-Diphospho-Alpha-D-Glucose | 97 | 98 | 94 | 1e-196 |
| Streptococcus dysgalactiae subsp. equisimilis 167 | cpsFQ | dTDP-glucose 4,6-dehydratase (EC 4.2.1.46) | Thymidine-5'-Diphospho-Beta-D-Xylose;2'deoxy-Thymidine-5'-Diphospho-Alpha-D-Glucose | 97 | 98 | 94 | 1e-195 |
| Streptococcus dysgalactiae subsp. equisimilis AC-2713 | rmlB | dTDP-glucose 4,6-dehydratase (EC 4.2.1.46) | Thymidine-5'-Diphospho-Beta-D-Xylose;2'deoxy-Thymidine-5'-Diphospho-Alpha-D-Glucose | 97 | 98 | 94 | 1e-195 |
| Streptococcus dysgalactiae subsp. equisimilis AKSDE4288 | rmlB | dTDP-glucose 4,6-dehydratase (EC 4.2.1.46) | Thymidine-5'-Diphospho-Beta-D-Xylose;2'deoxy-Thymidine-5'-Diphospho-Alpha-D-Glucose | 97 | 98 | 94 | 1e-195 |
| Streptococcus dysgalactiae subsp. equisimilis ATCC 12394 | rmlB | dTDP-glucose 4,6-dehydratase (EC 4.2.1.46) | Thymidine-5'-Diphospho-Beta-D-Xylose;2'deoxy-Thymidine-5'-Diphospho-Alpha-D-Glucose | 97 | 98 | 93 | 1e-195 |
| Streptococcus dysgalactiae subsp. equisimilis GGS_124 | cpsFQ | dTDP-glucose 4,6-dehydratase (EC 4.2.1.46) | Thymidine-5'-Diphospho-Beta-D-Xylose;2'deoxy-Thymidine-5'-Diphospho-Alpha-D-Glucose | 97 | 98 | 95 | 1e-197 |
| Streptococcus dysgalactiae subsp. equisimilis RE378 | cpsFQ | dTDP-glucose 4,6-dehydratase (EC 4.2.1.46) | Thymidine-5'-Diphospho-Beta-D-Xylose;2'deoxy-Thymidine-5'-Diphospho-Alpha-D-Glucose | 97 | 98 | 94 | 1e-195 |
| Streptococcus dysgalactiae subsp. equisimilis SK1249 | rffG | dTDP-glucose 4,6-dehydratase (EC 4.2.1.46) | Thymidine-5'-Diphospho-Beta-D-Xylose;2'deoxy-Thymidine-5'-Diphospho-Alpha-D-Glucose | 97 | 98 | 94 | 1e-195 |
| Streptococcus dysgalactiae subsp. equisimilis SK1250 | rfbB | dTDP-glucose 4,6-dehydratase (EC 4.2.1.46) | Thymidine-5'-Diphospho-Beta-D-Xylose;2'deoxy-Thymidine-5'-Diphospho-Alpha-D-Glucose | 97 | 98 | 94 | 1e-196 |
| Streptococcus dysgalactiae subsp. equisimilis strain ASDSE_96 | rmlB | dTDP-glucose 4,6-dehydratase (EC 4.2.1.46) | Thymidine-5'-Diphospho-Beta-D-Xylose;2'deoxy-Thymidine-5'-Diphospho-Alpha-D-Glucose | 97 | 98 | 94 | 1e-195 |
| Streptococcus dysgalactiae subsp. equisimilis strain ASDSE_99 | rmlB | dTDP-glucose 4,6-dehydratase (EC 4.2.1.46) | Thymidine-5'-Diphospho-Beta-D-Xylose;2'deoxy-Thymidine-5'-Diphospho-Alpha-D-Glucose | 97 | 98 | 94 | 1e-195 |
| Streptococcus dysgalactiae subsp. equisimilis strain C161L1 | rmlB | dTDP-glucose 4,6-dehydratase (EC 4.2.1.46) | Thymidine-5'-Diphospho-Beta-D-Xylose;2'deoxy-Thymidine-5'-Diphospho-Alpha-D-Glucose | 98 | 99 | 93 | 0 |
| Streptococcus dysgalactiae subsp. equisimilis strain KNZ01 | rmlB | dTDP-glucose 4,6-dehydratase (EC 4.2.1.46) | Thymidine-5'-Diphospho-Beta-D-Xylose;2'deoxy-Thymidine-5'-Diphospho-Alpha-D-Glucose | 97 | 98 | 94 | 1e-195 |
| Streptococcus dysgalactiae subsp. equisimilis strain KNZ03 | rmlB | dTDP-glucose 4,6-dehydratase (EC 4.2.1.46) | Thymidine-5'-Diphospho-Beta-D-Xylose;2'deoxy-Thymidine-5'-Diphospho-Alpha-D-Glucose | 97 | 98 | 93 | 1e-195 |
| Streptococcus dysgalactiae subsp. equisimilis strain KNZ04 | rmlB | dTDP-glucose 4,6-dehydratase (EC 4.2.1.46) | Thymidine-5'-Diphospho-Beta-D-Xylose;2'deoxy-Thymidine-5'-Diphospho-Alpha-D-Glucose | 97 | 98 | 93 | 1e-195 |
| Streptococcus dysgalactiae subsp. equisimilis strain KNZ06 | rmlB | dTDP-glucose 4,6-dehydratase (EC 4.2.1.46) | Thymidine-5'-Diphospho-Beta-D-Xylose;2'deoxy-Thymidine-5'-Diphospho-Alpha-D-Glucose | 97 | 98 | 94 | 1e-195 |
| Streptococcus dysgalactiae subsp. equisimilis strain KNZ07 | rmlB | dTDP-glucose 4,6-dehydratase (EC 4.2.1.46) | Thymidine-5'-Diphospho-Beta-D-Xylose;2'deoxy-Thymidine-5'-Diphospho-Alpha-D-Glucose | 97 | 98 | 94 | 1e-195 |
| Streptococcus dysgalactiae subsp. equisimilis strain KNZ10 | rmlB | dTDP-glucose 4,6-dehydratase (EC 4.2.1.46) | Thymidine-5'-Diphospho-Beta-D-Xylose;2'deoxy-Thymidine-5'-Diphospho-Alpha-D-Glucose | 97 | 98 | 94 | 1e-195 |
| Streptococcus dysgalactiae subsp. equisimilis strain KNZ12 | rmlB | dTDP-glucose 4,6-dehydratase (EC 4.2.1.46) | Thymidine-5'-Diphospho-Beta-D-Xylose;2'deoxy-Thymidine-5'-Diphospho-Alpha-D-Glucose | 97 | 98 | 94 | 1e-195 |
| Streptococcus dysgalactiae subsp. equisimilis strain KNZ15 | rmlB | dTDP-glucose 4,6-dehydratase (EC 4.2.1.46) | Thymidine-5'-Diphospho-Beta-D-Xylose;2'deoxy-Thymidine-5'-Diphospho-Alpha-D-Glucose | 97 | 98 | 94 | 1e-195 |
| Streptococcus dysgalactiae subsp. equisimilis strain KNZ16 | rmlB | dTDP-glucose 4,6-dehydratase (EC 4.2.1.46) | Thymidine-5'-Diphospho-Beta-D-Xylose;2'deoxy-Thymidine-5'-Diphospho-Alpha-D-Glucose | 97 | 98 | 93 | 1e-195 |
| Streptococcus dysgalactiae subsp. equisimilis strain NCTC10321 | rmlB | dTDP-glucose 4,6-dehydratase (EC 4.2.1.46) | Thymidine-5'-Diphospho-Beta-D-Xylose;2'deoxy-Thymidine-5'-Diphospho-Alpha-D-Glucose | 97 | 98 | 93 | 1e-195 |
| Streptococcus dysgalactiae subsp. equisimilis strain NCTC11554 | rmlB | dTDP-glucose 4,6-dehydratase (EC 4.2.1.46) | Thymidine-5'-Diphospho-Beta-D-Xylose;2'deoxy-Thymidine-5'-Diphospho-Alpha-D-Glucose | 97 | 98 | 93 | 1e-195 |
| Streptococcus dysgalactiae subsp. equisimilis strain NCTC11555 | rmlB | dTDP-glucose 4,6-dehydratase (EC 4.2.1.46) | Thymidine-5'-Diphospho-Beta-D-Xylose;2'deoxy-Thymidine-5'-Diphospho-Alpha-D-Glucose | 97 | 98 | 94 | 1e-195 |
| Streptococcus dysgalactiae subsp. equisimilis strain NCTC11556 | rmlB | dTDP-glucose 4,6-dehydratase (EC 4.2.1.46) | Thymidine-5'-Diphospho-Beta-D-Xylose;2'deoxy-Thymidine-5'-Diphospho-Alpha-D-Glucose | 97 | 98 | 93 | 1e-195 |
| Streptococcus dysgalactiae subsp. equisimilis strain NCTC11557 | rmlB | dTDP-glucose 4,6-dehydratase (EC 4.2.1.46) | Thymidine-5'-Diphospho-Beta-D-Xylose;2'deoxy-Thymidine-5'-Diphospho-Alpha-D-Glucose | 97 | 98 | 94 | 1e-195 |
| Streptococcus dysgalactiae subsp. equisimilis strain NCTC11564 | rmlB | dTDP-glucose 4,6-dehydratase (EC 4.2.1.46) | Thymidine-5'-Diphospho-Beta-D-Xylose;2'deoxy-Thymidine-5'-Diphospho-Alpha-D-Glucose | 97 | 98 | 94 | 1e-195 |
| Streptococcus dysgalactiae subsp. equisimilis strain NCTC5370 | rmlB | dTDP-glucose 4,6-dehydratase (EC 4.2.1.46) | Thymidine-5'-Diphospho-Beta-D-Xylose;2'deoxy-Thymidine-5'-Diphospho-Alpha-D-Glucose | 97 | 98 | 94 | 1e-195 |
| Streptococcus dysgalactiae subsp. equisimilis strain NCTC5371 | rmlB | dTDP-glucose 4,6-dehydratase (EC 4.2.1.46) | Thymidine-5'-Diphospho-Beta-D-Xylose;2'deoxy-Thymidine-5'-Diphospho-Alpha-D-Glucose | 97 | 98 | 94 | 1e-195 |
| Streptococcus dysgalactiae subsp. equisimilis strain NCTC6179 | rmlB | dTDP-glucose 4,6-dehydratase (EC 4.2.1.46) | Thymidine-5'-Diphospho-Beta-D-Xylose;2'deoxy-Thymidine-5'-Diphospho-Alpha-D-Glucose | 97 | 98 | 93 | 1e-195 |
| Streptococcus dysgalactiae subsp. equisimilis strain NCTC6181 | rmlB | dTDP-glucose 4,6-dehydratase (EC 4.2.1.46) | Thymidine-5'-Diphospho-Beta-D-Xylose;2'deoxy-Thymidine-5'-Diphospho-Alpha-D-Glucose | 97 | 98 | 93 | 1e-195 |
| Streptococcus dysgalactiae subsp. equisimilis strain NCTC6407 | rmlB | dTDP-glucose 4,6-dehydratase (EC 4.2.1.46) | Thymidine-5'-Diphospho-Beta-D-Xylose;2'deoxy-Thymidine-5'-Diphospho-Alpha-D-Glucose | 97 | 98 | 93 | 1e-195 |
| Streptococcus dysgalactiae subsp. equisimilis strain NCTC7136 | rmlB | dTDP-glucose 4,6-dehydratase (EC 4.2.1.46) | Thymidine-5'-Diphospho-Beta-D-Xylose;2'deoxy-Thymidine-5'-Diphospho-Alpha-D-Glucose | 97 | 98 | 94 | 1e-195 |
| Streptococcus dysgalactiae subsp. equisimilis strain NCTC8543 | rmlB | dTDP-glucose 4,6-dehydratase (EC 4.2.1.46) | Thymidine-5'-Diphospho-Beta-D-Xylose;2'deoxy-Thymidine-5'-Diphospho-Alpha-D-Glucose | 97 | 98 | 94 | 1e-195 |
| Streptococcus dysgalactiae subsp. equisimilis strain NCTC8546 | rmlB | dTDP-glucose 4,6-dehydratase (EC 4.2.1.46) | Thymidine-5'-Diphospho-Beta-D-Xylose;2'deoxy-Thymidine-5'-Diphospho-Alpha-D-Glucose | 97 | 98 | 94 | 1e-195 |
| Streptococcus dysgalactiae subsp. equisimilis strain NCTC9413 | rmlB | dTDP-glucose 4,6-dehydratase (EC 4.2.1.46) | Thymidine-5'-Diphospho-Beta-D-Xylose;2'deoxy-Thymidine-5'-Diphospho-Alpha-D-Glucose | 97 | 98 | 94 | 1e-195 |
| Streptococcus dysgalactiae subsp. equisimilis strain NCTC9414 | rmlB | dTDP-glucose 4,6-dehydratase (EC 4.2.1.46) | Thymidine-5'-Diphospho-Beta-D-Xylose;2'deoxy-Thymidine-5'-Diphospho-Alpha-D-Glucose | 97 | 98 | 94 | 1e-195 |
| Streptococcus dysgalactiae subsp. equisimilis strain NCTC9603 | rmlB | dTDP-glucose 4,6-dehydratase (EC 4.2.1.46) | Thymidine-5'-Diphospho-Beta-D-Xylose;2'deoxy-Thymidine-5'-Diphospho-Alpha-D-Glucose | 97 | 98 | 94 | 1e-195 |
| Streptococcus dysgalactiae subsp. equisimilis strain SS1575 | rmlB | dTDP-glucose 4,6-dehydratase (EC 4.2.1.46) | Thymidine-5'-Diphospho-Beta-D-Xylose;2'deoxy-Thymidine-5'-Diphospho-Alpha-D-Glucose | 97 | 98 | 94 | 1e-195 |
| Streptococcus dysgalactiae subsp. equisimilis strain T642 | rmlB | dTDP-glucose 4,6-dehydratase (EC 4.2.1.46) | Thymidine-5'-Diphospho-Beta-D-Xylose;2'deoxy-Thymidine-5'-Diphospho-Alpha-D-Glucose | 98 | 99 | 93 | 0 |
| Streptococcus dysgalactiae subsp. equisimilis strain UT_4031CC | rmlB | dTDP-glucose 4,6-dehydratase (EC 4.2.1.46) | Thymidine-5'-Diphospho-Beta-D-Xylose;2'deoxy-Thymidine-5'-Diphospho-Alpha-D-Glucose | 97 | 98 | 94 | 1e-195 |
| Streptococcus dysgalactiae subsp. equisimilis strain UT_4231_KK | rmlB | dTDP-glucose 4,6-dehydratase (EC 4.2.1.46) | Thymidine-5'-Diphospho-Beta-D-Xylose;2'deoxy-Thymidine-5'-Diphospho-Alpha-D-Glucose | 97 | 98 | 94 | 1e-195 |
| Streptococcus dysgalactiae subsp. equisimilis strain UT_4234_DH | rmlB | dTDP-glucose 4,6-dehydratase (EC 4.2.1.46) | Thymidine-5'-Diphospho-Beta-D-Xylose;2'deoxy-Thymidine-5'-Diphospho-Alpha-D-Glucose | 97 | 98 | 94 | 1e-195 |
| Streptococcus dysgalactiae subsp. equisimilis strain UT_4241_XS | rmlB | dTDP-glucose 4,6-dehydratase (EC 4.2.1.46) | Thymidine-5'-Diphospho-Beta-D-Xylose;2'deoxy-Thymidine-5'-Diphospho-Alpha-D-Glucose | 97 | 98 | 94 | 1e-195 |
| Streptococcus dysgalactiae subsp. equisimilis strain UT_4242_AB | rmlB | dTDP-glucose 4,6-dehydratase (EC 4.2.1.46) | Thymidine-5'-Diphospho-Beta-D-Xylose;2'deoxy-Thymidine-5'-Diphospho-Alpha-D-Glucose | 97 | 98 | 93 | 1e-195 |
| Streptococcus dysgalactiae subsp. equisimilis strain UT_4255RC | rmlB | dTDP-glucose 4,6-dehydratase (EC 4.2.1.46) | Thymidine-5'-Diphospho-Beta-D-Xylose;2'deoxy-Thymidine-5'-Diphospho-Alpha-D-Glucose | 97 | 98 | 94 | 1e-195 |
| Streptococcus dysgalactiae subsp. equisimilis strain UT_4277_BB | rmlB | dTDP-glucose 4,6-dehydratase (EC 4.2.1.46) | Thymidine-5'-Diphospho-Beta-D-Xylose;2'deoxy-Thymidine-5'-Diphospho-Alpha-D-Glucose | 97 | 98 | 95 | 1e-198 |
| Streptococcus dysgalactiae subsp. equisimilis strain UT_4966_RC | rmlB | dTDP-glucose 4,6-dehydratase (EC 4.2.1.46) | Thymidine-5'-Diphospho-Beta-D-Xylose;2'deoxy-Thymidine-5'-Diphospho-Alpha-D-Glucose | 97 | 98 | 94 | 1e-195 |
| Streptococcus dysgalactiae subsp. equisimilis strain UT-5345 | rmlB | dTDP-glucose 4,6-dehydratase (EC 4.2.1.46) | Thymidine-5'-Diphospho-Beta-D-Xylose;2'deoxy-Thymidine-5'-Diphospho-Alpha-D-Glucose | 97 | 98 | 94 | 1e-196 |
| Streptococcus dysgalactiae subsp. equisimilis strain UT-5354 | rmlB | dTDP-glucose 4,6-dehydratase (EC 4.2.1.46) | Thymidine-5'-Diphospho-Beta-D-Xylose;2'deoxy-Thymidine-5'-Diphospho-Alpha-D-Glucose | 97 | 98 | 94 | 1e-195 |
| Streptococcus dysgalactiae subsp. equisimilis strain UT-SS1069 | rmlB | dTDP-glucose 4,6-dehydratase (EC 4.2.1.46) | Thymidine-5'-Diphospho-Beta-D-Xylose;2'deoxy-Thymidine-5'-Diphospho-Alpha-D-Glucose | 97 | 98 | 94 | 1e-195 |
| Streptococcus dysgalactiae subsp. equisimilis strain UT-SS957 | rmlB | dTDP-glucose 4,6-dehydratase (EC 4.2.1.46) | Thymidine-5'-Diphospho-Beta-D-Xylose;2'deoxy-Thymidine-5'-Diphospho-Alpha-D-Glucose | 97 | 98 | 94 | 1e-195 |
| Streptococcus dysgalactiae subsp. equisimilis strain WCHSDSE-1 | rmlB | dTDP-glucose 4,6-dehydratase (EC 4.2.1.46) | Thymidine-5'-Diphospho-Beta-D-Xylose;2'deoxy-Thymidine-5'-Diphospho-Alpha-D-Glucose | 97 | 98 | 94 | 1e-196 |
| Streptococcus dysgalactiae subsp. equisimilis SD SCDR1 | rmlC | dTDP-4-dehydrorhamnose 3,5-epimerase (EC 5.1.3.13) | Thymidine-5'-Diphospho-Beta-D-Xylose;2'deoxy-Thymidine-5'-Diphospho-Alpha-D-Glucose | 100 | 100 | 83 | 4e-95 |
| Streptococcus dysgalactiae subsp. equisimilis 167 | cpsFP | dTDP-4-dehydrorhamnose 3,5-epimerase (EC 5.1.3.13) | Thymidine-5'-Diphospho-Beta-D-Xylose;2'deoxy-Thymidine-5'-Diphospho-Alpha-D-Glucose | 53 | 96 | 82 | 4e-46 |
| Streptococcus dysgalactiae subsp. equisimilis AC-2713 | rmlC | dTDP-4-dehydrorhamnose 3,5-epimerase (EC 5.1.3.13) | Thymidine-5'-Diphospho-Beta-D-Xylose;2'deoxy-Thymidine-5'-Diphospho-Alpha-D-Glucose | 100 | 100 | 82 | 5e-95 |
| Streptococcus dysgalactiae subsp. equisimilis AKSDE4288 | rmlC | dTDP-4-dehydrorhamnose 3,5-epimerase (EC 5.1.3.13) | Thymidine-5'-Diphospho-Beta-D-Xylose;2'deoxy-Thymidine-5'-Diphospho-Alpha-D-Glucose | 100 | 100 | 83 | 4e-95 |
| Streptococcus dysgalactiae subsp. equisimilis ATCC 12394 | rmlC | dTDP-4-dehydrorhamnose 3,5-epimerase (EC 5.1.3.13) | Thymidine-5'-Diphospho-Beta-D-Xylose;2'deoxy-Thymidine-5'-Diphospho-Alpha-D-Glucose | 100 | 100 | 82 | 1e-94 |
| Streptococcus dysgalactiae subsp. equisimilis GGS_124 | cpsFP | dTDP-4-dehydrorhamnose 3,5-epimerase (EC 5.1.3.13) | Thymidine-5'-Diphospho-Beta-D-Xylose;2'deoxy-Thymidine-5'-Diphospho-Alpha-D-Glucose | 100 | 100 | 83 | 4e-95 |
| Streptococcus dysgalactiae subsp. equisimilis RE378 | cpsFP | dTDP-4-dehydrorhamnose 3,5-epimerase (EC 5.1.3.13) | Thymidine-5'-Diphospho-Beta-D-Xylose;2'deoxy-Thymidine-5'-Diphospho-Alpha-D-Glucose | 100 | 100 | 83 | 4e-95 |
| Streptococcus dysgalactiae subsp. equisimilis SK1249 | rmlC | dTDP-4-dehydrorhamnose 3,5-epimerase (EC 5.1.3.13) | Thymidine-5'-Diphospho-Beta-D-Xylose;2'deoxy-Thymidine-5'-Diphospho-Alpha-D-Glucose | 100 | 100 | 83 | 4e-95 |
| Streptococcus dysgalactiae subsp. equisimilis SK1250 | rmlC | dTDP-4-dehydrorhamnose 3,5-epimerase (EC 5.1.3.13) | Thymidine-5'-Diphospho-Beta-D-Xylose;2'deoxy-Thymidine-5'-Diphospho-Alpha-D-Glucose | 100 | 100 | 83 | 4e-95 |
| Streptococcus dysgalactiae subsp. equisimilis strain ASDSE_96 | rmlC | dTDP-4-dehydrorhamnose 3,5-epimerase (EC 5.1.3.13) | Thymidine-5'-Diphospho-Beta-D-Xylose;2'deoxy-Thymidine-5'-Diphospho-Alpha-D-Glucose | 100 | 100 | 82 | 1e-94 |
| Streptococcus dysgalactiae subsp. equisimilis strain ASDSE_99 | rmlC | dTDP-4-dehydrorhamnose 3,5-epimerase (EC 5.1.3.13) | Thymidine-5'-Diphospho-Beta-D-Xylose;2'deoxy-Thymidine-5'-Diphospho-Alpha-D-Glucose | 100 | 100 | 83 | 4e-95 |
| Streptococcus dysgalactiae subsp. equisimilis strain C161L1 | rmlC | dTDP-4-dehydrorhamnose 3,5-epimerase (EC 5.1.3.13) | Thymidine-5'-Diphospho-Beta-D-Xylose;2'deoxy-Thymidine-5'-Diphospho-Alpha-D-Glucose | 100 | 100 | 82 | 0 |
| Streptococcus dysgalactiae subsp. equisimilis strain KNZ01 | rmlC | dTDP-4-dehydrorhamnose 3,5-epimerase (EC 5.1.3.13) | Thymidine-5'-Diphospho-Beta-D-Xylose;2'deoxy-Thymidine-5'-Diphospho-Alpha-D-Glucose | 100 | 100 | 83 | 4e-95 |
| Streptococcus dysgalactiae subsp. equisimilis strain KNZ03 | rmlC | dTDP-4-dehydrorhamnose 3,5-epimerase (EC 5.1.3.13) | Thymidine-5'-Diphospho-Beta-D-Xylose;2'deoxy-Thymidine-5'-Diphospho-Alpha-D-Glucose | 100 | 100 | 82 | 1e-94 |
| Streptococcus dysgalactiae subsp. equisimilis strain KNZ04 | rmlC | dTDP-4-dehydrorhamnose 3,5-epimerase (EC 5.1.3.13) | Thymidine-5'-Diphospho-Beta-D-Xylose;2'deoxy-Thymidine-5'-Diphospho-Alpha-D-Glucose | 100 | 100 | 82 | 1e-94 |
| Streptococcus dysgalactiae subsp. equisimilis strain KNZ06 | rmlC | dTDP-4-dehydrorhamnose 3,5-epimerase (EC 5.1.3.13) | Thymidine-5'-Diphospho-Beta-D-Xylose;2'deoxy-Thymidine-5'-Diphospho-Alpha-D-Glucose | 100 | 100 | 83 | 4e-95 |
| Streptococcus dysgalactiae subsp. equisimilis strain KNZ07 | rmlC | dTDP-4-dehydrorhamnose 3,5-epimerase (EC 5.1.3.13) | Thymidine-5'-Diphospho-Beta-D-Xylose;2'deoxy-Thymidine-5'-Diphospho-Alpha-D-Glucose | 100 | 100 | 83 | 4e-95 |
| Streptococcus dysgalactiae subsp. equisimilis strain KNZ10 | rmlC | dTDP-4-dehydrorhamnose 3,5-epimerase (EC 5.1.3.13) | Thymidine-5'-Diphospho-Beta-D-Xylose;2'deoxy-Thymidine-5'-Diphospho-Alpha-D-Glucose | 100 | 100 | 83 | 4e-95 |
| Streptococcus dysgalactiae subsp. equisimilis strain KNZ12 | rmlC | dTDP-4-dehydrorhamnose 3,5-epimerase (EC 5.1.3.13) | Thymidine-5'-Diphospho-Beta-D-Xylose;2'deoxy-Thymidine-5'-Diphospho-Alpha-D-Glucose | 100 | 100 | 83 | 4e-95 |
| Streptococcus dysgalactiae subsp. equisimilis strain KNZ15 | rmlC | dTDP-4-dehydrorhamnose 3,5-epimerase (EC 5.1.3.13) | Thymidine-5'-Diphospho-Beta-D-Xylose;2'deoxy-Thymidine-5'-Diphospho-Alpha-D-Glucose | 100 | 100 | 83 | 4e-95 |
| Streptococcus dysgalactiae subsp. equisimilis strain KNZ16 | rmlC | dTDP-4-dehydrorhamnose 3,5-epimerase (EC 5.1.3.13) | Thymidine-5'-Diphospho-Beta-D-Xylose;2'deoxy-Thymidine-5'-Diphospho-Alpha-D-Glucose | 100 | 100 | 82 | 1e-94 |
| Streptococcus dysgalactiae subsp. equisimilis strain NCTC10321 | rmlC | dTDP-4-dehydrorhamnose 3,5-epimerase (EC 5.1.3.13) | Thymidine-5'-Diphospho-Beta-D-Xylose;2'deoxy-Thymidine-5'-Diphospho-Alpha-D-Glucose | 100 | 100 | 83 | 2e-95 |
| Streptococcus dysgalactiae subsp. equisimilis strain NCTC11554 | rmlC | dTDP-4-dehydrorhamnose 3,5-epimerase (EC 5.1.3.13) | Thymidine-5'-Diphospho-Beta-D-Xylose;2'deoxy-Thymidine-5'-Diphospho-Alpha-D-Glucose | 100 | 100 | 82 | 1e-94 |
| Streptococcus dysgalactiae subsp. equisimilis strain NCTC11555 | rmlC | dTDP-4-dehydrorhamnose 3,5-epimerase (EC 5.1.3.13) | Thymidine-5'-Diphospho-Beta-D-Xylose;2'deoxy-Thymidine-5'-Diphospho-Alpha-D-Glucose | 100 | 100 | 83 | 4e-95 |
| Streptococcus dysgalactiae subsp. equisimilis strain NCTC11556 | rmlC | dTDP-4-dehydrorhamnose 3,5-epimerase (EC 5.1.3.13) | Thymidine-5'-Diphospho-Beta-D-Xylose;2'deoxy-Thymidine-5'-Diphospho-Alpha-D-Glucose | 100 | 100 | 82 | 1e-94 |
| Streptococcus dysgalactiae subsp. equisimilis strain NCTC11557 | rmlC | dTDP-4-dehydrorhamnose 3,5-epimerase (EC 5.1.3.13) | Thymidine-5'-Diphospho-Beta-D-Xylose;2'deoxy-Thymidine-5'-Diphospho-Alpha-D-Glucose | 100 | 100 | 83 | 4e-95 |
| Streptococcus dysgalactiae subsp. equisimilis strain NCTC11564 | rmlC | dTDP-4-dehydrorhamnose 3,5-epimerase (EC 5.1.3.13) | Thymidine-5'-Diphospho-Beta-D-Xylose;2'deoxy-Thymidine-5'-Diphospho-Alpha-D-Glucose | 100 | 100 | 83 | 4e-95 |
| Streptococcus dysgalactiae subsp. equisimilis strain NCTC5370 | rmlC | dTDP-4-dehydrorhamnose 3,5-epimerase (EC 5.1.3.13) | Thymidine-5'-Diphospho-Beta-D-Xylose;2'deoxy-Thymidine-5'-Diphospho-Alpha-D-Glucose | 100 | 100 | 82 | 1e-94 |
| Streptococcus dysgalactiae subsp. equisimilis strain NCTC5371 | rmlC | dTDP-4-dehydrorhamnose 3,5-epimerase (EC 5.1.3.13) | Thymidine-5'-Diphospho-Beta-D-Xylose;2'deoxy-Thymidine-5'-Diphospho-Alpha-D-Glucose | 100 | 100 | 83 | 4e-95 |
| Streptococcus dysgalactiae subsp. equisimilis strain NCTC6179 | rmlC | dTDP-4-dehydrorhamnose 3,5-epimerase (EC 5.1.3.13) | Thymidine-5'-Diphospho-Beta-D-Xylose;2'deoxy-Thymidine-5'-Diphospho-Alpha-D-Glucose | 100 | 100 | 83 | 2e-95 |
| Streptococcus dysgalactiae subsp. equisimilis strain NCTC6181 | rmlC | dTDP-4-dehydrorhamnose 3,5-epimerase (EC 5.1.3.13) | Thymidine-5'-Diphospho-Beta-D-Xylose;2'deoxy-Thymidine-5'-Diphospho-Alpha-D-Glucose | 100 | 100 | 83 | 2e-95 |
| Streptococcus dysgalactiae subsp. equisimilis strain NCTC6407 | rmlC | dTDP-4-dehydrorhamnose 3,5-epimerase (EC 5.1.3.13) | Thymidine-5'-Diphospho-Beta-D-Xylose;2'deoxy-Thymidine-5'-Diphospho-Alpha-D-Glucose | 100 | 100 | 83 | 2e-95 |
| Streptococcus dysgalactiae subsp. equisimilis strain NCTC7136 | rmlC | dTDP-4-dehydrorhamnose 3,5-epimerase (EC 5.1.3.13) | Thymidine-5'-Diphospho-Beta-D-Xylose;2'deoxy-Thymidine-5'-Diphospho-Alpha-D-Glucose | 100 | 100 | 83 | 4e-95 |
| Streptococcus dysgalactiae subsp. equisimilis strain NCTC8543 | rmlC | dTDP-4-dehydrorhamnose 3,5-epimerase (EC 5.1.3.13) | Thymidine-5'-Diphospho-Beta-D-Xylose;2'deoxy-Thymidine-5'-Diphospho-Alpha-D-Glucose | 100 | 100 | 82 | 1e-94 |
| Streptococcus dysgalactiae subsp. equisimilis strain NCTC8546 | rmlC | dTDP-4-dehydrorhamnose 3,5-epimerase (EC 5.1.3.13) | Thymidine-5'-Diphospho-Beta-D-Xylose;2'deoxy-Thymidine-5'-Diphospho-Alpha-D-Glucose | 100 | 100 | 83 | 4e-95 |
| Streptococcus dysgalactiae subsp. equisimilis strain NCTC9413 | rmlC | dTDP-4-dehydrorhamnose 3,5-epimerase (EC 5.1.3.13) | Thymidine-5'-Diphospho-Beta-D-Xylose;2'deoxy-Thymidine-5'-Diphospho-Alpha-D-Glucose | 100 | 100 | 83 | 4e-95 |
| Streptococcus dysgalactiae subsp. equisimilis strain NCTC9414 | rmlC | dTDP-4-dehydrorhamnose 3,5-epimerase (EC 5.1.3.13) | Thymidine-5'-Diphospho-Beta-D-Xylose;2'deoxy-Thymidine-5'-Diphospho-Alpha-D-Glucose | 100 | 100 | 83 | 4e-95 |
| Streptococcus dysgalactiae subsp. equisimilis strain NCTC9603 | rmlC | dTDP-4-dehydrorhamnose 3,5-epimerase (EC 5.1.3.13) | Thymidine-5'-Diphospho-Beta-D-Xylose;2'deoxy-Thymidine-5'-Diphospho-Alpha-D-Glucose | 100 | 100 | 83 | 4e-95 |
| Streptococcus dysgalactiae subsp. equisimilis strain SS1575 | rmlC | dTDP-4-dehydrorhamnose 3,5-epimerase (EC 5.1.3.13) | Thymidine-5'-Diphospho-Beta-D-Xylose;2'deoxy-Thymidine-5'-Diphospho-Alpha-D-Glucose | 100 | 100 | 83 | 4e-95 |
| Streptococcus dysgalactiae subsp. equisimilis strain T642 | rmlC | dTDP-4-dehydrorhamnose 3,5-epimerase (EC 5.1.3.13) | Thymidine-5'-Diphospho-Beta-D-Xylose;2'deoxy-Thymidine-5'-Diphospho-Alpha-D-Glucose | 100 | 100 | 83 | 0 |
| Streptococcus dysgalactiae subsp. equisimilis strain UT_4031CC | rmlC | dTDP-4-dehydrorhamnose 3,5-epimerase (EC 5.1.3.13) | Thymidine-5'-Diphospho-Beta-D-Xylose;2'deoxy-Thymidine-5'-Diphospho-Alpha-D-Glucose | 100 | 100 | 83 | 4e-95 |
| Streptococcus dysgalactiae subsp. equisimilis strain UT_4231_KK | rmlC | dTDP-4-dehydrorhamnose 3,5-epimerase (EC 5.1.3.13) | Thymidine-5'-Diphospho-Beta-D-Xylose;2'deoxy-Thymidine-5'-Diphospho-Alpha-D-Glucose | 100 | 100 | 82 | 1e-94 |
| Streptococcus dysgalactiae subsp. equisimilis strain UT_4234_DH | rmlC | dTDP-4-dehydrorhamnose 3,5-epimerase (EC 5.1.3.13) | Thymidine-5'-Diphospho-Beta-D-Xylose;2'deoxy-Thymidine-5'-Diphospho-Alpha-D-Glucose | 100 | 100 | 83 | 4e-95 |
| Streptococcus dysgalactiae subsp. equisimilis strain UT_4241_XS | rmlC | dTDP-4-dehydrorhamnose 3,5-epimerase (EC 5.1.3.13) | Thymidine-5'-Diphospho-Beta-D-Xylose;2'deoxy-Thymidine-5'-Diphospho-Alpha-D-Glucose | 100 | 100 | 82 | 1e-94 |
| Streptococcus dysgalactiae subsp. equisimilis strain UT_4242_AB | rmlC | dTDP-4-dehydrorhamnose 3,5-epimerase (EC 5.1.3.13) | Thymidine-5'-Diphospho-Beta-D-Xylose;2'deoxy-Thymidine-5'-Diphospho-Alpha-D-Glucose | 100 | 100 | 82 | 1e-94 |
| Streptococcus dysgalactiae subsp. equisimilis strain UT_4255RC | rmlC | dTDP-4-dehydrorhamnose 3,5-epimerase (EC 5.1.3.13) | Thymidine-5'-Diphospho-Beta-D-Xylose;2'deoxy-Thymidine-5'-Diphospho-Alpha-D-Glucose | 100 | 100 | 83 | 4e-95 |
| Streptococcus dysgalactiae subsp. equisimilis strain UT_4277_BB | rmlC | dTDP-4-dehydrorhamnose 3,5-epimerase (EC 5.1.3.13) | Thymidine-5'-Diphospho-Beta-D-Xylose;2'deoxy-Thymidine-5'-Diphospho-Alpha-D-Glucose | 100 | 100 | 82 | 1e-94 |
| Streptococcus dysgalactiae subsp. equisimilis strain UT_4966_RC | rmlC | dTDP-4-dehydrorhamnose 3,5-epimerase (EC 5.1.3.13) | Thymidine-5'-Diphospho-Beta-D-Xylose;2'deoxy-Thymidine-5'-Diphospho-Alpha-D-Glucose | 100 | 100 | 82 | 1e-94 |
| Streptococcus dysgalactiae subsp. equisimilis strain UT-5345 | rmlC | dTDP-4-dehydrorhamnose 3,5-epimerase (EC 5.1.3.13) | Thymidine-5'-Diphospho-Beta-D-Xylose;2'deoxy-Thymidine-5'-Diphospho-Alpha-D-Glucose | 100 | 100 | 83 | 4e-95 |
| Streptococcus dysgalactiae subsp. equisimilis strain UT-5354 | rmlC | dTDP-4-dehydrorhamnose 3,5-epimerase (EC 5.1.3.13) | Thymidine-5'-Diphospho-Beta-D-Xylose;2'deoxy-Thymidine-5'-Diphospho-Alpha-D-Glucose | 100 | 100 | 82 | 1e-94 |
| Streptococcus dysgalactiae subsp. equisimilis strain UT-SS1069 | rmlC | dTDP-4-dehydrorhamnose 3,5-epimerase (EC 5.1.3.13) | Thymidine-5'-Diphospho-Beta-D-Xylose;2'deoxy-Thymidine-5'-Diphospho-Alpha-D-Glucose | 77 | 100 | 84 | 1e-72 |
| Streptococcus dysgalactiae subsp. equisimilis strain UT-SS957 | rmlC | dTDP-4-dehydrorhamnose 3,5-epimerase (EC 5.1.3.13) | Thymidine-5'-Diphospho-Beta-D-Xylose;2'deoxy-Thymidine-5'-Diphospho-Alpha-D-Glucose | 100 | 100 | 83 | 4e-95 |
| Streptococcus dysgalactiae subsp. equisimilis strain WCHSDSE-1 | rmlC | dTDP-4-dehydrorhamnose 3,5-epimerase (EC 5.1.3.13) | Thymidine-5'-Diphospho-Beta-D-Xylose;2'deoxy-Thymidine-5'-Diphospho-Alpha-D-Glucose | 100 | 100 | 83 | 4e-95 |
| Streptococcus dysgalactiae subsp. equisimilis strain NCTC10321 | rplC | LSU ribosomal protein L3p (L3e) | Retapamulin;Spiramycin | 100 | 100 | 94 | 1e-110 |
| Streptococcus dysgalactiae subsp. equisimilis strain NCTC11554 | rplC | LSU ribosomal protein L3p (L3e) | Retapamulin;Spiramycin | 100 | 100 | 99 | 1e-114 |
| Streptococcus dysgalactiae subsp. equisimilis strain NCTC11555 | rplC | LSU ribosomal protein L3p (L3e) | Retapamulin;Spiramycin | 100 | 100 | 99 | 1e-114 |
| Streptococcus dysgalactiae subsp. equisimilis strain NCTC11556 | rplC | LSU ribosomal protein L3p (L3e) | Retapamulin;Spiramycin | 100 | 100 | 99 | 1e-114 |
| Streptococcus dysgalactiae subsp. equisimilis strain NCTC11557 | rplC | LSU ribosomal protein L3p (L3e) | Retapamulin;Spiramycin | 100 | 100 | 99 | 1e-114 |
| Streptococcus dysgalactiae subsp. equisimilis strain NCTC11564 | rplC | LSU ribosomal protein L3p (L3e) | Retapamulin;Spiramycin | 100 | 100 | 99 | 1e-114 |
| Streptococcus dysgalactiae subsp. equisimilis strain NCTC5370 | rplC | LSU ribosomal protein L3p (L3e) | Retapamulin;Spiramycin | 100 | 100 | 99 | 1e-114 |
| Streptococcus dysgalactiae subsp. equisimilis strain NCTC5371 | rplC | LSU ribosomal protein L3p (L3e) | Retapamulin;Spiramycin | 100 | 100 | 99 | 1e-114 |
| Streptococcus dysgalactiae subsp. equisimilis strain NCTC6179 | rplC | LSU ribosomal protein L3p (L3e) | Retapamulin;Spiramycin | 100 | 100 | 94 | 1e-109 |
| Streptococcus dysgalactiae subsp. equisimilis strain NCTC6181 | rplC | LSU ribosomal protein L3p (L3e) | Retapamulin;Spiramycin | 100 | 100 | 94 | 1e-109 |
| Streptococcus dysgalactiae subsp. equisimilis strain NCTC6407 | rplC | LSU ribosomal protein L3p (L3e) | Retapamulin;Spiramycin | 100 | 100 | 94 | 1e-110 |
| Streptococcus dysgalactiae subsp. equisimilis strain NCTC7136 | rplC | LSU ribosomal protein L3p (L3e) | Retapamulin;Spiramycin | 100 | 100 | 99 | 1e-114 |
| Streptococcus dysgalactiae subsp. equisimilis strain NCTC8543 | rplC | LSU ribosomal protein L3p (L3e) | Retapamulin;Spiramycin | 100 | 100 | 99 | 1e-114 |
| Streptococcus dysgalactiae subsp. equisimilis strain NCTC8546 | rplC | LSU ribosomal protein L3p (L3e) | Retapamulin;Spiramycin | 100 | 100 | 99 | 1e-114 |
| Streptococcus dysgalactiae subsp. equisimilis strain NCTC9413 | rplC | LSU ribosomal protein L3p (L3e) | Retapamulin;Spiramycin | 100 | 100 | 99 | 1e-114 |
| Streptococcus dysgalactiae subsp. equisimilis strain NCTC9414 | rplC | LSU ribosomal protein L3p (L3e) | Retapamulin;Spiramycin | 100 | 100 | 99 | 1e-114 |
| Streptococcus dysgalactiae subsp. equisimilis strain NCTC9603 | rplC | LSU ribosomal protein L3p (L3e) | Retapamulin;Spiramycin | 100 | 100 | 99 | 1e-114 |
| Streptococcus dysgalactiae subsp. equisimilis strain SS1575 | rplC | LSU ribosomal protein L3p (L3e) | Retapamulin;Spiramycin | 100 | 100 | 99 | 1e-114 |
| Streptococcus dysgalactiae subsp. equisimilis strain T642 | rplC | LSU ribosomal protein L3p (L3e) | Retapamulin;Spiramycin | 100 | 100 | 99 | 4e-148 |
| Streptococcus dysgalactiae subsp. equisimilis strain UT_4031CC | rplC | LSU ribosomal protein L3p (L3e) | Retapamulin;Spiramycin | 100 | 100 | 99 | 1e-114 |
| Streptococcus dysgalactiae subsp. equisimilis strain UT_4231_KK | rplC | LSU ribosomal protein L3p (L3e) | Retapamulin;Spiramycin | 100 | 100 | 99 | 1e-114 |
| Streptococcus dysgalactiae subsp. equisimilis strain UT_4234_DH | rplC | LSU ribosomal protein L3p (L3e) | Retapamulin;Spiramycin | 100 | 100 | 99 | 1e-114 |
| Streptococcus dysgalactiae subsp. equisimilis strain UT_4241_XS | rplC | LSU ribosomal protein L3p (L3e) | Retapamulin;Spiramycin | 100 | 100 | 99 | 1e-114 |
| Streptococcus dysgalactiae subsp. equisimilis strain UT_4242_AB | rplC | LSU ribosomal protein L3p (L3e) | Retapamulin;Spiramycin | 100 | 100 | 99 | 1e-114 |
| Streptococcus dysgalactiae subsp. equisimilis strain UT_4255RC | rplC | LSU ribosomal protein L3p (L3e) | Retapamulin;Spiramycin | 100 | 100 | 99 | 1e-114 |
| Streptococcus dysgalactiae subsp. equisimilis strain UT_4277_BB | rplC | LSU ribosomal protein L3p (L3e) | Retapamulin;Spiramycin | 100 | 100 | 99 | 1e-114 |
| Streptococcus dysgalactiae subsp. equisimilis strain UT_4966_RC | rplC | LSU ribosomal protein L3p (L3e) | Retapamulin;Spiramycin | 100 | 100 | 99 | 1e-114 |
| Streptococcus dysgalactiae subsp. equisimilis strain UT-5345 | rplC | LSU ribosomal protein L3p (L3e) | Retapamulin;Spiramycin | 100 | 100 | 99 | 1e-114 |
| Streptococcus dysgalactiae subsp. equisimilis strain UT-5354 | rplC | LSU ribosomal protein L3p (L3e) | Retapamulin;Spiramycin | 100 | 100 | 99 | 1e-114 |
| Streptococcus dysgalactiae subsp. equisimilis strain UT-SS1069 | rplC | LSU ribosomal protein L3p (L3e) | Retapamulin;Spiramycin | 100 | 100 | 99 | 1e-114 |
| Streptococcus dysgalactiae subsp. equisimilis strain UT-SS957 | rplC | LSU ribosomal protein L3p (L3e) | Retapamulin;Spiramycin | 100 | 100 | 99 | 1e-114 |
| Streptococcus dysgalactiae subsp. equisimilis strain WCHSDSE-1 | rplC | LSU ribosomal protein L3p (L3e) | Retapamulin;Spiramycin | 100 | 100 | 99 | 1e-114 |
| Streptococcus dysgalactiae subsp. equisimilis SK1249 |  | Catabolite control protein A | 2-Phenylamino-Ethanesulfonic Acid | 20 | 85 | 85 | 2e-25 |
| Streptococcus dysgalactiae subsp. equisimilis strain C161L1 |  | DNA gyrase subunit A (EC 5.99.1.3) |  | 98 | 97 | 82 | 0 |
| Streptococcus dysgalactiae subsp. equisimilis strain C161L1 |  | Enolase (EC 4.2.1.11) |  | 99 | 99 | 81 | 0 |
| Streptococcus dysgalactiae subsp. equisimilis strain C161L1 |  | Glyceraldehyde-3-phosphate dehydrogenase (NADP(+)) (EC 1.2.1.9) |  | 100 | 100 | 85 | 0 |
| Streptococcus dysgalactiae subsp. equisimilis strain C161L1 |  | hypothetical protein |  | 22 | 94 | 92 | 0.0 |
| Streptococcus dysgalactiae subsp. equisimilis strain C161L1 |  | hypothetical protein |  | 27 | 96 | 94 | 0.0 |
| Streptococcus dysgalactiae subsp. equisimilis strain C161L1 |  | hypothetical protein |  | 14 | 93 | 94 | 0.0 |
| Streptococcus dysgalactiae subsp. equisimilis strain C161L1 |  | hypothetical protein |  | 5 | 85 | 88 | 3e-107 |
| Streptococcus dysgalactiae subsp. equisimilis strain T642 |  | DNA gyrase subunit A (EC 5.99.1.3) |  | 98 | 97 | 82 | 0 |
| Streptococcus dysgalactiae subsp. equisimilis strain T642 |  | Enolase (EC 4.2.1.11) |  | 99 | 99 | 81 | 0.0 |
| Streptococcus dysgalactiae subsp. equisimilis strain T642 |  | Glyceraldehyde-3-phosphate dehydrogenase (NADP(+)) (EC 1.2.1.9) |  | 100 | 100 | 85 | 0 |
| Streptococcus dysgalactiae subsp. equisimilis strain T642 |  | Protein G-related alpha 2 macroglobulin-binding protein (GRAB) |  | 72 | 81 | 98 | 0.0 |
| Streptococcus dysgalactiae subsp. equisimilis strain KNZ06 | parC | DNA topoisomerase IV subunit A (EC 5.99.1.3) | Gatifloxacin;Besifloxacin | 47 | 100 | 87 | 1e-203 |
