## Supplemental File 6 for "Comprehensive comparative genomics analysis for the emerging human pathogen *Streptococcus dysgalactiae* subsp. *equisimilis* (SDSE): A case study and Pan-subspecies genomic analysis"

**Supplementary Table 6:** Detailed information for the complete phage genomes annotated in SDSE genomes.

| .Host Strain | Genome Name | Genome Length | GC Content | Consensus CDS | Best Hit of Blast |
| --- | --- | --- | --- | --- | --- |
| SDSE AC-2713 | Streptococcus phage Javan122 | 39241 | 40.79152 | 52 | Adenine-specific methyltransferase; DNA helicase, phage-associated; DNA primase, phage associated; DNA-cytosine methyltransferase; phage encoded DNA polymerase I; Holin; Phage capsid and scaffold; Phage endonuclease; Phage endopeptidase; Phage integrase; Phage major capsid protein; Phage portal protein; Phage protein, contains HNH endonuclease motif; Phage tail length tape-measure protein T; Phage tail tube protein; Phage terminase, large subunit; Phage terminase, small subunit; Phage-associated cell wall hydrolase; Prophage Clp protease-like protein; prophage pi2 protein 37; prophage pi2 protein 38; S-adenosylmethionine synthetase (3), Site-specific recombinase; hypothetical protein (21); phage protein (6) |
| SDSE AC-2713 | Streptococcus phage Javan123 | 37428 | 41.589184 | 49 | Adenine-specific methyltransferase; conserved hypothetical protein; DNA helicase (EC 3.6.4.12), phage-associated; DNA primase, phage associated; DNA-cytosine methyltransferase; FIG006036: phage encoded DNA polymerase I; Holin; Phage endonuclease; Phage head, head-tail preconnector protease C; Phage integrase; Phage major capsid protein; Phage portal protein; Phage tail length tape-measure protein T (2); Phage tail tube protein; Phage terminase, large subunit; Phage terminase, small subunit; Phage-associated cell wall hydrolase; hypothetical protein (19); phage protein (8); Plasmid stabilization system antitoxin protein; Plasmid stabilization system toxin protein |
| SDSE NS3396 | Streptococcus phage phi3396 | 38528 | 37.48183 | 72 | Adenine-specific methyltransferase; Hyaluronate lyase (phage associated); Paratox; Pathogenicity island SaPIn2; Phage antirepressor protein; Phage baseplate; Phage DNA replication protein O; Phage endopeptidase, Phage essential recombination function protein, Erf; Phage head, head-tail preconnector protease C; Phage holin; Phage hyaluronidase; Phage integrase; Phage integrase: site-specific recombinase; Phage major capsid protein; Phage major tail protein; Phage peptidoglycan hydrolase; Phage portal protein; Phage repressor (2); Phage tail; Phage tail length tape-measure protein T; Phage terminase, large subunit; Phage terminase, small subunit; Phage-associated homing endonuclease; putative protein; Transcriptional regulator, hypothetical protein (21), phage protein (24). |
| SDSE strain ASDSE_99 | Streptococcus phage Javan117 | 40234 | 37.744198 | 74 | CI-like repressor, phage associated; DNA modification methylase; DNA replication protein DnaD; DNA-cytosine methyltransferase; Phage capsid and scaffold; Phage capsid protein; Phage holin; Phage integrase; Phage portal; Phage replication protein; Phage tail length tape-measure protein T; Phage-associated cell wall hydrolase; Protein gp30; putative minor structural protein; Recombinational DNA repair protein RecT (prophage associated); terminase, large subunit, putative; Transcriptional regulator; Type II, 5-methyl-cytosine DNA methyltransferase; Type II, 5-methyl-cytosine DNA methyltransferase; hypothetical protein (34); phage protein (21) |
| SDSE UT_4231_KK | Streptococcus phage Javan161 | 42720 | 38.81086 | 75 | Abortive infection bacteriophage resistance protein; Adenine-specific methyltransferase; Bacteriophage replicative DNA helicase, repA; Bacteriophage resolvase; DNA helicase, phage associated; Type III restriction enzyme; Hyaluronate lyase (phage associated); Paratox; Phage capsid protein; Phage DNA packaging protein (ACLAME 138); Phage endonuclease; Phage endopeptidase; Phage holin; Phage hyaluronidase; Phage integrase, site-specific tyrosine recombinase; Phage major tail protein; Phage peptidoglycan hydrolase; Phage portal protein; Phage tail length tape-measure protein T; Phage terminase, large subunit; Phage terminase, small subunit; Pleiotropic regulator of exopolysaccharide synthesis competence and biofilm formation Ftr XRE family, Prophage Clp protease-like protein; Streptococcal extracellular nuclease 3 Mitogenic factor 3; hypothetical protein (29); phage protein (23). |
| SDSE UT_4231_KK | Streptococcus satellite phage Javan162 | 14338 | 36.392803 | 39 | CI-like repressor, phage associated; DNA integration/recombination/invertion protein; DNA primase, phage associated; kilA protein, putative phage-related DNA binding protein; Phage antirepressor protein; Phage encoded; transcriptional regulator, ArpU family; Putative DNA-binding protein in cluster with Type I restriction-modification system; hypothetical protein (27); phage protein (5) |
| SDSE UT_4242_AB | Streptococcus phage Javan163 | 40438 | 39.789307 | 60 | DNA helicase (phage-associated), DNA modification methyltransferase (2); DNA polymerase (phage-associated); DNA primase (phage associated); Hyaluronate lyase (phage associated); Integrase; Phage capsid and scaffold; Phage endopeptidase; Phage hyaluronidase; Phage lysin; Phage major capsid protein; Phage portal; Phage tail length tape-measure protein T; Phage terminase, large subunit; Phage terminase, small subunit; Phage transcriptional activator; Phage transcriptional regulator; Streptodornase D; hypothetical protein (9); Phage protein (32) |
| SDSE UT_4966_RC | Streptococcus phage Javan164 | 37145 | 37.493607 | 68 | Conserved hypothetical protein - phage associated; Hyaluronate lyase (phage associated); Paratox; Phage antirepressor protein; Phage baseplate; Phage DNA replication protein O; Phage DNA-binding protein; Phage endonuclease / Phage holin (ACLAME 9); Phage endopeptidase; Phage head, head-tail preconnector protease C; Phage holin (2); Phage hyaluronidase; Phage integrase (2); Phage integrase: site-specific recombinase; Phage major capsid protein; Phage major tail protein; Phage peptidoglycan hydrolase; Phage portal protein; Phage protein (ACLAME 893); Phage tail; Phage tail length tape-measure protein T; Phage terminase, large subunit; Phage terminase, small subunit; Phage-associated homing endonuclease (2); Predicted transcriptional regulator; Single-stranded DNA-binding protein; Streptodornase D; hypothetical protein (21); phage protein (17) |
| SDSE UT_4966_RC | Streptococcus satellite phage Javan165 | 13418 | 35.668507 | 38 | DNA integration/recombination/invertion protein (2); DNA primase, phage associated; Phage antirepressor protein; Phage encoded transcriptional regulator, ArpU family; Predicted transcriptional regulator; Putative DNA-binding protein in cluster with Type I restriction-modification system; hypothetical protein (27); Phage protein (4) |
| SDSE UT-5345 | Streptococcus phage Javan166 | 45241 | 38.597733 | 85 | CI-like repressor, phage associated; conserved hypothetical protein; DNA helicase (phage-associated); Hyaluronate lyase (phage associated); Paratox; Phage antirepressor protein; Phage capsid and scaffold; Phage DNA binding protein; Phage DNA helicase; Phage DNA primase/helicase; Phage endonuclease / Phage holin (ACLAME 9); Phage endopeptidase; Phage hyaluronidase; Phage integrase; Phage lysin; Phage major capsid protein (2); Phage major tail protein; Phage portal (connector) protein; Phage tail; Phage tail fibers; Phage tail length tape-measure protein T; Phage terminase, large subunit Phage terminase, small subunit; putative antirepressor - phage associated; SAM-dependent methyltransferase; Structural protein; hypothetical protein (30); phage protein (28) |
| SDSE UT-5345 | Streptococcus satellite phage Javan167 | 8144 | 33.349705 | 16 | conserved hypothetical protein (phage associated); Helicase loader DnaI; HTH DNA-binding protein; Phage head, head-tail preconnector protease C; Phage integrase; prophage ps3 protein 13; hypothetical protein (10) |
| SDSE UT-5345 | Streptococcus satellite phage Javan168 | 13967 | 35.81299 | 39 | DNA integration/recombination/invertion protein; DNA primase (phage associated); Phage antirepressor protein; Phage encoded transcriptional regulator, ArpU family; Predicted transcriptional regulator; Putative DNA-binding protein in cluster with Type I restriction-modification system, hypothetical protein (28); phage protein (5) |
| SDSE UT-5354 | Streptococcus phage Javan169 | 44153 | 38.500214 | 90 | Adenine-specific methyltransferase; Endodeoxyribonuclease RusA; Hyaluronate lyase (phage associated); Integrase; Paratox; Phage antirepressor protein; Phage capsid protein; Phage DNA packaging protein (ACLAME 138); Phage endonuclease; Phage endopeptidase; Phage essential recombination function protein, Erf; Phage hyaluronidase; Phage lysin; Phage lysin, N-acetylmuramoyl-L-alanine amidase; Phage major tail protein; Phage portal protein; Phage protein (ACLAME 87); Phage replication initiation protein; Phage tail length tape-measure protein T; Phage terminase, large subunit; Phage terminase, small subunit; Pleiotropic regulator of exopolysaccharide synthesis, competence and biofilm formation Ftr, XRE family; Prophage Clp protease-like protein; putative cro protein; putative DNA-binding phage protein; Single-stranded DNA-binding protein; Uncharacterized protein Npun_F4007; hypothetical protein (39); phage protein (24) |
| SDSE UT-SS1069 | Streptococcus phage Javan170 | 42762 | 38.81951 | 78 | Abortive infection bacteriophage resistance protein; Adenine-specific methyltransferase; Bacteriophage replicative DNA helicase, repA Bacteriophage resolvase; DNA helicase, phage associated; Type III restriction enzyme; DNA helicase, phage-associated; Type III restriction enzyme Hyaluronate lyase (phage associated); Paratox; Phage capsid protein; Phage cI repressor (ACLAME 5); Phage DNA packaging protein (ACLAME 138), Phage endonuclease; Phage endopeptidase; Phage holin; Phage hyaluronidase; Phage integrase, site-specific tyrosine recombinase; Phage major tail protein; Phage peptidoglycan hydrolase; Phage portal protein; Phage tail length tape-measure protein T; Phage terminase, large subunit; Phage terminase, small subunit; Pleiotropic regulator of exopolysaccharide synthesis, competence and biofilm formation Ftr, XRE family; Prophage Clp protease-like protein; Streptococcal extracellular nuclease 3 (Mitogenic factor 3); hypothetical protein (30); Phage protein (22) |
| SDSE UT-SS1069 | Streptococcus satellite phage Javan171 | 12900 | 36.263565 | 34 | DNA integration/recombination/invertion protein; DNA primase, phage associated (2); Phage antirepressor protein; Phage encoded transcriptional regulator, ArpU family; Predicted transcriptional regulator; hypothetical protein (24); Phage protein (4) |
| SDSE WCHSDSE-1 | Streptococcus phage Javan172 | 39869 | 36.923424 | 74 | DNA-cytosine methyltransferase; Hyaluronate lyase (phage associated); Paratox; Phage antirepressor protein; Phage baseplate; Phage DNA replication protein O; Phage DNA-binding protein; Phage endopeptidase; Phage head, head-tail preconnector protease C; Phage hyaluronidase; Phage integrase; Phage integrase: site-specific recombinase; Phage lysin; Phage major capsid protein; Phage major tail protein; Phage portal protein; Phage protein, conserved hypothetical protein - phage associated (2); Phage repressor; Phage tail length tape-measure protein T; Phage terminase, large subunit; Phage terminase, small subunit; Phage-associated homing endonuclease; Phage-associated homing endonuclease; Single-stranded DNA-binding protein; Transcriptional regulator; Type II, 5-methyl-cytosine DNA methyltransferase; Hypothetical protein (26); Phage protein (20) |
